# Using Temporal ICA to Selectively Remove Global Noise While Preserving Global Signal in Functional MRI Data

**DOI:** 10.1101/193862

**Authors:** Matthew F. Glasser, Timothy S. Coalson, Janine D. Bijsterbosch, Samuel J. Harrison, Michael P. Harms, Alan Anticevic, David C. Van Essen, Stephen M. Smith

## Abstract

Temporal fluctuations in functional Magnetic Resonance Imaging (fMRI) have been profitably used to study brain activity and connectivity for over two decades. Unfortunately, fMRI data also contain structured temporal “noise” from a variety of sources, including subject motion, subject physiology, and the MRI equipment. Recently, methods have been developed to automatically and selectively remove spatially specific structured noise from fMRI data using spatial Independent Components Analysis (ICA) and machine learning classifiers. Spatial ICA is particularly effective at removing spatially specific structured noise from high temporal and spatial resolution fMRI data of the type acquired by the Human Connectome Project and similar studies. However, spatial ICA is mathematically, by design, unable to separate spatially widespread “global” structured noise from fMRI data (e.g., blood flow modulations from subject respiration). No methods currently exist to selectively and completely remove global structured noise while retaining the global signal from neural activity. This has left the field in a quandary—to do or not to do global signal regression—given that both choices have substantial downsides. Here we show that temporal ICA can selectively segregate and remove global structured noise while retaining global neural signal in both task-based and resting state fMRI data. We compare the results before and after temporal ICA cleanup to those from global signal regression and show that temporal ICA cleanup removes the global positive biases caused by global physiological noise without inducing the network-specific negative biases of global signal regression. We believe that temporal ICA cleanup provides a “best of both worlds” solution to the global signal and global noise dilemma and that temporal ICA itself unlocks interesting neurobiological insights from fMRI data.

## Introduction

Functional MRI (fMRI) is a remarkably powerful tool for non-invasive mapping of brain function and for estimating the functional interactions between brain areas (i.e., “functional connectivity”). Although its indirectness has been criticized (Farah, 2014), both task-based and resting state fMRI are capable of precisely replicating known fine-scale patterns of functional organization and connectivity in the human visual cortex (Glasser et al., 2016a; Sereno et al., 1995) when analyzed so that spatial details are preserved by aligning areas across subjects on the cortical surface and avoiding smoothing (Glasser et al., 2016b). Such patterns closely reflect many fine details observed invasively in nonhuman primates (Brewer et al., 2002; Gattass et al., 1988; Van Essen et al., 1984), lending strong support for fMRI as a tool for non-invasively mapping the brain’s functional organization.

It is also known, however, that a variety of confounding and nuisance signals are present in the fMRI timeseries, particularly those arising from subject motion (Power et al., 2012; Power et al., 2014; Power et al., 2018; Satterthwaite et al., 2012; Yan et al., 2013a), and subject physiology, such as respiration and heart rate (Birn et al., 2006; Chang et al., 2009; Chang and Glover, 2009; Golestani et al., 2015; Power et al., 2018; Power et al., 2017b; Shmueli et al., 2007). If not removed systematically, such signals can lead to false positives, false negatives, and erroneous interpretations of fMRI results. Indeed, how to appropriately and comprehensively remove structured (i.e., non-Gaussian) temporal noise from fMRI data is a longstanding methodological controversy in the functional neuroimaging community (Aguirre et al., 1997; Aguirre et al., 1998; Anderson et al., 2011; Fox et al., 2009; Liu, 2016; Liu et al., 2017; Macey et al., 2004; Murphy et al., 2009; Murphy and Fox, 2017; Power et al., 2017a; Power et al., 2014; Power et al., 2015; Saad et al., 2012; Shmueli et al., 2007; Uddin, 2017; Zarahn et al., 1997). Analogous to our careful approach to removing spatial imaging artifacts and aligning brains across subjects without blurring the data (Glasser et al., 2016b; Glasser et al., 2013), it is desirable to use temporal cleanup methods that selectively remove artifacts while preserving as much of the neurally generated signal of interest as possible.

Significant progress has been made in developing automated methods for selectively removing spatially specific structured temporal noise—i.e., non-random time-varying artifacts that are also spatially nonuniform—using spatial ICA (sICA, e.g., sICA+FIX, FMRIB’s ICA component classifier, (Griffanti et al., 2014; Salimi-Khorshidi et al., 2014); see also ICA-AROMA, (Pruim et al., 2015a; Pruim et al., 2015b), and Multi-Echo ICA (Kundu et al., 2012)). Semi-global and global structured noise has presented a more difficult challenge, however. sICA is inherently unable to separate global temporal fluctuations into spatially orthogonal signal and noise components (see Main Supplementary Information Section #1), and thus the global fluctuations will remain mixed into all of the sICA component timeseries. Indeed, several studies have used indirect evidence to assert that there is residual structured noise in Human Connectome Project (HCP) resting state fMRI data after sICA+FIX cleanup (Burgess et al., 2016; Power, 2017; Power et al., 2017b; Siegel et al., 2017); however, this noise has not been selectively separated from the data or characterized as to its spatial and temporal properties. The most common approach for removing global structured noise is to remove the mean (across space) fMRI timecourse from the data either explicitly (using Global Signal Regression, GSR (Power et al., 2017b)) or implicitly (by including white matter and ventricle voxels that are likely contaminated by the mean grey-matter signal because of voxel size, unconstrained volumetric smoothing, and/or spatial proximity (Behzadi et al., 2007; Chai et al., 2012; Marx et al., 2013; Muschelli et al., 2014; Power et al., 2018; Power et al., 2017b).

Global signal regression is not selective, and its appropriateness is predicated on the assumption that global fluctuations are entirely artifactual. However, GSR will also remove or reduce any global or semi-global neural signal in the data, in particular impacting brain functional networks that are spatially widespread or have higher amplitude fluctuations such that they contribute more to the mean timeseries used in GSR (Glasser et al., 2016b). Indeed global fMRI fluctuations have some correlation with global electrophysiological signals in laboratory animals (Scholvinck et al., 2010) and humans (Wen and Liu, 2016) and have been related to the brain’s state of arousal (Chang et al., 2016; Wong et al., 2016; Wong et al., 2013). As a result, GSR and related approaches are controversial because the removal of neural signal may distort the resulting connectivity or activation measures in complex and network-specific ways (Glasser et al., 2016b; Gotts et al., 2013; Saad et al., 2012; Yang et al., 2016; Yang et al., 2014). That said, it is clear that non-neuronal physiological processes induce global fluctuations in fMRI timeseries through the T2* dependent BOLD mechanism, particularly those that globally change brain blood flow such as respiratory rate and depth (Power et al., 2018; Power et al., 2017b). Such artifactual fluctuations will lead to positive biases in apparent functional connectivity (or functional activation if they correlate with the task stimulus), which may vary across subjects and groups to create confounds for brain imaging studies (Glasser et al., 2016b; Hayasaka, 2013; Power et al., 2017b). These global fluctuations are incompletely removed by current models of physiological regressors based on separate recording of physiological parameters (Power et al., 2017b). Other intermediate strategies such as removing the first principal component (Carbonell et al., 2011), which is highly correlated with the mean global timecourse (Carbonell et al., 2011; He and Liu, 2012), or reducing its strength relative to the other principal components (He and Liu, 2012) are not selective as the first principal component contains a mix of global neural signal and global artifact. Thus, the field has lacked an effective and mathematically unbiased solution for separating and removing global and semi-global sources of structured temporal noise while retaining global, semi-global, or high amplitude BOLD neural signal (Glasser et al., 2016b; Power et al., 2018; Power et al., 2017b).

Here, we use a temporal ICA-based (tICA) approach to address this problem. Building upon recent success in separating spatially specific neural signal and structured noise by decomposing fMRI data into spatially orthogonal independent components (using sICA), we sought to further subdivide the resulting sICA-cleaned data into temporally orthogonal independent components (using tICA). Unlike sICA, which is mathematically blind to global signal and noise, tICA is able to identify and separate global and semi-global temporally independent components (Smith et al., 2012) (see Main Supplementary Information Section #1 for more on the distinction between tICA and sICA). We explore the relationship between tICA components and both physiological and motion parameters in task-based and resting state fMRI. We identify multiple specific components as non-neuronal structured noise, remove these components from the fMRI data, and compare the results of common task-based and resting state fMRI analyses before tICA cleanup, after tICA cleanup, and after global signal regression. While the tICA cleanup method is primarily intended for resting state fMRI data, we first consider task fMRI data, because we have explicit hypotheses about a portion of the subjects’ neural activity that allows us to objectively compare the resulting BOLD responses after each cleanup approach in relation to these hypotheses. We then discuss the implications of these results for task fMRI analyses and how they inform the resting state analyses that follow, where we do not have explicit hypotheses about our subjects’ neural activity and thus lack a ground truth.

At the outset, it is worth setting some expectations of what the proposed method aims to do and what we believe is outside of the purview of a data denoising approach. Our objective is to clean the fMRI timeseries of global and semi-global noise from physiological or other sources and also to remove any remaining spatially specific noise that may have been left behind by sICA+FIX so that the fMRI timeseries reflects, as accurately as possible, the “true” neural BOLD signal that occurred in each subject’s brain during their scans. As a result, we do not aim to remove the neurally driven BOLD effects of undesired subject behavior in the scanner. Some subjects may not have actually done the requested task in task-based fMRI, or they may not have kept their eyes open and fixated on the cross-hairs for resting state fMRI. They may have moved frequently and as a result have neurally driven BOLD effects arising from motion, and some subjects may have become drowsy or fallen asleep during the long HCP resting state scans, causing profound changes in their functional connectivity (Fukunaga et al., 2006; Horovitz et al., 2008; Laumann et al., 2017; Liu et al., 2017; Tagliazucchi and Laufs, 2014; Wong et al., 2016; Wong et al., 2013; Yeo et al., 2015). While it may be appropriate for some studies to set stringent data inclusion and exclusion criteria surrounding such “neural compliance” issues, addressing such issues is outside the scope of our method and this study (see Discussion), even though temporal ICA does shed some light on them. Importantly, neural activity and functional connectivity need not be the same during periods of non-compliance as they would have been if the subject had been compliant and such differences are expected to be present in both global and spatially specific neural signals. Accordingly, there are interpretational difficulties with several published data cleanup metrics (Burgess et al., 2016; Ciric et al., 2017; Power et al., 2017a; Power et al., 2015; Siegel et al., 2017) as these methods assume (explicitly or implicitly) that neural activity is the same during periods of subject compliance and non-compliance and in some cases that all other subject-wise parameters that may influence BOLD fMRI at any time during a scan are the same across compliant and non-compliant subjects. Thus, if one wishes to preserve as much neural signal as possible, these existing data cleanup metrics cannot in of themselves be used to determine whether one cleanup approach is superior to another. Nonetheless, we discuss and include some of these metrics after various cleanup stages so as to provide a historical perspective on prior literature (see Main Supplementary Information Section #6). Finally, we emphasize the particular relevance of our approach to the growing amount of high spatial and temporal resolution multi-band fMRI data being acquired as a part of the Human Connectome Project and related HCP-Style neuroimaging efforts (Glasser et al., 2016b). We focus our efforts on multi-band fMRI data because this approach is quickly becoming standard in the field, and such data work best with the powerful data-driven methods that we will use.

## Subjects and Methods

### 1.1 Subject Population

Data from 449 Human Connectome Project young healthy adults (ages 22 – 35) were used in this study, all from the HCP S500 data release. These data were acquired in accordance with the Washington University Institutional Review Board (Van Essen et al., 2013). Only subjects with complete fMRI acquisitions (resting-state and task) from the S500 HCP data release were included (Glasser et al., 2016a). Subject groups (all 449, 210P, and 210V) were determined in a prior study (Glasser et al., 2016a).

### 1.2 Images Acquired

T1-weighted and T2-weighted structural scans were acquired as previously described (Glasser et al., 2013). Resting-state fMRI data were acquired with 2.0mm isotropic resolution, TR=720ms, and 1200 frames (14.4 min) per run. Two runs with reversed phase encoding directions, RL or LR, with the order counterbalanced across each of two sessions, were acquired (Smith et al., 2013a) for a total of 4800 frames of resting state per subject. Task fMRI data were acquired with identical pulse sequence settings while subjects performed 7 tasks (Barch et al., 2013) with runs lasting between 2 and 5 minutes (176 - 405 frames), and totaling 22.6 min (1884 frames) in session 1 and 24.0 min (1996 frames) in session 2 for a total of 3880 frames of task fMRI per subject. Each task is comprised of a pair of runs with reversed phase encoding directions, RL and then LR. Task runs were halted if a subject stopped performing the task. Spin echo EPI scans were acquired (termed “spin echo field map” scans by the HCP) and used to correct the fMRI data for B0-induced geometric distortions and B1-receive-induced image intensity inhomogeneities (Glasser et al., 2016a; Glasser et al., 2013).

### 1.3 Image Preprocessing

The HCP’s spatial image preprocessing has been described previously in detail (Glasser et al., 2016a; Glasser et al., 2013). In brief, it involves minimizing smoothing while doing the following: 1) removing MR-induced image distortions so that each image represents the physical space of the subject; 2) removing the spatial effects of subject motion within and between modalities; and 3) projecting the data to a 2mm average spacing standard CIFTI grayordinates space after a FNIRT-based T1w nonlinear subcortical volume alignment and an MSMAll areal-feature-based cortical alignment, which uses myelin maps, resting state network maps, and resting state visuotopic maps for registration (Glasser et al., 2016a; Robinson et al., 2017; Robinson et al., 2014). For resting state fMRI, sICA+FIX was run (Griffanti et al., 2014; Salimi-Khorshidi et al., 2014) to identify and remove spatially specific noise components using a machine learning classifier trained on HCP data. Task fMRI data was processed using a modified sICA+FIX pipeline (“Multi-run sICA+FIX”) that utilized concatenation across runs and phase encoding directions within a single scanning session. The original and modified sICA+FIX pipelines are described in Main Supplemental Information Section #2. The result of these steps was the cleaned grayordinate-wise (“dense”) timeseries or voxelwise timeseries. Cross-run or cross-session timeseries were then concatenated after normalization of the unstructured noise variance (Glasser et al., 2016a). An overview of subsequent processing is shown in Figure 1.

**Figure 1.**
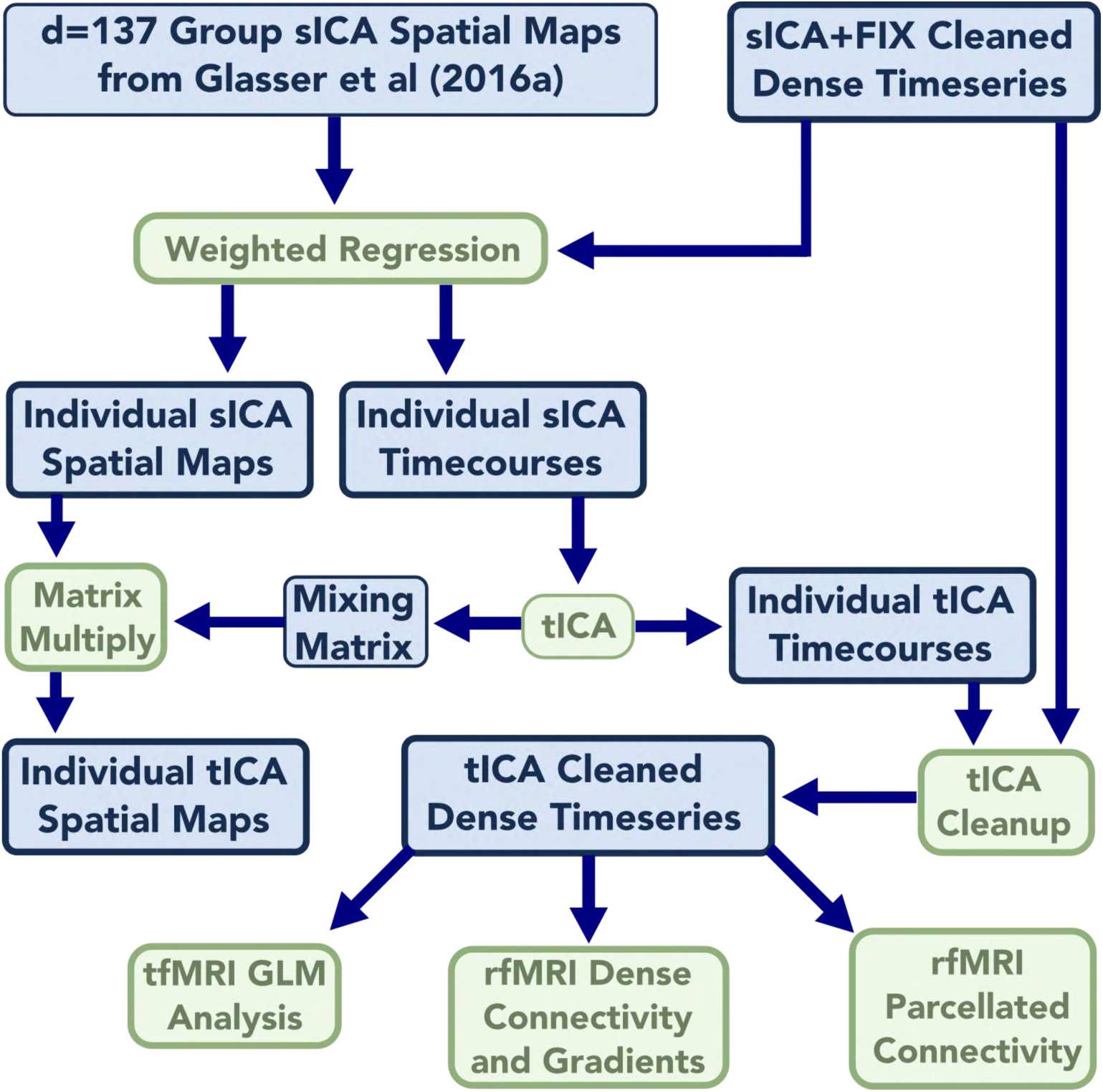
shows an overview of the methods of the paper from the preprocessed dense timeseries to the final analysis outputs. Data are blue and algorithms are green. Data or algorithms with a thicker outline were run on a per-subject or per run basis. All 449 subjects of both resting state and task data were run through the process; however, only the 210P subgroup was used in the group dense functional connectivity and gradient analysis for computational reasons (210P and 210V groups of subjects have a correlation of r=0.98 in their group functional connectivity, (Glasser et al., 2016a)).

### 1.4 Cross-Subject Consistent Spatial ICA Dimensionality Reduction

We reduced the dimensionality of each subject’s sICA+FIX cleaned, MSMAll aligned dense timeseries using a previously published group sICA decomposition of d=137 components (Glasser et al., 2016a). Performing an initial spatial ICA data reduction is similar to the approach previously used to perform temporal ICA on fMRI data (Smith et al., 2012), and it enables a matched dimensionality reduction to occur across all subjects. Individual subject sICA component timecourses were created using weighted spatial regression (Glasser et al., 2016a), a variant of dual regression (Filippini et al., 2009) that helps to further compensate for any residual misalignments after MSMAll areal-feature-based registration by weighting the spatial regression according to alignment quality. This process individualizes the group components to each subject’s resting state and task fMRI data. The first stage of weighted regression involves regressing the group sICA component spatial maps into each individual’s dense timeseries. The final stage of weighted regression involves temporally regressing the individual subject component timecourses into both the grayordinate and volume-based individual subject timeseries to produce individual subject component spatial maps for both CIFTI grayordinate and volume spaces. These spatial maps were then averaged across subjects. No smoothing was applied to either set of maps prior to averaging so as maximize the spatial sharpness of the group averages. Thus, all computations requiring spatial correspondence were performed in MSMAll aligned standard CIFTI grayordinates space (Glasser et al., 2016b), whereas steps requiring temporal correspondence, such as temporal regression, are unaffected by spatial misalignment and can occur in standard volume space.

Thus the final outputs for each subject of the spatial ICA dimensionality reduction for both resting state and task fMRI were: (1) a timecourse for each of the d=137 group-ICA components for each subject, (2) a subject-specific grayordinate-based spatial map for each component, and (3) a subject-specific volume-based spatial map for each component. Additionally, group averages of (2) and (3) were generated, and (1) was temporally concatenated across subjects into a 137 X ∼2 million timepoint matrix (one for rfMRI and one for tfMRI). Equation #1 summarizes the spatial ICA decomposition (either at the group level for the initial sICA computation or after projecting the decomposition to the individuals using weighted regression) with the Data_Nspace X Ntime_ representing a dense timeseries in an individual (or a PCA dimensionality reduction in the original group sICA decomposition, see Methods Section #1.14). sICA_Maps_Nspace X DsICA_ are the sICA spatial maps and sICA_TCS_DsICA X Ntime_ are the sICA timecourses (TCS). Error is the unstructured noise subspace left over after the dimensionality reduction, whereas the product of sICA_Maps_Nspace X DsICA_ and sICA_TCS_DsICA X Ntime_ represent the structured subspace of the data (e.g., d=137; see also Main Supplementary Information Section #3 on partitioning fMRI data into different subspaces and variance bins). The equation also applies to the individual volume dense timeseries and volume-based spatial maps (where space is voxels instead of grayordinates) when the volume dense timeseries is substituted for the last temporal regression stage of weighted regression (Glasser et al., 2016a):

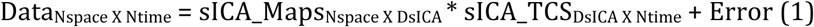

### 1.5 Temporal ICA

Temporal ICA was performed on the cross-subject concatenated individual subject sICA component timecourses using a method similar to (Smith et al., 2012) with the FAST ICA algorithm (Hyvarinen, 1999) implemented in Matlab. Although the maximum possible temporal ICA dimensionality was 137 (which is the number of concatenated sICA component timecourses), the temporal ICA decompositions were not reproducible at this dimensionality. We used binary search and the ICASSO (Himberg et al., 2004) algorithm to identify the dimensionality yielding the maximum number of clusters of reproducible components (greater than 0.5 in the ICASSO Iq cluster quality measure, range 0 to 1). We found 70 reproducible clusters for task fMRI and 84 for resting state fMRI. ICASSO was then used to find the component cluster centrotypes of (i.e., the component estimate closest to the cluster center), which were used as an initialization to a final temporal ICA run with FastICA to produce the final temporally orthogonal decomposition. The other FastICA parameter settings were: nonlinear function = tanh, estimation method = symm, and there was no further PCA-based dimensionality reduction prior to the 100 ICASSO iterations or the final ICA decomposition.

The results of the temporal ICA were a set of unmixed timecourses (representing the underlying temporally independent latent sources) and a mixing matrix (which represents how to combine the tICA timeseries to recover the estimate of the original sICA component timeseries). Equation #2 describes the group level tICA computation (on concatenated sICA component timecourses) or the single subject decomposition, with sICA_TCS_DsICA X Ntime_ representing either the concatenated group sICA timeseries or that of a single subject. Mixing_Matrix_DsICA X DtICA_ always represents the tICA mixing matrix computed by FastICA at the group level, and tICA_TCS_DtICA X Ntime_ represents the unmixed tICA component timecourses at either the group or individual subject level. The single subject tICA component timecourses are produced by deconcatenating the group timecourses. Note that for tICA there is no “error” term as there is no further PCA data reduction before running tICA.

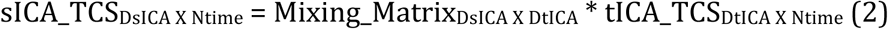

To understand how to get spatial maps of temporal ICA components it is necessary to write out the full equation that substitutes Equation #2 into Equation #1 and thus includes both spatial and temporal ICA decompositions: Equation #3. This equation can be rewritten as Equation #4 by introducing a term that represents the spatial maps of the temporal ICA components tICA_Maps_Nspace X DtICA_. At the group level Data_Nspace X Ntime_ would be an enormous file of concatenated dense timeseries and thus is never generated (though see Section 1.14 below for a Principal Components Analysis (PCA) approximation of this).

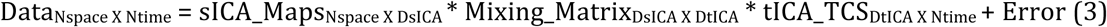

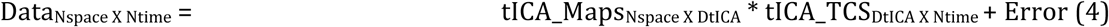

Thus, the spatial maps of temporal ICA are given by Equation 5, and these can be either the individual subject spatial maps or the mean spatial maps at the group level:

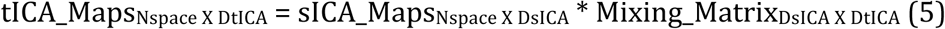

Thus, the group average spatial ICA maps in both grayordinate and volume spaces were multiplied by the temporal ICA mixing matrix, producing grayordinate-based and volume-based spatial maps for each temporal ICA component. These components were ordered according to the RMS (root mean square) of the mixing matrix, and the maximum spatial map value was set by convention to be positive. This imposes a specific order and sign convention on the components, making stronger components that explain more variance appear first and weaker components appear last (similar to how FSL’s Melodic orders components), though the order is not exactly by percent variance explained. Because the data used for temporal ICA was the 137 X ∼2 million timepoints concatenated sICA component timecourse matrix that has so many more timepoints than ‘spatial’ points, group tICA is able to outperform single subject temporal ICA (Smith et al., 2012). Because we have functionally aligned the data, have already cleaned the data of subject-specific, spatially-specific temporal artifacts, and are primarily interested in semi-global and global artifacts, we did not expect the group analysis to be substantially harmed by either residual misalignments or large numbers of uniquely subject-specific artifacts. That said, we did find some single-subject-specific artifacts of both spatially specific and global natures; however, these components did not use up the available degrees of freedom, as the number of reproducible components for resting state (d=84) and task (d=70) temporal ICA were substantially less than the d=137 maximum imposed by the spatial ICA dimensionality reduction.

### 1.6 Physiological Noise Modeling

Physiological data were acquired using a respiratory belt and a heart rate monitor (pulse oximeter on a finger) in most runs from most subjects (87%). We used FSL’s Physiological Noise Modelling (PNM) tool (Brooks et al., 2008) to convert these two traces into 14 physiological regressors, including 4 cardiac regressors, 4 respiratory regressors, 4 interaction regressors, a heart rate regressor and an RVT regressor (Respiration Volume per Time; (Birn et al., 2006), see also: https://fsl.fmrib.ox.ac.uk/fsl/fslwiki/PNM/UserGuide). These physiological measures were then compared with temporal ICA component timeseries in those subjects that had physiologic measures acquired. Because of the very large number of physiological traces, the HCP was unable to manually review and quality assure the peak detection of each trace for each subject. Therefore, these data represent a useful but imperfect ‘proof-of-principle’ measure of subject-specific physiology, and we rely on prior work in a smaller study that did manual quality control and shows the extent to which physiological regressors can remove artifacts from fMRI data (Power et al., 2017b).

In practice, the RVT trace was found to have the strongest relationship to the data (likely because the more spatially specific, higher frequency physiological artifacts had already been removed by sICA+FIX). Indeed this could be predicted from the group average beta maps of the physiological regressors after only detrending and motion regression (i.e., no sICA+FIX cleanup), as RVT has the strongest global relationship with the fMRI data and for simplicity we chose to focus on RVT. Because the quality of the HCP RVT data is variable and there is a variable amount of respiratory signal contamination in fMRI data (Power et al., 2017b), we chose to focus on those runs that had 1) good quality RVT data and 2) a substantial contamination of the fMRI timeseries by respiratory signal. These data will show the strongest relationship between the tICA components and respiration. To identify these runs, we correlated the RVT trace with the average (“parcellated”) timeseries from each of the 360 areas of the HCP-MMP1.0 multi-modal parcellation^1^ (Glasser et al., 2016a) after cleanup with sICA+FIX for each resting state fMRI run or concatenated fMRI task session^2^. For each subject, we then averaged the correlation value across parcels. We took the top 10% of runs or sessions where RVT had the most substantial relationship with the parcellated fMRI data and then correlated the temporal ICA component timeseries with RVT for these runs.

### 1.7 Modeling Motion

We used a newly developed measure of motion that we have termed “DVARS Dips” instead of more commonly used measures of motion such as Framewise Displacement (FD; Power et al., 2012) or DVARS (D referring to the temporal derivative of timecourses, VARS referring to RMS variance over voxels) (Burgess et al., 2016; Power et al., 2012; Smyser et al., 2010). We believe this measure is particularly relevant for fast TR fMRI data because it is more specific to genuine physical head motion that disrupts image intensities. The measure is not affected by “phantom motion” that results from the magnetic field fluctuations that become evident in ‘motion’ estimates when using fast TR fMRI nor by global fluctuations from other sources such as physiology or neural signal that may be present in DVARS (see Main Supplementary Information Section #4). Notably, both Burgess et al. (2016) and Power et al. (2017) have commented that the FD estimates in the HCP data do not always exhibit the same utility for flagging ‘motion corrupted’ time points as has been observed in traditional (slower TR) data.

DVARS Dips are computed as deviations below (or above) the median DVARS of a given run or concatenated session’s unstructured noise timeseries after regressing out all structured signals (i.e., both signal and noise components from sICA+FIX, see Main Supplementary Information Section #4). We used an empirically chosen threshold of +/- 25 to identify frames as “Dips”, because this maximized the correlation between the number of DVARS dips and the subject-wise standard deviation of an obviously motion-related tICA component (TC51/RC50); however, tested values between 15 and 50 had similar correlations, making any values in this range reasonable. For the concatenated task fMRI session DVARS we eliminated any differences in the median DVARS between the individual runs before identifying the Dips (or Spikes). DVARS Dips avoid the issues with phantom motion in FD that appear to be related to subject weight and BMI, though DVARS Dips are highly correlated with FD in the absence of phantom motion (see Supplementary Figures 1, 2, and 3 and Supplementary Table 1). Because DVARS Dips are computed on the unstructured noise timeseries, they are essentially unaffected by the global fluctuations that are under investigation here (as the global signal variance of the unstructured noise timeseries is only 6% of that of the sICA+FIX cleaned timeseries of the resting state data), and thus should not be biased by them. See Main Supplementary Information Section #4 for an explanation of how DVARS Dips are caused by motion and further discussion of the corruption of FD by phantom motion.

### 1.8 Examining the Effects of Sleep on Resting State fMRI

In addition to motion, sleep is another subject behavior that has potentially both neural and artifactual correlates. During sleep, respiratory patterns may change (Igasaki et al., 2016), which may lead to more artifactual physiologically driven BOLD fluctuations. In addition, sleeping or drowsy subjects may also exhibit different amounts of head motion. Also, arousal state is known to affect the amount of global signal in the brain (Laumann et al., 2017; Liu et al., 2017; Tagliazucchi and Laufs, 2014; Wong et al., 2016; Wong et al., 2013; Yeo et al., 2015), though it is not yet known to what extent this effect is due physiological confounds or genuine neural effects. Unfortunately the HCP was unable to implement eye tracking of its young-adult cohort; however, the personnel operating the scanner were instructed to document when subjects were obviously sleeping. These acquisition logs were extracted from the internally facing HCP database, and any subject who was noted to be sleeping during any resting-state scan was flagged in the current analysis as a ‘sleepy subject’. This allowed explicit comparisons between subjects who had been noted to be sleeping and those who had not. As with the physiology measures discussed above, this an imperfect but highly useful proof of concept metric.

### 1.9 Classification of Temporal ICA Components as Signal or Noise

All temporal ICA components were classified manually using multiple sources of information, as summarized for each component in the Supplementary Information Sections TC, TCr, and RC. The specific rationale for each component’s classification is reported at the bottom of its figure. The tICA Spatial Maps (Eq. (5)), both on the surface and in the volume, were the most important source of information for component classification, as components could usually be clearly determined as neural “Signal” based on (i) similarity to known resting state networks (Laumann et al., 2015; Yeo et al., 2011) or known task activation patterns (in the task fMRI data); (ii) existence of boundaries between positive and negative patches that matched known areal boundaries (Glasser et al., 2016a); or (iii) they matched known somatotopic or retinotopic topographic organization (Glasser et al., 2016a). Indeed, as has been found with spatial ICA (Griffanti et al., 2017), temporal ICA signal components are usually visibly distinct from structured noise components.

Prior work demonstrated that respiratory-related global structured noise often appears as pan-grey matter ‘greyplot stripes’ (see Section #1.11 on greyplots below) that are attenuated within white matter and CSF as one moves away from the greymatter (Power et al., 2018; Power et al., 2017b). Additionally, the global timecourse after sICA+FIX is known to be a grey matter specific signal (Glasser et al., 2016b). Therefore, we hypothesized that at least one component should have a globally positive spatial map across grey matter in both the resting-state and task fMRI and classified such components as noise. We also looked for components with significant white matter signal, venous signal, patterns that reflected vascular territories of the brain rather than functional networks, or other spatial patterns not compatible with neural signal.

To aid in classifying components whose spatial maps were not clearly consistent with signal or noise and to further explore the possible etiologies of the components, we computed several quantitative component-wise measures. One measure was each component’s direct temporal correlation with RVT as described above. Another was a globality index computed as the `abs(ln2(#PositiveGrayordinates / #NegativeGrayordinates))`. For each subject, we computed tICA component amplitudes, which were always defined as the standard deviations of the component timeseries, with this calculation sometimes confined to particular timeseries epochs. To relate each component’s association to physical head motion, we computed the difference in component amplitudes during periods of DVARS Dips compared to the non-DVARS Dips periods (`std(DipTimePoints)-std(NonDipTimePoints)`). We computed the variability of component amplitudes across subjects, the differences in component amplitudes between subjects who had been noted to be sleeping vs those who had not been noted to be sleeping, and components that were prominent in only one subject, run, or concatenated task session (i.e., where there was a large difference between highest component amplitude and second highest amplitude across runs or subjects). For task fMRI we also computed component amplitudes that increased during particular tasks relative to the other tasks in order to associate components with specific tasks. We searched for thresholds that best discriminated between components that had already been clearly identified as signal or noise based on spatial patterns so as to aid in classifying components where the spatial maps did not suggest an obvious classification. Additionally, we marked the few components where the classification ultimately remained uncertain or was disputed amongst the authors with the “Controversial” flag.

### 1.10 Temporal ICA-based Cleanup

sICA+FIX cleaned timeseries were further cleaned by removing the temporal-ICA components that were classified as noise (i.e., non-neural). The group concatenated tICA component timecourses were split according to single subject concatenated task sessions or resting state runs, and were then temporally regressed into the dense task or resting state fMRI timeseries data of each concatenated task fMRI session or resting state fMRI run to compute spatial beta maps. Then the tICA timecourses classified as noise and their associated beta maps were matrix multiplied to determine the portion of the dense timeseries that was best explained by the noise timecourses, and this noise dense timeseries was subtracted from the sICA+FIX cleaned timeseries data, producing the “sICA+FIX + tICA” cleaned dense timeseries data (see Equation #6 of Main Supplementary Information Section #3)^3^.

### 1.11 Generation of Greyplots

Greyplots, which display the timeseries intensities in grey scale using a compressed representation of space on the y axis and time on the x axis, have proven to be a useful method of visualizing the spatio-temporal structure of fMRI data and in particular have been used to highlight global fluctuations in timeseries data (Power, 2017; Power et al., 2014; Power et al., 2018; Power et al., 2017b). The greyplots used in the current study were generated for the data after each cleanup step by within-parcel averaging of the timeseries using the HCP-MMP1.0 multi-modal parcellation (Glasser et al., 2016a). This enables structured patterns in the data to be more easily seen (as unstructured noise is averaged out), but avoids obscuring features using unconstrained spatial or temporal smoothing of the data. Because simply displaying the 360 parcellated timeseries using one row per parcel would bias the resulting greyplot image towards smaller cortical areas and away from larger ones (by giving smaller areas relatively more space on the y axis than their size would dictate relative to an unparcellated greyplot), the smallest area (by surface area in mm) was assigned one row of the greyplot, and larger areas were assigned proportionally more rows based on surface area (subcortical grey matter was not included due to the lack of an areal parcellation of subcortical structures). Additionally, because semi-global signals will appear more global if randomly mixed along the spatial (i.e., y) axis of the greyplot, cortical areas were clustered according to the group-average full correlation resting state functional connectome (see Section #1.15 below) computed after tICA cleanup, so that areas with more similar timeseries are placed closer together. This also makes it easier to see the neurobiological structure in the data, as like rows will be averaged with like. All plots had these transformations applied to them identically, followed by Matlab’s default image downsampling to generate the final rasterized image.

### 1.12 Mean Grey Timecourse Regression

The global signal (across grey, white, and CSF), after sICA+FIX cleanup, is known to be a grey matter specific signal (see Supplementary Figure 5 in (Glasser et al., 2016b)). Furthermore, it is highly correlated with the mean grey signal (r=0.98+/-0.01 for the n=449 resting state subjects used in this study; see also (Power et al., 2014; Power et al., 2018) for a similar result). Additionally, the mean signal across parcels (Mean Parcel Timecourse) is very highly correlated with the MGT (r=0.99+/- 0.006). We used Mean Grey Timecourse Regression (MGTR)—the mean signal across grey matter within the CIFTI grayordinates standard space—for the task GLM and greyplot analyses, and the MPT—the mean signal across the parcels—for the parcellated connectome analysis, instead of the mean across the whole-brain mask (as has traditionally been done for global signal regression). For the group dense connectome and gradient analyses we regress the Mean PCA Series (MPS) out of the MIGP PCA series (see Section #1.14 below), which is a close approximation to MGTR on the full concatenated-across-subjects dense timeseries (Smith et al., 2014). We made these choices largely for computational convenience, as the different global timeseries are highly similar. We use the specific term MGTR in the methods and results of this study, but use the more general and well-known term Global Signal Regression (GSR) in the introduction and discussion when speaking about the technique in general.

### 1.13 Generation of Task fMRI Statistical Maps

After each cleanup methodology (including a without cleanup baseline) task fMRI dense timeseries data were analyzed as in (Glasser et al., 2016a) using an FSL-based (Woolrich et al., 2001) surface-enabled pipeline to produce mixed effects group z-statistical maps and intensity bias corrected beta maps for each task contrast. Cluster mass (the sum of above threshold z values multiplied by the surface vertex areas in mm) for each clean up approach and task contrast was computed using a Z=+/-5 threshold (roughly the two tailed Bonferroni corrected significance level across the 91282 grayordinate space, (Glasser et al., 2016a)). We analyzed 100 random subsets of 28 subjects from the 449-subject dataset (to match the sample size of the 28-subject registration optimization dataset previously used in a similar manner in (Glasser et al., 2016a)) to ensure that the effects of interest were reproducible across different subsets of subjects without increasing the z-stat values arbitrarily by using large numbers of subjects.

### 1.14 Generation of Dense Functional Connectomes and Gradients

After the sICA+FIX and sICA+FIX + tICA cleanup approaches, we generated dense functional connectomes from resting state fMRI data using the approach described in (Glasser et al., 2016a). This involved running MELODIC’s Incremental Group-PCA (MIGP), an iterative, distributed PCA data reduction algorithm (Smith et al., 2014). The purpose of MIGP is to provide a highly accurate approximation of the structured portion of the fully concatenated dense timeseries, as the vast majority of the information left out by MIGP is unstructured noise— see Main Supplementary Information Section #3. We did not compare conditions with and without sICA+FIX because it is already well established that sICA+FIX is strongly beneficial for HCP-Style resting state fMRI data (Griffanti et al., 2014; Salimi-Khorshidi et al., 2014; Smith et al., 2013a; Smith et al., 2013b). Because MIGP is a highly computationally intensive process, we analyzed the 210P data (however; see Glasser et al., 2016a, which showed that group resting state connectivity from 210P and 210V are highly correlated, r=0.98). Dense functional connectivity matrices were computed from this MIGP PCA series, and functional connectivity gradient maps were computed from the result for each cleanup approach as in (Glasser et al., 2016a).

### 1.15 Generation of Parcellated Functional Connectomes

Parcellated connectomes in Figure 12 were generated by averaging the dense MIGP PCA series within the parcels of the HCP-MMP1.0 multi-modal cortical parcellation and computing the full correlation values (i.e., standard Pearson correlation with no partialling). Parcellated connectomes in Figure 13 were generated in individual subjects by computing the full covariance values or Fisher Z-transformed regularized partial correlation values using FSLNets (ridge rho=0.23) and then averaging across subjects. The full correlation tICA-cleaned functional connectome was clustered using hierarchical clustering implemented in FSLNets (https://fsl.fmrib.ox.ac.uk/fsl/fslwiki/FSLNets) to order the cortical areas according to similarity of connectivity and enable grouping the areas into clusters having similar connectivity.

### 1.16 Data sharing

The data and annotations used in each brain image figure are stored as Connectome Workbench scenes and will be uploaded to the BALSA neuroimaging results database (http://balsa.wustl.edu) upon final publication. Scene-specific identifiers will be placed in each figure legend to allow easy previewing of scenes in BALSA and immediate download of individual scenes, complete scene files, or individual data files (Glasser et al., 2016b; Van Essen et al., 2017).

## Results

Although the issue of global structured noise is usually considered most problematic for resting state analyses, we will discuss task fMRI data first, as noted in the Introduction, because subjects’ underlying neural activity is explicitly manipulated by the task, and thus the task design provides a well-defined hypothesis about what a major portion of the subjects’ neural BOLD activation should look like, allowing objective tests of how well that hypothesis is matched by the data after differing cleanup approaches. In addition, subjects who are performing a cognitive task are less likely to fall asleep, and the scanner technicians were trained to halt the scan if a subject stopped performing the task. We will examine how task-induced neural BOLD activation manifests itself in temporal ICA, revealing important properties of this relatively unexplored method in fMRI data. We then compare and contrast task fMRI data with resting state fMRI data where we do not have the benefit of a prior hypothesis about the subjects’ neural BOLD activation and also where subject behavior is likely less well controlled. In this way, insights gained from the task-based analyses can directly inform our interpretation of the more challenging resting state analyses and we can draw parallels between findings in both types of data where possible.

Organizationally, within the task fMRI section (#2.1), we first explore the tICA components and their properties (2.1.1), then show the effect of tICA cleanup and MGTR on greyplots (2.1.2), and finally show the effects of sICA+FIX, sICA+FIX + tICA, and sICA+FIX + MGTR on the task GLM contrast maps themselves (2.1.3). We then discuss the implications of these results for task fMRI analyses and how they will inform our interpretation of resting state fMRI analyses (2.1.4). Within the resting state fMRI section (#2.2), we likewise first explore the tICA components and their properties (2.2.1), then discuss greyplots (2.2.2), followed by the effects of differing data cleanup methods on common resting state analyses (2.2.3).

### 2.1.1 Exploration of Temporal ICA Components in Task fMRI Data

We identified 70 reproducible temporal ICA components in the task fMRI data (as assessed by ICASSO) and ordered them approximately according to temporal variance explained (See Supplementary Task Components (TC)). The strongest of these components (6.7% of the tICA explained variance) was globally positive in grey matter (Figure 2, Supplementary Figure TC1), though with less intensity in regions with lower T2*-weighted signal intensity (i.e., gradient echo fMRI dropout regions), near zero in the white matter, and negative within the ventricles (see main Supplementary Information Section #5 for a hypothesis explaining the negative CSF signal of global or semi-global components regardless of whether they are non-neural or neural). Component TC1 also had the highest temporal correlation with the respiratory measure, RVT (r=0.30), of all 70 components (Panel A, Figure 3). It also exhibited higher amplitude during DVARS Dips (Panel B, Figure 3) and has high cross-subject variability in amplitude (Panel C, Figure 3). The grey matter specific spatial pattern and correlation with RVT of component TC1 suggests that it reflects global changes in grey matter blood flow from physiological sources, such as those arising from variations in breathing depth and rate, end tidal CO2, and/or heart rate (Birn et al., 2006; Chang et al., 2009; Chang and Glover, 2009; Golestani et al., 2015; Power et al., 2018; Power et al., 2017b). Though TC1 is correlated with DVARS Dips, this may be because motion may be more likely to co-occur with changes in respiration despite the fact that motion induces signal intensity changes in fMRI data via a different MR physics mechanism (s0 intensity mediated) than does respiration (T2* decay mediated) (Power et al., 2018; Power et al., 2017b). This component may partly or completely explain the global artifact that has been shown to persist after sICA+FIX in HCP resting state fMRI data (Burgess et al., 2016; Power, 2017; Power et al., 2017b; Siegel et al., 2017). Indeed its spatial pattern reflects what many practitioners of global signal regression believe that they are removing from their data (though see Supplementary Figure 25, below Figure 13, and Supplementary Figures 5 and 6 from (Glasser et al., 2016b) for what is actually removed by GSR).

**Figure 2.**
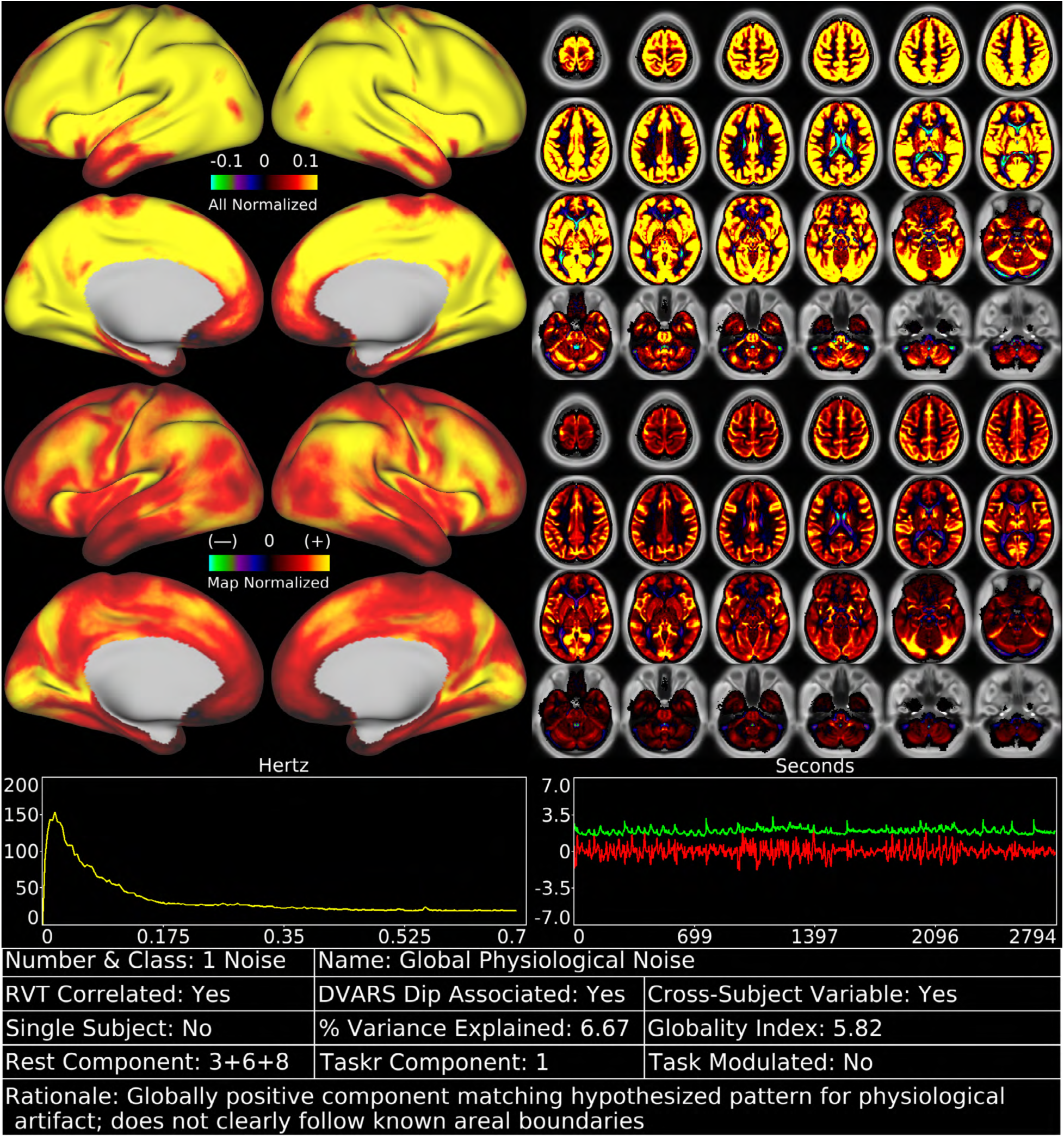
shows the first task fMRI tICA component in a standardized display format (see also Supplementary Figures TC 1-70). The top row of data has the color scale normalized and held constant across all 70 components (in percent BOLD), whereas the second row has the color scale set independently for each component. Hence the first view allows components to be compared with each other on the same scale, whereas the second view highlights patterns in the map of each specific component and is scaled between 2% and 98%. The left chart in the third row indicates the power spectrum of the component averaged across subjects, and the right chart indicates the average timeseries (red) and average absolute value of the timeseries (green). Both of these will show evidence of the task stimulus for task-modulated components because of consistent task timing across subjects. Additional information about the component is provided in the table along the bottom row (see Figures 3 and 4 for thresholds that determine Yes/No status, which is also sortable in the Supplementary Component Data Table). The rationale for classifying the component is listed along the bottom row of the table. Supplementary Figure 4 shows how the task fMRI runs were concatenated. Percent variance explained is computed using the ‘total variance’ of the data at the tICA modeling stage (i.e., after sICA+FIX cleaning and the d=137 sICA dimensionality reduction, and thus sums to 100% across all the tICA signal and noise components). The globality index is abs(ln2(# positive grayordinates/# negative grayordinates)).

**Figure 3.**
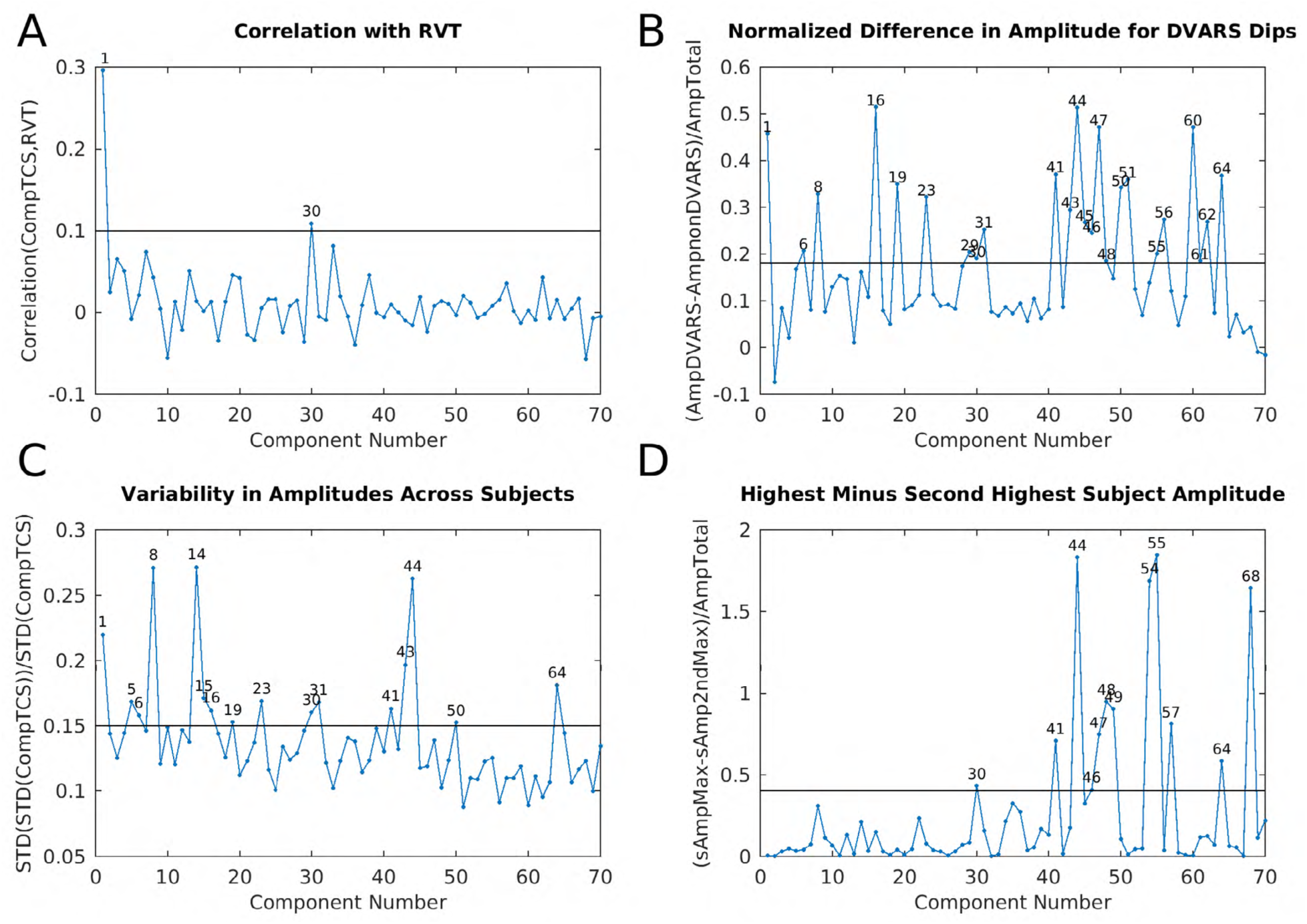
shows four plots that were helpful during task fMRI component classification into signal and noise. Panel A is the correlation between the component timeseries and RVT for the 70 concatenated task sessions that had the top 10% mean correlation between RVT and their parcellated fMRI data. This selects for subjects having both good quality RVT traces and substantial respiratory contamination of their data (see Methods Section #1.6). The line is at r=0.1. Panel B is the difference in component amplitude (standard deviation of the component timeseries) between frames with DVARS dips and those without DVARS dips normalized by the component amplitude across all frames (see Methods Section #1.9), so as to highlight those components that have stronger temporal fluctuations during DVARS dips. The line is at 0.18. Panel C shows the variability of component amplitudes across subjects normalized by the overall amplitude of each component. The line is at 0.15. Panel D shows the difference between the maximum subject’s component amplitude and the next highest subject’s component amplitude normalized by the overall amplitude of each component. This measure highlights those components that are particularly strong in a single subject. The line is at 0.4. The discriminatory thresholds in this figure and in Figure 7 and Supplementary Figure 16 were chosen as described in Methods Section #1.9. Those components above each threshold are numbered on each graph.

Together with the global noise component, we identified a total of 25 components having properties consistent with structured noise (Supplementary Figures TC1, 8, 30, 38, 41-52, 54-57, 59-60, 62, 64, and 68), which together accounted for 26.9% of the variance in the data at the tICA modeling stage (i.e., of the concatenated d=137 sICA component timeseries). (Notably this is much less than what is removed by sICA+FIX from the task fMRI data where 89% of the structured variance was noise). Noise components include those with similar patterns to the main global physiological component (Supplementary Figures TC30, 38, and 52), substantial extensions into white matter (Supplementary Figures TC8, 30, 42, 50-52, 60, and 64), inclusion of veins (Supplementary Figures TC8 and 62), similarity to movement regressor beta maps (Supplementary Figure TC51), high values around brain margins (Supplementary Figures TC38, 44, 46, 47, 48, and 52), similarity to known image reconstruction artifacts (Supplementary Figures TC41, 43, 45, 49, 54, 55, 56, and 59) or receive coil instabilities (Supplementary Figures TC50, 60, and 64), banding patterns (Supplementary Figures TC 56), amplitudes that are correlated with DVARS Dips (Supplementary Figures TC8, 30, 41, 43-48, 50, 51, 55, 56, 60, 62, and 64; 3 Panel B), and prominence only in one subject or one run (Supplementary Figures TC30, 41, 44, 46-49, 54, 55, 57, 64, and 68; 3 Panel D). While sICA+FIX classification performance exceeds 99% accuracy on HCP data (Griffanti et al., 2014; Salimi-Khorshidi et al., 2014) some spatially specific noise does slip through (either due to the rare misclassifications or due to being in the null subspace of the sICA+FIX dimensionality reduction, i.e., “error” term in Eq. (1) for each sICA+FIX run), and this structured noise is likely the source of some of these tICA components. As shown below, this residual spatially specific structured noise is removed along with the global structured noise by tICA cleanup.

The other 45 temporal ICA components had properties consistent with neural signal (Supplementary Figures TC2-7, 9-29, 31-37, 39, 40, 53, 58, 61, 63, 65-67, 69-70) and accounted for the other 73.1% of the tICA explained variance. The upper panel in Figure 4 shows the difference in amplitude for each task relative to all other tasks, with each of the 7 tasks in a different color, revealing the components that are specifically modulated by particular tasks. The amplitudes of 18 of the signal components were clearly modulated by a specific task (being stronger in one or more tasks than the other tasks, see Supplementary Figures TC2-5, 7, 9, 11, 13, 16, 18, 19, 21, 24, 28, 31, 33, 37, 58) and most of these were not present in the resting state (TC2-3, 7, 11, 18, 21, 24, 28, 33, 37, 58). In striking contrast, none of the 25 noise components’ amplitudes were clearly modulated by a specific task. Many of these 18 task-modulated components look very similar to their corresponding task fMRI GLM contrast beta maps (Supplementary Figures 5 6 7 8 9 10 11 12 13 14 15), which is to be expected if temporal ICA is sensibly decomposing the task fMRI data into neurobiologically meaningful temporally orthogonal components.

**Figure 4.**
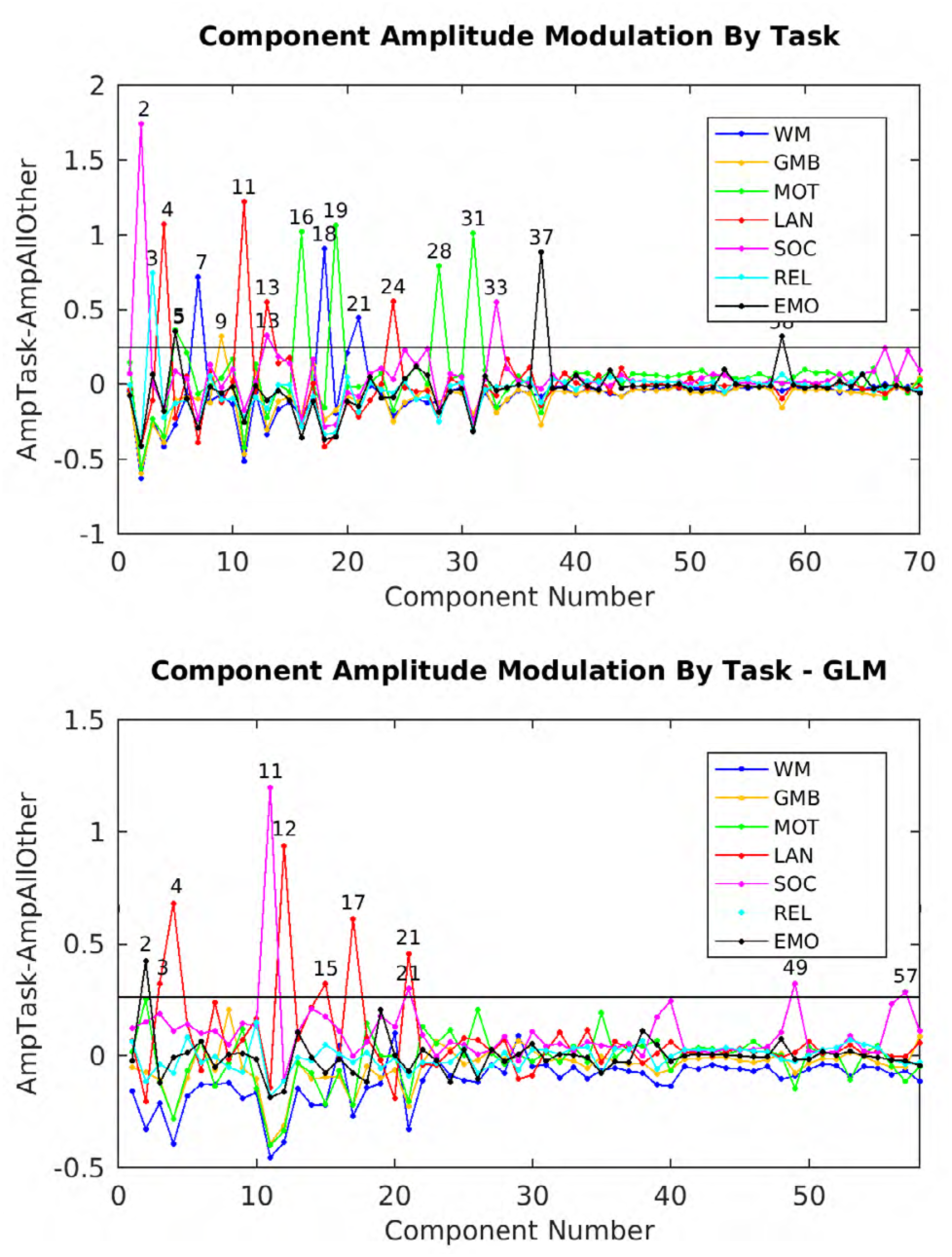
shows the component amplitudes modulated by task. The measure is the amplitude of a component (standard deviation over time) during a given task versus all other tasks (`std(SpecificTask)-std(AllOtherTasks)`). Task abbreviations are WM=Working Memory, GMB=Gambling, MOT=Motor, LAN=Language, SOC=Social, REL=Relational, and EMO=Emotion. The top panel shows the original task component amplitudes (70) whereas the bottom panel shows the component amplitudes after regressing out the task GLM (58 reproducible components). The line is at 0.25 in both cases.

A supplementary analysis was performed on the task fMRI data by rerunning weighted regression and temporal ICA on the residuals after fitting HCP’s task fMRI design matrix. 58 reproducible temporal ICA components were found (See Supplementary Task residual Components (TCr)). 38 were classified as signal, accounting for 71.7% of the tICA explained variance, and 20 were classified as noise, accounting for the remaining 28.3% of the tICA explained variance, with TCr1, the main global noise component, accounting for 7.9% of the tICA explained variance (Supplementary Figure 16, Supplementary Figures TCr1-58). Ten of the signal components’ amplitudes remained modulated by specific tasks (Figure 4, lower panel), suggesting that the residuals after the task design fitting still contain some task-driven effects in the underlying spontaneous fluctuations (i.e. that the task design is an imperfect model of the task driven neural activity). Additionally, some of these residually task-modulated components are not present during the resting-state (TCr4 and TCr11).

Five of the 45 signal components (Supplementary Figures TC6, 16, 19, 29, and 31) in the main task fMRI temporal ICA analysis represent topographically organized sensori-motor networks for the head (TC16), right hand/upper extremity (TC6), left hand/upper extremity (TC31), eyes/neck/trunk (TC29), and the feet/lower extremities (TC19). These components all include nodes in the primary sensori-motor cortex (M1 and S1), the supplementary sensory cortex (SII), the insular cortex, the supplementary and cingulate motor cortices, superior and inferior cerebellar motor areas, the thalamus, and the striatum (in addition the head sensori-motor component also appears to include brainstem cranial nerve nuclei). Notably, the hemispherically lateralized hand networks (TC6, 16) show correct hemispheric specificity (contralateral to the body part that they control, for all nodes except the cerebellum, which has ipsilateral nodes). All but the right hand sensori-motor network (TC6) and the eye/neck/trunk network (TC29) are specifically modulated by the MOTOR task. That the right hand motor component is not specifically modulated by only the motor task is expected because the right hand was used in the other tasks for button box pressing and thus was likely heavily engaged in all of them. The motor task also did not explicitly ask the subject to move their eyes, neck, or trunk. There is one additional predominantly cortical network (Supplementary Figure TC23) that spans all of S1/M1, SII, M2 and auditory cortex that is not specifically modulated by a task.

All six of these sensori-motor networks have higher amplitude during DVARS dips, a property shared with only one other signal component (Supplementary Figure TC61), but with many of the noise components. Thus, these sensori-motor networks have two interesting properties in common: 1) they are modulated by a MOTOR task and 2) they have higher amplitude during DVARS dips, which are usually the result of head motion (see Methods Section #1.7 and Supplementary Topic #4). Indeed, the MOTOR task has the highest rate of DVARS dips per frame of all the tasks at 0.029 dips per frame vs the average across other tasks of 0.014 dips per frame (resting state has 0.013 dips per frame). We can thus infer that during at least some periods of DVARS dips, subjects’ sensori-motor networks have higher BOLD signal amplitudes than usual. These results lead to the conclusion that we should not assume that the neural signal of subjects during DVARS dips is the same as during non-dip periods. The impact of these findings on the metrics that we should use for assessing data cleanup in this study is considered in the supplementary discussion (Supplementary Information Topic #6), and the impact on some study designs is considered in the main text discussion.

Several other interesting properties are present within the 45 signal components of the task fMRI temporal ICA analysis. Many networks that are reminiscent of canonical resting state networks are both present and less modulated by specific tasks. This includes components representing the default mode network (Supplementary Figures TC5, 12, 25-27, 40, 53, and 61), the fronto-parietal network (Supplementary Figures TC12, 32) the cingulo-opercular network (Supplementary Figure TC10), the language network that closely resembles recent reports (Glasser et al., 2016a; Spronk et al., 2017)(Supplementary Figure TC22), and the visual system (Supplementary Figure TC34). On the other hand, some signal networks do not clearly match canonical resting state networks, e.g., Supplementary Figures TC35, 36, and 39 (Cerebellum), 63 (left vs right network), and 66 (extra-striate visual network).

Interestingly, four signal networks show visuotopic organization with respect to polar angle, with positive vs. negative separations across the horizontal (Supplementary Figures TC69 and 70) and vertical meridians (Supplementary Figures TC65 and 67). These networks also have some visuotopic organization relative to eccentricity (foveal vs peripheral), as do three other signal components (Supplementary Figures TC15, 17, and 58). We suspect that these temporal ICA components underlie the ability to extract visuotopy from fMRI data that does not contain an explicit visuotopic task (Figure 8 in the Supplementary Methods and Figures 3, 4, 5, and 6 of the Supplementary Neuroanatomical Results, both from (Glasser et al., 2016a).

Finally, one component (Supplementary Figure TC 14) highlights specific interactions between V1 and the LGN that may account for the ability to sharply delineate area V1 using functional connectivity in our prior parcellation (Figure 2 in the Supplementary Neuroanatomical Results in (Glasser et al., 2016a)), as the V1 boundary is not evident in any of the other components. We suspect that this component is related to whether a subject’s eyes are open or closed, as identifying V1 as a separate parcel using automated winner-take-all approaches depends on whether a subjects’ eyes are open or closed (Laumann et al., 2015). Interestingly, this component is variable across subjects, perhaps reflecting differences in subjects’ eyes-open vs eyes-closed behavior in the scanner despite the presence of a task. Other signal components that showed high cross subject variability included the left hand sensori-motor component (TC31), the pan-motor and auditory component (TC23), and the strongest default mode component (TC5).

We also correlated the component amplitudes (temporal standard deviation of each component in each concatenated task session—in this case an entire tfMRI session on day 1 or day 2 for each subject). For each component, there were therefore 898 (2 * 449 subjects) amplitudes, and those amplitudes were correlated across tICA components. We then performed hierarchical clustering on the resulting correlation matrix (Supplementary Figure 17). In general, signal components cluster together with other signal components and noise components cluster with other noise components. Not surprisingly, components that are strongly modulated by task and components whose tasks occurred on a particular day cluster together, and are anti-correlated with the components whose tasks occurred on the other day, likely because they have strong amplitudes on one day and that are near zero on the other (and after removing the mean it is obvious how they will be anticorrelated). Between these are spontaneous components that are generally not specifically task modulated. For the residuals after regressing out the task designs, again signal and noise components tended to cluster together (Supplementary Figure 18). Components with residual task modulations tend to cluster together and spontaneous components generally form a separate cluster.

Finally, we computed the beta (effect size) of each tICA component on the mean grey timecourse after sICA+FIX cleanup to show which components make the greatest contribution to the global signal. Not surprisingly, the global components indeed are the biggest contributors (Supplementary Figure 19). Additionally in the same figure, we show the effect of tICA cleanup on the mean grey timecourse variances.

### 2.1.2 Effects of Temporal ICA Cleanup and MGTR on Task fMRI Greyplots

Figure 5 shows mean grey signal traces and parcellated greyplots of two concatenated task sessions of two subjects having particularly high global noise, after sICA+FIX under three cleanup conditions: after sICA+FIX only, after sICA+FIX plus temporal ICA cleanup (sICA+FIX + tICA), and after sICA+FIX plus MGTR (sICA+FIX + MGTR). Regressing out the components identified as noise removes the obvious bands in the greyplots (and the corresponding substantial mean grey signal fluctuations) that have been previously associated with respiratory changes (Power et al., 2018; Power et al., 2017b). Importantly, regressing out the noise temporal ICA components is not the same as removing the mean grey signal, as the mean grey timeseries still retains some fluctuations after temporal ICA cleanup, unlike (by definition) with MGTR. In task fMRI data, the variance removed by regressing out the mean gray timecourse (MGTVar) averaged across all subjects before sICA+FIX cleanup is 2161; after sICA+FIX it is 482, and after temporal ICA cleanup it is 235 (on data scaled to a grand mean across the volume of 10,000). Thus, in task fMRI data, the neural global signal variance is 11% (235/2161) of the original global timecourse variance and 49% (235/482) of the global timecourse variance after sICA+FIX. Given that there is minimal global signal in the unstructured noise (see above) and that the structured signal remaining is from temporal ICA components classified as signal, we feel that this is the best available estimate of the true neurally-related global signal variance. Across subjects, the variance of the mean grey timecourse (MGT) itself is substantially reduced by tICA cleanup (Supplementary Figure 19).

**Figure 5.**
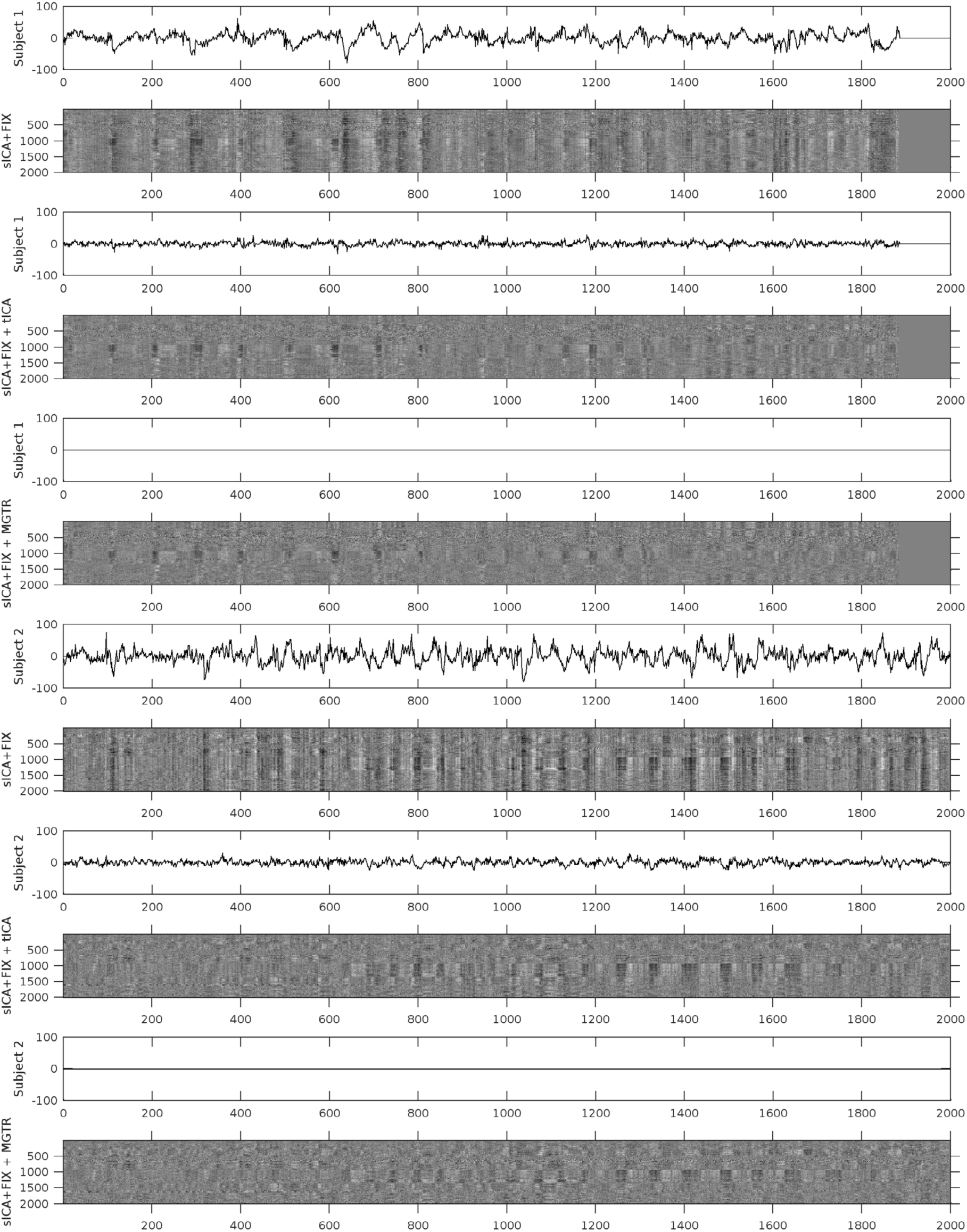
shows the mean grey signals and parcellated greyplots of two concatenated task fMRI timeseries from two subjects after sICA+FIX (Rows 1-2, 7-8), sICA+FIX + tICA (Rows 3-4, 9-10), and sICA+FIX + MGTR (Rows 5-6, 11-12). Note that concatenated session 1 is shorter than concatenated session 2 and so the first 3 rows include zero-padding on the far right. The data were parcellated as described in the methods, then displayed according to parcel surface area (with 1 row assigned for the smallest parcel and proportionally larger numbers of rows assigned for larger parcels such that there are more than 360 rows). Additionally, the data are ordered by hierarchical clustering of the group full correlation parcellated connectome so that parcels with more similar timeseries across the group are closer together (see the bottom row of Figure 13 which shows the “cognitive/task negative vs non-cognitive/task positive split that forms the primary clustering split about half way down the y-axis in the grey plots). tICA cleanup removes the vertical “stripes” (signal deviations of the same sign across the whole brain) from the greyplots that were present after sICA+FIX and their corresponding MGT fluctuations, but without removing the entire MGT as occurs with MGTR. The greyplots after tICA cleanup and MGTR look similar but not identical. The greyscale ranges from −2% to +2% BOLD.

### 2.1.3 Effects of sICA+FIX, tICA, and MGTR Cleanup on Task fMRI GLM Contrast Maps

We found a clear benefit in statistical sensitivity from cleaning task fMRI data with sICA+FIX with a median of 28/40 (70%; robust range of 22-32, see Figure 6 legend) contrasts increasing in cluster mass vs standard analysis across the 100 sub-sampled analyses, clearly above the 50% (20/40) chance line (Figure 6, Panel A). The median improvement in cluster mass was 12% (Figure 6, Panel B). Cluster mass did not increase for all task contrasts, however, and for these task contrasts, stimulus-correlated noise likely led to biases or false positives in the group effect size maps prior to sICA+FIX cleanup. For example, Supplementary Figure 20 shows a neurobiologically implausible false-positive deactivation in orbitofrontal cortex during tongue movement that is removed by sICA+FIX cleanup, and Supplementary Figure 21 shows a negative bias in the Language STORY contrast that is also removed (see the strongly negative CSF), revealing a positive activation that would otherwise have been near zero (due to the negative bias). The false positive deactivation during tongue movement is likely caused by stimulus-correlated head motion given its localization to orbitofrontal cortex, a site of strong MR susceptibility vs movement interaction (Griffanti et al., 2017) where the region of fMRI signal dropout changes with head motion. The language story contrast negative bias is likely caused by failure to reach T1 steady state during the baseline period, which happens to be located only at the very beginning of the language task, and which affects bright CSF signal most.

**Figure 6.**
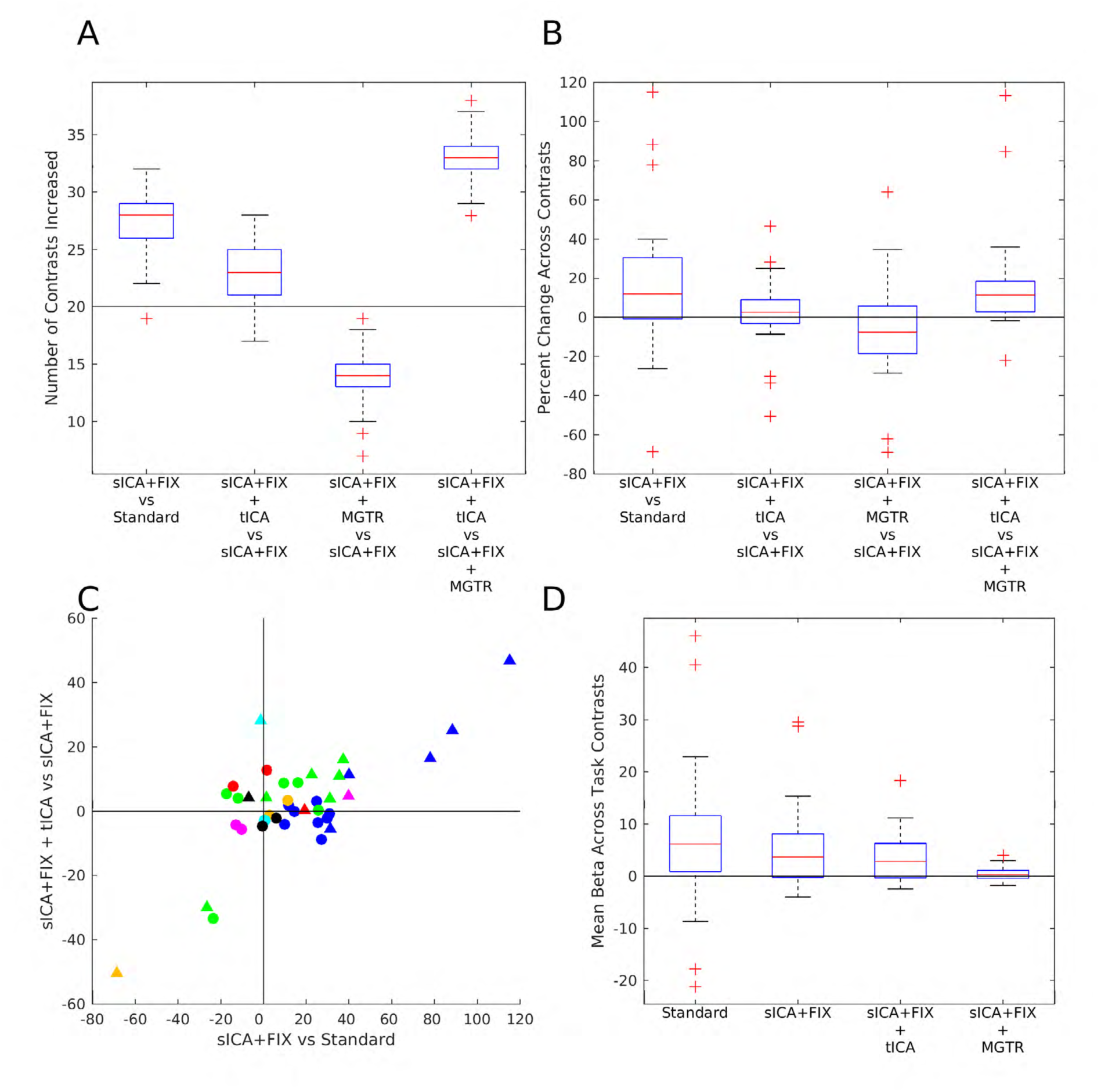
compares the statistical sensitivity (Panels A, B, and C) and mean beta values (Panel D) across contrasts and cleanup approaches. Statistical sensitivity was quantified via the cluster mass using a Z=+/- 5 threshold. Panel A shows the number of contrasts found to have increased cluster mass using a box and whisker plot for sICA+FIX vs Standard, sICA+FIX + tICA vs sICA+FIX, and sICA+FIX + MGTR vs sICA+FIX, and sICA+FIX + tICA vs sICA+FIX + MGTR for 100 random subsets of 28 subjects from the 449 total subjects. Only primary contrasts that are not averages of other primary contrasts and differential contrasts are plotted with no negative duplicates (n=40 contrasts out of the total of 86 released by the HCP). The Gambling REWARD-PUNISH contrast has minimal neural signal (Glasser et al., 2016a) and thus acts as a negative control (golden triangle in (C)). The red line is the median, the edges of the box are the 25^th^ and 75^th^ percentiles, the whiskers are at the data point closest to +/- 2.7 standard deviations (the robust range), and the outliers (+’s) are the data points beyond these thresholds. The horizontal black line is at 20 (of 40 total) contrasts improving (50%). Panel B shows the percent change in cluster mass for the tfMRI contrasts for the same comparisons, with a horizontal black line at 0% change (the percent change for each contrast is the average across 100 random subsets and the boxplot shows the distribution across contrasts). Panel C shows a scatter plot of the percent improvement of cluster mass from sICA+FIX over standard processing vs the improvement of sICA+FIX + tICA over sICA+FIX processing. Circles are primary contrasts and triangles are differential contrasts. The colors are the same as used in Figure 4 to represent the different tasks. Panel D shows the spatial means across the entire contrast beta maps for standard processing, sICA+FIX, sICA+FIX + tICA, and sICA+FIX + MGTR.

We next compare the effects of the two global noise cleanup methods, our tICA-based method and MGTR, both after sICA+FIX has already been applied. tICA cleanup did not clearly benefit or harm statistical sensitivity overall, with a median 23/40 (58%; robust range of 17-28) of contrasts improving, overlapping with the 50% chance line (Figure 6, Panel A). The median improvement in cluster mass for tICA cleanup across all contrasts was 3%. However, “primary” vs baseline (circles) and “differential” vs another contrast (triangles, Figure 6, Panel C) contrasts do not behave in the same way. Differential contrasts between task “on” periods will generally have similar physiological noise across the two periods being compared, as the subject will be engaged in task behavior in both cases (e.g., working memory of Tools vs Faces) and so differential contrasts will not typically be biased by physiology (see below). Indeed, we found that tICA cleanup improves 13/16 (81%) differential contrasts, with a median improvement in cluster mass of 8%. On the other hand, primary contrasts vs baseline are much less likely to improve 10/24 (42%), with a median change in cluster mass of −1%, as they are vulnerable to physiological noise biases arising from differences in physiology between task “on” and task “off” periods (e.g., respiration: the correlation for task “on” blocks vs the mean RVT across subjects is r=0.56, see Supplementary Figure 22). An extreme example is the 33% cluster mass reduction of the Motor CUE contrast (Supplementary Figure 23). Thus, we suspect that many of the statistical sensitivity decreases found with tICA cleanup are actually removal of physiological-noise-induced biases of different magnitudes, depending on the vulnerability of the task contrast type to such biases and how much the task modulates subject physiology between task on and task off blocks.

Contrary to tICA cleanup, applying MGTR after sICA+FIX clearly reduces statistical sensitivity across a majority of task contrasts, with a median of only 14/40 (35%, robust range of 10-18) contrasts improving, which is clearly below the 50% chance line (Figure 6, Panel A). The median cluster mass decrease is 8.0% (Figure 6, Panel B). More importantly, a median of 33/40 (83%; robust range of 29-37) of contrasts are higher with sICA+FIX + tICA than sICA+FIX + MGTR, clearly above the 50% chance line (Figure 6, Panel A). In addition, there is no predilection for differential or primary contrasts to be better with tICA cleanup vs MGTR (primary 20/24, 83%, differential 13/16, 81%), which argues against any effect of additional physiological noise removal by MGTR, but would be consistent with removal of some task-correlated neural signal across the board by MGTR. sICA+FIX + tICA has a median cluster mass that is 11% higher than sICA+FIX + MGTR (Figure 6, Panel B, primary contrasts are median 13% higher and differential contrasts are median 9% higher). Those few contrasts that are higher with MGTR may be because of removing task uncorrelated neural signal to a greater extent than task correlated neural signal (see below). MGTR also consistently shifts the mean of the activation beta maps across all tasks to be near zero, something that is highly improbable as a neurobiological ground truth (Figure 6, Panel D) and that does not occur with sICA+FIX and sICA+FIX + tICA. Indeed, if we compare the effect of tICA cleanup to the effect of MGTR (Supplementary Figure 25), we see that tICA cleanup removes a global positive bias that is highly spatially correlated with TC1 (r=0.93) and only modestly spatially correlated with the task activation pattern of interest (r=0.44, note that the Motor CUE activation map itself is semi-global, unavoidably leading to some correlation). On the other hand, MGTR removes a network-specific effect that is highly spatially correlated with the task activation pattern of interest (r=0.85). Thus, MGTR evidently removes a portion of the expected neural signal from fMRI data, in contrast to tICA cleanup.

We explicitly quantified the effect of MGTR after sICA+FIX + tICA and found that 26% of the global neural signal variance is task related and 55% of the overall neural signal variance is task related (see Main Supplementary Information Section #3 for details). The proportion of overall neural signal and global neural signal explained by the task GLM does vary across tasks, however, with the Working Memory task being highest in terms of overall neural signal explained at 70% overall / 31% global, then the Motor task at 67%/34%, the Gambling task at 53%/36%, the Relational task at 53%/36%, the Social task at 51%/35%, the Emotion task at 46%/13%, and the Language task at 41%/10%. The overall proportion of the neural signal that is explained by the task GLM will vary based on how well the task drives the BOLD fluctuations (to the exclusion of, or over and above the spontaneous fluctuations), how widespread the task induced fluctuations are spatially, and how well the task GLM model explains the task induced variance, which as noted above is incomplete. The global neural variance should depend on how much synchronous activity the task generates across the brain and the amplitude of this activity (particularly in the early non-cognitive/task positive areas that generate most of the global neural signal). The portion of the overall neural signal variance that is global in task fMRI data is 10% (i.e., the portion removed by MGTR after sICA+FIX and tICA cleanup).

### 2.1.4 Implications of Task MRI Cleanup Results for Task and Resting State Analyses

sICA+FIX is clearly beneficial for both task fMRI and resting state fMRI data, both improving statistical sensitivity and removing biases from stimulus correlated spatially specific artifacts in the case of task analyses and removing these biases in the case of correlational resting state analyses (Griffanti et al., 2014; Salimi-Khorshidi et al., 2014; Smith et al., 2013a; Smith et al., 2013b). Temporal ICA cleanup has modest benefits in the form of increasing statistical sensitivity for differential contrasts and removing biases from primary contrasts due to stimulus-correlated physiology that is consistent across subjects. Thus, the more conservative approach would be to clean task fMRI data with tICA. The effects of tICA cleanup are very different from the effects of MGTR, where in addition to removing global noise, neural signal is also removed, leading to clear statistical sensitivity decreases relative to sICA+FIX + tICA across both primary and differential contrasts (Figure 6 Panels A and B), removing a network-specific pattern of signal highly correlated with the task activation map (Supplementary Figure 25, and biasing the means of the contrast beta maps towards zero (Figure 6 Panel D). Thus, temporal ICA cleanup produces modest benefits for task fMRI data, particularly when stimulus correlated global physiological noise is present, but it does not remove task correlated neural signal as MGTR does.

Some of these properties of global signal regression (or similarly mean stabilization) have been previously reported for task fMRI data (Aguirre et al., 1997; Aguirre et al., 1998; Macey et al., 2004; Zarahn et al., 1997), but they did not prevent the subsequent widespread use of global signal regression as a cleanup technique for resting state fMRI data (perhaps because the effects were perceived as modest and the problems from global noise are thought to be much larger for resting state fMRI than for task fMRI). Correlational resting state analyses are analogous to task analyses that have substantial stimulus correlated global physiological noise. Unlike task fMRI, where there is at least the possibility that the task design will be uncorrelated with the global physiological noise, such physiological noise will always maximally impact univariate resting state analyses because there is no task design (model) against which the fMRI timeseries can be explicitly compared. As a result, any additive global or semi-global noise will positively bias correlations, and these biases may differ across subjects or groups, which is why global structured noise is considered such a pernicious problem for resting state fMRI.

### 2.2.1 Exploration of Temporal ICA Components in Resting State fMRI Data

We identified 84 reproducible (as assessed by ICASSO) temporal ICA components in the resting state fMRI data (See Supplementary Resting state Components (RC)). Three components were globally positive across the cerebral cortex, and two of those three were globally positive throughout the grey matter (the other was largely negative in the cerebellum; Supplementary Figures RC3, 6, and 8). Together, the three global components accounted for 8.5% of the tICA explained variance. Like the single global component in the task fMRI data, CSF was negative (see discussion on anti-correlations of CSF in global or semi-global components in Main Supplementary Information Section #5), and white matter was near zero. Interestingly, the two globally positive components (RC6 and 8) had some left/right asymmetries and were associated with the phase encoding direction of the data (Supplementary Figure 24). As with the task fMRI data, these global components were particularly correlated with RVT (Panel A, Figure 7), were higher in amplitude during DVARS dips (Panel B, Figure 7).

**Figure 7.**
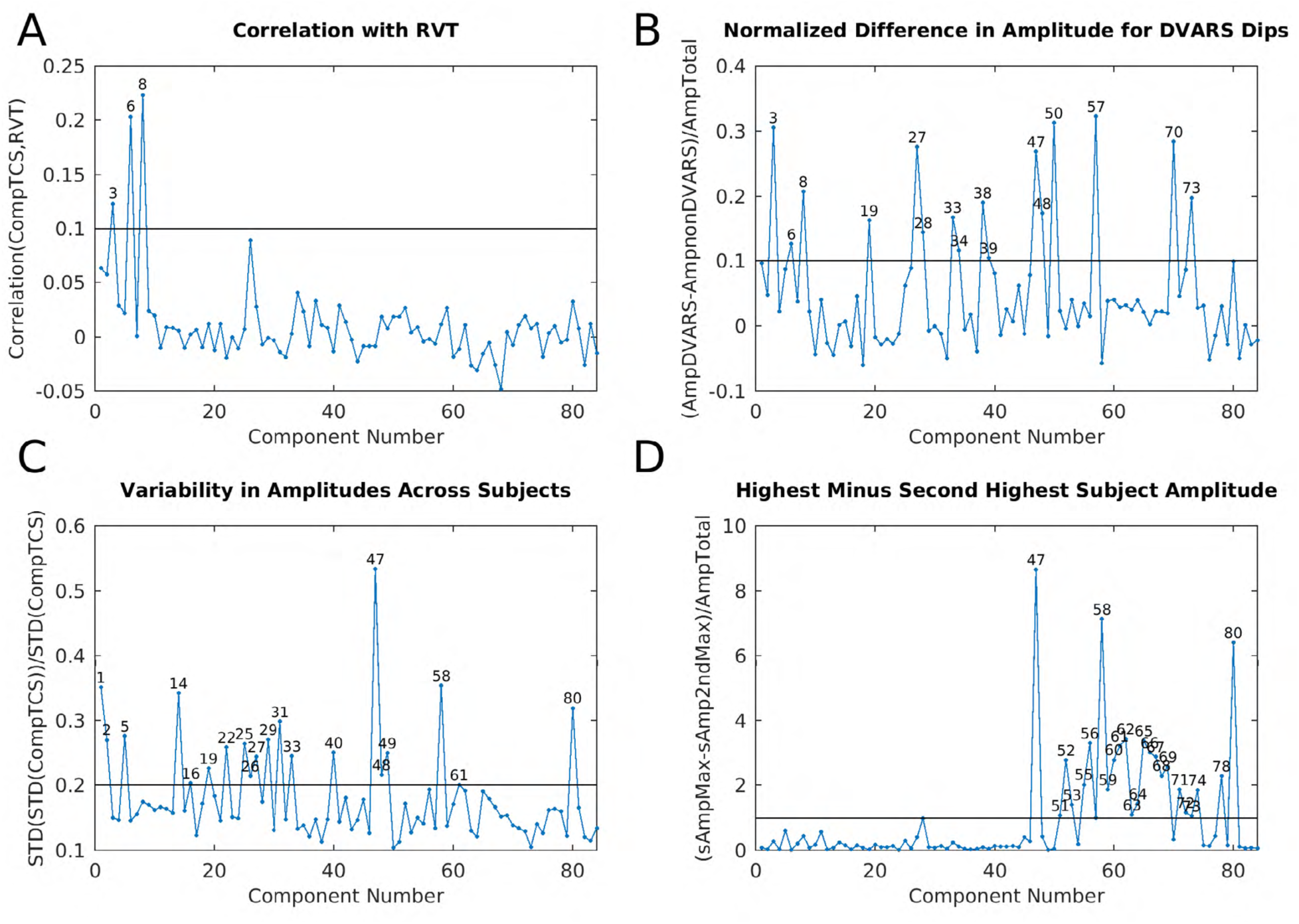
shows four plots that were helpful in classifying the resting state fMRI tICA components into signal and noise. Panel A shows the correlation between the component timeseries and RVT for the 157 runs with the top 10% mean correlation between RVT and their parcellated timeseries as in Figure 3. When the 3 global components timeseries (RC3, 6, and 8) are added together, the correlation with RVT is 0.28, very similar to the correlation of the single task global component (0.30) with RVT. The line is at r=0.1. Panel B is the difference in component amplitude (standard deviation of the component timeseries) between frames with DVARS dips and those without DVARS dips normalized by the standard deviation of each component across all frames as in Figure 3. The line is at 0.1. Panel C shows the variability of component amplitudes across subjects normalized by the standard deviation of each component as in Figure 3. The line is at 0.2. Panel D shows the difference between the maximum subject’s component amplitude and the next highest subject’s component amplitude normalized by the standard deviation of each component as in Figure 3. The line is at 1.0.

Together with these three global noise components, we identified a total of 32 components with properties suggestive/indicative of noise, which together with the global components accounted for 28.4% of the variance in the data at the tICA modeling stage (i.e., of the concatenated d=137 sICA component timeseries). (Notably this is much less than what is removed by sICA+FIX from the resting state fMRI data where 93% of the structured variance was noise). Noise components include those with substantial extensions into white matter (Supplemental Figures RC19, 25, 28, 44, 47, 50, and 70), inclusion of veins (Supplemental Figures RC19, and 28), similarity to movement regressor beta maps (Supplemental Figure RC50), high values around brain margins (Supplemental Figure RC57), similarity to known receive coil instabilities (Supplemental Figure RC47, 72, and 73) or image reconstruction artifacts (Supplemental Figures RC46, 53, and 64), higher amplitudes during DVARS dips (Figure 7, Supplementary Figures RC19, 28, 34, 38, 47, 50, 57, 70, and 73), and/or prominence in only one subject or run (Panel D, Figure 7, Supplemental Figures RC47, 51-53, 58-64, 66-68, 71-73, 78, and 80). Many of the noise components were similar to those found in the task fMRI data (Supplementary Figures TC/RC 1/3+6+8, 8/19, 8/28, 42/44, 43/46, 50/47, 51/50, 44/57, and 64/70) showing that much of the structured noise is common to resting state and task fMRI data. As with the task fMRI data, sICA+FIX’s classification accuracy is very good in HCP data (99%), but not perfect, and some noise does slip through. (Additionally, as before, some weakly structured noise may exist in the unstructured noise subspace of sICA+FIX that is only detectable at the group level). In fact, some of the noise found in an initial temporal ICA analysis was traced back to misclassification of a few sICA+FIX components in a few subjects and was fixed using manual reclassification of these components prior to the HCP 1200 subject release.

The other 52 temporal ICA components were consistent with neural signal (Supplementary Figures RC1, 2, 4, 5, 7, 9-18, 20-24, 26, 27, 29-33, 35-37, 39-43, 45, 48, 49, 54-56, 65, 69, 74-77, 79, and 81-84), and together accounted for 71.6% of the tICA explained variance. 20 of these components had spatial patterns that visually corresponded to patterns seen in the task fMRI analysis (Supplementary Figures TC/RC 14/2, 23/5, −9/9, 5/10, 4/12, 10/13, 26/15, 17/16, 22+36/20, 16/27, 6/33, 13/37, 29/39, 31/40, 34/41, 19/48, 65/76, 70/81, −67/82, 69/84; a negated number means that the sign of the component was reversed, but note that the signs of ICA components are arbitrary). Particularly notable correspondences were the five somatotopically organized sensori-motor networks, which retained the property of being the only signal components associated with DVARS dips (though the left hand component (RC40) was not formally labeled as “DVARS Dip Associated” because it did not exceed the threshold, Supplementary Figures RC27, 33, 39, 40, 48). The pan-sensori-motor component that was seen in the task fMRI data was also found in the resting state (Supplementary Figures TC23/RC5); however, it was much stronger, explaining 2.87% of the variance vs 1.57% in task data.

Visual components with task/rest homologues included the four visuotopic components with a polar angle relationship (Supplementary Figures TC/RC 65/76, 70/81, 67/82, and 69/84) and two with an eccentricity relationship (Supplementary Figures TC/RC 17/16 and 34/41). Five additional visuotopically organized components were also found (Supplementary Figures RC29, 42, 49, 77, and 83). The component associated with eyes open vs closed in the task fMRI data (1.95% of explained variance, Supplementary Figure TC14; (Laumann et al., 2015)), was the second strongest component in the resting state fMRI data (3.64% of explained variance, Supplementary Figure RC2), and was variable across subjects, suggesting (not surprisingly) that many subjects actually closed their eyes for part of the resting state scans despite instructions to keep them open and stare at the fixation cross-hairs.

Aside from the somatotopically organized sensori-motor components that were modulated by task fMRI and the primary default mode component (Supplementary Figure TC/RC 5/10) that is slightly modulated by the MOTOR and EMOTION tasks, no other resting state fMRI components closely matched task components that were modulated by task, though there were some similarities (e.g., the fronto-parietal network Supplementary Figure TC/RC 4/12, a subsidiary default mode network TC/RC −9/9 (inverted), and the memory retrieval (POS2/RSC/IPS) network TC/RC 13/37). A number of the resting state signal components have correspondence with task fMRI signal components that are not modulated by task, including the default mode network (Supplementary Figures TC/RC 26/15), the cingulo-opercular network (Supplementary Figures TC/RC 10/13), and the language network (Supplementary Figures TC/RC 22+36/20). Additional canonical resting state networks found only at rest included further subdivisions of the default mode network (Supplementary Figures RC7, 11, 24, 43), the fronto-parietal control network (Supplementary Figures RC17, 21), a component with the default mode network and fronto-parietal control network anti-correlated (Supplementary Figure RC23), the dorsal attention network (Supplementary Figure RC18), the parieto-occipital network (Laumann et al., 2015), Supplementary Figure RC32), the auditory network (Supplementary Figure RC36), and the language network (Supplementary Figure RC45). Other non-canonical resting state signal components included (Supplementary Figures RC4, 26, 30, 31, 35, 54, 74, 75, and 79).

In addition to the signal components described above, another set of signal components showed a variety of interesting properties potentially related to subjects’ arousal state. Most notably, the strongest component of the resting state data (Figure 8, Supplementary Figure RC1), which accounted for 8.5% of the tICA-explained variance, has visual, sensori-motor, and auditory regions with positive correlation and multi-modal cognitive regions generally with negative or near zero correlation. This component bears a strong resemblance to the beta map of the mean grey signal of sICA+FIX cleaned resting state fMRI data if the global positive background were removed (See Figure S6 of (Glasser et al., 2016b)). It is also notably variable across subjects (Figure 7). Being the strongest component and present only in the resting state data, this component presumably is linked to a behavior that is present during the resting state but not the task state. Drowsiness or sleep is a good candidate, as this component shows spatial map similarities to prior studies of the pattern of correlation during sleep vs wake (Tagliazucchi and Laufs, 2014). Also, increased amplitude of this semi-global neural component would contribute to the higher amplitude of global signal that is seen during sleep or decreased arousal that has been previously described (Laumann et al., 2017; Liu et al., 2017; Wong et al., 2016; Wong et al., 2013; Yeo et al., 2015).

**Figure 8.**
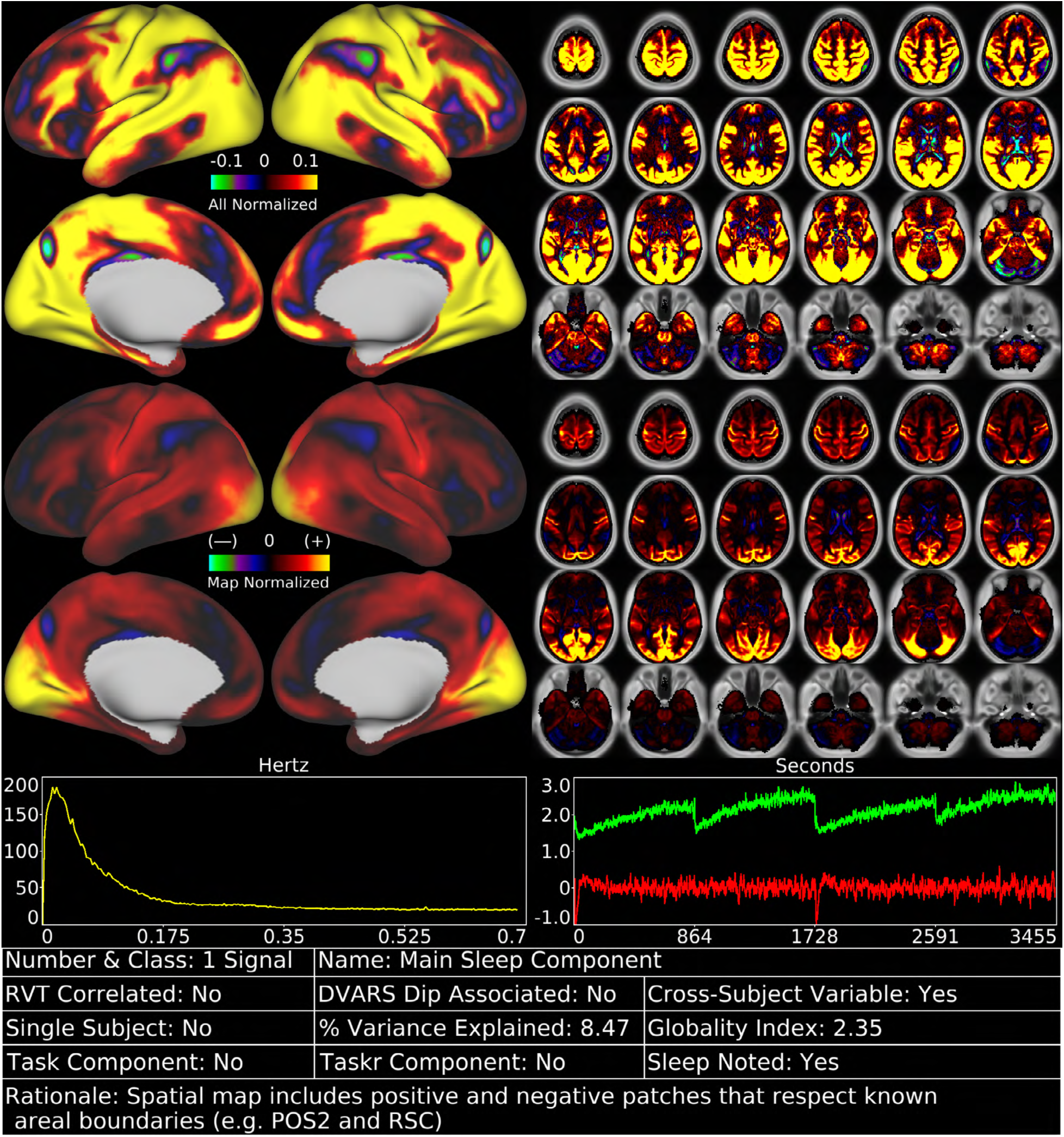
shows the first resting state fMRI tICA component in a standardized display format (see Supplementary Figures RC 1-84). The top row of data has the color scale normalized and held constant across all 84 components (in percent BOLD), whereas the second row has the color scale set independently for each component (scaled between 2nd and 98^th^ percentiles for that component). The first view allows components to be compared with each other on the same scale, whereas the second view highlights spatial variation in the map of each specific component. The left chart in the third row indicates the power spectra of the component averaged across subjects and the right chart indicates the average timeseries (red) and average absolute value of the timeseries (green). The increasing trend of the absolute average timeseries across the run (which is like an average instantaneous component amplitude) is likely indicative of subjects progressively becoming drowsy or falling asleep. The timeseries are concatenated in the order that the runs were typically acquired (REST1_RL, REST1_LR, REST2_LR, REST2_RL) to enable visualization of trends across the two sessions. Additional data on the components is provided in the table along the bottom row. The classification rationale appears along the bottom row of the table. Percent variance explained and the globality index are computed as explained in Figure 2.

Though sleep was not explicitly monitored in HCP subjects using eye tracking or EEG, technicians scanning the subjects were instructed to note if they noticed subjects were sleeping during resting state scans. When we compared the amplitude of the components for the 70 subjects who were noted to be sleeping vs the 379 who were not noted to be sleeping, component RC1 had the strongest association (Figure 9) and additionally had much stronger amplitude in the last 300 frames of the resting state run than the first 300 frames (Figure 9). Furthermore, in Figure 8, the green line showing the average absolute timeseries value is analogous to an average instantaneous amplitude that likely shows the effects of subjects on average becoming progressively drowsier or falling asleep over the course of each 14.4 minute-long resting state run. Five other components were associated both with sleep and with being stronger in the last 300 frames relative to the first 300 (Figure 9, Supplemental Figures RC5, 14, 22, 29, and 31) and had the same property of high cross-subject variability (Figure 7). Most of these components were not present in task fMRI data either, with the pan-sensori-motor component being the only exception (Supplemental Figures RC5). There were also 4 single subject components that appeared similar to RC1 or its spatial map negated (Supplemental Figures RC 55, 56, 65, and 69) and were thus classified as signal.

**Figure 9.**
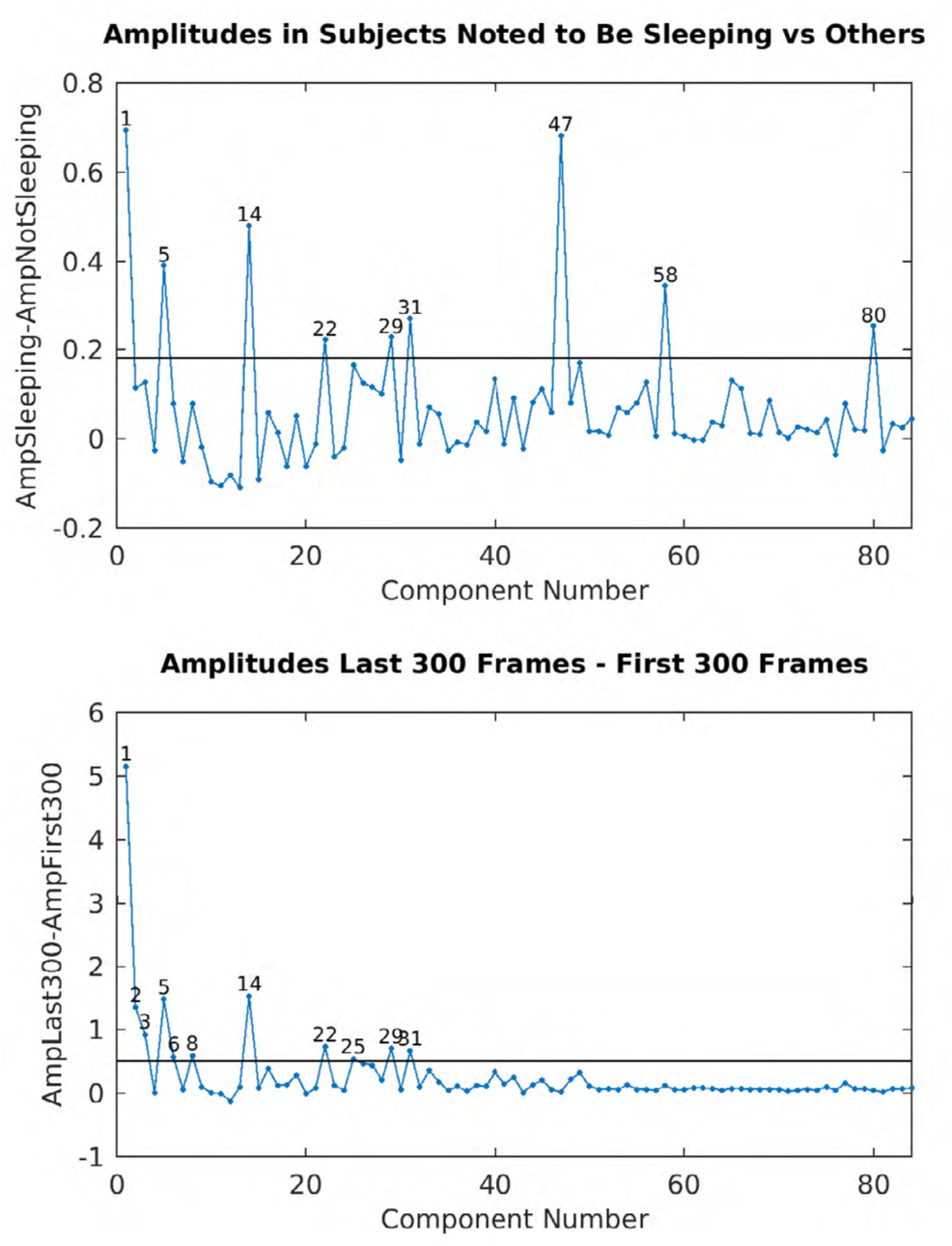
shows two useful sleep related component measures. The top row is the difference in component amplitude in subjects that were noted to be sleeping vs those who were not. The bottom row shows the difference in component amplitude between the last 300 frames and the first 300 frames of the 1200 frame runs, as subjects will presumably be more likely to be asleep at the end of a 14.4 minute run than at the beginning. Components 47, 58, and 80 are “single subject” components that are unlikely to be specifically sleep related. The 3 global physiological noise components (RC3, 6, and 8) are all also more likely to be higher at the end of a run than at the beginning, likely indicating greater respiratory variability at the end of runs when subjects are more likely to be drowsy. This is also true of noise component 25. Otherwise, these two measures agree on the components most likely to be related to sleep.

In aggregate sleep-related components accounted for 19.3% of the tICA explained variance. In resting state fMRI data, after removing the global noise components, the next two biggest contributors to the global signal are the semi-global sleep component and the pan-sensori-motor component (Supplementary Figure 26). Because of these components, resting state fMRI data overall has more global signal than does task fMRI data: MGTRVar=581 (mean variance removed by MGTR after sICA+FIX + tICA cleanup in grand mean 10000 scaled data) vs MGTRVar=235 for the task data. If sleep-related components are also removed, however, this difference disappears (MGTRVar=238 vs MGTRVar=235). Thus, there is no indication from the data that the global neural BOLD signal differs in magnitude between task fMRI and <u>awake</u> resting state fMRI (i.e., when subjects in the resting state paradigm are not sleeping) once the data have been appropriately cleaned using spatial and temporal ICA.

As with task fMRI, we clustered resting state fMRI components by run-wise amplitude cross-correlations (1796 runs = 4 runs * 449 subjects; Supplementary Figure 27). Again, we found that the clusters were dominated by a signal group and a noise group. Additionally, we found two groups of resting state signal components. One group tended to be correlated with RC1, and perhaps are networks that are generally more active in a drowsy or asleep state, whereas the other signal group was not as correlated with RC1 and may reflect networks that are more active in awake subjects. Additionally, we correlated the grayordinate spatial maps of the resting state fMRI, task fMRI, and task fMRI residual component maps (Supplementary Figure 28) and used this information when assigning component matches in the TC, TCr, and RC supplementary materials, the Supplementary Component Data Table and in the text above.

### 2.2.2 Effects of Temporal ICA Cleanup and MGTR on Resting State fMRI Greyplots

Figure 10 shows greyplots and mean grey signal of two runs of two subjects having particularly high global noise after sICA+FIX under three cleanup conditions: sICA+FIX alone, sICA+FIX + tICA, and sICA+FIX + MGTR. As with task fMRI, regressing out the components identified as noise removes the obvious bands in the greyplots (and substantial mean grey signal fluctuations at those times) that have previously been associated with respiratory changes (Power et al., 2018; Power et al., 2017b). Importantly, as with task fMRI, regressing out the noise temporal ICA components is not the same as removing the mean grey signal, as the mean grey signal after tICA cleanup still retains some fluctuations.

**Figure 10.**
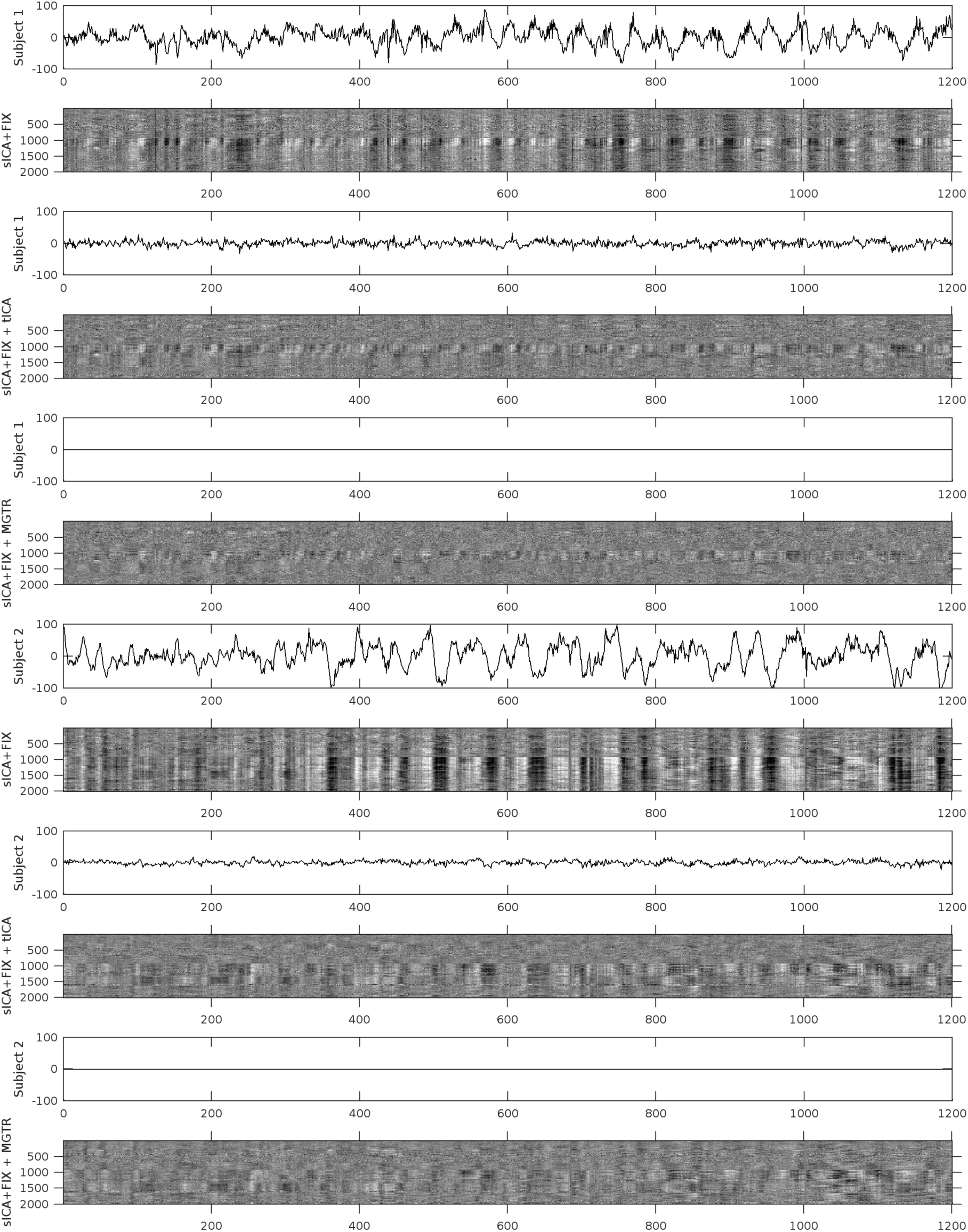
shows the mean grey signals and parcellated greyplots for two resting state fMRI runs from two subjects after sICA+FIX (Rows 1-2, 7-8), sICA+FIX + tICA (Rows 3-4, 9-10), and sICA+FIX + MGTR (Rows 5-6, 11-12). The data are parcellated, then displayed according to parcel surface area (with 1 row assigned for the smallest parcel and proportionally larger numbers of rows assigned for larger parcels such that there are more than 360 rows). Additionally, the data are ordered by hierarchical clustering of the group full correlation functional connectome after sICA+FIX + tICA so that parcels with more similar timeseries are closer together. tICA cleanup removes the vertical “stripes” from the grey plots after sICA+FIX and their corresponding MGT fluctuations without removing the entire MGT as occurs with MGTR. The greyplots after tICA cleanup and MGTR look very similar, although the tICA cleanup plots retain more semi-global structure.

In resting state fMRI data, the variance removed by regressing out the mean grey timecourse (MGTVar) before sICA+FIX cleanup is MGTRVar=9975; after sICA+FIX it is MGTRVar=1114; and after temporal ICA cleanup it is 581 (but see above point about increased sleep-related global variance). Thus, the neural global signal variance is 5.8% (581/9975) of the original global signal variance and is 52% (581/1114) of the global signal variance after sICA+FIX in resting state data. (For comparison, Liu et al found that the mean global timecourse represented ∼7% of the variance remaining after removing drift, motion, physiology, and WM/CSF regressors (Liu et al., 2017)). The neural global signal variance is 13.3% of the overall neural signal variance (see above for task fMRI values per task, and Supplemental Information Section #3), though this is increased somewhat because of sleeping subjects (7.4% after removal of sleep components). In contrast to task fMRI, there is no task design with which to assess the extent of neural signal removed using temporal ICA cleanup or MGTR. This lack of a standard makes assessing the effect of neural signal removal more challenging. It is the primary reason that we present the resting state data after the task fMRI data, even though the majority of the recent debate regarding fMRI denoising has been centered on resting-state fMRI because of the greater impact that residual global structured noise has on some resting state analyses.

### 2.2.3 Effects of Temporal ICA Cleanup and MGTR on Resting State fMRI Analyses

Figure 11 compares the three types of cleanup for resting state fMRI data. The top row shows group dense functional connectivity maps for a seed in area V1 after sICA+FIX (left column), after sICA+FIX + tICA cleanup (middle column), and after sICA+FIX + MGTR (right column). For this and other visual cortex seeds, the three methods differ substantially, with temporal ICA appearing to remove a global positive bias in functional connectivity and MGTR appearing to induce a negative bias in functional connectivity (anti-correlations that were not present in either the original or the tICA cleaned data). This mathematical effect of MGTR has long been known (Murphy et al., 2009; Saad et al., 2012) and is one of the arguments against use of MGTR, as it is neurobiologically implausible for the “true” functional connectome to have zero mean correlation. On the other hand, the large global positive bias in the correlations without temporal ICA cleanup shows why others have argued for the necessity of using something akin to MGTR (Fox et al., 2009; Hayasaka, 2013; Power et al., 2014). The second row shows dense functional connectivity maps for a seed in a cognitive/task-negative network with strong anti-correlation with other regions, including the ‘task-positive’ network (Figure 11). Here, the three methods yield more similar maps, with temporal ICA cleanup and MGTR being particularly similar. Thus, not all networks are equivalently biased by MGTR (though for this seed a global bias still appears to be removed by temporal ICA relative to no additional cleanup beyond sICA+FIX).

**Figure 11.**
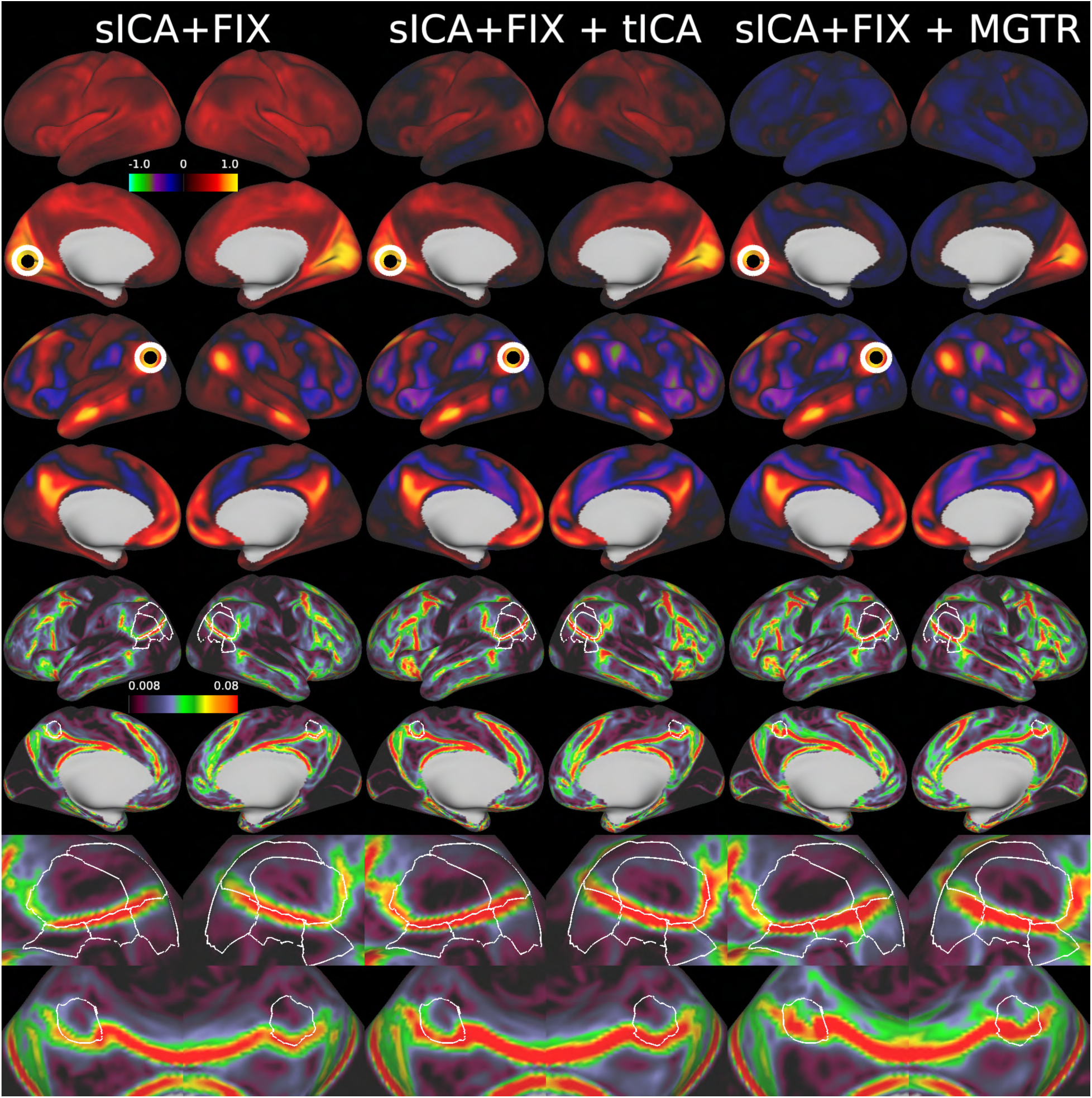
shows the results of group grayordinate-wise functional connectivity and functional connectivity gradient maps after sICA+FIX, sICA+FIX + tICA, and sICA+FIX + MGTR for the 210P subject group. The top row shows a seed in left hemisphere area V1 (circled) that has generally positive correlation with other grayordinates after sICA+FIX, positive to zero correlation after sICA+FIX + tICA, and widespread induced negative correlation after sICA+FIX + MGTR. The second row shows a seed in left hemisphere Area PGi (circled), which shows much more similar maps in an already anti-correlated network that is not as affected by MGTR. The third row shows the mean gradients after each kind of processing, with the gradients after sICA+FIX and sICA+FIX + tICA matching well (with an increase in gradient strength after tICA cleanup), whereas the gradients after MGTR shift in several regions highlighted by the outlined cortical areas (in white). The fourth row zooms in on several regions showing gradient shift after MGTR.

The bottom two rows of Figure 11 shows mean functional connectivity gradients computed for each method (see Methods Section #1.14), plus areal boundaries for six selected cortical areas (white contours, 5 lateral, 1 medial) in regions where methodological differences in gradient ridge location are particularly prominent. Both temporal ICA cleanup and MGTR increase the gradient magnitudes due to the removal of global noise. Unfortunately, MGTR also introduces a substantial shift in functional connectivity gradients relative to both no global noise cleanup and the temporal ICA-based cleanup solution. (This gradient shift is why MGTR was not used for the HCP’s multi-modal cortical parcellation (Glasser et al., 2016a), as a homogeneous positive bias would not affect areal boundary locations, only gradient strength, whereas a network-specific negative bias would lead to biased areal boundaries). This altering of the gradients is apparent quantitatively as well, as the spatial correlation between gradient maps is r=0.87 between sICA+FIX and sICA+FIX + tICA, vs. r=0.69 between sICA+FIX and sICA+FIX + MGTR, and r=0.77 between sICA+FIX + tICA and sICA+FIX + MGTR. Such shifting and altering of gradients is not consistent with a pure global artifact removal by MGTR (which would not affect some networks more than others) but is consistent with removal of network-specific neural signal as demonstrated above for the task fMRI data. Temporal ICA cleanup, on the other hand, removes the same global noise as MGTR without introducing a gradient shift because it does not remove semi-global neural signal.

When one parcellates the data and examines the functional connectivity between all pairs of cortical areas, it is apparent that the temporal ICA solution lies roughly halfway between no global noise cleanup and MGTR (Figures 12 and 13). On the one hand, if no additional cleanup is done after sICA+FIX, there is a positive bias in the data (Figure 12 left hand panel and Figure 13 Panel 1), and this bias may be correlated with other subject traits like motion, arousal, or body mass index (Siegel et al., 2017). On the other hand, MGTR removes both the global physiological noise and some of the neural signal in a network-specific way (Figure 13 compare Panel 2 vs 3 and 4 vs 5) as previously predicted (Glasser et al., 2016b; Saad et al., 2012). In particular, neural signal is mainly removed from both non-cognitive and task-positive regions, whereas there is less impact on other cognitive, task-negative regions (Figure 13 Panel 6). We believe that the temporal ICA cleanup approach offers a principled “best of both worlds” solution to the problem of separating global structured noise from global neural signal (see Discussion). The effect of tICA cleanup (Figure 13 Panel 4) is much more spatially homogeneous than the effect of MGTR (Panel 5), and spatial homogeneity is arguably the desired behavior when cleaning up the global artifacts that have prompted the use of MGTR. The homogeneous global effect of tICA cleanup vs. the network-specific effect of MGTR in resting state data mirrors what we saw in task fMRI data above (Supplementary Figure 25). Partial correlation is an alternative approach for establishing functional connectivity relationships between cortical areas that does not suffer from the positive bias that exists with full correlation after applying only sICA+FIX. We have previously recommended this approach (including in cases where the tICA cleanup approach has not been applied, (Glasser et al., 2016b), in part because partial correlation estimation is much less sensitive to global confounds than is full correlation. Indeed, consistent with this, the benefits from tICA cleanup for partial correlation are much more modest than for full correlation (Figure 13 Panels 7-9).

**Figure 12.**
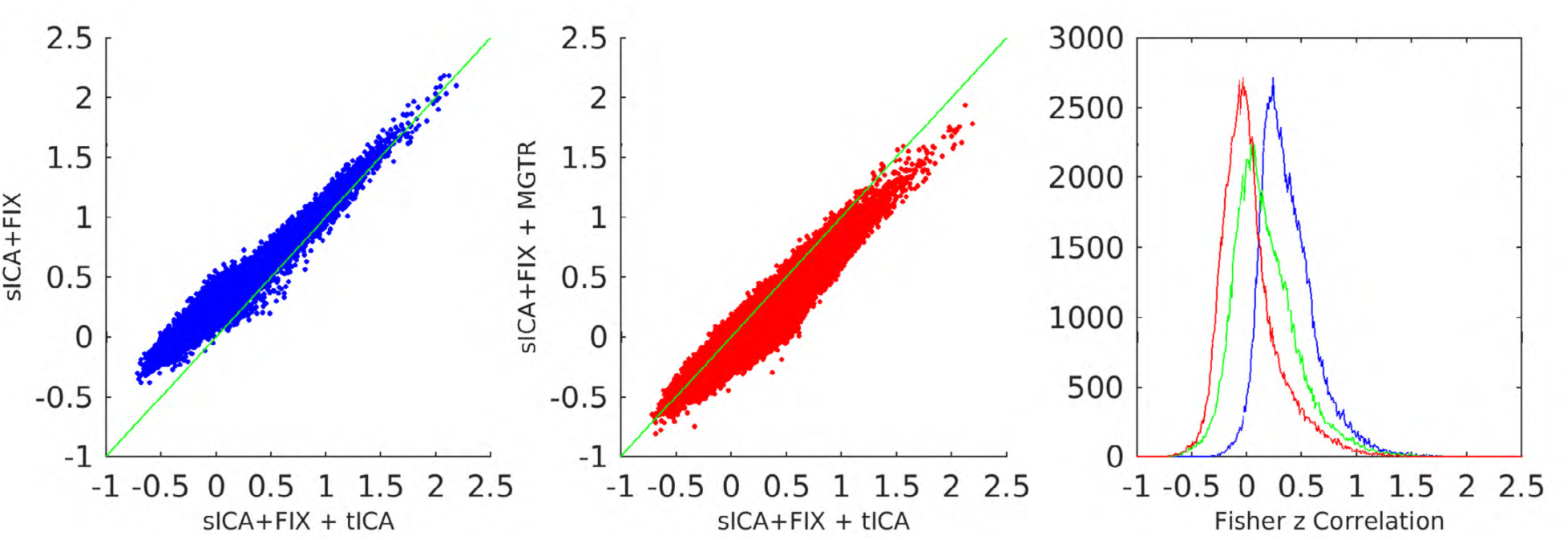
shows the entire full correlation parcellated connectome computed by parcellating the MIGP PCA series that was the input to the analyses in Figure 11 to show an overall summary of the trends in this data. It has sICA+FIX + tICA (x-axis) plotted vs sICA+FIX (y-axis; blue) on the left, showing a positive bias in sICA+FIX, and sICA+FIX + tICA plotted vs sICA+FIX + MGTR (red) in the middle, showing a negative bias in sICA+FIX + MGTR. Histograms of the correlation values are shown on the right with sICA+FIX blue, sICA+FIX + tICA green, and sICA+FIX + MGTR red, illustrating that sICA+FIX + tICA falls roughly halfway between sICA+FIX and sICA+FIX + MGTR.

**Figure 13.**
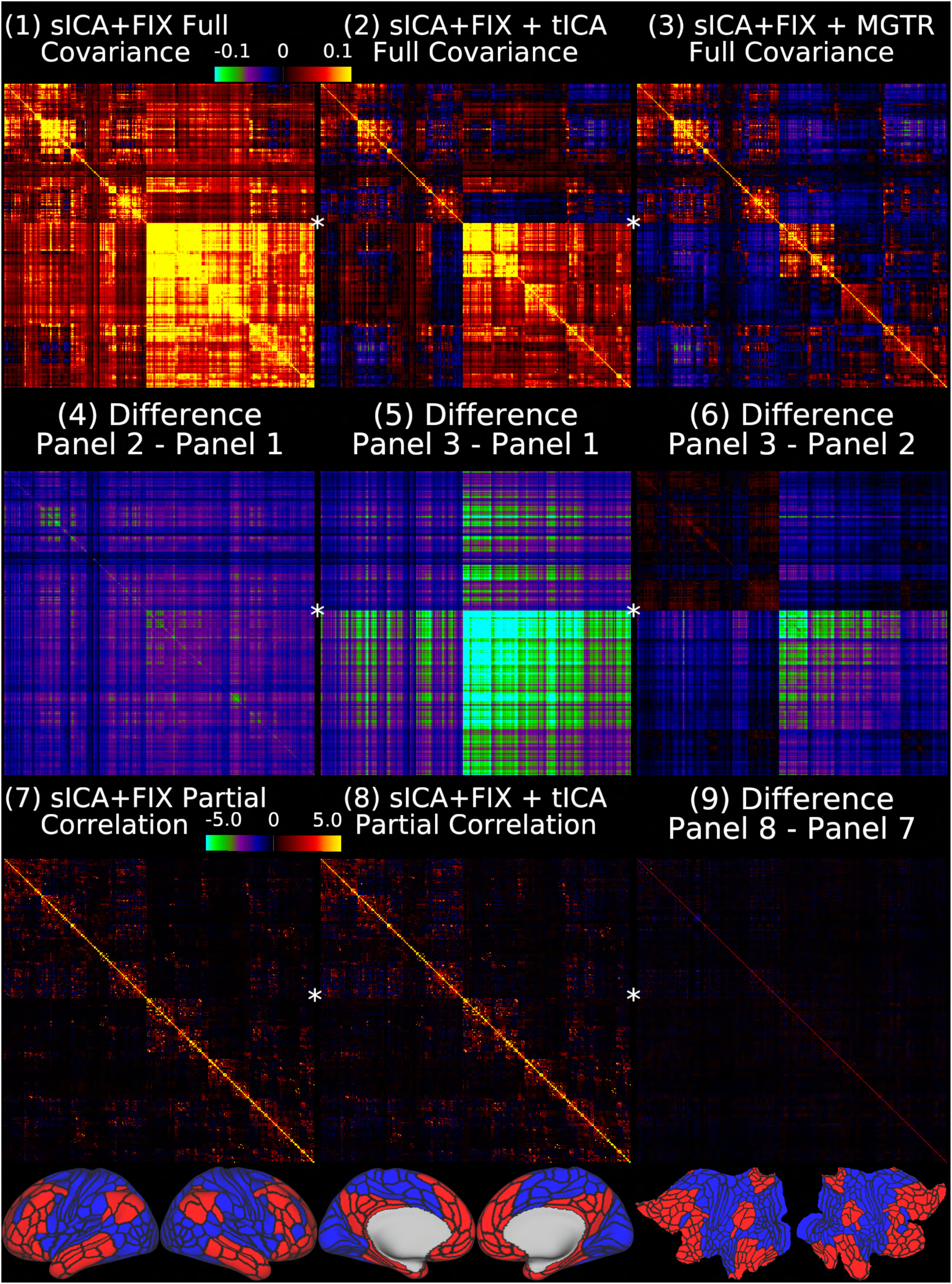
shows the group average full covariance matrices after sICA+FIX, sICA+FIX + tICA, and sICA+FIX + MGTR in Panels 1-3. We use covariance here because, like variances, covariances are additive and represent the absolute amount of variance shared by any two pairs of ROIs (scaled from −0.1 to 0.1 in percent BOLD for Panels 1-6). As in other figures, a global positive bias is removed by tICA cleanup (Panel 4 shows the difference between Panel 2 and Panel 1), but MGTR also removes additional signal in the bottom right quadrant of the matrix relative to the upper left quadrant with the off-diagonal quadrants in between (Panel 5 shows the difference between Panel 3 and Panel 1). Importantly, the difference between the tICA cleanup and MGTR (Panel 6 shows the difference between Panel 3 and Panel 2) is highly network specific, including small increases in cognitive/task-negative regions (bottom row, parcels shown in red) and large decreases in primarily non-cognitive/task positive regions (bottom row, blue parcels), with connections between parcels of these two broad groups of regions showing smaller decreases. Panels 7 and 8 show that mean across subjects partial correlation regularized with ridge regression (rho=0.23, which was optimal in matching the individual matrices to the group matrix computed with no regularization; scaled Z=+/-5) is much less affected by tICA cleanup, as it already controls for global artifacts (Glasser et al., 2016b). Thus Panel 9 (difference between Panel 8 and Panel 7) does not reveal substantial differences. The 360 cortical areas are ordered according to the same hierarchical clustering as the grey plots, and the first split, into cognitive/task negative (red) and non-cognitive and task positive (blue) regions, is shown in the bottom row and noted by a star on the netmats, with red parcels in the upper left quadrant of the netmats, and the blue parcels in the lower right quadrant. Note that it would be inappropriate to use partial correlation after MGTR, as any dataset that has zero global signal is rank deficient, because each parcel’s timeseries equals the negated sum of all other parcels’ timeseries.

## Discussion

We have shown that it is possible to separate global noise from global neural signal using temporal ICA and then to selectively remove the noise, something that was urgently needed but previously lacking in the literature (Glasser et al., 2016b; Power et al., 2017a; Power et al., 2018; Power et al., 2017b). Based on its spatial pattern (specific to grey matter but global) and temporal characteristics (BOLD power spectrum, correlated with RVT), much of this global noise is likely due to global fluctuations in blood flow from changes in respiratory rate and depth, leading to changes in pCO_2_, or from other physiological sources like heart rate variations, as previously predicted (Glasser et al., 2016b; Power et al., 2017a; Power et al., 2018). This global noise is left behind by spatial-ICA-based cleanup approaches like sICA+FIX (Burgess et al., 2016; Glasser et al., 2016b; Power et al., 2017b; Siegel et al., 2017). Selectively removing this noise without removing the global neural signal addresses a major issue in functional MRI denoising by allowing for an attractive intermediate solution between not performing global signal regression (and thus tolerating a homogeneous positive bias from global noise in univariate connectivity measures that may differ across subjects and groups), or performing global signal regression (and thus inducing network-specific negative biases in connectivity measures, as was previously predicted in the literature (Glasser et al., 2016b; Gotts et al., 2013; Saad et al., 2012; Yang et al., 2016; Yang et al., 2014)). Thus, with temporal ICA cleanup we enable the generation of univariate connectivity measures that are both free of global physiological noise and not biased by global signal regression. Additionally, by separating the global signal and global noise fluctuations we are able to measure the size of each independently, finding in both task and resting state fMRI data that after sICA+FIX cleanup about half the global signal variance is noise and half is neural. Thus, the non-GSR and GSR solutions both are roughly equidistant from the tICA solution, though they have very different kinds of biases relative to it (global positive vs network-specific negative) that have differing implications for downstream analyses as discussed below. That said, multivariate approaches such as partial correlation, which we have previously recommended as a way around the global noise dilemma (Glasser et al., 2016b), are substantially less affected by global artifacts or tICA cleanup.

We first considered task fMRI data, where we have a prediction of what a portion of the subjects’ BOLD signal should look like due to the task and thus can objectively determine if a cleanup method is removing neural signal. In the task fMRI data, we identified a single global component that is correlated with the respiratory physiology parameter RVT and represents a major source of global noise (e.g., this component has a correlation of r=0.68 with the mean grey timecourse after sICA+FIX). We then selectively removed this component, together with other noise components, and compared the results of task fMRI statistics before and after this processing, and to those after global signal regression. Because it had not previously been done, we also compared the effects of sICA+FIX cleanup vs no cleanup on task fMRI data. We found that sICA+FIX clearly increases statistical sensitivity and removes biases. Temporal ICA cleanup increases statistical sensitivity for differential contrasts between different task “on” blocks and removes biases for primary task “on” vs baseline contrasts that arise from stimulus-correlated physiology. On the other hand, global signal regression clearly reduces statistical sensitivity across both primary and differential contrasts relative to temporal ICA cleanup, removes a network-specific spatial pattern that is highly correlated with the task activation map, and biases the contrast beta maps towards a mean of zero by removing neural signal. Importantly, our tICA cleanup approach also removed the “stripes” associated with global artifacts in greyplots (Figures 5 and 10), something that physiological regressors have only incompletely achieved to date and that has otherwise required global signal regression to achieve (Power et al., 2017b).

We then applied the same temporal ICA approach to resting state fMRI data, finding three global components, the sum of whose timecourses correlated strongly with the mean grey signal after sICA+FIX (r=0.71) and with RVT. Using tICA cleanup, we demonstrated the removal of a global positive bias in univariate connectivity measures without inducing a network-specific negative bias in these measures as found with global signal regression. Additionally, we showed that whereas functional connectivity gradients shift after global signal regression (consistent with removing neural signal in a network-specific way), they do not shift with temporal ICA-based global noise cleanup, consistent with removing only a global artifact. Indeed, the effect of tICA cleanup (Figure 13 Panel 4) is arguably more like the spatially homogeneous effect that practitioners of global signal regression aspire to apply to their data, whereas the effect of global signal regression (Figure 13 Panel 5) is actually quite spatially inhomogeneous and network specific.

In addition to these successful denoising results, the temporal ICA approach – run on millions of task fMRI and resting state fMRI timepoints – generates several interesting neurobiological findings. We identified a number of neural signal components that are modulated by tasks in task fMRI data, many of which have spatial maps that are very similar to the task fMRI contrast beta maps. Additionally, we identified components present in both task and resting state fMRI data that are not modulated by task. We found a set of five somatotopically organized sensori-motor networks that are modulated by the MOTOR task, but are also correlated with DVARS Dips. DVARS Dips are a more robust indicator of subject head movement in the high spatial and temporal resolution fMRI data that we used in this study than are common measures based on movement parameter traces such as FD (Main Supplementary Information Section #4). The correlation between the somatotopically organized sensorimotor networks and DVARS Dips is notable because it explains how some data cleanup benchmarks that do not take into account the expected neural changes of different subject behaviors (such as movement or sleep) may suggest an apparent benefit for approaches that remove neural signal (e.g., scrubbing or global signal regression, see Main Supplementary Information Section #6 for further discussion of the limitations of such metrics and examples of some of these metrics applied to this study to help provide a historical perspective on the denoising literature). There may also be subject-wise correlations between noncompliant behavior in the scanner and other behavioral variables collected outside the scanner (e.g., putative head motion and BMI), or subject behavior in the scanner (e.g., movement and sleep), or other subject-varying properties that affect the BOLD signal such as subject breathing patterns or heart rate variability. We also identify a set of components present only in the resting state that are associated with subjects noted to be sleeping and, in particular, a semi-global neural component that likely underlies the higher global signal that has been found in sleep or drowsy states (Laumann et al., 2017; Liu et al., 2017; Tagliazucchi and Laufs, 2014; Wong et al., 2016; Wong et al., 2013; Yeo et al., 2015). Future work may explore whether these arousal-associated components can be experimentally manipulated independently from the global structured noise components that are correlated with RVT.

In general, a number of components are shared between task and resting state fMRI data. But there are clear differences as well—above and beyond the task-modulated components. That said, we found no evidence that task and resting state fMRI have differential amounts of global neural signal once sleep is taken into account. Therefore, using first spatial and then temporal ICA allows for selective structured noise removal from both kinds of fMRI data. We believe that the converging evidence from task and resting state fMRI presented in this study support temporal ICA cleanup as an attractive “best of both worlds” approach that avoids both the pitfalls of failing to clean global structured noise from the data, and of removing global neural signal from the data with overly aggressive methods.

The method described in this study is designed to work with high spatial and temporal resolution fMRI data at the group level (including a large number of subjects) after precise functional alignment across subjects. Many studies (often by necessity) acquire relatively small amounts of resting state fMRI data in small numbers of subjects at low spatial and temporal resolutions. It is already known that subject-wise methods like sICA+FIX do not perform as well on such datasets because of the small number of timepoints (Griffanti et al., 2014; Salimi-Khorshidi et al., 2014). Temporal ICA will be even more affected by these issues (Smith et al., 2012), so the methods presented in this study are intended for use in HCP-Style neuroimaging acquisitions (i.e., better than 2.6mm isotropic spatial resolution, better than 1s temporal resolution, and large total numbers of timepoints (#timepoints per subject * #subjects), e.g., something greater than the 20,000 timepoints used in (Smith et al., 2012)). However, the minimum number of timepoints for robust performance is currently unknown, as is the rate at which performance improves with increasing numbers of timepoints. Thus, these methods are particularly applicable to large datasets such as the existing Young Adult HCP data, the ongoing HCP Lifespan (Development and Aging) projects, Connectomes Related to Human Disease (“Disease Connectome”) data, HCP-Style studies of individual investigators, the ABCD (Adolescent Brain and Cognitive Development) study, and the UK Biobank (Miller et al., 2016). In general, it will be particularly important in the future to assess tICA denoising performance as a function of the number of subjects included in a study and how many timepoints are available per subject. A corollary approach that also warrants careful examination is to perform tICA on task and resting-state data combined, thereby potentially increasing the number of data points available for analysis in many projects. Additionally, it may be possible to use the temporal ICA decomposition from a larger group of subjects to correct other individual subjects who were not a part of the original group; however, investigating and validating this approach is left for future work.

The approach presented here has several limitations. First, it currently relies on manual classification of temporal ICA components at the group level before cleanup can proceed. However, this is likely not a long-term impediment, given the success of machine learning classifiers at classifying spatial ICA components. Also, not all investigators may agree with the specific component classifications used here, particularly for the small number of controversial components (e.g., TC42+TC59/TCr36/RC44 in particular engendered debate among the coauthors and are indeed in need of further study), and further understanding and experience with temporal ICA may lead to improved classifications in the future. Indeed, the tICA approach makes the global signal/global noise problem one of component classification, rather than whether or not to do global signal regression. However, generalizing the approach outside of HCP-Style datasets may be challenging because of the data quality and quantity requirements of temporal ICA, which are even more demanding than those for spatial ICA (which itself also works less well on more traditional data (Griffanti et al., 2014; Salimi-Khorshidi et al., 2014)). That said, we have shown that the method works in two large datasets, a task fMRI dataset and a resting state fMRI dataset from the Young Adult HCP.

The importance of the temporal ICA cleanup step might be underestimated because of the relatively small amount of noise remaining after sICA+FIX (27% or 28% of the structured variance remaining after sICA+FIX is noise vs. 89% or 93% of the structured variance before sICA+FIX), and relatively small amount of neural global signal relative to the overall neural signal variance (10% or 13%). Thus, the question might reasonably be asked: Given the complexity of the tICA cleanup method and its demanding data requirements, why not just analyze the data after sICA+FIX and ignore the positive bias from the spatially homogeneous global noise? Or conversely, why not just use GSR and not worry about the neural signal lost? After sICA+FIX^4^ about half of the global signal variance is artifact and half is neural (49% or 52%). As a result, the size of the positive bias if tICA cleanup is not done is about the same size as the negative bias if GSR is performed. Thus, it would be logically inconsistent to argue that it is critical to remove the positive bias by using GSR while dismissing the negative bias of similar magnitude induced by using GSR.

Moreover, the global artifact removed by tICA is evenly distributed across the grey matter, scaled mainly by the T2* decay rate (being lower in dropout regions). Such a uniform global artifact will lead to a harmless global increase in functional connectivity for many analyses, e.g., boundary based brain areal parcellation (Glasser et al., 2016a) or clustering-based functional network parcellation, and the effects on graph theoretic measures could be dealt with by increasing thresholds (or if necessary by using connectivity matrix demeaning). GSR, on the other hand, distorts maps of connectivity gradients and induces a network-specific negative bias in both task fMRI and resting state fMRI activity, as predicted in the literature (Glasser et al., 2016b; Gotts et al., 2013; Saad et al., 2012; Yang et al., 2016; Yang et al., 2014). Such a bias will change functional connectivity measures non-uniformly across the edges, affecting some networks more than others, which would distort quantification or parcellation of functional networks or graph theoretic relationships. Further, GSR could cause an underestimation of brain dynamics, as functional networks that increase their amplitude transiently during increased utilization would contribute more to the global neural signal and thus these dynamic changes would be blunted by GSR (e.g., the sensori-motor networks during the motor task). At the same time, for a study of connectivity dynamics it would be critical to clean the data with tICA so that physiological noise does not give rise to spurious dynamics. Another challenging scenario involves a study of patients and controls when the groups differ in amount of global physiological noise. Rather than using GSR in this situation and altering the connectivity patterns in the two groups, a safer solution (if partial correlation is not an option) would be to use the amplitude of the global signal, the mean connectivity value across the connectome (Saad et al., 2013), or another similar measure (Hahamy et al., 2014) as a subject-wise covariate of no interest in the group statistical analysis. Thus, although group differences in the overall amplitude of neural global signal or mean correlation value would no longer be available, the connectivity relationships between brain areas in functional networks would remain undistorted. Indeed, we struggled to conceive of a situation in which the network-specific biases induced by GSR would be preferable to an alternative analysis strategy using some sort of post-hoc standardization (Yan et al., 2013b) that takes advantage of the global (spatially homogeneous) nature of the positive connectivity bias induced by physiological noise, even in the absence of tICA cleanup being widely available at the present time.

Finally, we reiterate that our approach is intended only to clean data of physiological, movement, and MR physics related artifacts, not to alter the BOLD neural signal present in subjects’ scans. Subjects who do not follow instructions and fail to perform the assigned task, who move, or fall asleep, will have differing neural BOLD activity and connectivity patterns than those who are compliant. We also cannot assume that these differences will be limited to the global neural signal. As a result, we do not recommend regressing out the somatotopic sensorimotor networks or the sleep components as a standard approach, because these represent real neural signals. That said, investigators might rightly wish to exclude portions of scans or subjects who engage in off-task behaviors. The occurrence of off-task behaviors is generally more prevalent in resting state fMRI data than task fMRI data, as we have shown here for sleep. Such behaviors may also be more prevalent in some groups of subjects than others, which is an important confound for resting state fMRI studies, particularly those comparing two or more groups of subjects. Other paradigms, such as naturalistic movies may do a better job of maintaining subject “neural compliance” with the study design. Nevertheless, it will still be important to selectively remove spatially specific and global structured noise from such fMRI data using spatial and temporal ICA.

## Acknowledgements

Supported in part by the Human Connectome Project, WU-Minn-Ox Consortium (1U54MH091657) funded by the 16 NIH Institutes and Centers that support the NIH Blueprint for Neuroscience Research; the McDonnell Center for Systems Neuroscience at Washington University; and NIH F30 MH097312 (M.F.G.), RO1 MH-60974 (D.C.V.E.).

Funding to SS, JB, SH gratefully acknowledged via Wellcome Trust strategic award 098369/Z/12/Z. The authors would also like to thank Andreas Bartsch for helpful references related to negative CSF correlations in global and semi-global components.

## Footnotes

1 Throughout the manuscript, the “HCP-MMP1.0 multi-modal cortical parcellation” refers specifically to the one derived using the maximum probability map (MPM) in the 210 validation subjects (210V group): https://balsa.wustl.edu/file/show/3VLx

2 Weighted regression was run using all of the data of a given kind (task or resting state), however cleanup was done at the level of individual 1200 timepoint resting state runs or 1884 or 1996 timepoint concatenated task fMRI sessions to match how the data were cleaned with sICA+FIX.

3 Note that the inclusion of all tICA timecourses (i.e., both “signal” and “noise”) in the multiple regression for computing the betas makes this a “non-aggressive” cleanup similar to what we use for sICA+FIX cleanup, leaving any variance associated with the tICA signal components intact. In practice, however, the issue of components having non-orthogonal variance will be a much smaller effect for tICA than sICA, because tICA component timecourses are defined to be temporally orthogonal at the group level and this property should largely persist at the individual run or concatenated task session level.

4 sICA+FIX removes the spatially specific non-BOLD noise (i.e. S0 intensity mediated effects; Power et al 2018) related to motion, coil artifacts, etc and thus the sICA+FIX cleaned data is the most appropriate stage for assessing the relative proportion of global neural signal vs global physiological noise (both processes mediated by T2* signal decay; Power et al 2018). Additionally, sICA+FIX has been demonstrated to be robust for routine use on HCP-Style data (Glasser et al., 2016b; Salimi-Khorshidi et al., 2014), which is the same kind of data we recommend for tICA cleanup.

## References

1. Aguirre, G.K., E. Zarahn, and M. D’Esposito. 1997. Empirical analyses of BOLD fMRI statistics. II. Spatially smoothed data collected under null-hypothesis and experimental conditions. NeuroImage. 5:199–212.

2. Aguirre, G.K., E. Zarahn, and M. D’Esposito. 1998. The inferential impact of global signal covariates in functional neuroimaging analyses. NeuroImage. 8:302–306.

3. Anderson, J.S., T.J. Druzgal, M. Lopez-Larson, E.K. Jeong, K. Desai, and D. Yurgelun-Todd. 2011. Network anticorrelations, global regression, and phase-shifted soft tissue correction. Human brain mapping. 32:919–934.

4. Barch, D.M., G.C. Burgess, M.P. Harms, S.E. Petersen, B.L. Schlaggar, M. Corbetta, M.F. Glasser, S. Curtiss, S. Dixit, C. Feldt, D. Nolan, E. Bryant, T. Hartley, O. Footer, J.M. Bjork, R. Poldrack, S. Smith, H. Johansen-Berg, A.Z. Snyder, and D.C. Van Essen. 2013. Function in the human connectome: task-fMRI and individual differences in behavior. NeuroImage. 80:169–189.

5. Behzadi, Y., K. Restom, J. Liau, and T.T. Liu. 2007. A component based noise correction method (CompCor) for BOLD and perfusion based fMRI. NeuroImage. 37:90–101.

6. Birn, R.M., J.B. Diamond, M.A. Smith, and P.A. Bandettini. 2006. Separating respiratory-variation-related fluctuations from neuronal-activity-related fluctuations in fMRI. NeuroImage. 31:1536–1548.

7. Brewer, A.A., W.A. Press, N.K. Logothetis, and B.A. Wandell. 2002. Visual areas in macaque cortex measured using functional magnetic resonance imaging. The Journal of neuroscience : the official journal of the Society for Neuroscience. 22:10416–10426.

8. Brooks, J.C., C.F. Beckmann, K.L. Miller, R.G. Wise, C.A. Porro, I. Tracey, and M. Jenkinson. 2008. Physiological noise modelling for spinal functional magnetic resonance imaging studies. NeuroImage. 39:680–692.

9. Burgess, G.C., S. Kandala, D. Nolan, T.O. Laumann, J.D. Power, B. Adeyemo, M.P. Harms, S.E. Petersen, and D.M. Barch. 2016. Evaluation of Denoising Strategies to Address Motion-Correlated Artifacts in Resting-State Functional Magnetic Resonance Imaging Data from the Human Connectome Project. Brain connectivity. 6:669–680.

10. Carbonell, F., P. Bellec, and A. Shmuel. 2011. Global and system-specific resting-state fMRI fluctuations are uncorrelated: principal component analysis reveals anti-correlated networks. Brain connectivity. 1:496–510.

11. Chai, X.J., A.N. Castanon, D. Ongur, and S. Whitfield-Gabrieli. 2012. Anticorrelations in resting state networks without global signal regression. NeuroImage. 59:1420–1428.

12. Chang, C., J.P. Cunningham, and G.H. Glover. 2009. Influence of heart rate on the BOLD signal: the cardiac response function. NeuroImage. 44:857–869.

13. Chang, C., and G.H. Glover. 2009. Relationship between respiration, end-tidal CO2, and BOLD signals in resting-state fMRI. NeuroImage. 47:1381–1393.

14. Chang, C., D.A. Leopold, M.L. Scholvinck, H. Mandelkow, D. Picchioni, X. Liu, F.Q. Ye, J.N. Turchi, and J.H. Duyn. 2016. Tracking brain arousal fluctuations with fMRI. Proceedings of the National Academy of Sciences of the United States of America. 113:4518–4523.

15. Ciric, R., D.H. Wolf, J.D. Power, D.R. Roalf, G.L. Baum, K. Ruparel, R.T. Shinohara, M.A. Elliott, S.B. Eickhoff, C. Davatzikos, R.C. Gur, R.E. Gur, D.S. Bassett, and T.D. Satterthwaite. 2017. Benchmarking of participant-level confound regression strategies for the control of motion artifact in studies of functional connectivity. NeuroImage. 154:174–187.

16. Farah, M. 2014. Brain images, babies, and bathwater: Critiquing critiques of functional neuroimaging. The Hastings Center Report. 44.

17. Filippini, N., B.J. MacIntosh, M.G. Hough, G.M. Goodwin, G.B. Frisoni, S.M. Smith, P.M. Matthews, C.F. Beckmann, and C.E. Mackay. 2009. Distinct patterns of brain activity in young carriers of the APOE-epsilon4 allele. Proceedings of the National Academy of Sciences of the United States of America. 106:7209–7214.

18. Fox, M.D., D. Zhang, A.Z. Snyder, and M.E. Raichle. 2009. The global signal and observed anticorrelated resting state brain networks. Journal of neurophysiology. 101:3270–3283.

19. Fukunaga, M., S.G. Horovitz, P. van Gelderen, J.A. de Zwart, J.M. Jansma, V.N. Ikonomidou, R. Chu, R.H. Deckers, D.A. Leopold, and J.H. Duyn. 2006. Large-amplitude, spatially correlated fluctuations in BOLD fMRI signals during extended rest and early sleep stages. Magnetic resonance imaging. 24:979–992.

20. Gattass, R., A.P. Sousa, and C.G. Gross. 1988. Visuotopic organization and extent of V3 and V4 of the macaque. The Journal of neuroscience : the official journal of the Society for Neuroscience. 8:1831–1845.

21. Glasser, M.F., T.S. Coalson, E.C. Robinson, C.D. Hacker, J. Harwell, E. Yacoub, K. Ugurbil, J. Andersson, C.F. Beckmann, M. Jenkinson, S.M. Smith, and D.C. Van Essen. 2016a. A multi-modal parcellation of human cerebral cortex. Nature. 536:171–178.

22. Glasser, M.F., S.M. Smith, D.S. Marcus, J.L. Andersson, E.J. Auerbach, T.E. Behrens, T.S. Coalson, M.P. Harms, M. Jenkinson, S. Moeller, E.C. Robinson, S.N. Sotiropoulos, J. Xu, E. Yacoub, K. Ugurbil, and D.C. Van Essen. 2016b. The Human Connectome Project’s neuroimaging approach. Nature neuroscience. 19:1175–1187.

23. Glasser, M.F., S.N. Sotiropoulos, J.A. Wilson, T.S. Coalson, B. Fischl, J.L. Andersson, J. Xu, S. Jbabdi, M. Webster, J.R. Polimeni, D.C. Van Essen, and M. Jenkinson. 2013. The minimal preprocessing pipelines for the Human Connectome Project. NeuroImage. 80:105–124.

24. Golestani, A.M., C. Chang, J.B. Kwinta, Y.B. Khatamian, and J. Jean Chen. 2015. Mapping the end-tidal CO2 response function in the resting-state BOLD fMRI signal: spatial specificity, test-retest reliability and effect of fMRI sampling rate. NeuroImage. 104:266–277.

25. Gotts, S.J., Z.S. Saad, H.J. Jo, G.L. Wallace, R.W. Cox, and A. Martin. 2013. The perils of global signal regression for group comparisons: a case study of Autism Spectrum Disorders. Frontiers in human neuroscience. 7:356.

26. Griffanti, L., G. Douaud, J. Bijsterbosch, S. Evangelisti, F. Alfaro-Almagro, M.F. Glasser, E.P. Duff, S. Fitzgibbon, R. Westphal, D. Carone, C.F. Beckmann, and S.M. Smith. 2017. Hand classification of fMRI ICA noise components. NeuroImage. 154:188–205.

27. Griffanti, L., G. Salimi-Khorshidi, C.F. Beckmann, E.J. Auerbach, G. Douaud, C.E. Sexton, E. Zsoldos, K.P. Ebmeier, N. Filippini, C.E. Mackay, S. Moeller, J. Xu, E. Yacoub, G. Baselli, K. Ugurbil, K.L. Miller, and S.M. Smith. 2014. ICA-based artefact removal and accelerated fMRI acquisition for improved resting state network imaging. NeuroImage. 95:232–247.

28. Hahamy, A., V. Calhoun, G. Pearlson, M. Harel, N. Stern, F. Attar, R. Malach, and R. Salomon. 2014. Save the global: global signal connectivity as a tool for studying clinical populations with functional magnetic resonance imaging. Brain connectivity. 4:395–403.

29. Hayasaka, S. 2013. Functional connectivity networks with and without global signal correction. Frontiers in human neuroscience. 7:880.

30. He, H., and T.T. Liu. 2012. A geometric view of global signal confounds in resting-state functional MRI. NeuroImage. 59:2339–2348.

31. Himberg, J., A. Hyvarinen, and F. Esposito. 2004. Validating the independent components of neuroimaging time series via clustering and visualization. NeuroImage. 22:1214–1222.

32. Horovitz, S.G., M. Fukunaga, J.A. de Zwart, P. van Gelderen, S.C. Fulton, T.J. Balkin, and J.H. Duyn. 2008. Low frequency BOLD fluctuations during resting wakefulness and light sleep: a simultaneous EEG-fMRI study. Human brain mapping. 29:671–682.

33. Hyvarinen, A. 1999. Fast and robust fixed-point algorithms for independent component analysis. IEEE transactions on Neural Networks,. 10:626–634.

34. Igasaki, T., K. Nagasawa, I.A. Akbar, and N. Kubon. 2016. Sleepiness classification by thoracic respiration using support vector machine. Biomedical Engineering Interational Conference (BMEICON). 9th:pp. 1-5, IEEE

35. Kundu, P., S.J. Inati, J.W. Evans, W.M. Luh, and P.A. Bandettini. 2012. Differentiating BOLD and non-BOLD signals in fMRI time series using multi-echo EPI. NeuroImage. 60:1759–1770.

36. Laumann, T.O., E.M. Gordon, B. Adeyemo, A.Z. Snyder, S.J. Joo, M.Y. Chen, A.W. Gilmore, K.B. McDermott, S.M. Nelson, N.U. Dosenbach, B.L. Schlaggar, J.A. Mumford, R.A. Poldrack, and S.E. Petersen. 2015. Functional System and Areal Organization of a Highly Sampled Individual Human Brain. Neuron. 87:657–670.

37. Laumann, T.O., A.Z. Snyder, A. Mitra, E.M. Gordon, C. Gratton, B. Adeyemo, A.W. Gilmore, S.M. Nelson, J.J. Berg, D.J. Greene, J.E. McCarthy, E. Tagliazucchi, H. Laufs, B.L. Schlaggar, N.U.F. Dosenbach, and S.E. Petersen. 2017. On the Stability of BOLD fMRI Correlations. Cereb Cortex. 27:4719–4732.

38. Liu, T.T. 2016. Noise contributions to the fMRI signal: An overview. NeuroImage. 143:141–151.

39. Liu, T.T., A. Nalci, and M. Falahpour. 2017. The global signal in fMRI: Nuisance or Information? NeuroImage. 150:213–229.

40. Macey, P.M., K.E. Macey, R. Kumar, and R.M. Harper. 2004. A method for removal of global effects from fMRI time series. NeuroImage. 22:360–366.

41. Marx, M., K.B. Pauly, and C. Chang. 2013. A novel approach for global noise reduction in resting-state fMRI: APPLECOR. NeuroImage. 64:19–31.

42. Miller, K.L., F. Alfaro-Almagro, N.K. Bangerter, D.L. Thomas, E. Yacoub, J. Xu, A.J. Bartsch, S. Jbabdi, S.N. Sotiropoulos, J.L. Andersson, L. Griffanti, G. Douaud, T.W. Okell, P. Weale, I. Dragonu, S. Garratt, S. Hudson, R. Collins, M. Jenkinson, P.M. Matthews, and S.M. Smith. 2016. Multimodal population brain imaging in the UK Biobank prospective epidemiological study. Nature neuroscience. 19:1523–1536.

43. Murphy, K., R.M. Birn, D.A. Handwerker, T.B. Jones, and P.A. Bandettini. 2009. The impact of global signal regression on resting state correlations: are anti-correlated networks introduced? NeuroImage. 44:893–905.

44. Murphy, K., and M.D. Fox. 2017. Towards a consensus regarding global signal regression for resting state functional connectivity MRI. NeuroImage. 154:169–173.

45. Muschelli, J., M.B. Nebel, B.S. Caffo, A.D. Barber, J.J. Pekar, and S.H. Mostofsky. 2014. Reduction of motion-related artifacts in resting state fMRI using aCompCor. NeuroImage. 96:22–35.

46. Power, J.D. 2017. A simple but useful way to assess fMRI scan qualities. NeuroImage. 154:150–158.

47. Power, J.D., K.A. Barnes, A.Z. Snyder, B.L. Schlaggar, and S.E. Petersen. 2012. Spurious but systematic correlations in functional connectivity MRI networks arise from subject motion. NeuroImage. 59:2142–2154.

48. Power, J.D., T.O. Laumann, M. Plitt, A. Martin, and S.E. Petersen. 2017a. On Global fMRI Signals and Simulations. Trends in cognitive sciences. 21:911–913.

49. Power, J.D., A. Mitra, T.O. Laumann, A.Z. Snyder, B.L. Schlaggar, and S.E. Petersen. 2014. Methods to detect, characterize, and remove motion artifact in resting state fMRI. NeuroImage. 84:320–341.

50. Power, J.D., M. Plitt, S.J. Gotts, P. Kundu, V. Voon, P.A. Bandettini, and A. Martin. 2018. Ridding fMRI data of motion-related influences: Removal of signals with distinct spatial and physical bases in multiecho data. Proceedings of the National Academy of Sciences of the United States of America. 115:E2105–E2114.

51. Power, J.D., M. Plitt, T.O. Laumann, and A. Martin. 2017b. Sources and implications of whole-brain fMRI signals in humans. NeuroImage. 146:609–625.

52. Power, J.D., B.L. Schlaggar, and S.E. Petersen. 2015. Recent progress and outstanding issues in motion correction in resting state fMRI. NeuroImage. 105:536–551.

53. Pruim, R.H., M. Mennes, J.K. Buitelaar, and C.F. Beckmann. 2015a. Evaluation of ICA-AROMA and alternative strategies for motion artifact removal in resting state fMRI. NeuroImage. 112:278–287.

54. Pruim, R.H., M. Mennes, D. van Rooij, A. Llera, J.K. Buitelaar, and C.F. Beckmann. 2015b. ICA-AROMA: A robust ICA-based strategy for removing motion artifacts from fMRI data. NeuroImage. 112:267–277.

55. Robinson, E.C., K. Garcia, M.F. Glasser, Z. Chen, T.S. Coalson, A. Makropoulos, J. Bozek, R. Wright, A. Schuh, M. Webster, J. Hutter, A. Price, L.C. Grande, E. Hughes, N. Tusor, P.V. Bayly, D.C. Van Essen, S.M. Smith, A.D. Edwards, J. Hajnal, M. Jenkinson, B. Glocker, and D. Rueckert. 2017. Multimodal surface matching with higher-order smoothness constraints. NeuroImage.

56. Robinson, E.C., S. Jbabdi, M.F. Glasser, J. Andersson, G.C. Burgess, M.P. Harms, S.M. Smith, D.C. Van Essen, and M. Jenkinson. 2014. MSM: a new flexible framework for Multimodal Surface Matching. NeuroImage. 100:414–426.

57. Saad, Z.S., S.J. Gotts, K. Murphy, G. Chen, H.J. Jo, A. Martin, and R.W. Cox. 2012. Trouble at rest: how correlation patterns and group differences become distorted after global signal regression. Brain connectivity. 2:25–32.

58. Saad, Z.S., R.C. Reynolds, H.J. Jo, S.J. Gotts, G. Chen, A. Martin, and R.W. Cox. 2013. Correcting brain-wide correlation differences in resting-state FMRI. Brain connectivity. 3:339–352.

59. Salimi-Khorshidi, G., G. Douaud, C.F. Beckmann, M.F. Glasser, L. Griffanti, and S.M. Smith. 2014. Automatic denoising of functional MRI data: combining independent component analysis and hierarchical fusion of classifiers. NeuroImage. 90:449–468.

60. Satterthwaite, T.D., K. Ruparel, J. Loughead, M.A. Elliott, R.T. Gerraty, M.E. Calkins, H. Hakonarson, R.C. Gur, R.E. Gur, and D.H. Wolf. 2012. Being right is its own reward: load and performance related ventral striatum activation to correct responses during a working memory task in youth. NeuroImage. 61:723–729.

61. Scholvinck, M.L., A. Maier, F.Q. Ye, J.H. Duyn, and D.A. Leopold. 2010. Neural basis of global resting-state fMRI activity. Proceedings of the National Academy of Sciences of the United States of America. 107:10238–10243.

62. Sereno, M.I., A.M. Dale, J.B. Reppas, K.K. Kwong, J.W. Belliveau, T.J. Brady, B.R. Rosen, and R.B. Tootell. 1995. Borders of multiple visual areas in humans revealed by functional magnetic resonance imaging. Science. 268:889–893.

63. Shmueli, K., P. van Gelderen, J.A. de Zwart, S.G. Horovitz, M. Fukunaga, J.M. Jansma, and J.H. Duyn. 2007. Low-frequency fluctuations in the cardiac rate as a source of variance in the resting-state fMRI BOLD signal. NeuroImage. 38:306–320.

64. Siegel, J.S., A. Mitra, T.O. Laumann, B.A. Seitzman, M. Raichle, M. Corbetta, and A.Z. Snyder. 2017. Data Quality Influences Observed Links Between Functional Connectivity and Behavior. Cereb Cortex. 27:4492–4502.

65. Smith, S.M., C.F. Beckmann, J. Andersson, E.J. Auerbach, J. Bijsterbosch, G. Douaud, E. Duff, D.A. Feinberg, L. Griffanti, M.P. Harms, M. Kelly, T. Laumann, K.L. Miller, S. Moeller, S. Petersen, J. Power, G. Salimi-Khorshidi, A.Z. Snyder, A.T. Vu, M.W. Woolrich, J. Xu, E. Yacoub, K. Ugurbil, D.C. Van Essen, and M.F. Glasser. 2013a. Resting-state fMRI in the Human Connectome Project. NeuroImage. 80:144–168.

66. Smith, S.M., A. Hyvarinen, G. Varoquaux, K.L. Miller, and C.F. Beckmann. 2014. Group-PCA for very large fMRI datasets. NeuroImage. 101:738–749.

67. Smith, S.M., K.L. Miller, S. Moeller, J. Xu, E.J. Auerbach, M.W. Woolrich, C.F. Beckmann, M. Jenkinson, J. Andersson, M.F. Glasser, D.C. Van Essen, D.A. Feinberg, E.S. Yacoub, and K. Ugurbil. 2012. Temporally-independent functional modes of spontaneous brain activity. Proceedings of the National Academy of Sciences of the United States of America. 109:3131–3136.

68. Smith, S.M., D. Vidaurre, C.F. Beckmann, M.F. Glasser, M. Jenkinson, K.L. Miller, T.E. Nichols, E.C. Robinson, G. Salimi-Khorshidi, M.W. Woolrich, D.M. Barch, K. Ugurbil, and D.C. Van Essen. 2013b. Functional connectomics from resting-state fMRI. Trends in cognitive sciences. 17:666–682.

69. Spronk, M., J.L. Ji, K. Kulkarni, G. Repovs, A. Anticevic, and M.W. Cole. 2017. Mapping the human brain’s cortical-subcortical functional network organization. bioRxiv:p. 206292.

70. Tagliazucchi, E., and H. Laufs. 2014. Decoding wakefulness levels from typical fMRI resting-state data reveals reliable drifts between wakefulness and sleep. Neuron. 82:695–708.

71. Uddin, L.Q. 2017. Mixed Signals: On Separating Brain Signal from Noise. Trends in cognitive sciences. 21:405–406.

72. Van Essen, D.C., W.T. Newsome, and J.H.R. Maunsell. 1984. The visual field representation in striate cortex of the macaque monkey: asymmetries, anisotropies and individual variability. Vision Res. 24:429–448.

73. Van Essen, D.C., J. Smith, M.F. Glasser, J. Elam, C.J. Donahue, D.L. Dierker, E.K. Reid, T. Coalson, and J. Harwell. 2017. The Brain Analysis Library of Spatial maps and Atlases (BALSA) database. NeuroImage. 144:270–274.

74. Van Essen, D.C., S.M. Smith, D.M. Barch, T.E. Behrens, E. Yacoub, and K. Ugurbil. 2013. The WU-Minn Human Connectome Project: an overview. NeuroImage. 80:62–79.

75. Wen, H., and Z. Liu. 2016. Broadband Electrophysiological Dynamics Contribute to Global Resting-State fMRI Signal. The Journal of neuroscience : the official journal of the Society for Neuroscience. 36:6030–6040.

76. Wong, C.W., P.N. DeYoung, and T.T. Liu. 2016. Differences in the resting-state fMRI global signal amplitude between the eyes open and eyes closed states are related to changes in EEG vigilance. NeuroImage. 124:24–31.

77. Wong, C.W., V. Olafsson, O. Tal, and T.T. Liu. 2013. The amplitude of the resting-state fMRI global signal is related to EEG vigilance measures. NeuroImage. 83:983–990.

78. Woolrich, M.W., B.D. Ripley, M. Brady, and S.M. Smith. 2001. Temporal autocorrelation in univariate linear modeling of FMRI data. NeuroImage. 14:1370–1386.

79. Yan, C.G., B. Cheung, C. Kelly, S. Colcombe, R.C. Craddock, A. Di Martino, Q. Li, X.N. Zuo, F.X. Castellanos, and M.P. Milham. 2013a. A comprehensive assessment of regional variation in the impact of head micromovements on functional connectomics. NeuroImage. 76:183–201.

80. Yan, C.G., R.C. Craddock, X.N. Zuo, Y.F. Zang, and M.P. Milham. 2013b. Standardizing the intrinsic brain: towards robust measurement of inter-individual variation in 1000 functional connectomes. NeuroImage. 80:246–262.

81. Yang, G.J., J.D. Murray, M. Glasser, G.D. Pearlson, J.H. Krystal, C. Schleifer, G. Repovs, and A. Anticevic. 2016. Altered Global Signal Topography in Schizophrenia. Cereb Cortex.

82. Yang, G.J., J.D. Murray, G. Repovs, M.W. Cole, A. Savic, M.F. Glasser, C. Pittenger, J.H. Krystal, X.J. Wang, G.D. Pearlson, D.C. Glahn, and A. Anticevic. 2014. Altered global brain signal in schizophrenia. Proceedings of the National Academy of Sciences of the United States of America. 111:7438–7443.

83. Yeo, B.T., F.M. Krienen, J. Sepulcre, M.R. Sabuncu, D. Lashkari, M. Hollinshead, J.L. Roffman, J.W. Smoller, L. Zollei, J.R. Polimeni, B. Fischl, H. Liu, and R.L. Buckner. 2011. The organization of the human cerebral cortex estimated by intrinsic functional connectivity. Journal of neurophysiology. 106:1125–1165.

84. Yeo, B.T., J. Tandi, and M.W. Chee. 2015. Functional connectivity during rested wakefulness predicts vulnerability to sleep deprivation. NeuroImage. 111:147–158.

85. Zarahn, E., G.K. Aguirre, and M. D’Esposito. 1997. Empirical analyses of BOLD fMRI statistics. I. Spatially unsmoothed data collected under null-hypothesis conditions. NeuroImage. 5:179–197.

