## Supplementary Materials for "Using Temporal ICA to Selectively Remove Global Noise While Preserving Global Signal in Functional MRI Data"

[1]Department of Neuroscience, Washington University Medical School, Saint Louis, Missouri 63110, USA. [2]St. Luke's Hospital, Saint Louis, Missouri 63017, USA. [3]Centre for Functional MRI of the Brain (FMRIB), Wellcome Centre for Integrative Neuroimaging, Nuffield Department of Clinical Neurosciences, University of Oxford. John Radcliffe Hospital, Headley Way, Oxford, OX3 9DU, UK. [4]Department of Psychiatry, Washington University Medical School, Saint Louis, MO. [5]Department of Psychiatry, Yale University School of Medicine, 300 George Street, New Haven, CT 06511, USA

### Introduction to Supplementary Material

The supplementary material is divided into five different documents: the Main Supplementary Information (this document), the Supplementary Task Components (TC), the Supplementary Task Components from the residuals after fitting the task GLM (TCr), the Supplementary Resting State Components (RC), and the Supplementary Component Data Table (a spreadsheet named tICA_Component_Data_Table.xlsx) containing information that is available as text associated with each component in the TC, TCr, and RC, documents, but which is provided additionally as a separate spreadsheet for convenient reference and sorting.

This document contains (i) a Supplementary Introduction on Independent Components Analysis (ICA; Topic #1); (ii) Supplementary Methods on modifications of the sICA+FIX pipeline to clean multiple fMRI runs at the same time (Topic #2), a model for variance partitioning of fMRI data (Topic #3), and a discussion of motion metrics (Topic #4); (iii) Supplementary Results and Figures, which includes a discussion of negative CSF in both non-neural and neural semi-global and global components (Topic #5) and numerous supplementary figures mentioned in the Main Text; and (iv) a Supplementary Discussion on data cleanup validation metrics (Topic #6) that includes at historical perspective on the denoising literature and some metrics historically used in the literature, but that we believe are challenging to interpret because they lack positive controls.

The Supplementary TC document contains 70 pages, one for each tICA component, presented in a standardized way similar to Main Text Figures 2 and 8. The Supplementary TCr document has 58 pages, and the Supplementary RC document has 84 pages presented in the same standardized format. For each component a description of the classification rationale is provided and all this information is also available in the Supplementary Component Data Table.

**Supplementary Introduction**

*1. Understanding ICA: The Difference Between Spatial and Temporal ICA*

To explain why temporal ICA can help with separating and removing global noise, whereas spatial ICA cannot, we briefly summarize key aspects of how ICA works. We first note that unlike ICA, Principal Components Analysis (PCA) decomposes the data so that each component is both spatially and temporally orthogonal to all others (and gives the same decomposition regardless of the orientation of the space-by-time matrix when it is presented to the PCA algorithm—see the relationship between PCA and Singular Value Decomposition, SVD). PCA is also generally a deterministic algorithm, and one can compute as many non-zero principal components as the rank of the dataset. ICA relaxes the orthogonality constraint along one of the axes, depending on how the space-by-time matrix is oriented when it is presented to the ICA algorithm. ICA algorithms are generally non-deterministic machine learning algorithms that attempt to find a rotation (the mixing matrix) that will separate the data into components that are statistically independent (and hence also orthogonal) along either the space or the time axis. PCA is typically applied before ICA to segregate the data into structured and unstructured subspaces (see Section #3). ICA is then run on the structured subspace so that algorithm convergence is more likely.

ICA works best if the axis for which component independence is optimized has a) many samples (voxels or timepoints), and b) strongly structured (i.e., non-Gaussian) components. For both of these reasons, spatial ICA has typically been used to separate fMRI data into independent components (Beckmann et al., 2005; Beckmann and Smith, 2004). In spatial ICA, the algorithm generates a set of spatial component maps that are independent (and hence also orthogonal, i.e., their pairwise spatial correlation matrix is the identity matrix). The mixing matrix of spatial ICA is composed of a timeseries for each component, and, importantly, these timeseries are not orthogonal to each other, though they are non-collinear. In fact, their correlation will increase with increasing amounts of global fluctations in the original fMRI data.

The constraint (in most ICA algorithms) that components must be spatially orthogonal and sparse effectively prohibits spatial ICA from identifying global or semi-global timeseries fluctuations as one or more separable components. This is because global components are difficult to make spatially uncorrelated with other components or with one another. Spatial ICA does do an excellent job of separating structured noise that has a specific temporal pattern localized to a specific location or locations (e.g., from movement, from vascular pulsation, or from localized MRI equipment-related artifacts). Global structured noise cannot be easily separated from other signals by spatial ICA, and the non-aggressive cleanup approach typically used by ICA-based denoising algorithms ensures that shared variance between signal and noise components also will not be removed (Griffanti et al., 2014; Salimi-Khorshidi et al., 2014). As a result, while spatial ICA has been very successful at removing spatially specific artifacts, it does not succeed at removing global artifacts.

As fMRI acquisition methods have become more advanced, allowing greater temporal resolution and larger numbers of timepoints per subject, temporal ICA has become a feasible analysis strategy (Smith et al., 2012). Temporal ICA is performed by transposing the fMRI data matrix relative to its orientation in Spatial ICA before presenting it to the ICA algorithm. This matrix flip results in temporally independent timeseries components (meaning that their temporal correlation matrix, i.e., their "Functional Connectome," is the identity matrix). Spatial maps constitute the mixing matrix, and, importantly, these spatial maps are not orthogonal to each other, though they are non-collinear. Unlike with spatial ICA, one or more global or semi-global spatial maps can be found if they are present in the data. (However, note that temporal ICA would have difficulty separating signals that are either constant or nearly constant across time, even if they were spatially localized). Additionally, when cleanup is done using temporal ICA components, there will be minimal shared temporal variance between the components (in our implementation there is zero shared variance across the whole concatenated group dataset, but this is not strictly true for each individual resting state run or concatenated task session). Thus, temporal ICA is inherently better suited for cleaning fMRI data of global artifacts than is spatial ICA.

Additionally, whereas spatial ICA behaves like a weighted parcellation of the brain into functional regions or network nodes when performed at high dimensionality, temporal ICA behaves more like a weighted parcellation of the brain into uncorrelated functional networks (Smith et al., 2012). These networks will tend to be more spatially widespread than the component maps of high dimensional spatial ICA and will contain all of a functional network's nodes in a single component spatial map. Such nodes may of course be shared by more than one network, but the temporal ICA component spatial maps of signal components represent temporally independent functional networks. All in all, temporal ICA is an as yet underutilized tool for cleaning and investigating fMRI data, and thus this study explores its properties in detail.

**Supplementary Methods**

*2. sICA+FIX and a Modified Pipeline for Task fMRI*

The original sICA+FIX pipeline begins by removing linear trends and means from the fMRI timeseries (mean across time, not space as in GSR) and from the 24 movement regressors (Friston et al., 1996). Then Melodic's implementation of FastICA is run (Beckmann et al., 2005; Beckmann and Smith, 2004), and the FIX algorithm automatically classifies the resulting sICA components as "signal" or "noise" using training weights previously established with the FIX machine learning classifier (Griffanti et al., 2014; Salimi-Khorshidi et al., 2014). Then the 24 movement regressors are "aggressively" regressed out of the data and sICA component timeseries (i.e., removing all variance spanning the space of the movement regressors[1]) (Smith et al., 2013). Finally the sICA components classified as noise are "non-aggressively" regressed out of the data (i.e., removing the unique variance associated with the noise sICA components while leaving in any shared variance that is also associated with the signal components (Smith et al., 2013). Importantly, while spatial ICA can reasonably converge using short (e.g., 5 min) fMRI runs (due to the large number of voxels in the spatial dimension), sICA nonetheless still benefits from having large numbers of time points per subject to enable finer and more accurate splitting of components, leading to better separation of signal and noise and easier component classification. Unpublished HCP pilot analyses of running sICA+FIX on single task fMRI runs showed that, while it reduced some false positives, it did not produce a marked statistical sensitivity benefit (improvement in Z scores). Importantly, the shorter the task fMRI run, the more there appeared to be a statistical sensitivity penalty (i.e., lower Z scores). As a result, sICA+FIX had not been previously performed for the released HCP task fMRI data. In the current study, the objective was to explore task fMRI data in parallel with resting-state fMRI data using temporal ICA. Therefore, despite the above noted caveats, it was essential to process the task fMRI runs similarly to the resting-state fMRI data. Thus, as a part of this study we modified the sICA+FIX pipeline to combine across task fMRI runs to enable robust sICA+FIX cleaning despite the individual runs having substantially shorter length than the resting state fMRI runs.

As before, linear trends and means were removed from each run's timeseries and 24 movement regressors. Then the data and movement regressors were concatenated across runs within a single task fMRI session (of which there were two in the HCP: Working Memory + Gambling + Motor with 1884 timepoints and Language + Social + Relational + Emotion with 1996 timepoints). Melodic sICA was run on the concatenated timeseries, and the FIX classification and cleanup pipelines were then run on the concatenated data. Finally, the timeseries were split back apart, and the corresponding single run mean images were added back. Using the same classifier weights as were used for the resting state fMRI data, we found that the performance of the FIX classifier on the concatenated task fMRI data was similar (99.5% classification accuracy vs manual classification, assessed on the 2 concatenated runs in each of 28 subjects) to what we previously had found with single resting state fMRI runs of 1200 timepoints (greater than 99% classification accuracy (Griffanti et al., 2014; Salimi-Khorshidi et al., 2014)). Given that both resting state and task fMRI perform well with the same classifier and training data, it would be reasonable to combine across resting state and task fMRI data for future studies that acquire less fMRI data, such as the HCP Lifespan Development and Aging projects.

---

[1]Note that this step was implemented before temporal ICA cleanup was developed and may not be necessary or desirable in the context of combined sICA and tICA cleanup, as aggressively regressing out the movement regressors from both the data and the sICA component timeseries could remove some neural signal, and the main advantage of including this aggressive movement parameter regression step in addition to the non-aggressive sICA noise component timeseries regression is to clean the global timecourse of artifactual movement related fluctuations. We will leave exploration the optimal use of external movement regressors for future work.

*3. A Model of fMRI Data Subspaces and Variance Partitions*

The signal of interest in an fMRI BOLD timeseries is comprised of its temporal fluctuations and the relationship of these fluctuations to either a task stimulus or between different brain regions. Thus, we can think of the fMRI timeseries in terms of different kinds of additive fluctuations (i.e., data subspaces or variances) some of which are desirable (neurally-related BOLD signals) and some of which are undesirable (i.e., fluctuations induced by subject motion, subject physiology, the MRI equipment and environment, or random thermal noise). When we clean an fMRI timeseries, the goal is to remove artifactual fluctuations selectively without modifying the BOLD fluctuations from neural signals. Thus, we can characterize all forms of data cleanup in terms of the variance that they remove, and we do this as implemented in a development version of the 'Resting State Stats' Pipeline in the HCP Pipelines. The temporal variance averaged across the grayordinates and then the 449 subjects of the 'spatially minimally-preprocessed' data prior to any temporal preprocessing is 115984 in resting state fMRI data and 92909 in task fMRI data (the data are grand mean scaled to 10,000 in the volume).

We can partition the variance of an fMRI timeseries into additive variance bins in a variety of ways, but perhaps the most fundamental partition is between structured variance and unstructured variance (Equation #1). By structured variance, we mean variance of the subspace of the data that is structured in space and/or time (i.e., non-random variance). We can define structured variance as the PCA components that have eigenvalues above those expected in the eigenvalue distribution of a random noise matrix of the same size as the data (i.e. a Wishart distribution) (Glasser et al., 2016a; Glasser et al., 2016b). Using this measure as estimated by FSL's Melodic (see below), the structured variance averages 47.1% of the total variance in resting state fMRI and 38.5% of the total variance in task fMRI data, with the remaining 52.9% and 61.5% of the variance being unstructured. These values will be sensitive to a variety of protocol factors, including field strength, head coil, voxel size, and repetition time, with high-resolution HCP data having a higher proportion of unstructured noise relative to traditional acquisition protocols with larger voxels and slower temporal sampling. Operationally, the variance attributable to a specific data subspace can be computed by taking the variance of the difference between the timeseries before and after regressing that subspace out of the data (e.g., the sICA component timeseries that make up the structured signal in Equation 1). In this way, it is possible to determine the proportion of the total variance (or the proportion of a subdivision of the total variance) that any given data subspace comprises.

$$Var_{orig} = Var_{struct} + Var_{unstruct} \quad (1)$$

A spatial independent components analysis (sICA, as implemented in FSL's Melodic software (Beckmann et al., 2005; Beckmann and Smith, 2004), achieves this separation by first performing a PCA, ordering the resulting principal components by eigenvalue, and finding the threshold at which the eigenvalues are higher than what would be expected in a null eigenvalue (i.e. Wishart) distribution of completely unstructured data with similar spatial and temporal autocorrelation. The subsequent ICA is then performed on the $Var_{struct}$ subspace of the fMRI timeseries data, and the $Var_{unstruct}$ subspace is ignored. The $Var_{unstruct}$ subspace is largely Gaussian or other random noise, and there are a variety of ways to reduce the impact of the $Var_{unstruct}$ subspace on other analyses of grayordinate-wise resting state and task fMRI data. For example, spatial or temporal smoothing, a Wishart Rolloff, or parcellation will all reduce unstructured noise (Glasser et al., 2016a; Glasser et al., 2016b). Here, we will focus on further partitioning the $Var_{struct}$ subspace and leave a comparison of the methods for reducing or removing the $Var_{unstruct}$ subspace for future work.

The $Var_{struct}$ subspace contains both the structured BOLD signal, which we term $Var_{BOLD}$, and the structured noise $Var_{structnoise}$. Many denoising approaches aim to separate the signal and noise fluctuations as cleanly as possible (Equation #2). Indeed, the structured variance averages 93.0% noise and 7.0% signal in HCP's resting state fMRI data and 89.0% noise and 11.1% signal in HCP's task fMRI data. We note that variance has a nonlinear relationship to amplitude (a signal with twice the amplitude has four times the variance), and we use variance here because of the convenient property that the total variance is the sum of the signal variance when adding orthogonal signals (we note below when this orthogonality assumption does not completely hold).

$$Var_{struct} = Var_{BOLD} + Var_{structnoise} \quad (2)$$

The sICA+FIX approach (Griffanti et al., 2014; Salimi-Khorshidi et al., 2014) classifies sICA components into $Var_{BOLDICA}$ and $Var_{noiseICA}$ (as do similar approaches like ICA-AROMA, (Pruim et al., 2015a; Pruim et al., 2015b) and Multi-Echo ICA (Kundu et al., 2012)). Prior to removing the noise components, sICA+FIX runs two additional cleanup stages: A high-pass filter that is for all practical purposes a linear detrending operation and a 24-movement parameter regression. Thus, the full variance partition model of sICA+FIX cleanup is shown in Equation #3. Of the original variance in HCP rest/task fMRI data, the detrend averages 18.9%/9.2%, the 24 movement parameters are 16.4%/5.7%, the sICA noise components are 13.1%/20.6%, the sICA signal components are 3.7%/2.8%, and the unstructured noise 52.9%/61.5%.

$$Var_{orig} = Var_{detrend} + Var_{mvm24} + Var_{noiseICA} + Var_{BOLDICA} + Var_{unstruct} \quad (3)$$

The cleaned fMRI timeseries produced by sICA+FIX is shown by Equation #4. In the cleaned timeseries the BOLD variance is 7.0%/4.3% of the remaining variance. Because sICA does not create temporally orthogonal components, there is the possibility for some shared variance between $Var_{noiseICA}$ and $Var_{BOLDICA}$. But, the sum of variances on the right side of Equation #3 is only 1.0%/0.4% larger than the variance of the original data, indicating that the magnitude of the shared variance is small (i.e., this particular variance partitioning is a reasonable model).

$$Var_{clean} = Var_{BOLDICA} + Var_{unstruct} \quad (4)$$

Although it would be possible during sICA+FIX cleanup to simply reconstruct the data from the estimated $Var_{BOLDICA}$ subspace and thereby eliminate the unstructured variance ($Var_{unstruct}$) entirely, this is not done because some weakly structured signals may not be detected at the individual subject level, and thus will exist in the unstructured noise subspace. These weakly structured signals may still be detectable at the group level, making it important to postpone discarding this subspace until as late in the analysis as possible (see footnote #6 in (Smith et al., 2013)). Similarly, the point at which the data eigenspectrum intersects with the null eigenspectrum is often much higher than the dimensionality estimated by melodic using the Laplacian approximation (Beckmann et al., 2005; Beckmann and Smith, 2004).

As mentioned in the Main Text and above, spatial ICA is mathematically blind to variance that is spatially semi-global or global in nature, and indeed one does not find semi-global or global components with the spatial ICA algorithm used here. That means that the spatially global structured variance must be shared across all of the spatial ICA components to some extent (importantly, all of the timeseries variance must be represented in at least one of the variance partitions, as the sum of the partitions approximately equals the original timeseries variance, subject to the shared variance caveat above). Global structured variance that is present in $Var_{detrend}$ and $Var_{mvm24}$ is of no concern, as that variance is removed entirely by the "aggressive" cleanup (see above) of those regressors during sICA+FIX. Similarly, global structured variance that is uniquely present in $Var_{noiseICA}$ is of no concern, as it is removed by the "non-aggressive" cleanup of the identified noise sICA components. Thus, if global structured artifactual variance remains in the data, it must largely be represented in the $Var_{BOLDICA}$ portion of the data. Thus, we can rewrite Equation #4 as Equation #5 where $Var_{BOLDICA}$ contains both global, $Var_{BOLDGlobalNeuralSignal}$, and spatially specific, $Var_{BOLDSpatiallySpecificNeuralSignal}$, neurally related BOLD signal along with the residual global noise $Var_{BOLDGlobalNoise}$.

$$Var_{clean} = Var_{BOLDGlobalNoise} + Var_{BOLDGlobalNeuralSignal} + Var_{BOLDSpatiallySpecificNeuralSignal} + Var_{unstruct} \quad (5)$$

The goal of our temporal ICA method is to separate and remove $Var_{BOLDGlobalNoise}$ selectively, leaving us with Equation #6.

$$\mathrm{Var}_{tclean} = \mathrm{Var}_{BOLDGlobalNeuralSignal} + \mathrm{Var}_{BOLDSpatiallySpecificNeuralSignal} + \mathrm{Var}_{unstruct} \quad (6)$$

Of the variance remaining after sICA+FIX cleanup, tICA cleanup removes 6.6%/2.9% (rest/task) of the variance, which is $\mathrm{Var}_{BOLDGlobalNoise}$ (which here technically also includes any residual spatially specific noise left behind by sICA+FIX). tICA cleanup leaves 0.9%/0.4% of the variance remaining after sICA+FIX that is attributable to global BOLD neural signal, 5.1%/3.3% to spatially specific BOLD neural signal, and the remaining 87.3%/93.5% of the variance to unstructured noise.

In addition to partitioning the timeseries variance into signal and noise, one can similarly regress out the task design to determine what proportion of the total BOLD neural signal or the global BOLD neural signal is task related to the task design. However, this will underestimate the task driven signal insofar as the task design does not fully model the task driven signal (see Main Text Figure 4 bottom panel), so those results in the Main Text should be considered a lower bound on the amount of task related signal.

*4. Is All that Moves Head Motion?*

It might seem relatively straightforward to model motion in the fMRI timeseries: Simply realign the data (as a standard preprocessing step) and then use the realignment parameters to calculate a measure of motion. Indeed, several QC measures of movement computed from realignment parameters have been proposed (e.g., FD (Power et al., 2012), absolute or relative RMS (Jenkinson, 1999)) and have proven very useful with traditional fMRI data (with larger voxels and slower TR). However, displacements in putative head position can arise from more than just true physical head motion, and the realignment algorithm will still correct for this 'apparent' motion. In particular, respiration (i.e., air entering the lungs) causes changes in the magnetic field inside the scanner bore that can lead to shifts in the reconstructed image position (as gradients in the magnetic field are used to encode image position (Raj et al., 2001)). When the TR becomes half or less than respiration interval, the effect of these field changes is no longer aliased into the data, and becomes visible as periodic oscillations in the realignment parameters. This effect was variably present within the main HCP cohort as described below, but importantly it means that realignment-parameter-based measures such as frame displacement (FD) do not represent veridical measures of movement across all HCP subjects because the realignment parameters include both real movements and a notable degree of 'phantom' (i.e., false) movements related to respiration induced magnetic field fluctuations. Notably, both (Burgess et al., 2016) and (Power, 2017) have similarly commented that FD estimates in HCP data do not exhibit the same utility for flagging 'motion corrupted' time points as has been observed in traditional (slower TR, larger voxel) data.

Another approach to identifying movement is the use of "DVARS" (Burgess et al., 2016; Power et al., 2012; Smyser et al., 2010), which equals the root-mean-square (RMS) of the frame-to-frame image intensity derivatives. DVARS is attractive because what we care most about from an image denoising perspective are image intensity changes, not whether variations in the magnetic field are causing spatial shifts in the image. Such image intensity changes often show up as "spikes" in the original timeseries DVARS trace. Conversely, however, motion is not the only potential source of DVARS spikes. These might also be caused by a variety of MR physics etiologies, including radio-frequency interference and coil or other scanner instabilities. Additionally, DVARS may be biased by global or semi-global fluctuations from either physiological or neural sources as these cause coordinated intensity changes across the brain.

We are particularly interested in those artifacts that potentially influence the data even after sICA+FIX cleanup. After sICA+FIX cleanup, some DVARS spikes disappear entirely, returning the DVARS to its baseline level, whereas others become "dips" instead of spikes (a few spikes remain and we include them in our measure to be more conservative). These dips occur because sICA+FIX is removing substantial variance from their corresponding timepoints, above and beyond that which would be needed to return the spike to the baseline DVARS value. Indeed, this is an inevitable and expected outcome when modeling a frame-specific artifact, as not just the artifact, but some or even all of the residuals are also removed. As an illustration in the limit, a simplified regressor in a 10-timepoint timeseries of 0000100000 (i.e., a “spike regressor”) is functionally equivalent to scrubbing timepoint 5 from the data and replacing it with the mean image of the other frames, thus completely removing any temporal variance that was previously specific to timepoint 5. Indeed, if all of the structured signal is removed from the data (see Section #3), it becomes clear that these DVARS dips occur as the result of removing some of the unstructured (Gaussian) noise from these timepoints. Panels 2 and 3 of Supplementary Figure 2 illustrate the similarity between DVARS computed from the unstructured noise timeseries versus the sICA+FIX cleaned timeseries (see Section #3 above); however, using the unstructured noise timeseries has the benefit that global structured fluctuations will have minimal influence on the DVARS measure, as the global signal variance of the unstructured noise timeseries is only 6% of that of the sICA+FIX cleaned timeseries of the resting state data. These DVARS dips reflect time segments in which the sICA+FIX cleanup solution begins to approach the scrubbing solution. However, an important benefit of sICA+FIX is that, unlike scrubbing, it is not constrained to make an all-or-nothing binary decision based on a chosen threshold (of FD or DVARS). Rather, it behaves like a weighted (“softer”) form of scrubbing, while at the same time handling subtler artifacts that would be below the scrubbing threshold.

Supplementary Figure 1 illustrates the run-wise relationship between the number of DVARS Dips and mean FD, while the color of each point on the scatter plot reflects the within-run (Pearson) correlation between FD and DVARS of the unstructured noise timeseries. (This is an inverse correlation because FD spikes are positive and DVARS Dips are largely negative). Most runs have both few DVARS dips (e.g., fewer than 20, out of a complete run length of 1200 frames) and low mean FD (e.g., less than 0.2) and thus have little physical head motion and are unaffected by phantom head motion. Some runs with more significant physical head motion but little phantom head motion show high within-run correlation between DVARS and FD (e.g., absolute value of r=0.5 or greater). Other runs have minimal physical head motion and substantial phantom head motion and low correlation between DVARS and FD at the run level. Still other runs fall in between, having both real physical head motion and phantom head motion (i.e., mean FD increased above what would be expected by the DVARS dips). Conspicuously absent are runs with high DVARS Dips but very low mean FD (i.e., upper left of figure), suggesting that DVARS Dips are a more specific measure of physical head motion than is FD.

Supplementary figures 2 and 3 show exemplar runs from various locations on the scatter plot that span the parameter space. Visual inspection of the original unprocessed volume timeseries of these exemplar runs (which are available at db.humanconnectome.org) indicate that DVARS dips represent locations where there are obvious disruptions of the raw image signal intensity due to head motion, whereas runs with high mean FD unaccompanied by DVARS Dips show rapidly fluctuating small shifts in image position (with a periodicity consistent with respiratory frequency) without obvious disruptions of image intensity. Further investigation suggests that body weight or BMI may substantially influence the amount of phantom head motion much more than real physical head motion. Table 1 shows that mean FD is substantially more correlated with body weight and BMI than are DVARS Dips. For these reasons, we use DVARS Dips as our primary index of physical head motion in this study. The time periods of the DVARS Dips (or if there are any residual positive DVARS spikes) warrant special attention to ensure that no residual structured noise remains. As shown in the main results, noise tICA components are often stronger during DVARS Dips. Also this analysis suggests that the correlation between phantom head motion and weight or BMI may have led to inflated estimates of the relationship between motion (as measured by FD) and weight or BMI in HCP data (Hodgson et al., 2016).

**Table 1** shows the run-wise Pearson correlations of mean FD, DVARS Dips, subject weight and subject BMI for the resting state fMRI data of the 449 subjects in this study.

| | mean FD | DVARS Dips | Weight | BMI |
|---|---|---|---|---|
| mean FD | 1.0000 | 0.3480 | 0.4730 | 0.5809 |
| DVARS Dips | 0.3480 | 1.0000 | 0.1204 | 0.0823 |
| Weight | 0.4730 | 0.1204 | 1.0000 | 0.8496 |
| BMI | 0.5809 | 0.0823 | 0.8496 | 1.0000 |

**Figure 1** shows the relationship between mean FD and the number of DVARS Dips (+/-25 from median DVARS) across all resting state runs. The color of each dot indicates the within run correlation of FD and DVARS of the unstructured noise timeseries. Because FD is positive and DVARS Dips are generally negative, these measures are inversely correlated. The circled dots indicate runs whose FD and unstructured noise DVARS are shown below in Figures 2 and 3.

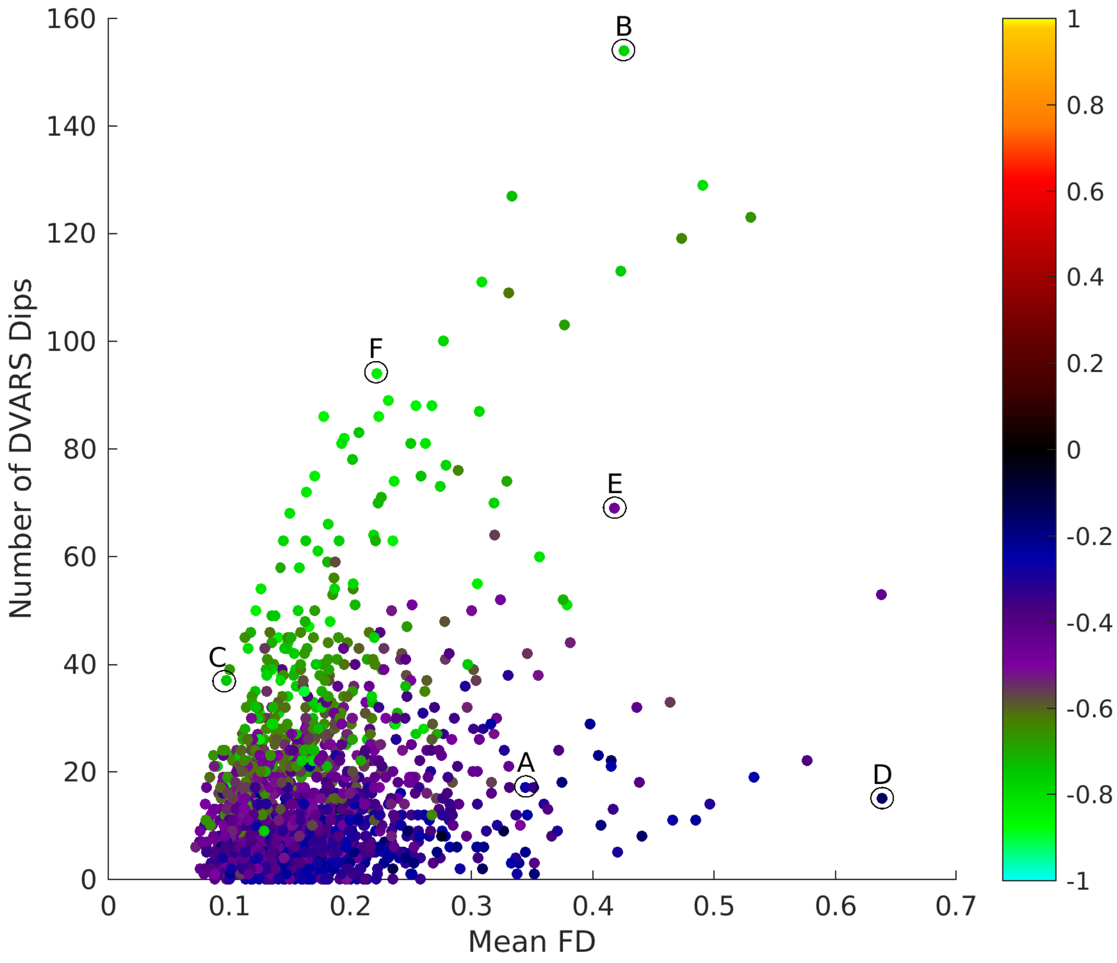

**Supplementary Figure 2** shows FD and DVARS traces (with +/-25 DVARS thresholds) from three resting state fMRI runs that were circled with letters in the above scatter plot. The first run (A) shows the discrepancy between high mean FD (Row 1) but only a few DVARS spikes (Rows 2 and 3) in a subject with "phantom" motion (likely induced by respiration-related magnetic field changes in a subject with high body mass index (BMI=35)—see Section #4 above). Rows 2 and 3 also illustrate that the DVARS of the sICA+FIX cleaned data and the unstructured noise are very similar. Rows 4 and 5 show FD and DVARS traces from a subject (BMI=26) who moved a lot, but in which FD and DVARS agree relatively well (B). Rows 6 and 7 show FD and DVARS traces from a subject (BMI=20) who only moved a little, and in which FD and DVARS again have good correspondence (C).

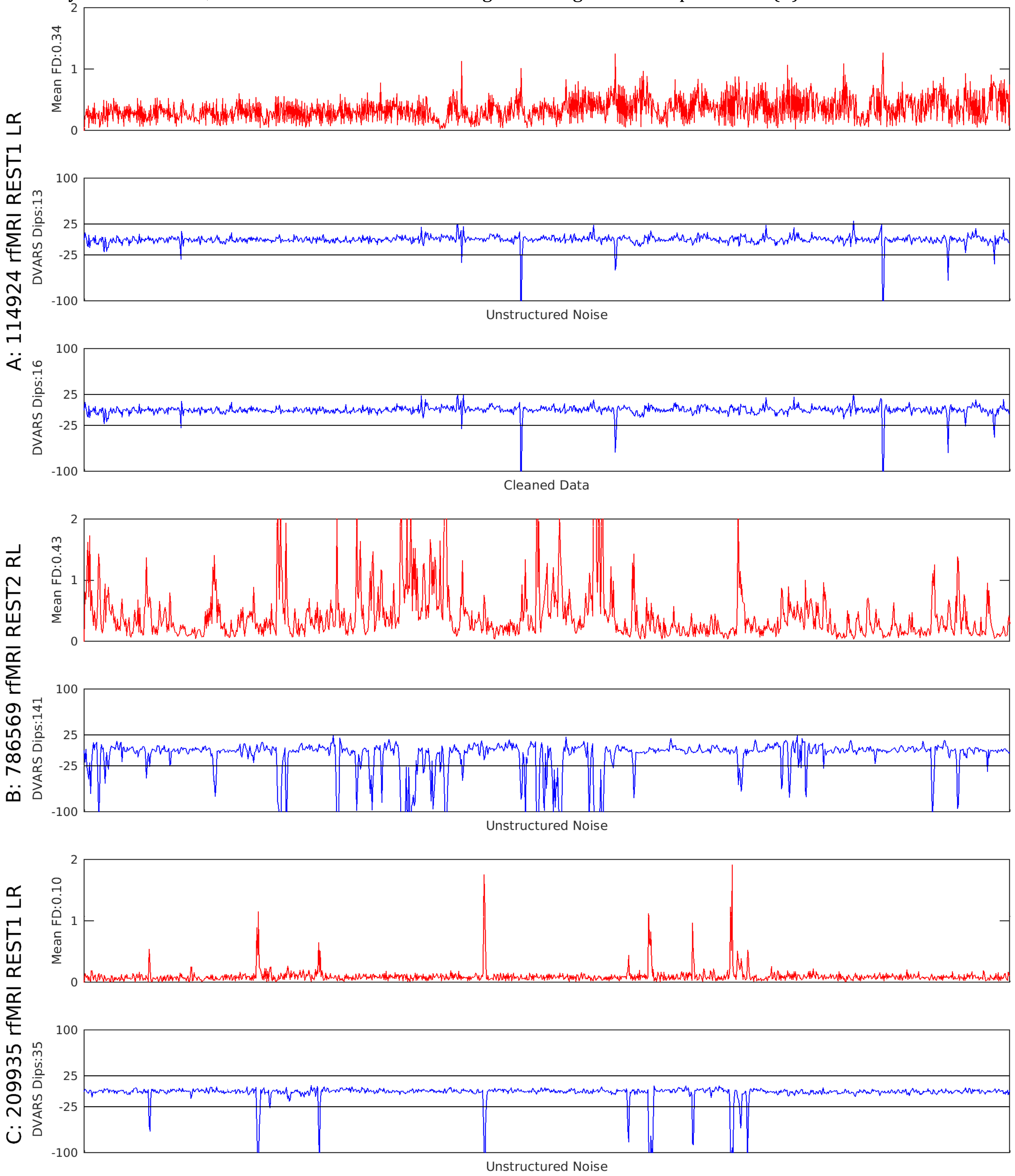

**Supplementary Figure 3** shows FD and DVARS traces (with +/-25 DVARS thresholds) from three more resting state fMRI runs that were circled in the above scatter plot. The first run (D) shows the discrepancy between high mean FD (Row 1) but only a few DVARS spikes (Row 2) in a subject with phantom motion and BMI=34. Rows 3 and 4 (E) show FD and DVARS traces from a subject (BMI=27) who moved a lot, but also had substantial phantom motion. Rows 5 and 6 (F) show FD and DVARS traces from a subject (BMI=21) who moved a moderate amount, and in which FD and DVARS again have good correspondence.

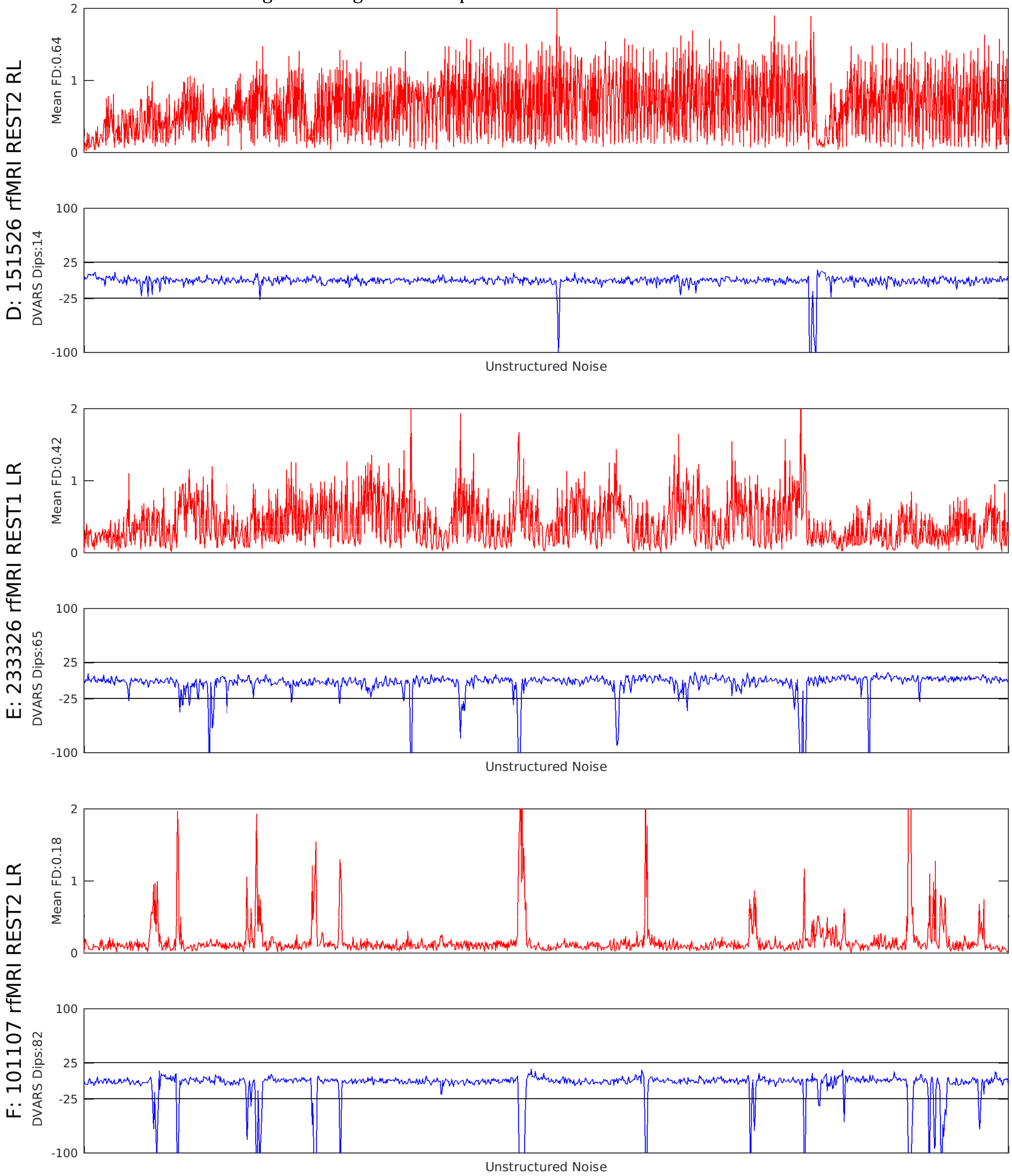

**Supplementary Results and Figures**

*5. Anti-correlated CSF in Global or Semi-global Components:* One initially puzzling feature of the global and semi-global components like TC1 is the anti-correlation between the ventricles and the grey matter. This kind of anti-correlation is also seen in other global or semi-global components that are categorized as noise (TC1, TC8, TC30, TC38), signal (TC6, TC15, TC20), or even are specifically task modulated (TC2-3, TC9, TC28, TC33). We suspect this anti-correlation is related to changes in global or semi-global blood flow in the brain due either to physiological or neural effects. When there is more blood flow (i.e., increase in T2* image intensity in grey matter), the brain tissue expands (Bright et al., 2014; Thomas et al., 2013), and this pushes CSF with bright signal on T2* images out of the ventricles and into the space around the spinal cord. This would lead to a decrease in ventricular signal intensity, particularly around the ventricular margins. Conversely, when blood flow drops, the CSF volume is restored and the ventricular signal intensity around the margins would increase (Bright et al., 2014; Thomas et al., 2013) at the same time the T2* single intensity in the grey matter was decreasing. Additionally, flowing CSF will have a lower T2* image intensity because of dephasing in the centers of CSF spaces (Chen et al., 2015; Yildiz et al., 2017). Analogous correlations among T2* BOLD intensity, blood flow, and CSF volume have previously been reported in cat visual cortex in response to visual stimulation (Jin and Kim, 2010). Thus, we do not consider anti-correlation in the CSF by itself to be an indicator of a component being either noise (i.e., of physiological origin) or signal (i.e., of neural origin); rather it may be a function of the spatial extent of the blood changes related to a given component.

**Various Supplementary Figures mentioned in the Main Text follow:**

**Supplementary Figure 4** shows the concatenation of the task fMRI design matrices (for ease of visualization, the linear trends and means were not removed), which is the order in which the task fMRI scans were concatenated. The rows alternate regressors and their temporal derivatives. For each task, there are two phase encoding directions ordered RL first and then LR (each black x-axis tick is a task run boundary).

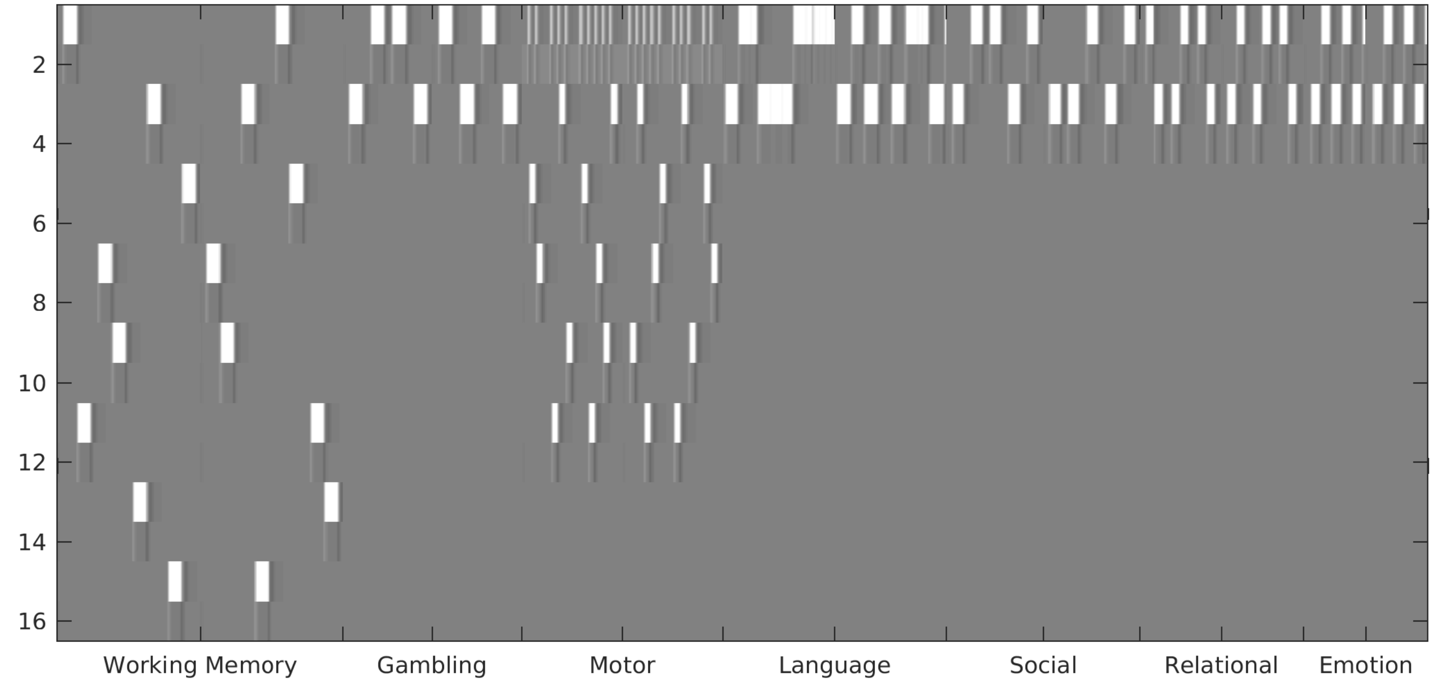

**Supplementary Figure 5** compares task fMRI tICA Component 7 and the Working Memory Face contrast (spatial correlation=0.97; other spatial correlations greater than 0.95 include the Working Memory Body contrast and the Working Memory 2BK contrast). In all the figures of this form, each map is scaled with a min/max corresponding to the 2nd/98th percentiles.

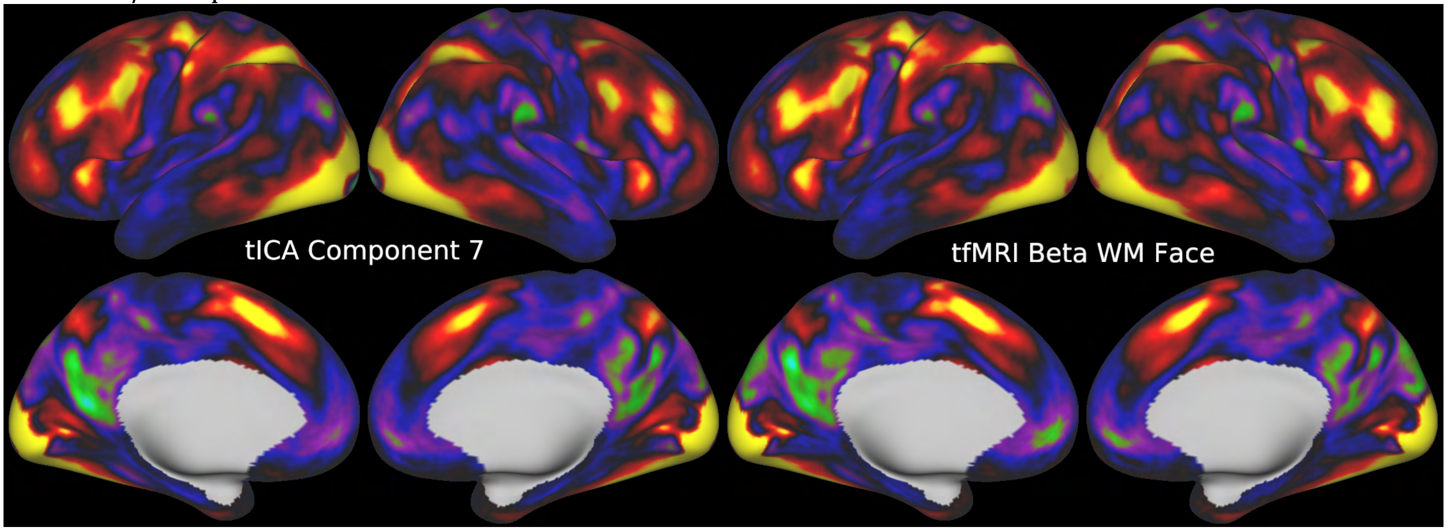


**Supplementary Figure 6** compares task fMRI tICA Component 18 and the Working Memory Place - Average contrast (spatial correlation=0.93).

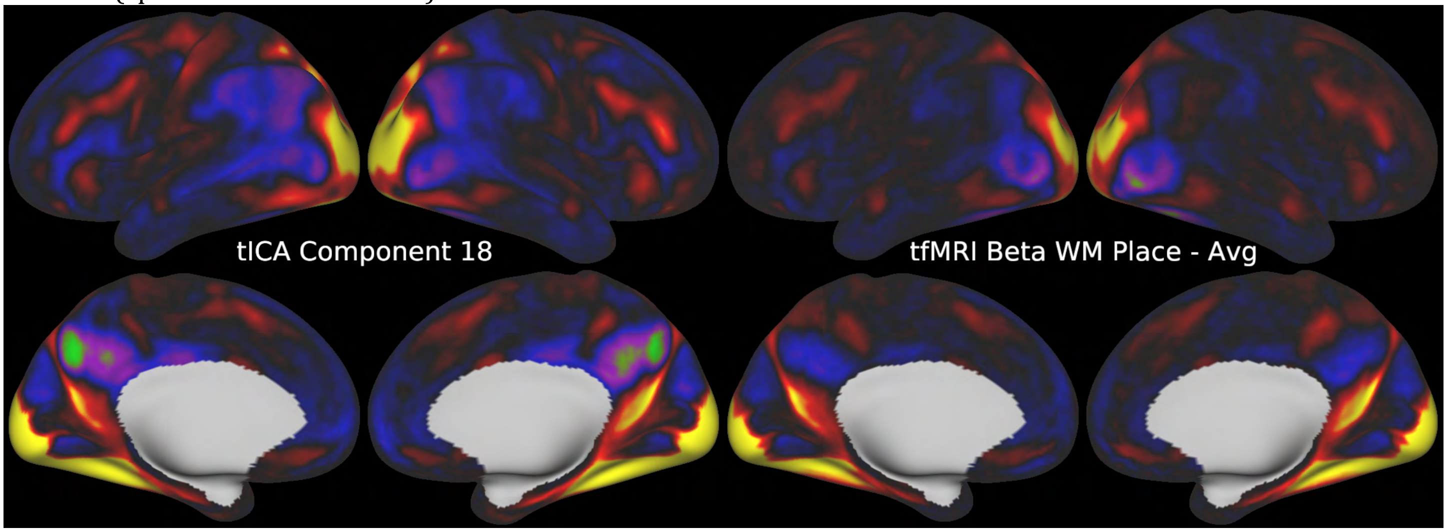


**Supplementary Figure 7** compares task fMRI tICA Component 16 and the Motor Face - Average contrast (spatial correlation=0.92).

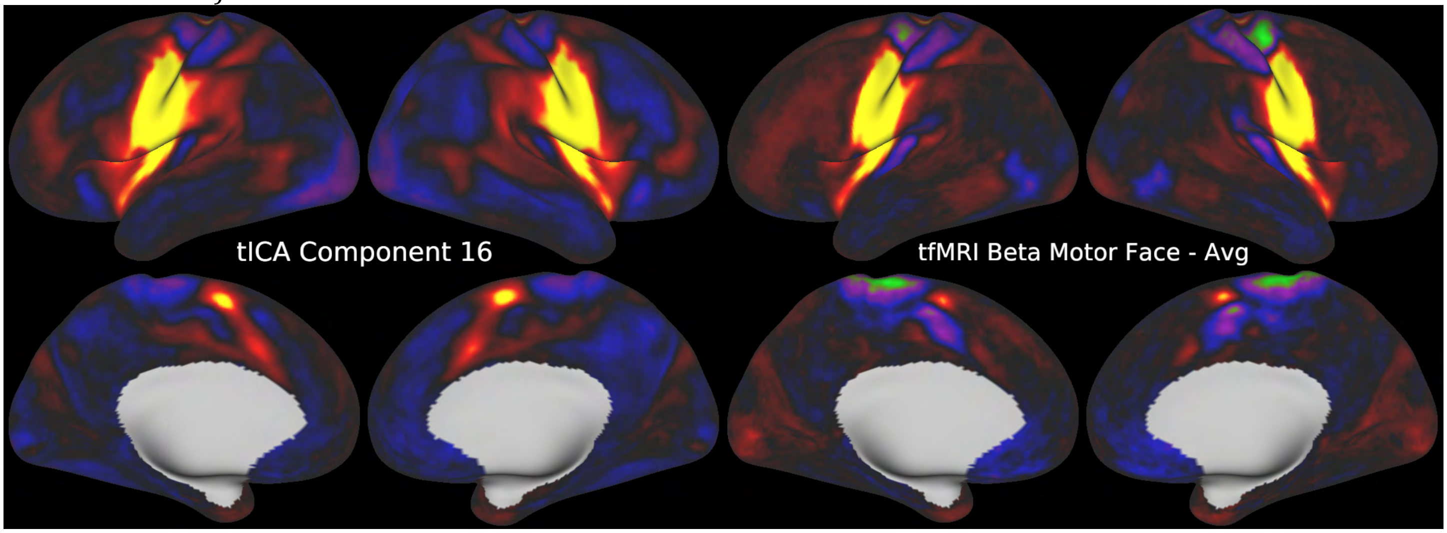

**Supplementary Figure 8** compares task fMRI tICA Component 19 and the average of the Motor Left Foot and Right Foot contrasts (spatial correlation=0.91).

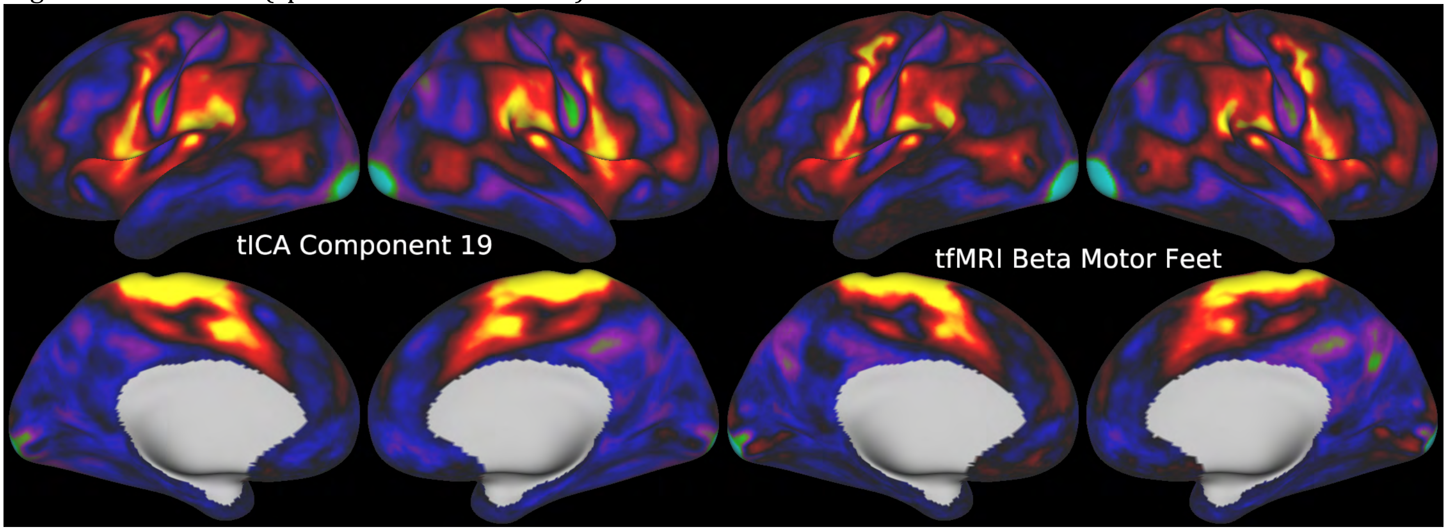


**Supplementary Figure 9** compares task fMRI tICA Component 31 and the Motor Left Hand contrast (spatial correlation=0.85).

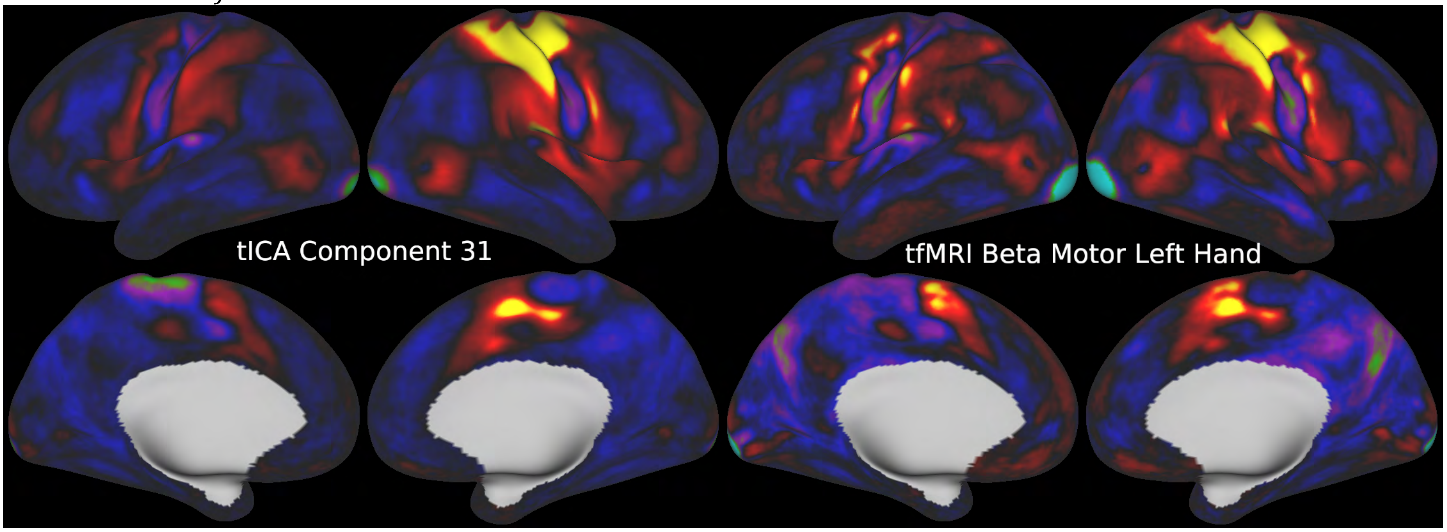


**Supplementary Figure 10** compares task fMRI tICA Component 6 and the Motor Right Hand contrast (spatial correlation=0.53).

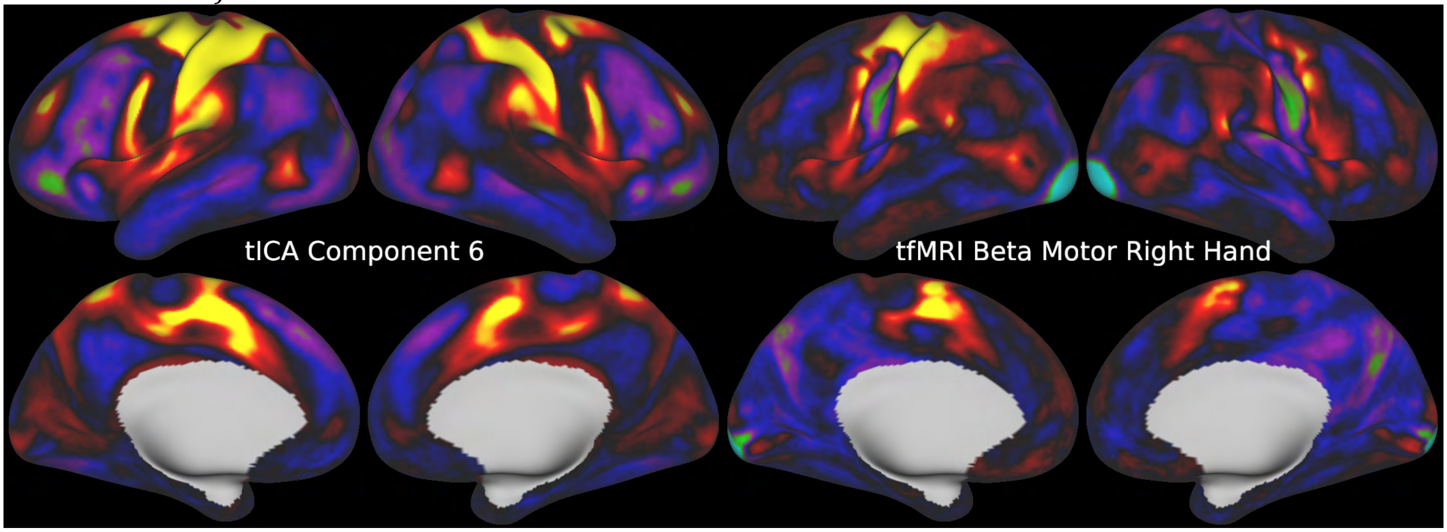

**Supplementary Figure 11** compares task fMRI tICA Component 11 and the Language Story contrast (spatial correlation=0.89).

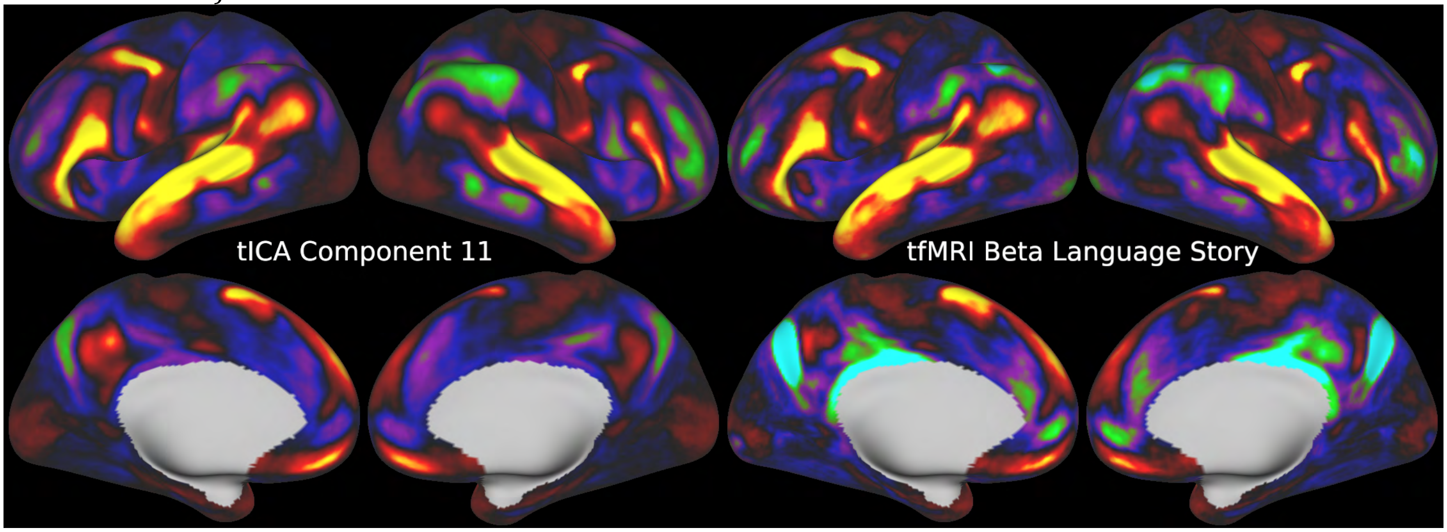


**Supplementary Figure 12** compares task fMRI tICA Component 4 and the Language Math-Story contrast (spatial correlation=0.88).

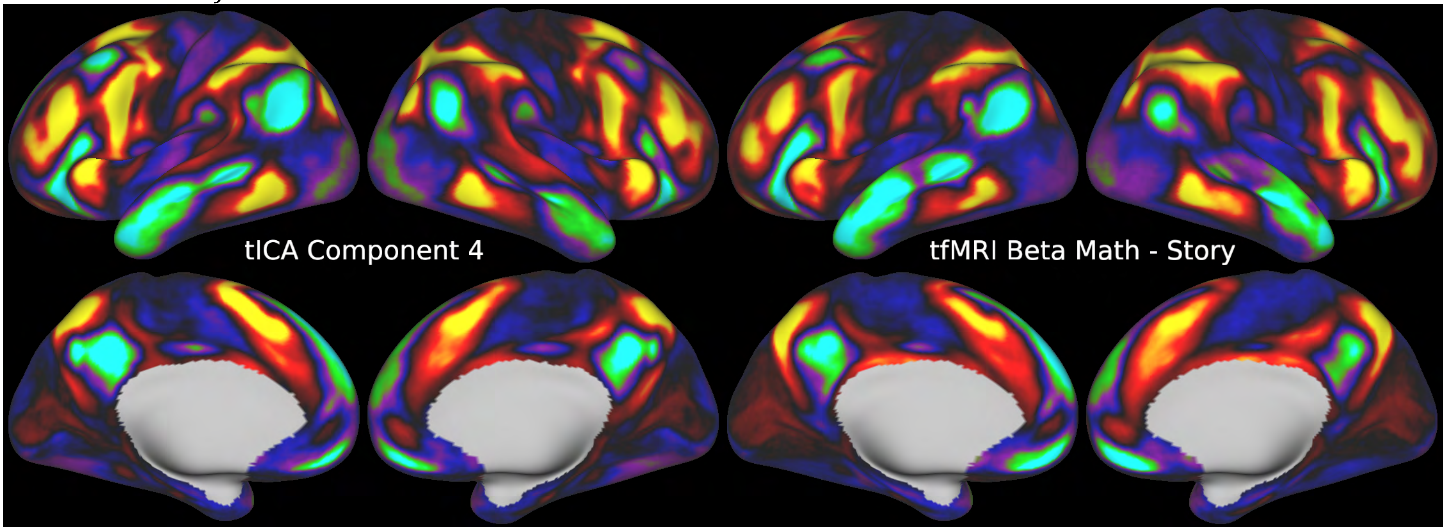


**Supplementary Figure 13** compares task fMRI tICA Component 2 and the average of the Social task Random and TOM contrasts (spatial correlation=0.97).

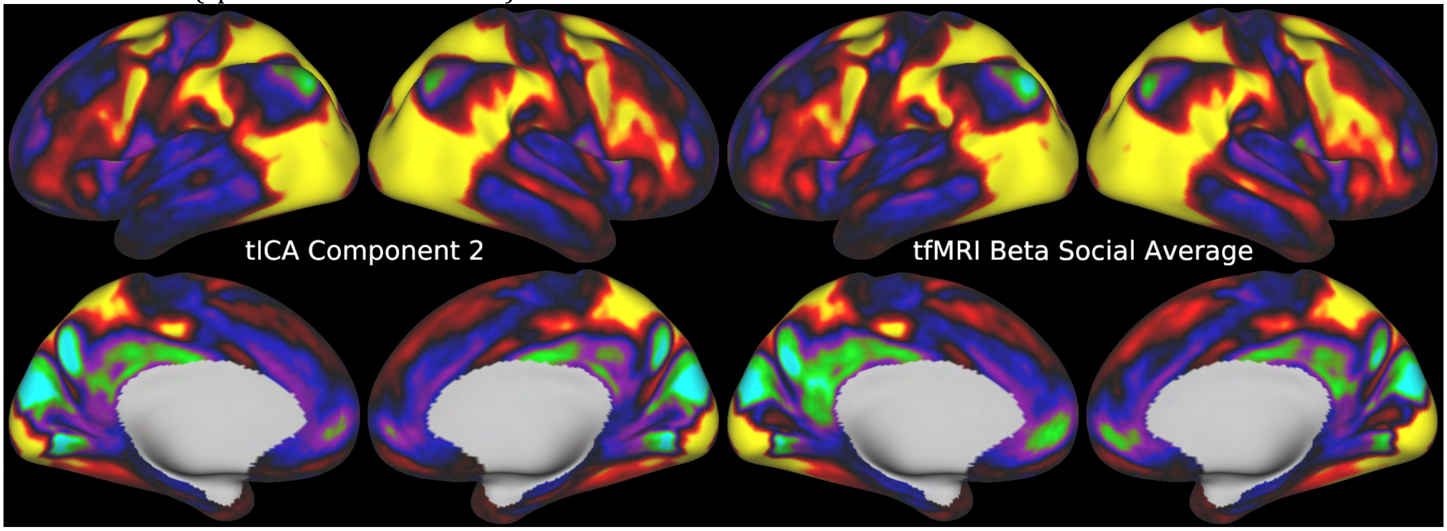

**Supplementary Figure 14** compares task fMRI tICA Component 3 and the average of the Relational task Relational and Match contrasts (spatial correlation=0.95).

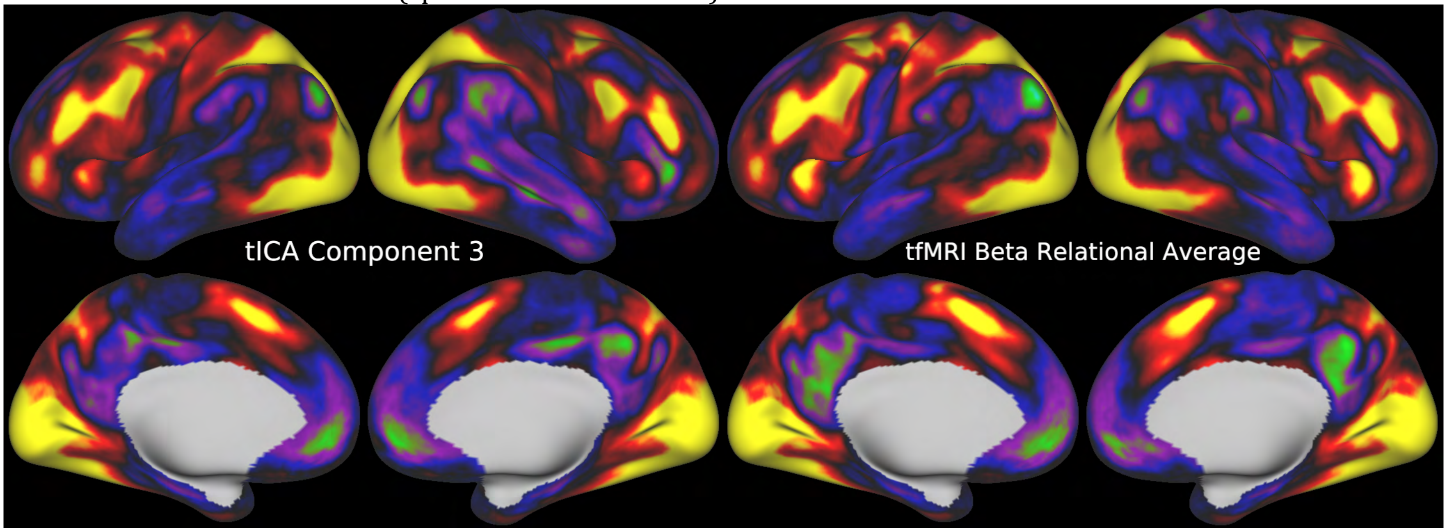


**Supplementary Figure 15** compares task fMRI tICA Component 37 and the Emotion Faces - Shapes contrast (spatial correlation=0.89).

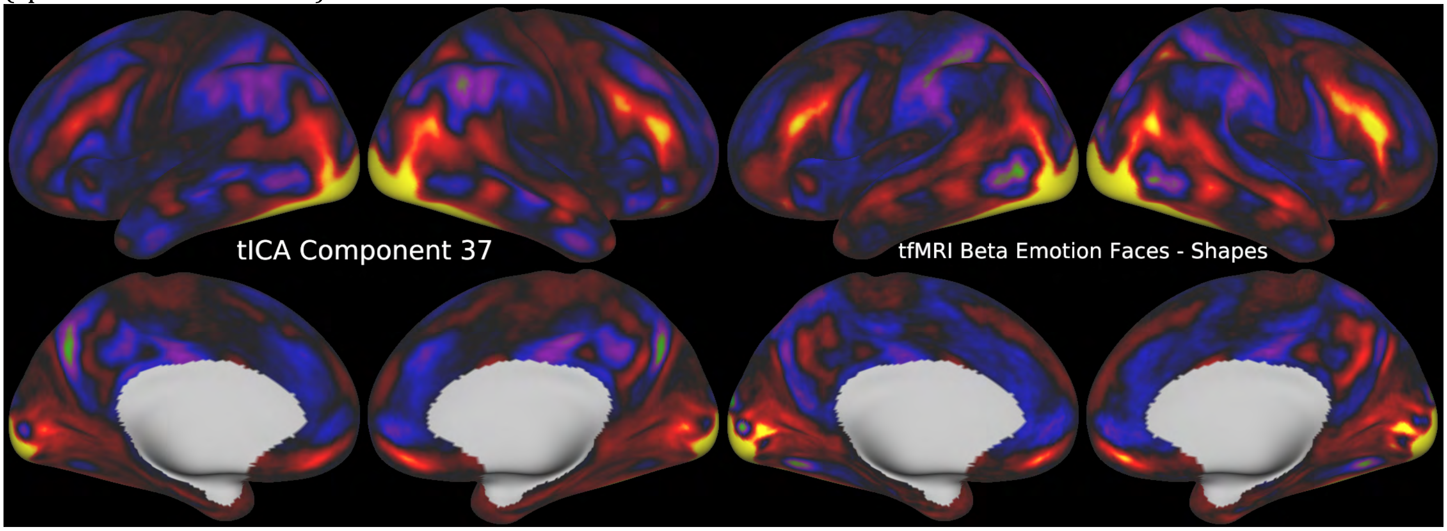

**Figure 16** shows the same four useful plots that were shown in Main Text Figure 3 for the tICA components generated from the residuals after fitting the task design. In RVT correlation (Panel A) the line is at 0.1. In DVARS dips normalized amplitude difference (Panel B) the line is at 0.25. In cross-subject component amplitude variability (Panel C) the line is at 0.15. In single subject component identification (Panel D) the line is at 0.4. The results are consistent with Main Text Figure 3 for the components that have similar spatial maps to those found before regressing out the task GLM and rerunning the weighted regression and temporal ICA.

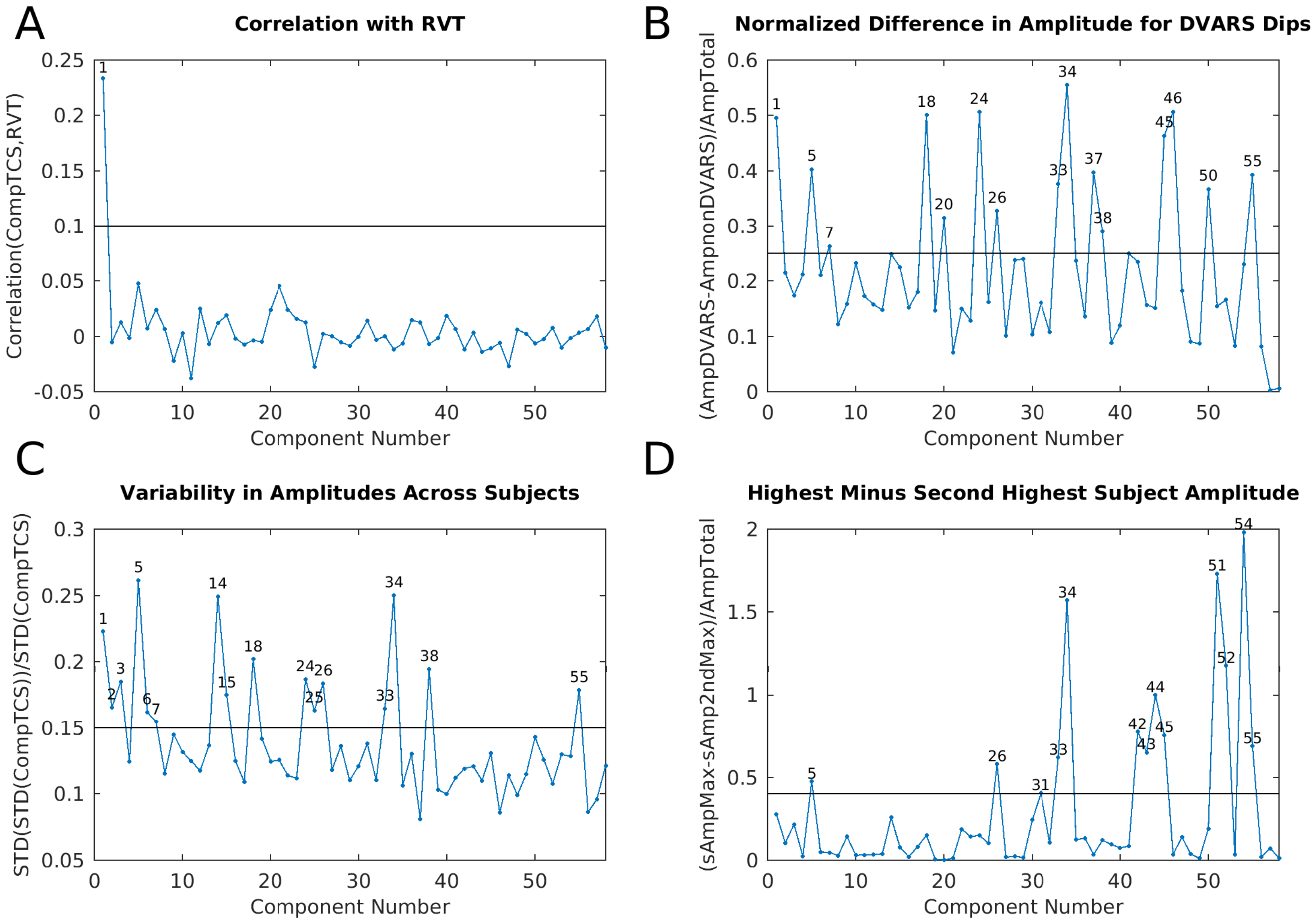

**Supplementary Figure 17** shows the correlation matrix of component amplitudes reordered using hierarchical clustering for the task fMRI data (using FSLNets with default settings). For each component, there were 898 (2 sessions * 449 subjects) amplitudes, and those amplitudes were correlated across tICA components to generate the correlation matrix of component amplitudes. The components are numbered according to the TC Supplementary Figures and whether they were classified as signal or noise is indicated. Components generally cluster according to signal (purple, neon green, blue) vs noise (red) classification, though there are some exceptions (i.e., the green cluster). Not surprisingly, components that are strongly task modulated and were scanned on a particular day cluster together and are anti-correlated with the components scanned on the other day (purple vs blue), as each day is a separate concatenated task session. Between these are spontaneous components that are generally not modulated by a specific task (neon green).

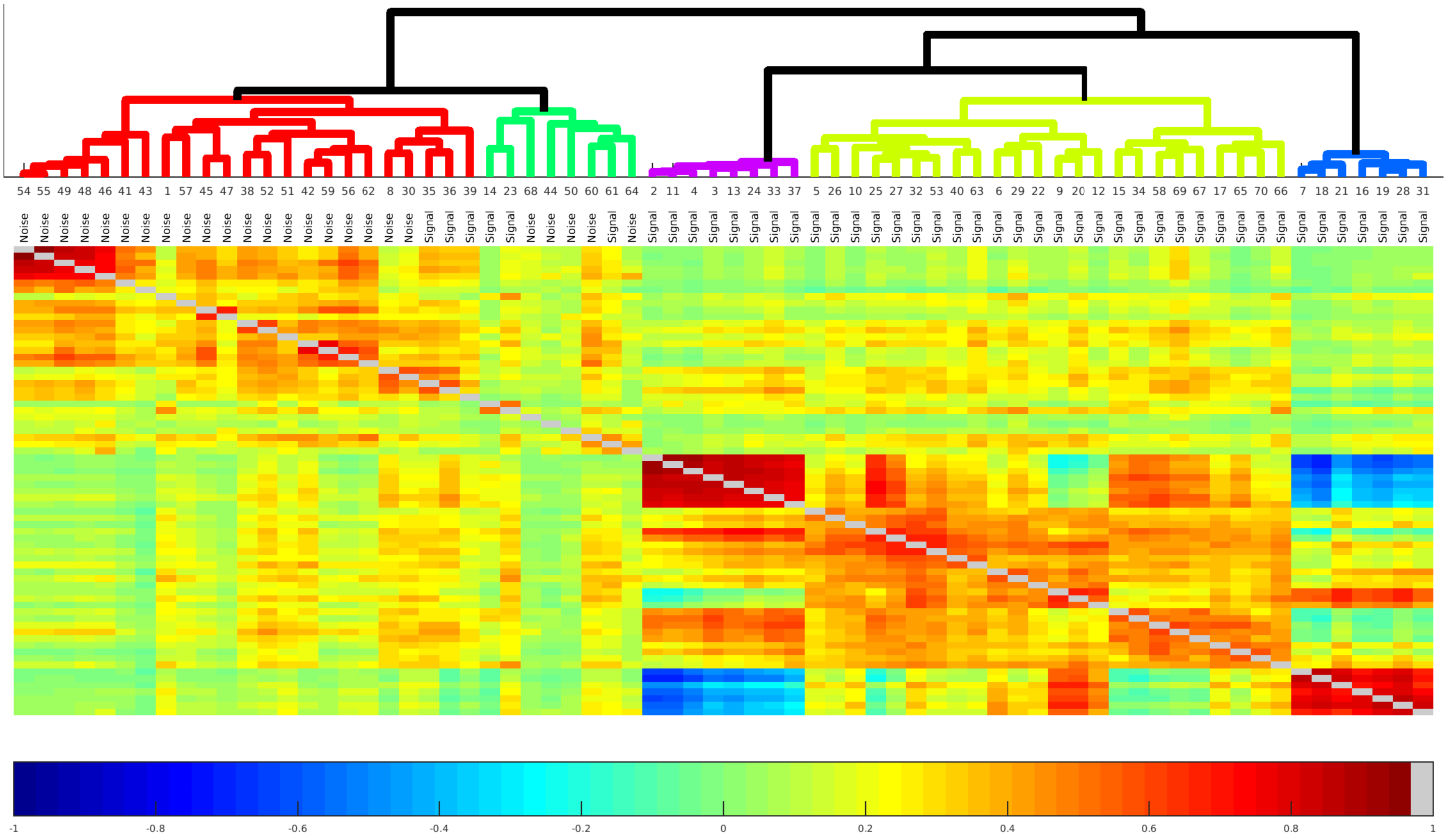

**Supplementary Figure 18** shows the correlation matrix of component amplitudes reordered using hierarchical clustering of the tfMRI residual data after fitting the task design. The components are numbered according to the TCr Supplementary Figures. Again, components generally cluster together according to their signal (purple and neon green) vs noise (cyan) classification (the red cluster is an exception). The components that remain task modulated after regressing out the GLM cluster together (purple), as do the spontaneous components (neon green).

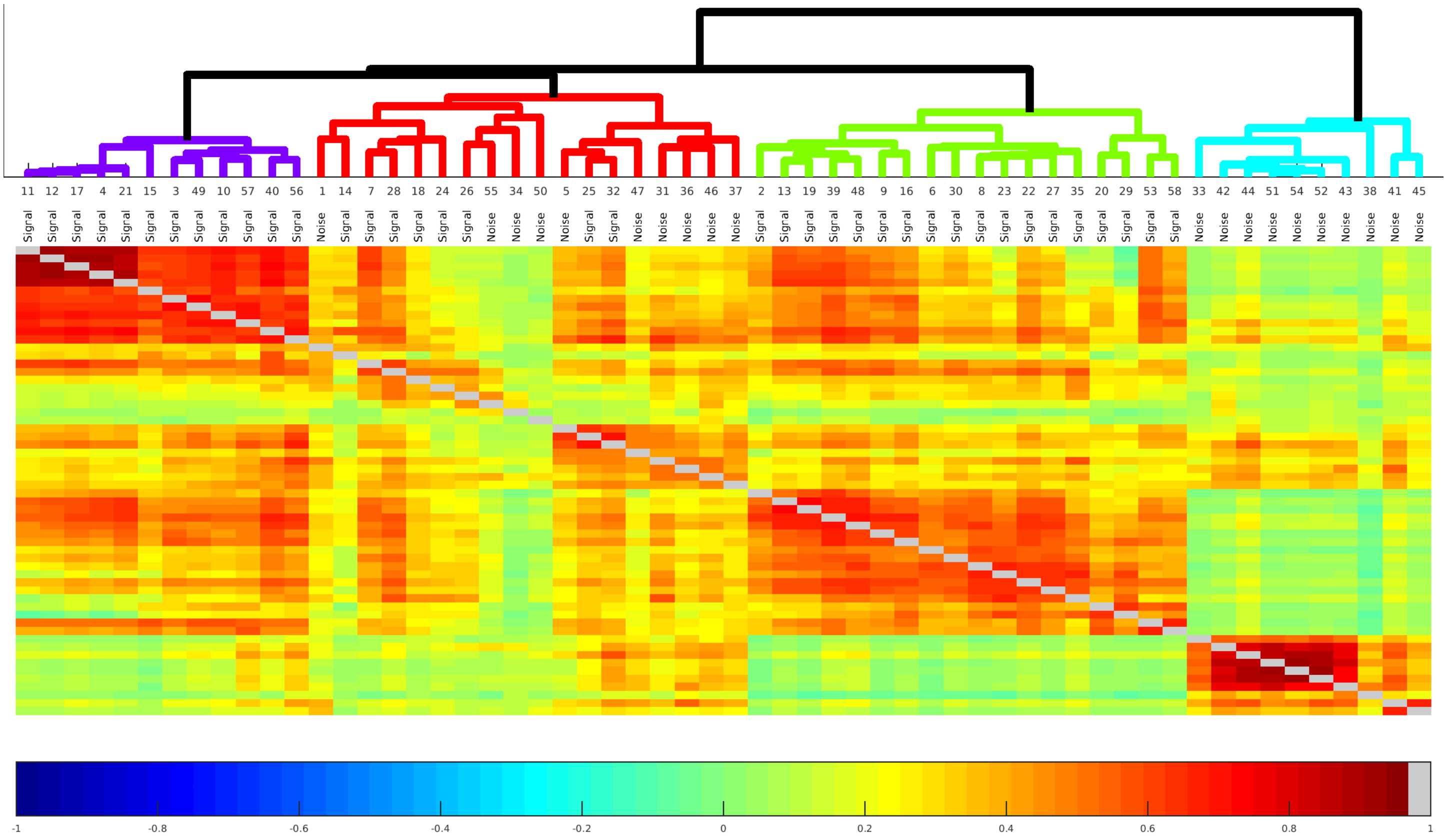

**Supplementary Figure 19** shows the betas of the tICA components from a regression onto the mean grey timecourse of the sICA+FIX cleaned task fMRI data (red) and the sICA+FIX + tICA cleaned task fMRI data (green). Global noise components 1, 8, 30, 38, 52, and 62 in particular are reduced nearly to zero by the tICA cleanup (though not exactly zero because the temporal ICA is only orthogonal at the group level, not for each individual concatenated session), but a number of non-global components that contribute to the mean grey timecourse are also removed by the tICA cleanup (e.g., see several of the components between 40-50 for examples). In contrast, signal components that contribute strongly to the mean grey signal have betas that are minimally affected (as intended) by the tICA cleanup of the task data (see components 20, 28, and 33). One the right side, the scatter plot shows mean grey timecourse variances before vs after tICA cleanup (i.e., sICA+FIX data vs. sICA+FIX + tICA data) of each of the concatenated task sessions of the 449 subjects (with hotter colors representing higher data point density). Note that these values are the variance of the mean grey timecourse (MGT) itself; not the variance removed by mean grey timecourse regression (MGTR) as is reported in the main text (as "MGTRVar"). The mean grey timecourse variance is always reduced by tICA cleanup, but there is still modest variability across subjects. Across almost all subjects, the mean grey timecourse variance is less than 100 after tICA cleanup. The plot is scaled comparable to Supplementary Figure 26.

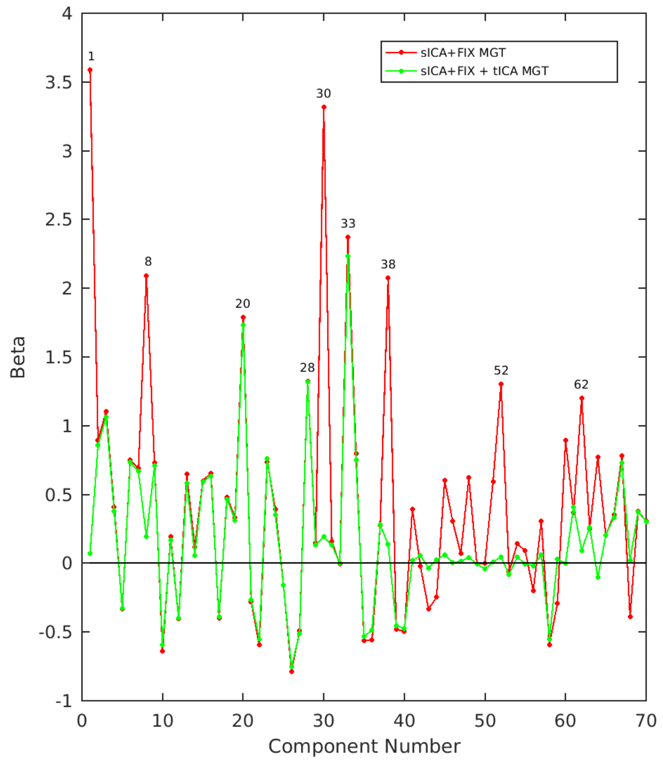


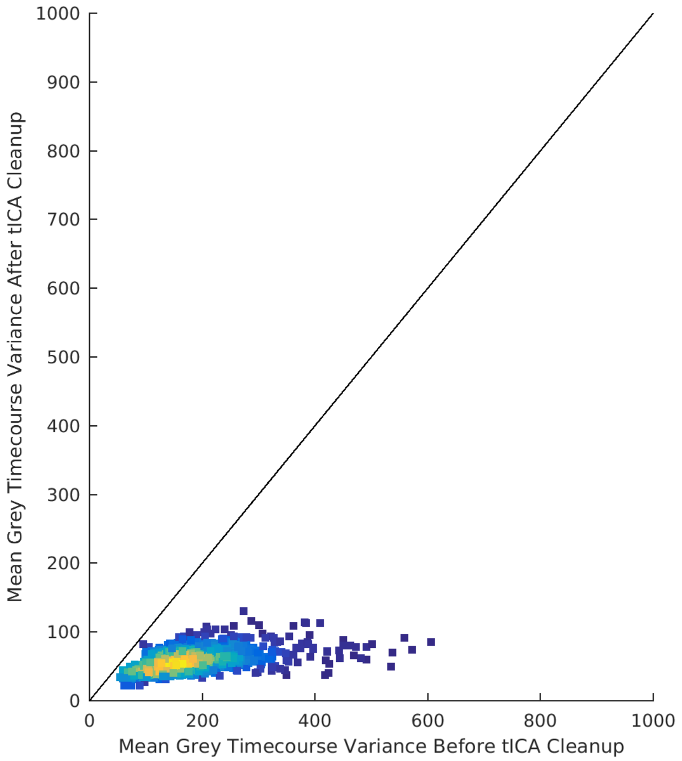

**Supplementary Figure 20** shows the effect of sICA+FIX cleanup on the Motor Tongue contrast. Top row is the standard analysis (i.e., 'minimal preprocessing,' 'MPP'). Middle Row is after sICA+FIX. The false positive "motor" deactivation in the orbitofrontal cortex is eliminated by sICA+FIX (red oval). This along with several other false positive activations are also visible in the volume slices and are likely related to stimulus-correlated motion. The difference between MPP and sICA+FIX is shown in the bottom row.

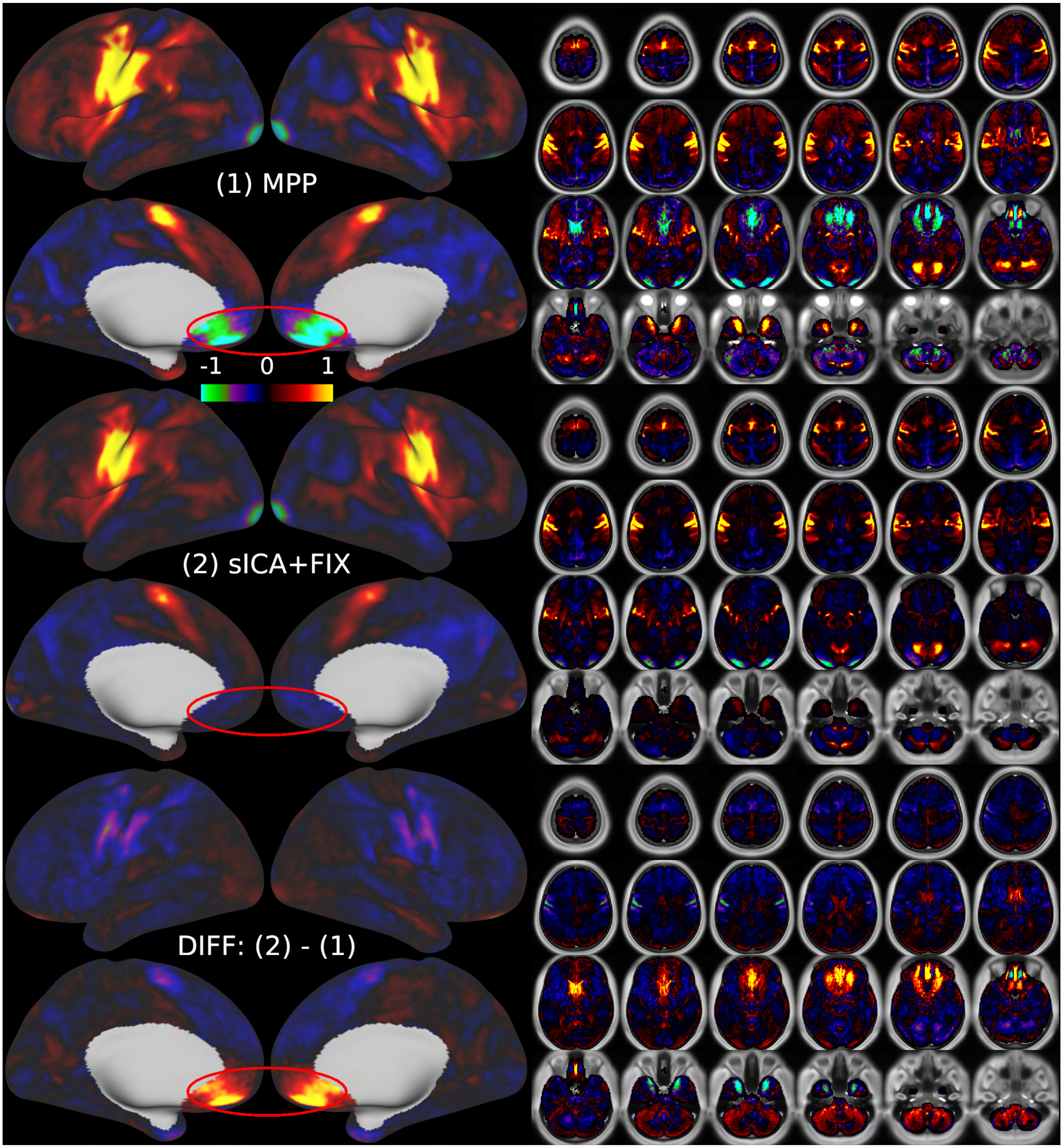

**Supplementary Figure 21** shows the effect of sICA+FIX cleanup on the Language Story contrast. Top row is the standard analysis (i.e., 'minimal preprocessing,' 'MPP'). Middle Row is after sICA+FIX. The negative bias in the contrast map is eliminated and consequently a positive activation in area 9m in the superior medial frontal cortex is revealed (red arrows). The volume slices also demonstrate how CSF signal is especially affected by the bias (likely from incomplete T1 steady state relaxation in the fMRI data because an insufficient number of volumes were discarded from the beginning of the run, which also is the only baseline period in this particular task). Even if the grey matter were not at all affected (it does appear to be somewhat affected), partial volume effects would lead to a bias within some grey matter voxels near the surface. The bottom row shows the difference between MPP and sICA+FIX.

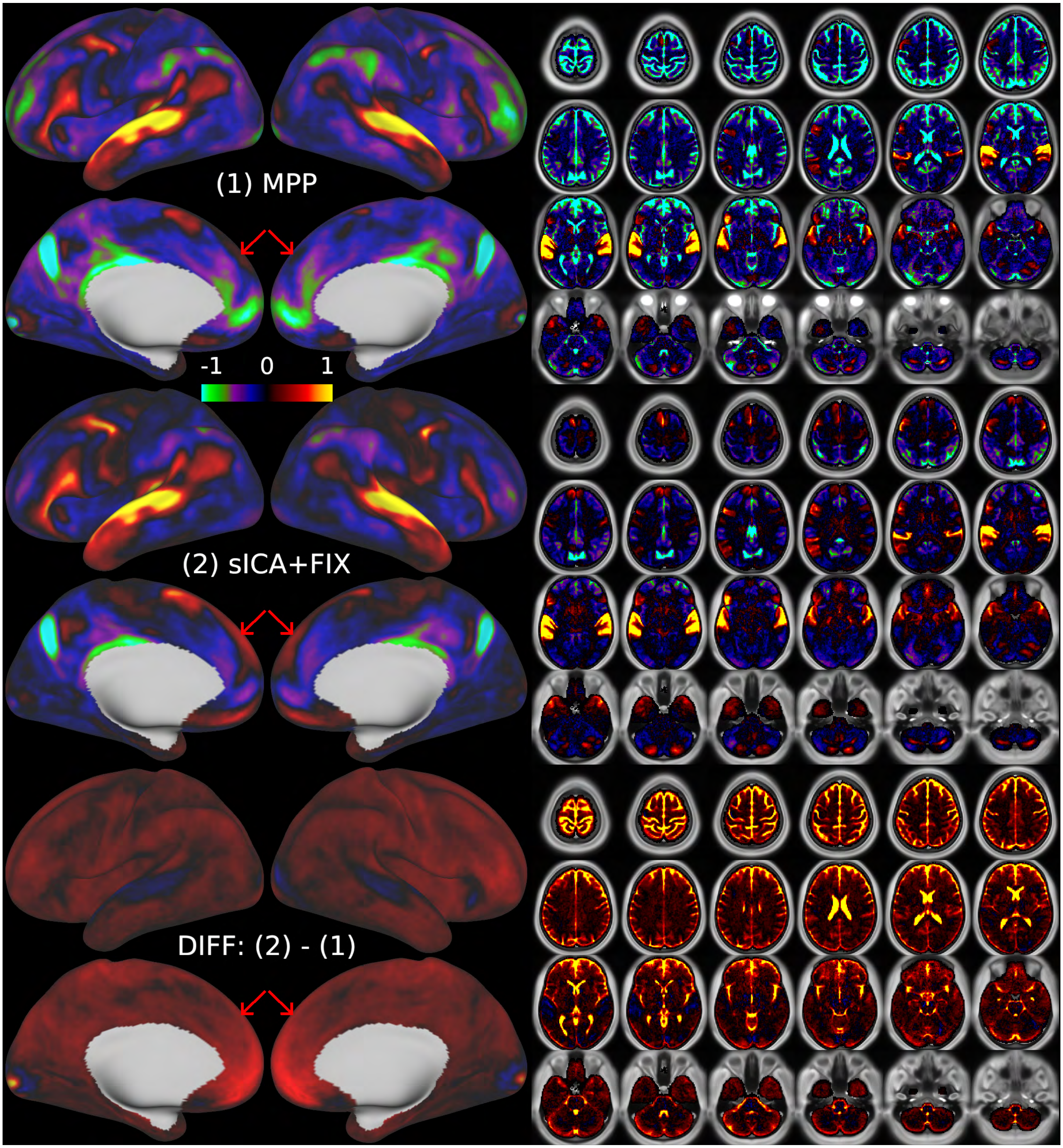

**Supplementary Figure 22** top panel shows the correlation between mean RVT across subjects during the task fMRI runs (blue) and the task "on" blocks from the design matrix (note that the blocks are convolved with the hemodynamic response function used for the HCP's task data). The correlation is r=0.56. The bottom panel shows the consistency of RVT across subjects. Subjects are ordered according to the correlation of their RVT with the mean RVT (highest at top). The 43 subjects at the bottom were missing RVT data entirely. Scale is black = -1, white = 1. Thus, the tasks induce stimulus correlated breathing across many subjects, though the magnitude of this correlation varies from subject to subject.

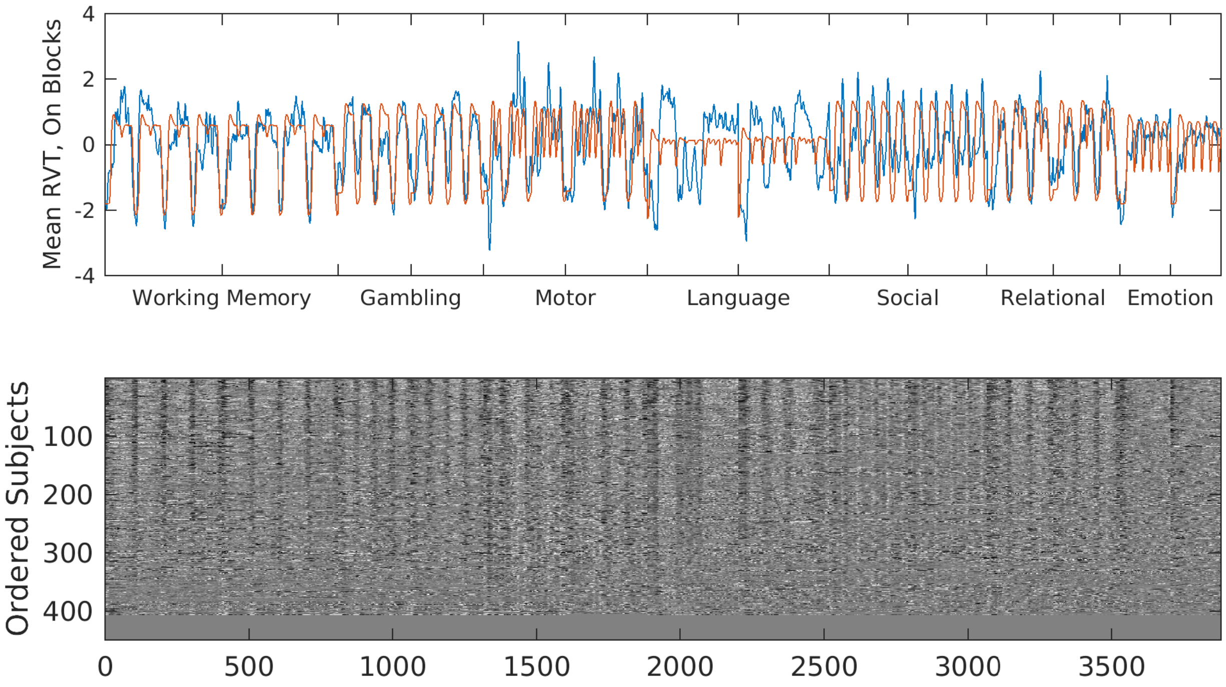

**Supplementary Figure 23** shows the effect of tICA cleanup on the Motor Cue contrast. The Cue period is the brief period in which the subject is instructed which movement to perform (immediately prior to executing the movement itself). Top row is the sICA+FIX analysis. Middle Row is sICA+FIX + tICA. A positive bias caused by stimulus-correlated respiration in the contrast map is reduced. The bottom row shows the difference between sICA+FIX and sICA+FIX + tICA.

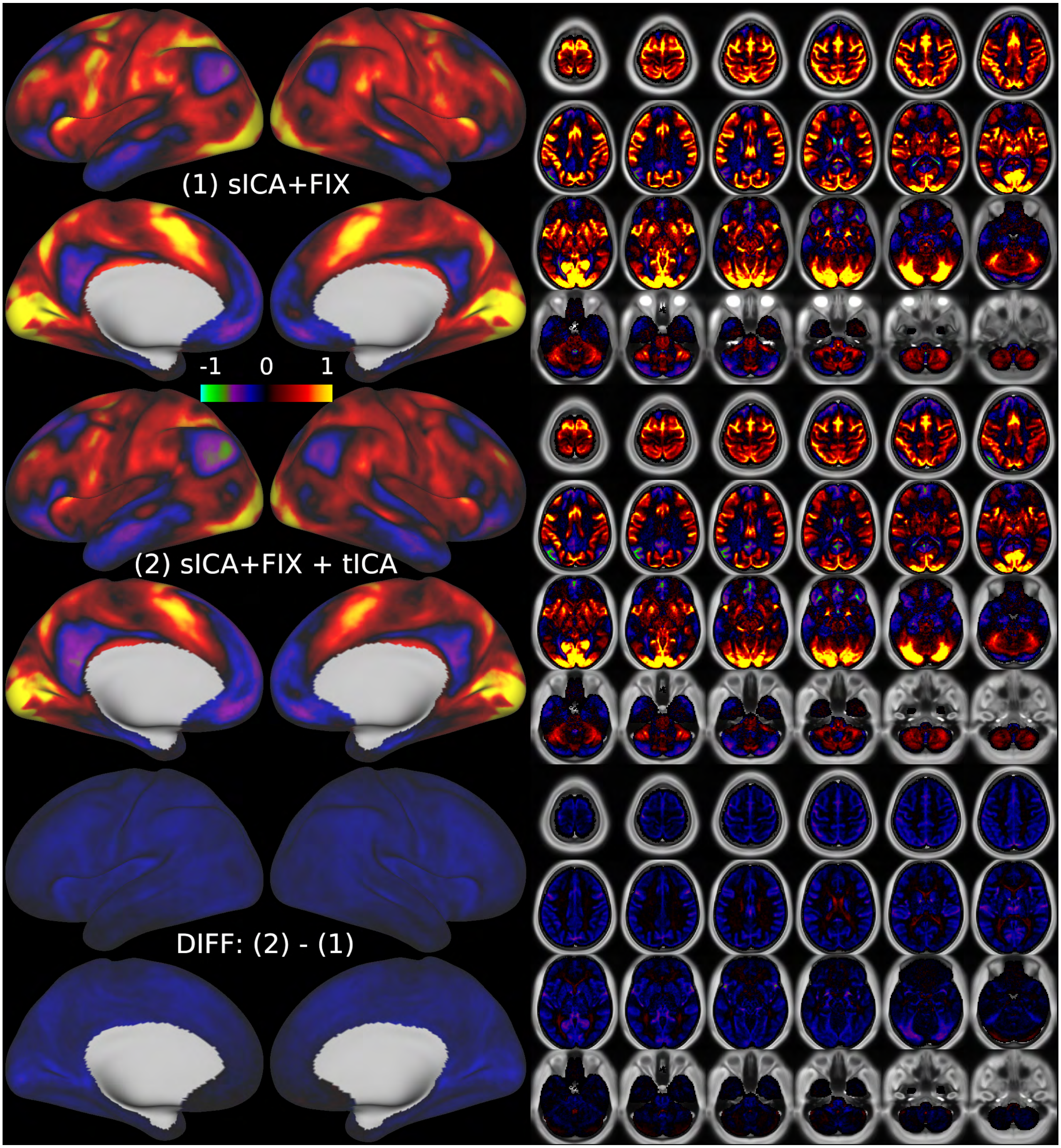

**Supplementary Figure 25** shows a comparison between tICA cleanup and MGTR for the Motor Cue contrast. The top row shows (A) sICA+FIX, (B) sICA+FIX + tICA, and (C) the difference between the two (also shown in Figure 23). A spatially global effect is removed by the tICA cleanup that is very similar to TC1 (spatial correlation r=0.93 with TC1), and has only a modest, unavoidable spatial correlation with the task activation map (r=0.44) because it is globally positive and the task activation map is semi-global. The bottom row shows (D) sICA+FIX + tICA, (E) sICA+FIX + MGTR and (F) the difference between the two. A network-specific effect is removed by MGTR that has a high spatial correlation to the task activation map (r=0.85). The spatial correlation between A and C is r=0.57, between B and C is r=0.44, between D and F is r=0.85, between E and F is r=0.73. Because panels B and D (the same) are our best estimate of the true neural activation in the task, the most informative comparison of spatial correlations is between r=0.44 for the tICA activation map vs. the signal removed by tICA cleanup (i.e., B vs. C) and r=0.85 for the tICA activation map vs. the signal removed by MGTR (i.e., D vs. F) that is in addition to what tICA removes. The higher spatial correlation of D vs F indicates that the effect of MGTR above and beyond that of tICA is to remove neural signal from the task fMRI data.

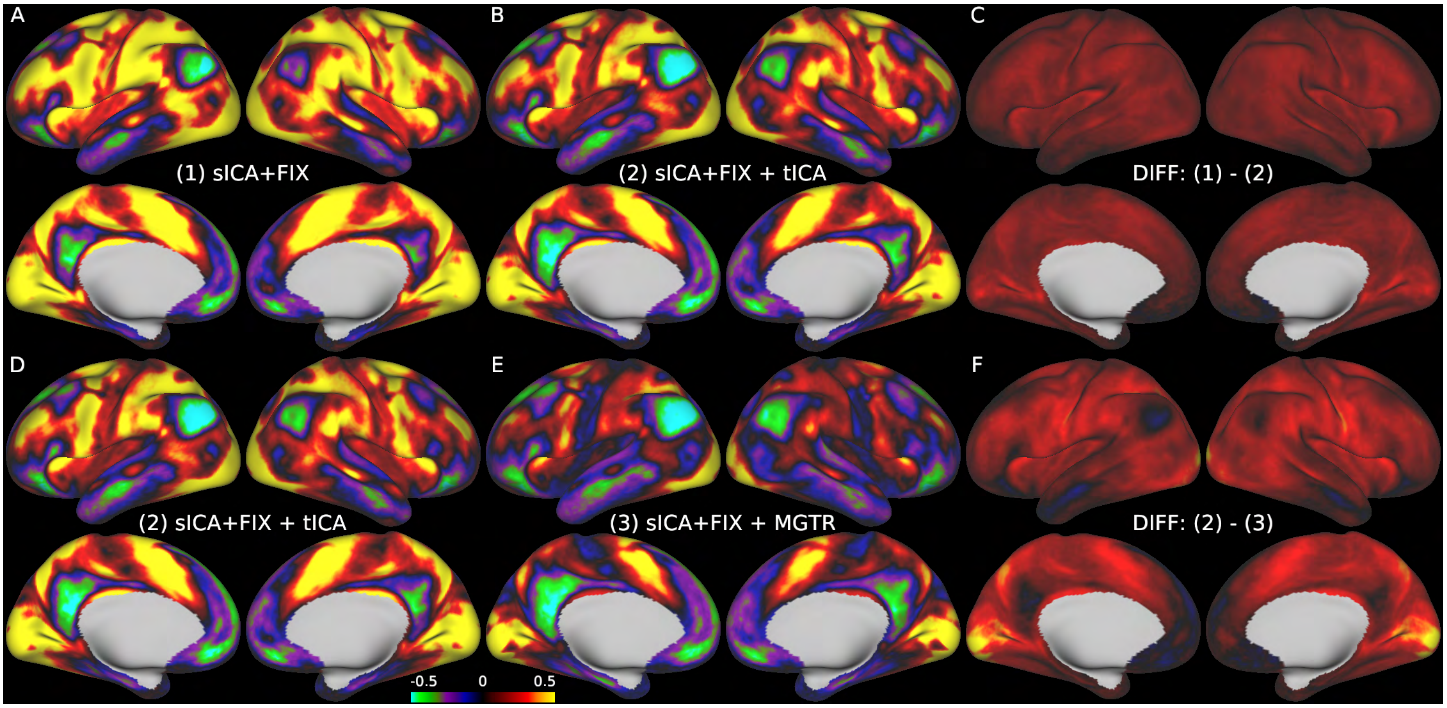


**Supplementary Figure 24** shows the components that have strongly different amplitudes across phase encoding directions in the resting state fMRI data. Components 6 and 8, two of the global components, particularly stand out in this measure. Components 7 and 17 are two signal components for the default mode network and component 47 is a large coil artifact present in only one run of one subject because of an sICA+FIX misclassification that has since been resolved in the released HCP data (as of the 'S1200' release).

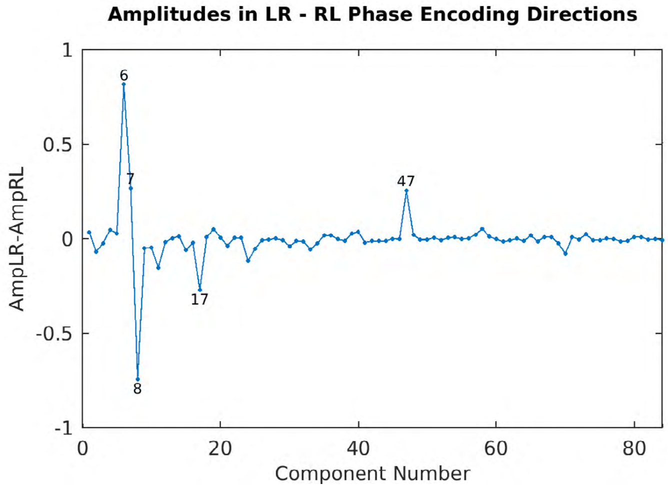

**Supplementary Figure 26.** Left panel shows the betas of the tICA components from a regression into the mean grey timecourse of the sICA+FIX cleaned resting state fMRI data (red) and the sICA+FIX + tICA cleaned resting state fMRI data (green). Global noise components 3, 6, 8, 34, 58, and 80 in particular are large contributors to the mean grey timecourse that are reduced nearly to zero by tICA cleanup, but a number of the non-global components that are removed also contribute to the mean grey timecourse. Components that contribute strongly to the mean grey timecourse, but are neural signal, include 1, 5, and 26, and, as intended, their betas are not much affected by the tICA cleanup. The right panel shows a scatter plot of mean grey timecourse variances before and after tICA cleanup (i.e., sICA+FIX data vs. sICA+FIX + tICA data) across the 4 resting state runs of the 449 subjects (with hotter colors representing higher data point densities). Note that these values are the variance of the mean grey timecourse (MGT) itself; not the variance removed by mean grey timecourse regression (MGTR) as is reported in the main text (as "MGTRVar"). The mean grey timecourse variance is always reduced by tICA cleanup, but there is still substantial variability across subjects (likely due to drowsiness and sleeping). The variance of the global signal across subjects in resting state fMRI data is much higher than in task fMRI data (note that the axes extend to 2000 for rfMRI but only to 1000 for the analogous tfMRI figure above). Still, for the majority of subjects (likely the awake ones) the mean grey timecourse variance is less than 100 after cleanup, as in task fMRI.

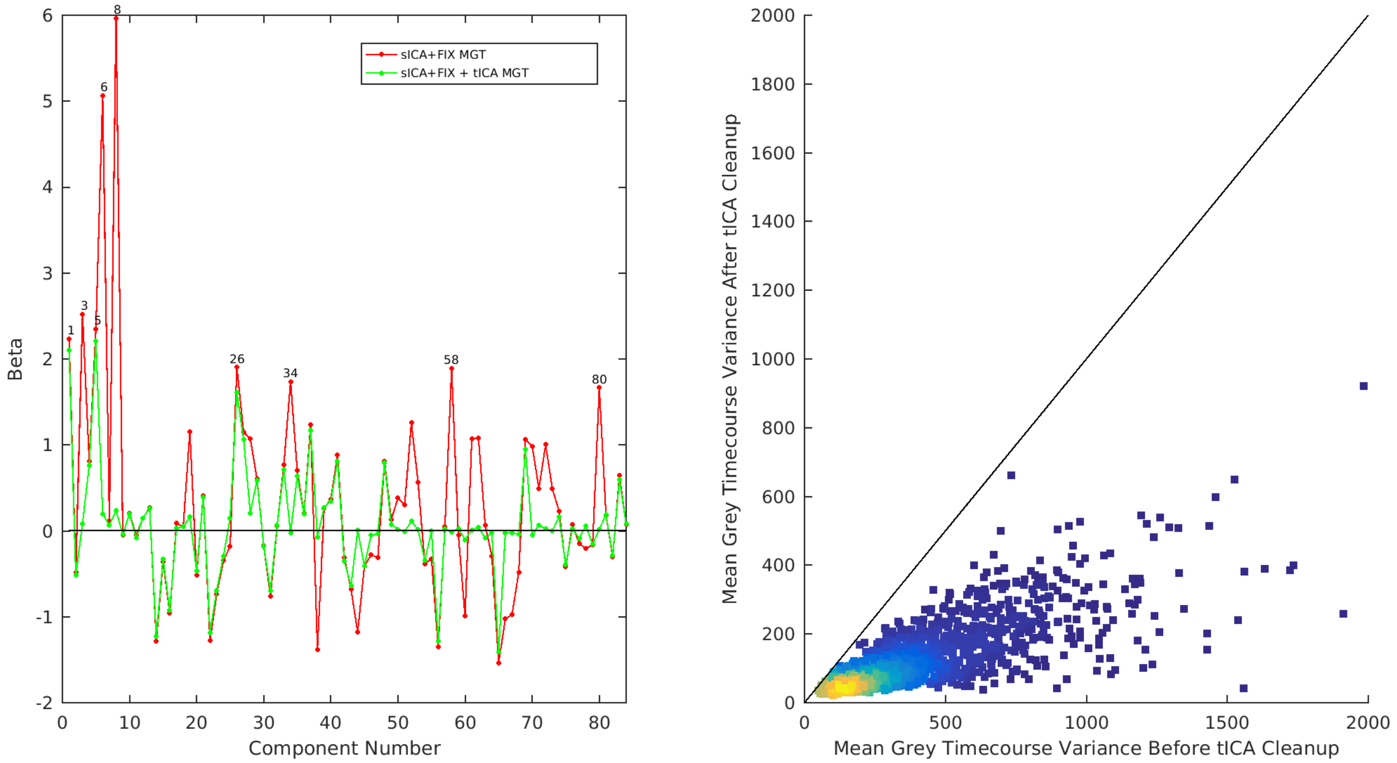

**Supplementary Figure 27** shows the correlation matrix of component amplitudes reordered using hierarchical clustering for the resting state components (using FSLNets with default settings). For each component, there were 1796 (4 runs * 449 subjects) amplitudes, and those amplitudes were correlated across tICA components and clustered. As with the task fMRI data, signal (here, red, cyan, and purple) and noise (green) components tend to cluster together. Additionally, signal components that correlate with sleep component RC1 form a group (purple), while the signal components that do not correlate with RC1 form a separate group (red and cyan). The run-wise variances of components 6 and 8 were summed and the sum entered into the matrix for both components (far left) to ensure that the splitting across RL and LR runs of those two global physiological noise components did not leave them with less correlation to the other components than they would have had if they had not split.

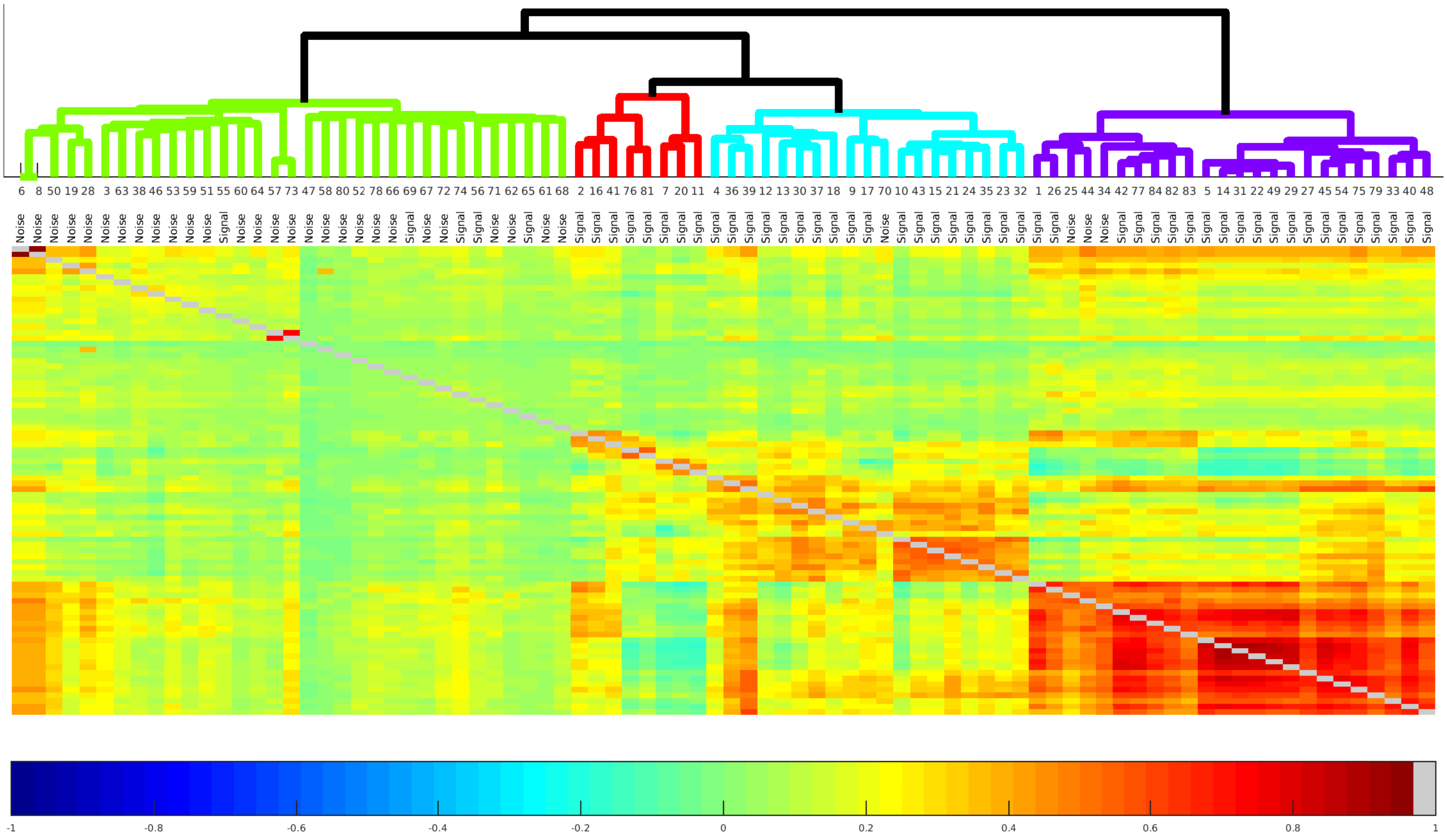

**Supplementary Figure 28** shows the absolute value of the spatial correlations between the grayordinate spatial maps of resting state fMRI (rfMRI), task fMRI (tfMRI), and the task fMRI residuals (tfMRIr) after fitting the task model. Absolute value was used because the sign of components is arbitrary. Correlations are scaled from 0 (black) to 1 (yellow). These data were referenced when assigning component similarities in the TC, TCr, and RC supplementary materials; however, the original sign was preserved in the Supplementary maps. Additionally, components that, when combined, would be similar to another map were also assigned via manual inspection.

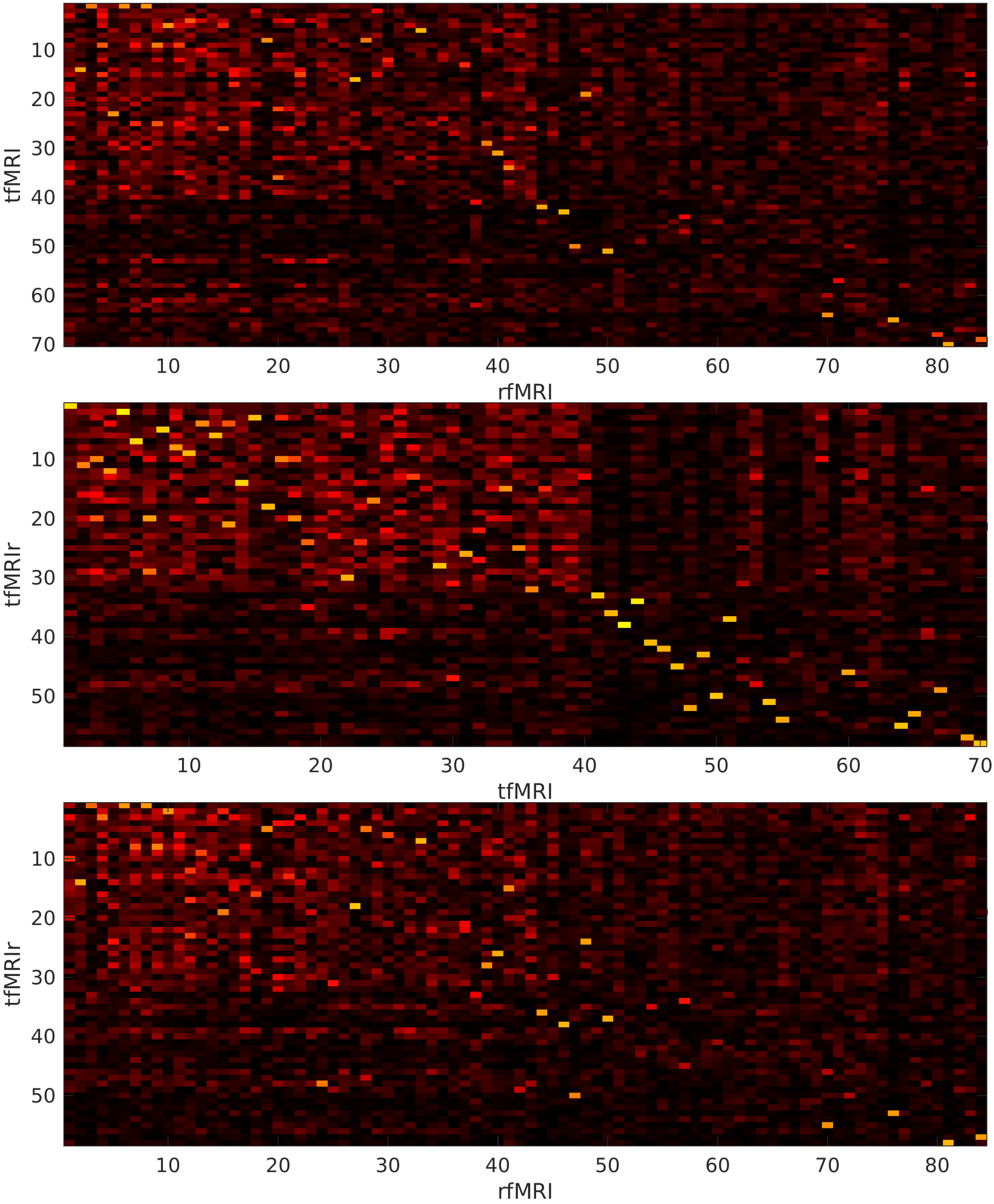


### Supplementary Discussion

*6. Interpretability of Metrics for the Evaluation of fMRI Data Cleanup Methods.*

Defining metrics for evaluating fMRI data cleanup methods is challenging because of the difficulty in assessing ground truth and because of the complex correlations between subject behavior in the scanner (particularly movement and sleep), subject neural activity, subject physiology, and the resulting fluctuations in fMRI signal intensity. For this reason, we made our initial evaluations of tICA cleanup on task fMRI data, where we had an explicit hypothesis about a portion of the true fluctuations of the fMRI signal that are a direct result of known subject behavior in the scanner (performing the task). With resting state fMRI data, we lack an explicit hypothesis about the true neural BOLD fluctuations that were ongoing during the scan, and thus we are limited to characterizing the effects of cleanup and assessing whether these effects are generally desirable or undesirable. In keeping with tenets of HCP-Style neuroimaging analysis (Glasser et al., 2016b), we consider it preferable to use data-driven approaches that selectively remove structured (and unstructured) noise while minimally affecting the properties of the neural signal, even if some of that neural signal is attributable to non-compliant subject behavior (see Main Introduction and Discussion). As a result, the publicly released HCP data avoids non-selective methods of data cleanup such as temporal filtering, frame censoring (scrubbing), and global signal regression (or equivalently regression of WM or CSF timeseries containing signal from locations within 2 voxels (4mm) of grey matter (Power et al., 2018; Power et al., 2017)). Instead, we have aimed here to develop a data driven method for removing global artifacts shown to be present in the released HCP data (Burgess et al., 2016; Power et al., 2017; Siegel et al., 2017) as well as any residual spatially-specific structured noise.

Because we choose to avoid unselective cleanup methods, it is important to carefully consider which data cleanup metrics to use and their underlying assumptions. In particular, we believe that metrics that would be improved by removing both artifactual fluctuations and true neural signal are problematic to interpret, because they have as an implicit underlying null assumption that subjects' true neural signal will be indistinguishable across periods of resting quietly while awake with eyes open, versus periods of eyes closed, movement, or drowsiness/sleep. Further, methods that rely on subject-wise correlations with external measures make this assumption across the entire scanning session, not just during periods of subject non-compliance. Prior studies have shown that these null assumptions are invalid for eyes-open vs. eyes-closed, as there are differences in winner-take-all functional network parcellations in visual cortex between the two states (Laumann et al., 2015). In the present study, we show that these null assumptions are incorrect for subject motion as well, insofar as we find that somatotopically organized sensori-motor temporal ICA components, which are highly similar to task GLM maps in both task fMRI and resting state fMRI data, have higher amplitude both during a motor task and during spontaneous movement as assessed by DVARS Dips in both task and resting state fMRI data. In addition, (Zeng et al., 2014) asserted that subjects who tend to move may have differing functional connectivity than those who do not, even when they are not actively moving. This finding is likely due to many factors including subject-wise correlations between behaviors in the scanner (e.g., movement and sleep) with distinct neural signatures, neural effects of movement extending beyond the immediate movement epoch because of the hemodynamic response function, subject-wise correlations with physiology (e.g., respiratory patterns or heart rate variations), and possibly other causes. Our study also shows that the null assumption of no neural difference between states is incorrect for the neural effects of drowsiness and sleeping behavior, as there are strong components present only at rest that are associated with sleeping behavior as recorded by the research assistants running the MRI scanner. This finding is consistent with prior literature on differences between functional connectivity for different levels of arousal (Laumann et al., 2017; Liu et al., 2017; Tagliazucchi and Laufs, 2014; Wong et al., 2016; Wong et al., 2013; Yeo et al., 2015). Metrics that assume neural equality between periods of movement vs. non-movement or sleep vs. awake will give an advantage to methods such as scrubbing and global signal regression that reduce or remove the true neural differences that exist between those conditions. Put another way, such metrics are one-sided and do not have a positive control that will alert us to when neural signal is being removed, which is in contrast to the task fMRI-based analysis we used here for primary evaluation of data cleanup.

Thus, in this study we did not use any of six metrics that have been used previously in the literature for primary evaluation of data cleanup (Burgess et al., 2016; Ciric et al., 2017; Power et al., 2014; Power et al., 2017; Siegel et al., 2017), however we provide some of them below for historical perspective on prior literature: 1) Statistical tests of differences in network matrices between high and low motion groups or sleeping and not sleeping subjects, as this metric would be improved by removing true neural signal differences related to motion or sleep. 2) "QC-FC" plots in which different methods are compared as to whether or not they reduce or eliminate correlations between some QC measure, such as a measure of subject motion, and functional connectivity (FC), for the same reason as #1 (QC-FC plots are actually even more limited because subject-wise connectivity differences related to neural activity differences during periods when subjects are not moving will still drive correlations with movement). 3) Distance-dependent functional connectivity correlations in which different methods of cleanup are evaluated based on whether they eliminate functional connectivity differences in a distance-dependent fashion. If specific neural activity patterns are more prevalent during periods of sleep, motion, etc., they may have some distance dependence in their connectivity relative to awake and motionless resting periods, particularly if the differences in activity are spatially widespread. 4) Global signal variance explained by movement and physiological parameters, which may correlate with subject behavior in the scanner and consequently also correlate with the global signal variance from neural sources. For example, sleeping may influence both the amount of movement and physiological parameters such as respiration and heart rate, together with causing increased global signal from semi-global neural components—particularly components RC1 and RC5 (see Figure 8 in Main Text). Thus, one might remove all of the artifactual global timecourse variance and still be left with correlations between neural global timecourse variance and the original motion and/or physiological parameters. 5) Correlation of resting state functional connectivity with behavioral variables—these correlations may still be increased by methods that reduce the neural effects of non-compliant behavior in the scanner by reducing overall cross-subject variability. Also, subject behavioral variables measured outside the scanner may still correlate with subject behavior in the scanner (e.g., body weight with motion, or sleep quality with likelihood of falling asleep in the scanner) and thus the neural effects of this behavior. These issues make interpretation of the causal link in such behavioral/connectivity correlations challenging, and make them non-optimal for use as a metric of data cleanup quality. 6) In addition, we avoided metrics that, by virtue of their mathematical definition, intrinsically prefer that the mean grey timeseries be minimized. Although we do not know the true amount of neural global signal (though we believe we are measuring it more accurately in this paper than in previous studies), it is neurobiologically implausible that it is zero, and indeed prior reports have identified correlations between fMRI global signal and electrophysiology (Scholvinck et al., 2010; Wen and Liu, 2016). Hence assuming that it will be zero is unwarranted. Maximizing modularity (Q) will tend to prefer cleanup methods that generate less global or semi-global signal because the resulting modules will be smaller and have fewer between-network connections relative to within-network connections. Q should in general be higher when global signal is lower, other things being equal. Because we do not have a ground truth measure of the amount of global or semi-global neural signal in the data, and because this may also vary according to subjects' within-scanner behavior, we do not know the optimal value of Q.

Because each of the above measures will tend to give an advantage to non-selective methods that minimize the global signal or reduce real differences in neural signal during different patterns of subject behavior, we approached the problem differently (though it is worth noting that we found several other widely used tools very helpful, such as DVARS for highlighting locations of strong signal deviations and greyplots for exploring the spatio-temporal structure of the data). As mentioned above, we focused first on task fMRI data, where we have specific hypotheses about a portion of the neural signal (55% of the overall neural signal and 26% of the global neural signal after sICA+FIX + tICA cleanup was task-related, as defined by the variance removed by regressing out the GLM design matrix). In Main Text Section 2.1.3 we show with the task data that sICA+FIX and tICA improve statistical sensitivity or remove biases in the statistics arising from stimulus correlated motion or physiology (Figure 6, and Supplementary Figures 20, 21, 22, 23), while not removing spatial patterns strongly correlated with the task activation pattern (Supplementary Figure 25). In contrast, MGTR harms task sensitivity (Figure 6) by removing spatial patterns strongly correlated with the task activation pattern (Figure 25), and also drives all of the means of the task fMRI contrast maps to zero (Figure 6), something that is neurobiologically implausible. It might be argued that this is something of a 'straw man' insofar as relatively few studies actually carry out GSR or MGTR on task fMRI data. However, we consider it to be an appropriate comparison, because the issues involved in using or not using GSR are largely the same for task and resting-state fMRI. Additionally, in Main Text Sections 2.1.2 and 2.2.2 (Figures 5 and 10) we show that tICA cleanup is as effective as MGTR at removing banding patterns in spatio-temporal "greyplots" of both task fMRI and resting state fMRI timeseries – banding patterns that have been convincingly linked to respiration (Power et al., 2018; Power et al., 2017). Because we lack a specific hypothesis about the temporal pattern of neural signal in resting-state fMRI data, there is no objective measure upon which to compute statistical sensitivity and to compare different cleanup approaches. We can nonetheless characterize the *effects* of differing cleanup approaches on functional connectivity maps. Like task fMRI data, however, a zero-mean resting-state correlation matrix is highly improbable as a ground truth. Indeed, in Main Text Section 2.2.3 (Figures 11 and 13) we show that while sICA+FIX data without temporal ICA cleanup has a positive bias, MGTR induces network-specific negative biases in the functional connectivity maps, just like the network-specific bias in task fMRI data in Supplementary Figure 25 above, a finding that was predicted in the literature (Gotts et al., 2013; Saad et al., 2012). Unlike the relatively uniform cleanup from tICA, the MGTR bias in resting state fMRI data affects non-cognitive and task positive regions much more than other cognitive regions and task negative regions. It is thus not surprising that while both approaches increase gradient magnitude, MGTR also shifts gradients by virtue of spatially non-uniform removal of neural signal. Indeed, it was concerns over this shift in functional connectivity gradients that led MGTR not to be used in a prior neuroanatomical study (Glasser et al., 2016a), and led the authors to explore the global signal in considerable detail.

That said, having outlined these interpretational issues, there is still value generating some of the metrics (Burgess et al., 2016; Ciric et al., 2017; Power et al., 2014; Power et al., 2017; Siegel et al., 2017) on our data to provide a historical perspective on the prior literature so long as we do not attempt to interpret them as telling us that one method or another is "better," using the problematic null assumptions discussed above. In the following four figures we illustrate (1) the difference in correlation between DVARS Dips timepoints and non-DVARS Dips timepoints and (2) QC-FC correlations between subject-wise functional connectivity ('FC') of all timepoints and the number of DVARS Dips as the 'QC' measure. All correlations were computed using Pearson correlations, although we found highly similar results when using non-parametric Spearman correlations for the QC-FC plots. These analyses were performed on each subject's concatenated task or resting state fMRI data after minimal preprocessing (MPP), minimal preprocessing + mean grey timecourse regression (MPP + MGTR), sICA+FIX, sICA+FIX + MGTR, and sICA+FIX + tICA. Each panel of each figure is a scatterplot of correlation vs Euclidean distance between parcels in millimeters (see combined figure legend below for details). In addition we computed the slopes of linear regression lines, the medians of the distributions, and the widths (standard deviations) of the distributions, which are tabulated in the sixth panel. Because all four figures show the same trends, we will describe them together.

Importantly, though we use a new, more robust QC metric, DVARS Dips (Figures 1, 2 and 3) for fMRI data with high temporal and spatial resolution, the previously described dependence of functional connectivity on distance in the presence of motion is replicated in the MPP and MPP + MGTR results for both task fMRI and resting state fMRI. The slopes of the regression lines are more different from zero for MGTR-based approaches than non-MGTR-based approaches, suggesting that MGTR universally increases the distance dependence of these measures while sICA+FIX and sICA+FIX + tICA reduce distance dependence (consistent with what was found with GSR and ICA-AROMA in (Ciric et al., 2017). Indeed, it appears that the use of GSR together with a lack of wide availability of automated spatial ICA-based cleanup methods was largely or completely responsible for the now thought to be erroneous findings of distance dependent connectivity changes with age (Dosenbach et al., 2010; Fair et al., 2008; Fair et al., 2009; Fair et al., 2007) that led to an increased focus on motion and other structured noise in fMRI data (Power et al., 2012; Satterthwaite et al., 2012; Van Dijk et al., 2012). Further, from the literature it appears that the spatial ICA-based cleanup methods (Griffanti et al., 2014; Salimi-Khorshidi et al., 2014) (Pruim et al., 2015a; Pruim et al., 2015b) (Kundu et al., 2012) produce similar reductions in distance dependent connectivity as the "scrubbing" technique (Power et al., 2012; Power et al., 2014) though spatial ICA-based cleanup removes such effects selectively without requiring complete removal of the problematic frames. Importantly, a major limitation of scrubbing relative to ICA-based cleanup is that scrubbing is binary decision dependent on the chosen QC metric thresholds, whereas ICA will clean the entire timeseries of artifacts, including timepoints that would be below a scrubbing threshold but will undoubtedly include some spatially-specific structured noise from motion, physiology, or MR physics sources. ICA-based cleanup produces weighted "softer" scrubbing-like changes (i.e., DVARS Dips) only when required to clean the data.

In the figures below, the two analysis variants involving MGTR generally have the median closest to zero which is not surprising given that MGTR centers the functional connectivity around zero; however, sICA+FIX and sICA+FIX + tICA also generally make the median closer to zero (relative to MPP and sICA+FIX respectively). The distributions of the data are generally tighter for the non-MGTR methods than the MGTR methods, indicating that MGTR-based methods either have greater difference in correlation at the extremes between DVARS Dips and non-Dip timepoints or stronger correlation with number of DVARS Dips at the extremes. In agreement with the overall higher percentage of frames with DVARS Dips in the task fMRI data (1.7%) compared to the resting state fMRI data (1.3%), the motion-related effects are stronger in the task fMRI data than the resting state fMRI data in the HCP subject population of young adults (i.e., slopes farther from zero and wide distributions). It is worth noting that we do not think the decrease in the median QC-FC value with sICA+FIX + tICA (relative to sICA+FIX) is actually related to motion per se, but rather to removing global physiological noise arising from respiration that is subject-wise correlated with motion, as motion itself is unlikely to produce greymatter specific global signal intensity changes of the type removed by tICA (Glasser et al., 2016b). Indeed, in a recent paper using multi-echo acquisitions (which allow separation of BOLD T2* decay dependent and non-BOLD S0 signal intensity changes) global respiratory effects of the kind removed by tICA cleanup were shown to occur via the BOLD T2* decay dependent mechanism, in contrast to the spatially specific effects of head motion that are removed by sICA+FIX, which were shown to occur via modulations of non-BOLD S0 intensity fluctuations (Power et al., 2018). Similarly, we believe the greater offset between the median QC-FC and zero in the resting state data vs. the task fMRI data after sICA+FIX + tICA is due to drowsy/sleeping subjects in the resting state fMRI data that do not occur in the task fMRI data, a behavior that may also be subject-wise correlated with motion. Thus, any difference in the offset from zero (captured by the median value) between sICA+FIX + tICA and the MGTR variants is reflective of the impact of global neural signal, and that the sICA+FIX + tICA plots represent the best available current estimate of what these sorts of metrics should look like when global neural signal is preserved.

**Combined legend for Figures 29, 30, 31, 32.** Either the correlation difference between DVARS Dips timepoints and non-Dips timepoints (Figures 29 and 30) or "QC-FC" correlation between subject-wise connectivity of all timepoints and the number of DVARS Dips (Figures 31 and 32) is displayed for task fMRI (Figures 29 and 31) or resting state fMRI (30 and 32). Correlations were computed between all parcels in the HCP-MMP1.0 multi-modal cortical parcellation (such that each 'edge' between parcels constitutes one data point). Five different analyses are shown ranging from minimal preprocessed data (MPP), MPP + mean grey timecourse regression (MPP + MGTR), sICA+FIX, sICA+FIX + MGTR, and sICA+FIX + tICA. In each case the plot is a scatter plot across all 'edges' with correlation difference or QC-FC correlation on the y-axis and Euclidean distance between parcels in millimeters on the x-axis. Distances were computed between the average midthickness coordinates of each of the HCP's multi-modal parcellation's cortical areas. Hotter colors indicate a higher density of data points. A straight black line marks 0, a wiggling black line represents a fitted curve (specifically, 1mm width mean correlation values), and a red line marks the linear regression fit of the entire point cloud. The sixth panel shows a table containing the slope of the regression line, the median of the point cloud along the y axis (collapsed across all distances), and the width of the point cloud (measured as the standard deviation (STDev) collapsed across all distances).

**Supplementary Figure 29** shows the DVARS Dips-non Dips difference in correlation for task fMRI data.

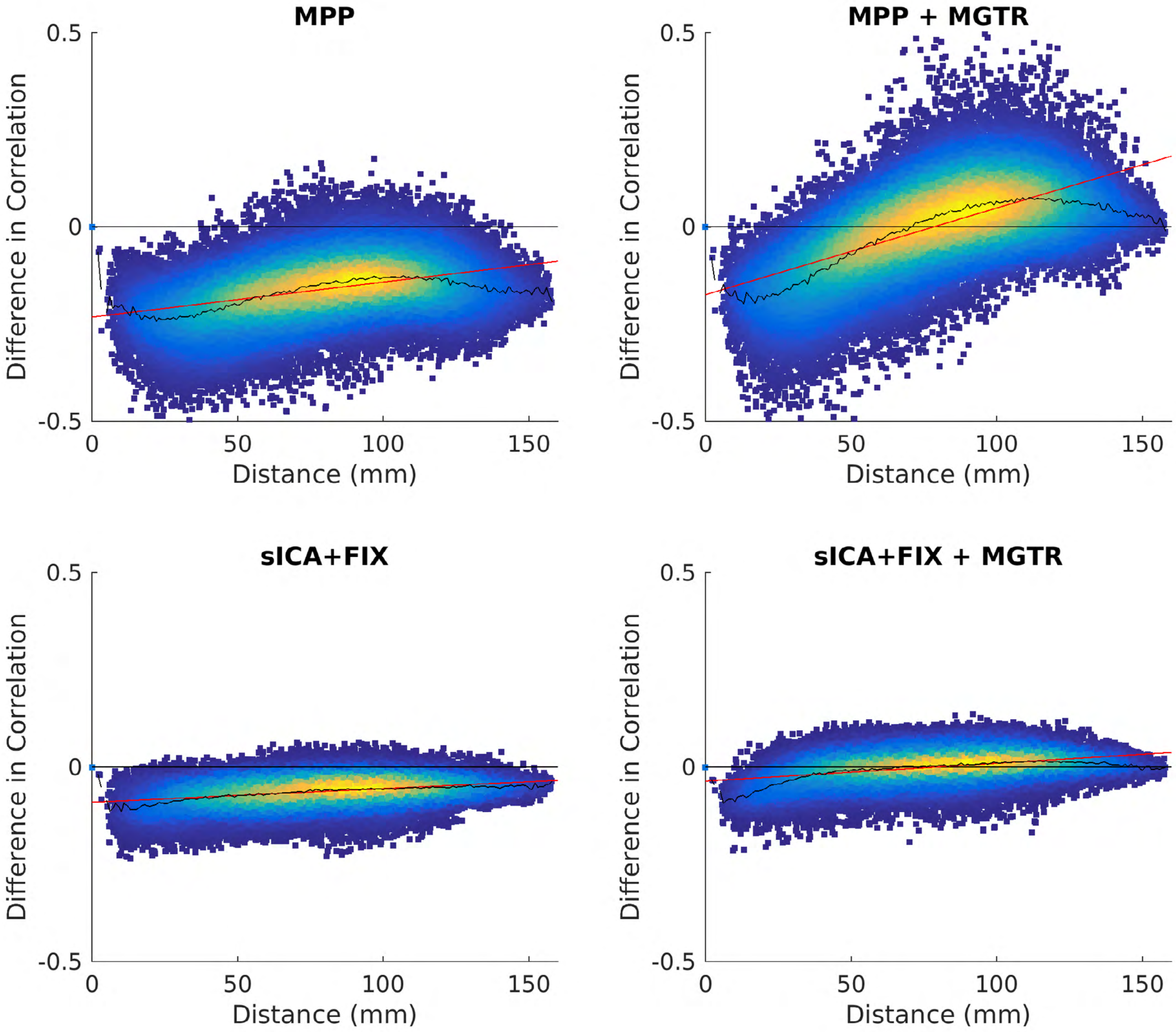


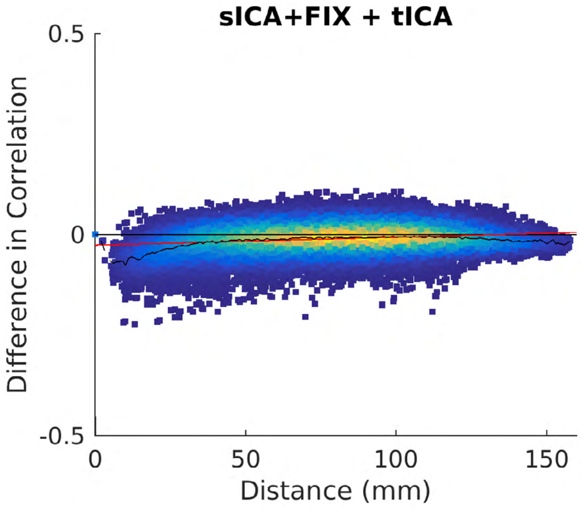


| Analysis | Slope | Median | STDev |
|---|---|---|---|
| **MPP** | 9.00e-04 | 1.59e-01 | 7.38e-02 |
| **MPP + MGTR** | 2.23e-03 | -9.09e-03 | 1.09e-01 |
| **sICA+FIX** | 3.51e-04 | 6.03e-02 | 3.39e-02 |
| **sICA+FIX + MGTR** | 4.59e-04 | -3.21e-03 | 3.63e-02 |
| **sICA+FIX + tICA** | 1.97e-04 | 9.49e-03 | 2.97e-02 |

**Supplementary Figure 30** shows the DVARS Dips-non Dips difference in correlation for resting state fMRI data.

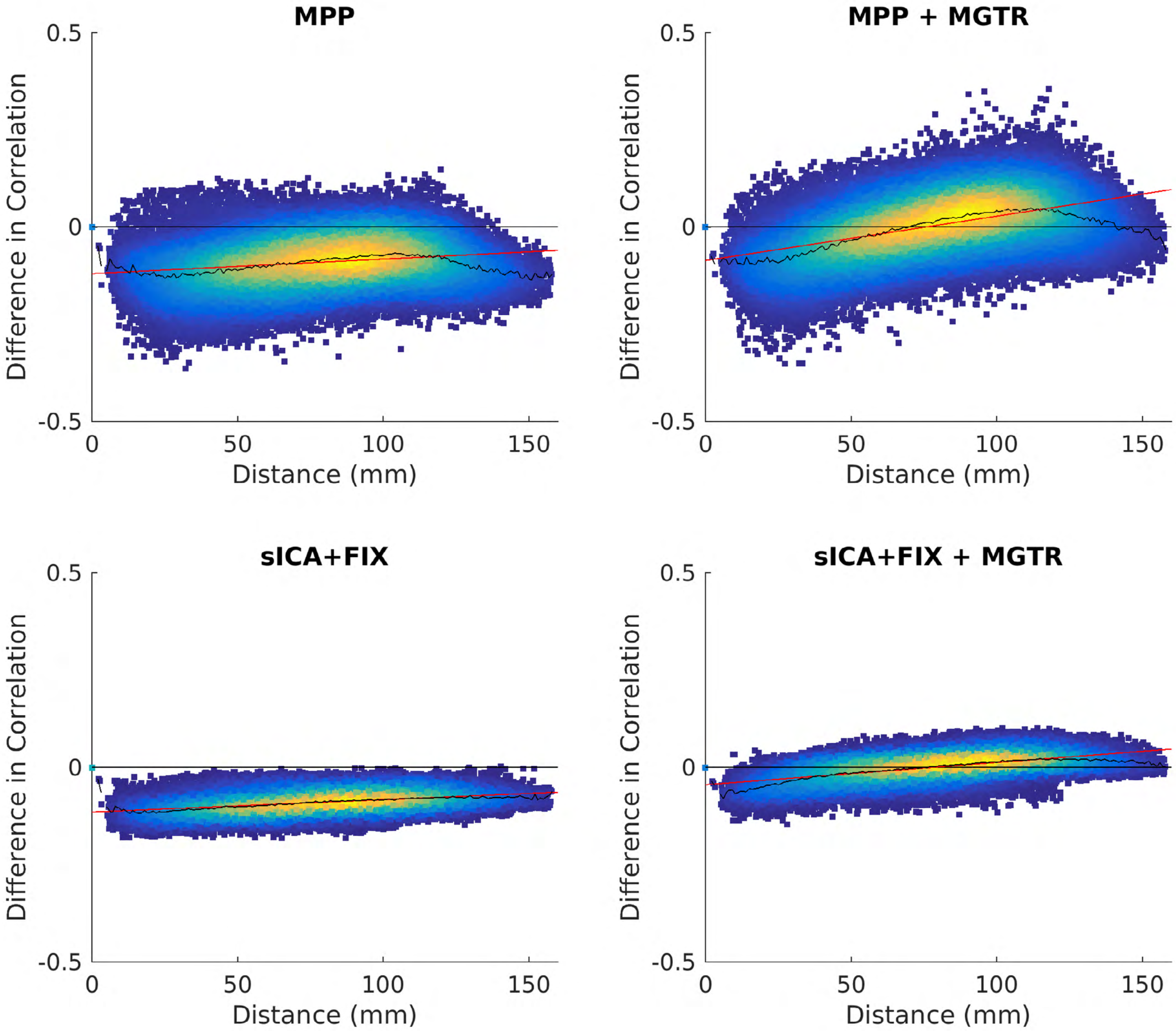


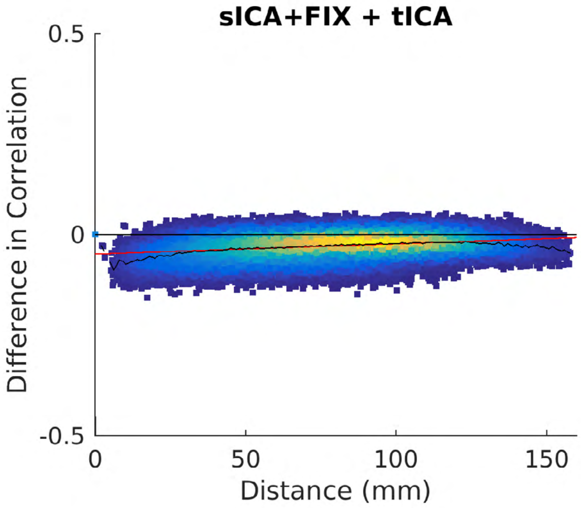


| Analysis | Slope | Median | STDev |
|---|---|---|---|
| **MPP** | 3.74e-04 | 9.06e-02 | 6.11e-02 |
| **MPP + MGTR** | 1.14e-03 | -5.39e-03 | 7.43e-02 |
| **sICA+FIX** | 3.18e-04 | 9.08e-02 | 2.68e-02 |
| **sICA+FIX + MGTR** | 5.72e-04 | -3.36e-03 | 3.20e-02 |
| **sICA+FIX + tICA** | 2.55e-04 | 2.51e-02 | 2.83e-02 |

**Supplementary Figure 31** shows the DVARS Dips QC-FC plots for task fMRI data.

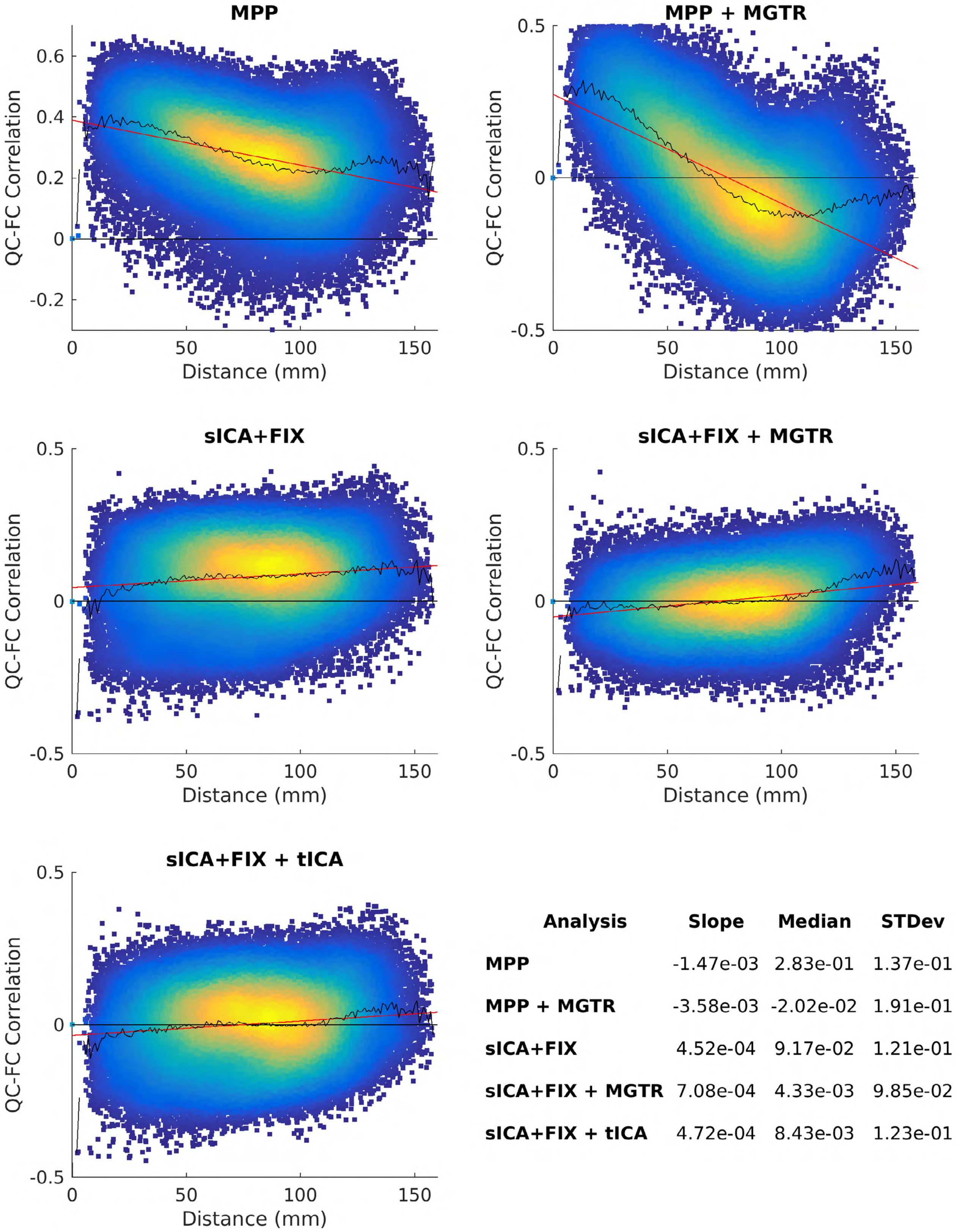


| Analysis | Slope | Median | STDev |
|---|---|---|---|
| **MPP** | -1.47e-03 | 2.83e-01 | 1.37e-01 |
| **MPP + MGTR** | -3.58e-03 | -2.02e-02 | 1.91e-01 |
| **sICA+FIX** | 4.52e-04 | 9.17e-02 | 1.21e-01 |
| **sICA+FIX + MGTR** | 7.08e-04 | 4.33e-03 | 9.85e-02 |
| **sICA+FIX + tICA** | 4.72e-04 | 8.43e-03 | 1.23e-01 |

**Supplementary Figure 32** shows the DVARS Dips QC-FC plots for resting state fMRI data.

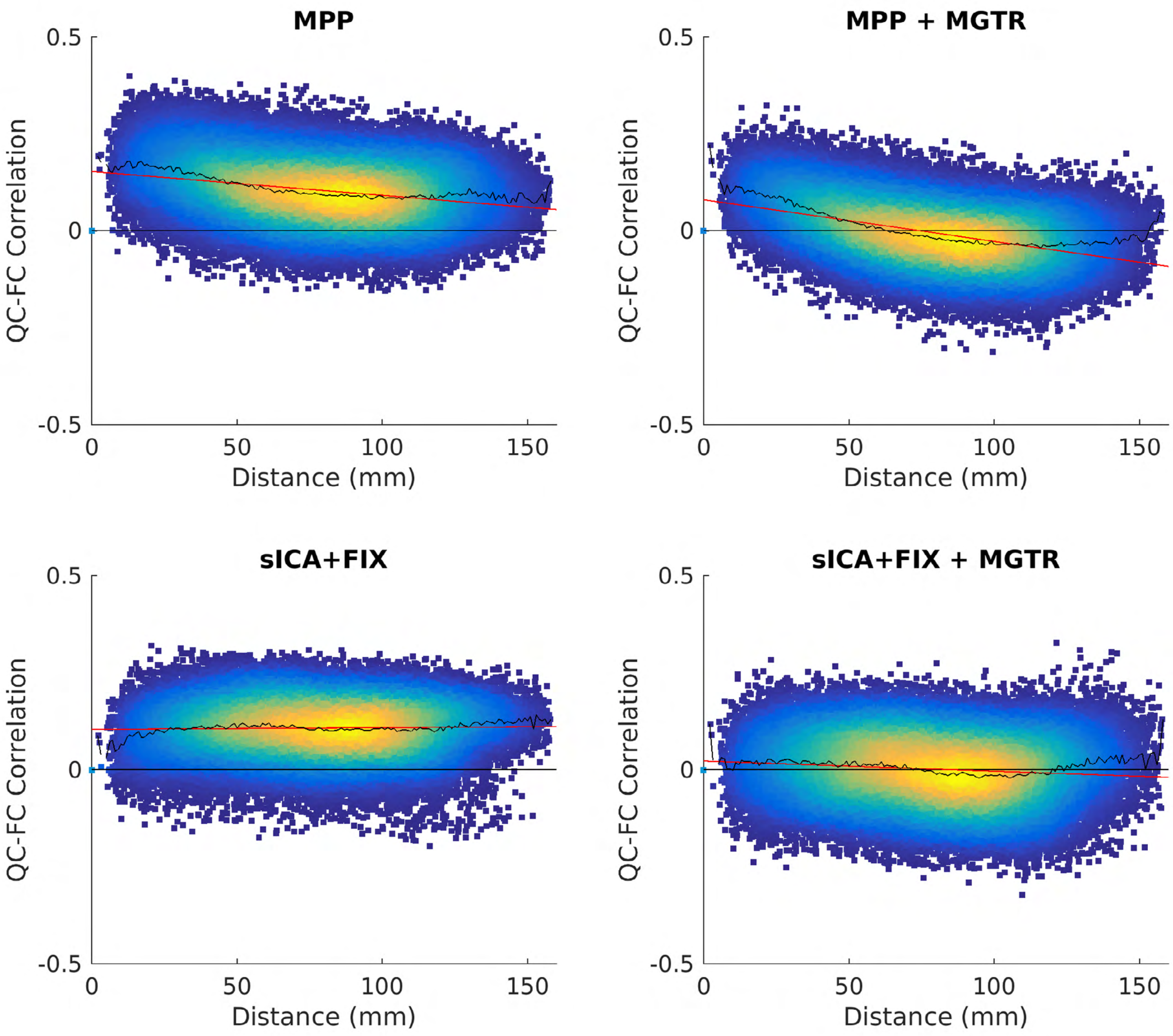


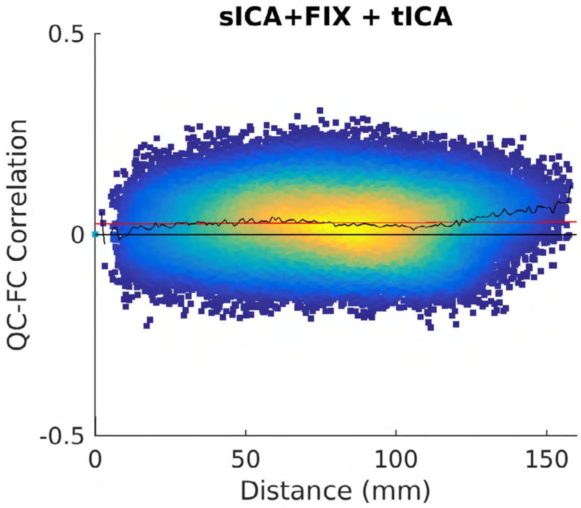


| Analysis | Slope | Median | STDev |
|---|---|---|---|
| **MPP** | -6.15e-04 | 1.04e-01 | 7.57e-02 |
| **MPP + MGTR** | -1.07e-03 | -5.98e-03 | 7.60e-02 |
| **sICA+FIX** | 4.55e-05 | 1.10e-01 | 6.65e-02 |
| **sICA+FIX + MGTR** | -2.62e-04 | 2.59e-03 | 8.23e-02 |
| **sICA+FIX + tICA** | 3.70e-05 | 2.97e-02 | 7.84e-02 |

**Supplementary References**


Beckmann, C.F., M. DeLuca, J.T. Devlin, and S.M. Smith. 2005. Investigations into resting-state connectivity using independent component analysis. *Philosophical transactions of the Royal Society of London. Series B, Biological sciences*. 360:1001-1013.

Beckmann, C.F., and S.M. Smith. 2004. Probabilistic independent component analysis for functional magnetic resonance imaging. *IEEE transactions on medical imaging*. 23:137-152.

Bright, M.G., M. Bianciardi, J.A. de Zwart, K. Murphy, and J.H. Duyn. 2014. Early anti-correlated BOLD signal changes of physiologic origin. *NeuroImage*. 87:287-296.

Burgess, G.C., S. Kandala, D. Nolan, T.O. Laumann, J.D. Power, B. Adeyemo, M.P. Harms, S.E. Petersen, and D.M. Barch. 2016. Evaluation of Denoising Strategies to Address Motion-Correlated Artifacts in Resting-State Functional Magnetic Resonance Imaging Data from the Human Connectome Project. *Brain connectivity*. 6:669-680.

Chen, L., A. Beckett, A. Verma, and D.A. Feinberg. 2015. Dynamics of respiratory and cardiac CSF motion revealed with real-time simultaneous multi-slice EPI velocity phase contrast imaging. *NeuroImage*. 122:281-287.

Ciric, R., D.H. Wolf, J.D. Power, D.R. Roalf, G.L. Baum, K. Ruparel, R.T. Shinohara, M.A. Elliott, S.B. Eickhoff, C. Davatzikos, R.C. Gur, R.E. Gur, D.S. Bassett, and T.D. Satterthwaite. 2017. Benchmarking of participant-level confound regression strategies for the control of motion artifact in studies of functional connectivity. *NeuroImage*. 154:174-187.

Dosenbach, N.U., B. Nardos, A.L. Cohen, D.A. Fair, J.D. Power, J.A. Church, S.M. Nelson, G.S. Wig, A.C. Vogel, C.N. Lessov-Schlaggar, K.A. Barnes, J.W. Dubis, E. Feczko, R.S. Coalson, J.R. Pruett, Jr., D.M. Barch, S.E. Petersen, and B.L. Schlaggar. 2010. Prediction of individual brain maturity using fMRI. *Science*. 329:1358-1361.

Fair, D.A., A.L. Cohen, N.U. Dosenbach, J.A. Church, F.M. Miezin, D.M. Barch, M.E. Raichle, S.E. Petersen, and B.L. Schlaggar. 2008. The maturing architecture of the brain's default network. *Proceedings of the National Academy of Sciences of the United States of America*. 105:4028-4032.

Fair, D.A., A.L. Cohen, J.D. Power, N.U. Dosenbach, J.A. Church, F.M. Miezin, B.L. Schlaggar, and S.E. Petersen. 2009. Functional brain networks develop from a "local to distributed" organization. *PLoS computational biology*. 5:e1000381.

Fair, D.A., N.U. Dosenbach, J.A. Church, A.L. Cohen, S. Brahmbhatt, F.M. Miezin, D.M. Barch, M.E. Raichle, S.E. Petersen, and B.L. Schlaggar. 2007. Development of distinct control networks through segregation and integration. *Proceedings of the National Academy of Sciences of the United States of America*. 104:13507-13512.

Friston, K.J., S. Williams, R. Howard, R.S. Frackowiak, and R. Turner. 1996. Movement-related effects in fMRI time-series. *Magnetic resonance in medicine*. 35:346-355.

Glasser, M.F., T.S. Coalson, E.C. Robinson, C.D. Hacker, J. Harwell, E. Yacoub, K. Ugurbil, J. Andersson, C.F. Beckmann, M. Jenkinson, S.M. Smith, and D.C. Van Essen. 2016a. A multi-modal parcellation of human cerebral cortex. *Nature*. 536:171-178.

Glasser, M.F., S.M. Smith, D.S. Marcus, J.L. Andersson, E.J. Auerbach, T.E. Behrens, T.S. Coalson, M.P. Harms, M. Jenkinson, S. Moeller, E.C. Robinson, S.N. Sotiropoulos, J. Xu, E. Yacoub, K. Ugurbil, and D.C. Van Essen. 2016b. The Human Connectome Project's neuroimaging approach. *Nature neuroscience*. 19:1175-1187.

Gotts, S.J., Z.S. Saad, H.J. Jo, G.L. Wallace, R.W. Cox, and A. Martin. 2013. The perils of global signal regression for group comparisons: a case study of Autism Spectrum Disorders. *Frontiers in human neuroscience*. 7:356.

Griffanti, L., G. Salimi-Khorshidi, C.F. Beckmann, E.J. Auerbach, G. Douaud, C.E. Sexton, E. Zsoldos, K.P. Ebmeier, N. Filippini, C.E. Mackay, S. Moeller, J. Xu, E. Yacoub, G. Baselli, K. Ugurbil, K.L. Miller, and S.M. Smith. 2014. ICA-based artefact removal and accelerated fMRI acquisition for improved resting state network imaging. *NeuroImage*. 95:232-247.

Hodgson, K., R. Poldrack, J.E. Curran, E.E. Knowles, S. Mathias, H.H. Goring, N. Yao, R.L. Olvera, P.T. Fox, L. Almasy, and R. Duggirala. 2016. Shared genetic factors influence head motion during MRI and body mass index. . *Cereb Cortex*. 27:5539-5546.

Jenkinson, M. 1999. Measuring Transformation Error by RMS Deviation. *fMRIB technical report TR99MJ1*.

Jin, T., and S.G. Kim. 2010. Change of the cerebrospinal fluid volume during brain activation investigated by T(1rho)-weighted fMRI. *NeuroImage*. 51:1378-1383.

Kundu, P., S.J. Inati, J.W. Evans, W.M. Luh, and P.A. Bandettini. 2012. Differentiating BOLD and non-BOLD signals in fMRI time series using multi-echo EPI. *NeuroImage*. 60:1759-1770.

Laumann, T.O., E.M. Gordon, B. Adeyemo, A.Z. Snyder, S.J. Joo, M.Y. Chen, A.W. Gilmore, K.B. McDermott, S.M. Nelson, N.U. Dosenbach, B.L. Schlaggar, J.A. Mumford, R.A. Poldrack, and S.E. Petersen. 2015. Functional System and Areal Organization of a Highly Sampled Individual Human Brain. *Neuron*. 87:657-670.

Laumann, T.O., A.Z. Snyder, A. Mitra, E.M. Gordon, C. Gratton, B. Adeyemo, A.W. Gilmore, S.M. Nelson, J.J. Berg, D.J. Greene, J.E. McCarthy, E. Tagliazucchi, H. Laufs, B.L. Schlaggar, N.U.F. Dosenbach, and S.E. Petersen. 2017. On the Stability of BOLD fMRI Correlations. *Cereb Cortex*. 27:4719-4732.

Liu, T.T., A. Nalci, and M. Falahpour. 2017. The global signal in fMRI: Nuisance or Information? *NeuroImage*. 150:213-229.

Power, J.D. 2017. A simple but useful way to assess fMRI scan qualities. *NeuroImage*. 154:150-158.

Power, J.D., K.A. Barnes, A.Z. Snyder, B.L. Schlaggar, and S.E. Petersen. 2012. Spurious but systematic correlations in functional connectivity MRI networks arise from subject motion. *NeuroImage*. 59:2142-2154.

Power, J.D., A. Mitra, T.O. Laumann, A.Z. Snyder, B.L. Schlaggar, and S.E. Petersen. 2014. Methods to detect, characterize, and remove motion artifact in resting state fMRI. *NeuroImage*. 84:320-341.

Power, J.D., M. Plitt, S.J. Gotts, P. Kundu, V. Voon, P.A. Bandettini, and A. Martin. 2018. Ridding fMRI data of motion-related influences: Removal of signals with distinct spatial and physical bases in multiecho data. *Proceedings of the National Academy of Sciences of the United States of America*. 115:E2105-E2114.

Power, J.D., M. Plitt, T.O. Laumann, and A. Martin. 2017. Sources and implications of whole-brain fMRI signals in humans. *NeuroImage*. 146:609-625.

Pruim, R.H., M. Mennes, J.K. Buitelaar, and C.F. Beckmann. 2015a. Evaluation of ICA-AROMA and alternative strategies for motion artifact removal in resting state fMRI. *NeuroImage*. 112:278-287.

Pruim, R.H., M. Mennes, D. van Rooij, A. Llera, J.K. Buitelaar, and C.F. Beckmann. 2015b. ICA-AROMA: A robust ICA-based strategy for removing motion artifacts from fMRI data. *NeuroImage*. 112:267-277.

Raj, D., A.W. Anderson, and J.C. Gore. 2001. Respiratory effects in human functional magnetic resonance imaging due to bulk susceptibility changes. *Physics in medicine and biology*. 46:3331-3340.

Saad, Z.S., S.J. Gotts, K. Murphy, G. Chen, H.J. Jo, A. Martin, and R.W. Cox. 2012. Trouble at rest: how correlation patterns and group differences become distorted after global signal regression. *Brain connectivity*. 2:25-32.

Salimi-Khorshidi, G., G. Douaud, C.F. Beckmann, M.F. Glasser, L. Griffanti, and S.M. Smith. 2014. Automatic denoising of functional MRI data: combining independent component analysis and hierarchical fusion of classifiers. *NeuroImage*. 90:449-468.

Satterthwaite, T.D., D.H. Wolf, J. Loughead, K. Ruparel, M.A. Elliott, H. Hakonarson, R.C. Gur, and R.E. Gur. 2012. Impact of in-scanner head motion on multiple measures of functional connectivity: relevance for studies of neurodevelopment in youth. *NeuroImage*. 60:623-632.

Scholvinck, M.L., A. Maier, F.Q. Ye, J.H. Duyn, and D.A. Leopold. 2010. Neural basis of global resting-state fMRI activity. *Proceedings of the National Academy of Sciences of the United States of America*. 107:10238-10243.

Siegel, J.S., A. Mitra, T.O. Laumann, B.A. Seitzman, M. Raichle, M. Corbetta, and A.Z. Snyder. 2017. Data Quality Influences Observed Links Between Functional Connectivity and Behavior. *Cereb Cortex*. 27:4492-4502.

Smith, S.M., C.F. Beckmann, J. Andersson, E.J. Auerbach, J. Bijsterbosch, G. Douaud, E. Duff, D.A. Feinberg, L. Griffanti, M.P. Harms, M. Kelly, T. Laumann, K.L. Miller, S. Moeller, S. Petersen, J. Power, G. Salimi-Khorshidi, A.Z. Snyder, A.T. Vu, M.W. Woolrich, J. Xu, E. Yacoub, K. Ugurbil, D.C. Van Essen, and M.F. Glasser. 2013. Resting-state fMRI in the Human Connectome Project. *NeuroImage*. 80:144-168.

Smith, S.M., K.L. Miller, S. Moeller, J. Xu, E.J. Auerbach, M.W. Woolrich, C.F. Beckmann, M. Jenkinson, J. Andersson, M.F. Glasser, D.C. Van Essen, D.A. Feinberg, E.S. Yacoub, and K. Ugurbil. 2012. Temporally-independent functional modes of spontaneous brain activity. *Proceedings of the National Academy of Sciences of the United States of America*. 109:3131-3136.

Smyser, C.D., T.E. Inder, J.S. Shimony, J.E. Hill, A.J. Degnan, A.Z. Snyder, and J.J. Neil. 2010. Longitudinal analysis of neural network development in preterm infants. *Cereb Cortex*. 20:2852-2862.

Tagliazucchi, E., and H. Laufs. 2014. Decoding wakefulness levels from typical fMRI resting-state data reveals reliable drifts between wakefulness and sleep. *Neuron*. 82:695-708.

Thomas, B.P., P. Liu, S. Aslan, K.S. King, M.J. van Osch, and H. Lu. 2013. Physiologic underpinnings of negative BOLD cerebrovascular reactivity in brain ventricles. *NeuroImage*. 83:505-512.

Van Dijk, K.R., M.R. Sabuncu, and R.L. Buckner. 2012. The influence of head motion on intrinsic functional connectivity MRI. *NeuroImage*. 59:431-438.

Wen, H., and Z. Liu. 2016. Broadband Electrophysiological Dynamics Contribute to Global Resting-State fMRI Signal. *The Journal of neuroscience : the official journal of the Society for Neuroscience*. 36:6030-6040.

Wong, C.W., P.N. DeYoung, and T.T. Liu. 2016. Differences in the resting-state fMRI global signal amplitude between the eyes open and eyes closed states are related to changes in EEG vigilance. *NeuroImage*. 124:24-31.

Wong, C.W., V. Olafsson, O. Tal, and T.T. Liu. 2013. The amplitude of the resting-state fMRI global signal is related to EEG vigilance measures. *NeuroImage*. 83:983-990.

Yeo, B.T., J. Tandi, and M.W. Chee. 2015. Functional connectivity during rested wakefulness predicts vulnerability to sleep deprivation. *NeuroImage*. 111:147-158.

Yildiz, S., S. Thyagaraj, N. Jin, X. Zhong, S. Heidari Pahlavian, B.A. Martin, F. Loth, J. Oshinski, and K.G. Sabra. 2017. Quantifying the influence of respiration and cardiac pulsations on cerebrospinal fluid dynamics using real-time phase-contrast MRI. *Journal of magnetic resonance imaging : JMRI*. 46:431-439.

Zeng, L.L., D. Wang, M.D. Fox, M. Sabuncu, D. Hu, M. Ge, R.L. Buckner, and H. Liu. 2014. Neurobiological basis of head motion in brain imaging. *Proceedings of the National Academy of Sciences of the United States of America*. 111:6058-6062.
