## Supplementary Materials for "Using Temporal ICA to Selectively Remove Global Noise While Preserving Global Signal in Functional MRI Data"

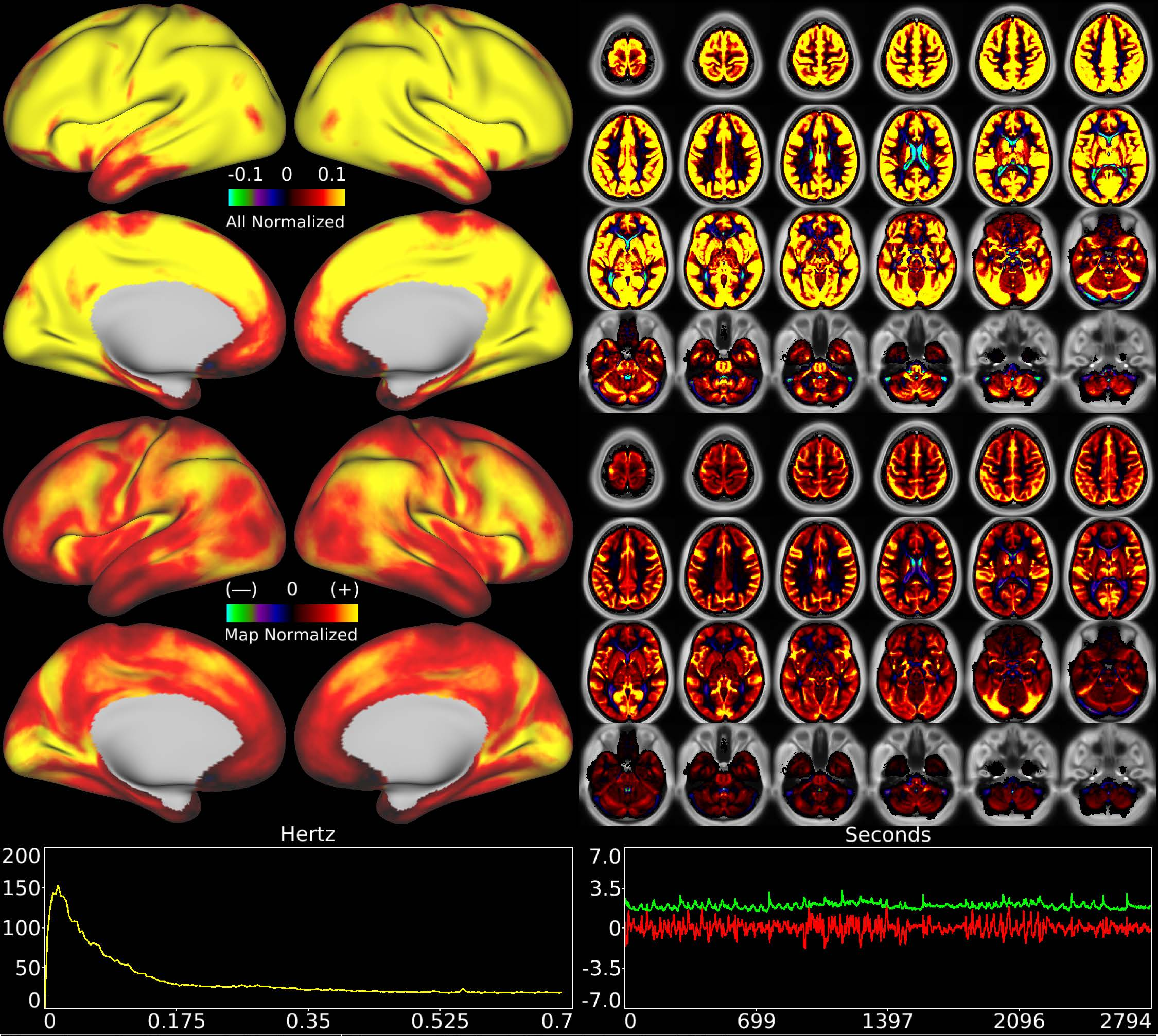


| | | |
|---|---|---|
| Number & Class: 1 Noise | Name: Global Physiological Noise | |
| RVT Correlated: Yes | DVARS Dip Associated: Yes | Cross-Subject Variable: Yes |
| Single Subject: No | % Variance Explained: 6.67 | Globality Index: 5.82 |
| Rest Component: 3+6+8 | Taskr Component: 1 | Task Modulated: No |
| Rationale: Globally positive component matching hypothesized pattern for physiological artifact; does not clearly follow known areal boundaries | | |

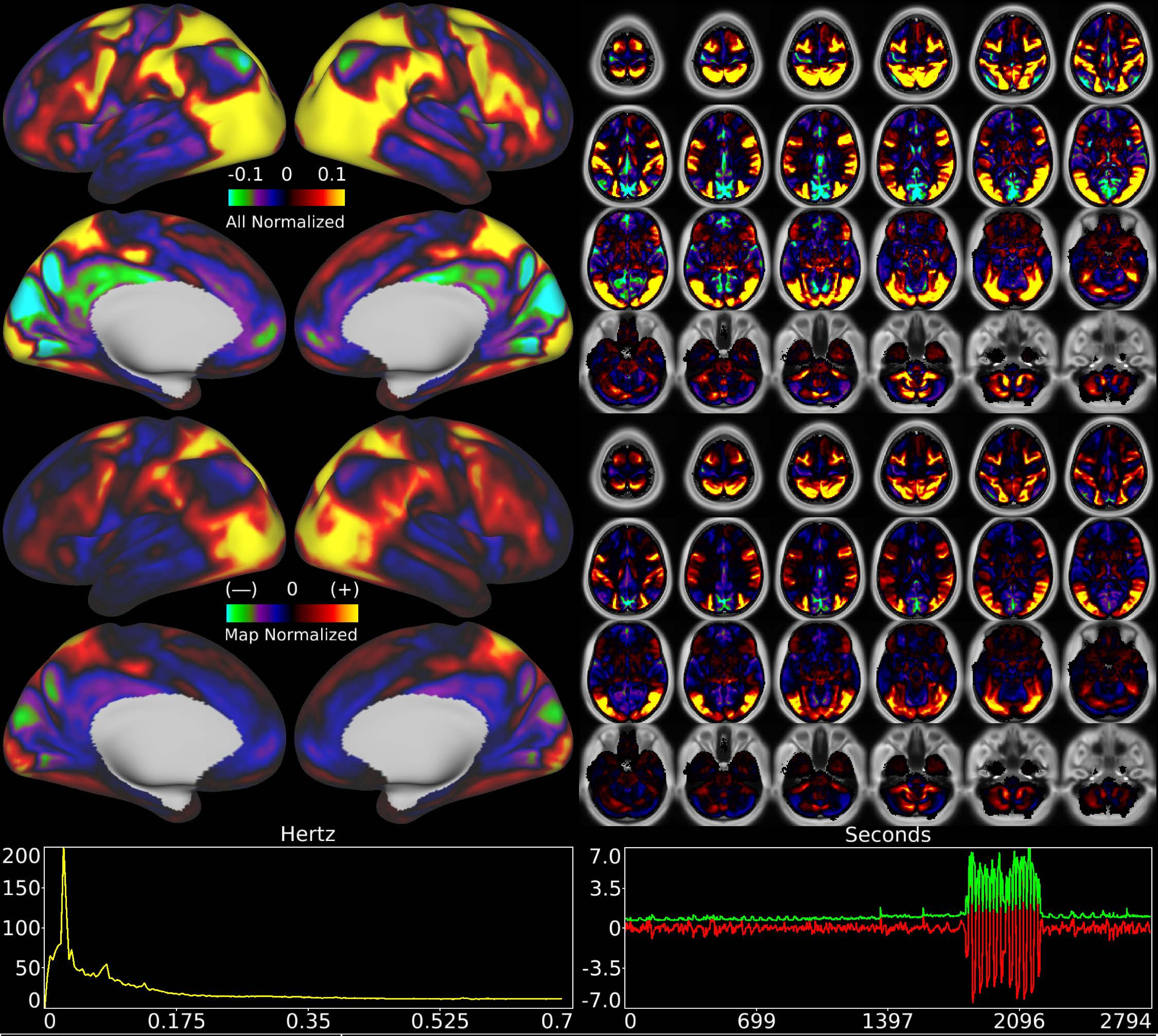


| | | |
|---|---|---|
| Number & Class: 2 Signal | Name: Social Task Main | |
| RVT Correlated: No | DVARS Dip Associated: No | Cross-Subject Variable: No |
| Single Subject: No | % Variance Explained: 4.78 | Globality Index: 0.25 |
| Rest Component: No | Taskr Component: 11 | Task Modulated: Social |

Rationale: Spatial map includes positive and negative patches that are correlated with a specific task design

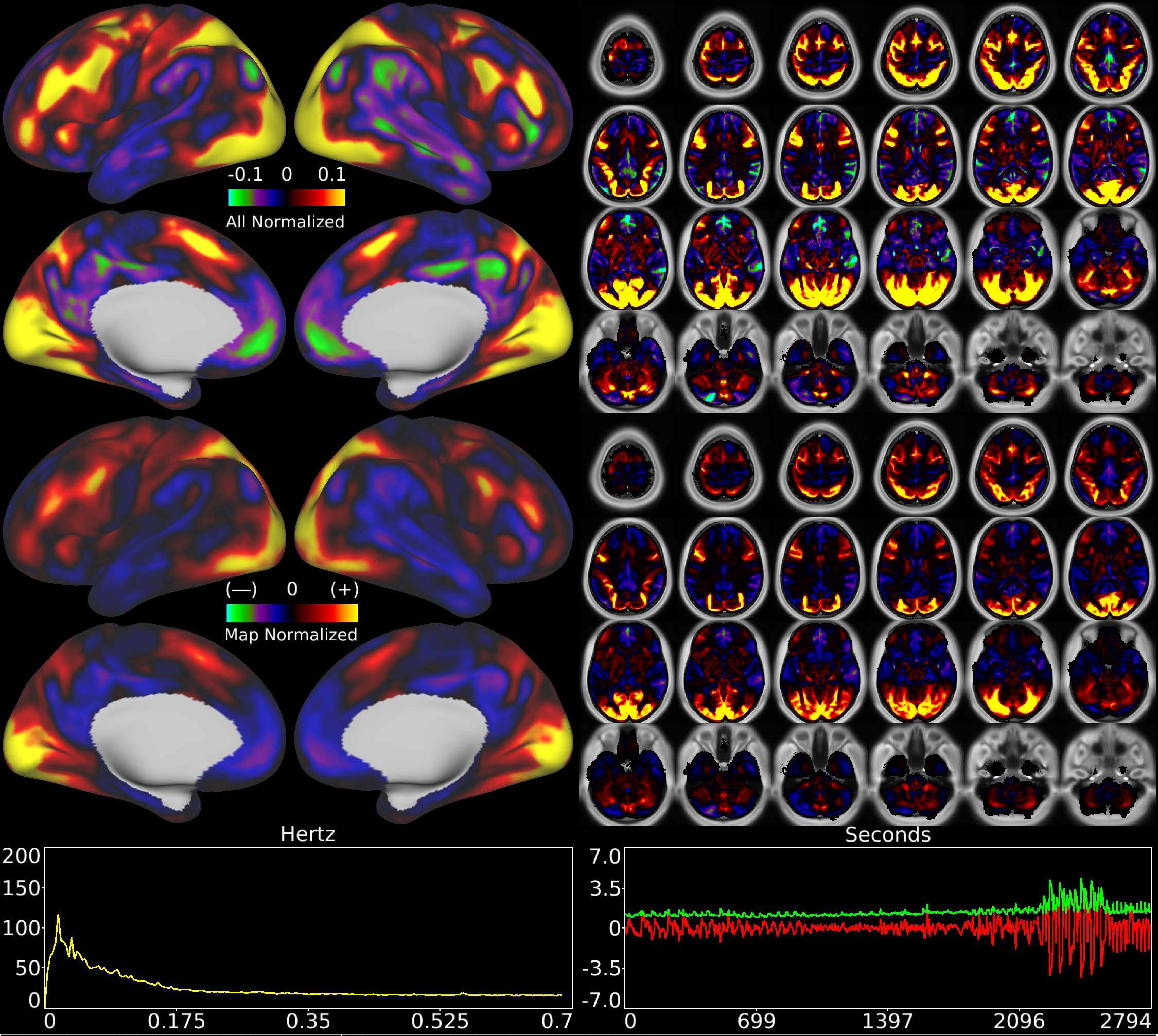


| | | |
|---|---|---|
| Number & Class: 3 Signal | Name: Relational Task Main | |
| RVT Correlated: No | DVARS Dip Associated: No | Cross-Subject Variable: No |
| Single Subject: No | % Variance Explained: 3.63 | Globality Index: 0.42 |
| Rest Component: No | Taskr Component: 10 | Task Modulated: Relational |
| Rationale: Spatial map includes positive and negative patches that are correlated with a specific task design | | |

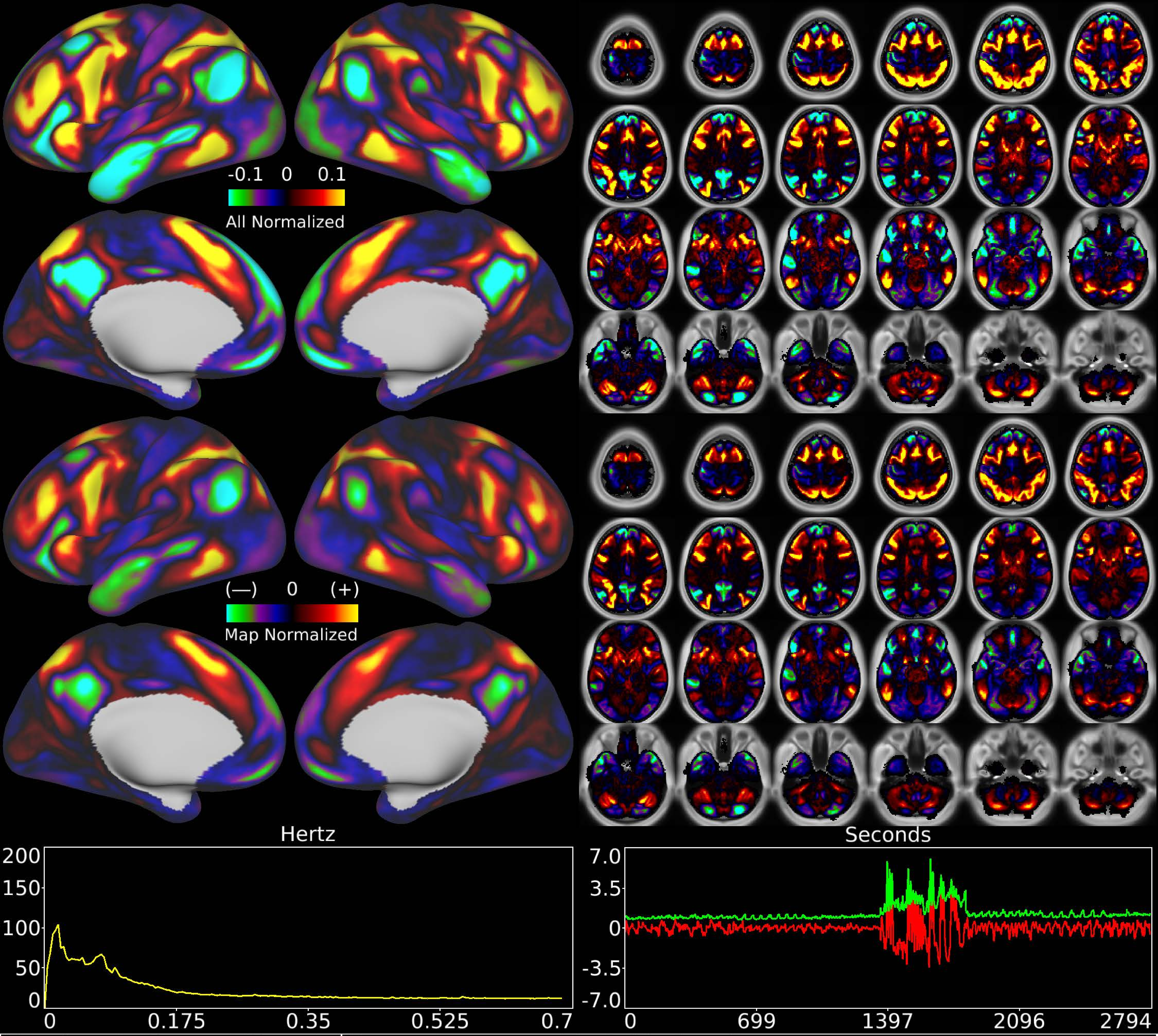


| | | |
|---|---|---|
| Number & Class: 4 Signal | Name: Language Task Math > Story | |
| RVT Correlated: No | DVARS Dip Associated: No | Cross-Subject Variable: No |
| Single Subject: No | % Variance Explained: 3.32 | Globality Index: 0.08 |
| Rest Component: 12 | Taskr Component: 12 | Task Modulated: Language |
| Rationale: Spatial map includes positive and negative patches that are correlated with a specific task design | | |

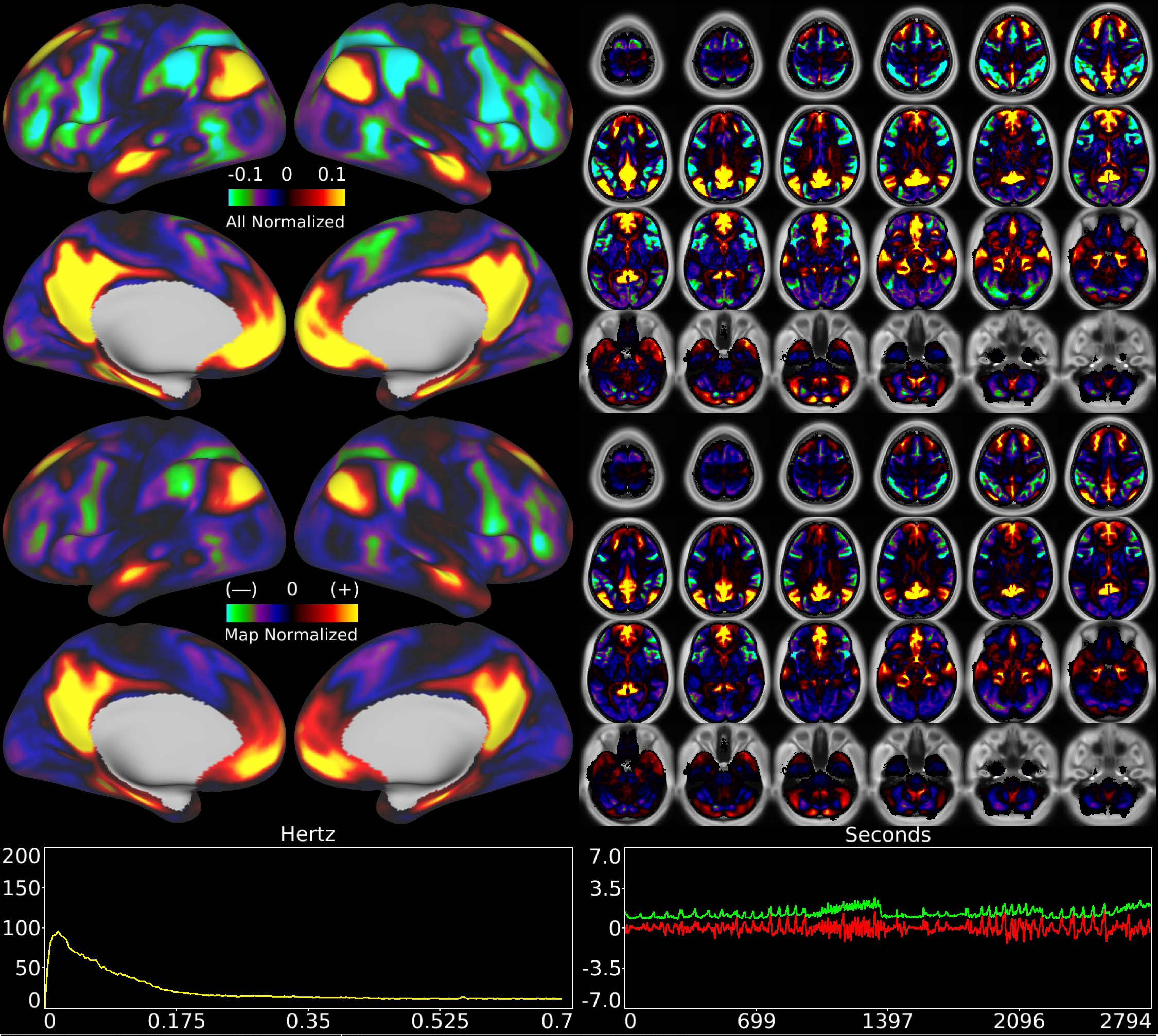


| | | |
|---|---|---|
| Number & Class: 5 Signal | Name: Primary Default Mode | |
| RVT Correlated: No | DVARS Dip Associated: No | Cross-Subject Variable: Yes |
| Single Subject: No | % Variance Explained: 3.3 | Globality Index: 0.95 |
| Rest Component: 10 | Taskr Component: 2 | Task Modulated: Motor + Emotion |
| Rationale: Spatial map includes positive and negative patches that respect known RSNs (e.g. Default Mode Network) | | |

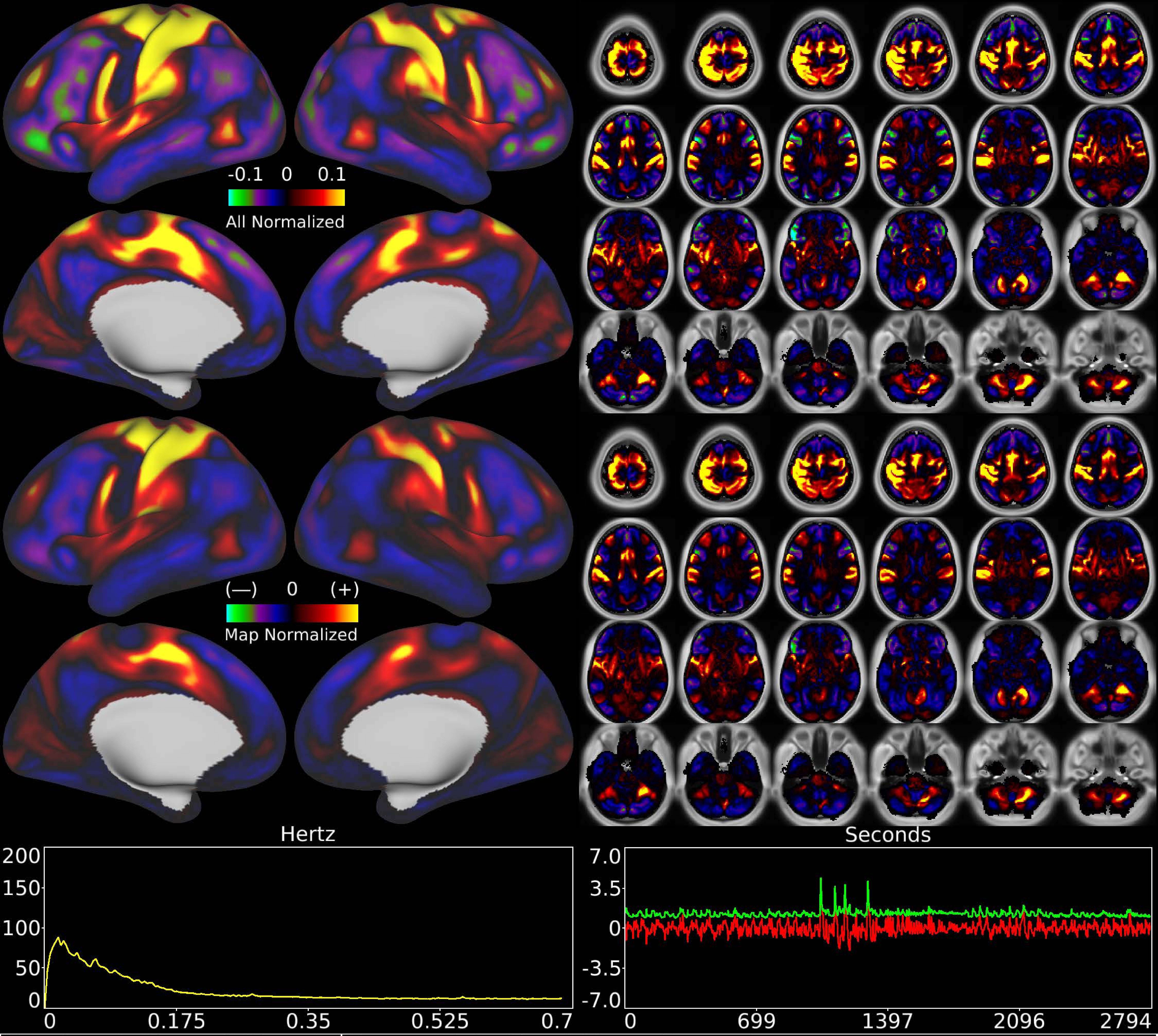


| | | |
|---|---|---|
| Number & Class: 6 Signal | Name: Right Hand Motor Network | |
| RVT Correlated: No | DVARS Dip Associated: Yes | Cross-Subject Variable: Yes |
| Single Subject: No | % Variance Explained: 2.81 | Globality Index: 0.03 |
| Rest Component: 33 | Taskr Component: 7 | Task Modulated: No |
| Rationale: Spatial map includes positive and negative patches that respect known somatotopic sensori-motor organization (Right Hand) | | |

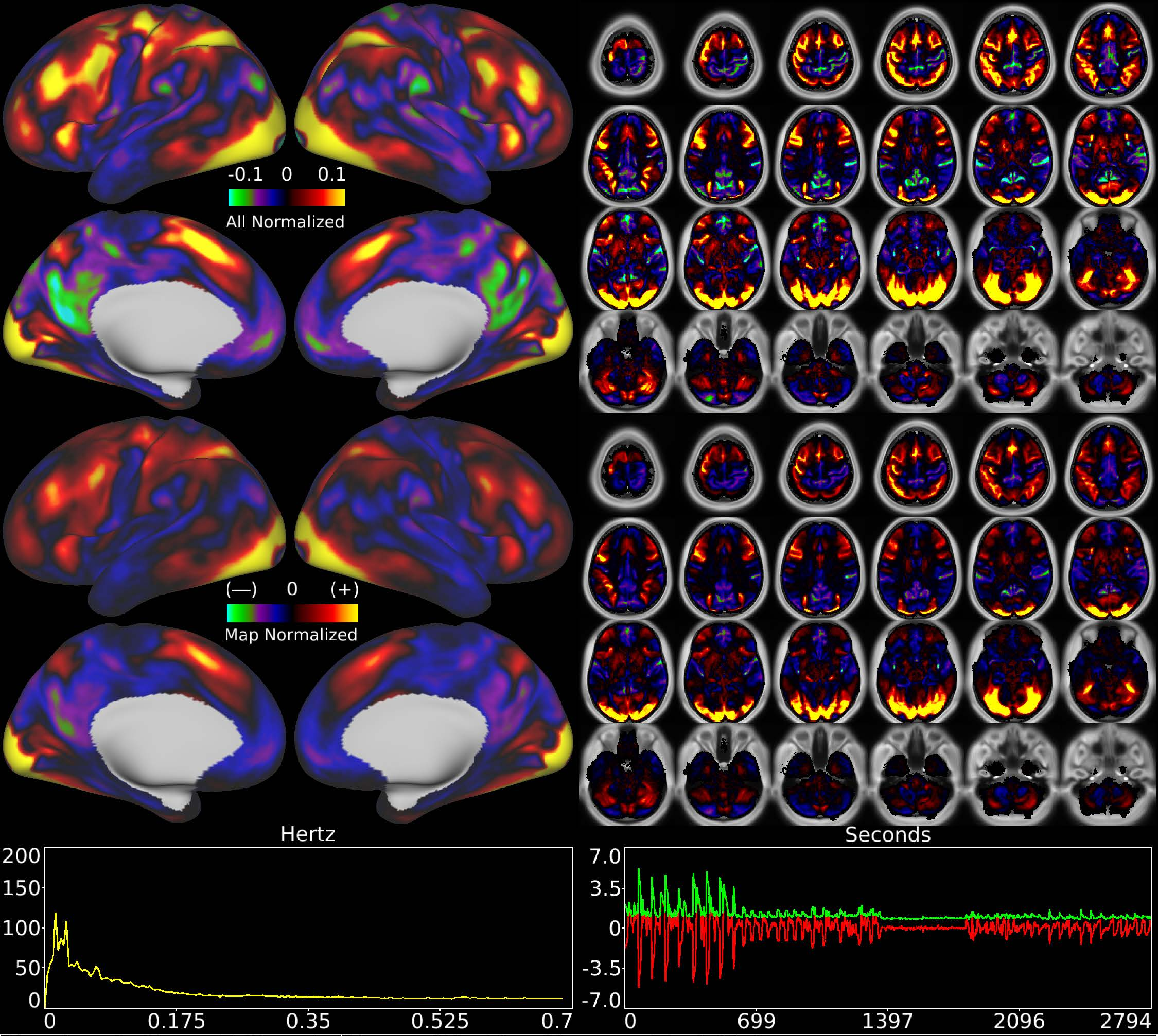


| | | |
|---|---|---|
| Number & Class: 7 Signal | Name: Working Memory Task Main | |
| RVT Correlated: No | DVARS Dip Associated: No | Cross-Subject Variable: No |
| Single Subject: No | % Variance Explained: 2.74 | Globality Index: 0.1 |
| Rest Component: No | Taskr Component: 20+29 | Task Modulated: Working Memory |
| Rationale: Spatial map includes positive and negative patches that are correlated with a specific task design | | |

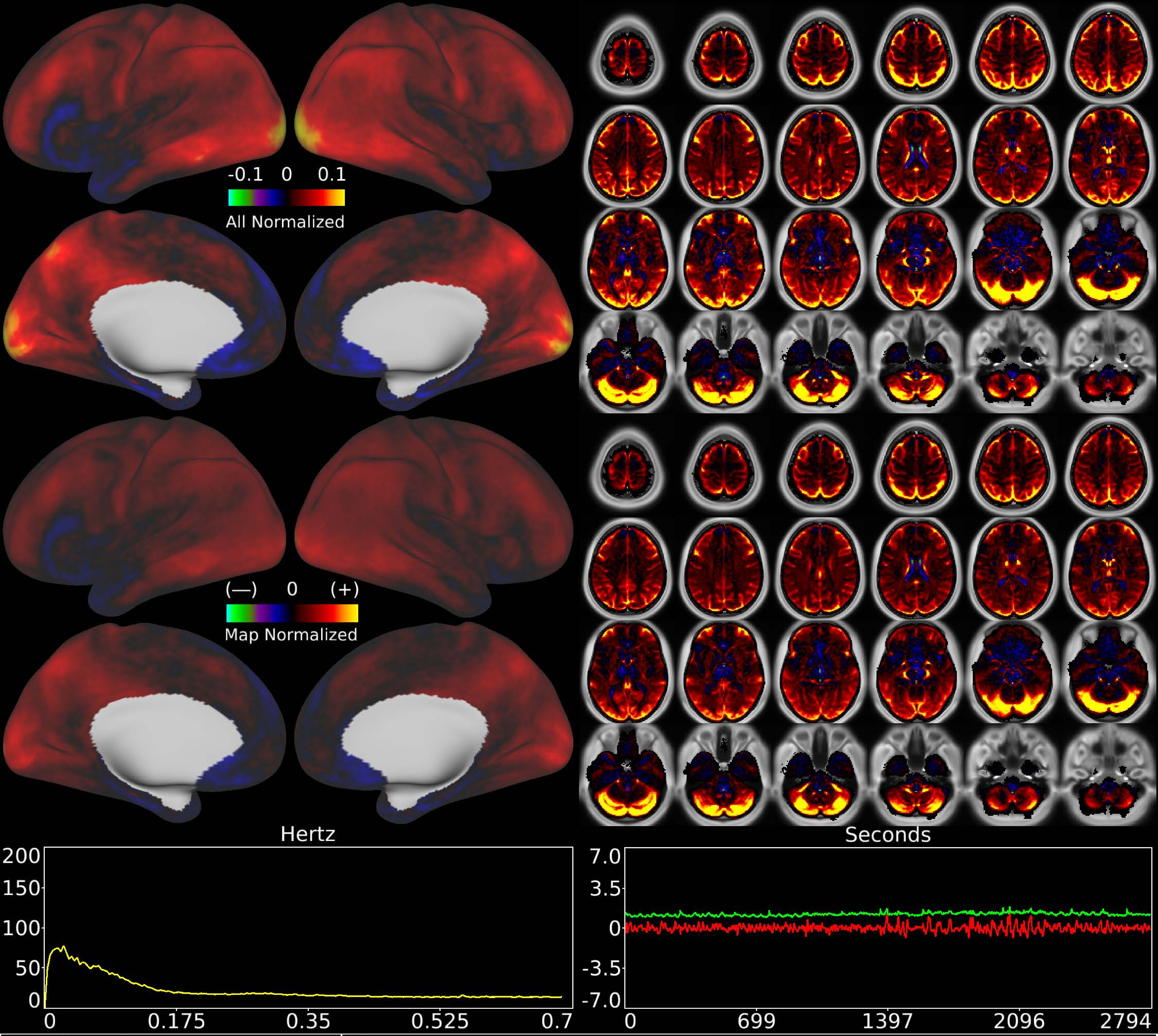


| | | |
|---|---|---|
| Number & Class: 8 Noise | Name: Veins + WM + Cerebellum | |
| RVT Correlated: No | DVARS Dip Associated: Yes | Cross-Subject Variable: Yes |
| Single Subject: No | % Variance Explained: 2.5 | Globality Index: 2.28 |
| Rest Component: 19+28 | Taskr Component: 5 | Task Modulated: No |
| Rationale: Spatial map contains substantial white matter and venous signal | | |

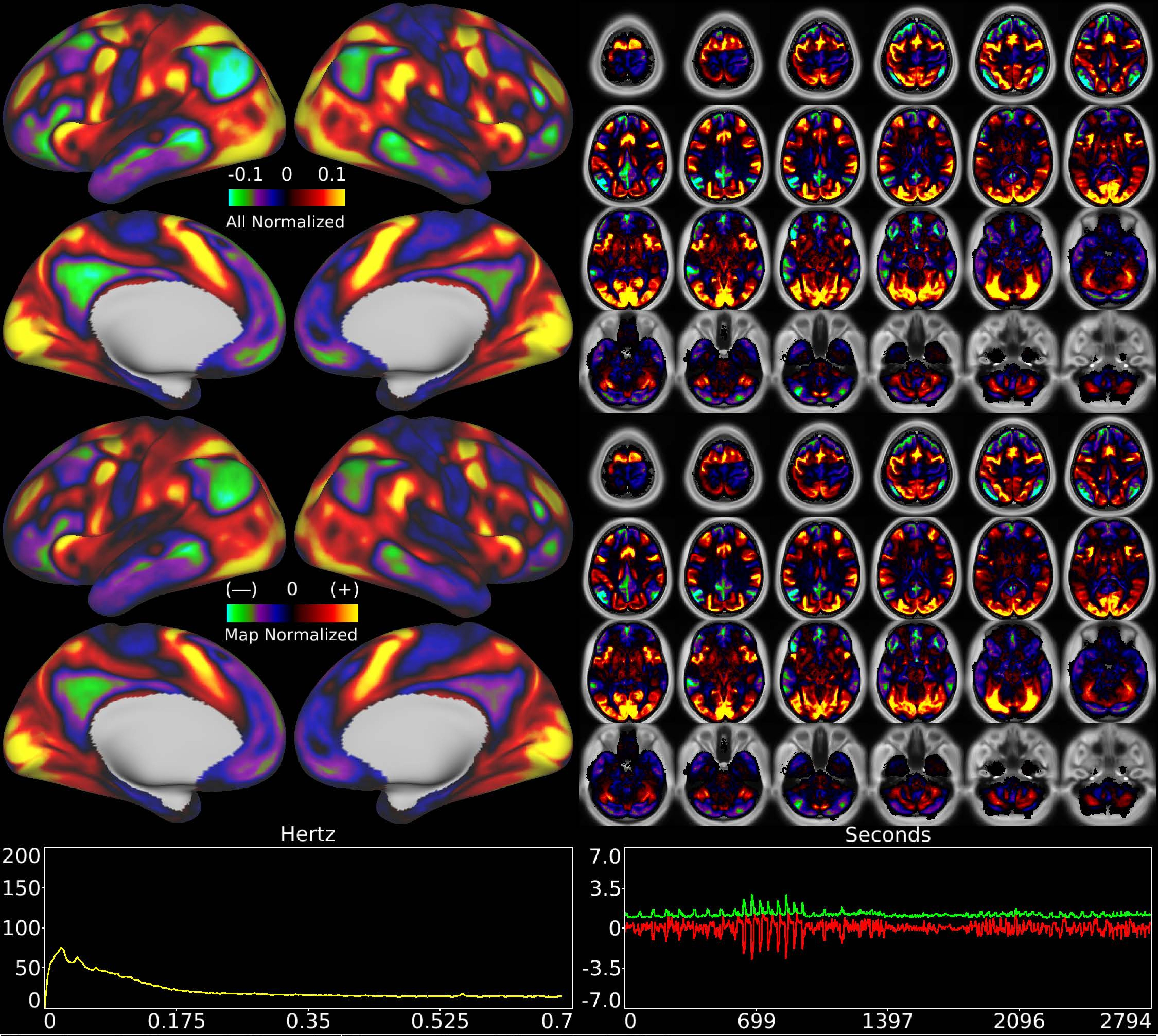


| | | |
|---|---|---|
| Number & Class: 9 Signal | Name: Gambling Task Main | |
| RVT Correlated: No | DVARS Dip Associated: No | Cross-Subject Variable: No |
| Single Subject: No | % Variance Explained: 2.47 | Globality Index: 0.42 |
| Rest Component: -9 | Taskr Component: -8 | Task Modulated: Gambling |

Rationale: Spatial map includes positive and negative patches that are correlated with a specific task design

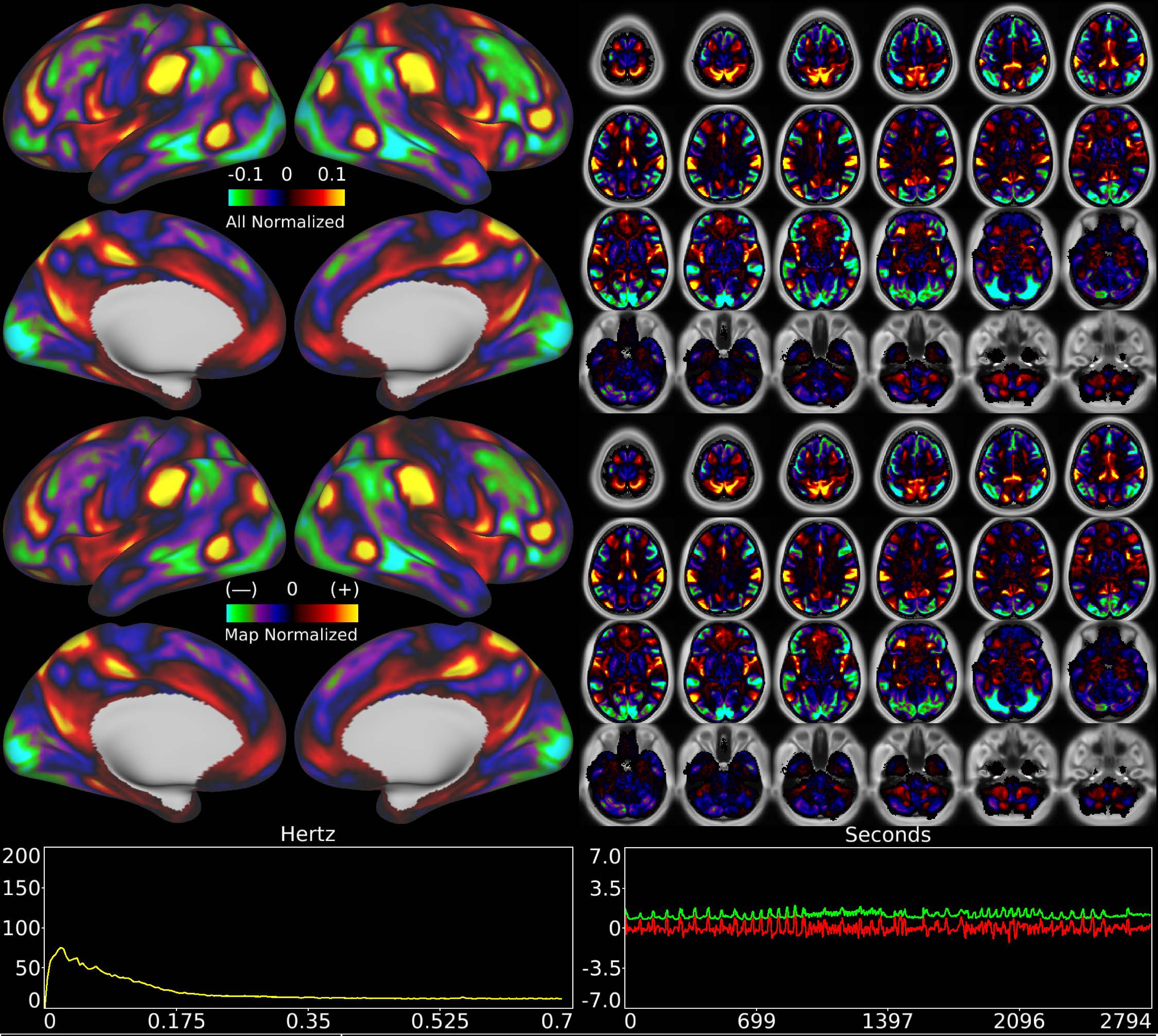


| | | |
|---|---|---|
| Number & Class: 10 Signal | Name: Cingulo-Opercular Network | |
| RVT Correlated: No | DVARS Dip Associated: No | Cross-Subject Variable: No |
| Single Subject: No | % Variance Explained: 2.38 | Globality Index: 0.71 |
| Rest Component: 13 | Taskr Component: 9 | Task Modulated: No |
| Rationale: Spatial map includes positive and negative patches that respect known RSNs (e.g. Cingulo-Opercular Network) | | |

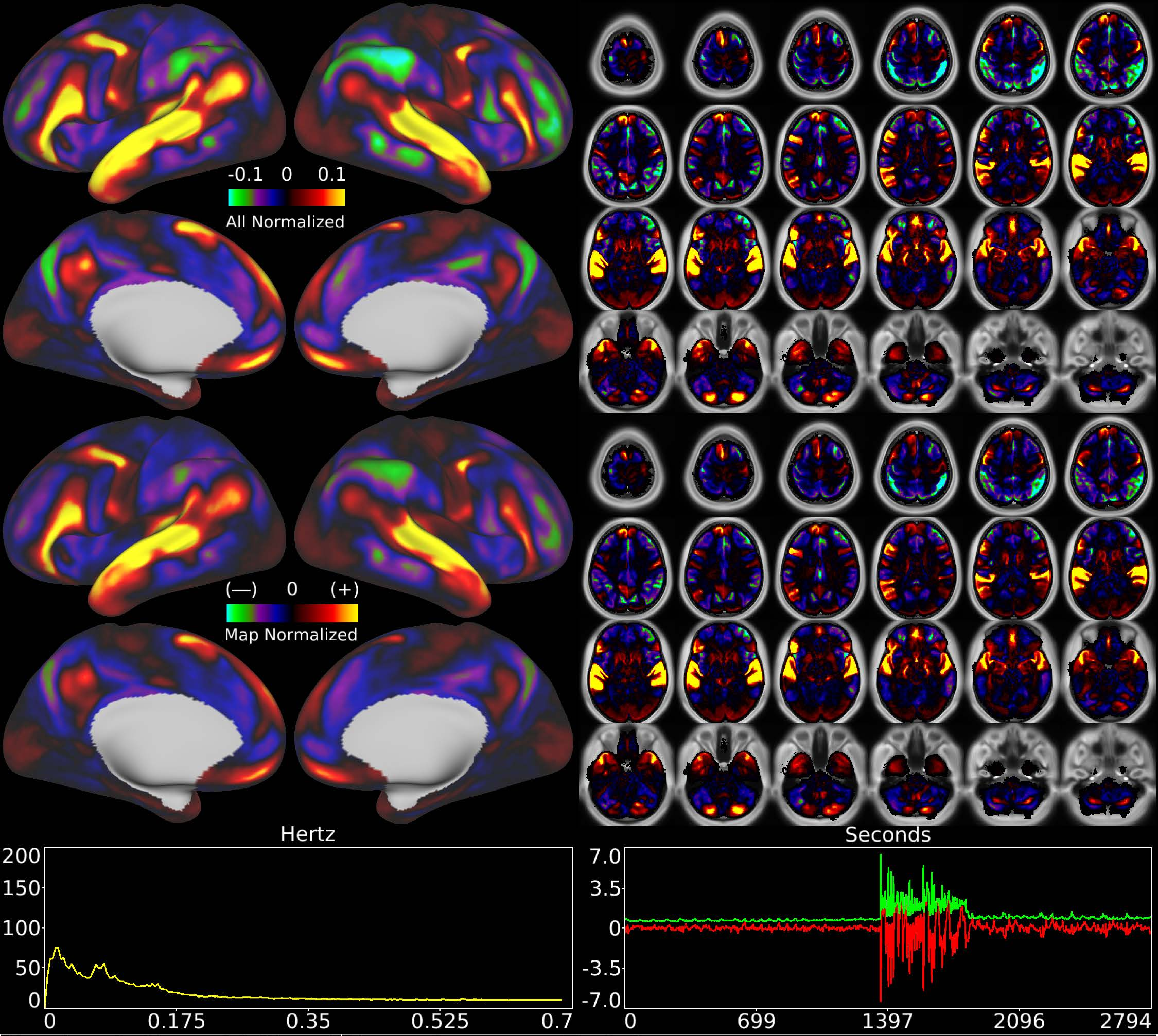


| | | |
|---|---|---|
| Number & Class: 11 Signal | Name: Language Task Story | |
| RVT Correlated: No | DVARS Dip Associated: No | Cross-Subject Variable: No |
| Single Subject: No | % Variance Explained: 2.22 | Globality Index: 0.26 |
| Rest Component: No | Taskr Component: -4 | Task Modulated: Language |
| Rationale: Spatial map includes positive and negative patches that are correlated with a specific task design | | |

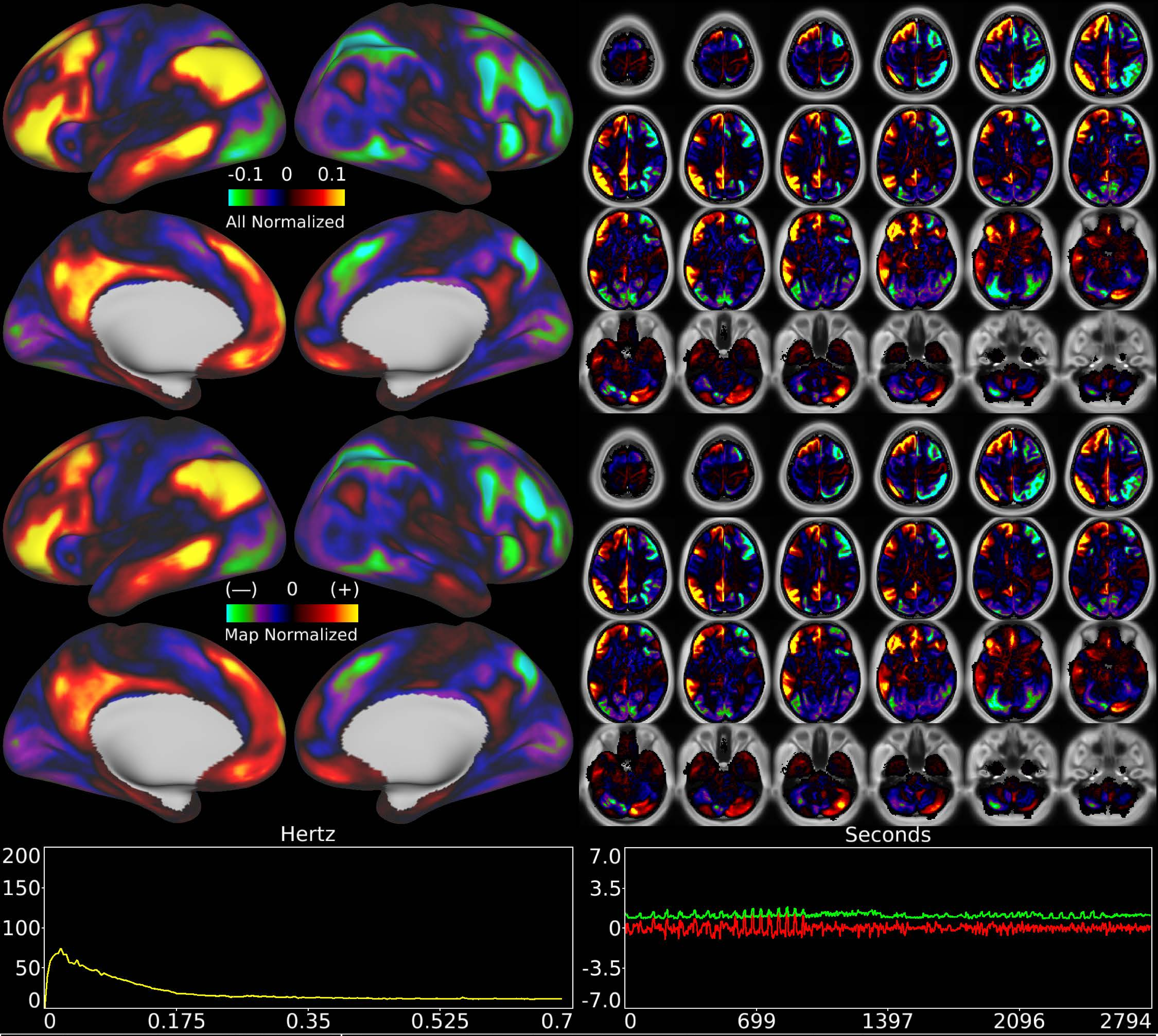


| | | |
|---|---|---|
| Number & Class: 12 Signal | Name: L Default Mode > R Fronto-parietal | |
| RVT Correlated: No | DVARS Dip Associated: No | Cross-Subject Variable: No |
| Single Subject: No | % Variance Explained: 2.11 | Globality Index: 0.56 |
| Rest Component: No | Taskr Component: -6 | Task Modulated: No |
| Rationale: Spatial map includes positive and negative patches that respect known RSNs (e.g. Default Mode Network and Fronto-parietal Network) | | |

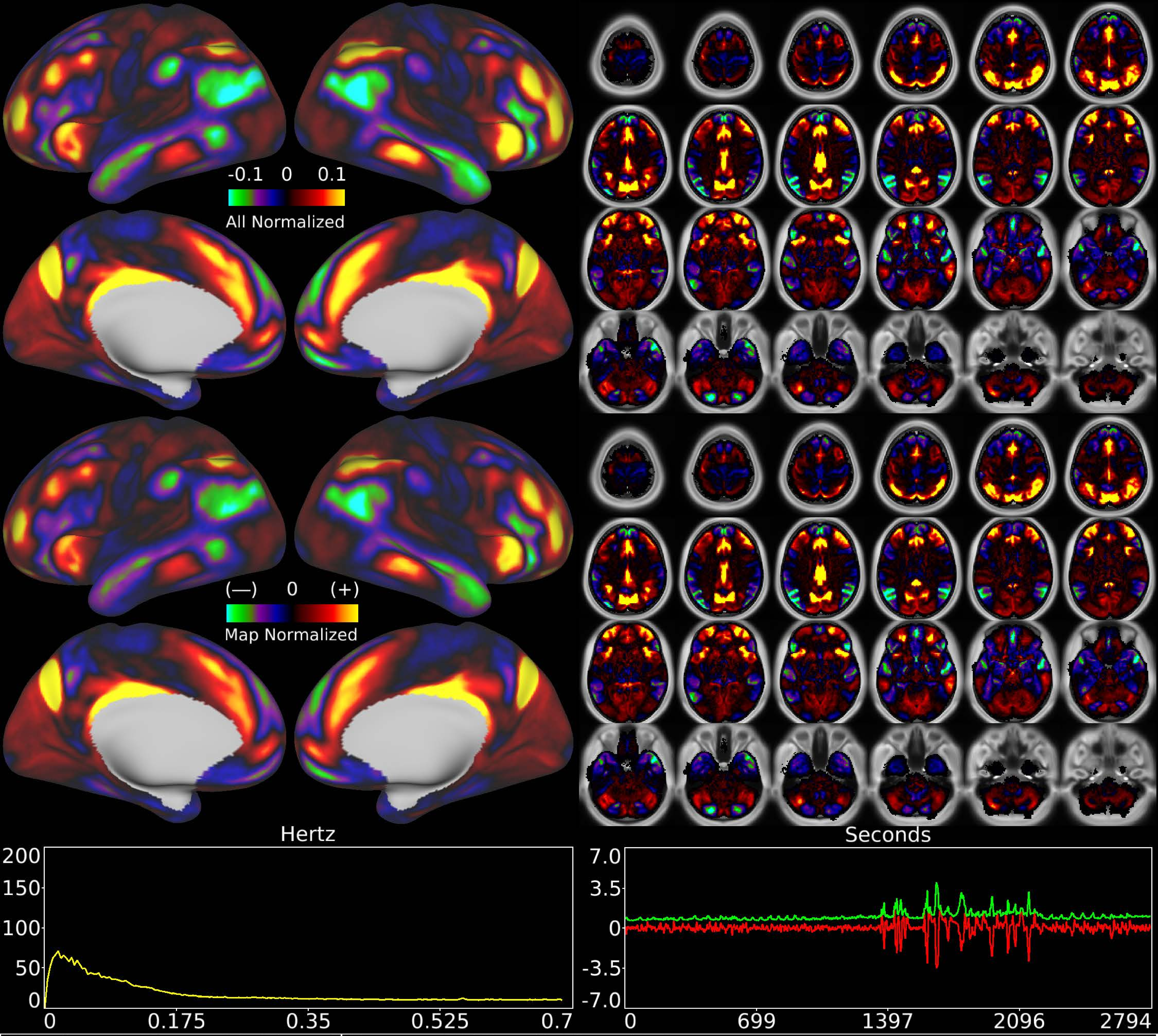


| | | |
|---|---|---|
| Number & Class: 13 Signal | Name: POS2 + RSC vs Unknown | |
| RVT Correlated: No | DVARS Dip Associated: No | Cross-Subject Variable: No |
| Single Subject: No | % Variance Explained: 2.08 | Globality Index: 0.59 |
| Rest Component: 37 | Taskr Component: 21 | Task Modulated: Language + Social |

Rationale: Spatial map includes positive and negative patches that are correlated with two specific task designs

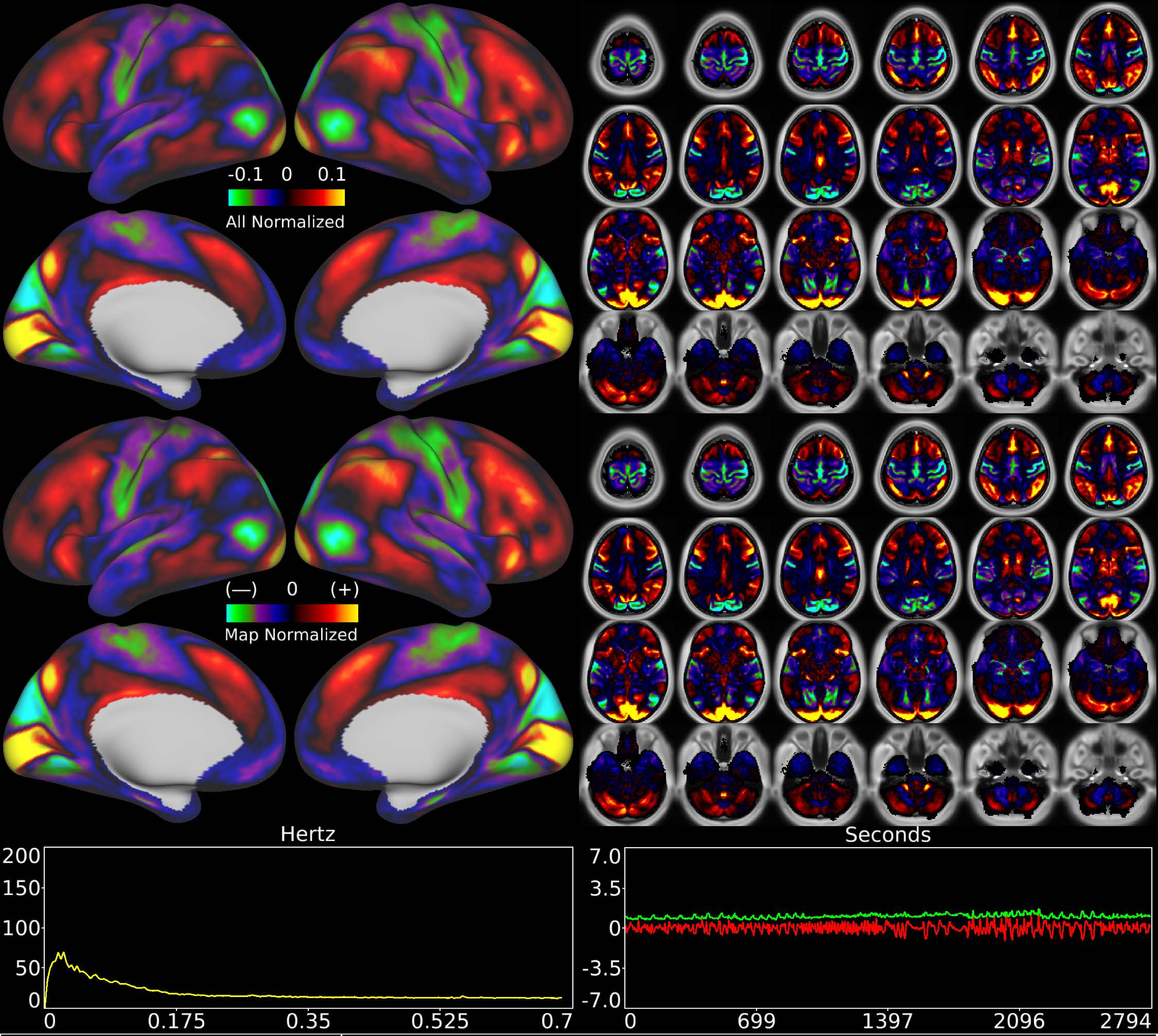


| | | |
|---|---|---|
| Number & Class: 14 Signal | Name: LGN to V1 Variable Component | |
| RVT Correlated: No | DVARS Dip Associated: No | Cross-Subject Variable: Yes |
| Single Subject: No | % Variance Explained: 1.95 | Globality Index: 0.13 |
| Rest Component: 2 | Taskr Component: 14 | Task Modulated: No |
| Rationale: Spatial map includes positive and negative patches that respect known areal boundaries (e.g. V1 and V2) | | |

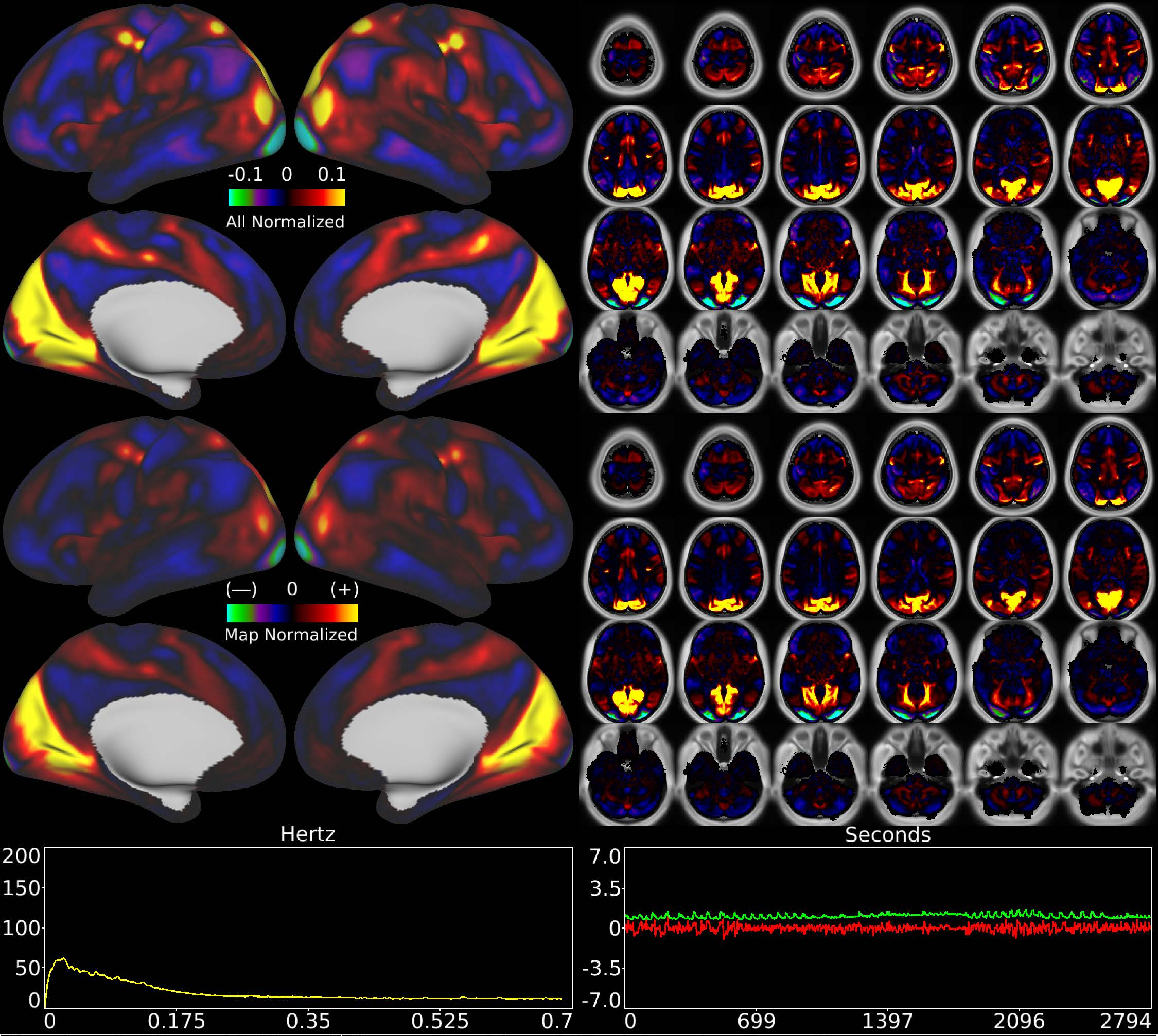


| | | |
|---|---|---|
| Number & Class: 15 Signal | Name: Pan-Visual (Peripheral > Foveal) | |
| RVT Correlated: No | DVARS Dip Associated: No | Cross-Subject Variable: Yes |
| Single Subject: No | % Variance Explained: 1.95 | Globality Index: 0.16 |
| Rest Component: No | Taskr Component: 3 | Task Modulated: No |
| Rationale: Spatial map includes positive and negative patches that respect known retinotopic visual organization (Peripheral vs Foveal) | | |

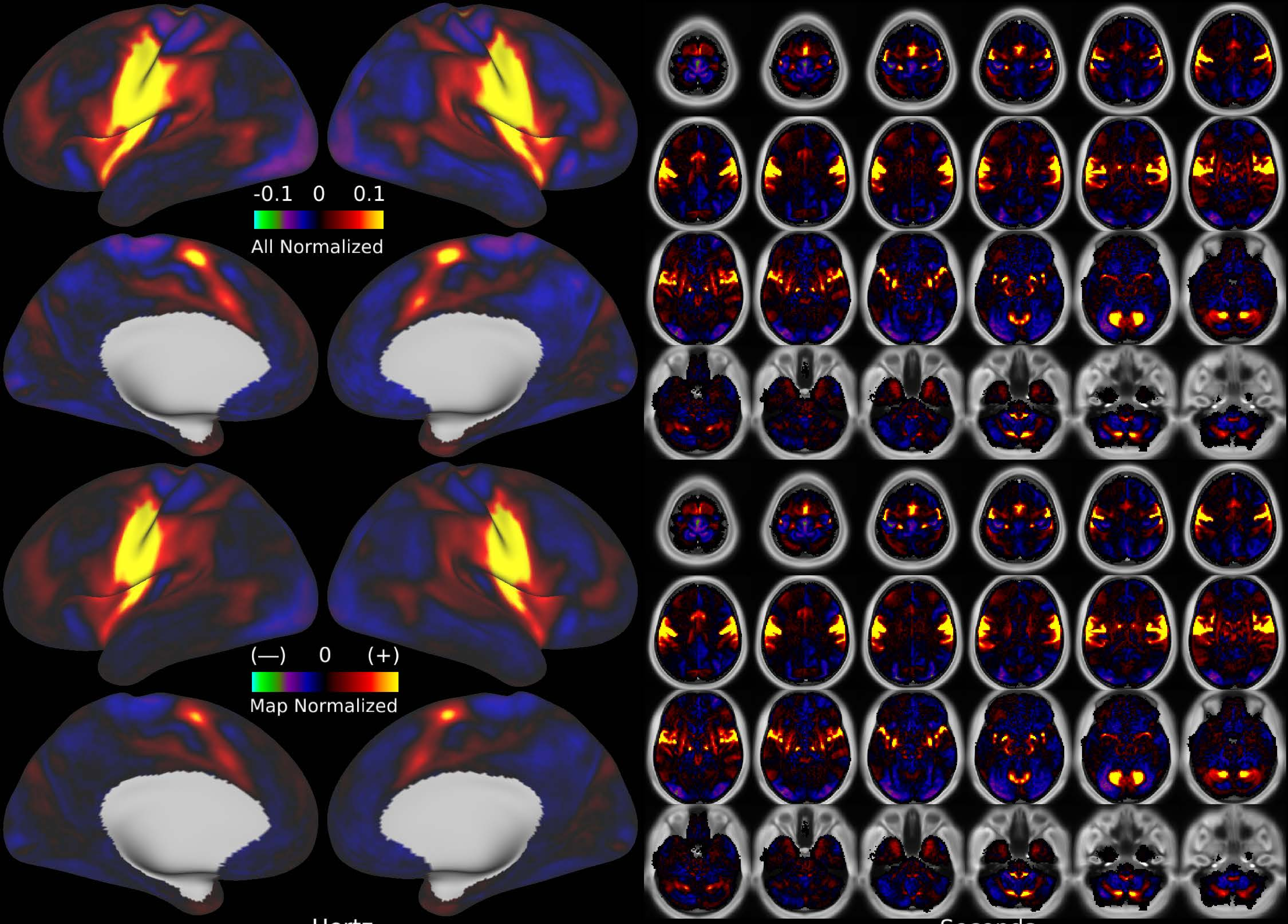


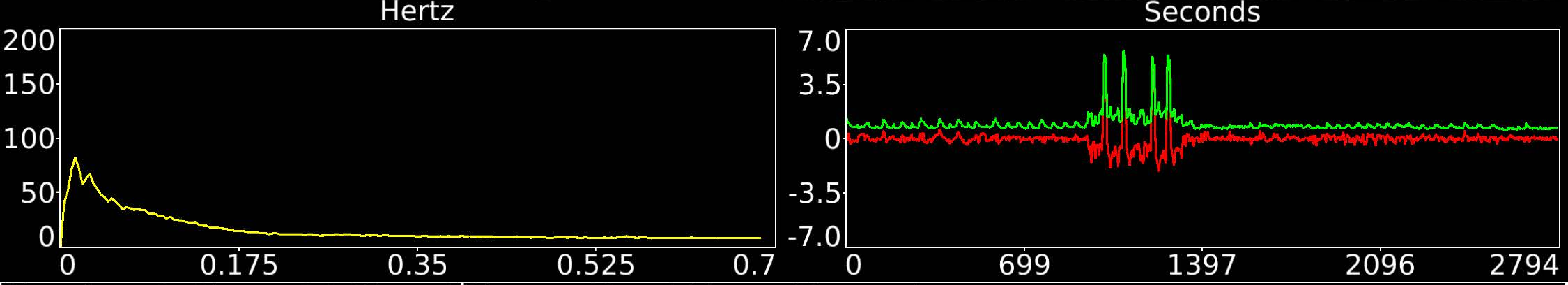


| | | |
|---|---|---|
| Number & Class: 16 Signal | Name: Head Motor Network | |
| RVT Correlated: No | DVARS Dip Associated: Yes | Cross-Subject Variable: Yes |
| Single Subject: No | % Variance Explained: 1.92 | Globality Index: 0.13 |
| Rest Component: 27 | Taskr Component: 18 | Task Modulated: Motor |
| Rationale: Spatial map includes positive and negative patches that respect known somatotopic sensori-motor organization (Face) | | |

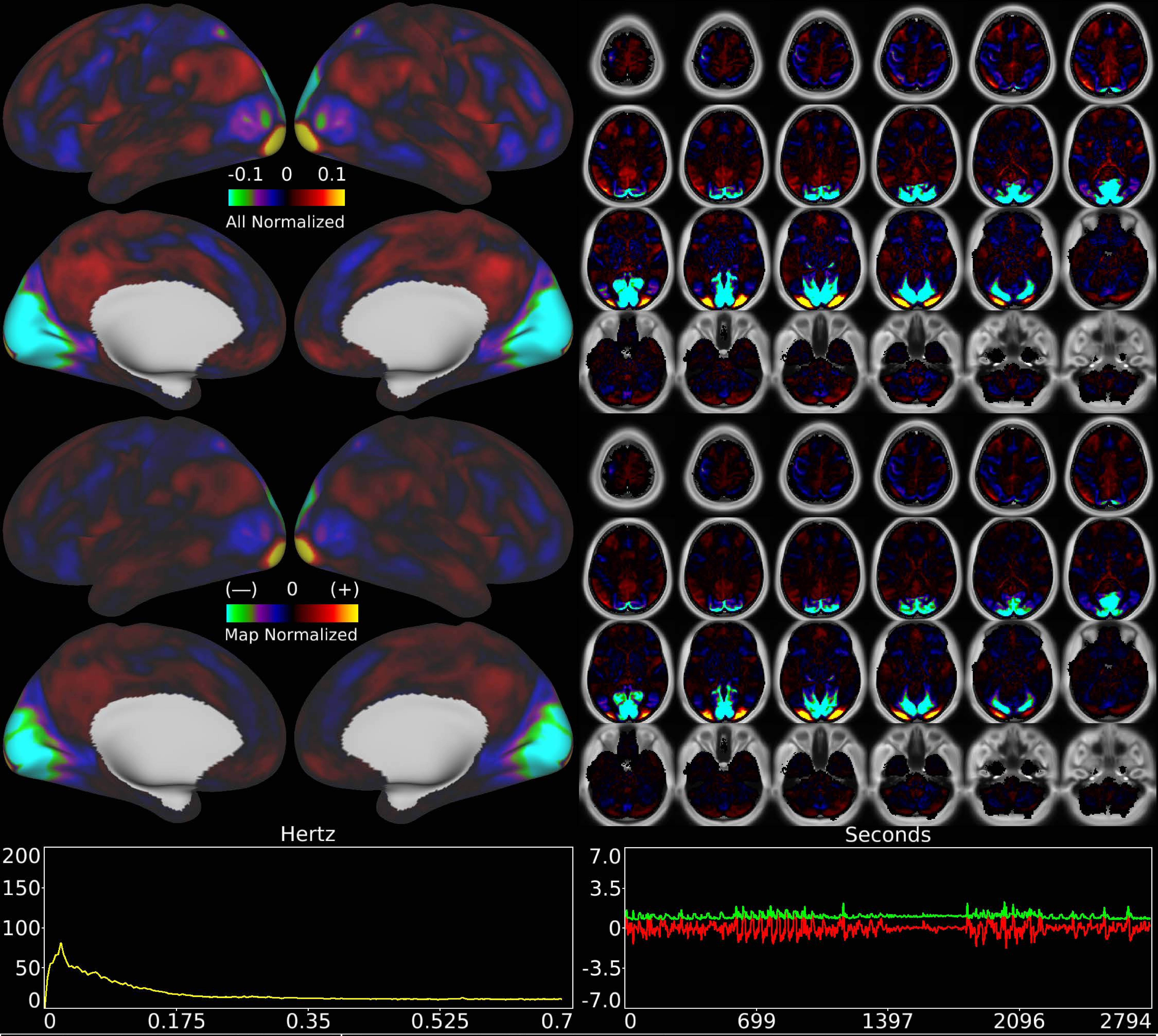


| | | |
|---|---|---|
| Number & Class: 17 Signal | Name: Early Visual Foveal > Peripheral | |
| RVT Correlated: No | DVARS Dip Associated: No | Cross-Subject Variable: No |
| Single Subject: No | % Variance Explained: 1.87 | Globality Index: 0.04 |
| Rest Component: 16 | Taskr Component: 10 | Task Modulated: No |

Rationale: Spatial map includes positive and negative patches that respect known retinotopic visual organization (Foveal vs Peripheral)

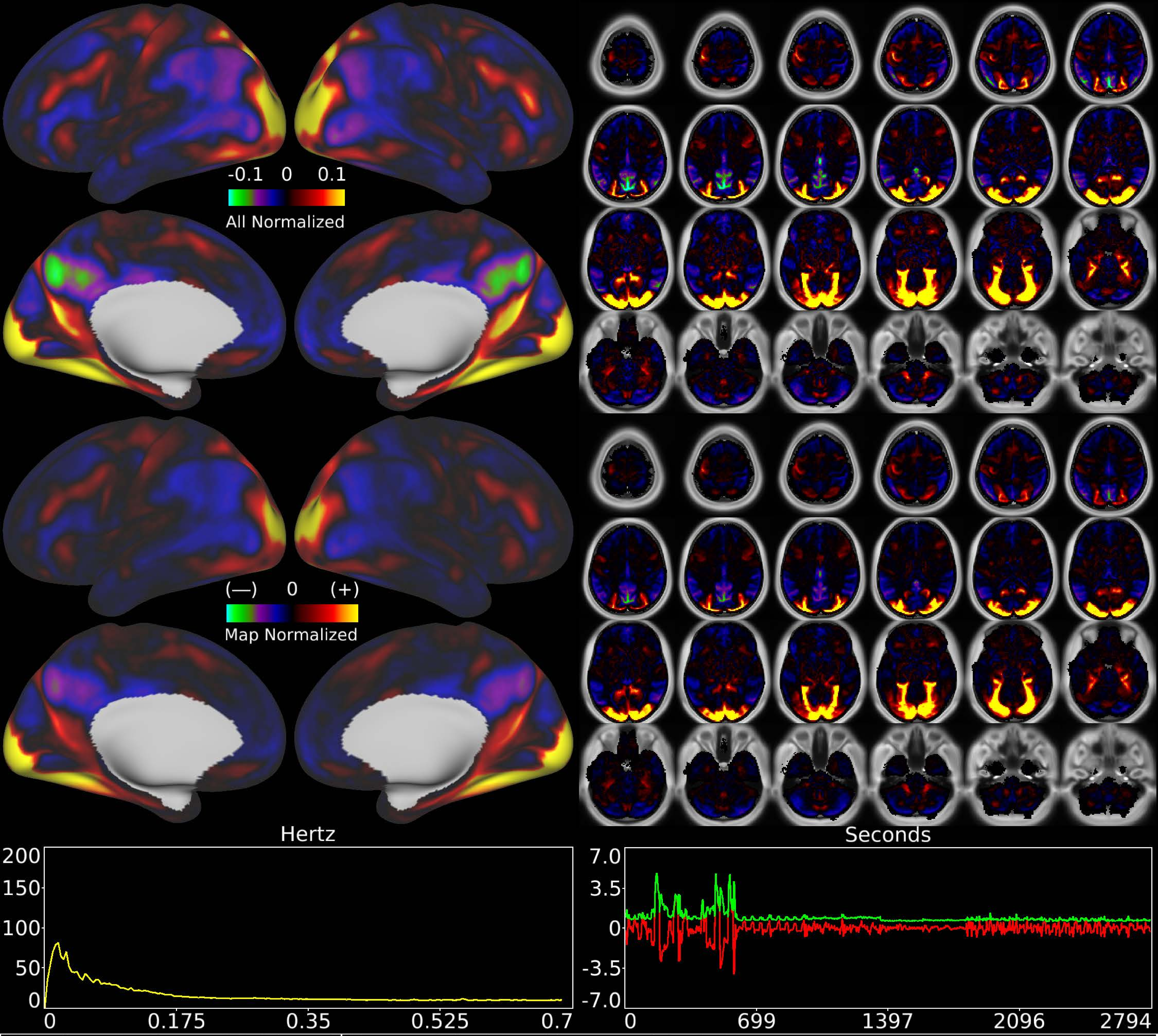


| | | |
|---|---|---|
| Number & Class: 18 Signal | Name: Working Memory Task Place | |
| RVT Correlated: No | DVARS Dip Associated: No | Cross-Subject Variable: No |
| Single Subject: No | % Variance Explained: 1.79 | Globality Index: 0.27 |
| Rest Component: No | Taskr Component: 7 | Task Modulated: Working Memory |

Rationale: Spatial map includes positive and negative patches that are correlated with a specific task design

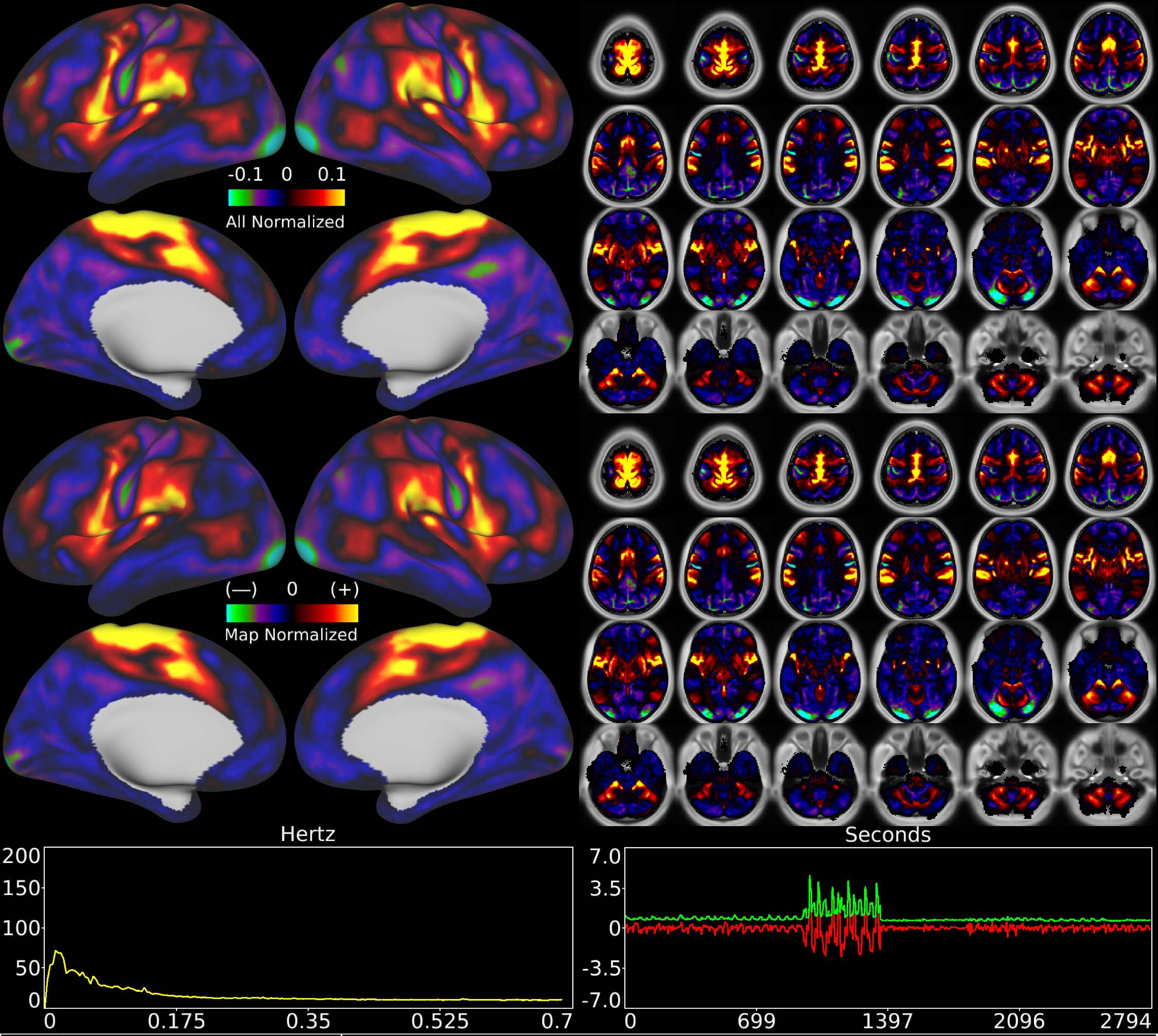


| | | |
|---|---|---|
| Number & Class: 19 Signal | Name: Feet Motor Network | |
| RVT Correlated: No | DVARS Dip Associated: Yes | Cross-Subject Variable: Yes |
| Single Subject: No | % Variance Explained: 1.7 | Globality Index: 0.48 |
| Rest Component: 48 | Taskr Component: 24 | Task Modulated: Motor |
| Rationale: Spatial map includes positive and negative patches that respect known somatotopic sensori-motor organization (Feet) | | |

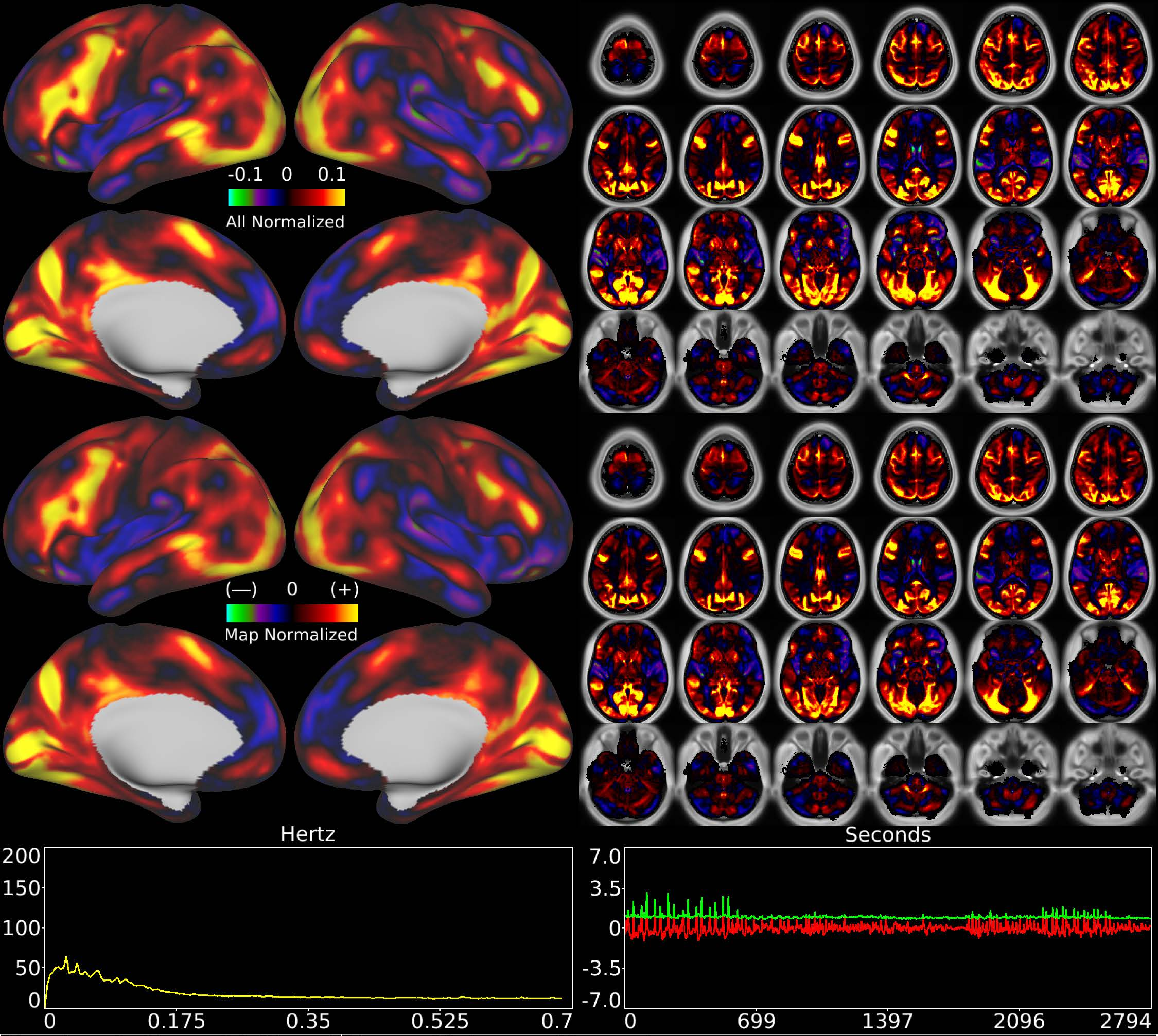


| | | |
|---|---|---|
| Number & Class: 20 Signal | Name: Unknown Network | |
| RVT Correlated: No | DVARS Dip Associated: No | Cross-Subject Variable: No |
| Single Subject: No | % Variance Explained: 1.62 | Globality Index: 1.36 |
| Rest Component: No | Taskr Component: No | Task Modulated: No |
| Rationale: Spatial map includes positive and negative patches that respect known areal boundaries (e.g. POS2) | | |

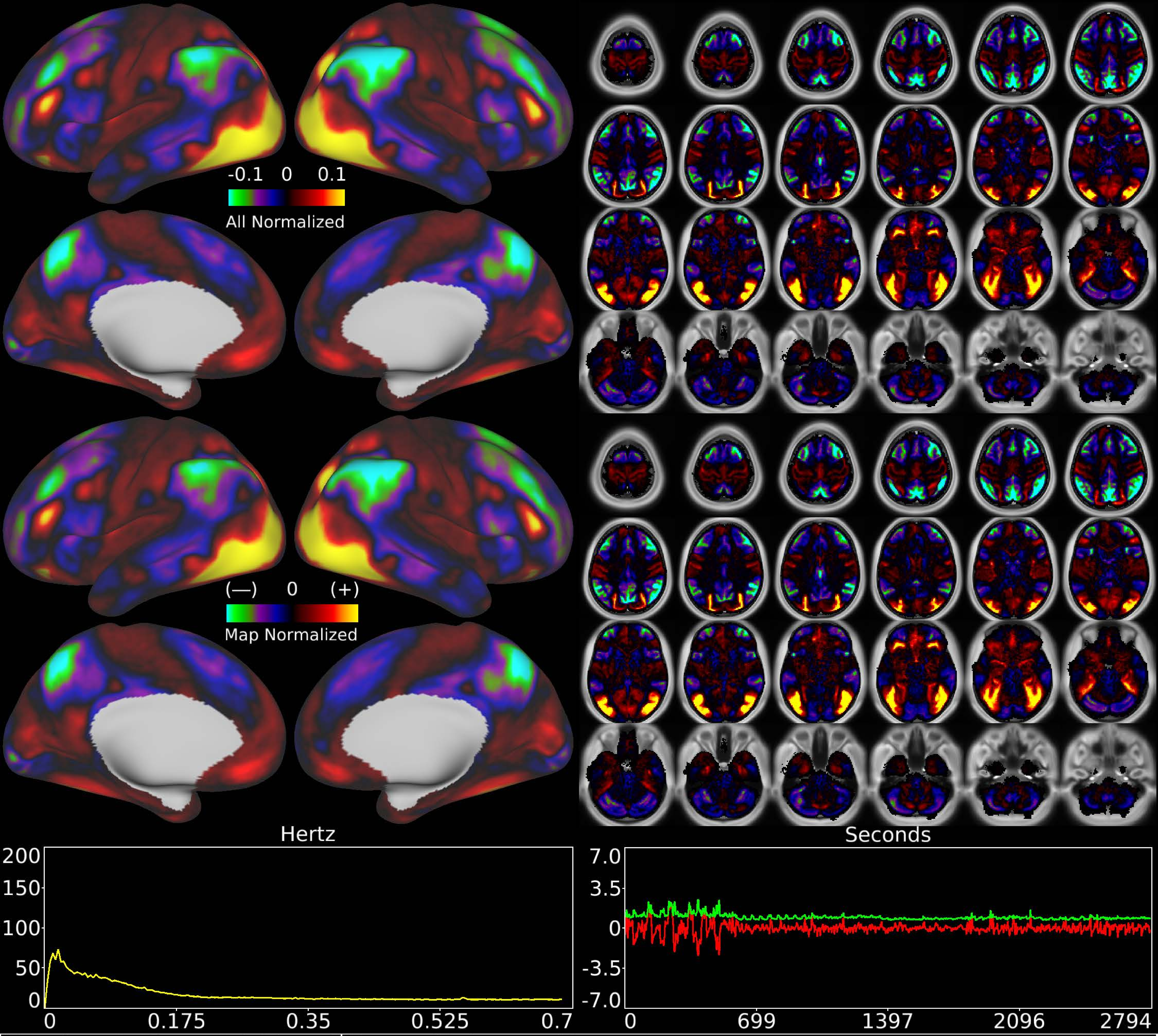


| | | |
|---|---|---|
| Number & Class: 21 Signal | Name: Unknown Network | |
| RVT Correlated: No | DVARS Dip Associated: No | Cross-Subject Variable: No |
| Single Subject: No | % Variance Explained: 1.63 | Globality Index: 0.06 |
| Rest Component: No | Taskr Component: No | Task Modulated: Working Memory |

Rationale: Spatial map includes positive and negative patches that are correlated with a specific task design

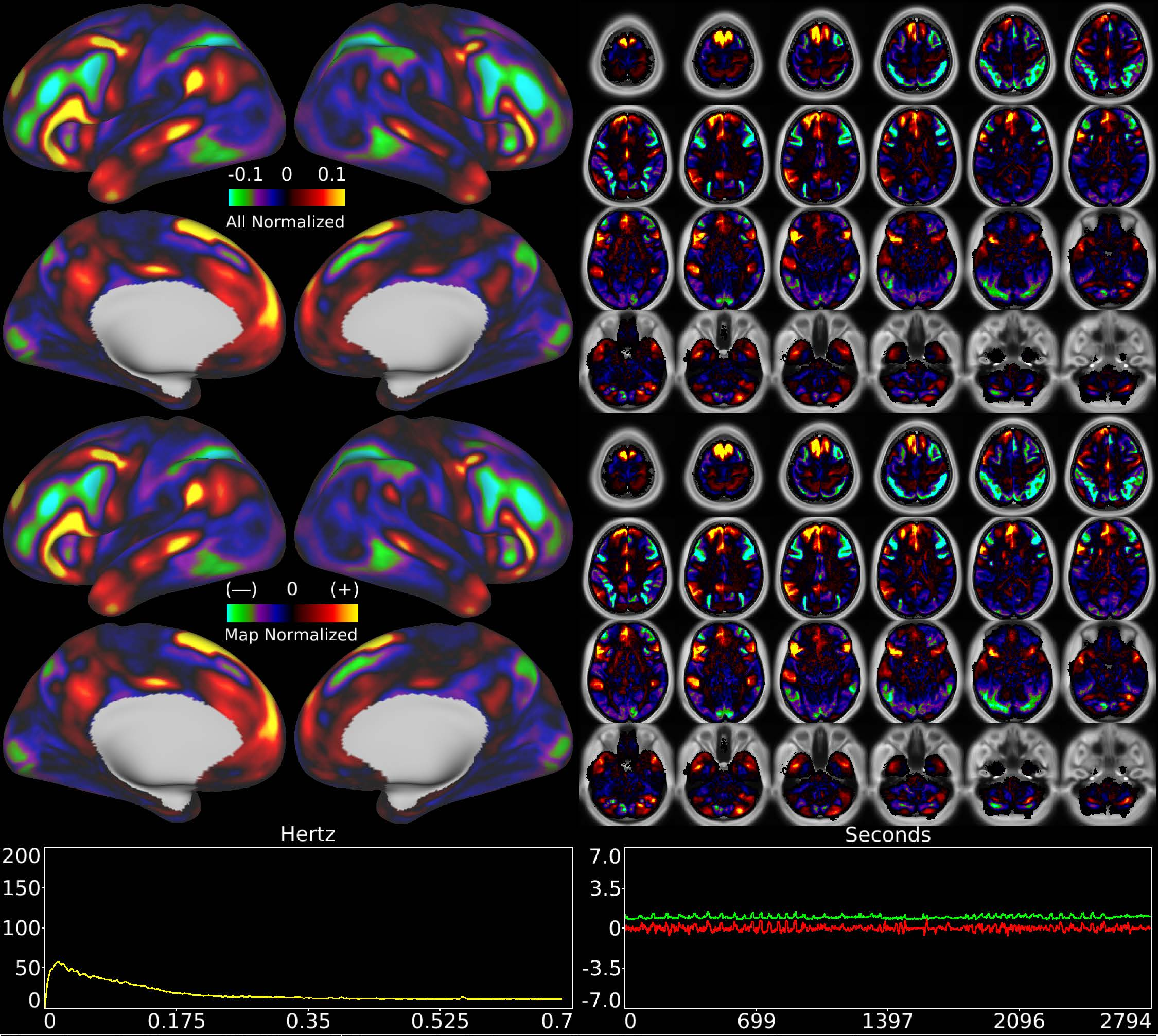


| | | |
|---|---|---|
| Number & Class: 22 Signal | Name: Left Lateralized Language Network | |
| RVT Correlated: No | DVARS Dip Associated: No | Cross-Subject Variable: No |
| Single Subject: No | % Variance Explained: 1.6 | Globality Index: 0.68 |
| Rest Component: 20 | Taskr Component: 30 | Task Modulated: No |

Rationale: Spatial map includes positive and negative patches that respect known RSNs (e.g. Language Network) and areal boundaries (44; 45; 55b; PSL; SFL)

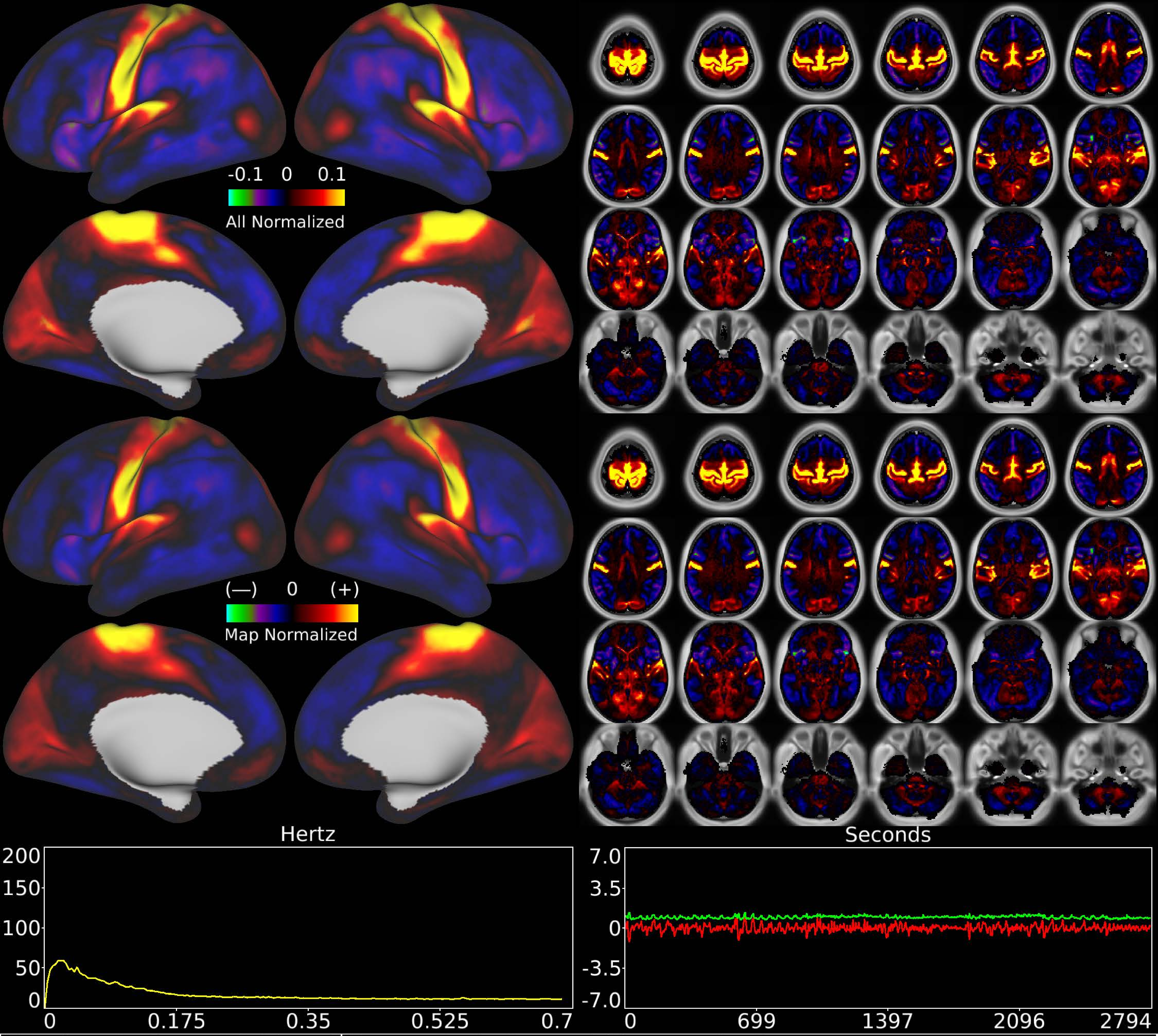


| | | |
|---|---|---|
| Number & Class: 23 Signal | Name: Early Sensori-Motor + Auditory | |
| RVT Correlated: No | DVARS Dip Associated: Yes | Cross-Subject Variable: Yes |
| Single Subject: No | % Variance Explained: 1.57 | Globality Index: 0.02 |
| Rest Component: 5 | Taskr Component: No | Task Modulated: No |
| Rationale: Spatial map includes positive and negative patches that respect known areal boundaries (e.g. around MT+); most specific to sensori-motor cortex | | |

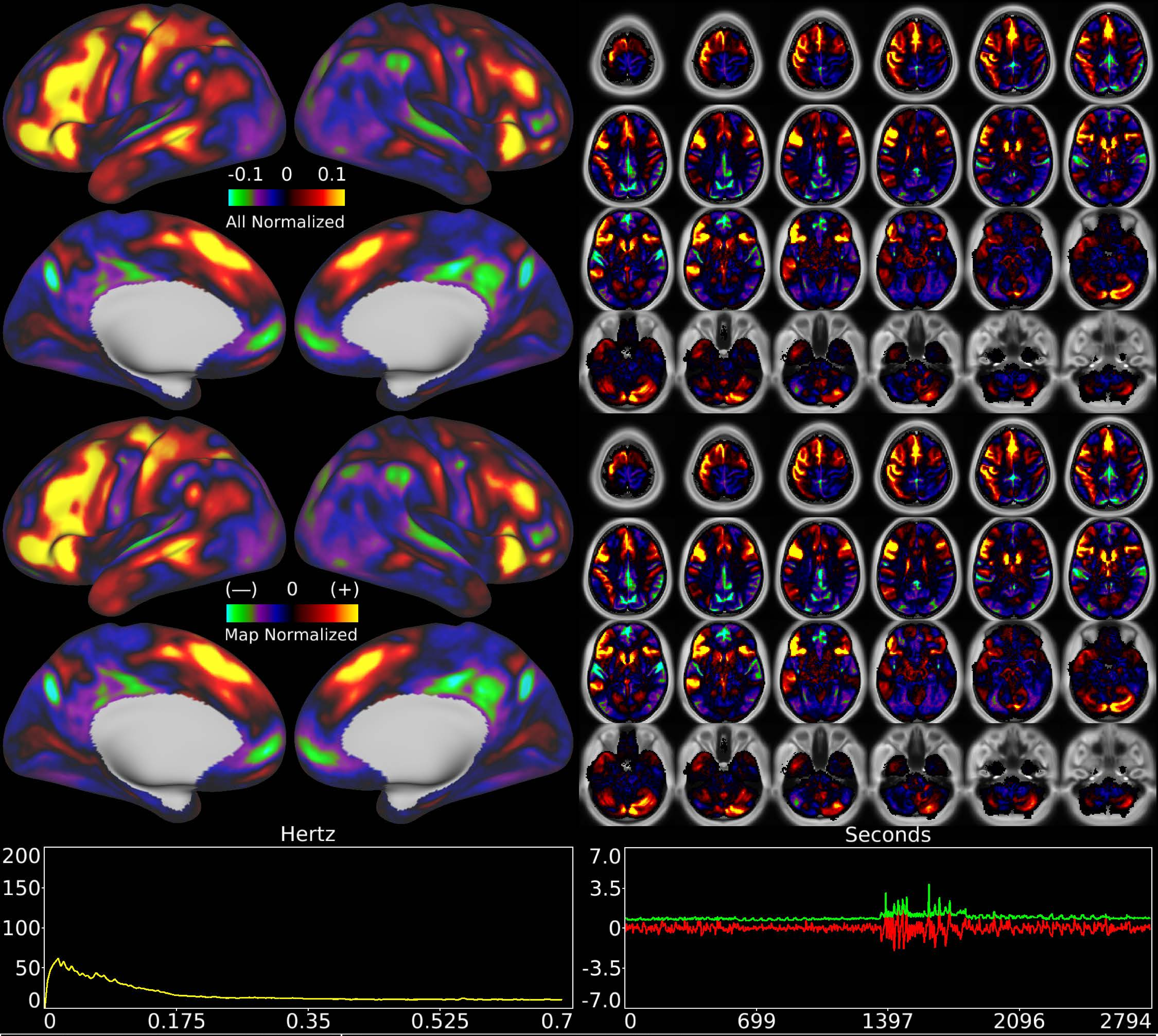


| | | |
|---|---|---|
| Number & Class: 24 Signal | Name: Language Task Math | |
| RVT Correlated: No | DVARS Dip Associated: No | Cross-Subject Variable: No |
| Single Subject: No | % Variance Explained: 1.53 | Globality Index: 0.19 |
| Rest Component: No | Taskr Component: 17 | Task Modulated: Language |
| Rationale: Spatial map includes positive and negative patches that are correlated with a specific task design | | |

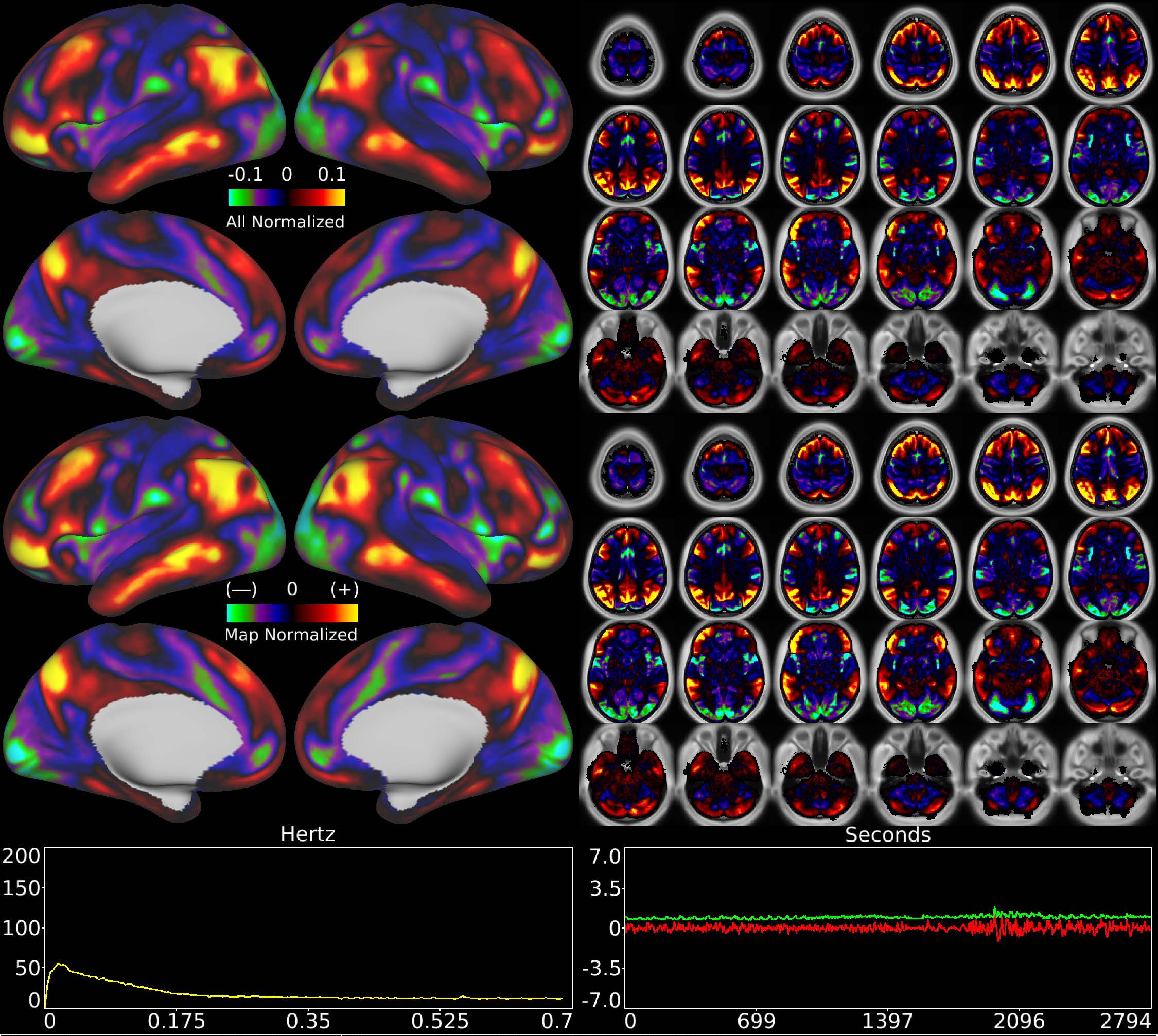


| Number & Class: 25 Signal | Name: Subsidiary Default Mode | |
|---|---|---|
| RVT Correlated: No | DVARS Dip Associated: No | Cross-Subject Variable: No |
| Single Subject: No | % Variance Explained: 1.52 | Globality Index: 0.32 |
| Rest Component: No | Taskr Component: No | Task Modulated: No |

Rationale: Spatial map includes positive and negative patches that respect known RSNs (e.g. Default Mode Network)

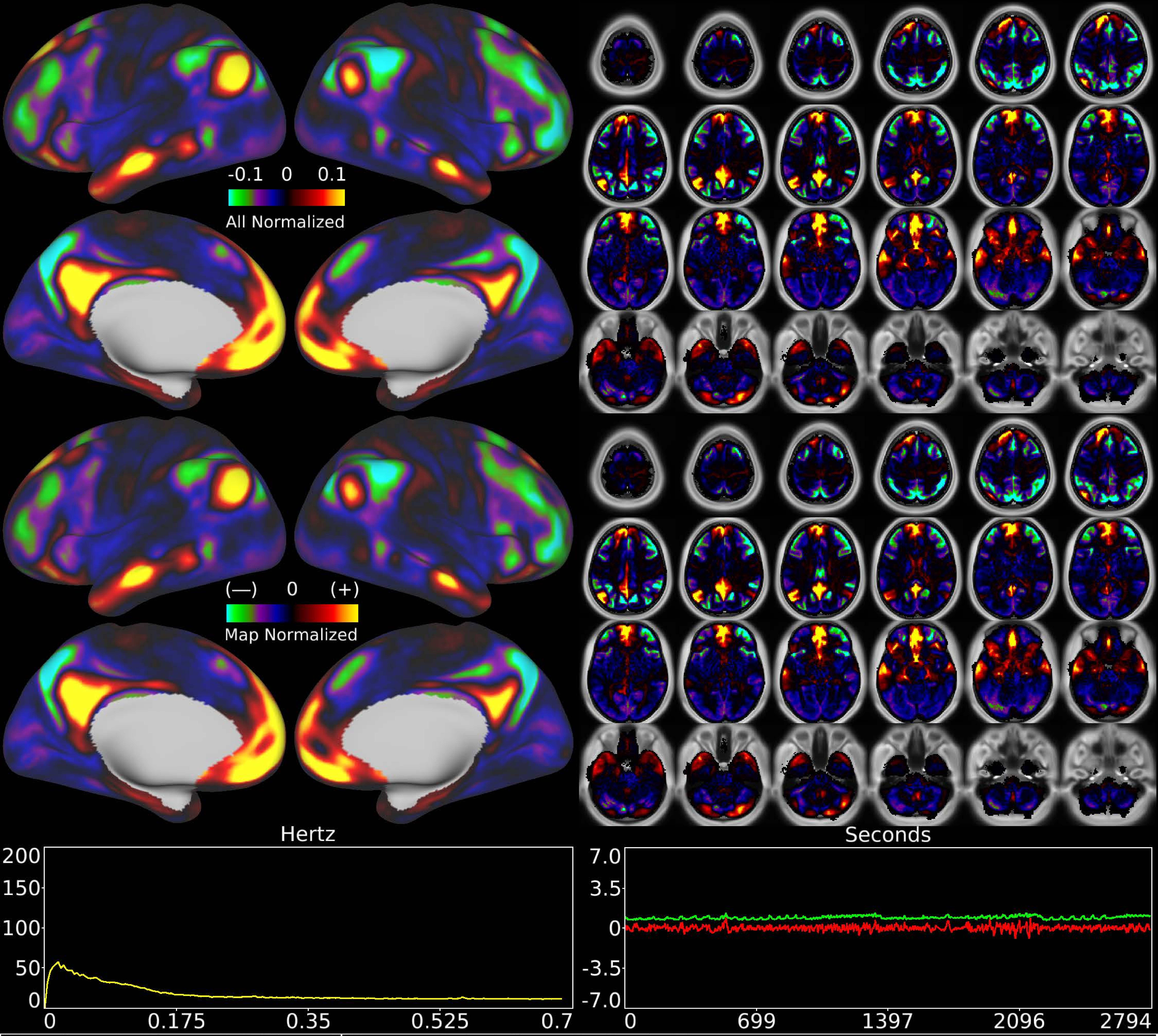


| | | |
|---|---|---|
| Number & Class: 26 Signal | Name: Subsidiary Default Mode | |
| RVT Correlated: No | DVARS Dip Associated: No | Cross-Subject Variable: No |
| Single Subject: No | % Variance Explained: 1.45 | Globality Index: 1.4 |
| Rest Component: 15 | Taskr Component: No | Task Modulated: No |
| Rationale: Spatial map includes positive and negative patches that respect known RSNs (e.g. Default Mode Network) | | |

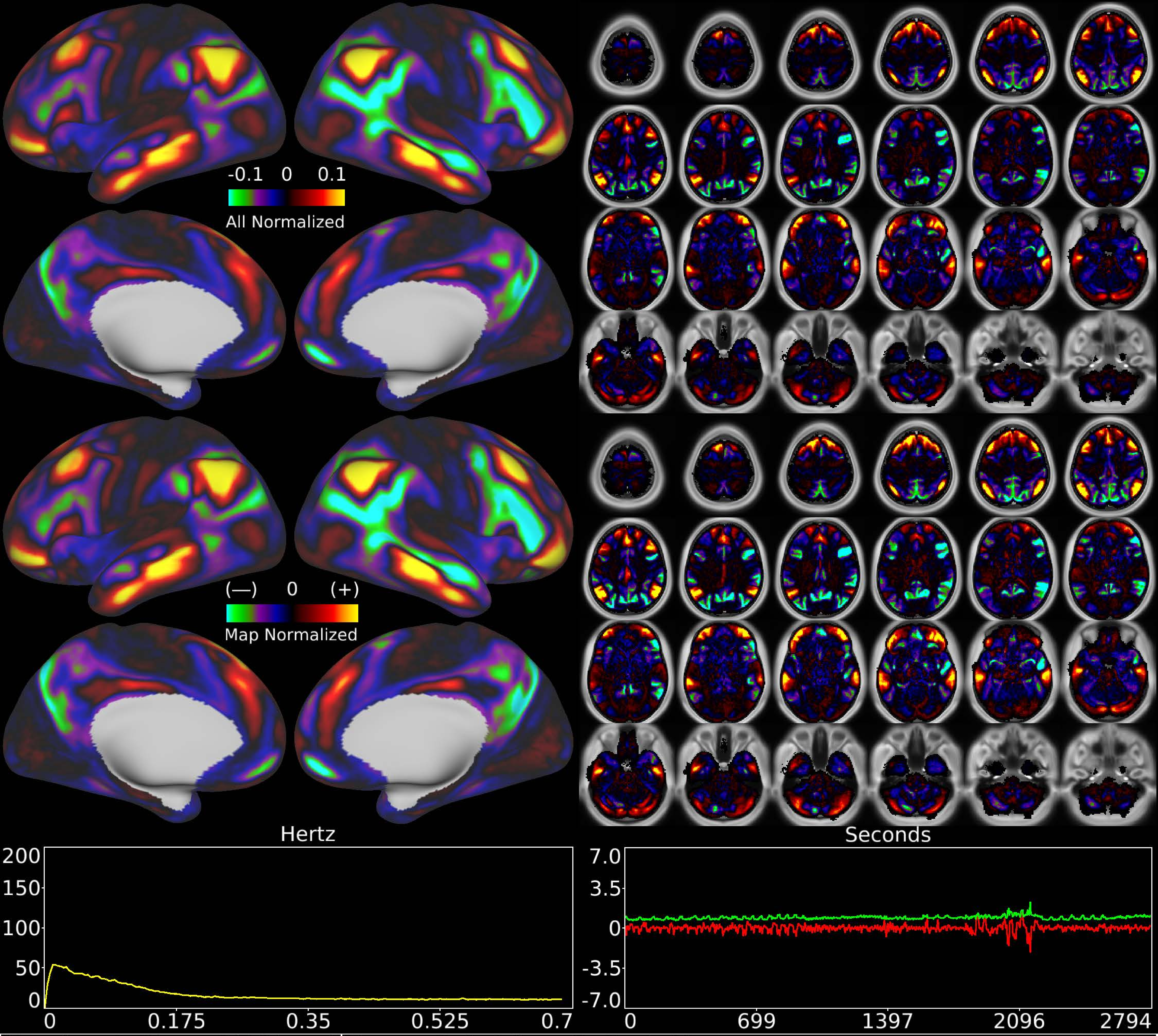


| | | |
|---|---|---|
| Number & Class: 27 Signal | Name: Subsidiary Default Mode | |
| RVT Correlated: No | DVARS Dip Associated: No | Cross-Subject Variable: No |
| Single Subject: No | % Variance Explained: 1.42 | Globality Index: 0.64 |
| Rest Component: No | Taskr Component: No | Task Modulated: No |
| Rationale: Spatial map includes positive and negative patches that respect known RSNs (e.g. Default Mode Network) | | |

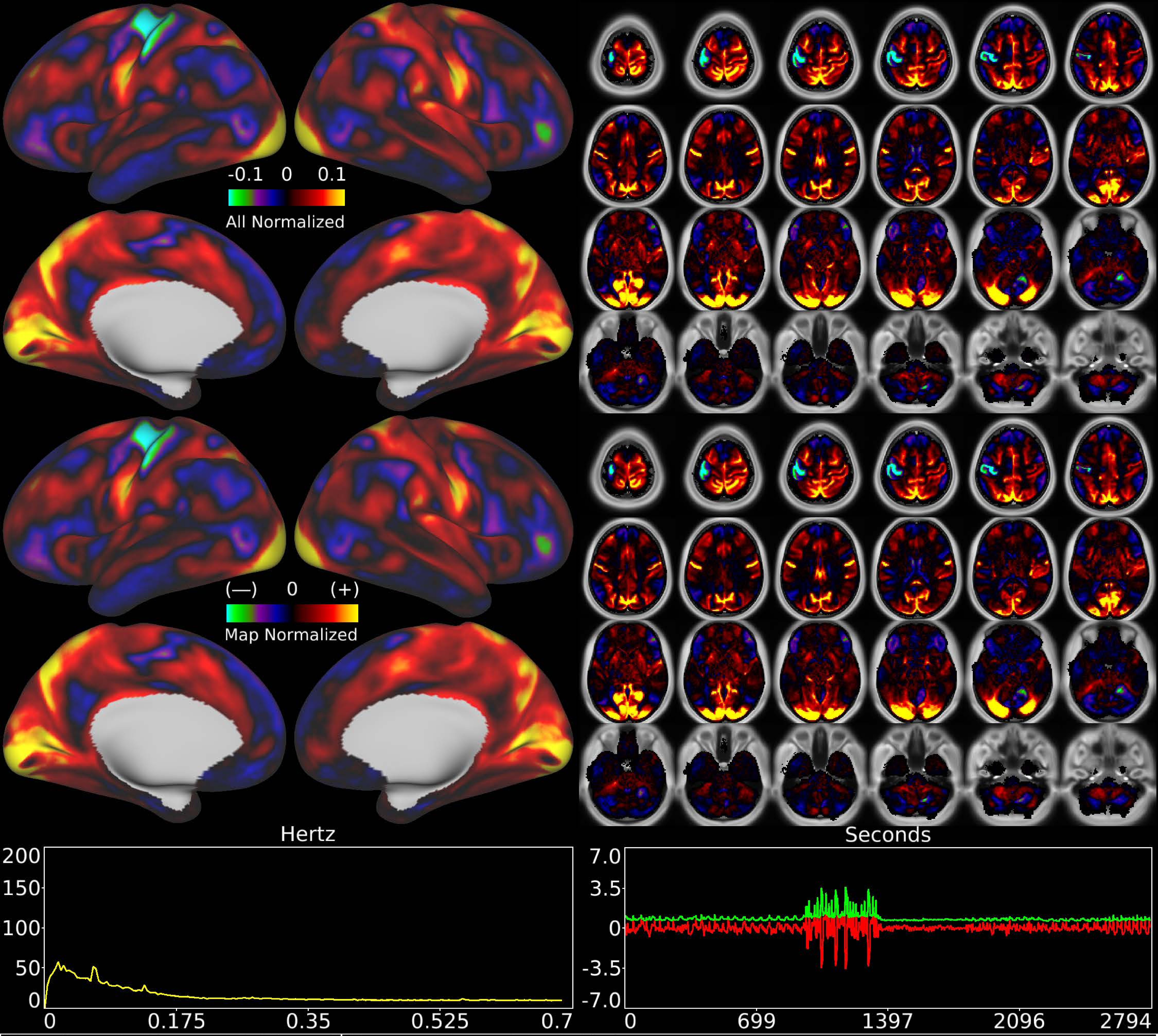


| Number & Class: 28 Signal | Name: Motor Task Unknown | |
|---|---|---|
| RVT Correlated: No | DVARS Dip Associated: No | Cross-Subject Variable: No |
| Single Subject: No | % Variance Explained: 1.36 | Globality Index: 1.16 |
| Rest Component: No | Taskr Component: No | Task Modulated: Motor |

Rationale: Spatial map includes positive and negative patches that are correlated with a specific task design

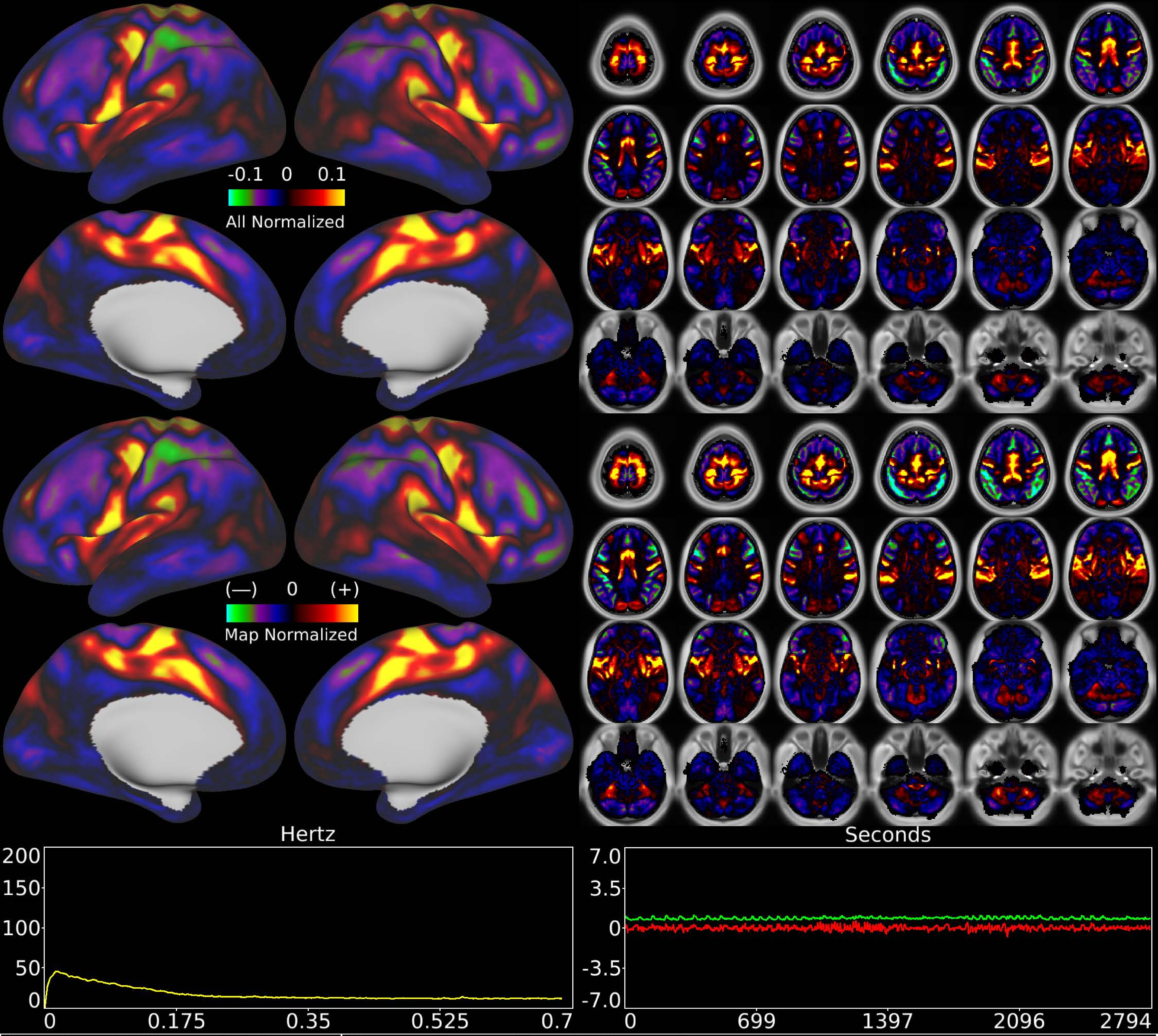


| | | |
|---|---|---|
| Number & Class: 29 Signal | Name: Eye + Trunk Motor Network | |
| RVT Correlated: No | DVARS Dip Associated: Yes | Cross-Subject Variable: No |
| Single Subject: No | % Variance Explained: 1.28 | Globality Index: 0.55 |
| Rest Component: 39 | Taskr Component: 28 | Task Modulated: No |
| Rationale: Spatial map includes positive and negative patches that respect known somatotopic sensori-motor organization (Eye and Trunk) | | |

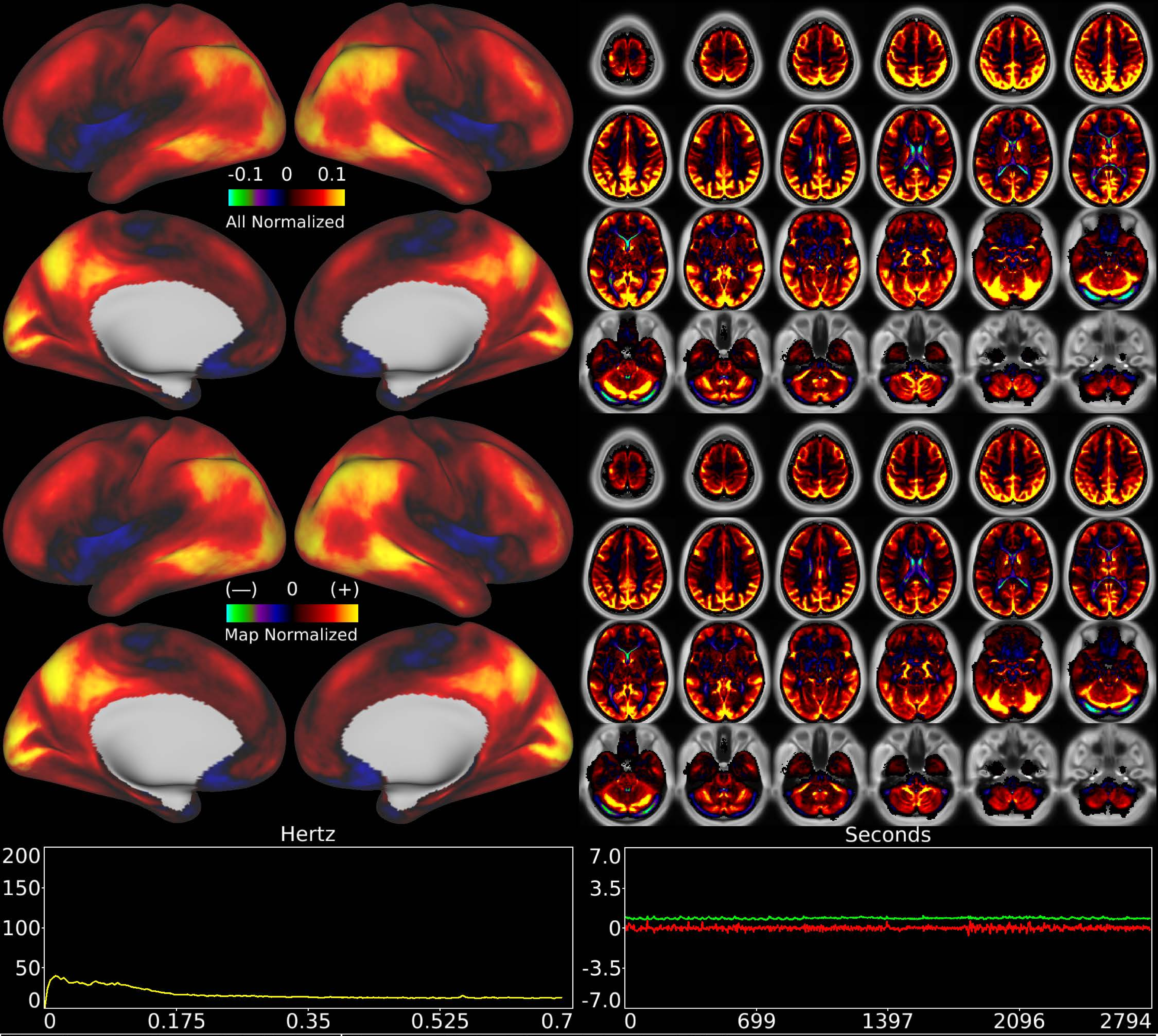


| | | |
|---|---|---|
| Number & Class: 30 Noise | Name: Global Physiological Noise | |
| RVT Correlated: Yes | DVARS Dip Associated: Yes | Cross-Subject Variable: Yes |
| Single Subject: Yes | % Variance Explained: 1.24 | Globality Index: 2.98 |
| Rest Component: No | Taskr Component: 31 | Task Modulated: No |
| Rationale: Globally positive component that is DVARS dips and RVT correlated and does not reflect known RSNs or areal boundaries | | |

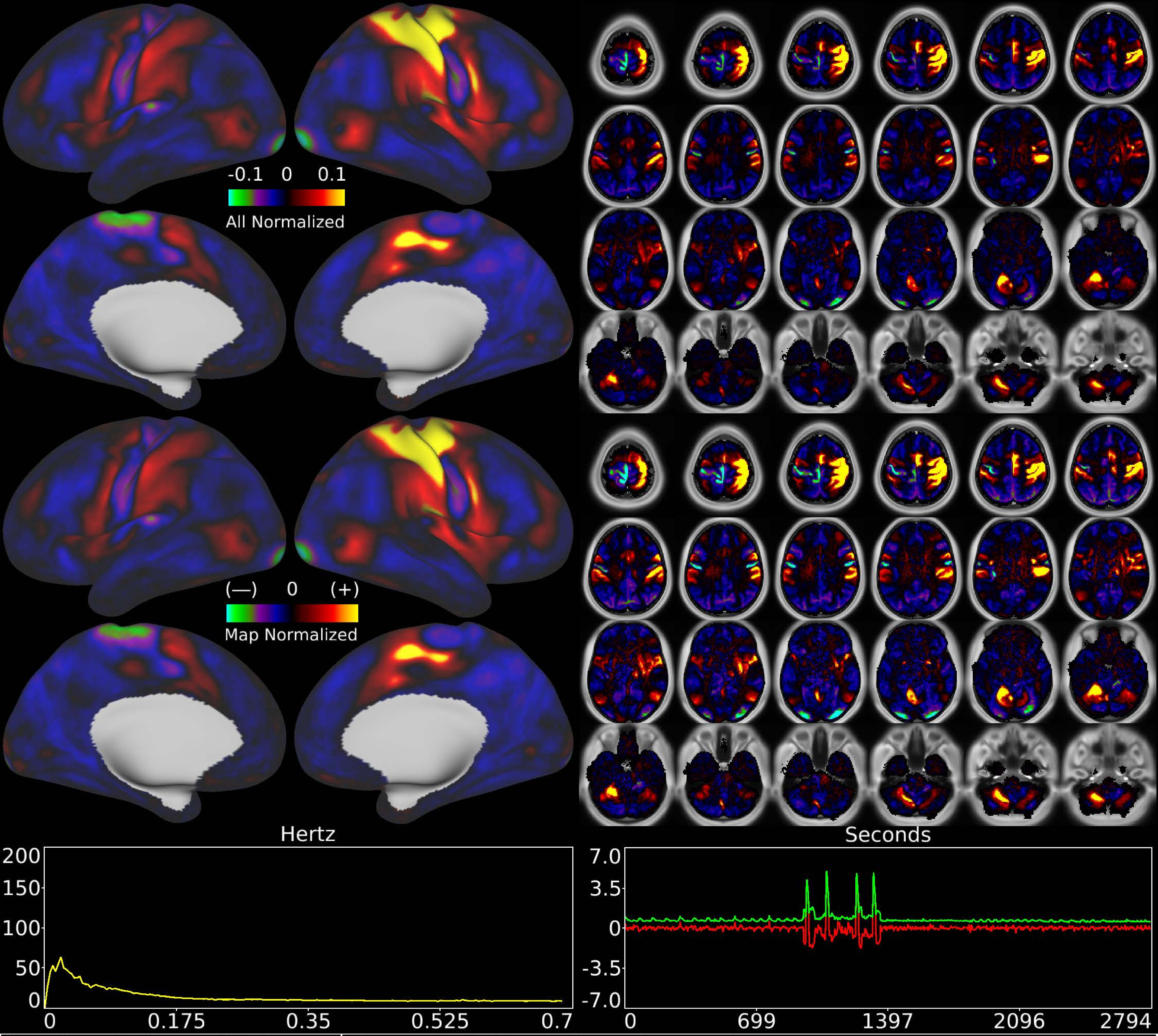


| | | |
|---|---|---|
| Number & Class: 31 Signal | Name: Left Hand Motor Network | |
| RVT Correlated: No | DVARS Dip Associated: Yes | Cross-Subject Variable: Yes |
| Single Subject: No | % Variance Explained: 1.25 | Globality Index: 0.9 |
| Rest Component: 40 | Taskr Component: 26 | Task Modulated: Motor |
| Rationale: Spatial map includes positive and negative patches that respect known somatotopic sensori-motor organization (Left Hand) | | |

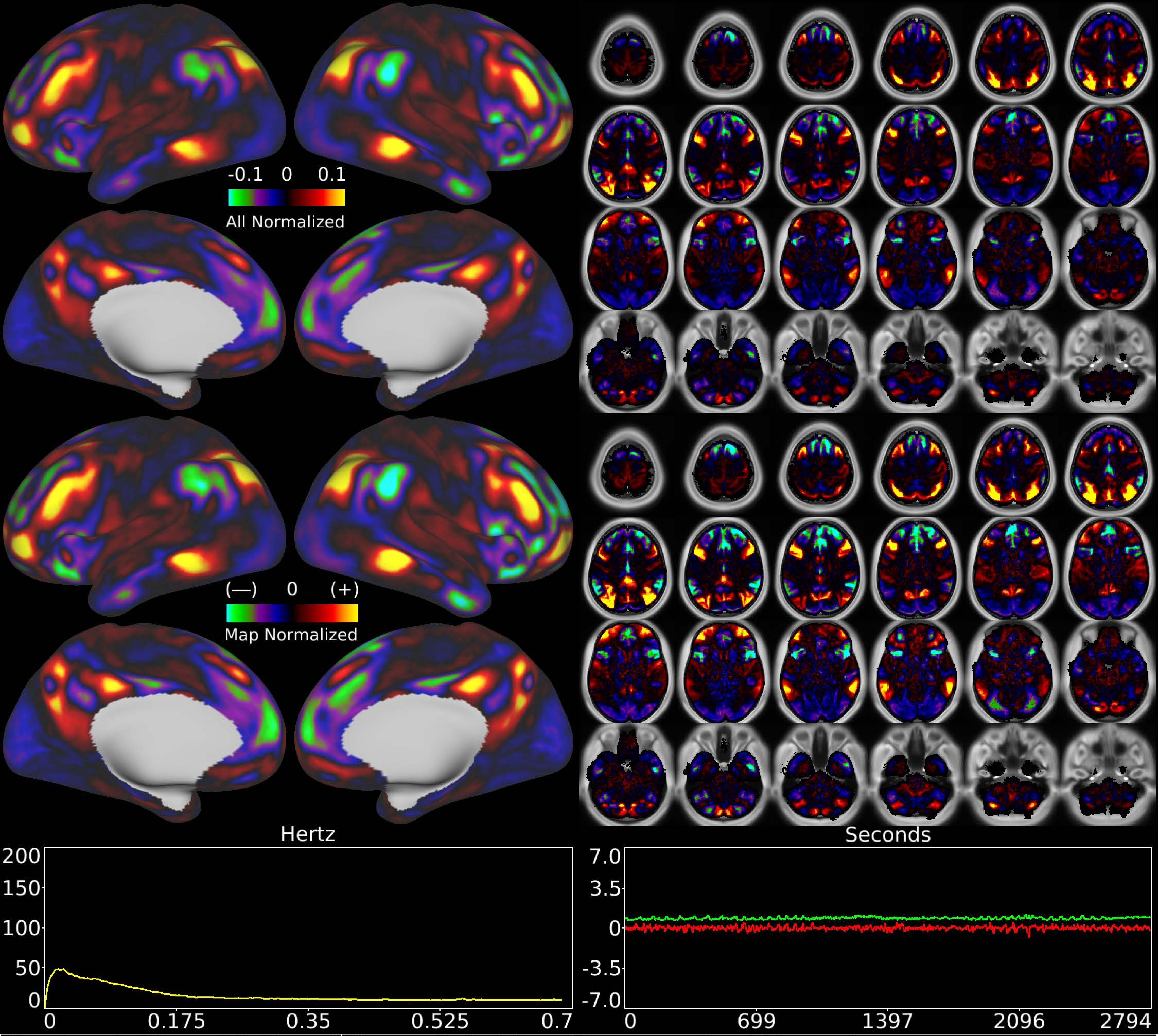


| | | |
|---|---|---|
| Number & Class: 32 Signal | Name: Subsidiary Fronto-parietal Network | |
| RVT Correlated: No | DVARS Dip Associated: No | Cross-Subject Variable: No |
| Single Subject: No | % Variance Explained: 1.25 | Globality Index: 0.11 |
| Rest Component: No | Taskr Component: No | Task Modulated: No |
| Rationale: Spatial map includes positive and negative patches that respect known RSNs (e.g. Fronto-Parietal Network) | | |

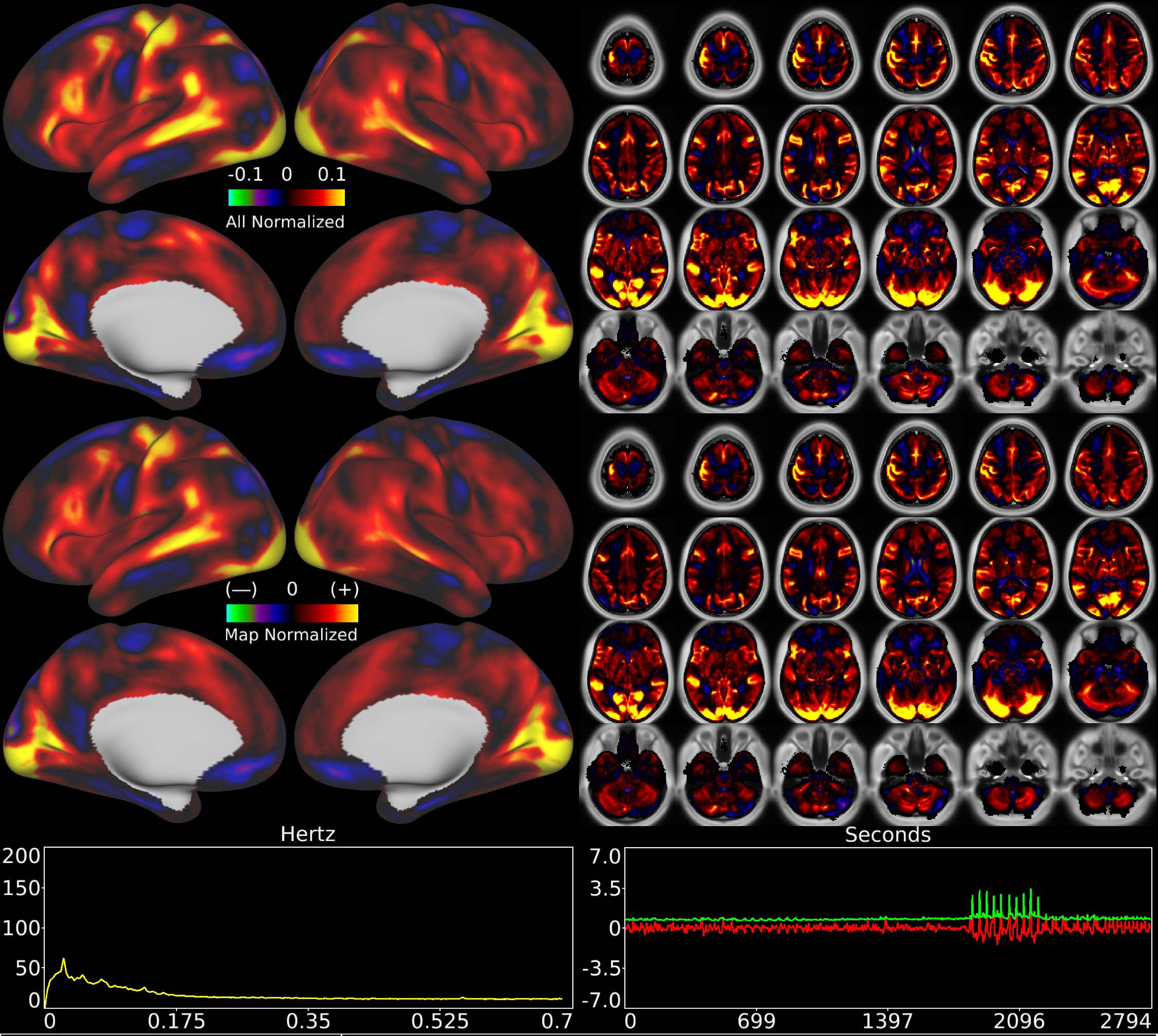


| | | |
|---|---|---|
| Number & Class: 33 Signal | Name: Unknown Network | |
| RVT Correlated: No | DVARS Dip Associated: No | Cross-Subject Variable: No |
| Single Subject: No | % Variance Explained: 1.24 | Globality Index: 2.19 |
| Rest Component: No | Taskr Component: No | Task Modulated: Social |

Rationale: Spatial map includes positive and negative patches that are correlated with a specific task design

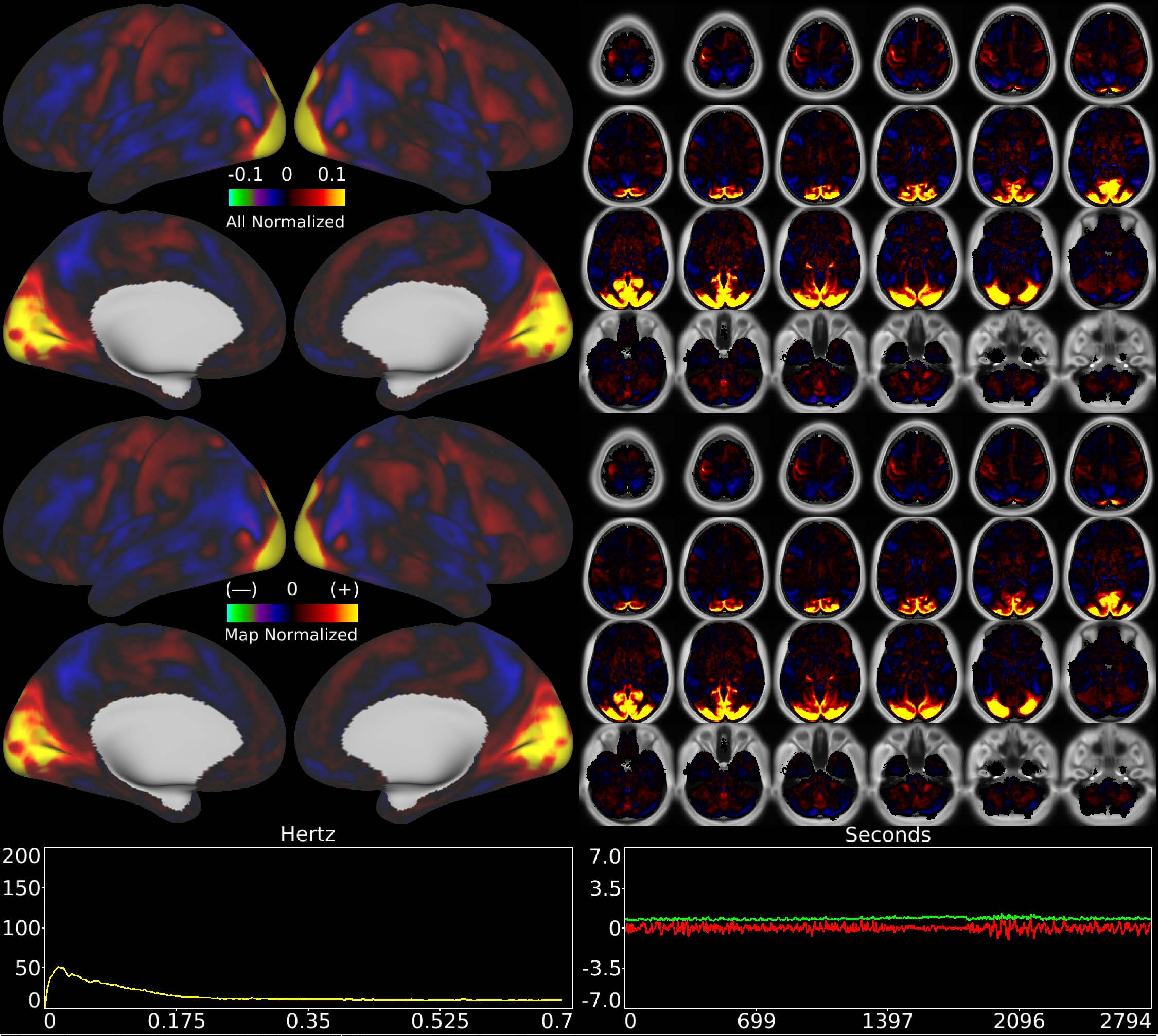


| | | |
|---|---|---|
| Number & Class: 34 Signal | Name: Early Visual | |
| RVT Correlated: No | DVARS Dip Associated: No | Cross-Subject Variable: No |
| Single Subject: No | % Variance Explained: 1.18 | Globality Index: 0.64 |
| Rest Component: 41 | Taskr Component: 15 | Task Modulated: No |
| Rationale: Spatial map includes positive and negative patches that respect known RSNs (e.g. Early Visual) | | |

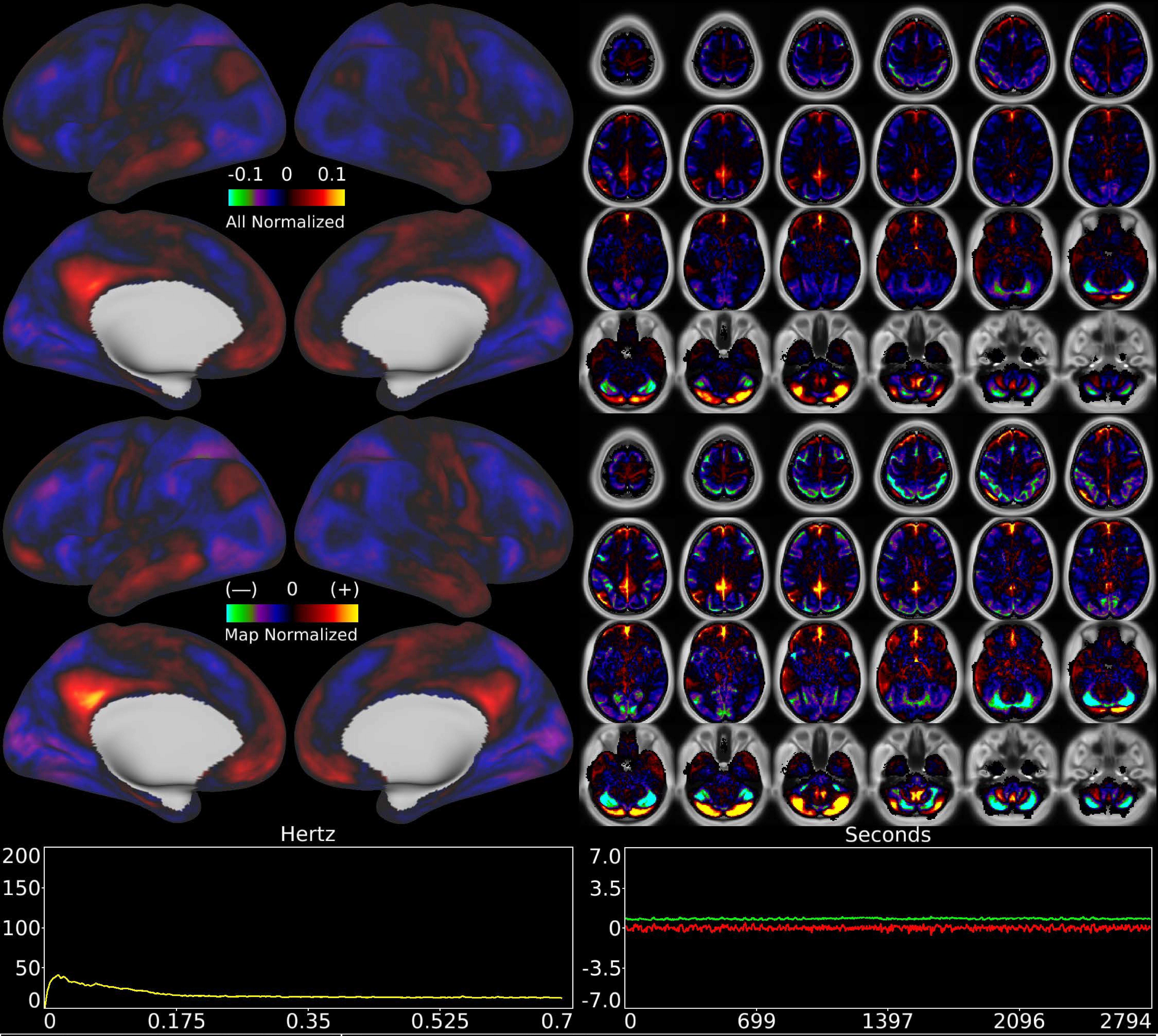


| | | |
|---|---|---|
| Number & Class: 35 Signal | Name: Cerebellar Unknown | |
| RVT Correlated: No | DVARS Dip Associated: No | Cross-Subject Variable: No |
| Single Subject: No | % Variance Explained: 1.11 | Globality Index: 0.94 |
| Rest Component: No | Taskr Component: 25 | Task Modulated: No |

Rationale: Spatial map includes positive and negative patches that respect known RSN boundaries in the cerebellum

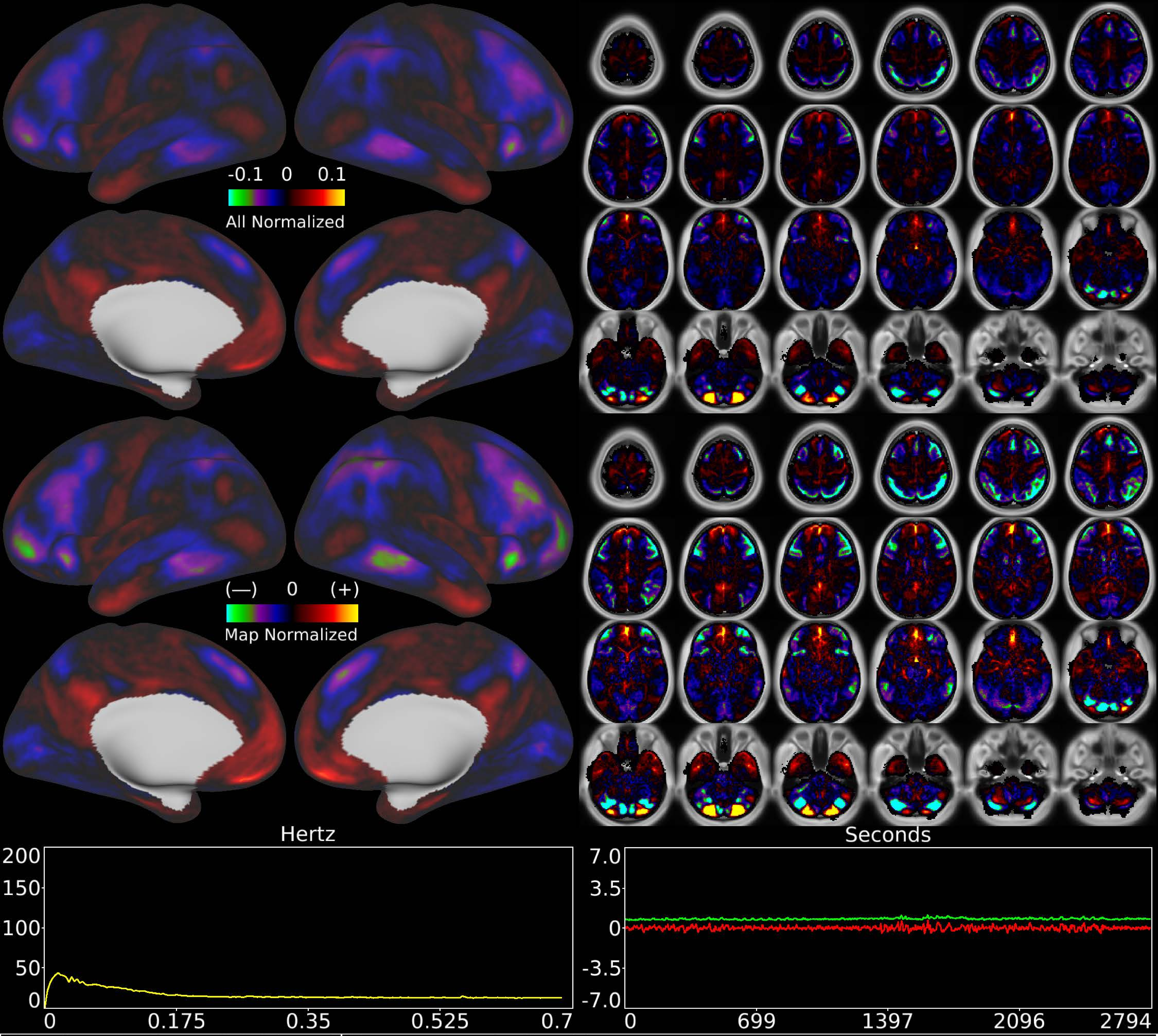


| | | |
|---|---|---|
| Number & Class: 36 Signal | Name: Cerebellar Unknown | |
| RVT Correlated: No | DVARS Dip Associated: No | Cross-Subject Variable: No |
| Single Subject: No | % Variance Explained: 1.08 | Globality Index: 0.56 |
| Rest Component: 20 | Taskr Component: -32 | Task Modulated: No |
| Rationale: Spatial map includes positive and negative patches that respect known RSN boundaries in the cerebellum | | |

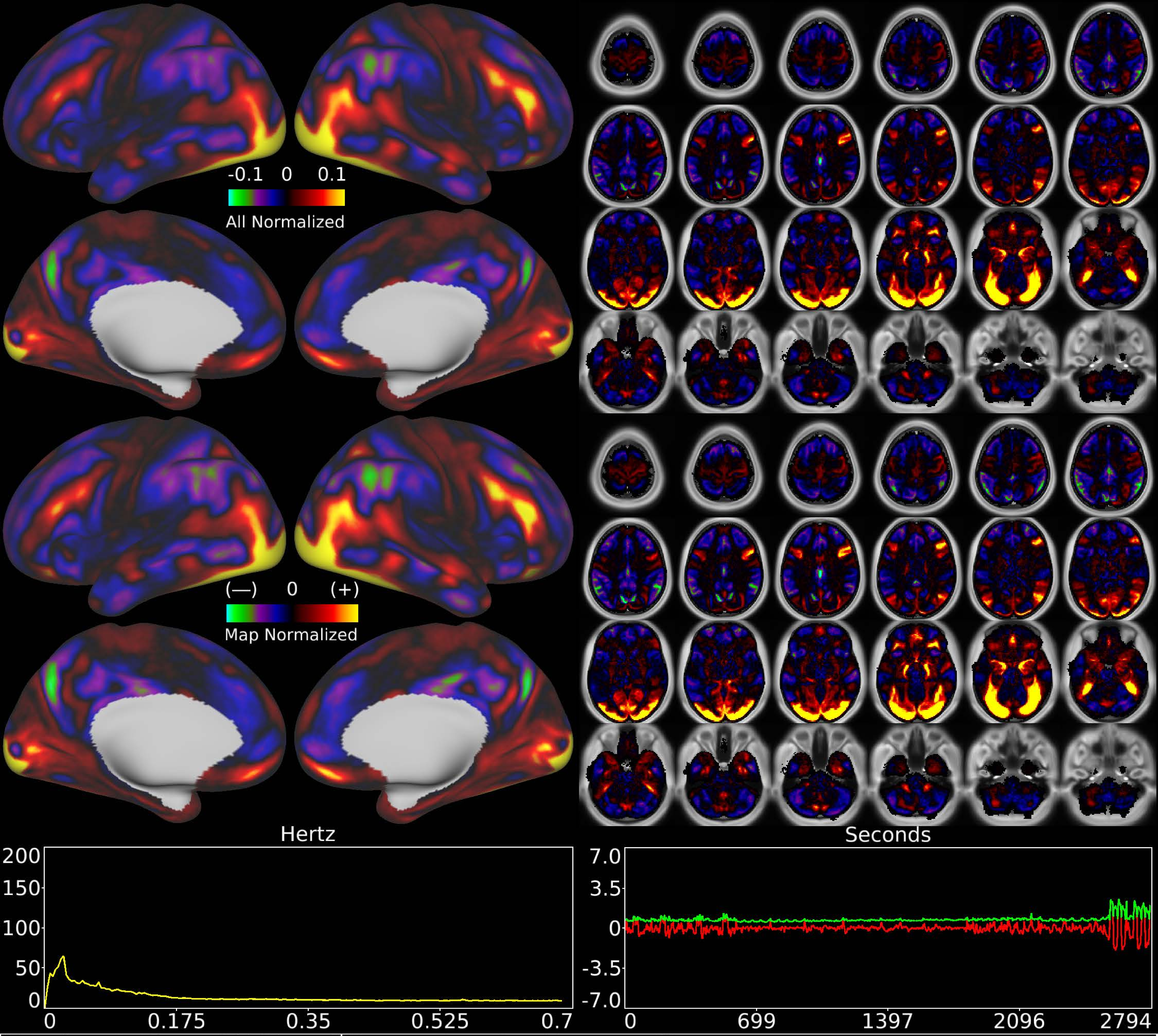


| | | |
|---|---|---|
| Number & Class: 37 Signal | Name: Emotion Faces > Shapes | |
| RVT Correlated: No | DVARS Dip Associated: No | Cross-Subject Variable: No |
| Single Subject: No | % Variance Explained: 1.06 | Globality Index: 0.16 |
| Rest Component: No | Taskr Component: No | Task Modulated: Emotion |

Rationale: Spatial map includes positive and negative patches that are correlated with a specific task design

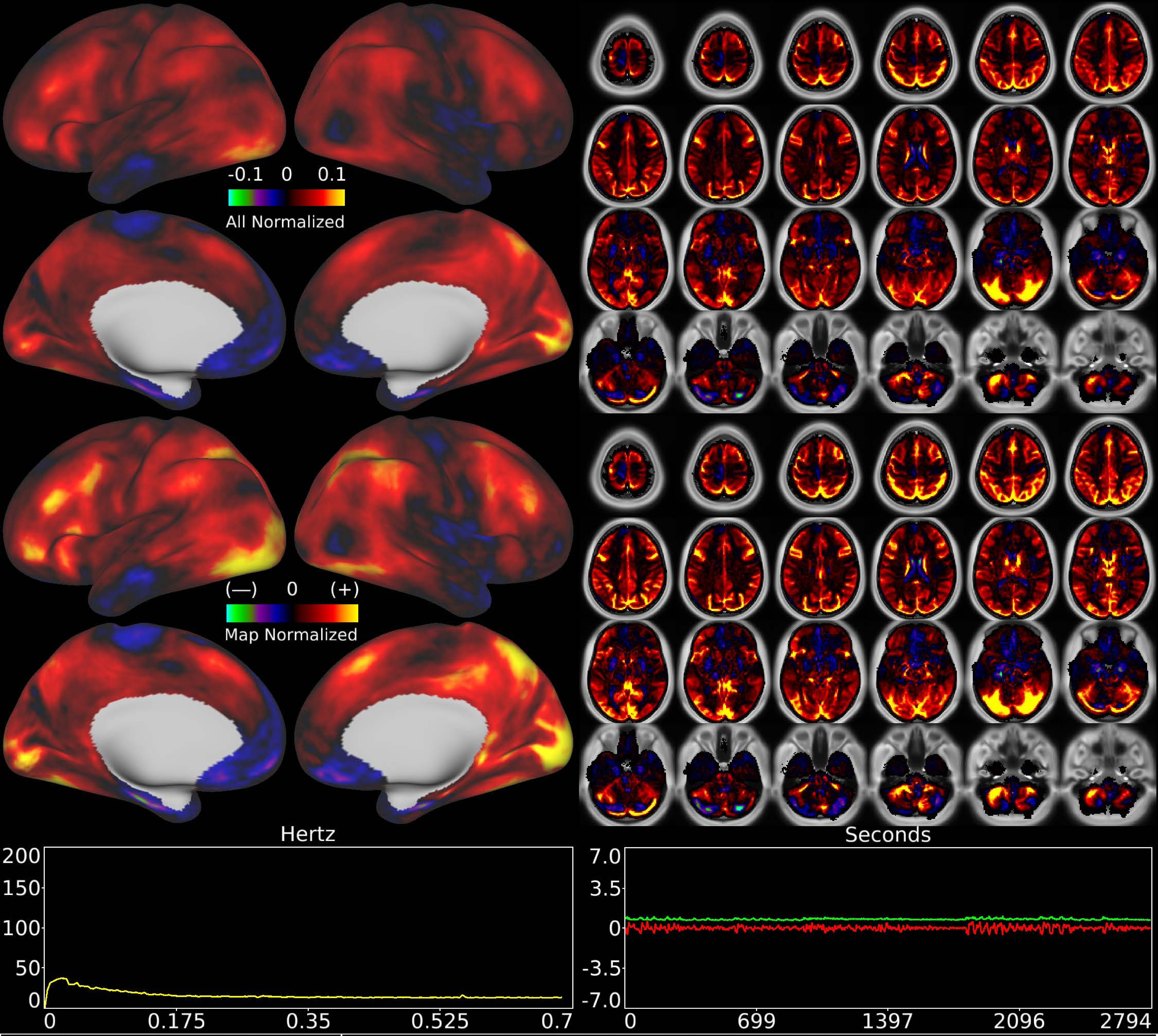


| | | |
|---|---|---|
| Number & Class: 38 Noise | Name: Global (Likely Physiological) Noise | |
| RVT Correlated: No | DVARS Dip Associated: No | Cross-Subject Variable: No |
| Single Subject: No | % Variance Explained: 1.02 | Globality Index: 2.03 |
| Rest Component: No | Taskr Component: No | Task Modulated: No |
| Rationale: Globally positive component without positive and negative patches that respect known RSNs or areas; some white matter signal | | |

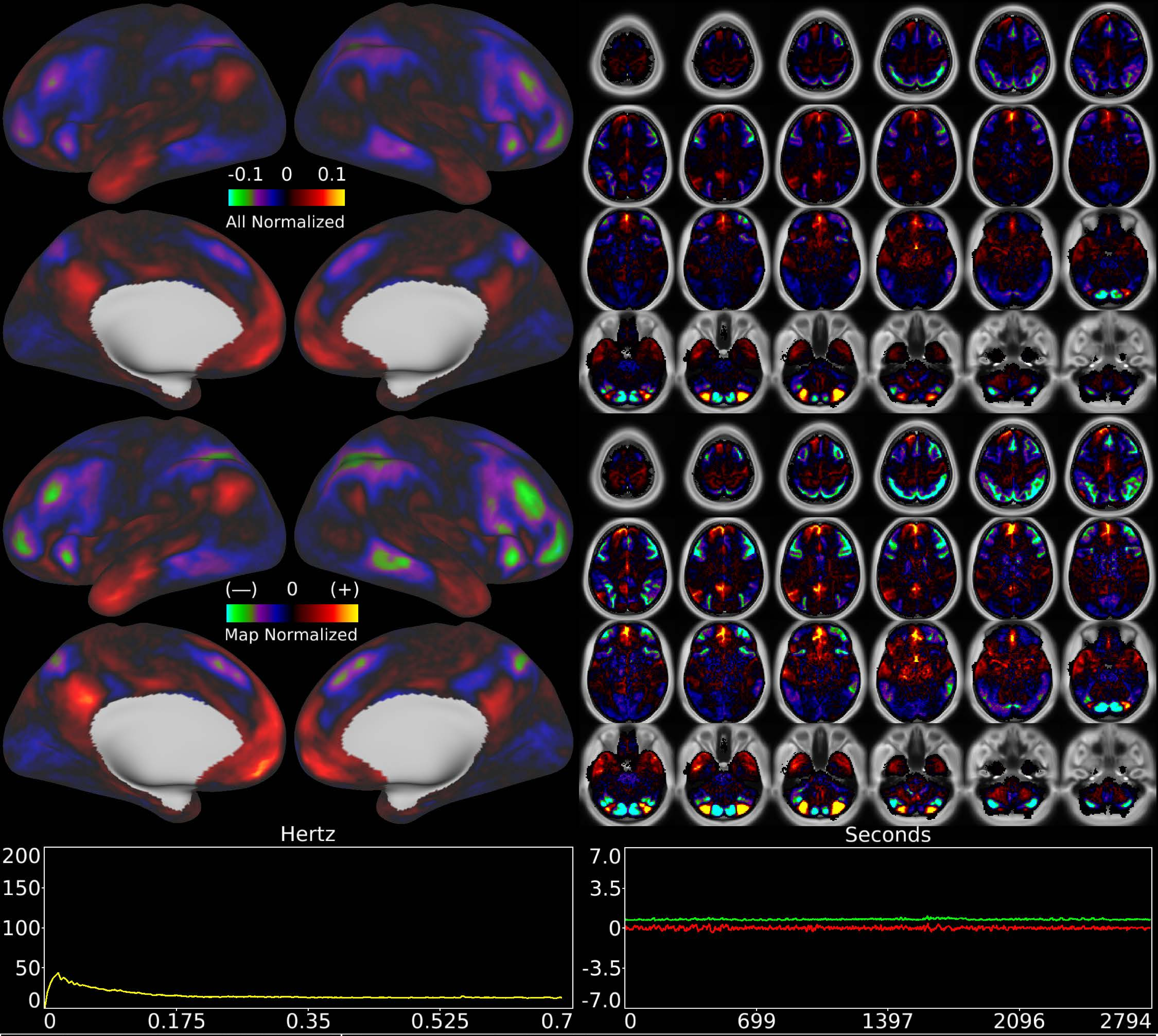


| | | |
|---|---|---|
| Number & Class: 39 Signal | Name: Cerebellar Unknown | |
| RVT Correlated: No | DVARS Dip Associated: No | Cross-Subject Variable: No |
| Single Subject: No | % Variance Explained: 0.96 | Globality Index: 0.19 |
| Rest Component: No | Taskr Component: No | Task Modulated: No |
| Rationale: Spatial map includes positive and negative patches that respect known RSN boundaries in the cerebellum | | |

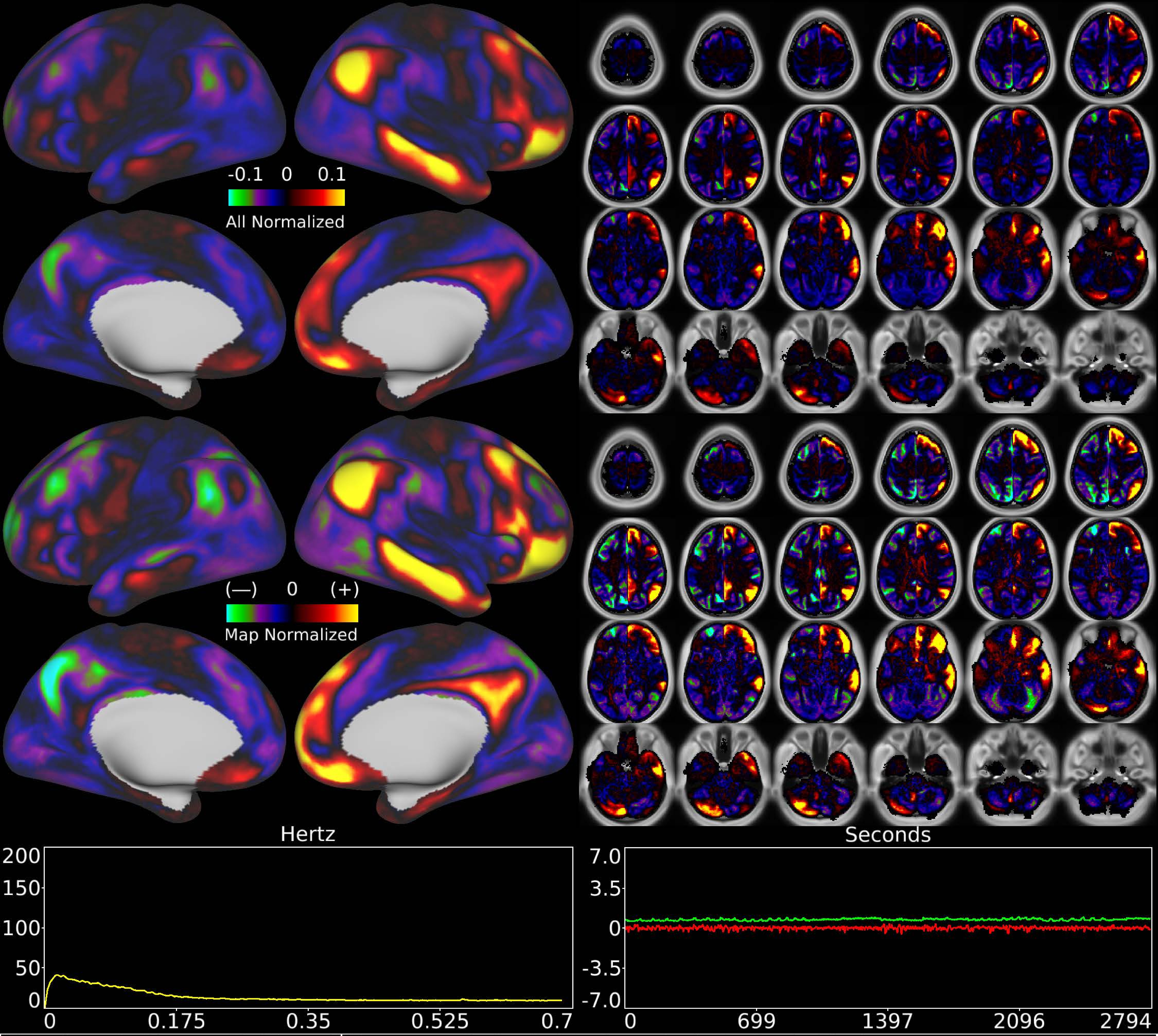


| | | |
|---|---|---|
| Number & Class: 40 Signal | Name: R Default Mode > Left Default Mode | |
| RVT Correlated: No | DVARS Dip Associated: No | Cross-Subject Variable: No |
| Single Subject: No | % Variance Explained: 0.93 | Globality Index: 1.13 |
| Rest Component: No | Taskr Component: No | Task Modulated: No |
| Rationale: Spatial map includes positive and negative patches that respect known RSNs (e.g. Default Mode Network) | | |

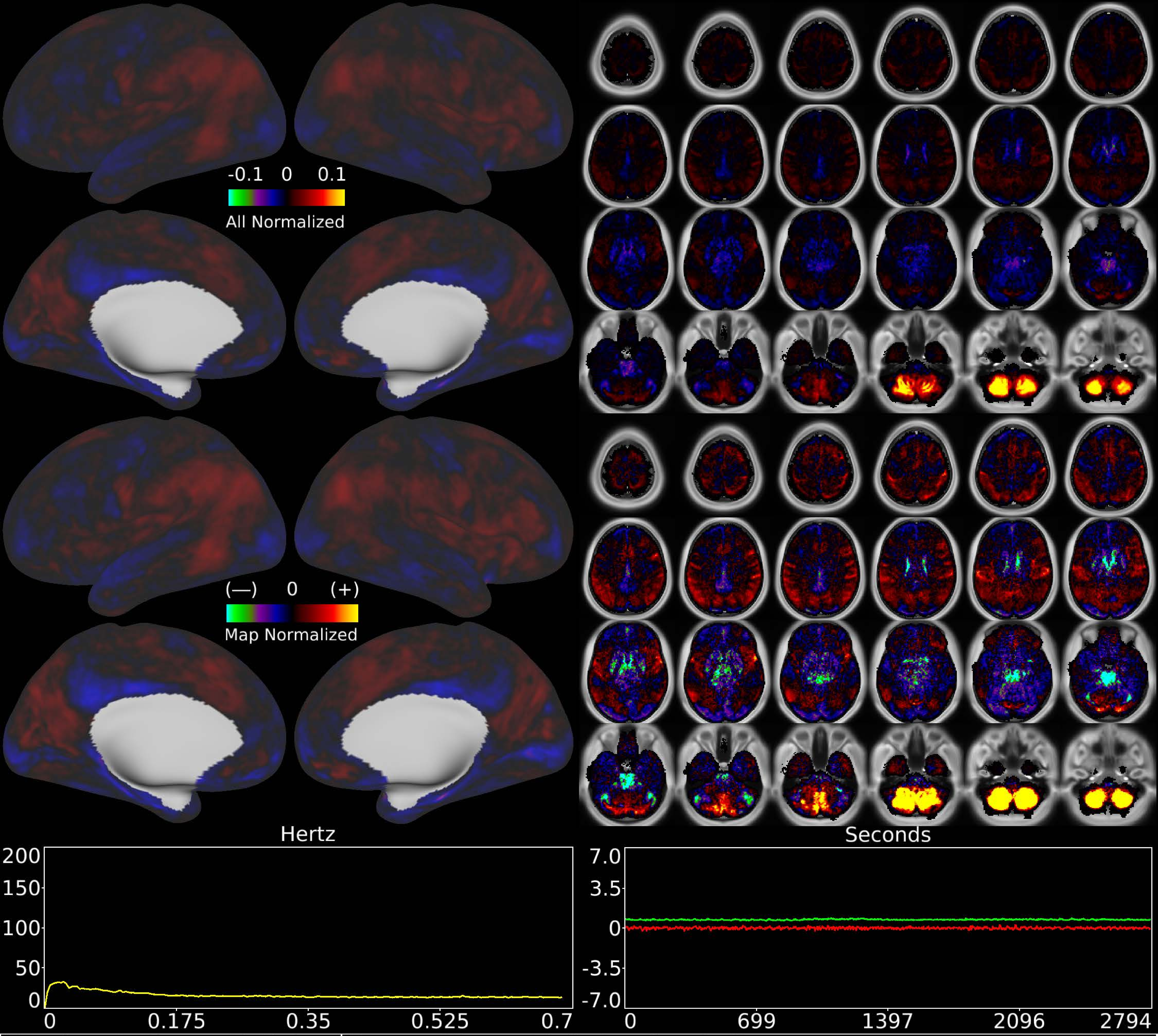


| | | |
|---|---|---|
| Number & Class: 41 Noise | Name: Bilateral Inferior Cerebellum | |
| RVT Correlated: No | DVARS Dip Associated: Yes | Cross-Subject Variable: Yes |
| Single Subject: Yes | % Variance Explained: 0.9 | Globality Index: 0.09 |
| Rest Component: No | Taskr Component: 33 | Task Modulated: No |

Rationale: Component is DVARS Dips associated and single subject; perhaps related to motion; reconstruction artifact; or unstructured noise

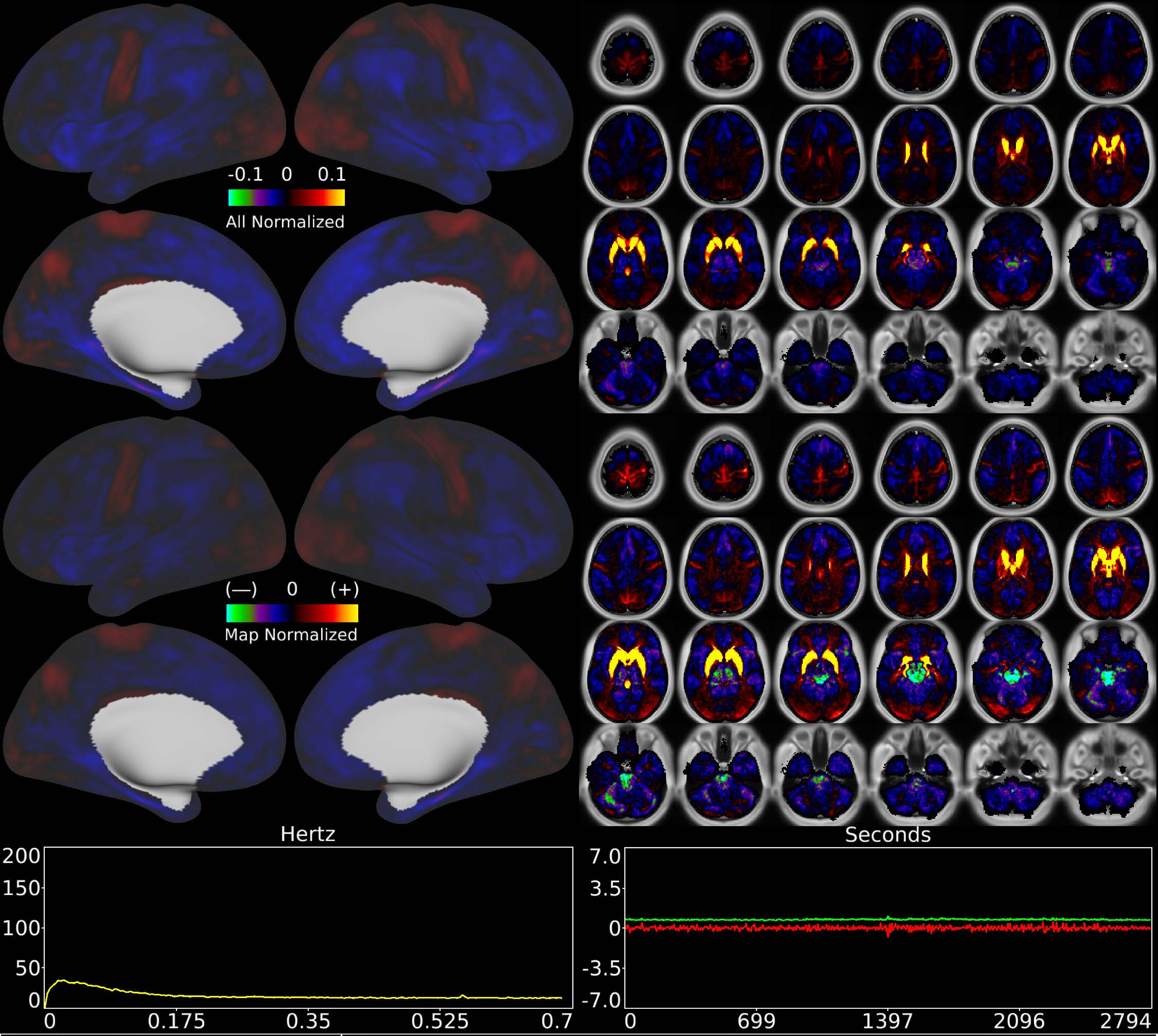


| | | |
|---|---|---|
| Number & Class: 42 Noise | Name: Striatum and Dorsal + Anterior Thalamus + WM | |
| RVT Correlated: No | DVARS Dip Associated: No | Cross-Subject Variable: No |
| Single Subject: No | % Variance Explained: 0.91 | Globality Index: 1.48 |
| Rest Component: 44 | Taskr Component: 36 | Task Modulated: No |

Rationale: Controversial: Diencephalon together with surrounding white matter positive vs rest of brain negative; could be due to differing vascular supplies

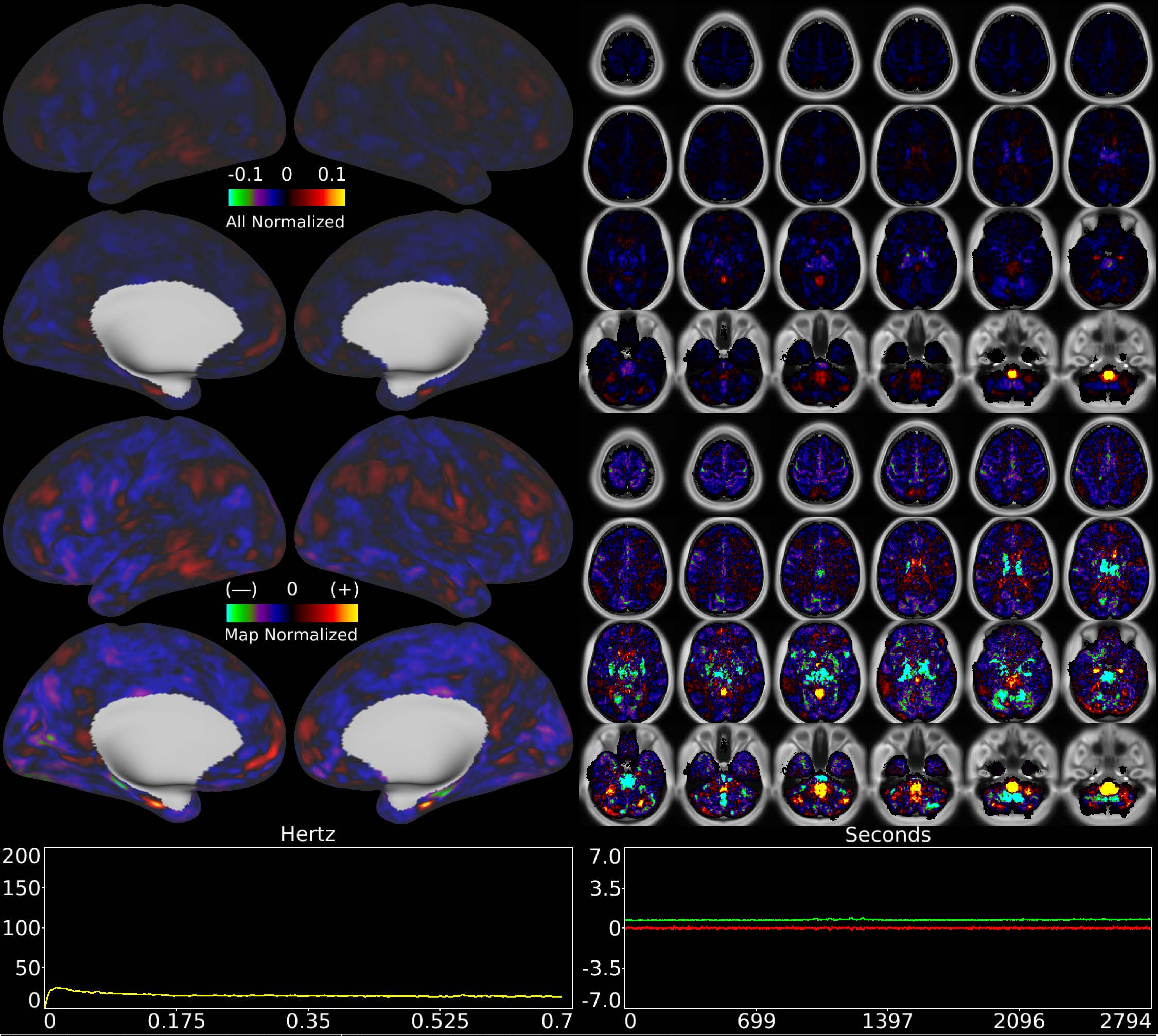


| | | |
|---|---|---|
| Number & Class: 43 Noise | Name: Medulla Recon Artifact | |
| RVT Correlated: No | DVARS Dip Associated: Yes | Cross-Subject Variable: Yes |
| Single Subject: No | % Variance Explained: 0.89 | Globality Index: 1.64 |
| Rest Component: 46 | Taskr Component: 38 | Task Modulated: No |
| Rationale: Spatial map not reflective of known areas or RSNs without connectivity other brain structures; some banding in sagittal plane suggestive of multi-band recon | | |

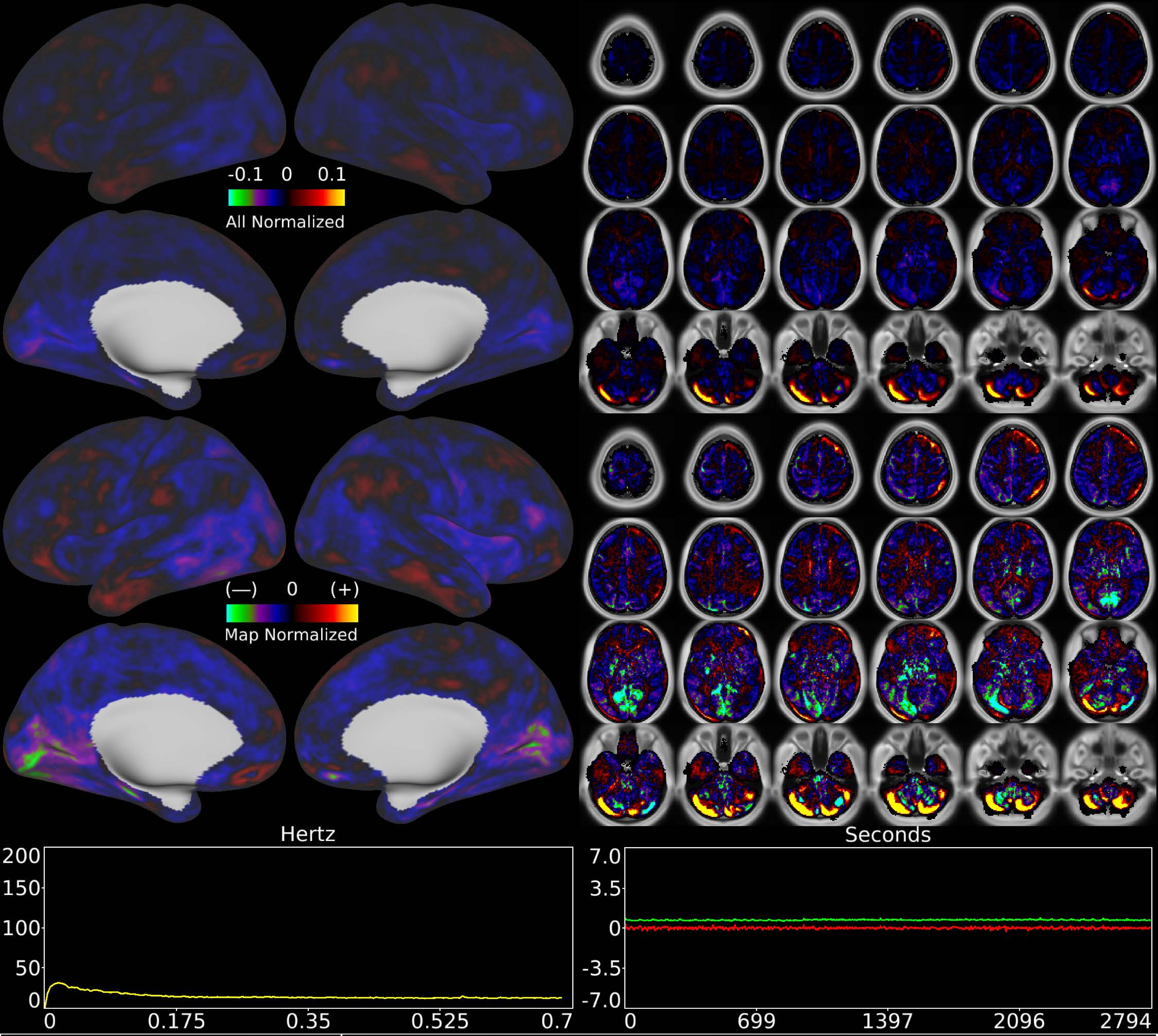


| | | |
|---|---|---|
| Number & Class: 44 Noise | Name: Cerebellar Movement Artifact Left | |
| RVT Correlated: No | DVARS Dip Associated: Yes | Cross-Subject Variable: Yes |
| Single Subject: Yes | % Variance Explained: 0.86 | Globality Index: 1.64 |
| Rest Component: 57 | Taskr Component: 34 | Task Modulated: No |

Rationale: Cerebellar edge motion component with high correlation to DVARS dips; looks like movement regressor beta map (derivative of X translation)

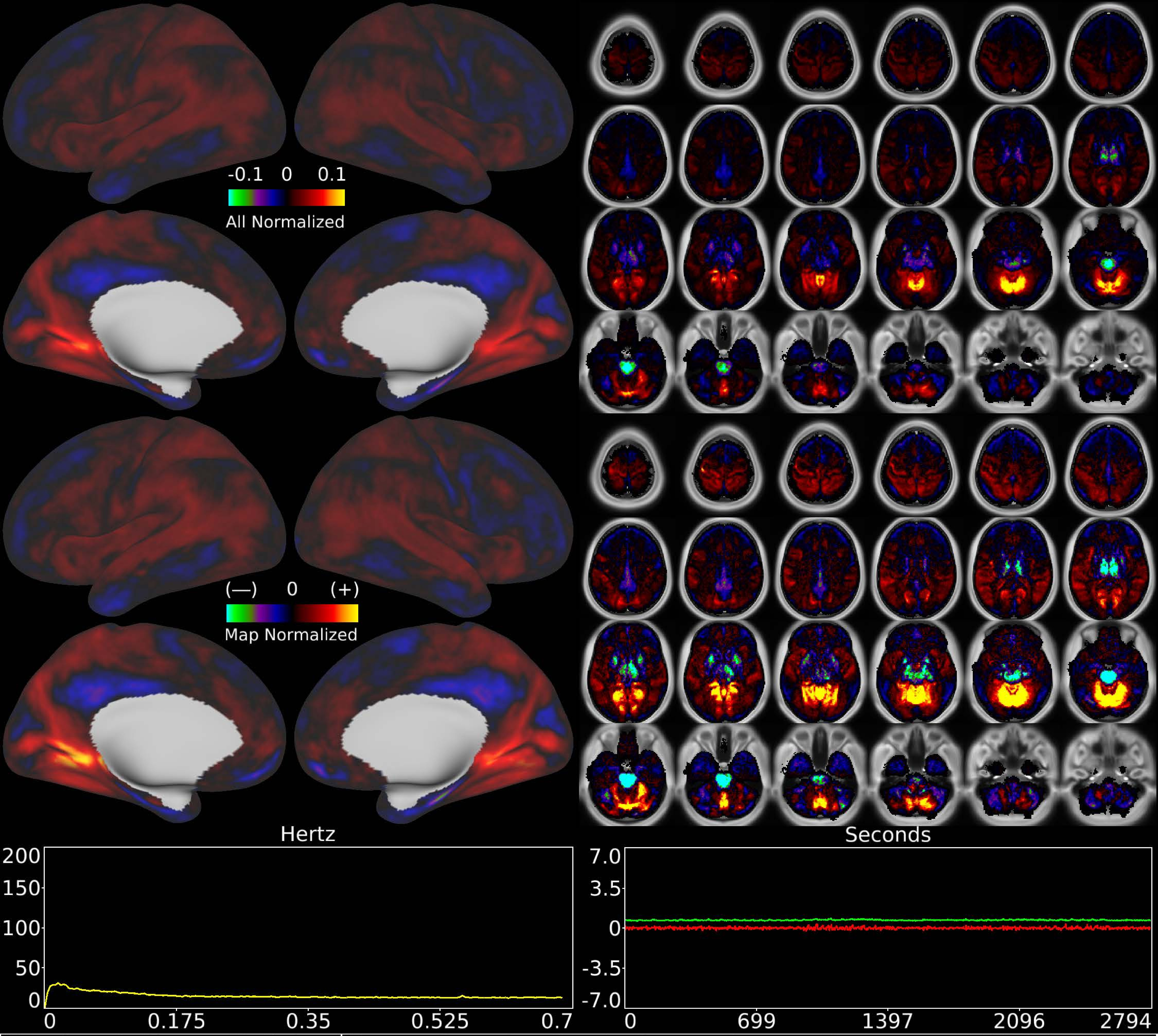


| | | |
|---|---|---|
| Number & Class: 45 Noise | Name: Superior Cerebellum > Pons + Thalamus DVARS Assoc | |
| RVT Correlated: No | DVARS Dip Associated: Yes | Cross-Subject Variable: No |
| Single Subject: No | % Variance Explained: 0.8 | Globality Index: 0.79 |
| Rest Component: No | Taskr Component: 41 | Task Modulated: No |

Rationale: Spatial map not reflective of known areas or RSNs
DVARS dips associated

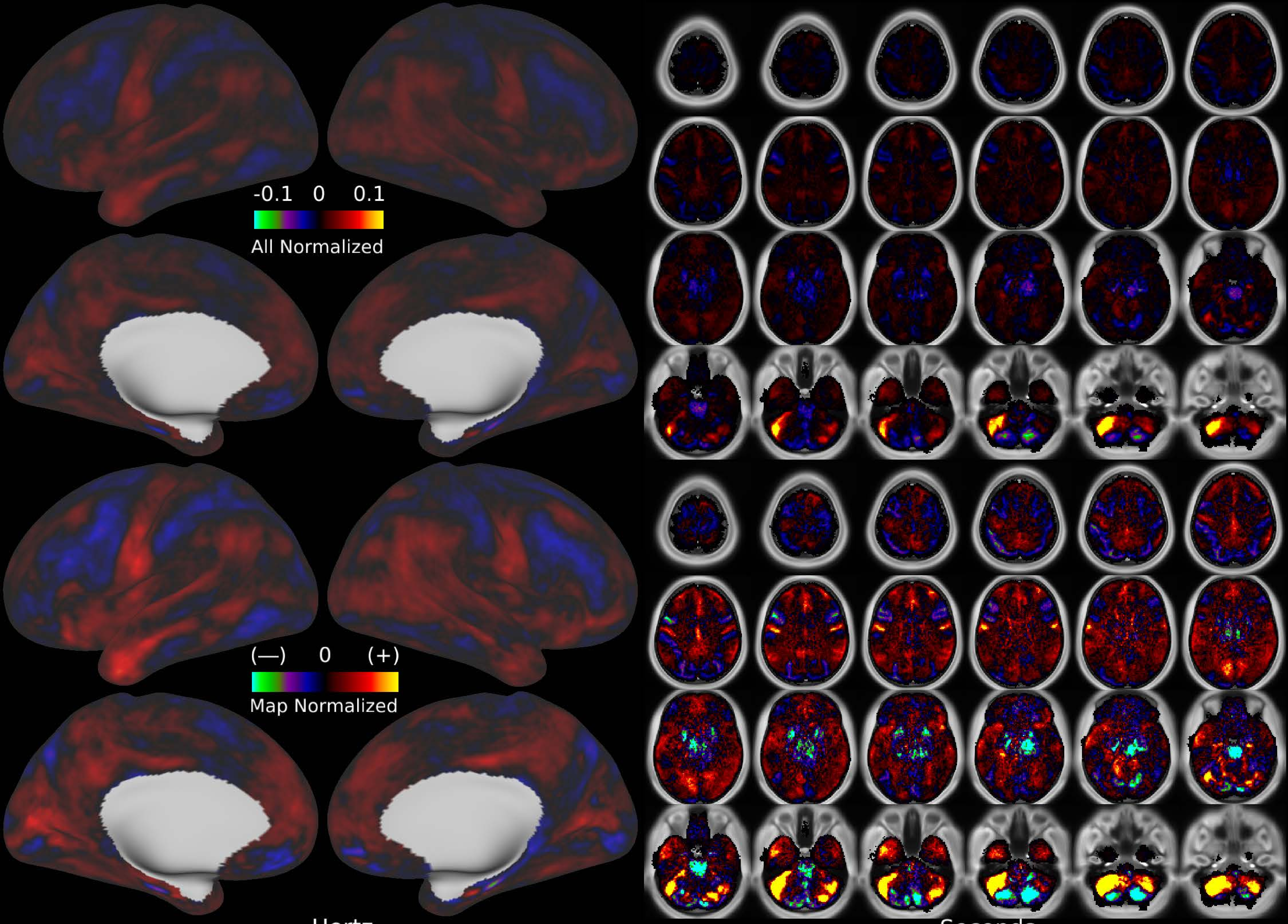


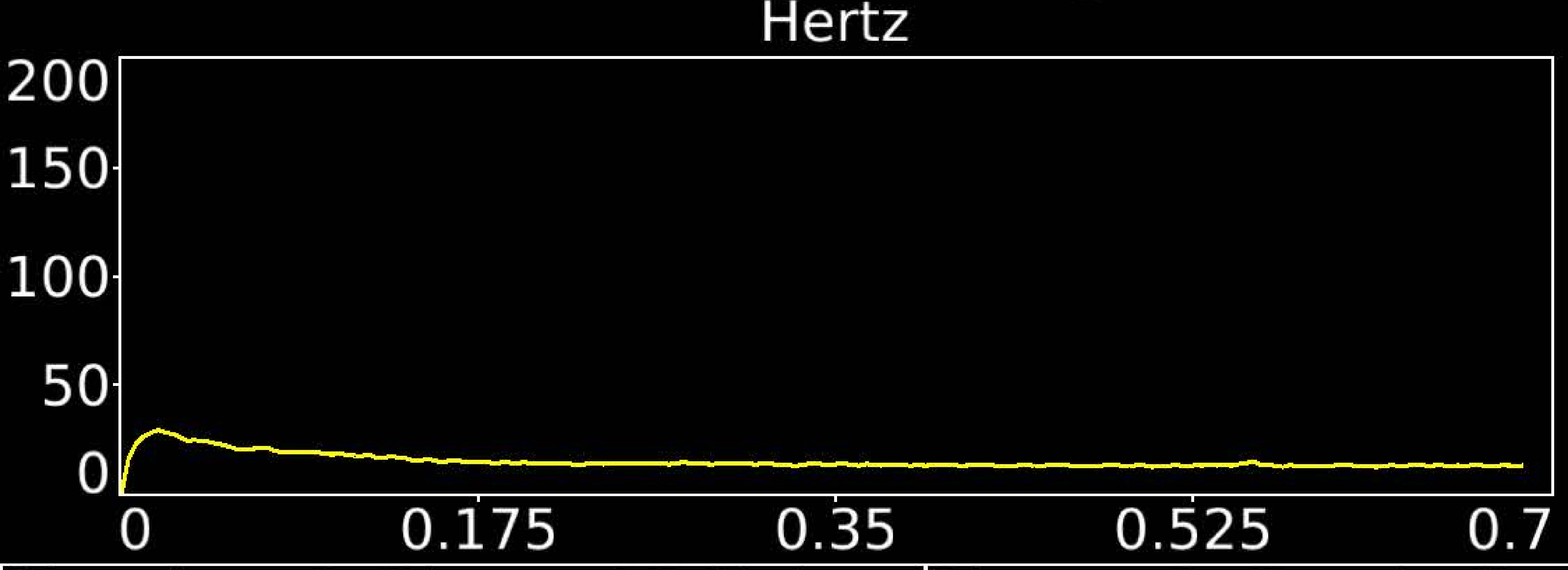


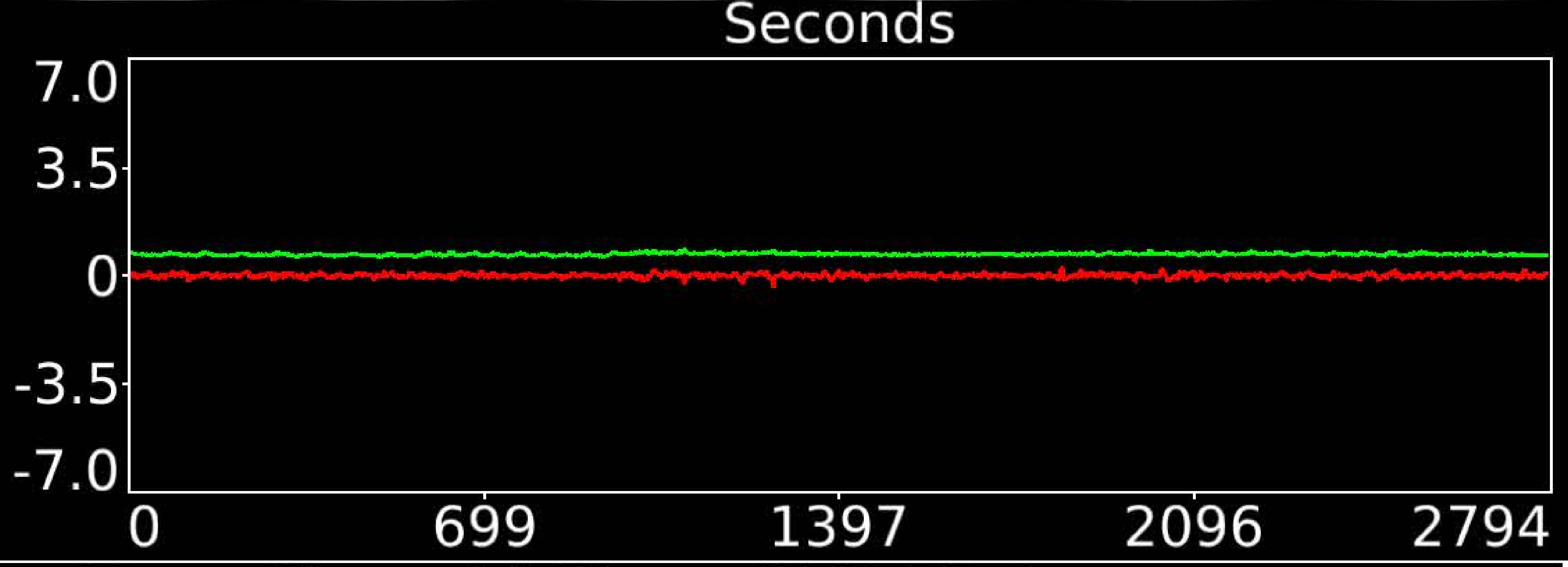


| | | |
|---|---|---|
| Number & Class: 46 Noise | Name: L Cerebellum Near Sigmoid Sinus DVARS Assoc | |
| RVT Correlated: No | DVARS Dip Associated: Yes | Cross-Subject Variable: No |
| Single Subject: Yes | % Variance Explained: 0.79 | Globality Index: 0.44 |
| Rest Component: No | Taskr Component: 42 | Task Modulated: No |
| Rationale: Cerebellar edge motion component with high correlation to DVARS dips; likely motion related | | |

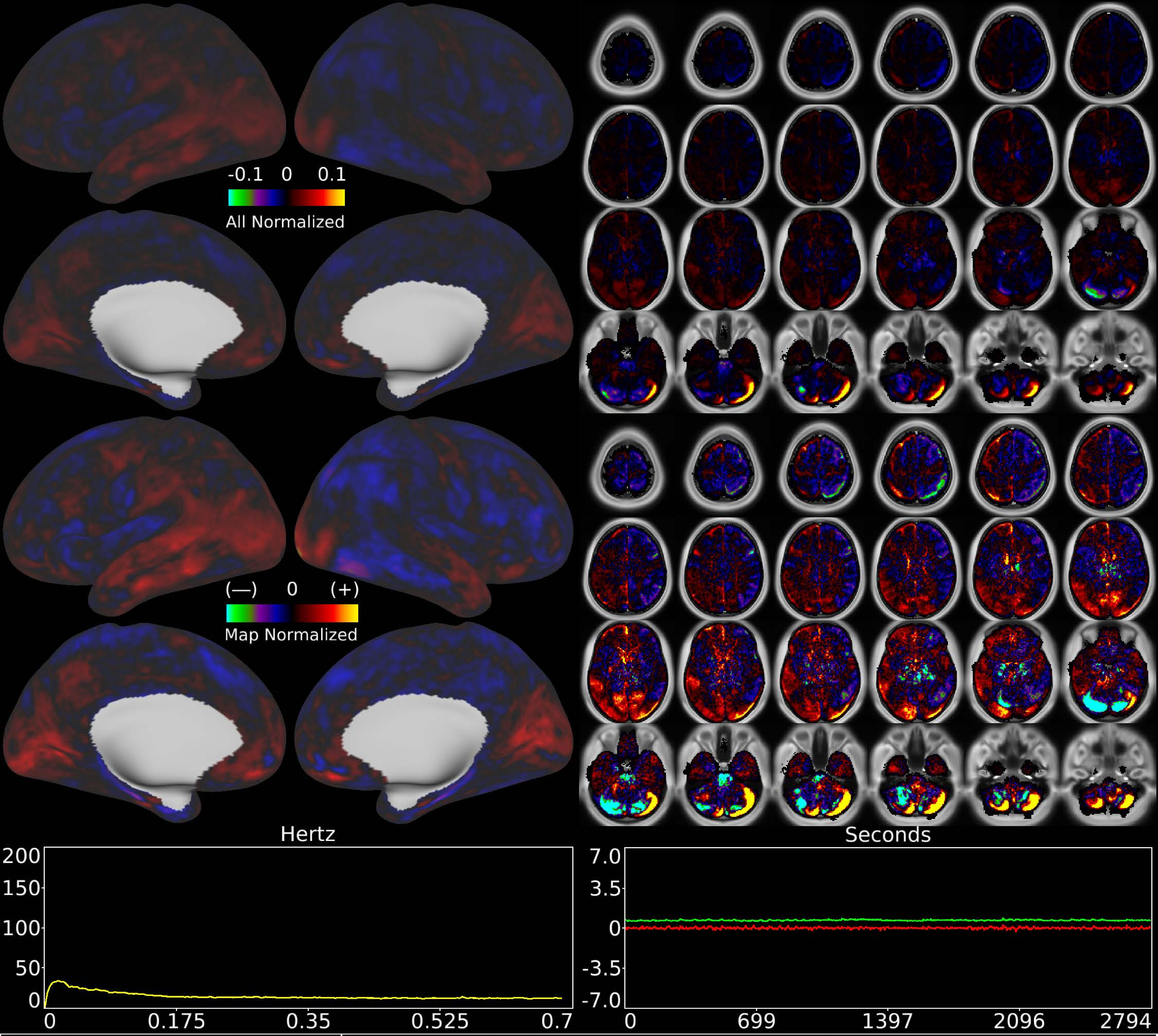


| | | |
|---|---|---|
| Number & Class: 47 Noise | Name: Cerebellar Movement Artifact Right | |
| RVT Correlated: No | DVARS Dip Associated: Yes | Cross-Subject Variable: No |
| Single Subject: Yes | % Variance Explained: 0.78 | Globality Index: 0.33 |
| Rest Component: No | Taskr Component: 45 | Task Modulated: No |
| Rationale: Cerebellar edge motion component with high correlation to DVARS dips; looks like movement regressor beta map (derivative of X translation) | | |

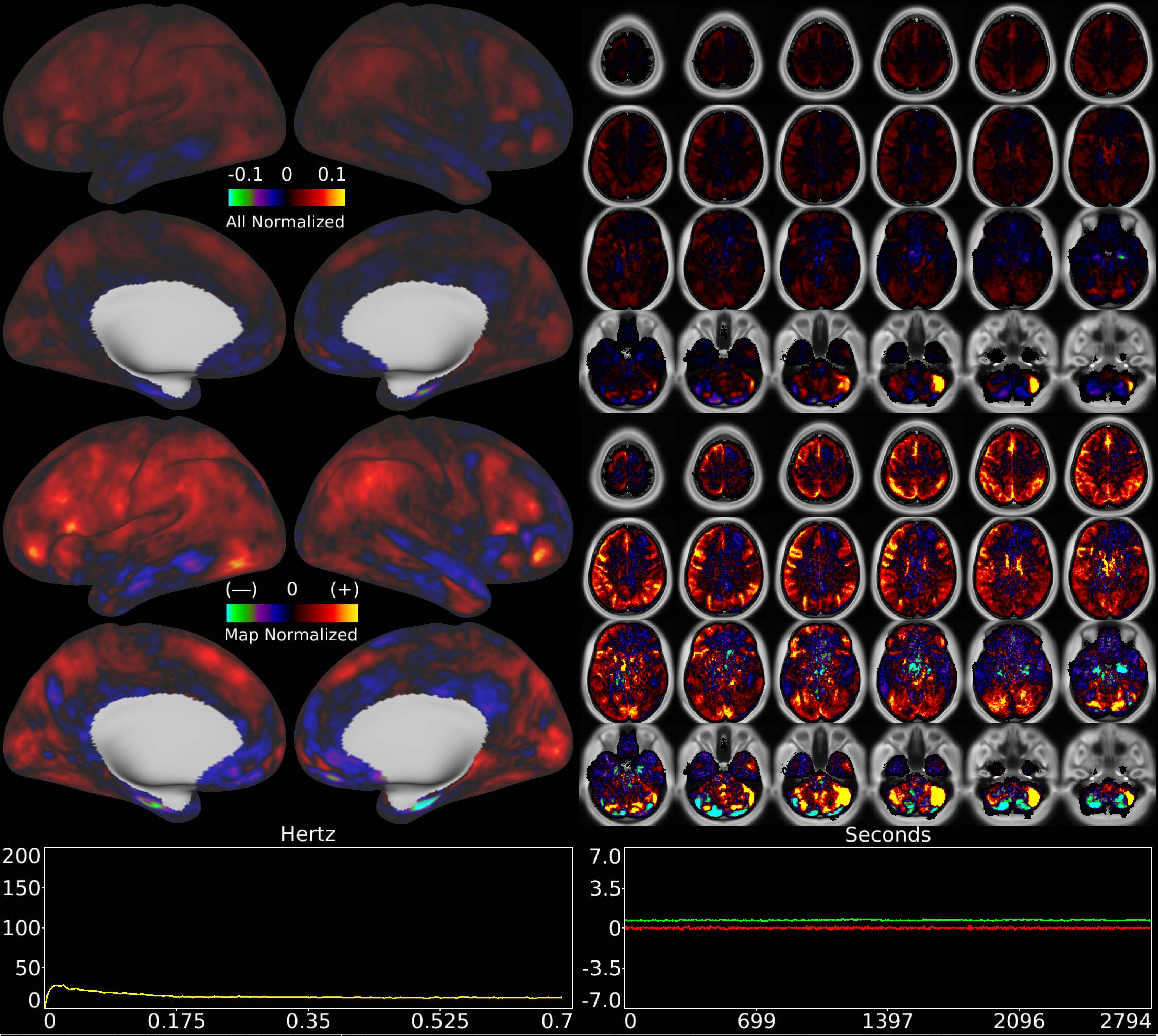


| | | |
|---|---|---|
| Number & Class: 48 Noise | Name: R Cerebellum Near Sigmoid Sinus DVARS Assoc | |
| RVT Correlated: No | DVARS Dip Associated: Yes | Cross-Subject Variable: No |
| Single Subject: Yes | % Variance Explained: 0.77 | Globality Index: 1.01 |
| Rest Component: No | Taskr Component: 52 | Task Modulated: No |
| Rationale: Cerebellar edge motion component with high correlation to DVARS dips; likely motion related | | |

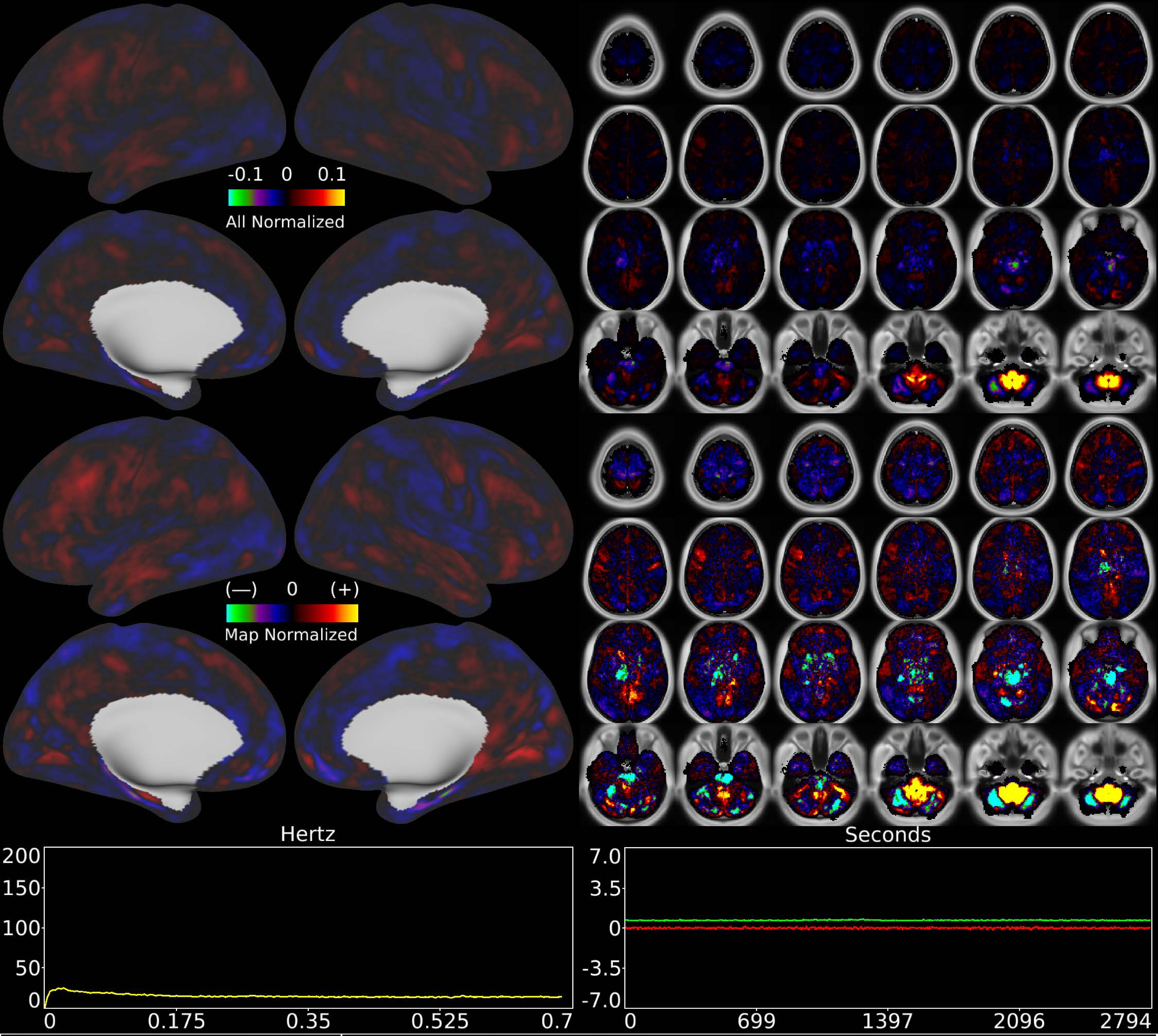


| | | |
|---|---|---|
| Number & Class: 49 Noise | Name: Cerebellar + Brainstem Recon Artifact | |
| RVT Correlated: No | DVARS Dip Associated: No | Cross-Subject Variable: No |
| Single Subject: Yes | % Variance Explained: 0.77 | Globality Index: 0.4 |
| Rest Component: No | Taskr Component: 43 | Task Modulated: No |

Rationale: Spatial map not reflective of known areas or RSNs without connectivity other brain structures; some banding in sagittal plane suggestive of multi-band recon

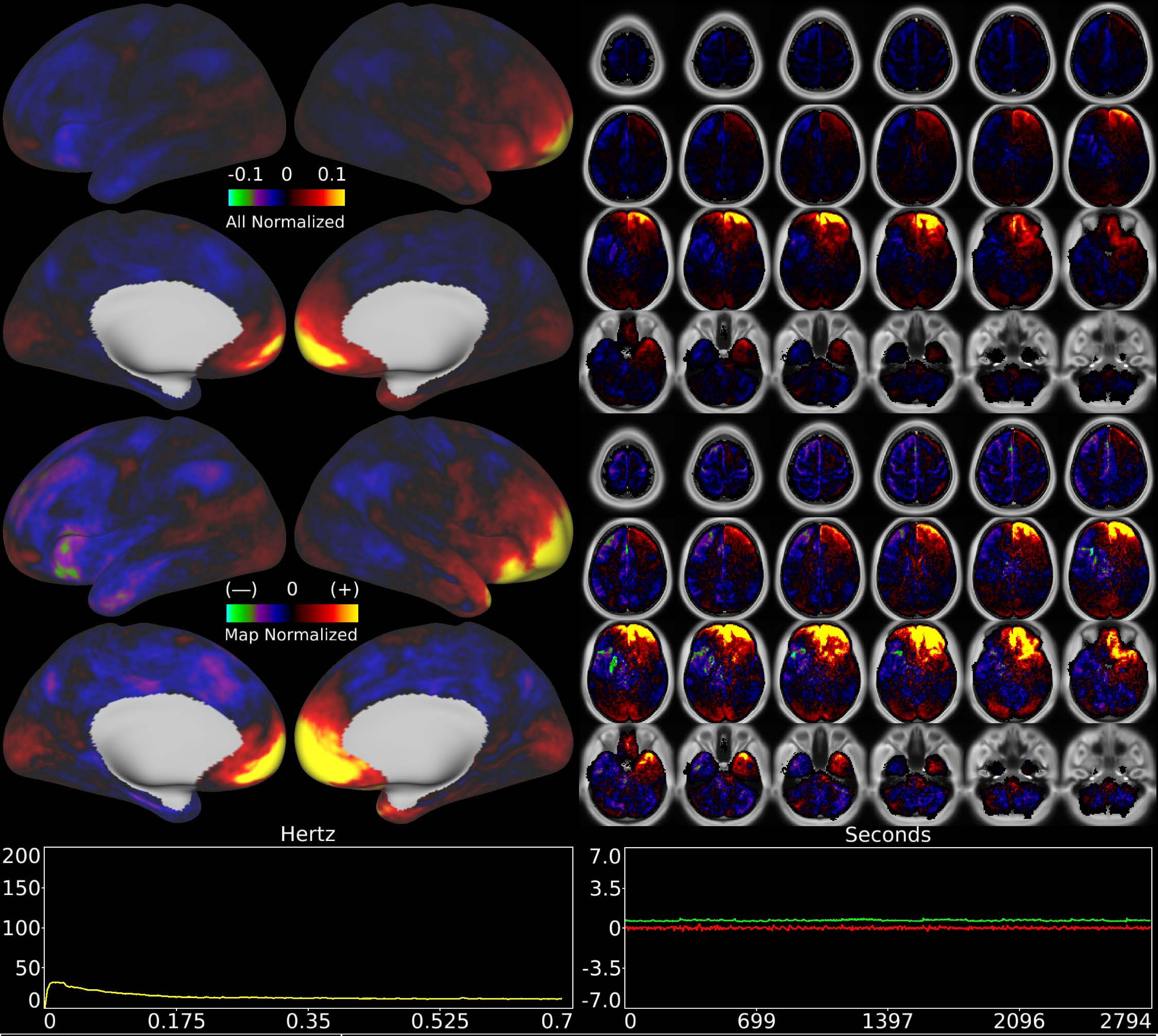


| | | |
|---|---|---|
| Number & Class: 50 Noise | Name: Coil | |
| RVT Correlated: No | DVARS Dip Associated: Yes | Cross-Subject Variable: Yes |
| Single Subject: No | % Variance Explained: 0.77 | Globality Index: 0.8 |
| Rest Component: 47 | Taskr Component: 50 | Task Modulated: No |
| Rationale: Known coil noise component | | |

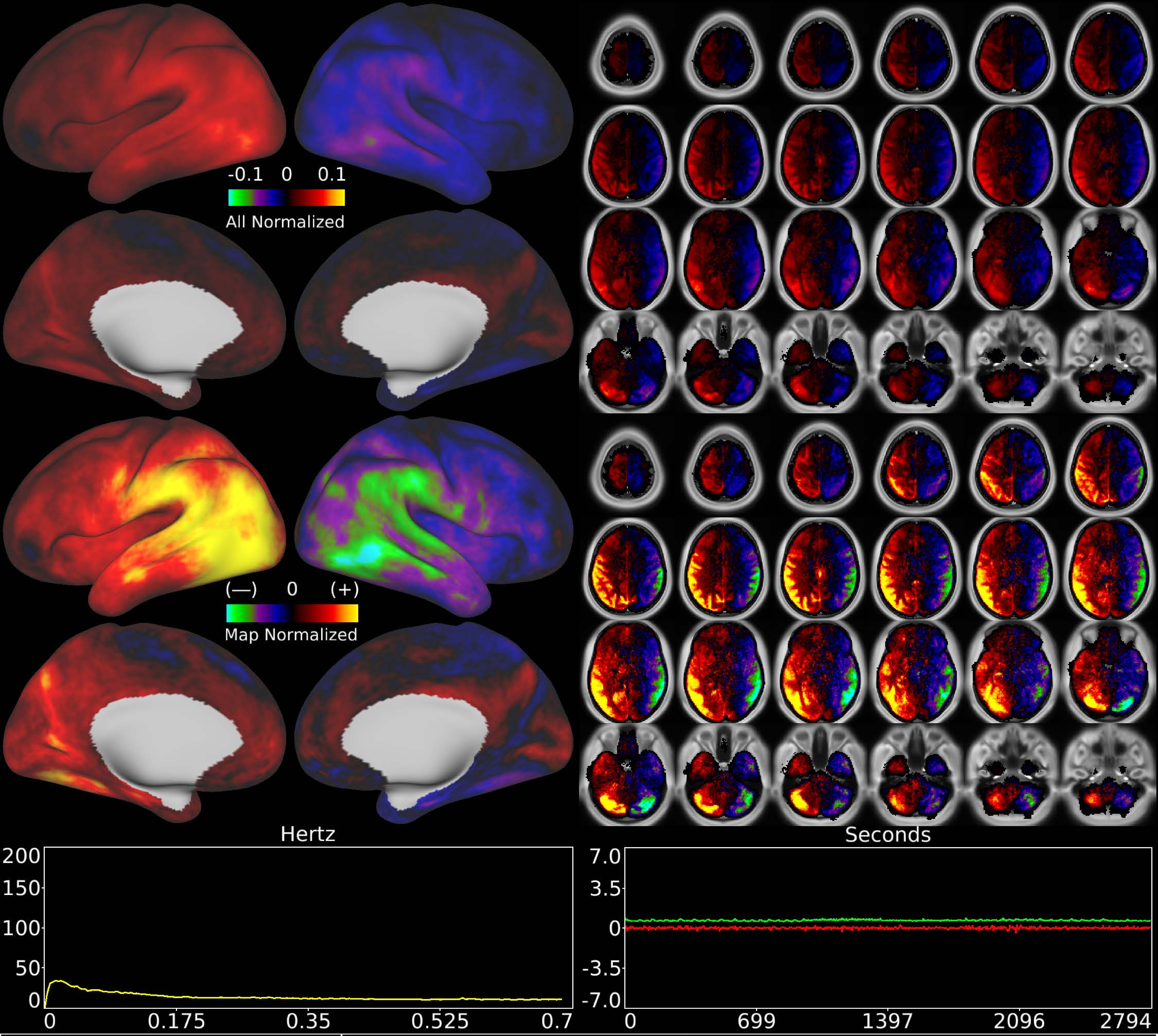


| | | |
|---|---|---|
| Number & Class: 51 Noise | Name: L>R Movement Artifact | |
| RVT Correlated: No | DVARS Dip Associated: Yes | Cross-Subject Variable: No |
| Single Subject: No | % Variance Explained: 0.75 | Globality Index: 0.5 |
| Rest Component: 50 | Taskr Component: 37 | Task Modulated: No |
| Rationale: Highly DVARS Dips associated component; looks like movement regressor beta map (Z rotation) | | |

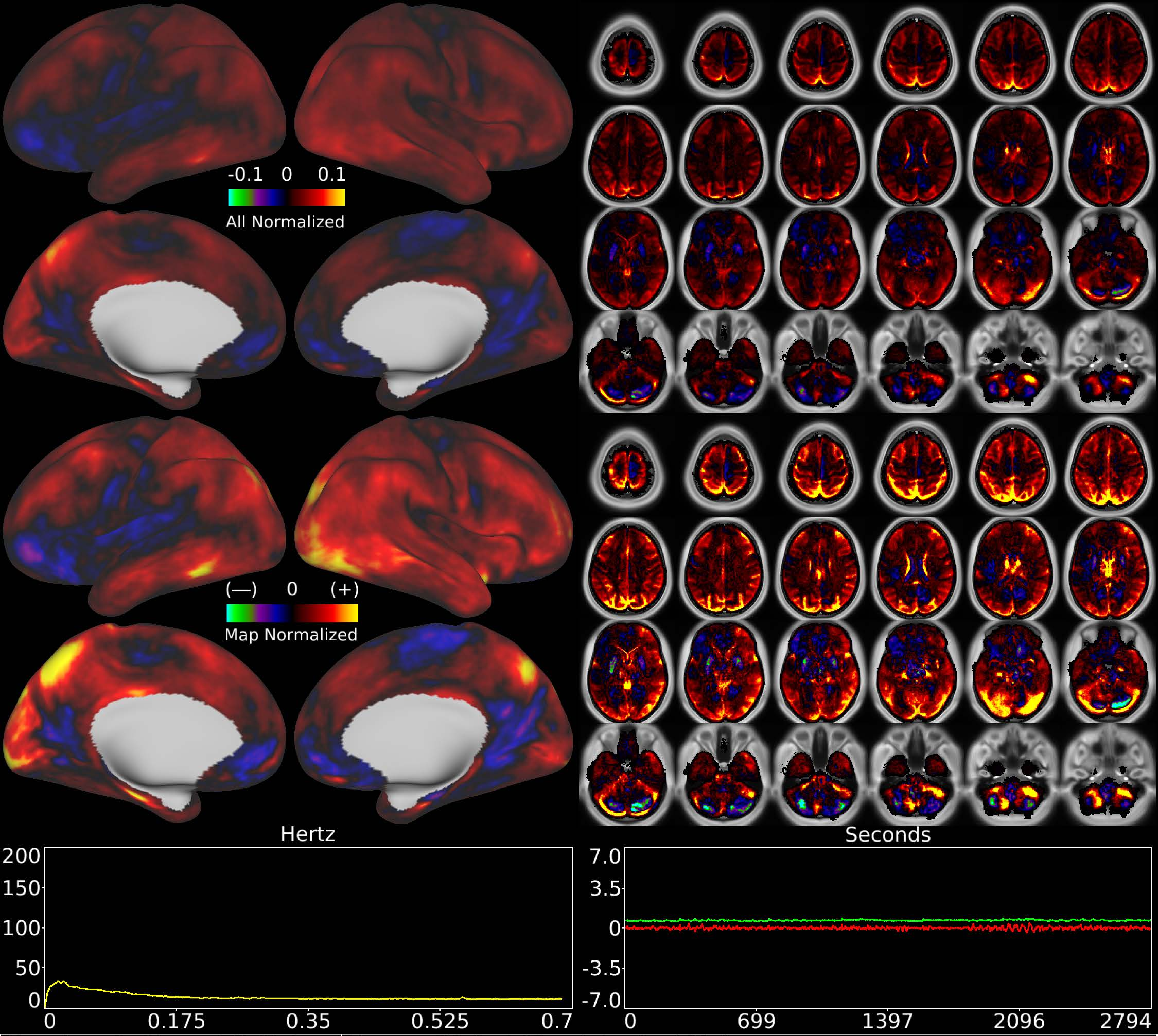


| | | |
|---|---|---|
| Number & Class: 52 Noise | Name: Global Physiological Noise | |
| RVT Correlated: No | DVARS Dip Associated: No | Cross-Subject Variable: No |
| Single Subject: No | % Variance Explained: 0.75 | Globality Index: 1.59 |
| Rest Component: No | Taskr Component: No | Task Modulated: No |

Rationale: Globally positive component without positive and negative patches that respect known RSNs or areas; some white matter signal

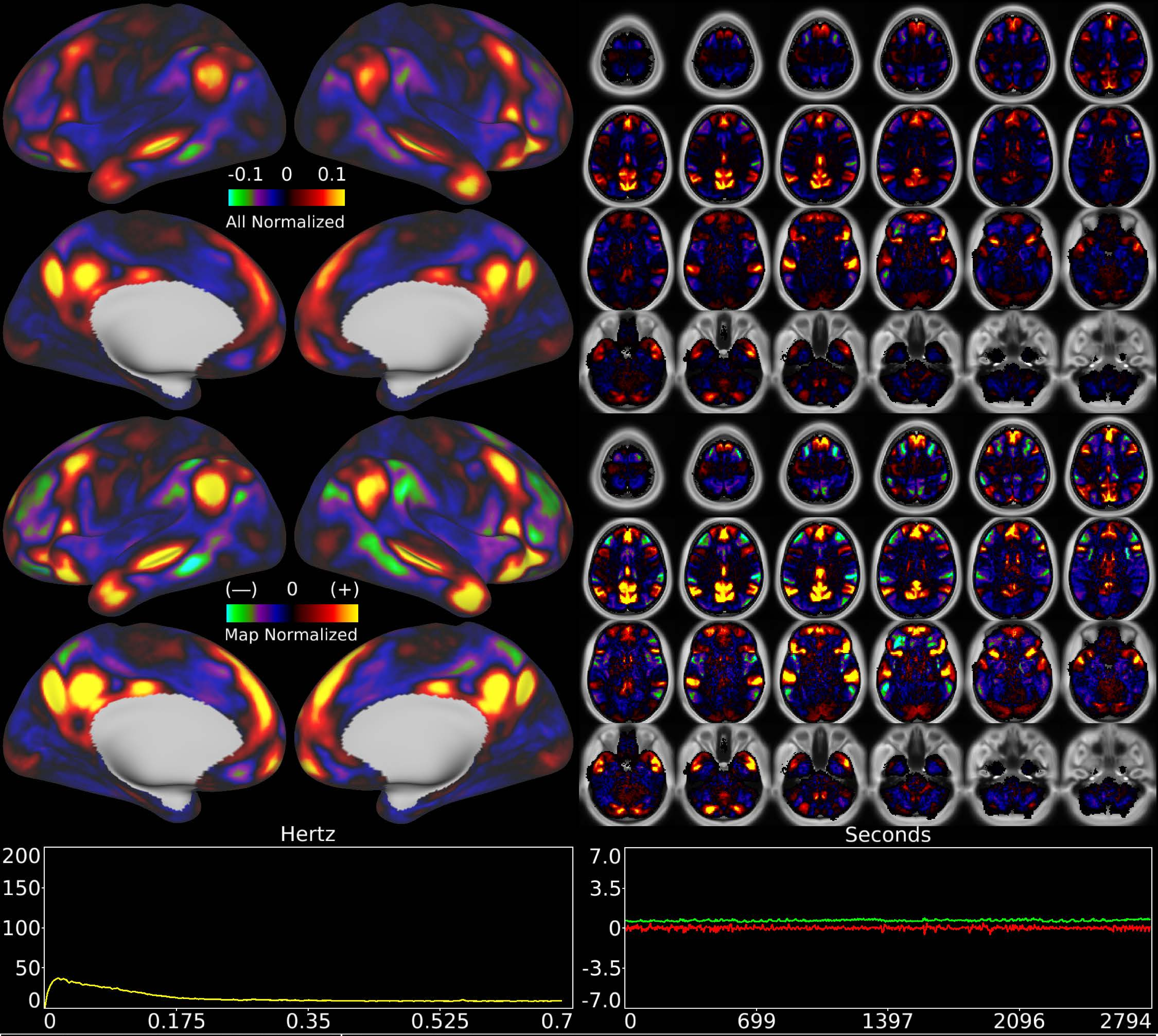


| | | |
|---|---|---|
| Number & Class: 53 Signal | Name: Unknown Network | |
| RVT Correlated: No | DVARS Dip Associated: No | Cross-Subject Variable: No |
| Single Subject: No | % Variance Explained: 0.74 | Globality Index: 0.61 |
| Rest Component: No | Taskr Component: No | Task Modulated: No |
| Rationale: Spatial map includes positive and negative patches that respect known RSNs (e.g. Default Mode Network) | | |

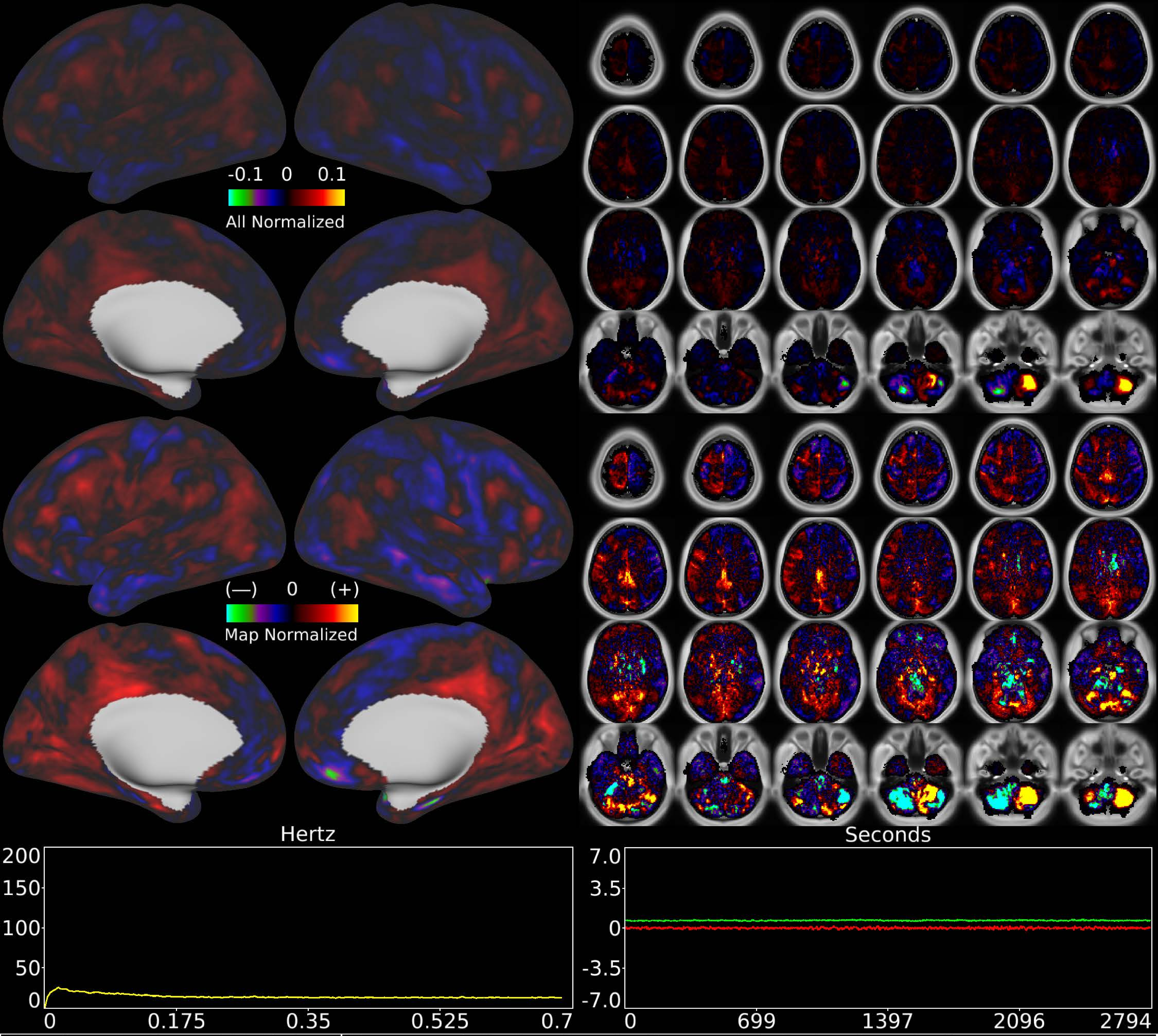


| | | |
|---|---|---|
| Number & Class: 54 Noise | Name: R Inferior Cerebellum > L Inferior Cerebellum | |
| RVT Correlated: No | DVARS Dip Associated: No | Cross-Subject Variable: No |
| Single Subject: Yes | % Variance Explained: 0.73 | Globality Index: 0.16 |
| Rest Component: No | Taskr Component: 51 | Task Modulated: No |
| Rationale: Single subject component whose spatial map is not reflective of known RSNs or Areas | | |

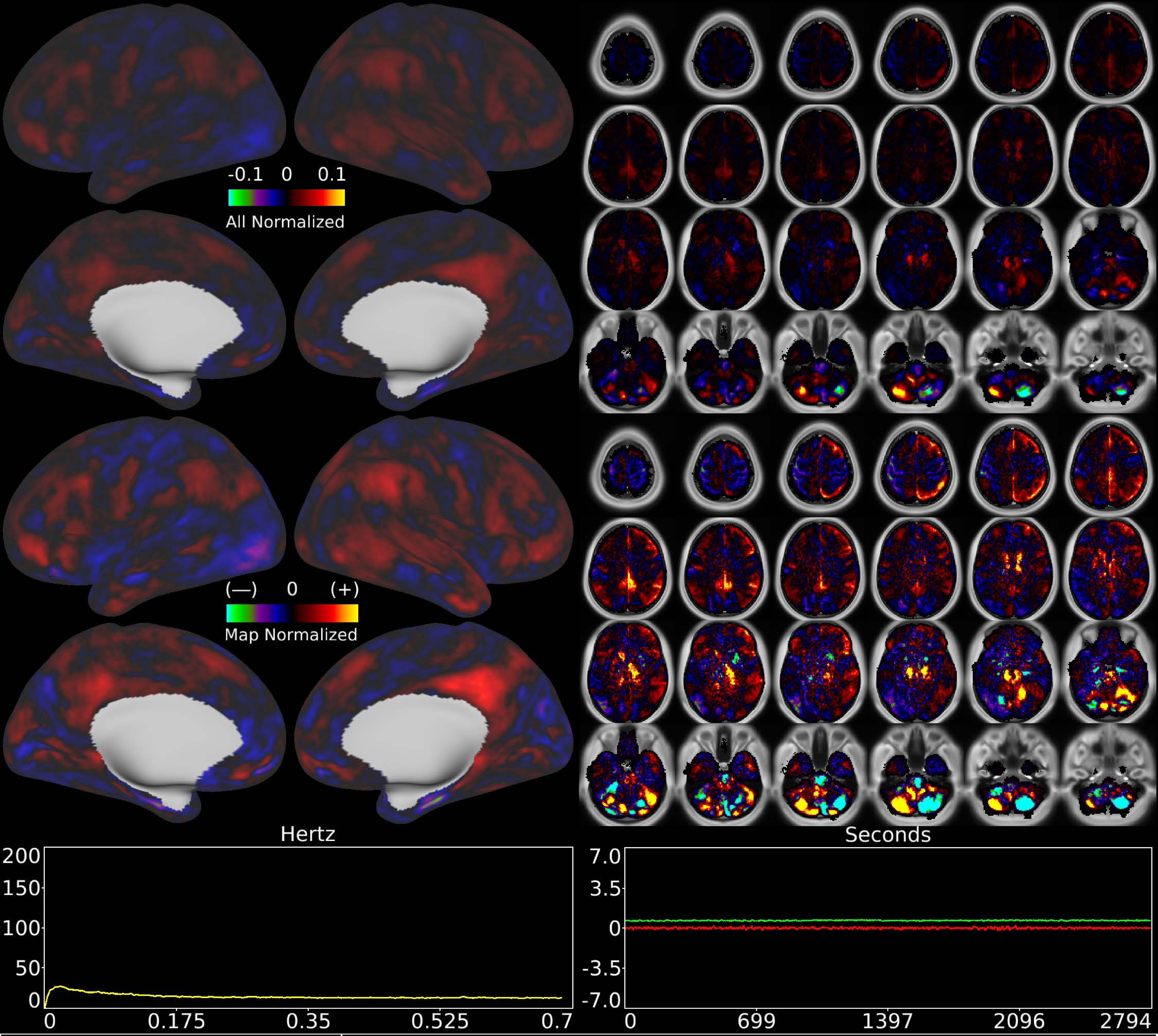


| | | |
|---|---|---|
| Number & Class: 55 Noise | Name: L Inferior Cerebellum > R Inferior Cerebellum | |
| RVT Correlated: No | DVARS Dip Associated: Yes | Cross-Subject Variable: No |
| Single Subject: Yes | % Variance Explained: 0.72 | Globality Index: 0.15 |
| Rest Component: No | Taskr Component: 54 | Task Modulated: No |
| Rationale: DVARS Dip associated single subject component whose spatial map is not reflective of known RSNs or Areas | | |

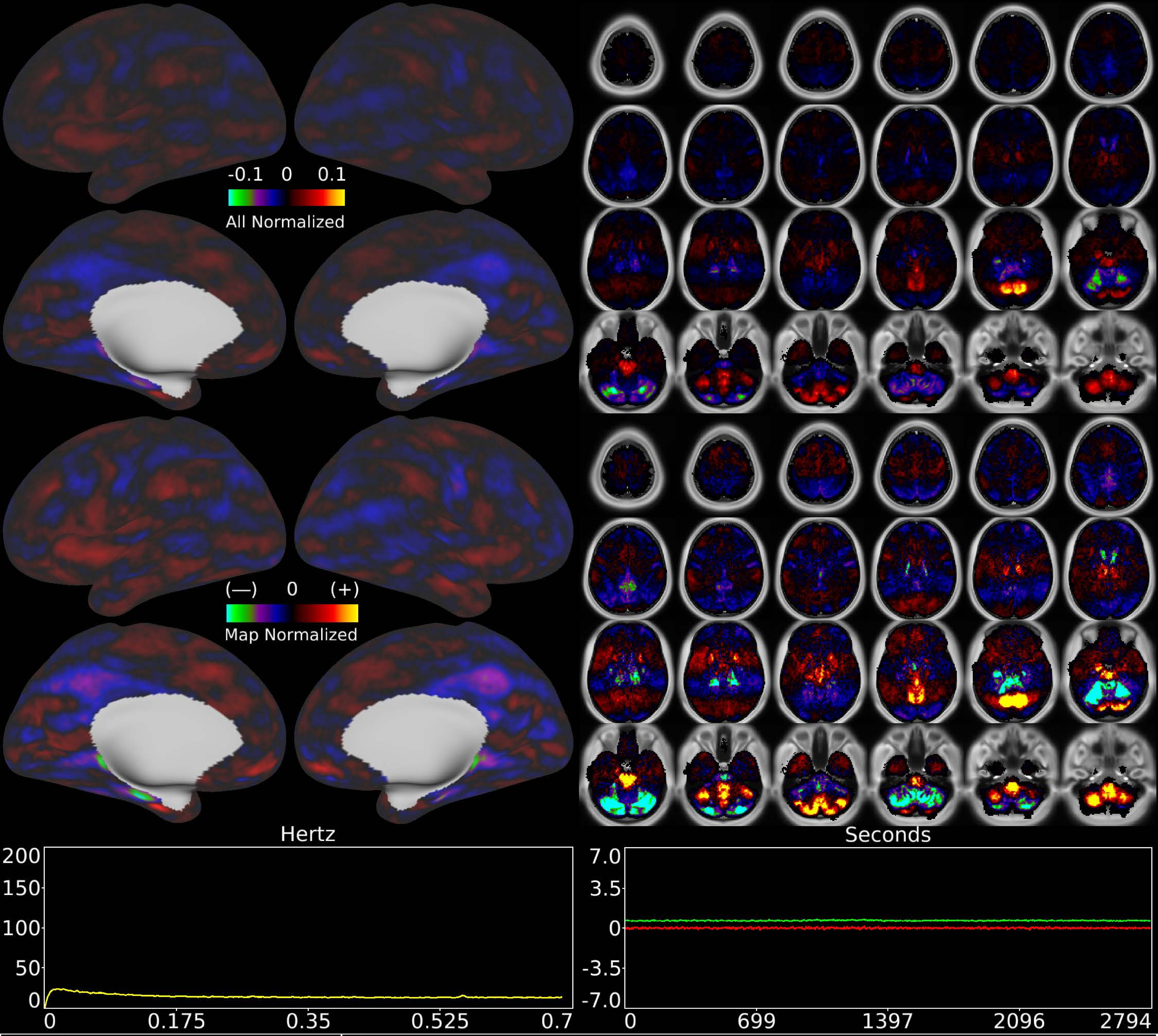


| Number & Class: 56 Noise | Name: Recon Artifact | |
|---|---|---|
| RVT Correlated: No | DVARS Dip Associated: Yes | Cross-Subject Variable: No |
| Single Subject: No | % Variance Explained: 0.7 | Globality Index: 0.17 |
| Rest Component: No | Taskr Component: No | Task Modulated: No |
| Rationale: Obvious banding in sagittal plane suggestive of multi-band reconstruction artifact; DVARS dips associated | | |

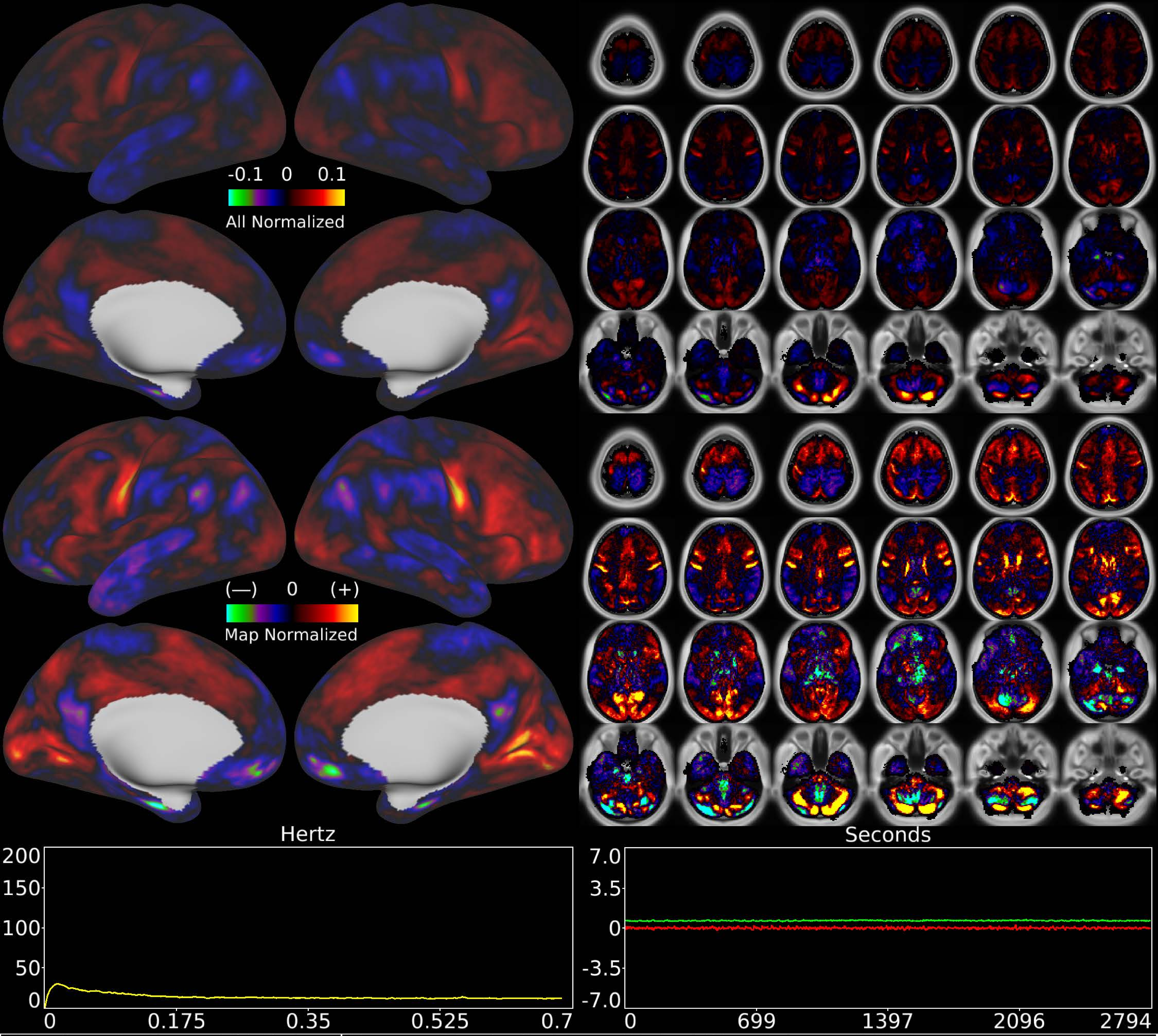


| | | |
|---|---|---|
| Number & Class: 57 Noise | Name: Single Subject Global Physiological Noise | |
| RVT Correlated: No | DVARS Dip Associated: No | Cross-Subject Variable: No |
| Single Subject: Yes | % Variance Explained: 0.69 | Globality Index: 0.25 |
| Rest Component: No | Taskr Component: No | Task Modulated: No |
| Rationale: Single subject component with high correlation to global timecourse | | |

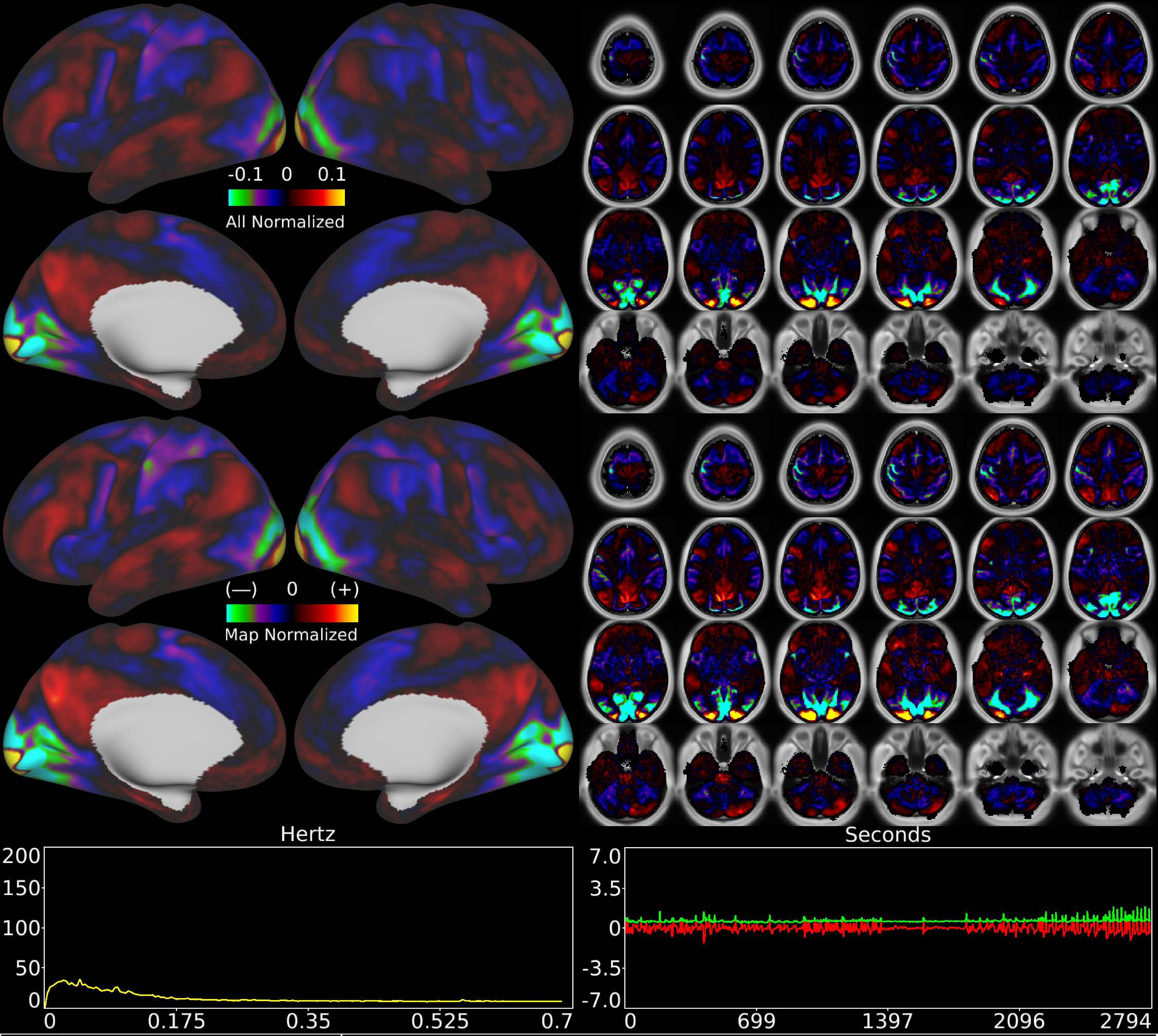


| | | |
|---|---|---|
| Number & Class: 58 Signal | Name: Complex Foveal > Peripheral Visual | |
| RVT Correlated: No | DVARS Dip Associated: No | Cross-Subject Variable: No |
| Single Subject: No | % Variance Explained: 0.69 | Globality Index: 0.37 |
| Rest Component: No | Taskr Component: No | Task Modulated: Emotion |
| Rationale: Spatial map includes positive and negative patches that respect known retinotopic visual organization (Foveal vs Peripheral) | | |

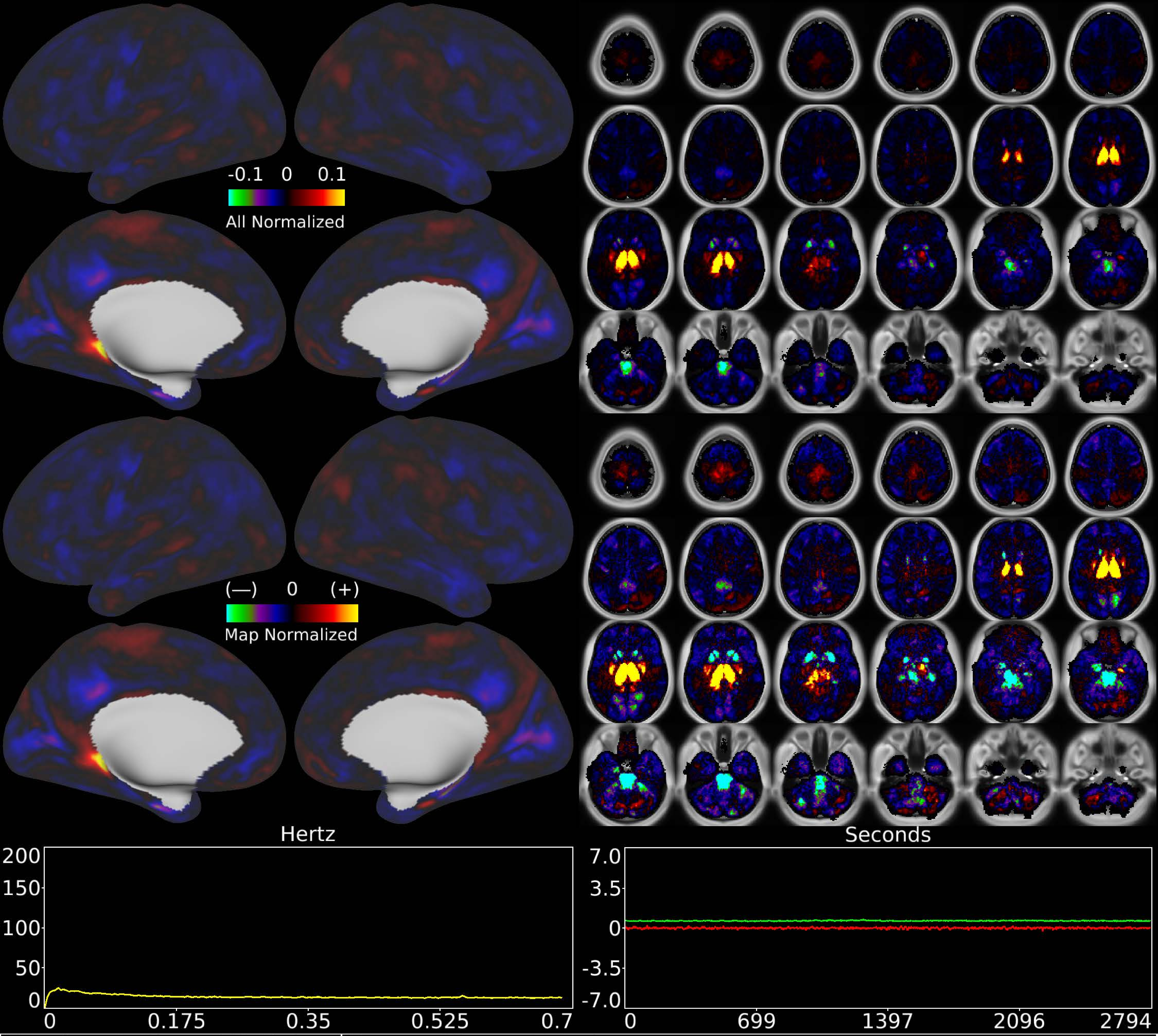


| | | |
|---|---|---|
| Number & Class: 59 Noise | Name: Bilateral Thalamus | |
| RVT Correlated: No | DVARS Dip Associated: No | Cross-Subject Variable: No |
| Single Subject: No | % Variance Explained: 0.66 | Globality Index: 1.13 |
| Rest Component: No | Taskr Component: No | Task Modulated: No |
| Rationale: Controversial: Variance correlated with other noise components; spatial map not reflective of known RSNs or areal boundaries | | |

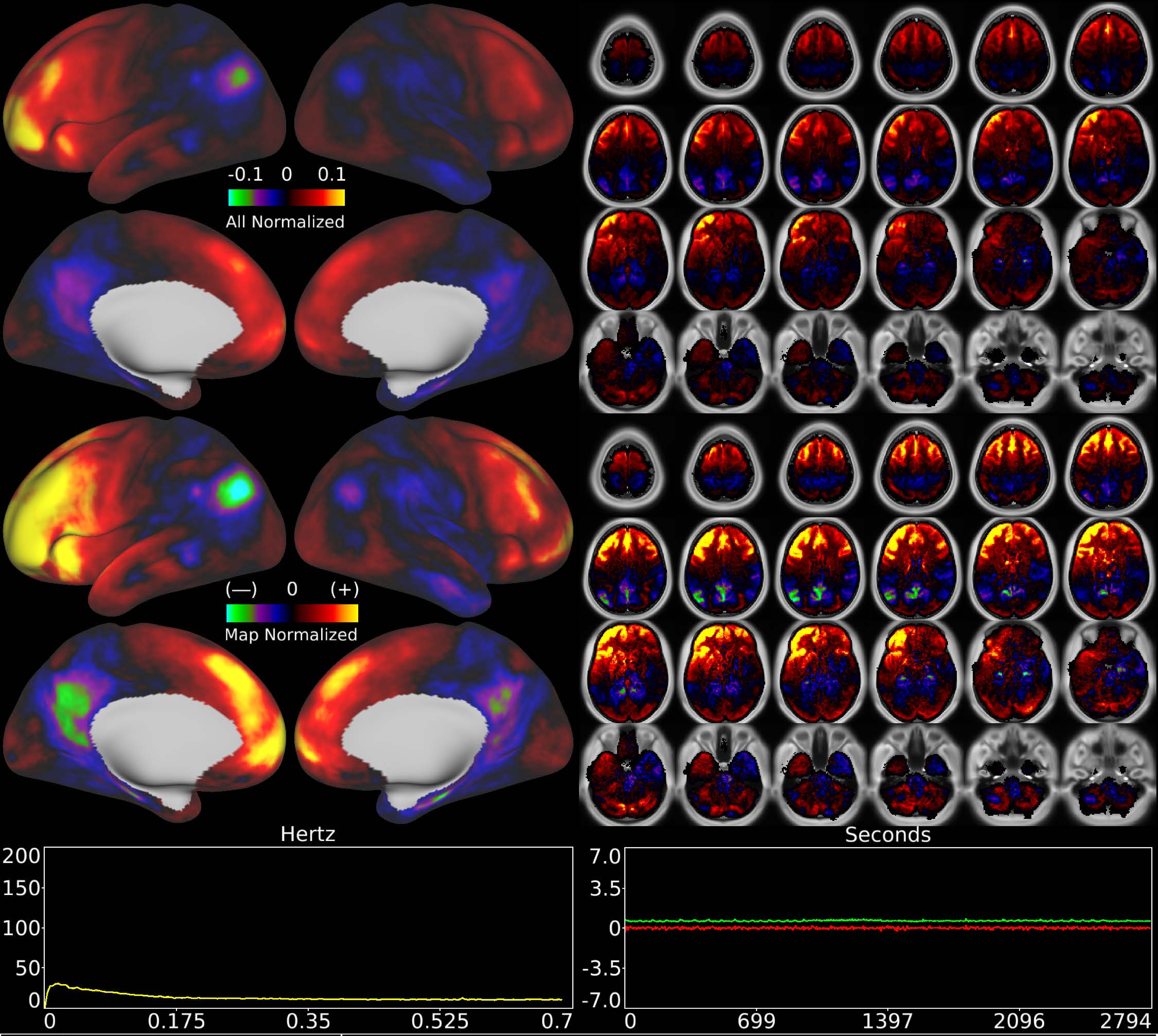


| | | |
|---|---|---|
| Number & Class: 60 Noise | Name: Coil or Movement | |
| RVT Correlated: No | DVARS Dip Associated: Yes | Cross-Subject Variable: No |
| Single Subject: No | % Variance Explained: 0.66 | Globality Index: 0.78 |
| Rest Component: No | Taskr Component: 46 | Task Modulated: No |

Rationale: Substantial white matter signal; associated with DVARS dips; perhaps related to coil or motion

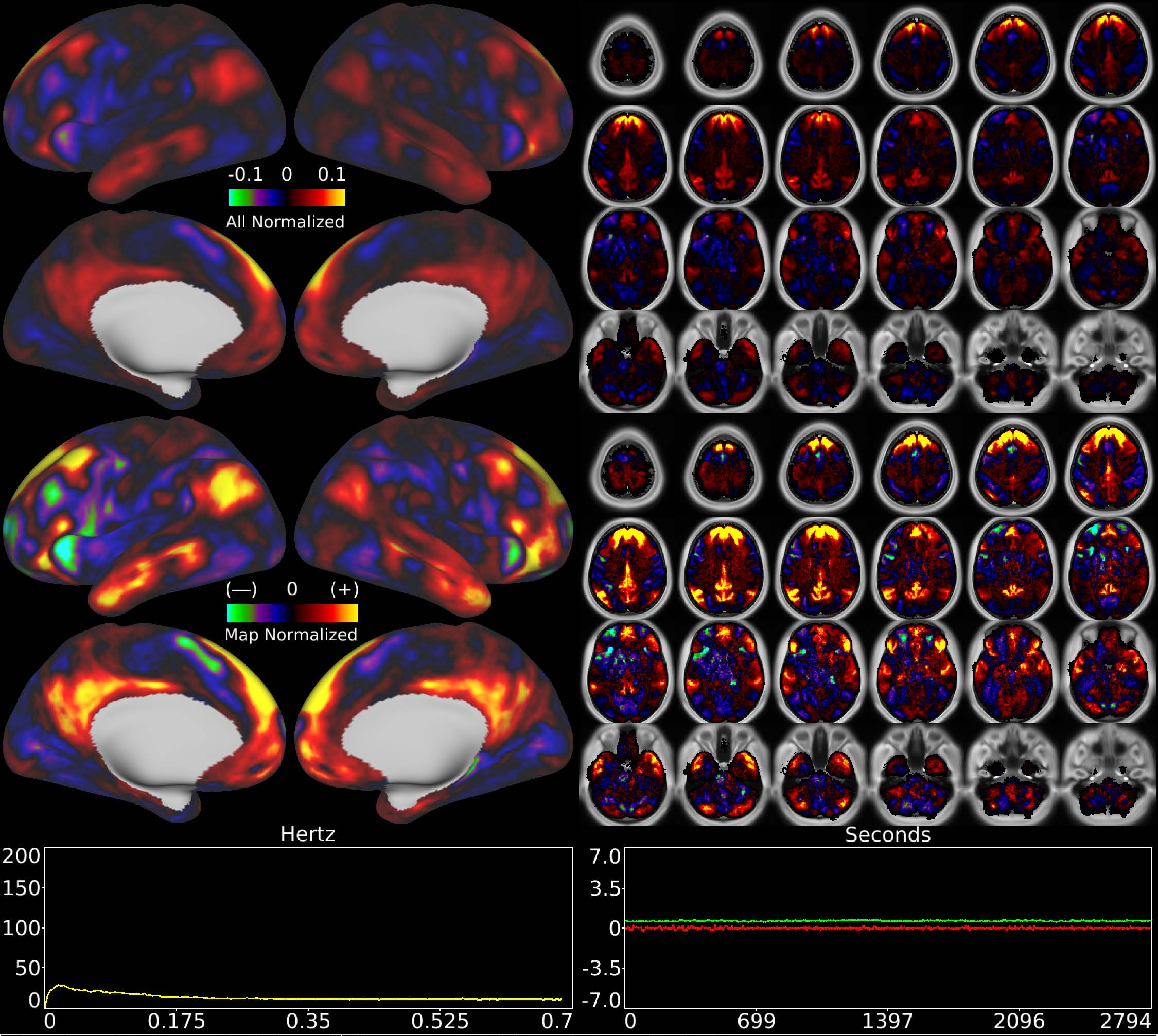


| | | |
|---|---|---|
| Number & Class: 61 Signal | Name: Subsidiary Default Mode | |
| RVT Correlated: No | DVARS Dip Associated: Yes | Cross-Subject Variable: No |
| Single Subject: No | % Variance Explained: 0.66 | Globality Index: 0.29 |
| Rest Component: No | Taskr Component: No | Task Modulated: No |
| Rationale: Spatial map includes positive and negative patches that respect known RSNs (e.g. Default Mode Network) | | |

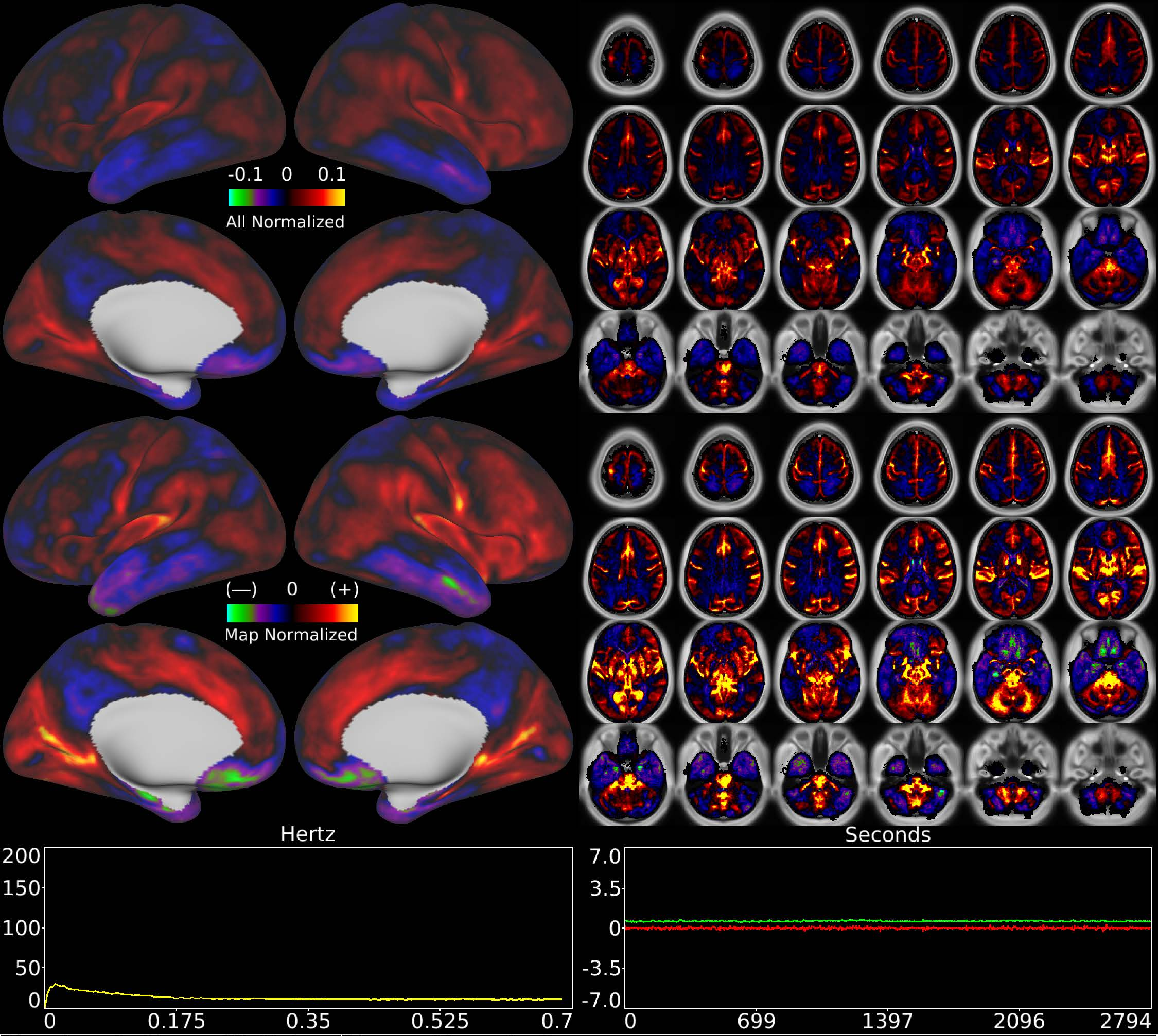


| | | |
|---|---|---|
| Number & Class: 62 Noise | Name: Vascular? | |
| RVT Correlated: No | DVARS Dip Associated: Yes | Cross-Subject Variable: No |
| Single Subject: No | % Variance Explained: 0.55 | Globality Index: 0.74 |
| Rest Component: No | Taskr Component: No | Task Modulated: No |

Rationale: Component more prominent in volume than on surface in cortical regions suggestive of vascular artifact; also DVARS dips associated

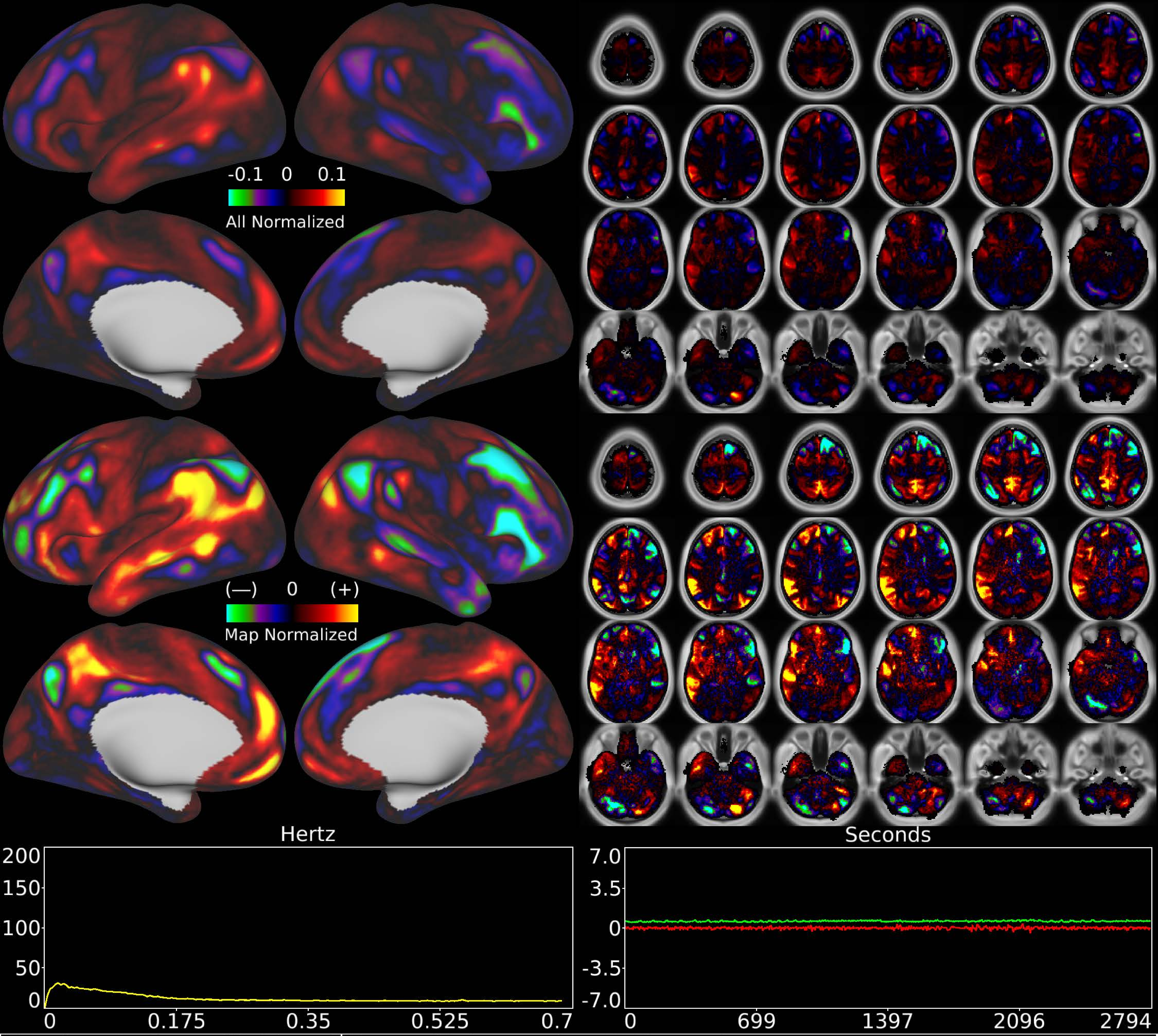


| | | |
|---|---|---|
| Number & Class: 63 Signal | Name: Unknown L > R Network | |
| RVT Correlated: No | DVARS Dip Associated: No | Cross-Subject Variable: No |
| Single Subject: No | % Variance Explained: 0.59 | Globality Index: 0.64 |
| Rest Component: No | Taskr Component: No | Task Modulated: No |
| Rationale: Spatial map includes positive and negative patches that respect known areal boundaries (e.g. POS2; 44; 45) | | |

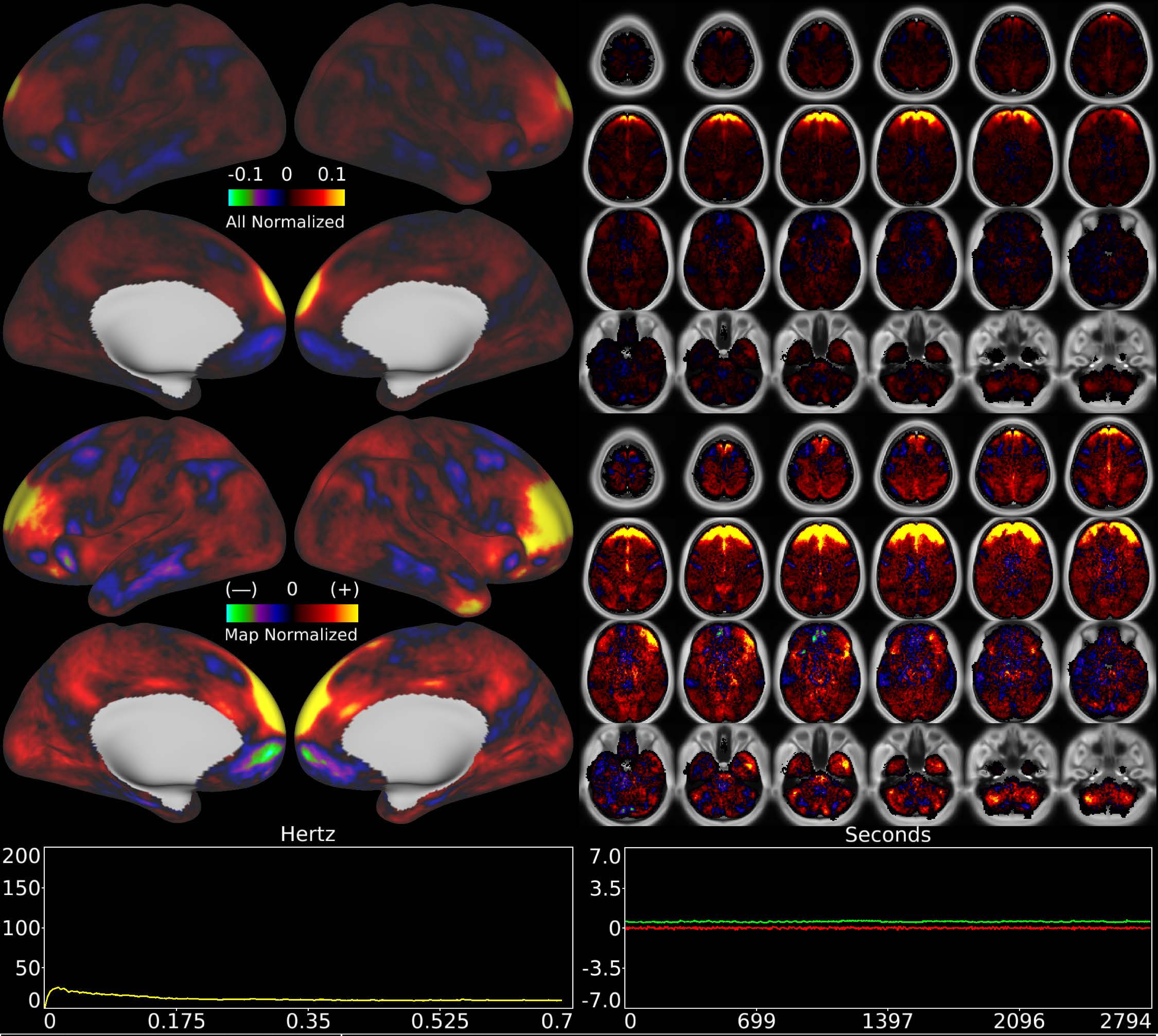


| | | |
|---|---|---|
| Number & Class: 64 Noise | Name: Coil? | |
| RVT Correlated: No | DVARS Dip Associated: Yes | Cross-Subject Variable: Yes |
| Single Subject: Yes | % Variance Explained: 0.54 | Globality Index: 1.82 |
| Rest Component: 70 | Taskr Component: 55 | Task Modulated: No |

Rationale: Spatial map is a band across frontal cortex and white matter; could be related to coil; head motion; or eye motion

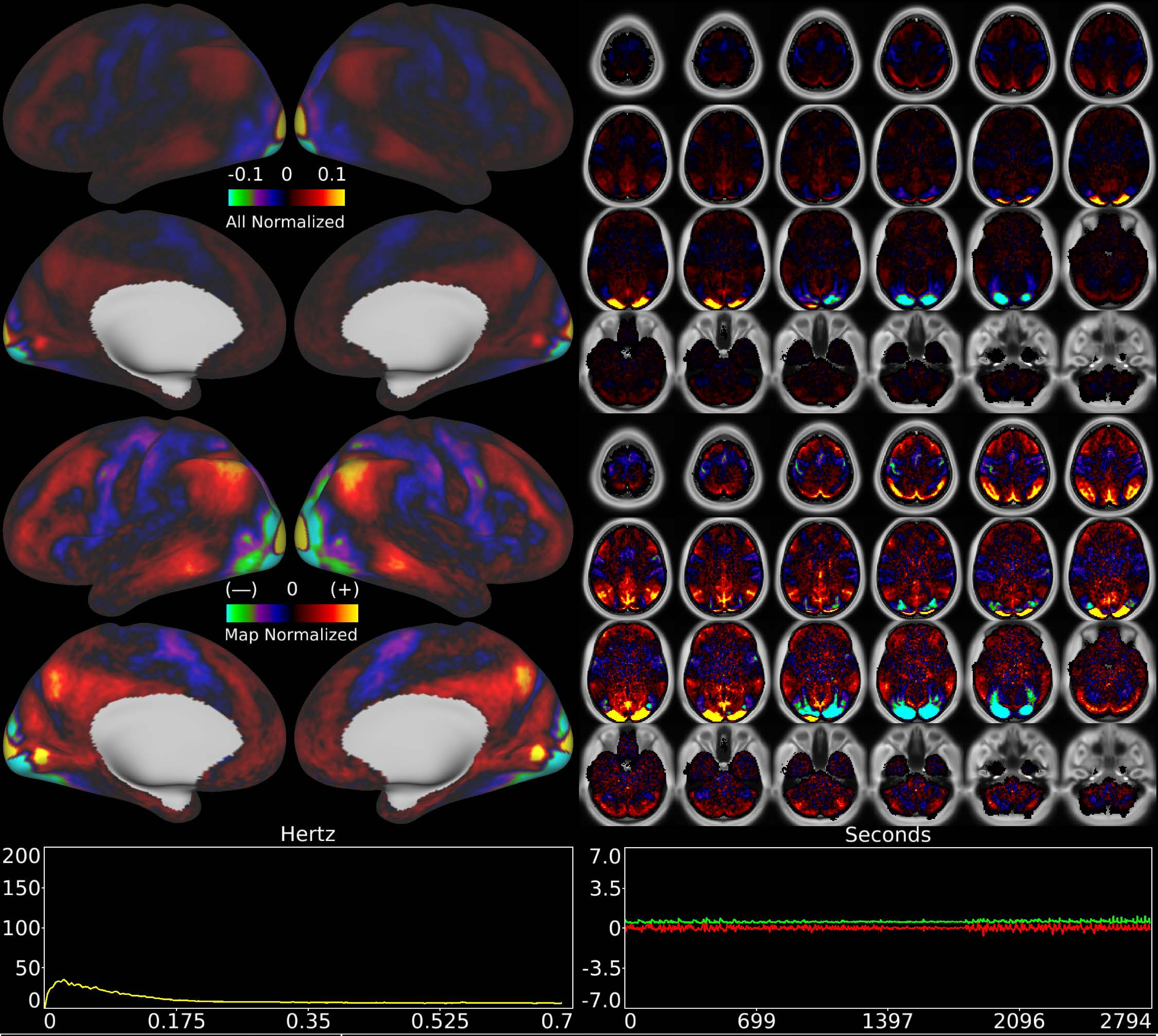


| Number & Class: 65 Signal | Name: Visuotopic: Foveal Dorsal > Ventral | |
|---|---|---|
| RVT Correlated: No | DVARS Dip Associated: No | Cross-Subject Variable: No |
| Single Subject: No | % Variance Explained: 0.54 | Globality Index: 0.62 |
| Rest Component: 76 | Taskr Component: 53 | Task Modulated: No |

Rationale: Spatial map includes positive and negative patches that respect known retinotopic visual organization (Foveal and Lower vs Upper)

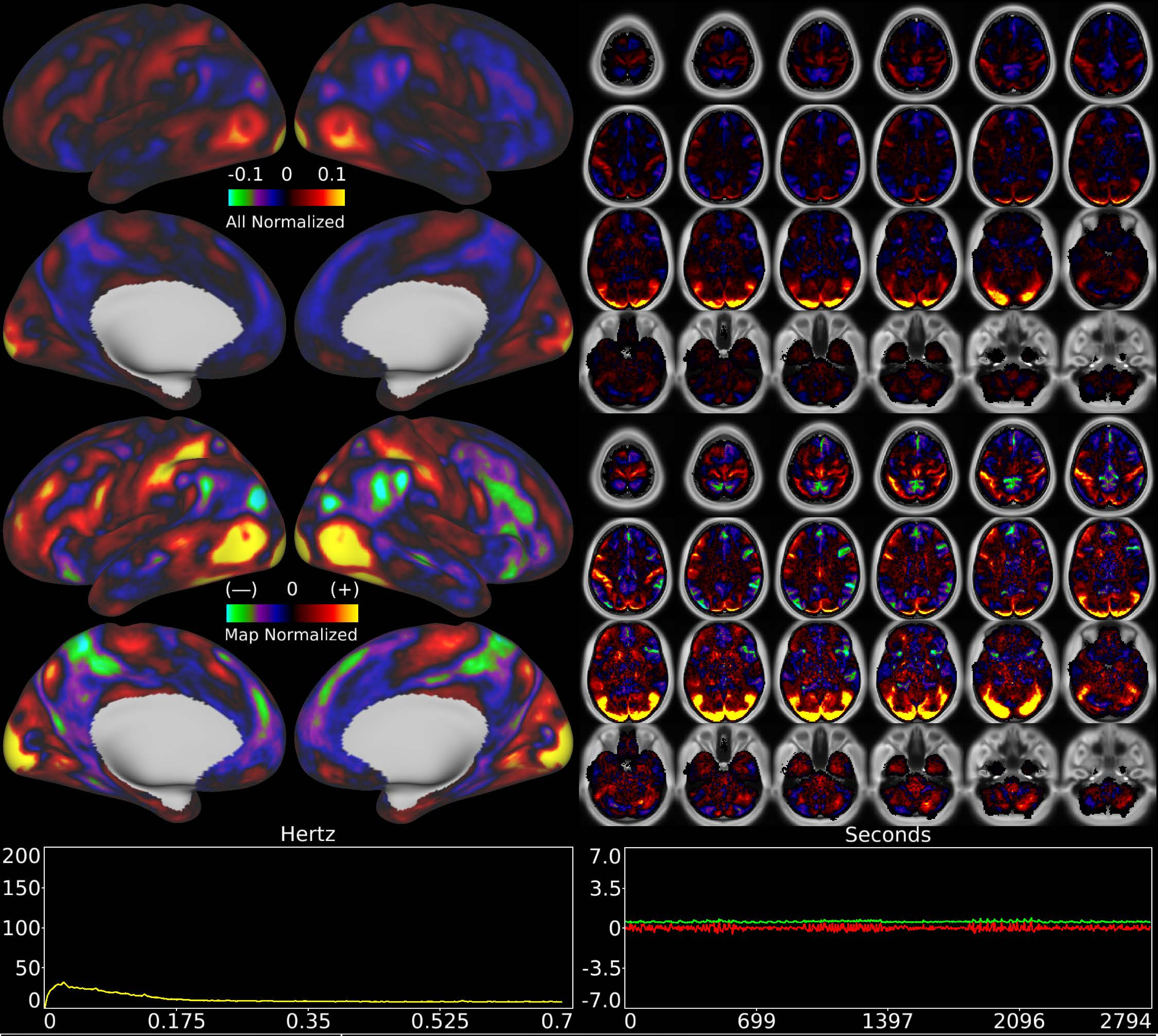


| | | |
|---|---|---|
| Number & Class: 66 Signal | Name: Extrastriate Visual Network | |
| RVT Correlated: No | DVARS Dip Associated: No | Cross-Subject Variable: No |
| Single Subject: No | % Variance Explained: 0.52 | Globality Index: 0.46 |
| Rest Component: No | Taskr Component: No | Task Modulated: No |
| Rationale: Spatial map has elements of visuotopy and known RSNs though in an atypical arrangement | | |

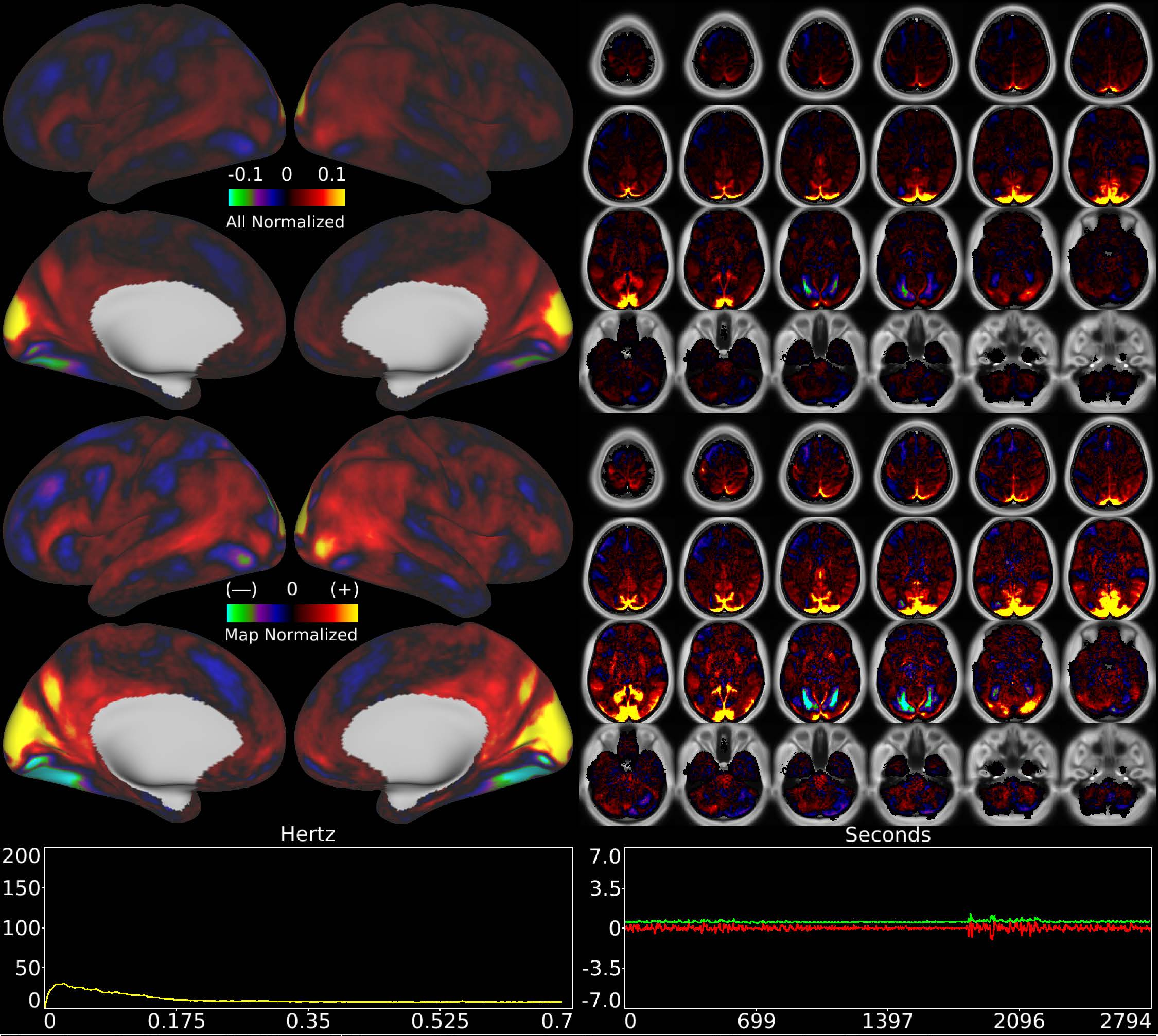


| | | |
|---|---|---|
| Number & Class: 67 Signal | Name: Visuotopic: Paracentral Dorsal > Ventral | |
| RVT Correlated: No | DVARS Dip Associated: No | Cross-Subject Variable: No |
| Single Subject: No | % Variance Explained: 0.52 | Globality Index: 1.42 |
| Rest Component: -82 | Taskr Component: 49 | Task Modulated: No |
| Rationale: Spatial map includes positive and negative patches that respect known retinotopic visual organization (Lower vs Upper) | | |

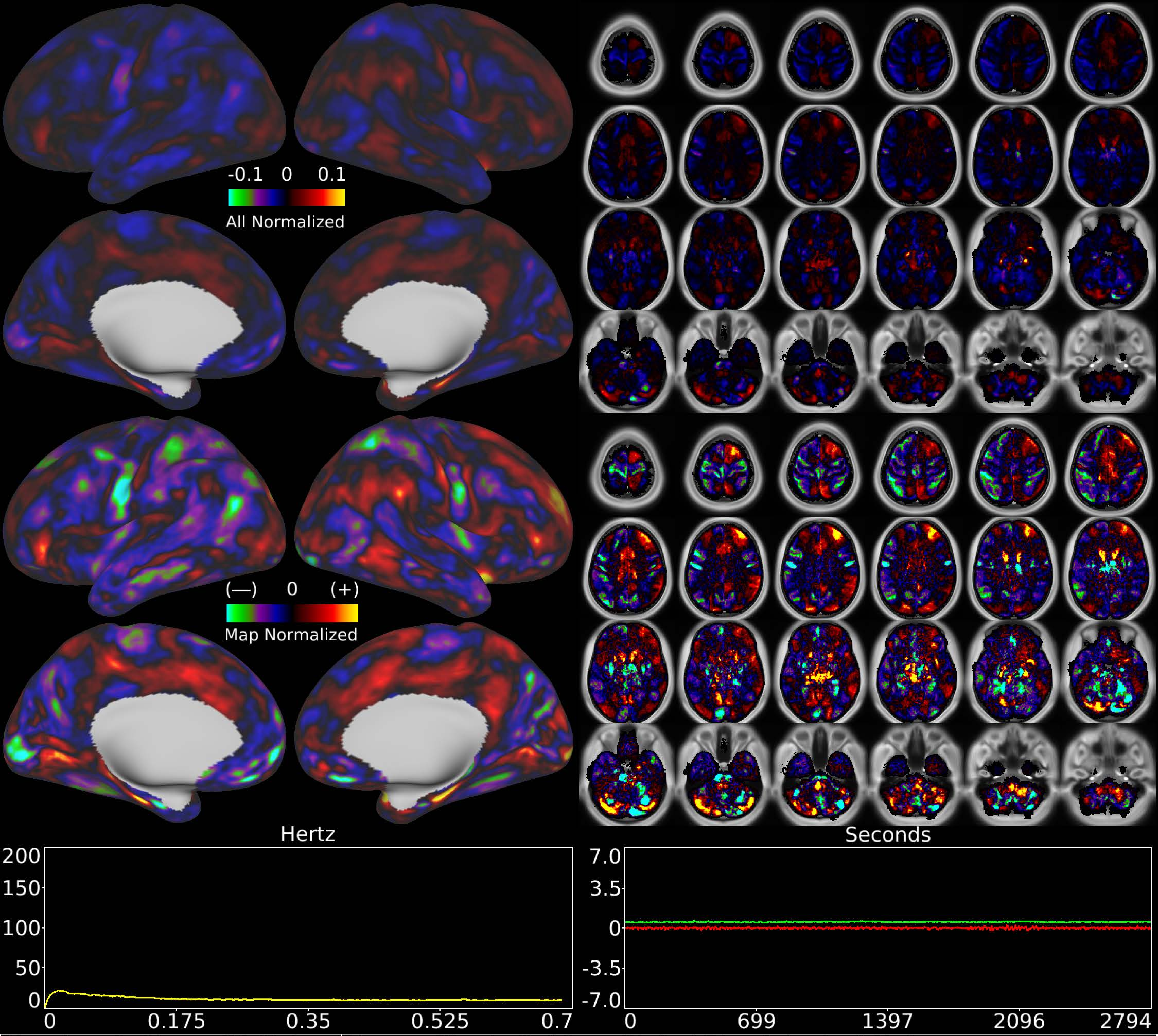


| | | |
|---|---|---|
| Number & Class: 68 Noise | Name: Single Subject Global Physiological Noise | |
| RVT Correlated: No | DVARS Dip Associated: No | Cross-Subject Variable: No |
| Single Subject: Yes | % Variance Explained: 0.46 | Globality Index: 0.41 |
| Rest Component: No | Taskr Component: No | Task Modulated: No |
| Rationale: Single subject component with high correlation to global timecourse | | |

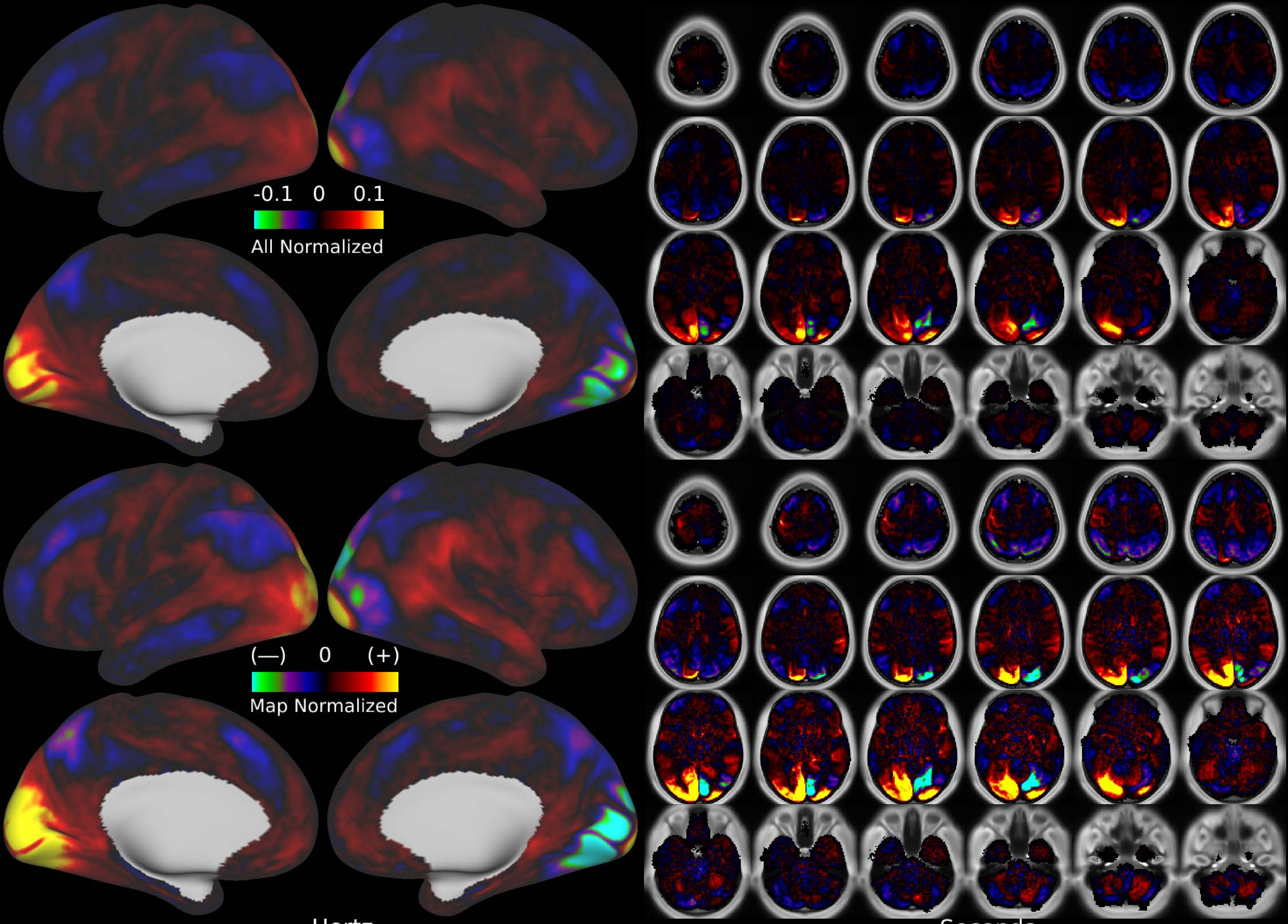


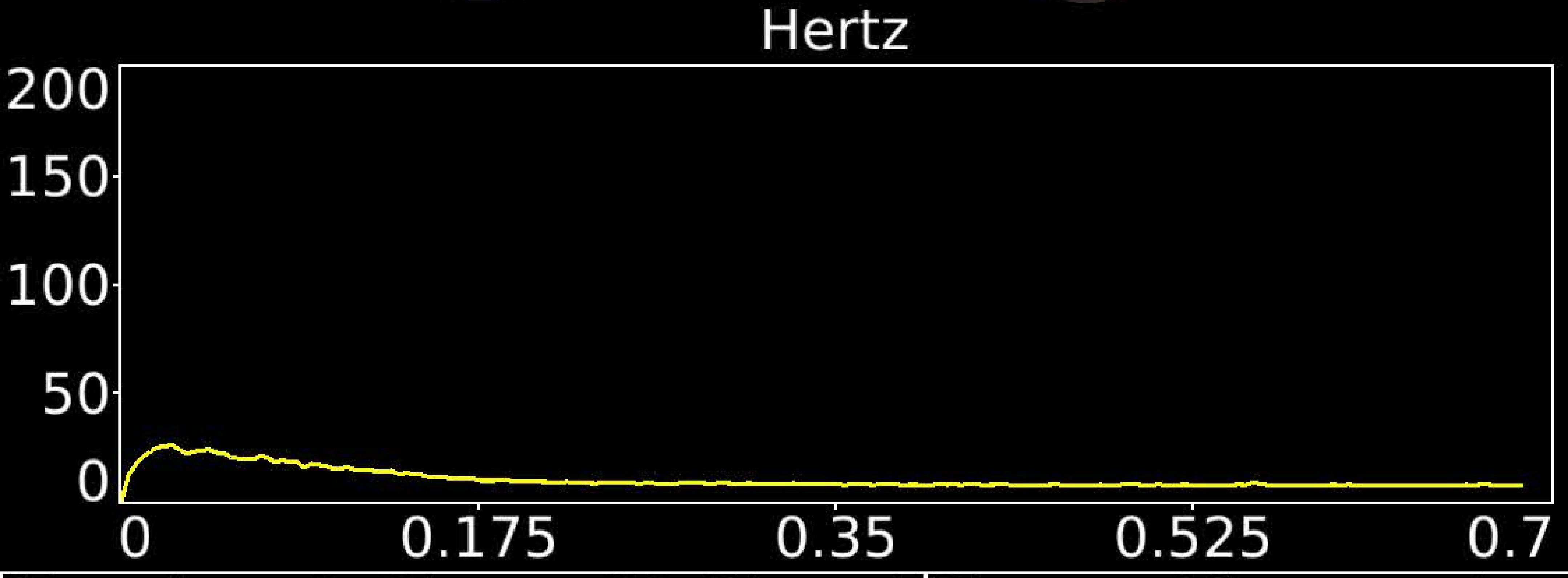


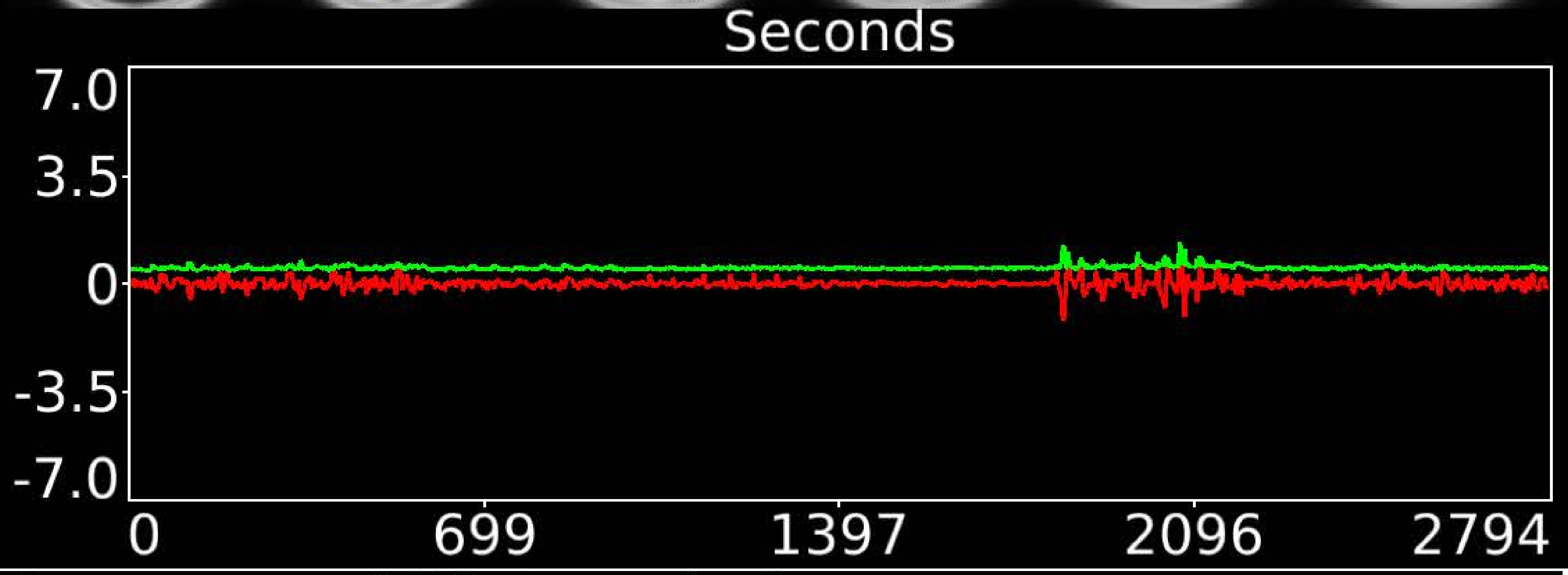


| | | |
|---|---|---|
| Number & Class: 69 Signal | Name: Visuotopic: Paracentral Right > Left | |
| RVT Correlated: No | DVARS Dip Associated: No | Cross-Subject Variable: No |
| Single Subject: No | % Variance Explained: 0.45 | Globality Index: 0.65 |
| Rest Component: 84 | Taskr Component: -57 | Task Modulated: No |
| Rationale: Spatial map includes positive and negative patches that respect known retinotopic visual organization (Foveal Right vs Left) | | |

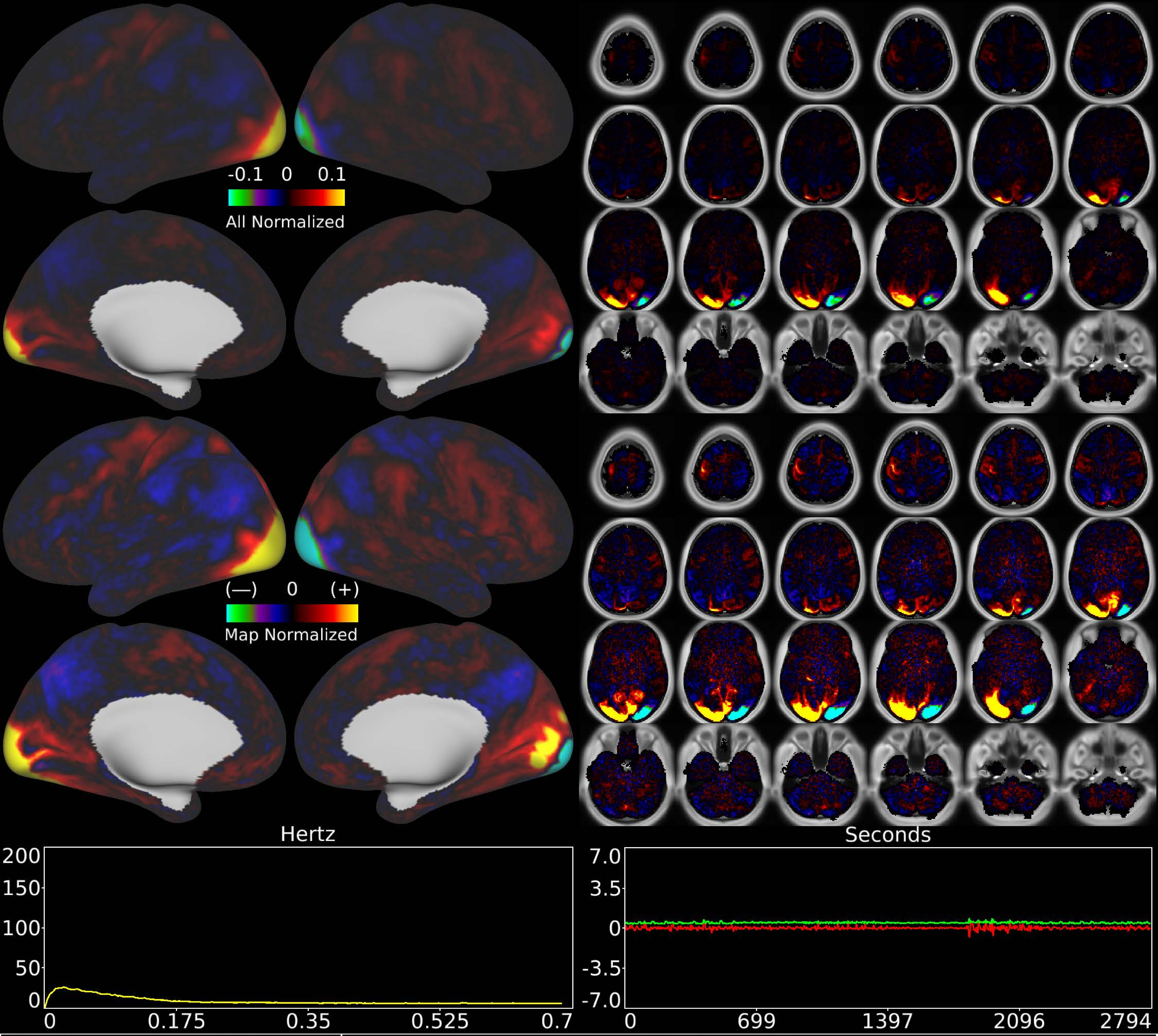


| | | |
|---|---|---|
| Number & Class: 70 Signal | Name: Visuotopic: Foveal Left > Right | |
| RVT Correlated: No | DVARS Dip Associated: No | Cross-Subject Variable: No |
| Single Subject: No | % Variance Explained: 0.37 | Globality Index: 0.26 |
| Rest Component: 81 | Taskr Component: 58 | Task Modulated: No |
| Rationale: Spatial map includes positive and negative patches that respect known retinotopic visual organization (Peripheral Right vs Left) | | |
