## Supplementary Materials for "Using Temporal ICA to Selectively Remove Global Noise While Preserving Global Signal in Functional MRI Data"

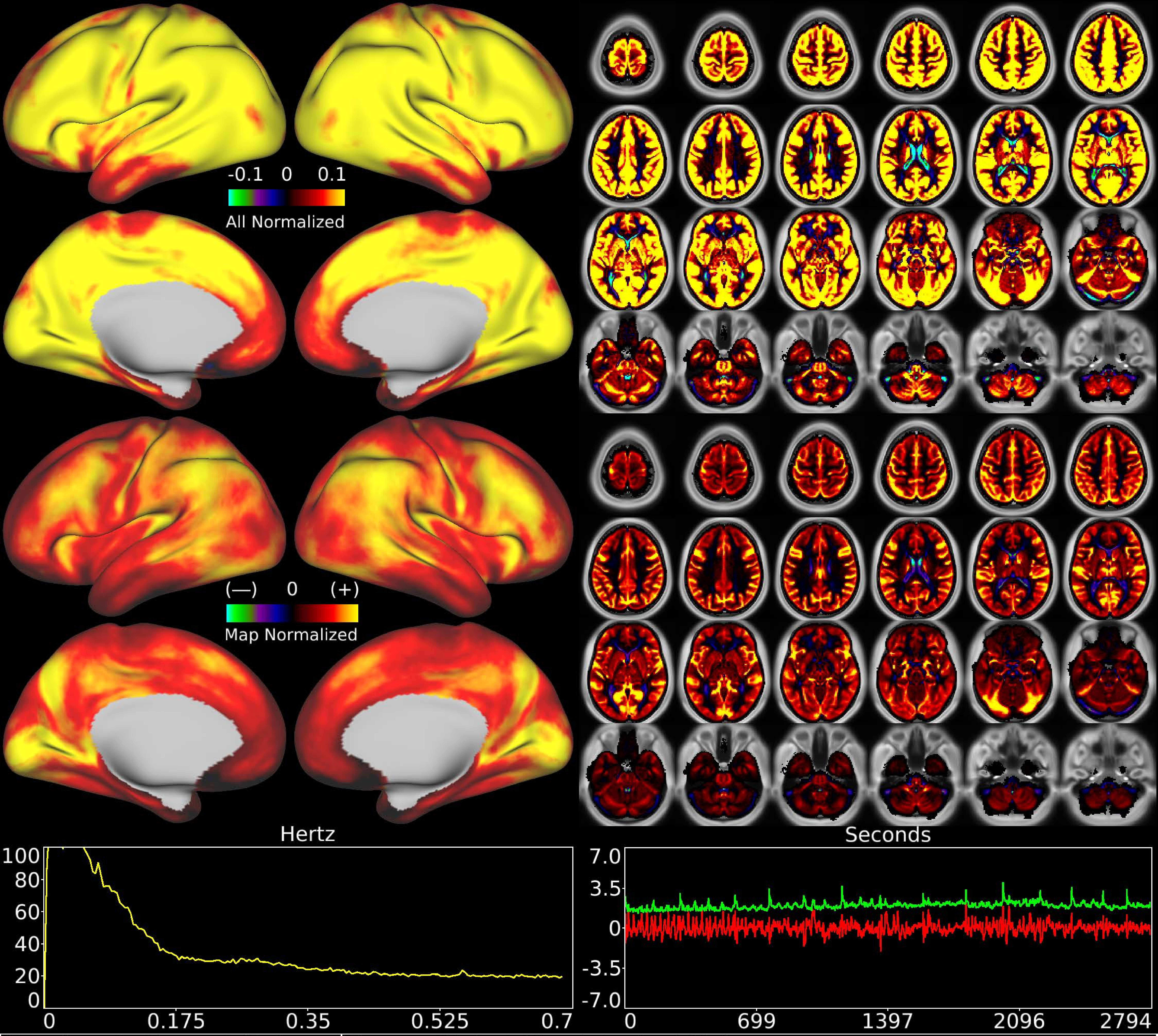


| | | |
|---|---|---|
| Number & Class: 1 Noise | Name: Global Physiological Noise | |
| RVT Correlated: Yes | DVARS Dip Associated: Yes | Cross-Subject Variable: Yes |
| Single Subject: No | % Variance Explained: 7.85 | Globality Index: 5.91 |
| Task Component: 1 | Rest Component: 3+6+8 | Task Modulated: No |
| Rationale: Globally positive component matching hypothesized pattern for physiological artifact and does not clearly follow known areal boundaries | | |

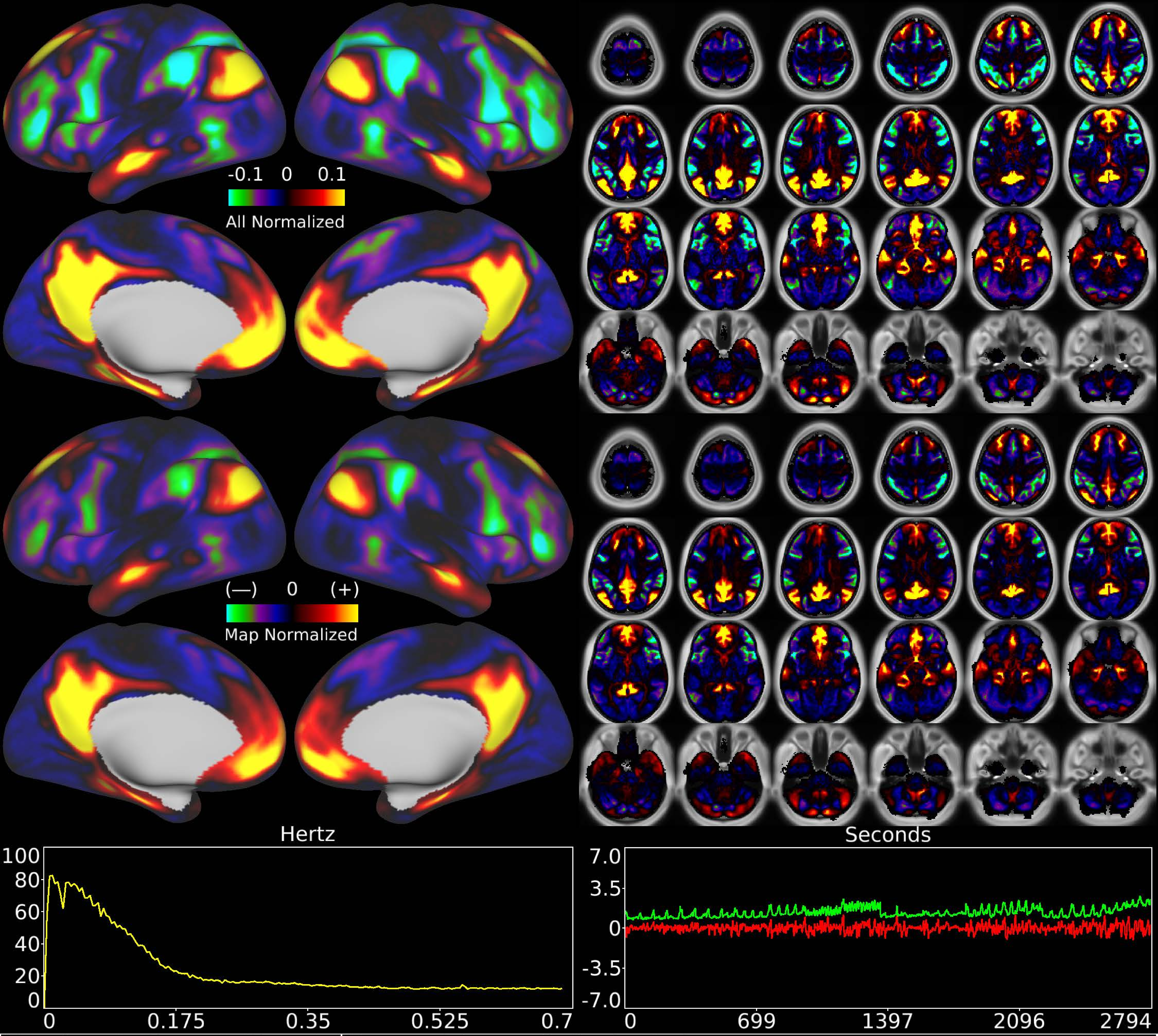


| | | |
|---|---|---|
| Number & Class: 2 Signal | Name: Primary Default Mode | |
| RVT Correlated: No | DVARS Dip Associated: No | Cross-Subject Variable: Yes |
| Single Subject: No | % Variance Explained: 4.1 | Globality Index: 1.04 |
| Task Component: 5 | Rest Component: 10 | Task Modulated: Emotion |
| Rationale: Spatial map includes positive and negative patches that respect known RSNs (e.g. Default Mode Network) | | |

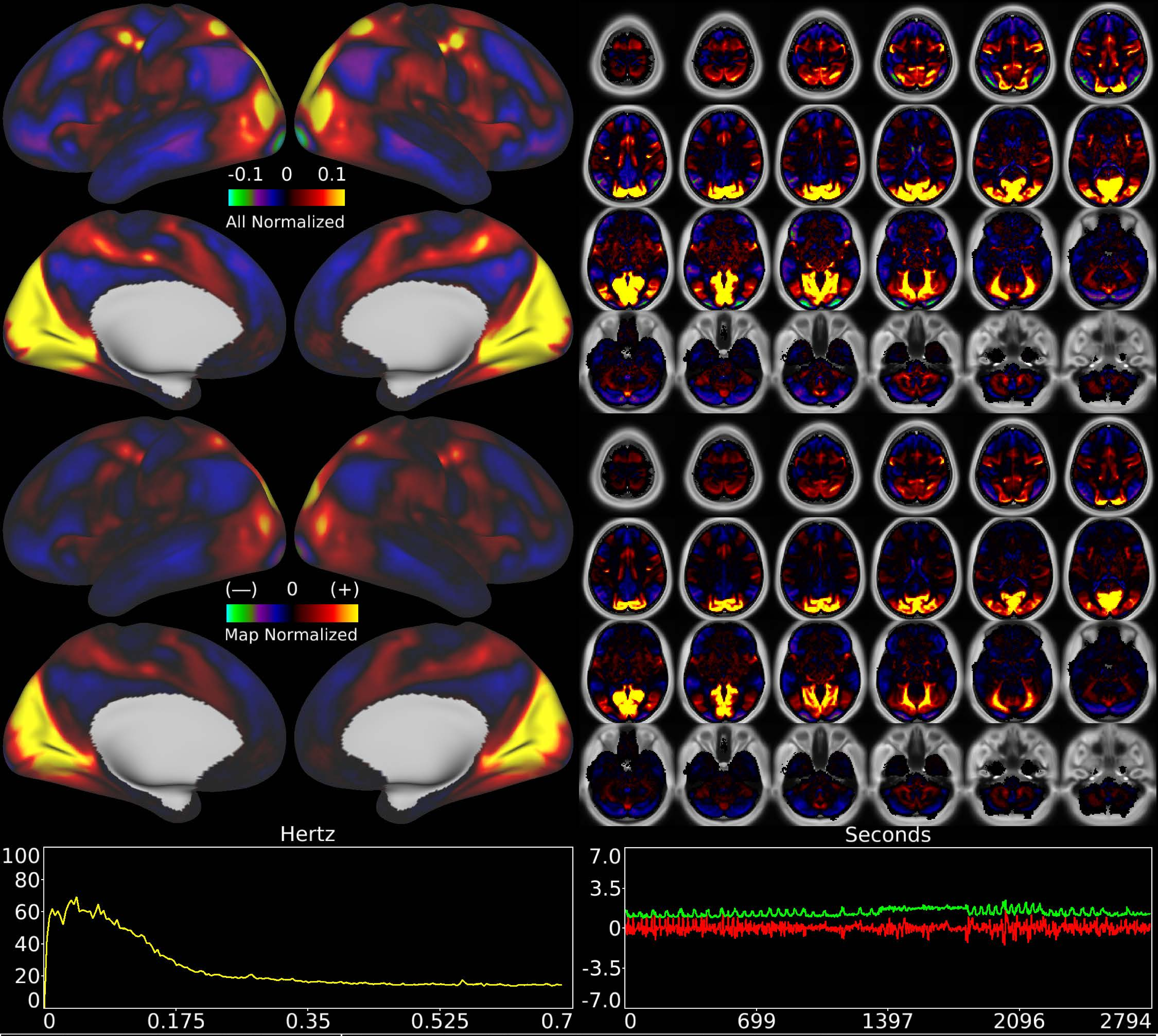


| | | |
|---|---|---|
| Number & Class: 3 Signal | Name: Pan-Visual (Peripheral > Foveal) | |
| RVT Correlated: No | DVARS Dip Associated: No | Cross-Subject Variable: Yes |
| Single Subject: No | % Variance Explained: 3.82 | Globality Index: 0.4 |
| Task Component: 15 | Rest Component: 4 | Task Modulated: Language |
| Rationale: Spatial map includes positive and negative patches that respect known retinotopic visual organization (Peripheral vs Foveal) | | |

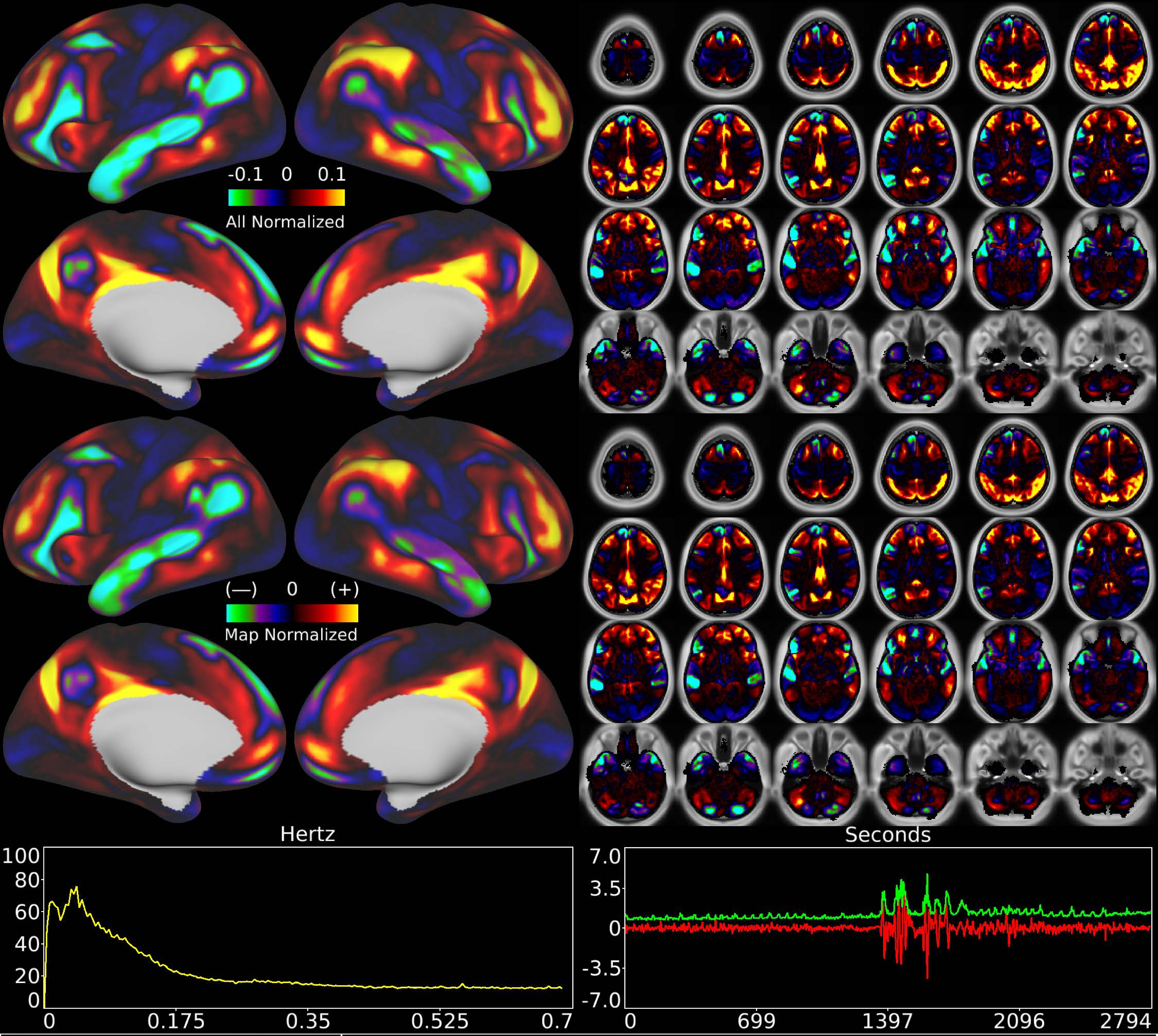


| | | |
|---|---|---|
| Number & Class: 4 Signal | Name: Language Task Story | |
| RVT Correlated: No | DVARS Dip Associated: No | Cross-Subject Variable: No |
| Single Subject: No | % Variance Explained: 3.23 | Globality Index: 0.41 |
| Task Component: -11 | Rest Component: No | Task Modulated: Language |
| Rationale: Spatial map includes positive and negative patches that are correlated with a specific task design | | |

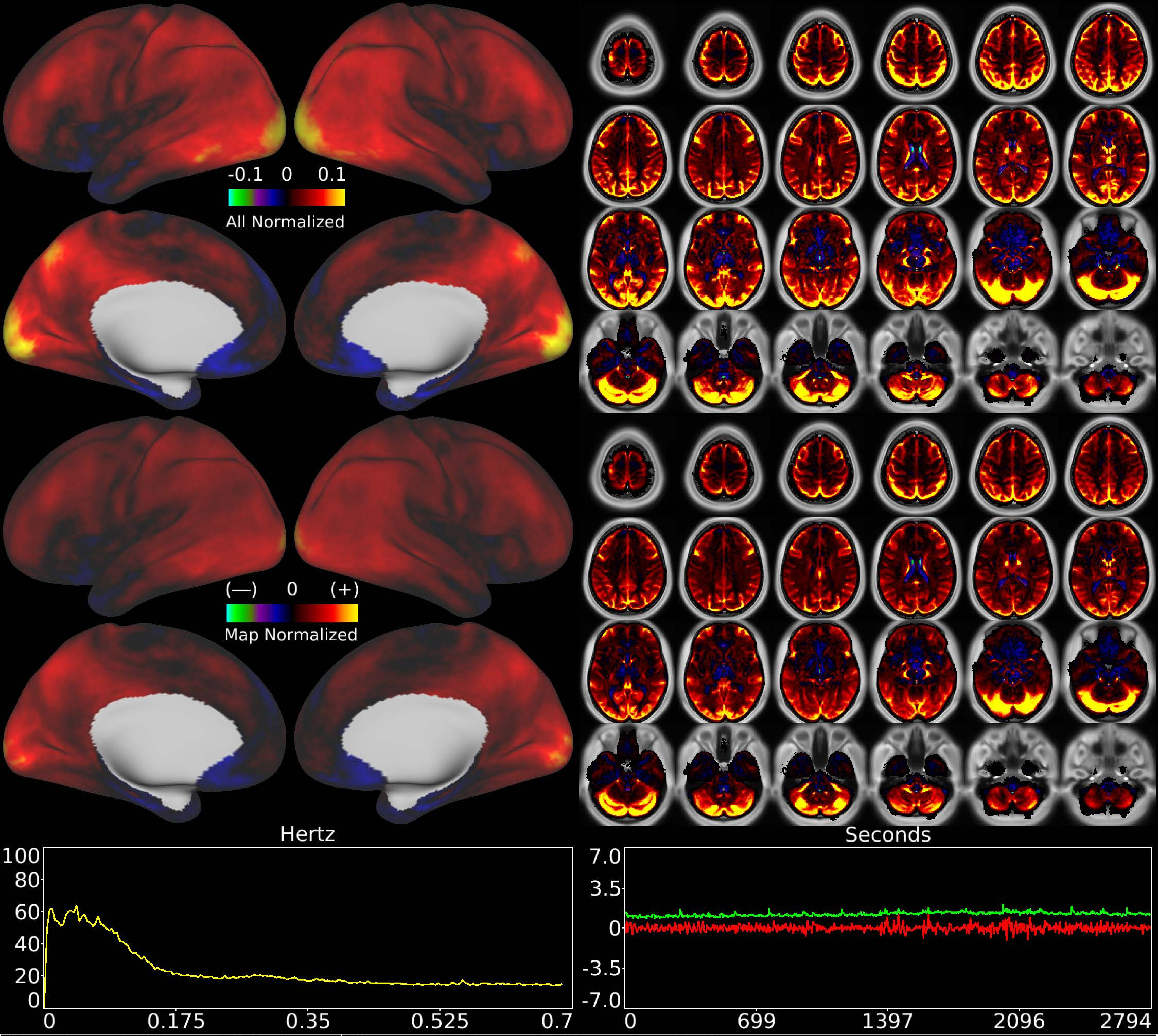


| | | |
|---|---|---|
| Number & Class: 5 Noise | Name: Veins + WM + Cerebellum | |
| RVT Correlated: No | DVARS Dip Associated: Yes | Cross-Subject Variable: Yes |
| Single Subject: Yes | % Variance Explained: 2.83 | Globality Index: 2.61 |
| Task Component: 8 | Rest Component: 19+28 | Task Modulated: No |
| Rationale: Spatial map contains substantial white matter and venous signal | | |

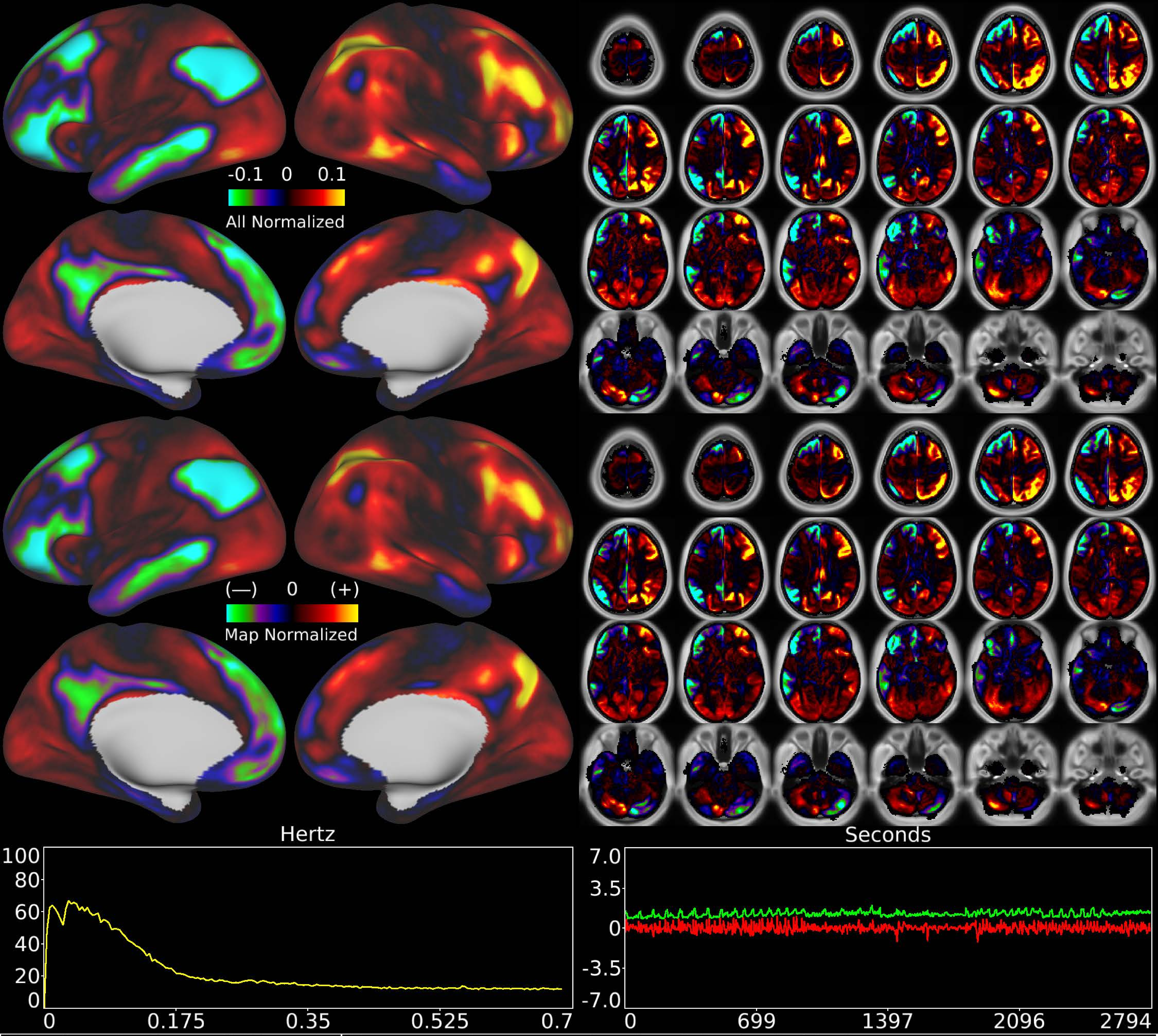


| | | |
|---|---|---|
| Number & Class: 6 Signal | Name: R Fronto-parietal > L Default Mode | |
| RVT Correlated: No | DVARS Dip Associated: No | Cross-Subject Variable: Yes |
| Single Subject: No | % Variance Explained: 3.04 | Globality Index: 1.02 |
| Task Component: -12 | Rest Component: No | Task Modulated: No |
| Rationale: Spatial map includes positive and negative patches that respect known RSNs (e.g. Default Mode Network and Fronto-parietal Network) | | |

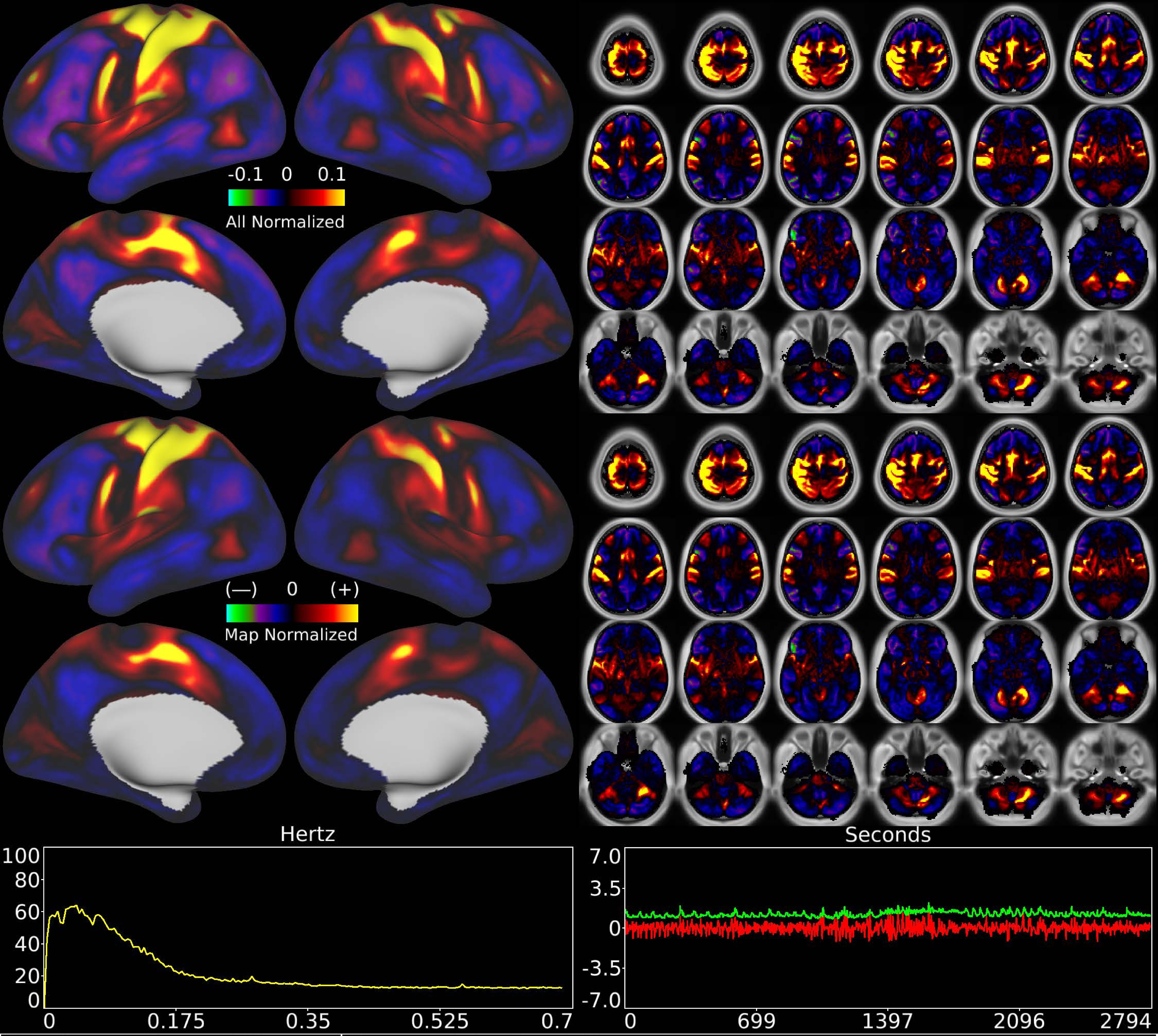


| | | |
|---|---|---|
| Number & Class: 7 Signal | Name: Right Hand Motor Network | |
| RVT Correlated: No | DVARS Dip Associated: Yes | Cross-Subject Variable: Yes |
| Single Subject: No | % Variance Explained: 2.97 | Globality Index: 0.16 |
| Task Component: 6 | Rest Component: 33 | Task Modulated: No |
| Rationale: Spatial map includes positive and negative patches that respect known somatotopic sensori-motor organization (Right Hand) | | |

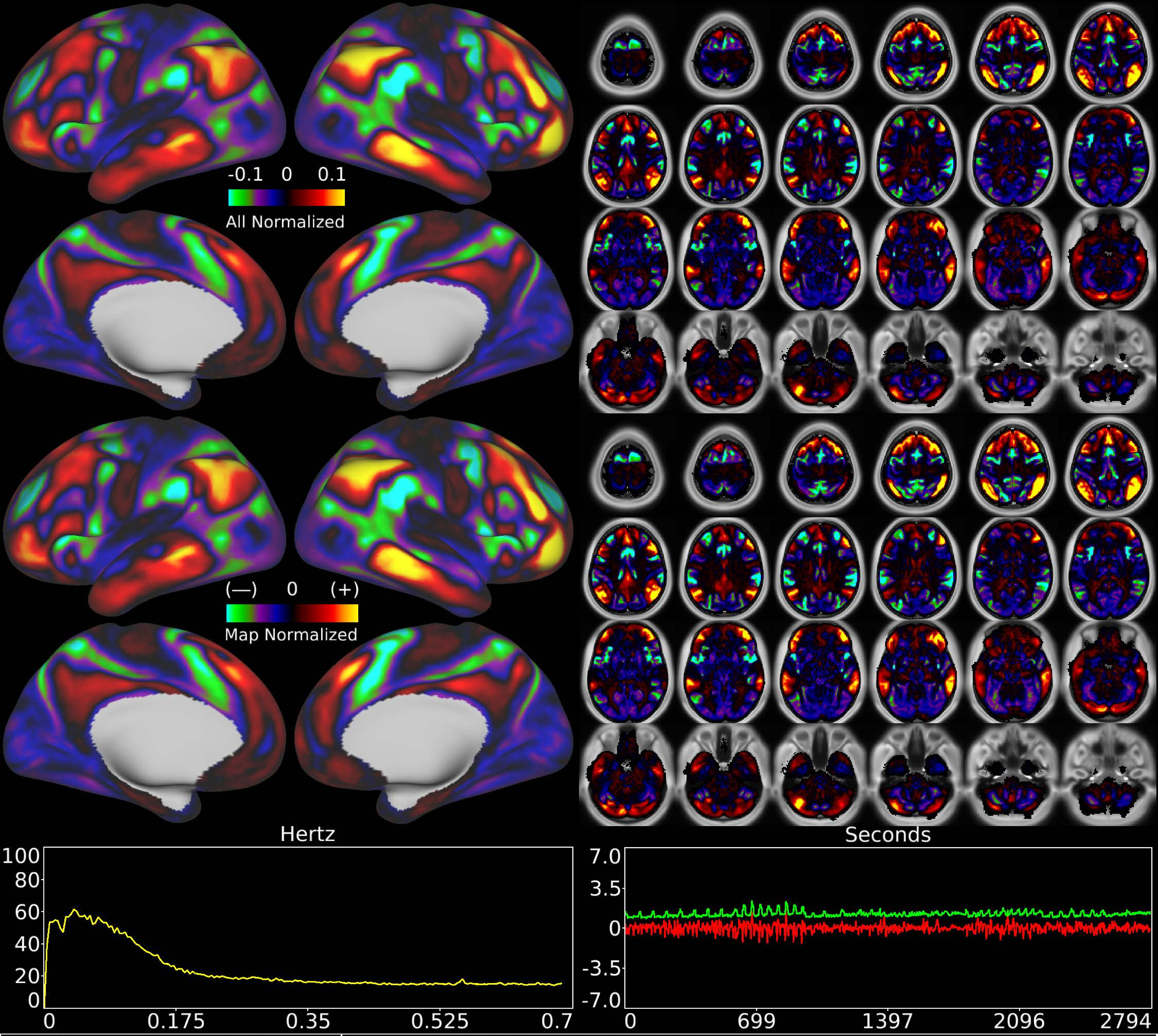


| | | |
|---|---|---|
| Number & Class: 8 Signal | Name: Subsidiary Default Mode | |
| RVT Correlated: No | DVARS Dip Associated: No | Cross-Subject Variable: No |
| Single Subject: No | % Variance Explained: 2.91 | Globality Index: 0.78 |
| Task Component: -9 | Rest Component: 9 | Task Modulated: No |

Rationale: Spatial map includes positive and negative patches that respect known RSNs (e.g. Default Mode Network)

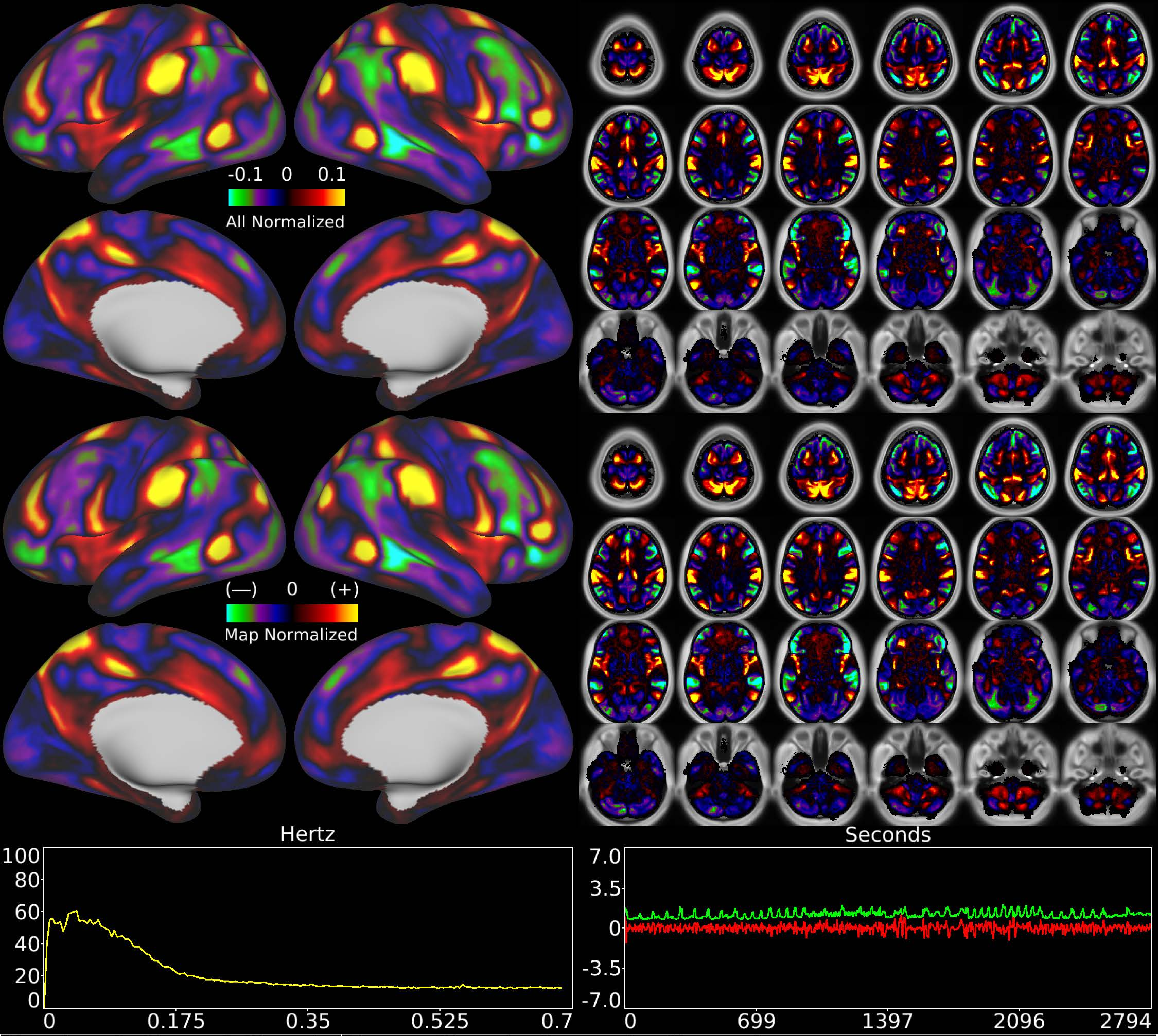


| | | |
|---|---|---|
| Number & Class: 9 Signal | Name: Cingulo-Opercular Network | |
| RVT Correlated: No | DVARS Dip Associated: No | Cross-Subject Variable: No |
| Single Subject: No | % Variance Explained: 2.8 | Globality Index: 0.57 |
| Task Component: 10 | Rest Component: 13 | Task Modulated: No |
| Rationale: Spatial map includes positive and negative patches that respect known RSNs (e.g. Cingulo-Opercular Network) | | |

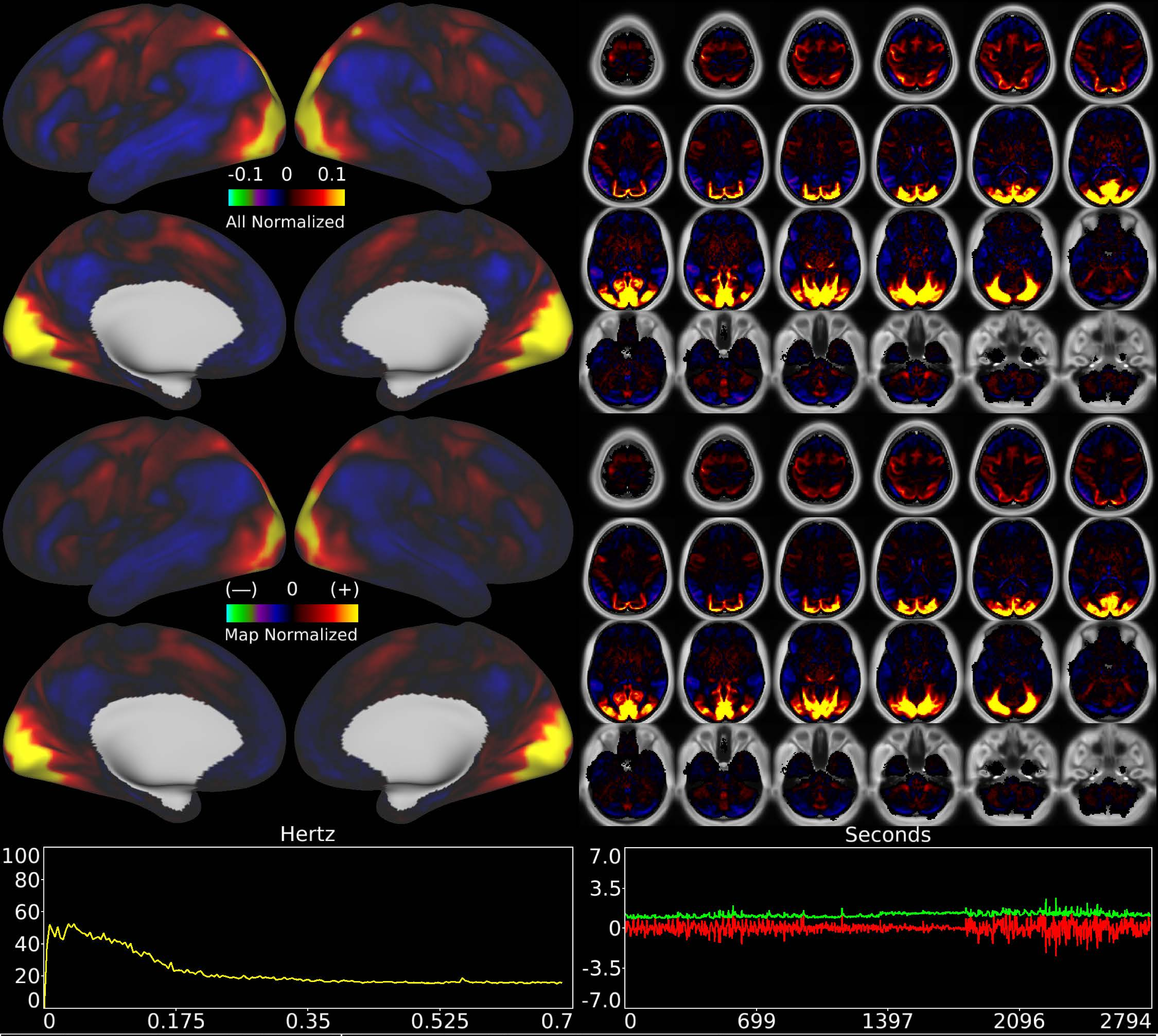


| | | |
|---|---|---|
| Number & Class: 10 Signal | Name: Pan-Visual (Paracentral) | |
| RVT Correlated: No | DVARS Dip Associated: No | Cross-Subject Variable: No |
| Single Subject: No | % Variance Explained: 2.6 | Globality Index: 0.19 |
| Task Component: 3+17 | Rest Component: No | Task Modulated: No |
| Rationale: Spatial map includes positive and negative patches that respect known retinotopic visual organization (Paracentral vs Foveal and Peripheral) | | |

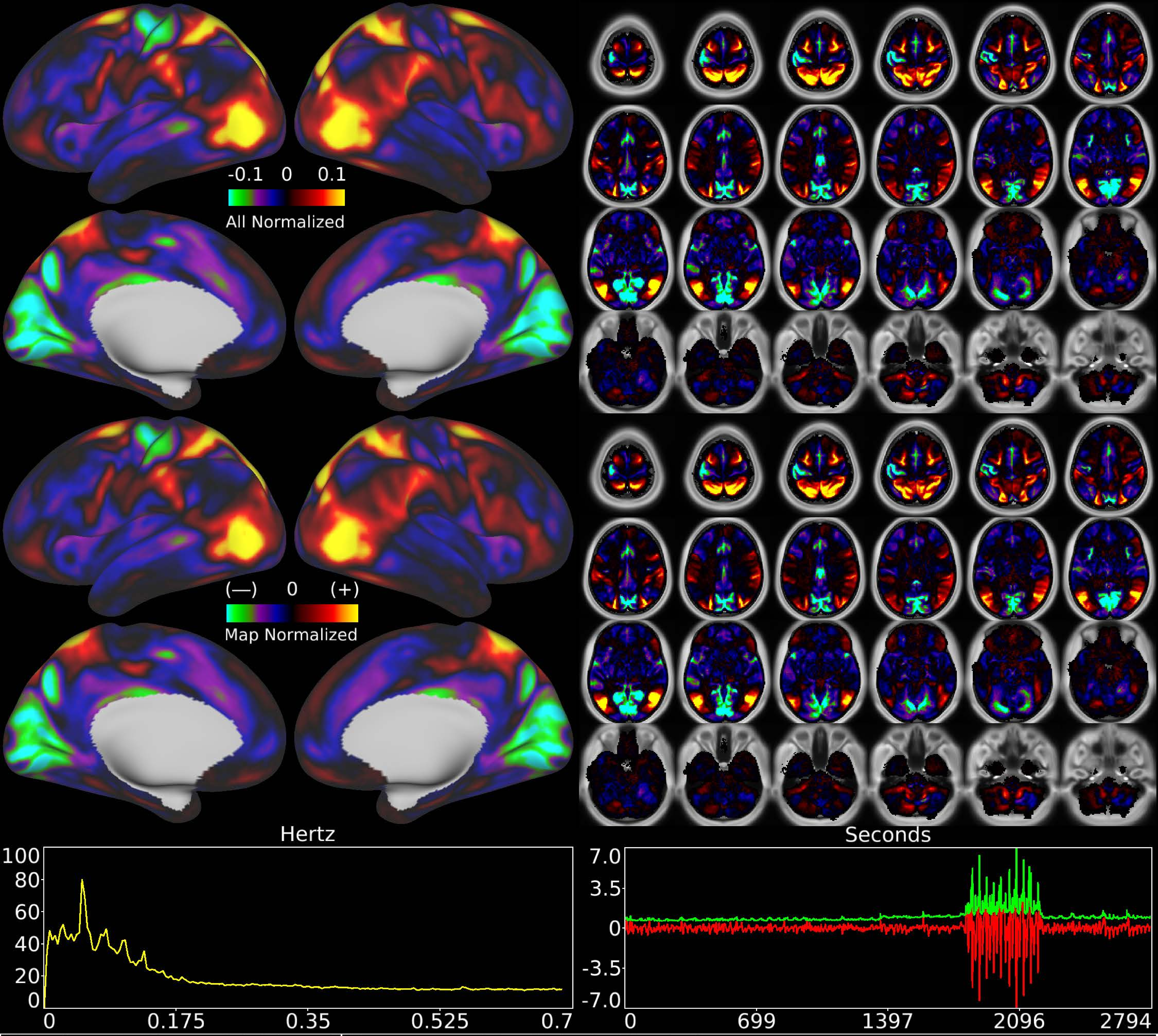


| | | |
|---|---|---|
| Number & Class: 11 Signal | Name: Residual Task (Social) | |
| RVT Correlated: No | DVARS Dip Associated: No | Cross-Subject Variable: No |
| Single Subject: No | % Variance Explained: 2.53 | Globality Index: 0.72 |
| Task Component: 2 | Rest Component: No | Task Modulated: Social |
| Rationale: Spatial map includes positive and negative patches that are correlated with a specific task design | | |

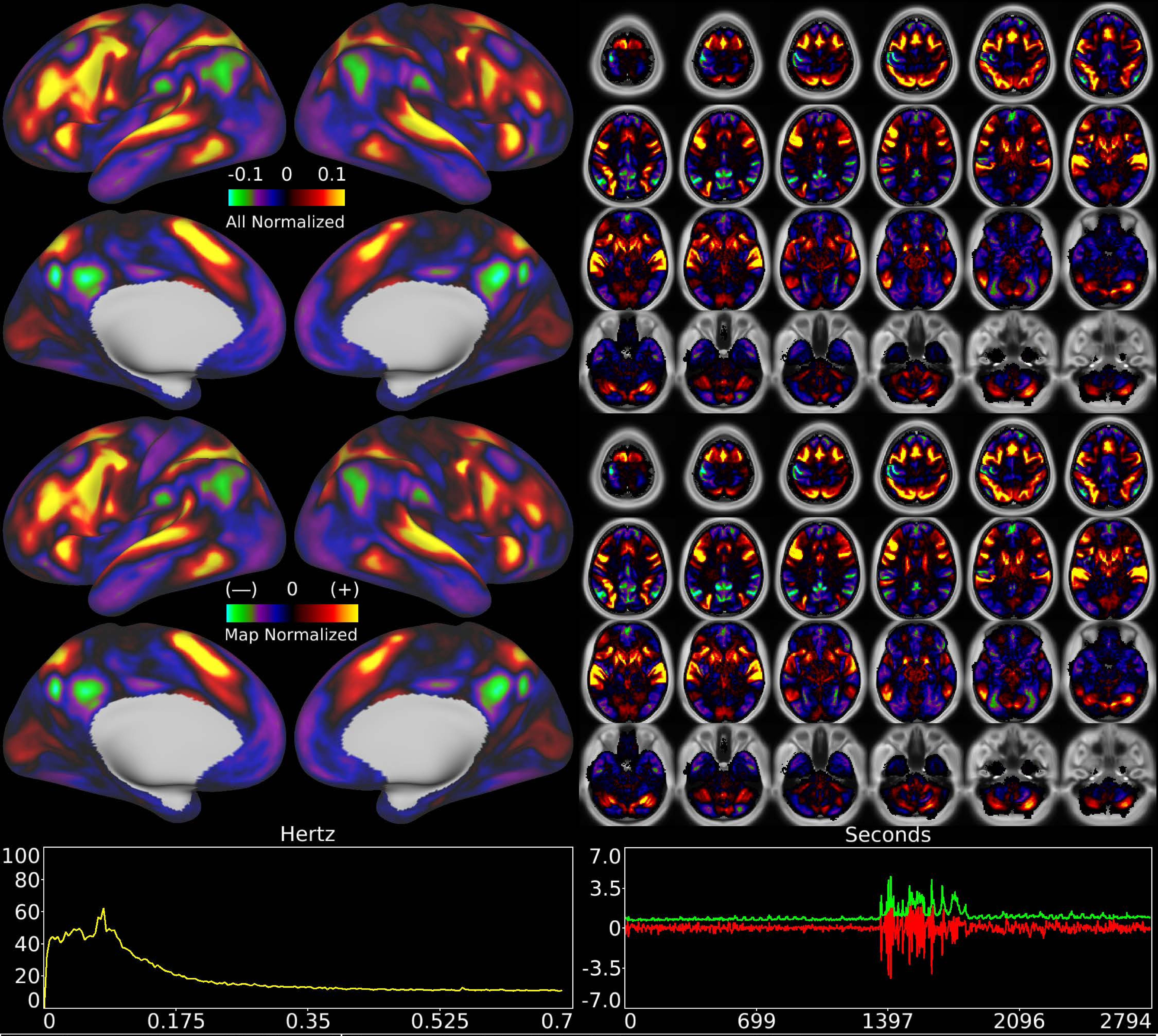


| | | |
|---|---|---|
| Number & Class: 12 Signal | Name: Residual Task (Language Math) | |
| RVT Correlated: No | DVARS Dip Associated: No | Cross-Subject Variable: No |
| Single Subject: No | % Variance Explained: 2.34 | Globality Index: 0.03 |
| Task Component: No | Rest Component: 12 | Task Modulated: Language |
| Rationale: Spatial map includes positive and negative patches that are correlated with a specific task design | | |

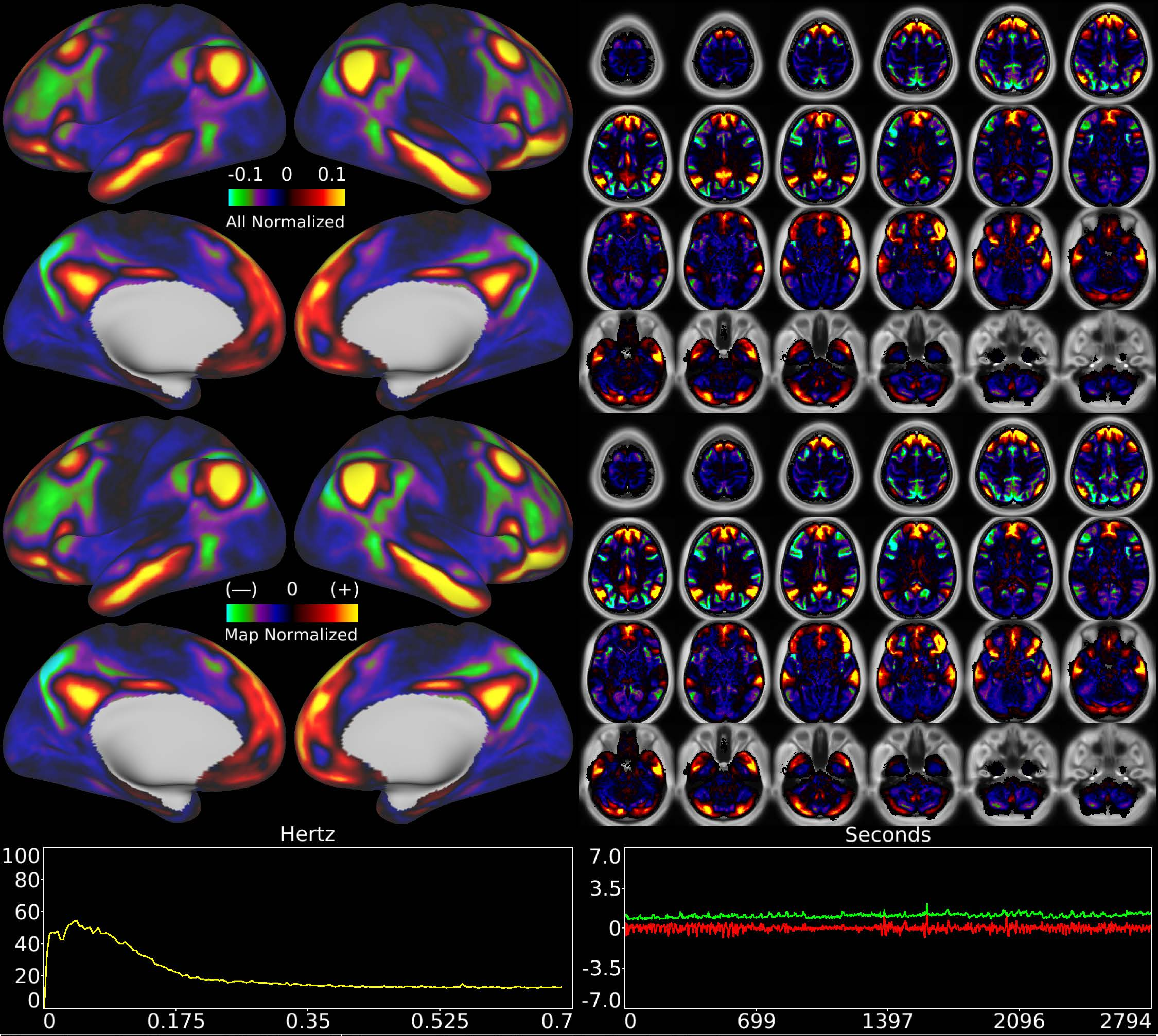


| | | |
|---|---|---|
| Number & Class: 13 Signal | Name: Subsidiary Default Mode | |
| RVT Correlated: No | DVARS Dip Associated: No | Cross-Subject Variable: No |
| Single Subject: No | % Variance Explained: 2.34 | Globality Index: 1.52 |
| Task Component: No | Rest Component: -21 | Task Modulated: No |
| Rationale: Spatial map includes positive and negative patches that respect known RSNs (e.g. Default Mode Network) | | |

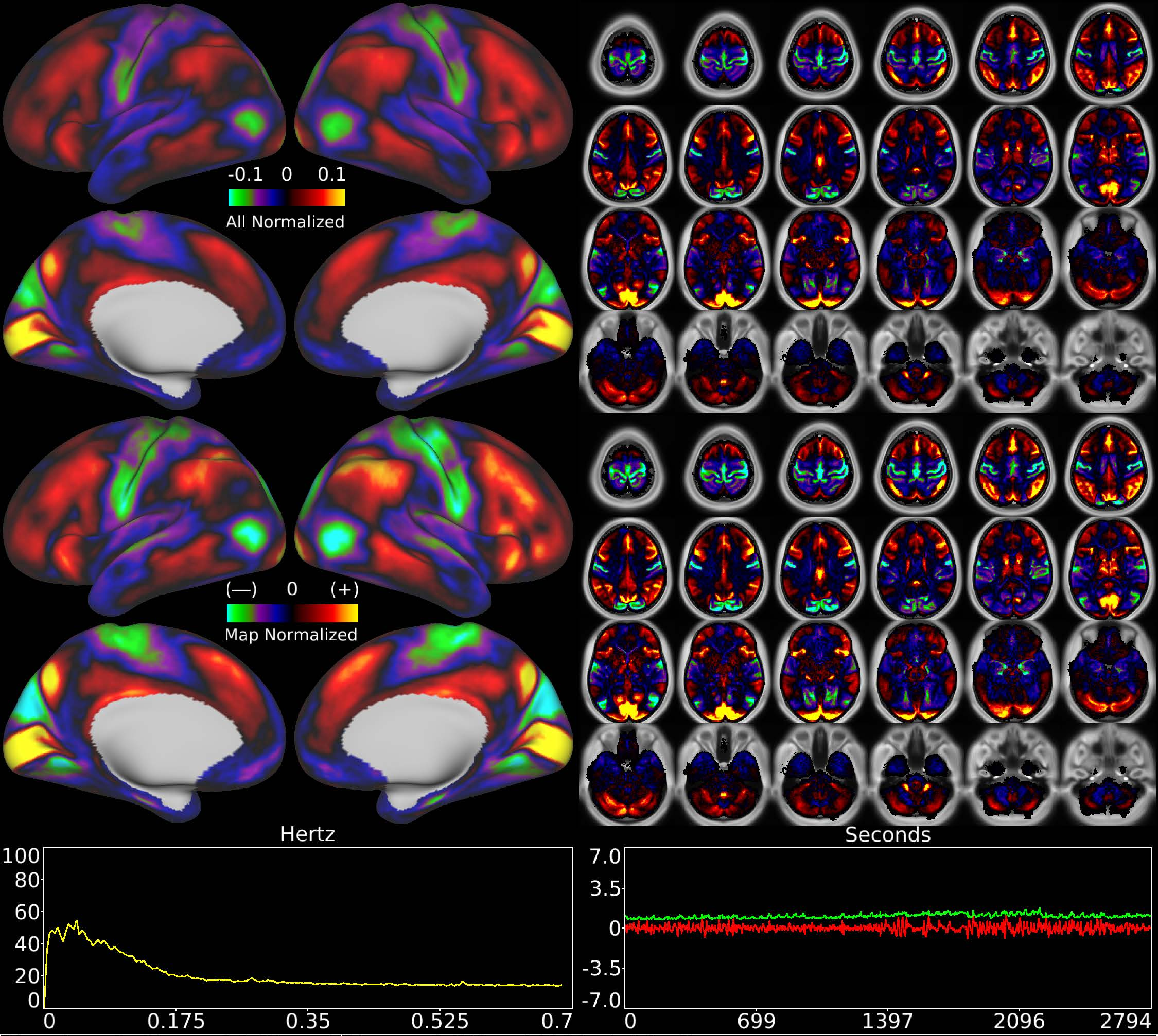


| | | |
|---|---|---|
| Number & Class: 14 Signal | Name: LGN to V1 Variable Component | |
| RVT Correlated: No | DVARS Dip Associated: No | Cross-Subject Variable: Yes |
| Single Subject: No | % Variance Explained: 2.27 | Globality Index: 0.02 |
| Task Component: 14 | Rest Component: 2 | Task Modulated: No |
| Rationale: Spatial map includes positive and negative patches that respect known areal boundaries (e.g. V1 and V2) | | |

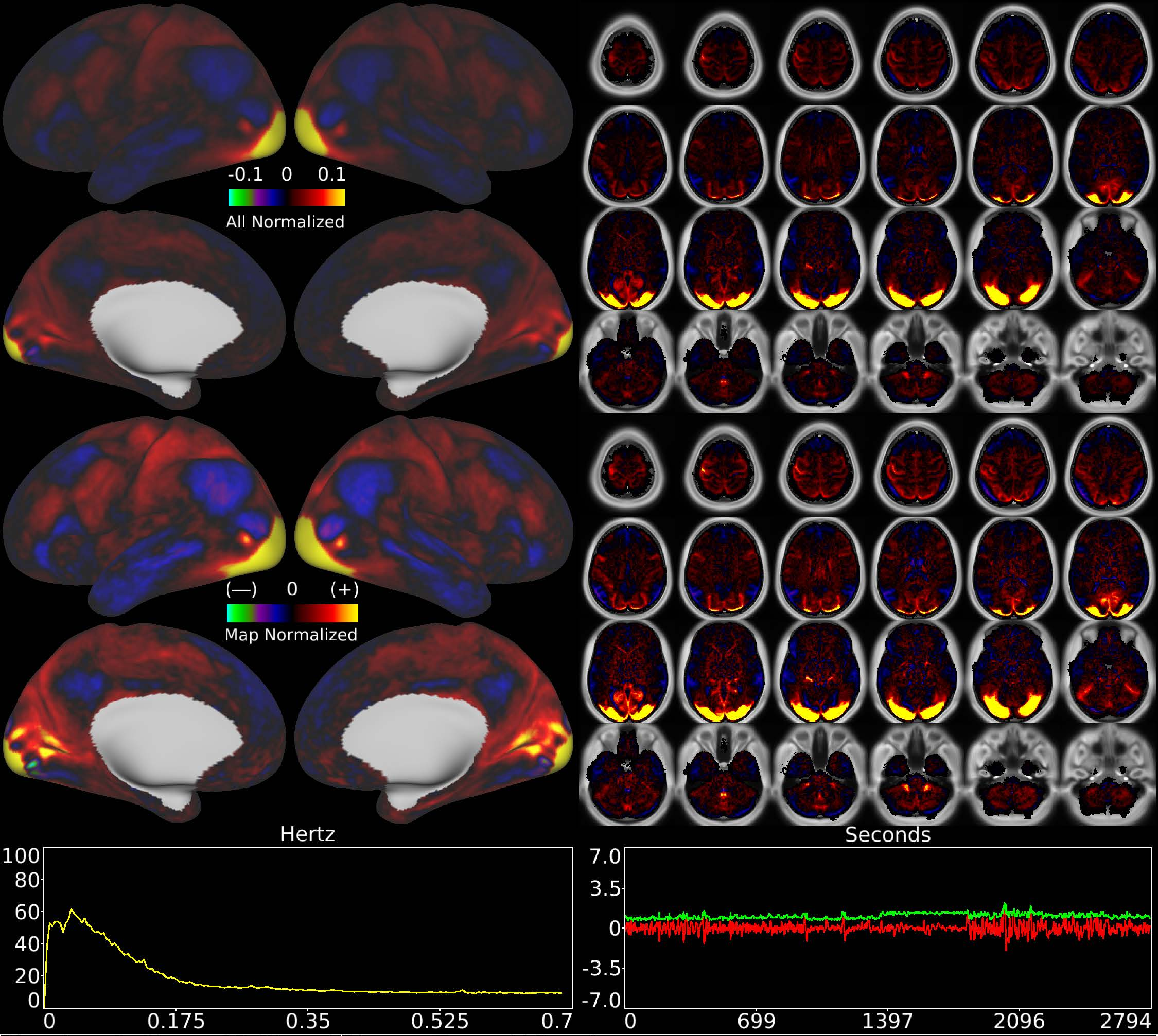


| | | |
|---|---|---|
| Number & Class: 15 Signal | Name: Foveal Visual | |
| RVT Correlated: No | DVARS Dip Associated: No | Cross-Subject Variable: Yes |
| Single Subject: No | % Variance Explained: 2.26 | Globality Index: 1.59 |
| Task Component: 34 | Rest Component: 41 | Task Modulated: Language |

Rationale: Spatial map includes positive and negative patches that respect known retinotopic visual organization (Foveal vs ParaCentral)

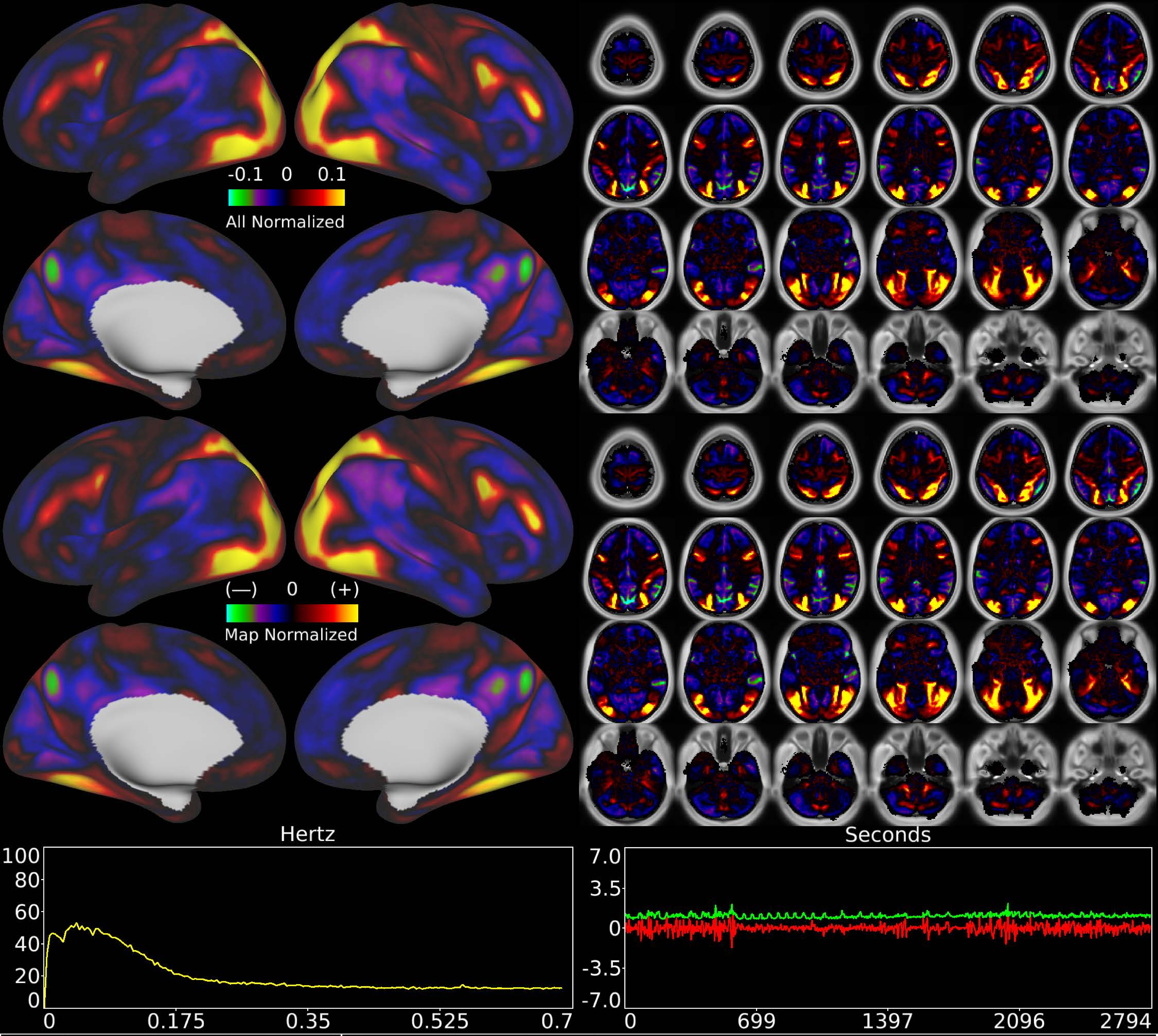


| | | |
|---|---|---|
| Number & Class: 16 Signal | Name: Higher Visual | |
| RVT Correlated: No | DVARS Dip Associated: No | Cross-Subject Variable: No |
| Single Subject: No | % Variance Explained: 2.21 | Globality Index: 0.3 |
| Task Component: No | Rest Component: 18 | Task Modulated: No |
| Rationale: Spatial map includes positive and negative patches that respect known RSNs (e.g. Dorsal Attention Network) | | |

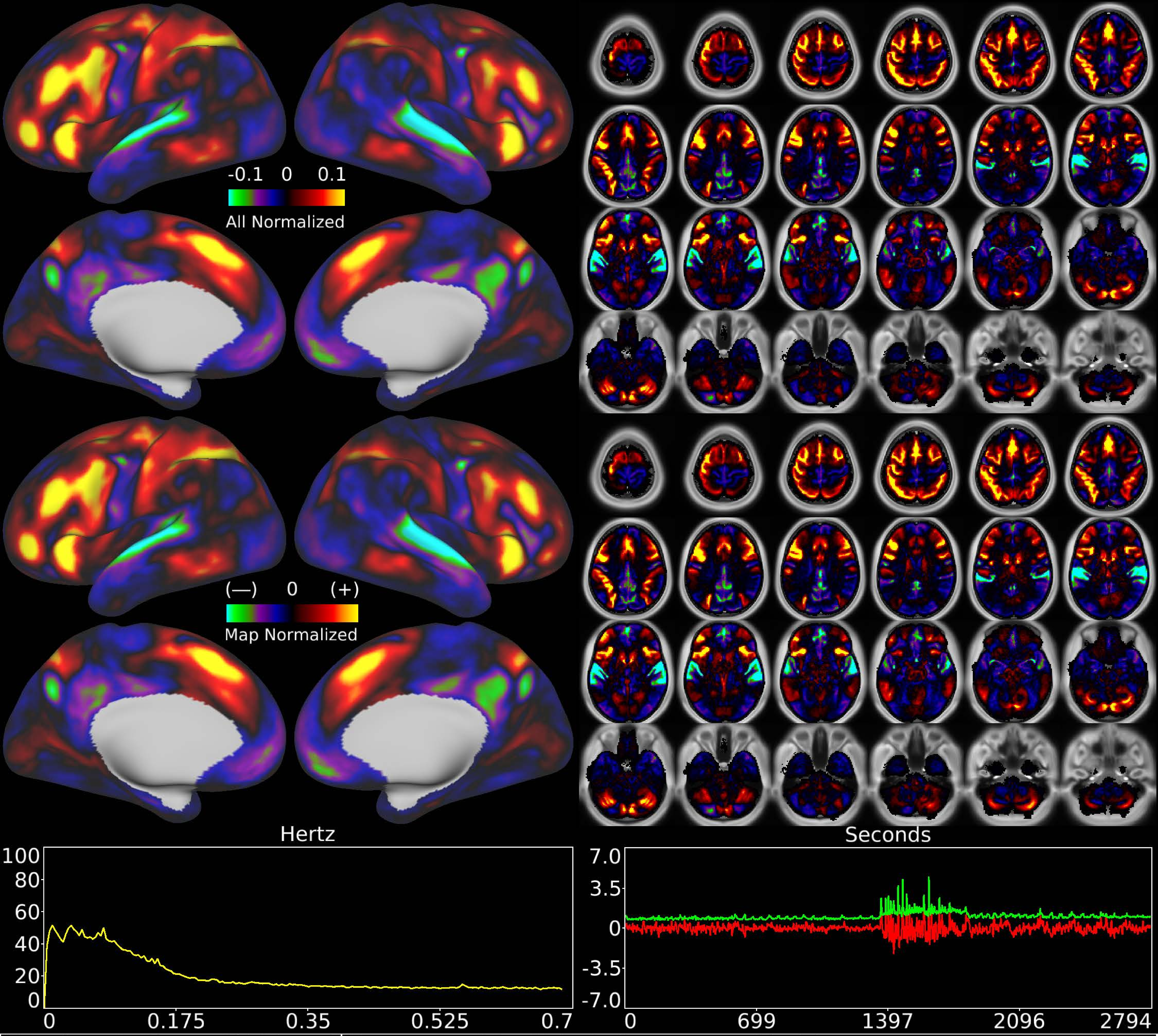


| | | |
|---|---|---|
| Number & Class: 17 Signal | Name: Residual Task (Language Math) | |
| RVT Correlated: No | DVARS Dip Associated: No | Cross-Subject Variable: No |
| Single Subject: No | % Variance Explained: 2.17 | Globality Index: 0 |
| Task Component: 24 | Rest Component: 12 | Task Modulated: Language |
| Rationale: Spatial map includes positive and negative patches that are correlated with a specific task design | | |

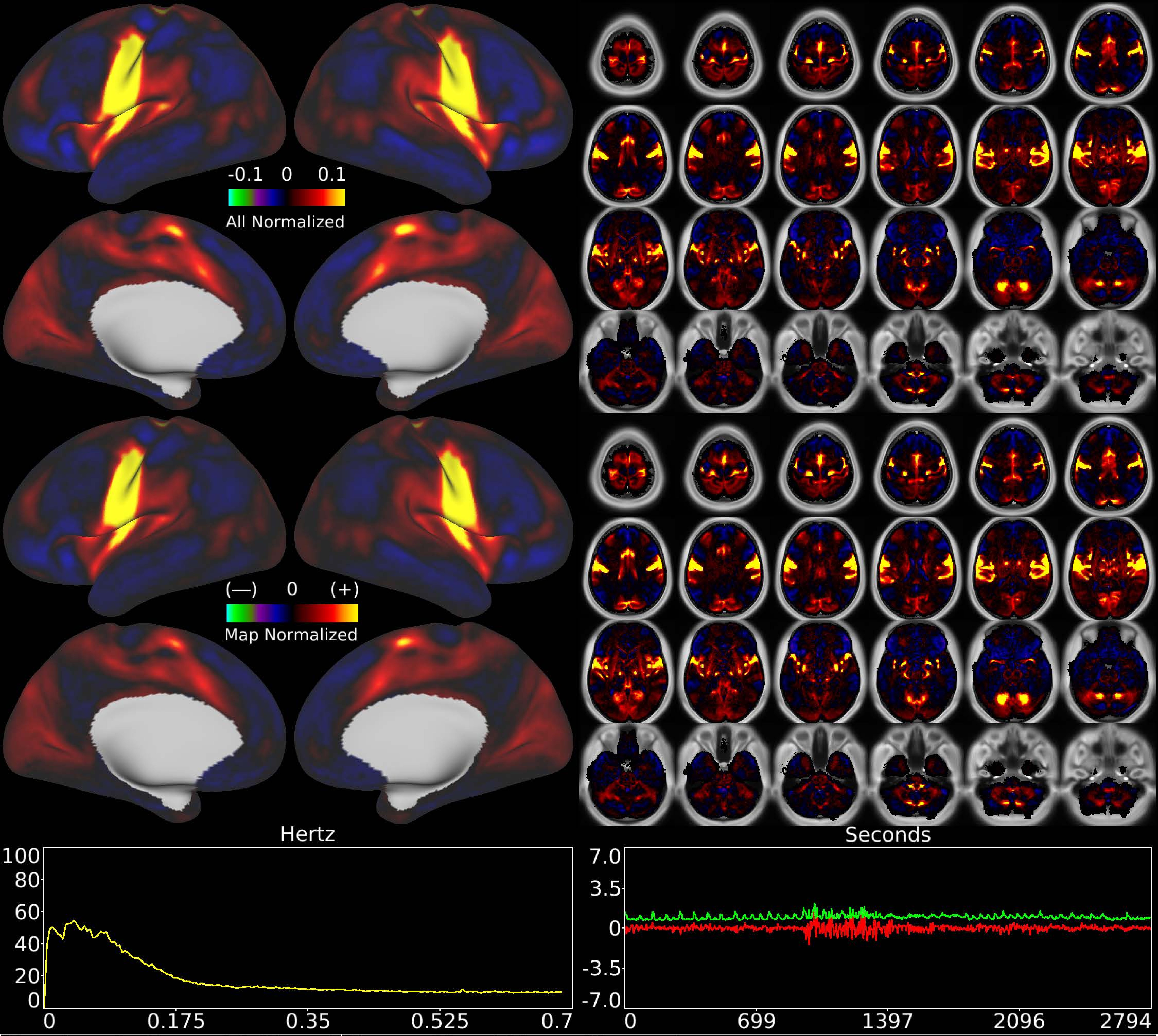


| | | |
|---|---|---|
| Number & Class: 18 Signal | Name: Head Motor Network | |
| RVT Correlated: No | DVARS Dip Associated: Yes | Cross-Subject Variable: Yes |
| Single Subject: No | % Variance Explained: 2.13 | Globality Index: 0.81 |
| Task Component: 16 | Rest Component: 27 | Task Modulated: No |
| Rationale: Spatial map includes positive and negative patches that respect known somatotopic sensori-motor organization (Face) | | |

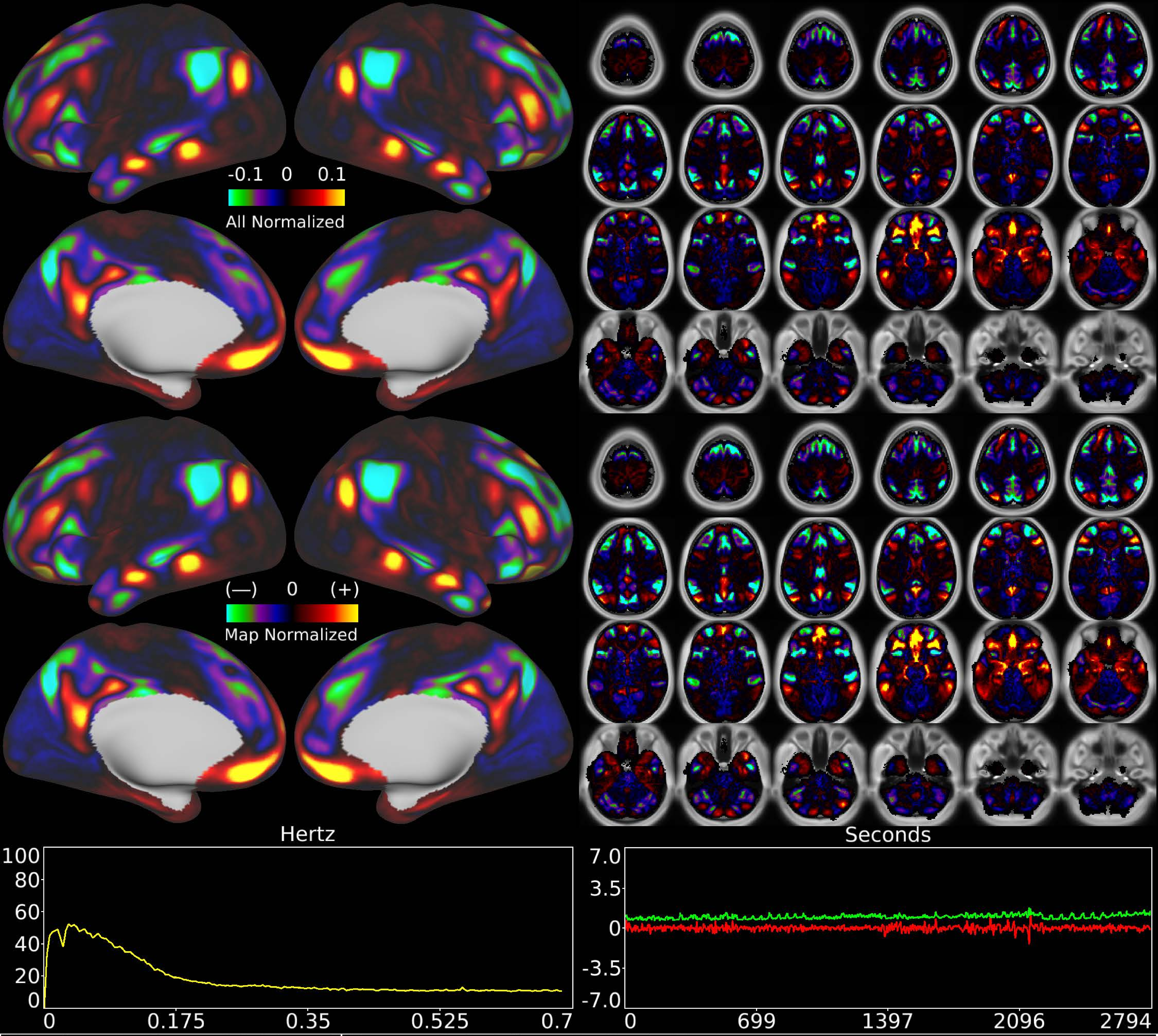


| | | |
|---|---|---|
| Number & Class: 19 Signal | Name: Subsidiary Default Mode | |
| RVT Correlated: No | DVARS Dip Associated: No | Cross-Subject Variable: No |
| Single Subject: No | % Variance Explained: 1.93 | Globality Index: 0.52 |
| Task Component: No | Rest Component: 15 | Task Modulated: No |
| Rationale: Spatial map includes positive and negative patches that respect known RSNs (e.g. Default Mode Network) | | |

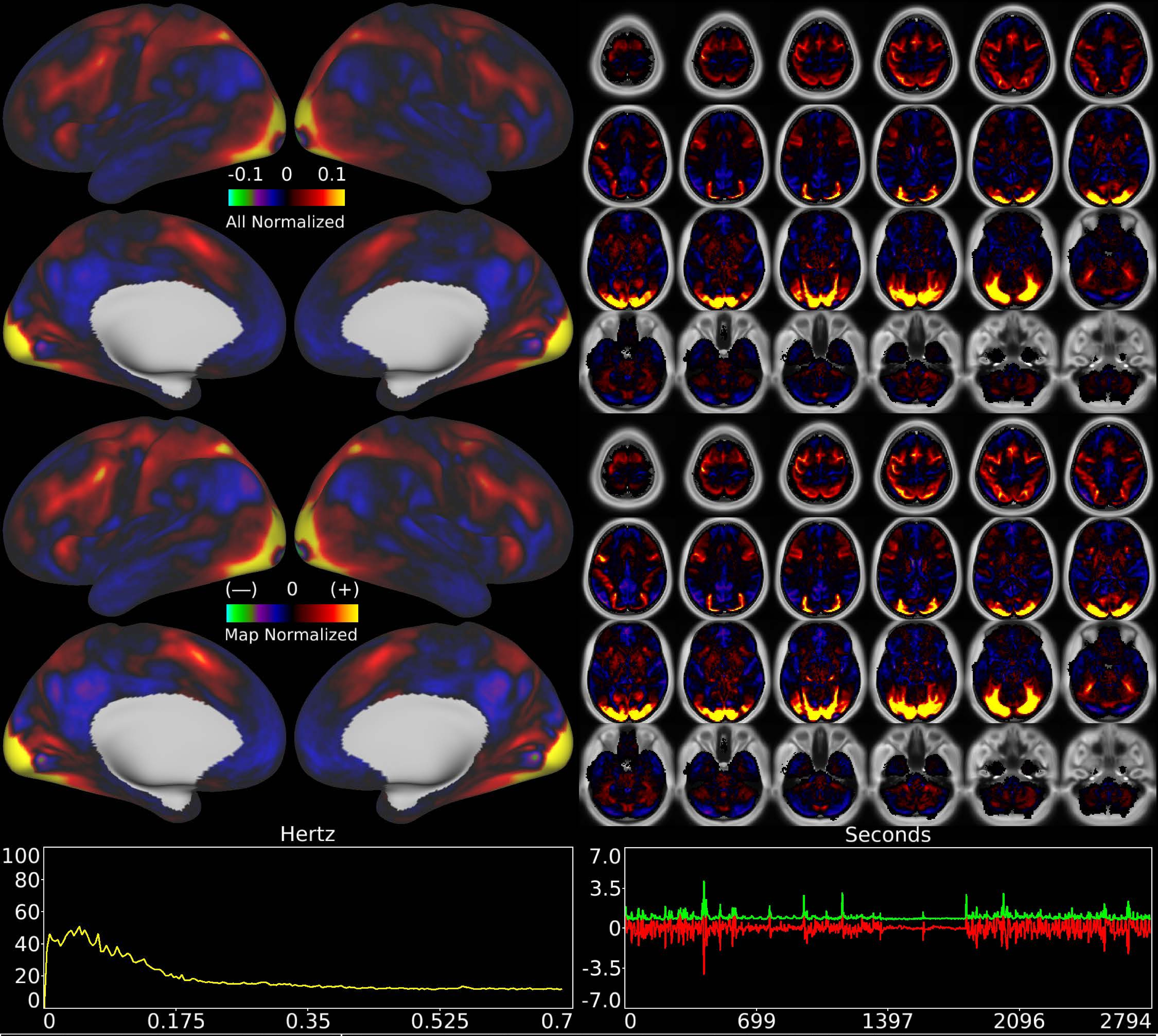


| | | |
|---|---|---|
| Number & Class: 20 Signal | Name: Pan-Visual (Parafoveal) Plus R Hand Motor | |
| RVT Correlated: No | DVARS Dip Associated: Yes | Cross-Subject Variable: No |
| Single Subject: No | % Variance Explained: 1.84 | Globality Index: 0.32 |
| Task Component: 7+18 | Rest Component: No | Task Modulated: No |
| Rationale: Spatial map includes positive and negative patches that respect known retinotopic visual organization (Paracentral vs Foveal and Peripheral) | | |

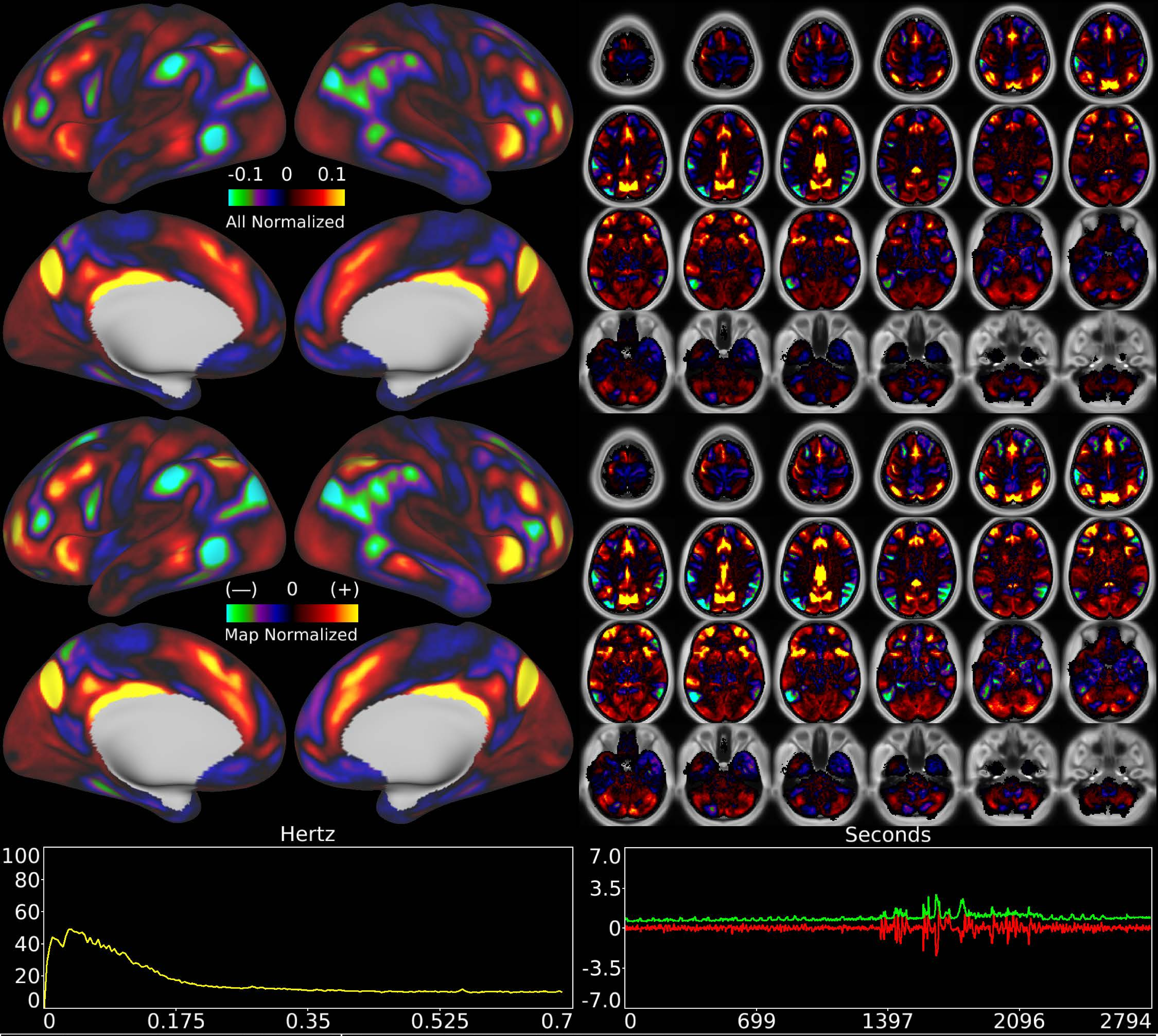


| | | |
|---|---|---|
| Number & Class: 21 Signal | Name: POS2 + RSC vs Unknown | |
| RVT Correlated: No | DVARS Dip Associated: No | Cross-Subject Variable: No |
| Single Subject: No | % Variance Explained: 1.74 | Globality Index: 0.7 |
| Task Component: 13 | Rest Component: 37 | Task Modulated: Language + Social |
| Rationale: Spatial map includes positive and negative patches that respect known areal boundaries (e.g. POS2 and RSC) | | |

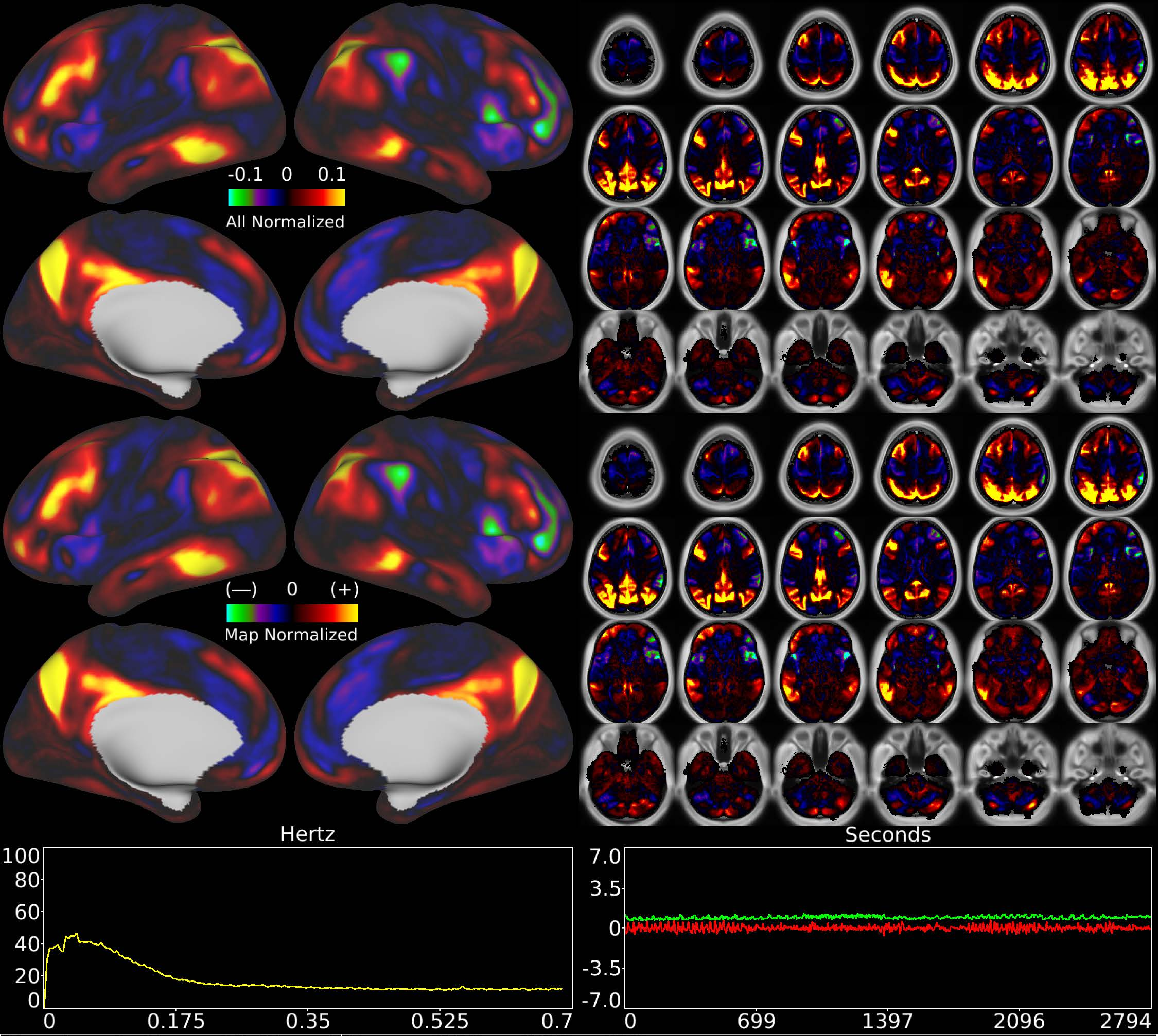


| | | |
|---|---|---|
| Number & Class: 22 Signal | Name: POS + RSC + IPS | |
| RVT Correlated: No | DVARS Dip Associated: No | Cross-Subject Variable: No |
| Single Subject: No | % Variance Explained: 1.64 | Globality Index: 0.42 |
| Task Component: No | Rest Component: No | Task Modulated: No |
| Rationale: Spatial map includes positive and negative patches that respect known areal boundaries (e.g. POS2 and RSC) | | |

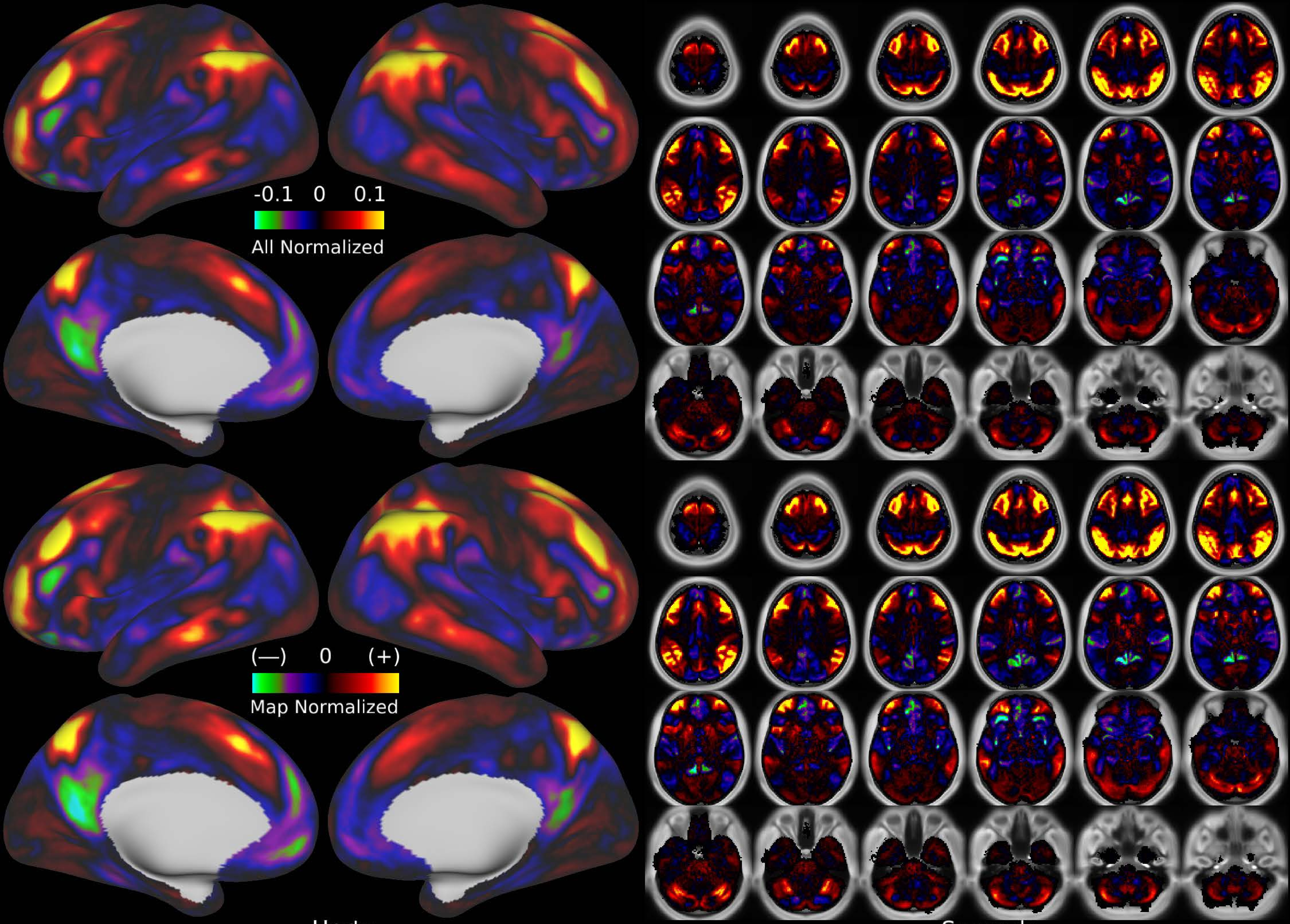


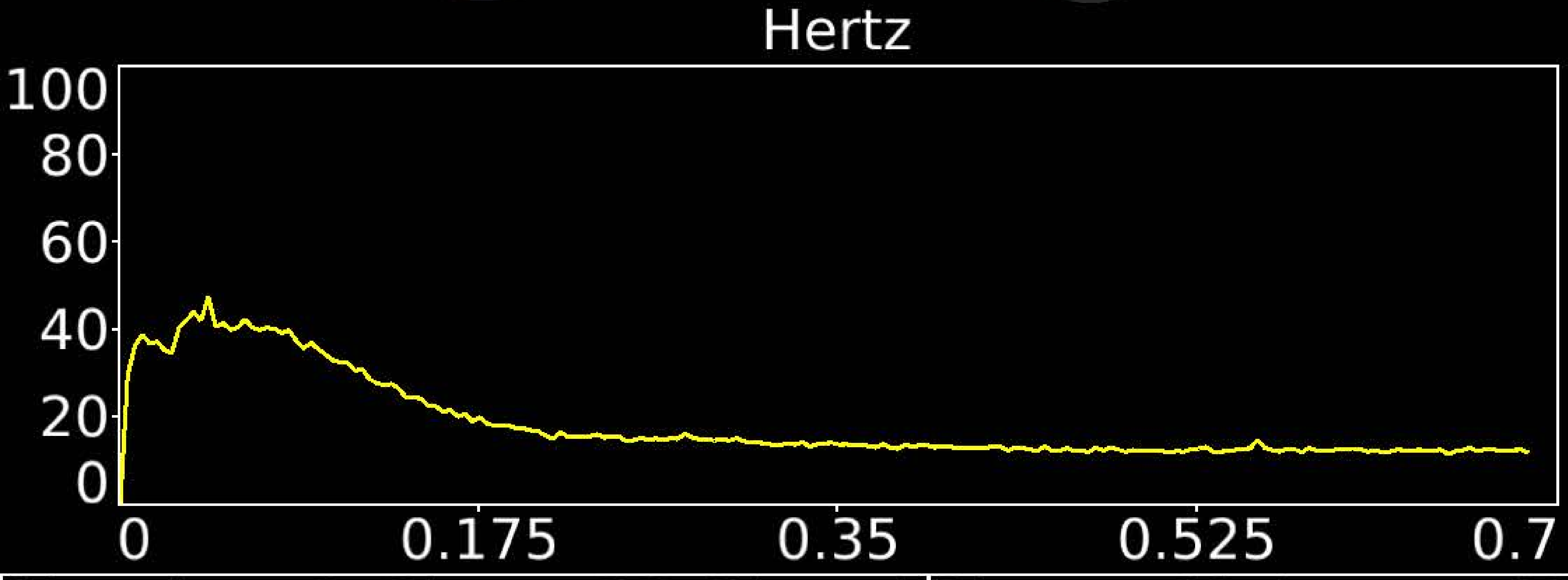


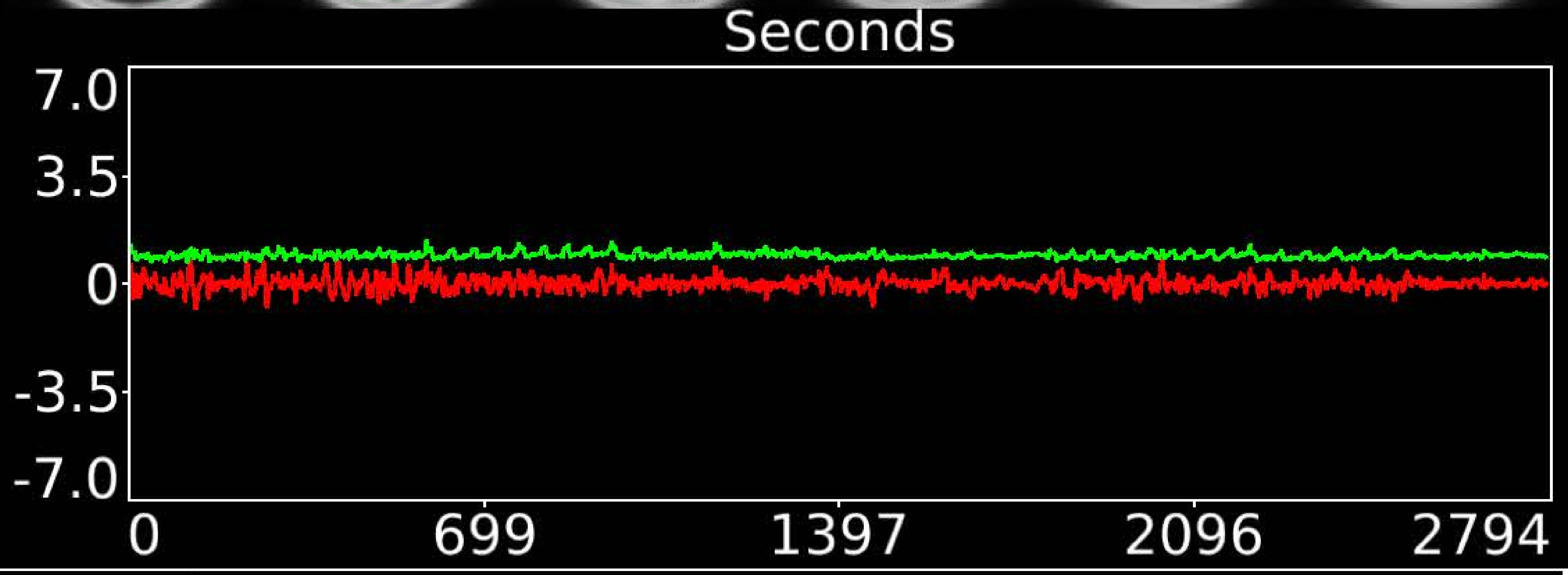


| | | |
|---|---|---|
| Number & Class: 23 Signal | Name: Unknown Network | |
| RVT Correlated: No | DVARS Dip Associated: No | Cross-Subject Variable: No |
| Single Subject: No | % Variance Explained: 1.62 | Globality Index: 0.49 |
| Task Component: No | Rest Component: 12 | Task Modulated: No |
| Rationale: Spatial map includes positive and negative patches that respect known RSNs (e.g. Fronto-Parietal Network) | | |

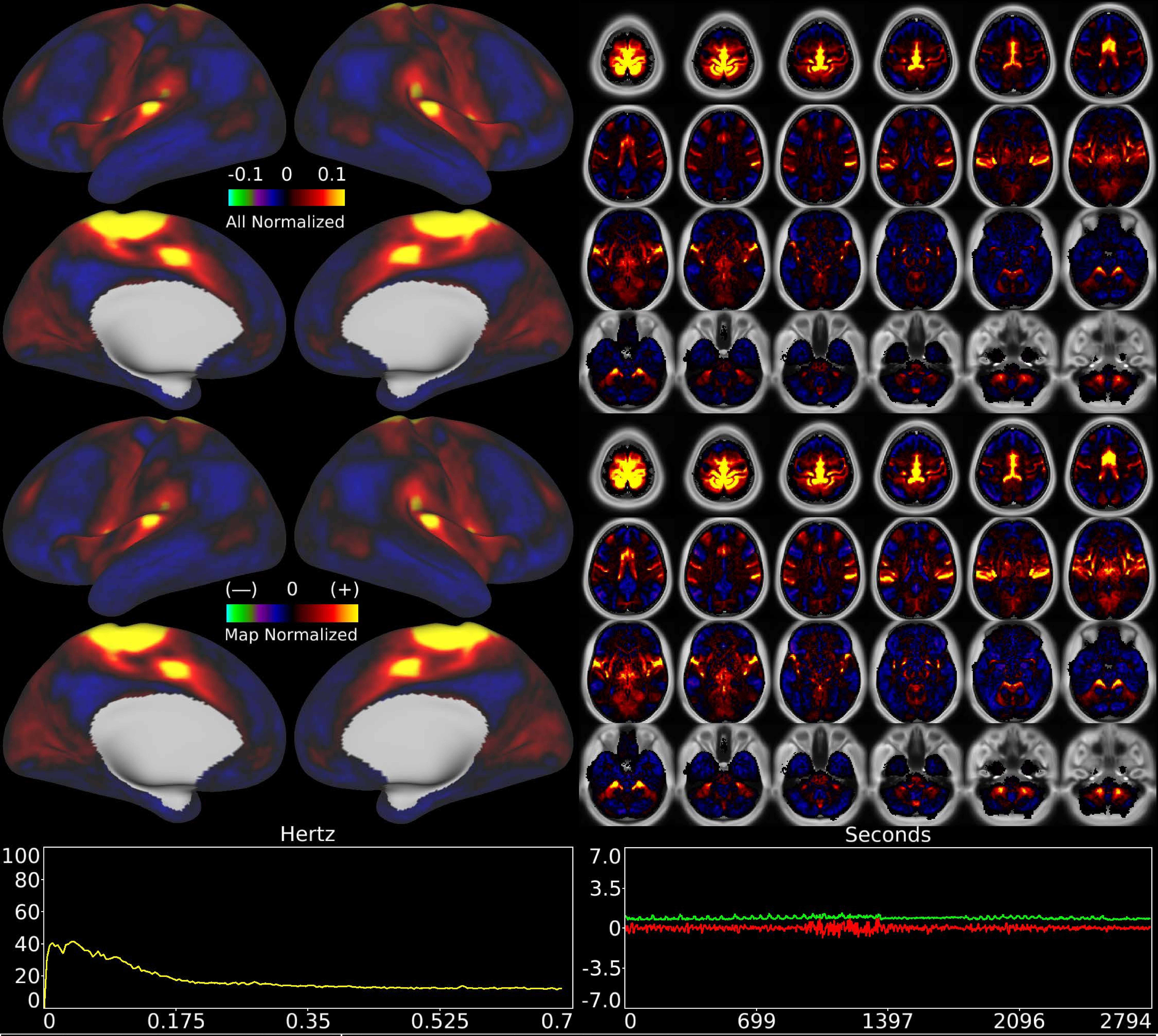


| | | |
|---|---|---|
| Number & Class: 24 Signal | Name: Feet Motor Network | |
| RVT Correlated: No | DVARS Dip Associated: Yes | Cross-Subject Variable: Yes |
| Single Subject: No | % Variance Explained: 1.61 | Globality Index: 0.02 |
| Task Component: 19 | Rest Component: 48 | Task Modulated: No |
| Rationale: Spatial map includes positive and negative patches that respect known somatotopic sensori-motor organization (Feet) | | |

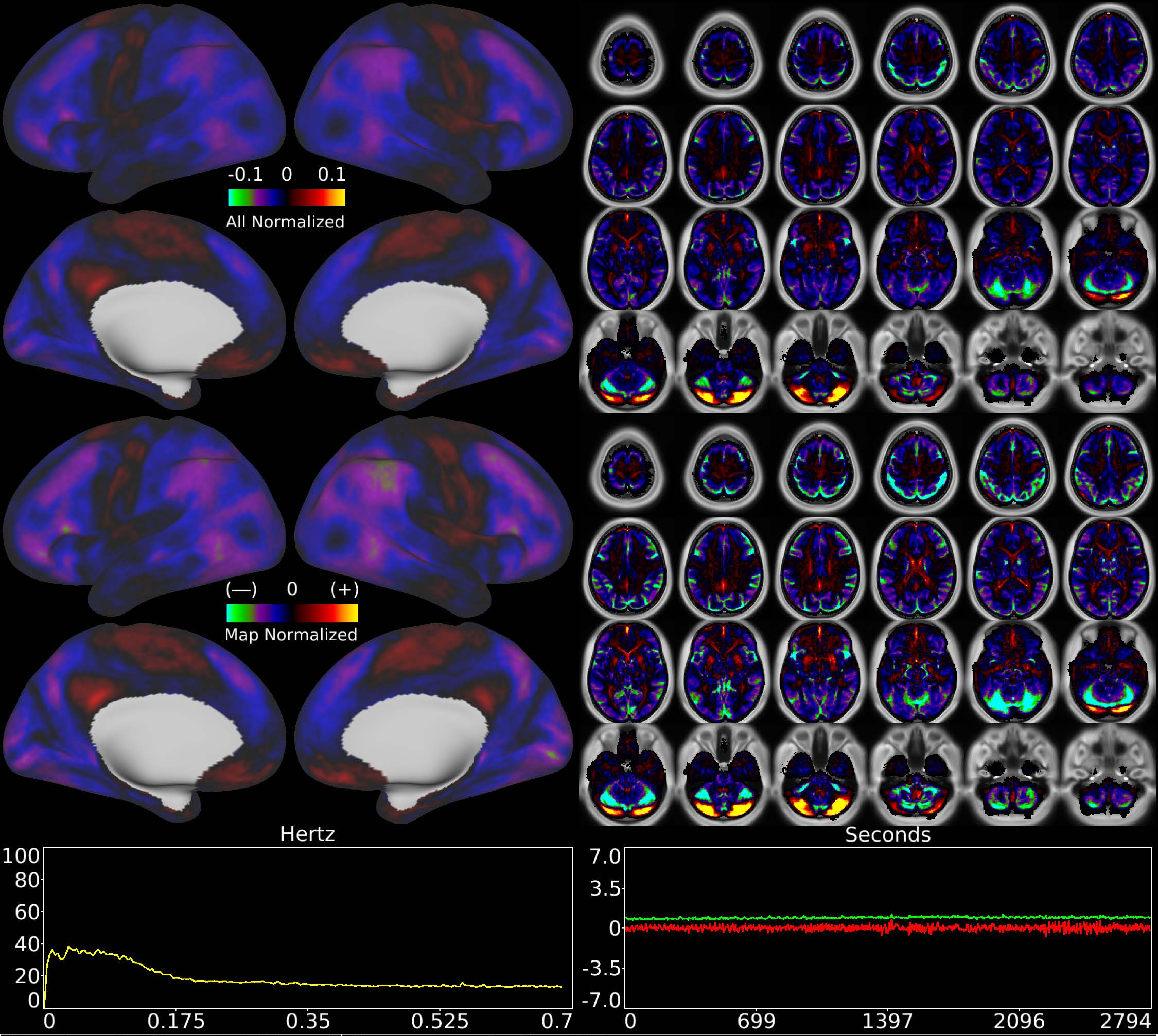


| | | |
|---|---|---|
| Number & Class: 25 Signal | Name: Cerebellar Unknown | |
| RVT Correlated: No | DVARS Dip Associated: No | Cross-Subject Variable: Yes |
| Single Subject: No | % Variance Explained: 1.57 | Globality Index: 1.74 |
| Task Component: 35 | Rest Component: No | Task Modulated: No |
| Rationale: Spatial map includes positive and negative patches that respect known RSN boundaries in the cerebellum | | |

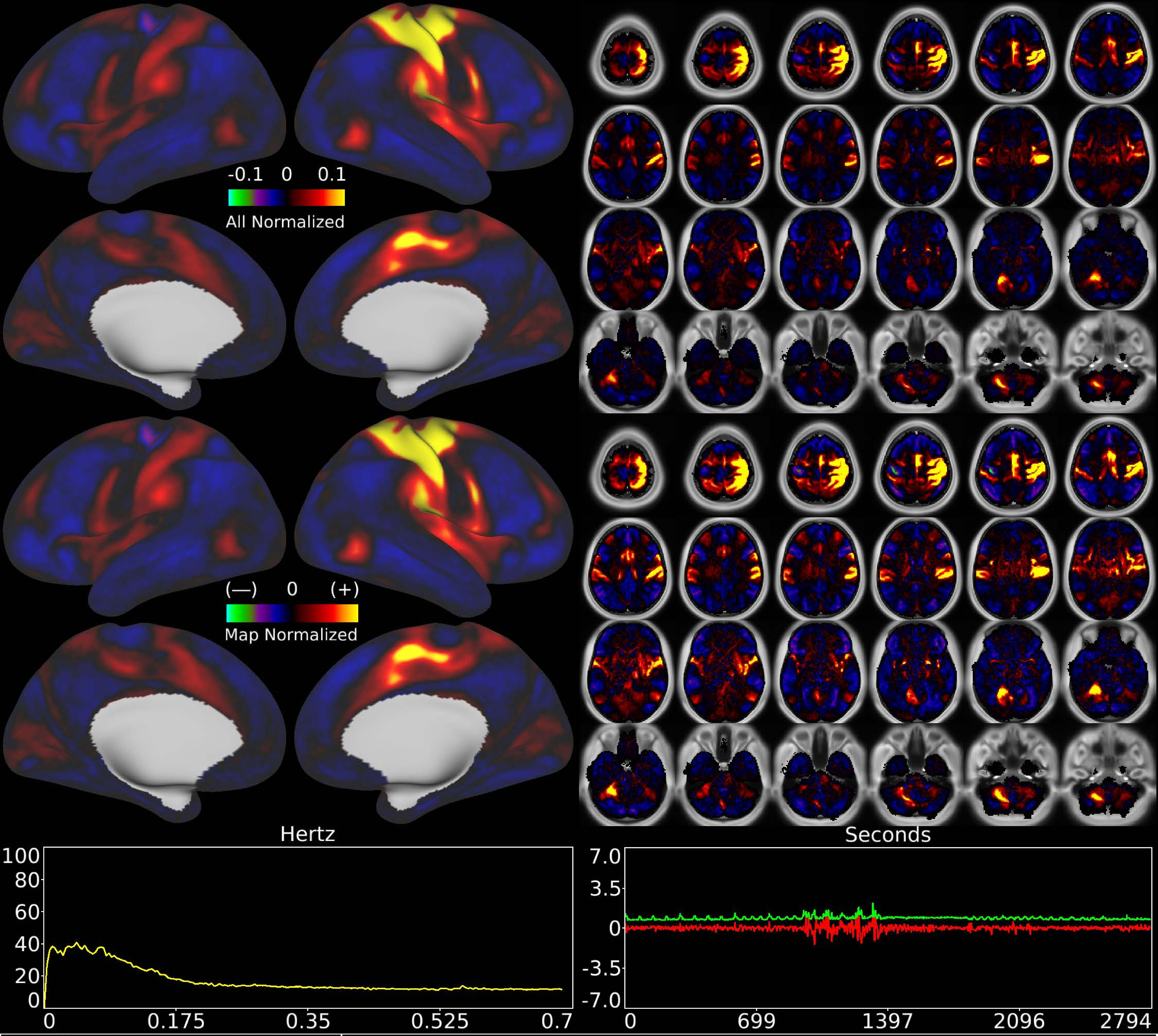


| | | |
|---|---|---|
| Number & Class: 26 Signal | Name: Left Hand Motor Network | |
| RVT Correlated: No | DVARS Dip Associated: Yes | Cross-Subject Variable: Yes |
| Single Subject: Yes | % Variance Explained: 1.55 | Globality Index: 0.26 |
| Task Component: 31 | Rest Component: 40 | Task Modulated: No |

Rationale: Spatial map includes positive and negative patches that respect known somatotopic sensori-motor organization (Left Hand)

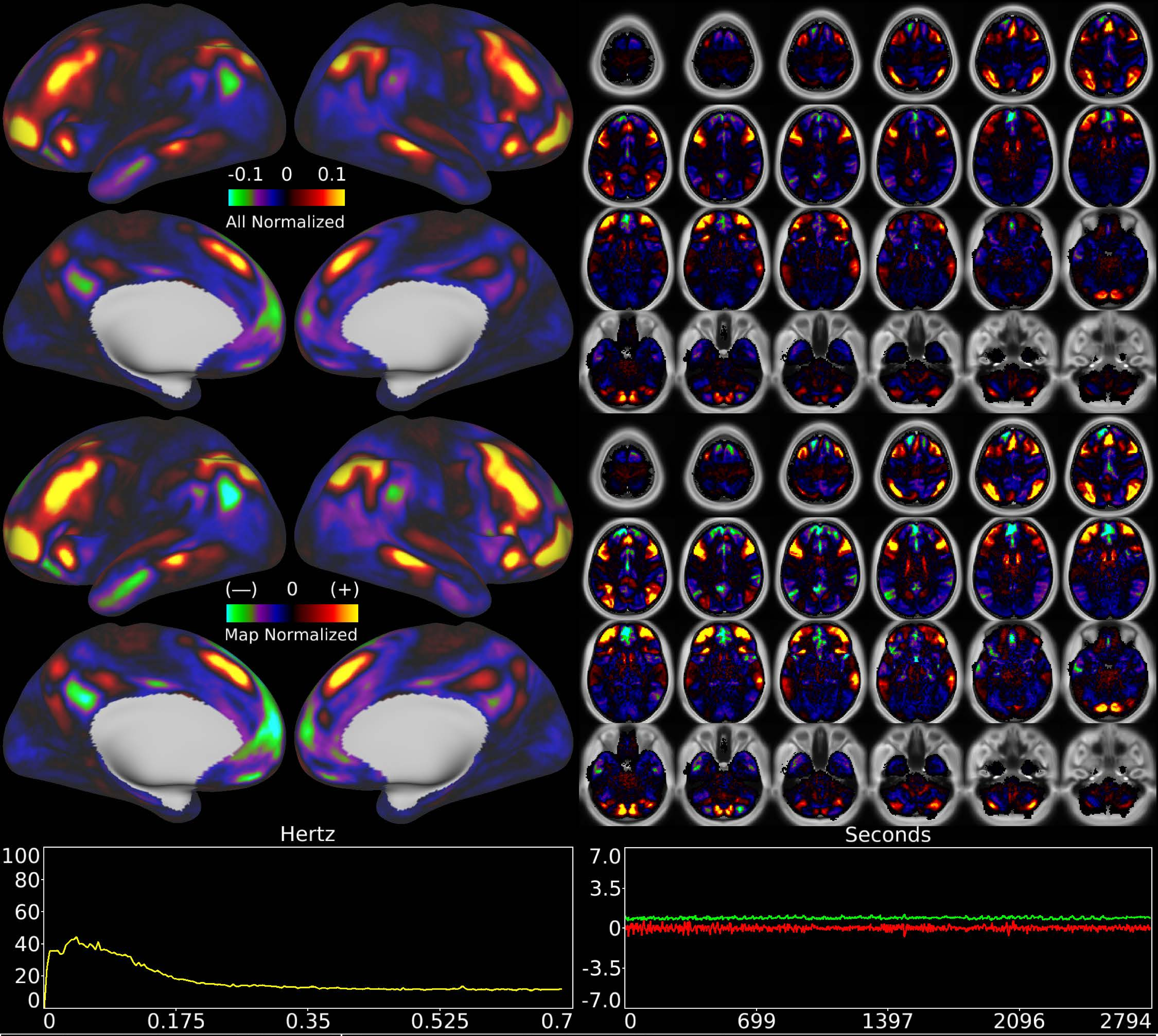


| | | |
|---|---|---|
| Number & Class: 27 Signal | Name: Fronto-Parietal Bilateral | |
| RVT Correlated: No | DVARS Dip Associated: No | Cross-Subject Variable: No |
| Single Subject: No | % Variance Explained: 1.49 | Globality Index: 0.79 |
| Task Component: No | Rest Component: No | Task Modulated: No |
| Rationale: Spatial map includes positive and negative patches that respect known RSNs (e.g. Fronto-Parietal Network) | | |

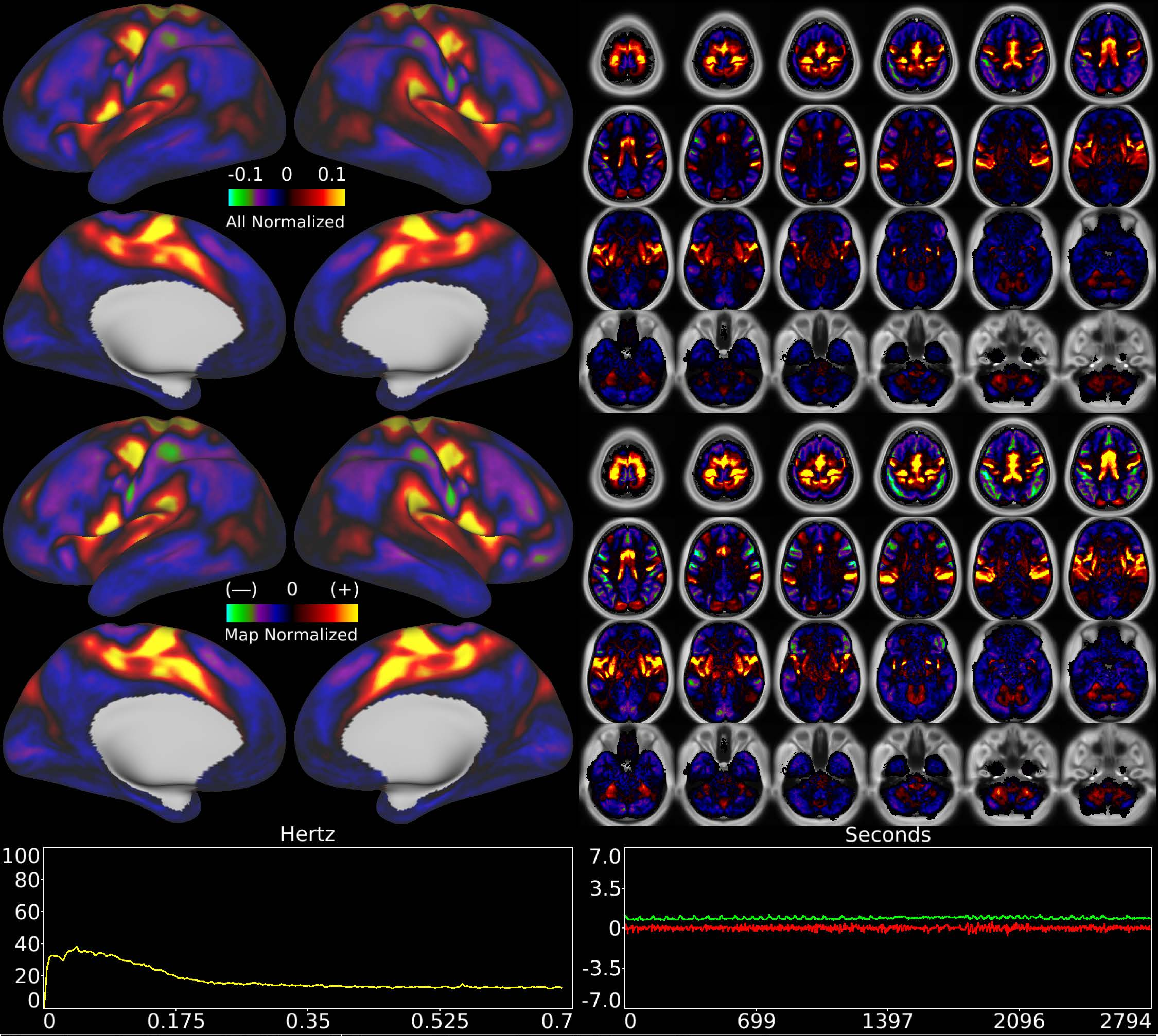


| | | |
|---|---|---|
| Number & Class: 28 Signal | Name: Eye + Trunk Motor Network | |
| RVT Correlated: No | DVARS Dip Associated: No | Cross-Subject Variable: No |
| Single Subject: No | % Variance Explained: 1.48 | Globality Index: 0.69 |
| Task Component: 29 | Rest Component: 39 | Task Modulated: No |
| Rationale: Spatial map includes positive and negative patches that respect known somatotopic sensori-motor organization (Eye and Trunk) | | |

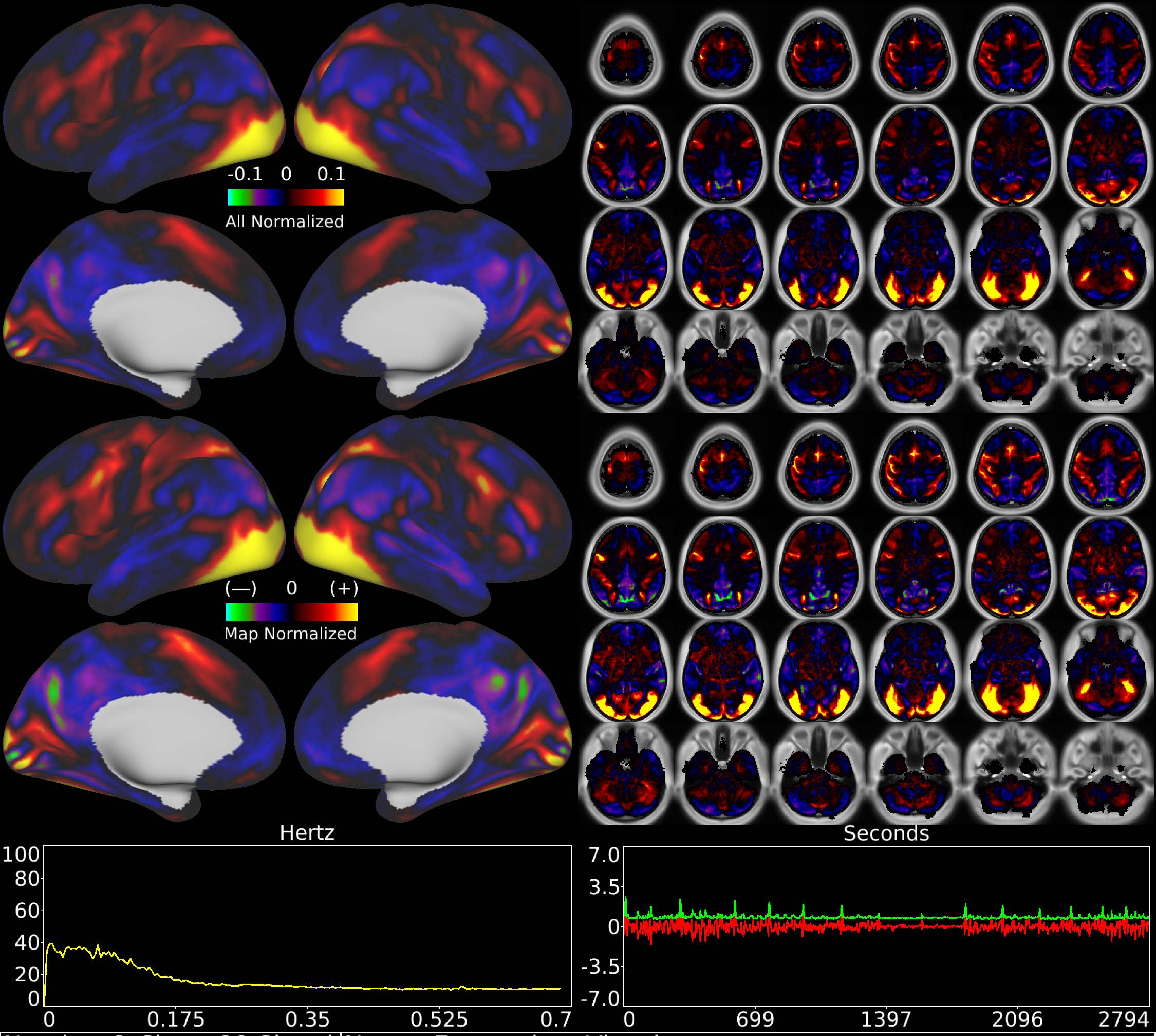


| | | |
|---|---|---|
| Number & Class: 29 Signal | Name: Extrastriate Visual | |
| RVT Correlated: No | DVARS Dip Associated: No | Cross-Subject Variable: No |
| Single Subject: No | % Variance Explained: 1.31 | Globality Index: 0.04 |
| Task Component: 7 | Rest Component: No | Task Modulated: No |

Rationale: Spatial map includes positive and negative patches that respect known retinotopic visual organization (Vertical vs Horizontal Meridians); Extrastriate Visual RSN

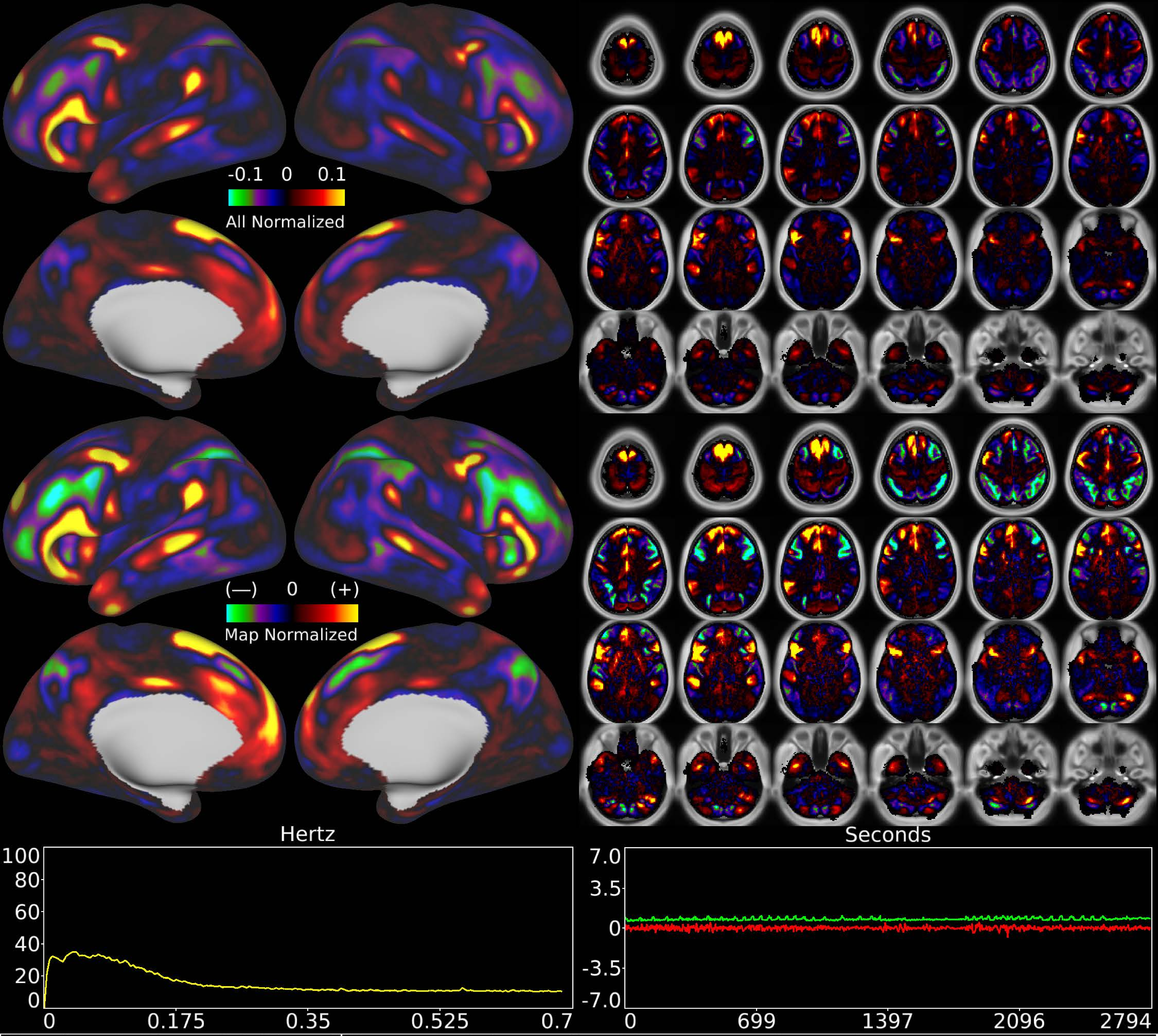


| | | |
|---|---|---|
| Number & Class: 30 Signal | Name: Left Lateralized Language Network | |
| RVT Correlated: No | DVARS Dip Associated: No | Cross-Subject Variable: No |
| Single Subject: No | % Variance Explained: 1.19 | Globality Index: 0.01 |
| Task Component: 22 | Rest Component: 20 | Task Modulated: No |

Rationale: Spatial map includes positive and negative patches that respect known RSNs (e.g. Language Network and areal boundaries (44; 45; 55b; PSL; SFL)

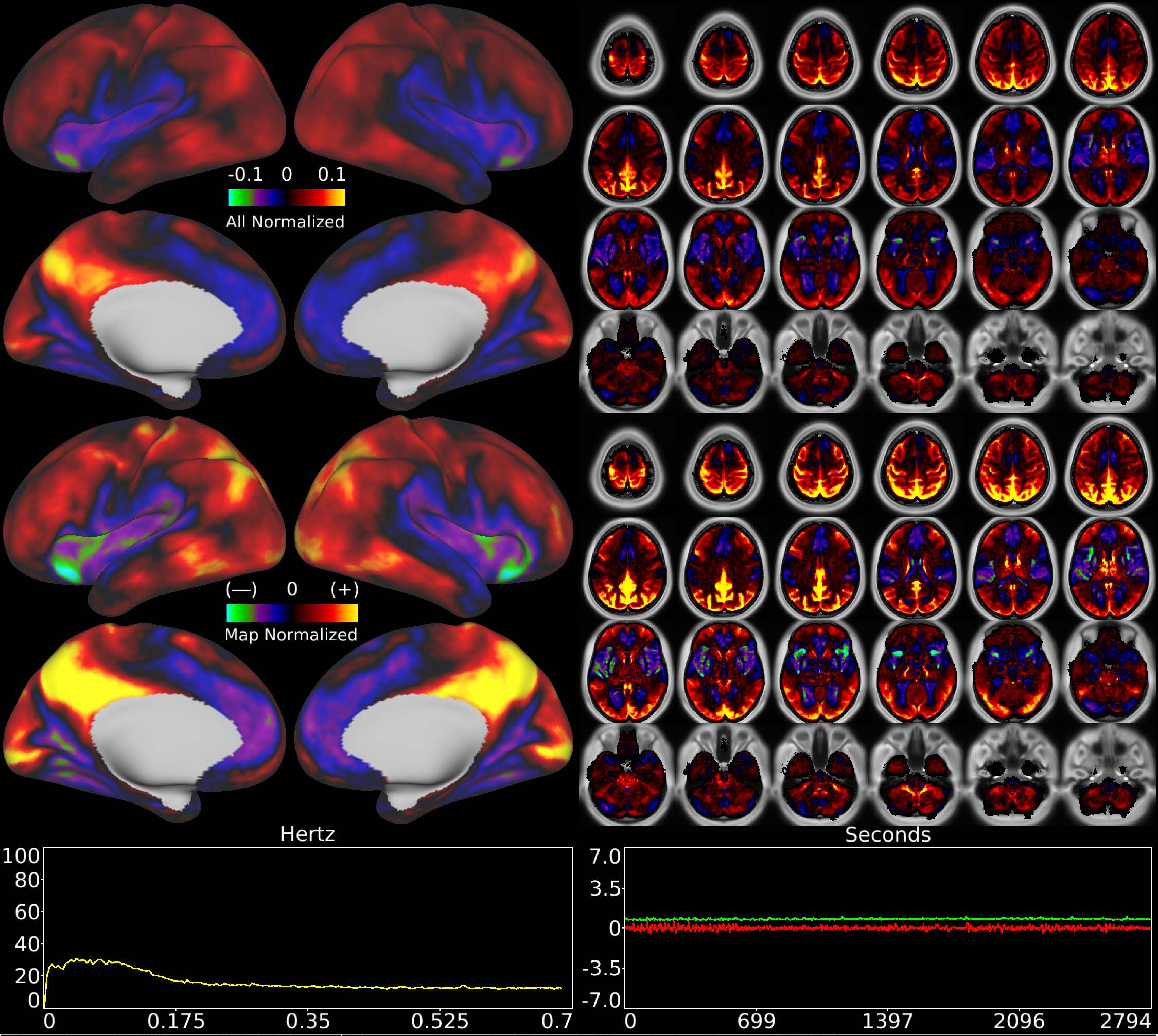


| | | |
|---|---|---|
| Number & Class: 31 Noise | Name: Semi-Global Physiological BOLD? | |
| RVT Correlated: No | DVARS Dip Associated: No | Cross-Subject Variable: No |
| Single Subject: Yes | % Variance Explained: 1.19 | Globality Index: 1.21 |
| Task Component: 30 | Rest Component: 25 | Task Modulated: No |
| Rationale: Controversial: Bears some resemblance to RC25 and TC30; positive and negative patches do not clearly reflect RSNs or areal borders | | |

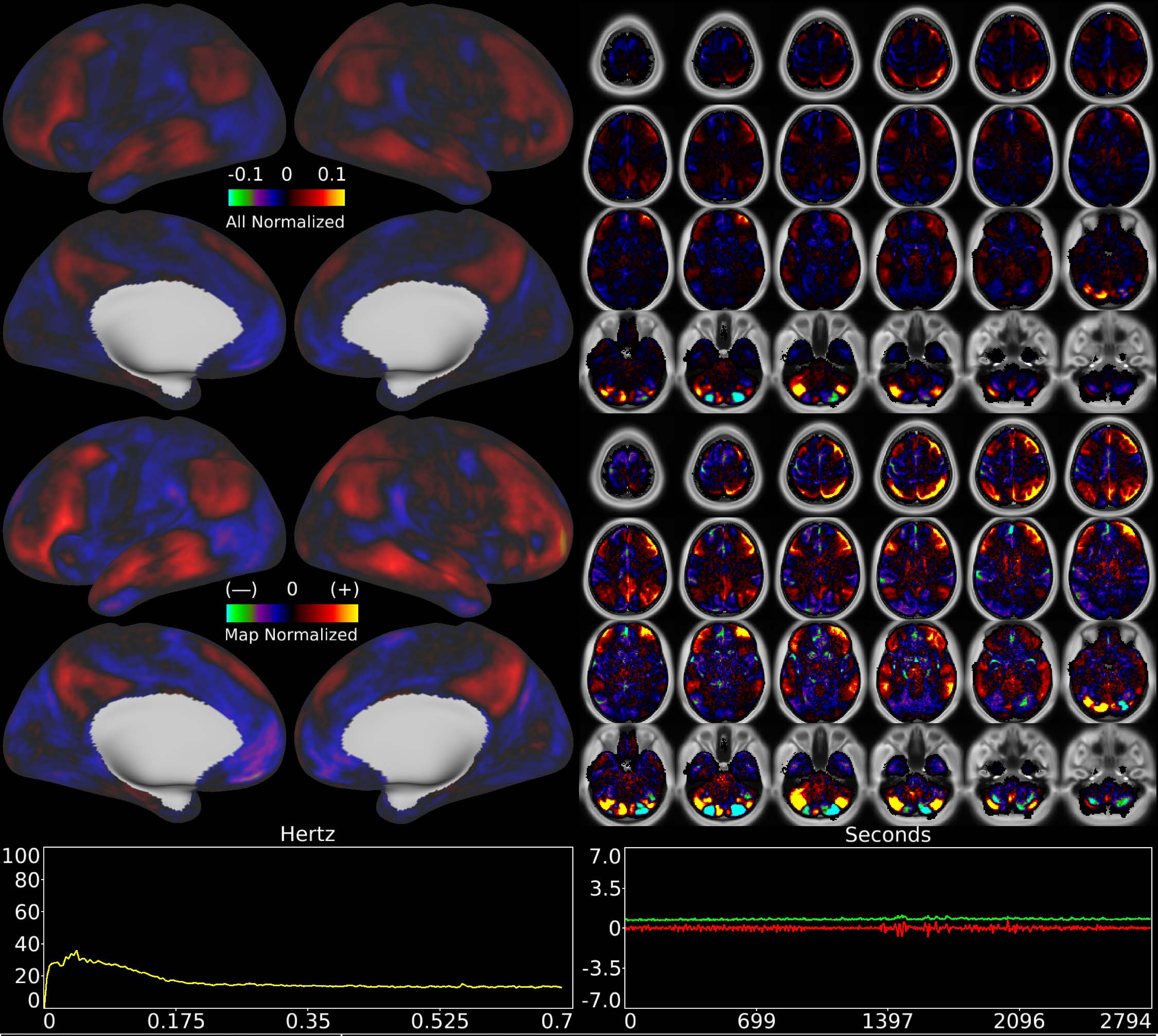


| | | |
|---|---|---|
| Number & Class: 32 Signal | Name: Cerebellar Unknown | |
| RVT Correlated: No | DVARS Dip Associated: No | Cross-Subject Variable: No |
| Single Subject: No | % Variance Explained: 1.16 | Globality Index: 0.28 |
| Task Component: -36 | Rest Component: 20 | Task Modulated: No |
| Rationale: Spatial map includes positive and negative patches that respect known RSN boundaries in the cerebellum | | |

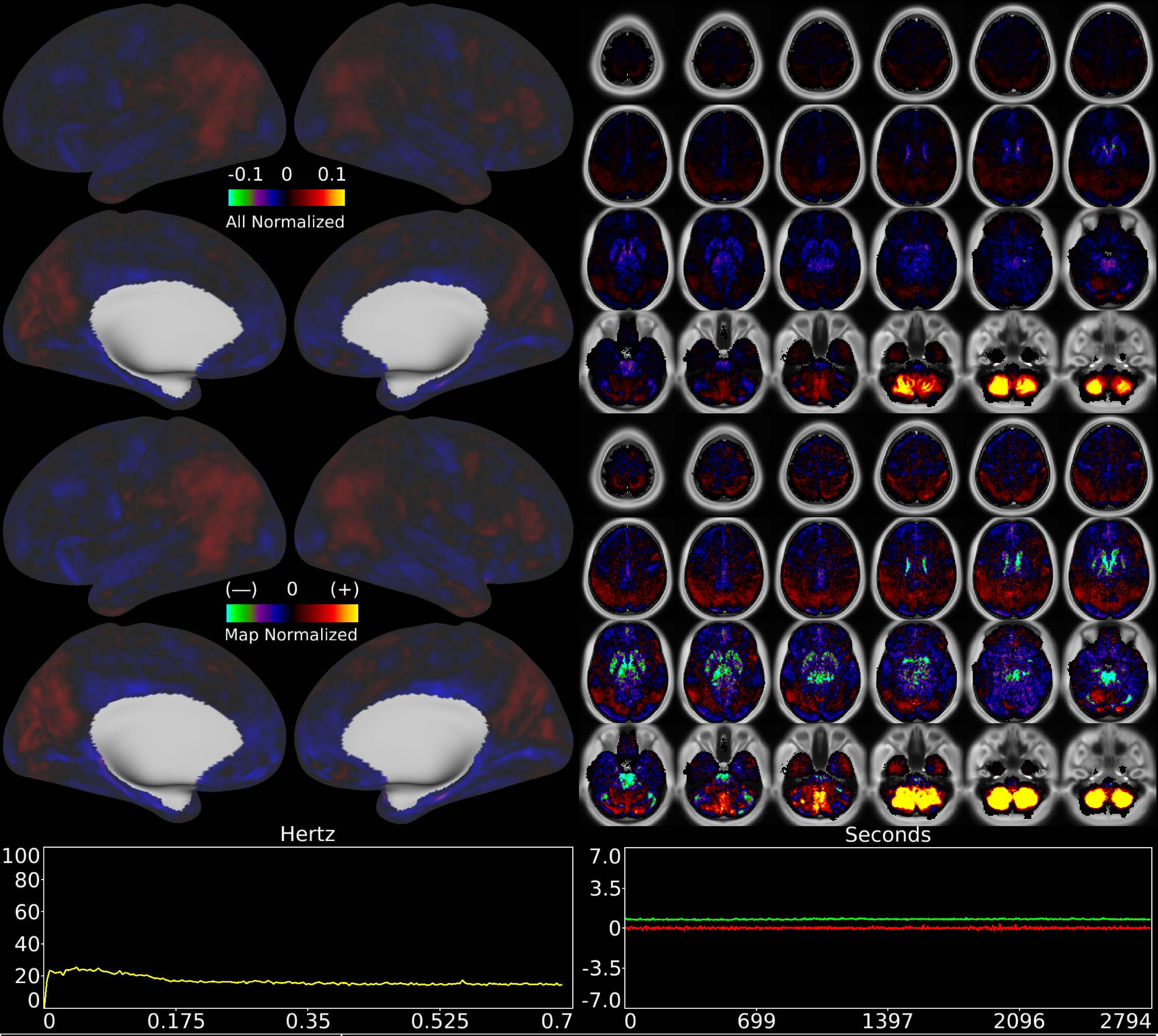


| | | |
|---|---|---|
| Number & Class: 33 Noise | Name: Bilateral Inferior Cerebellum | |
| RVT Correlated: No | DVARS Dip Associated: Yes | Cross-Subject Variable: Yes |
| Single Subject: Yes | % Variance Explained: 1.11 | Globality Index: 0.61 |
| Task Component: 41 | Rest Component: No | Task Modulated: No |
| Rationale: Component is DVARS Dips associated and single subject; perhaps related to motion; reconstruction; or unstructured noise | | |

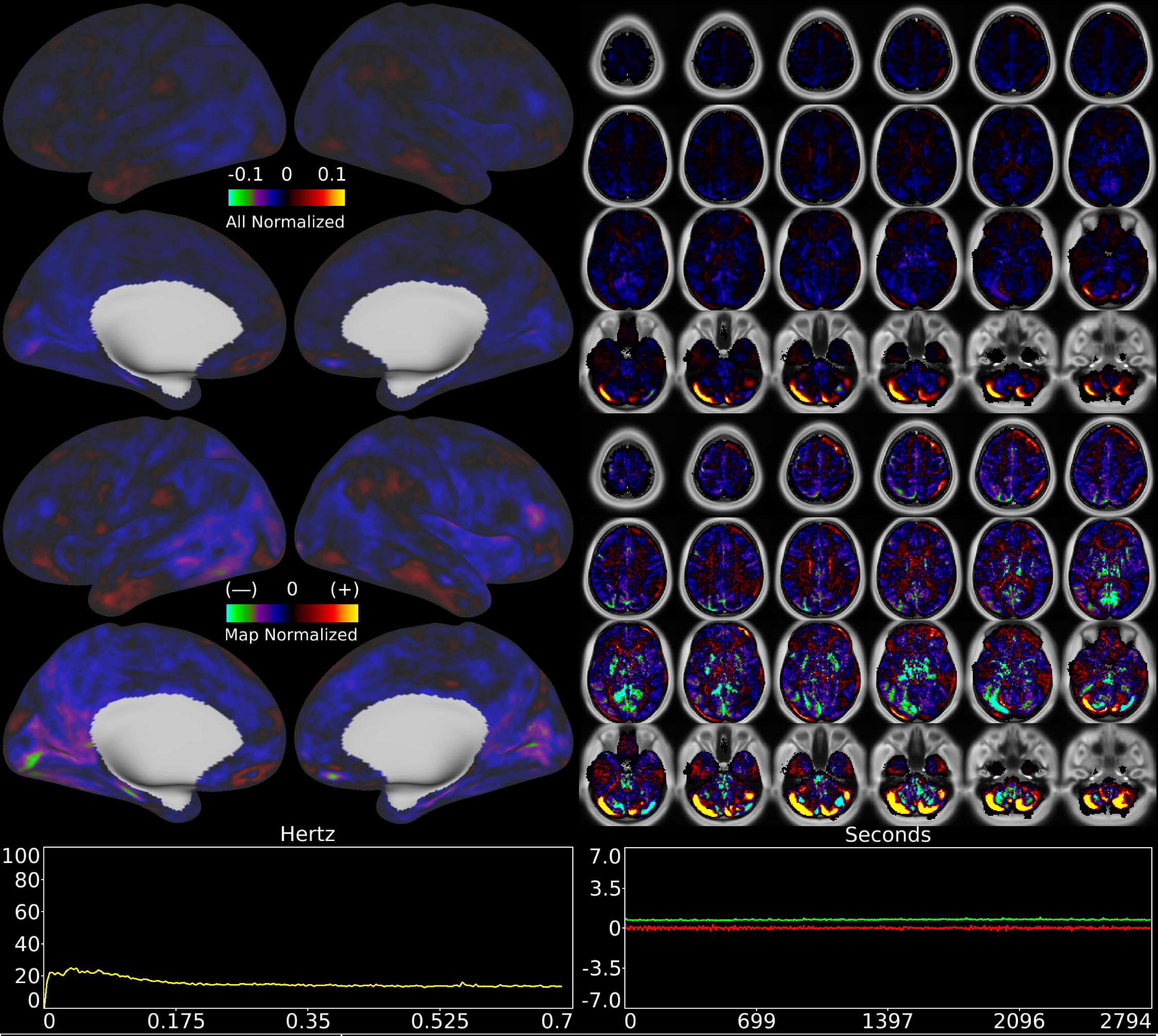


| | | |
|---|---|---|
| Number & Class: 34 Noise | Name: Cerebellar Movement Artifact Left | |
| RVT Correlated: No | DVARS Dip Associated: Yes | Cross-Subject Variable: Yes |
| Single Subject: Yes | % Variance Explained: 1.05 | Globality Index: 1.82 |
| Task Component: 44 | Rest Component: 57 | Task Modulated: No |

Rationale: Cerebellar edge motion component with high correlation to DVARS dips; looks like movement regressor beta map (derivative of X translation)

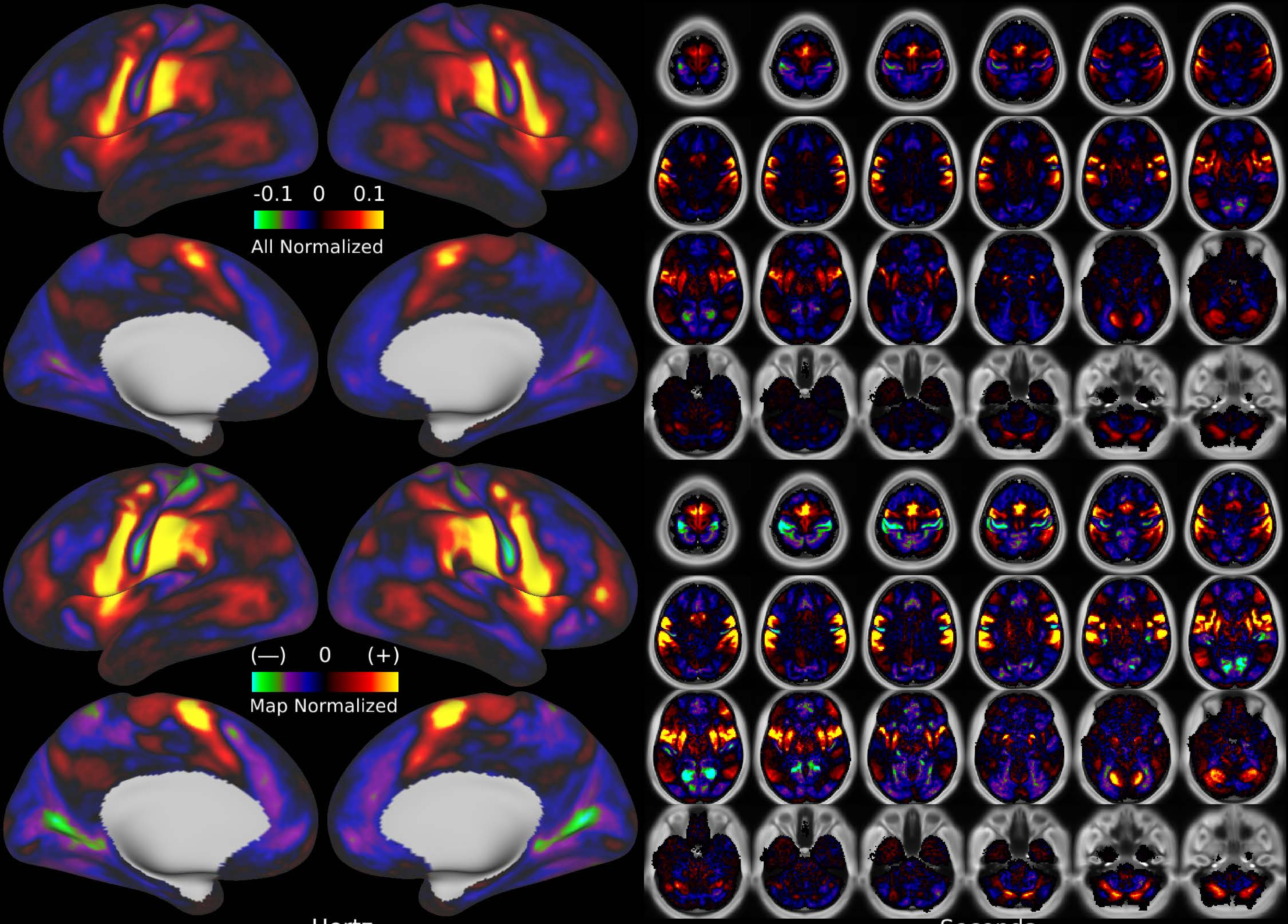


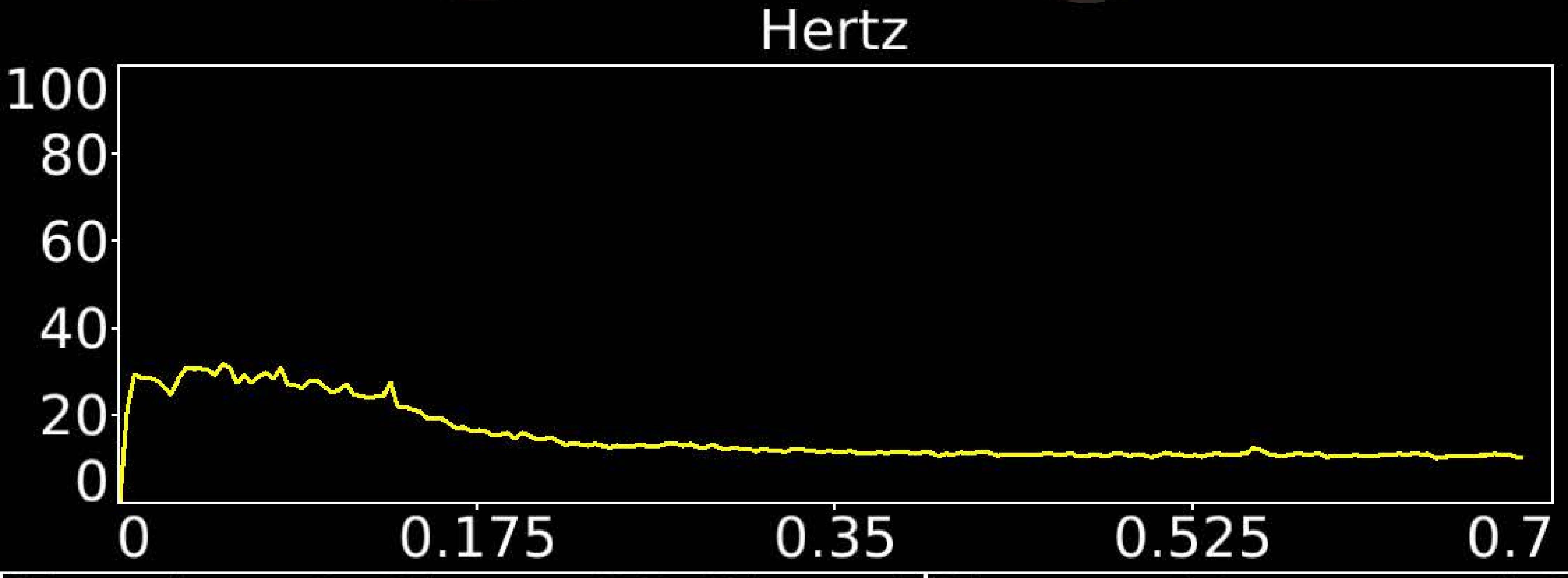


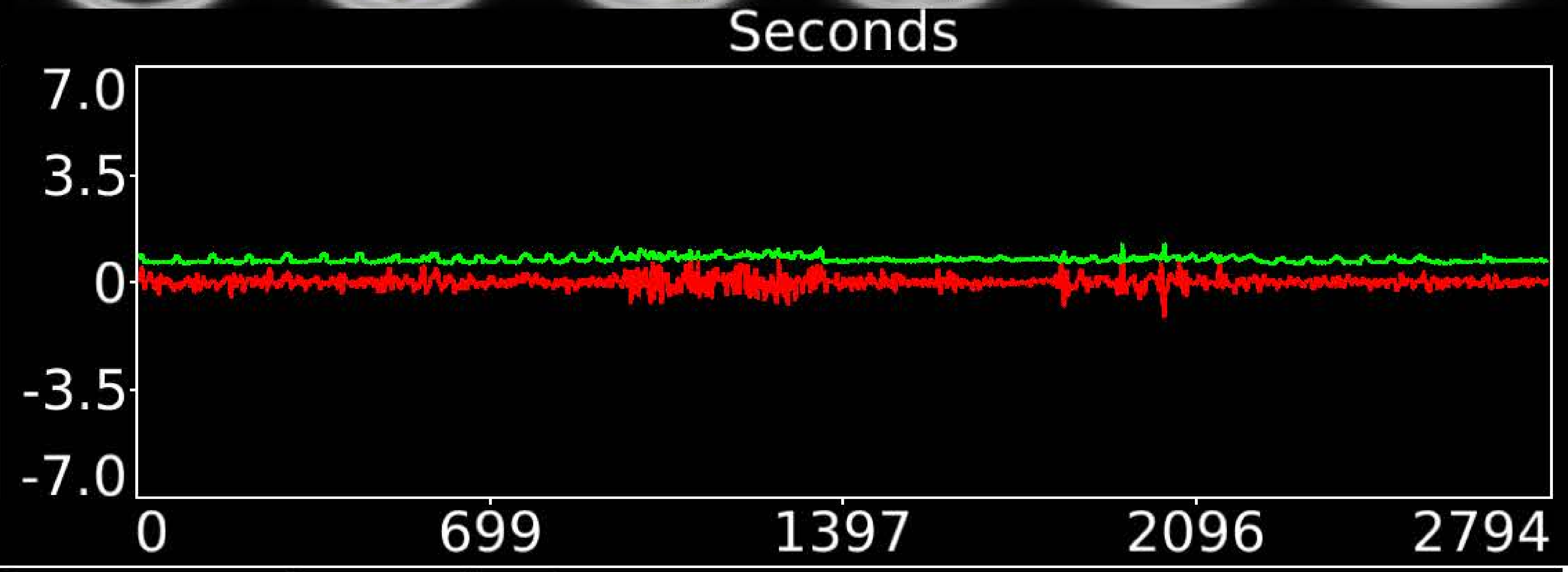


| | | |
|---|---|---|
| Number & Class: 35 Signal | Name: Motor and Sensory Association | |
| RVT Correlated: No | DVARS Dip Associated: No | Cross-Subject Variable: No |
| Single Subject: No | % Variance Explained: 1.06 | Globality Index: 0.39 |
| Task Component: No | Rest Component: 54 | Task Modulated: No |
| Rationale: Spatial map includes positive and negative patches that respect known areal boundaries (e.g. 4 and 1) | | |

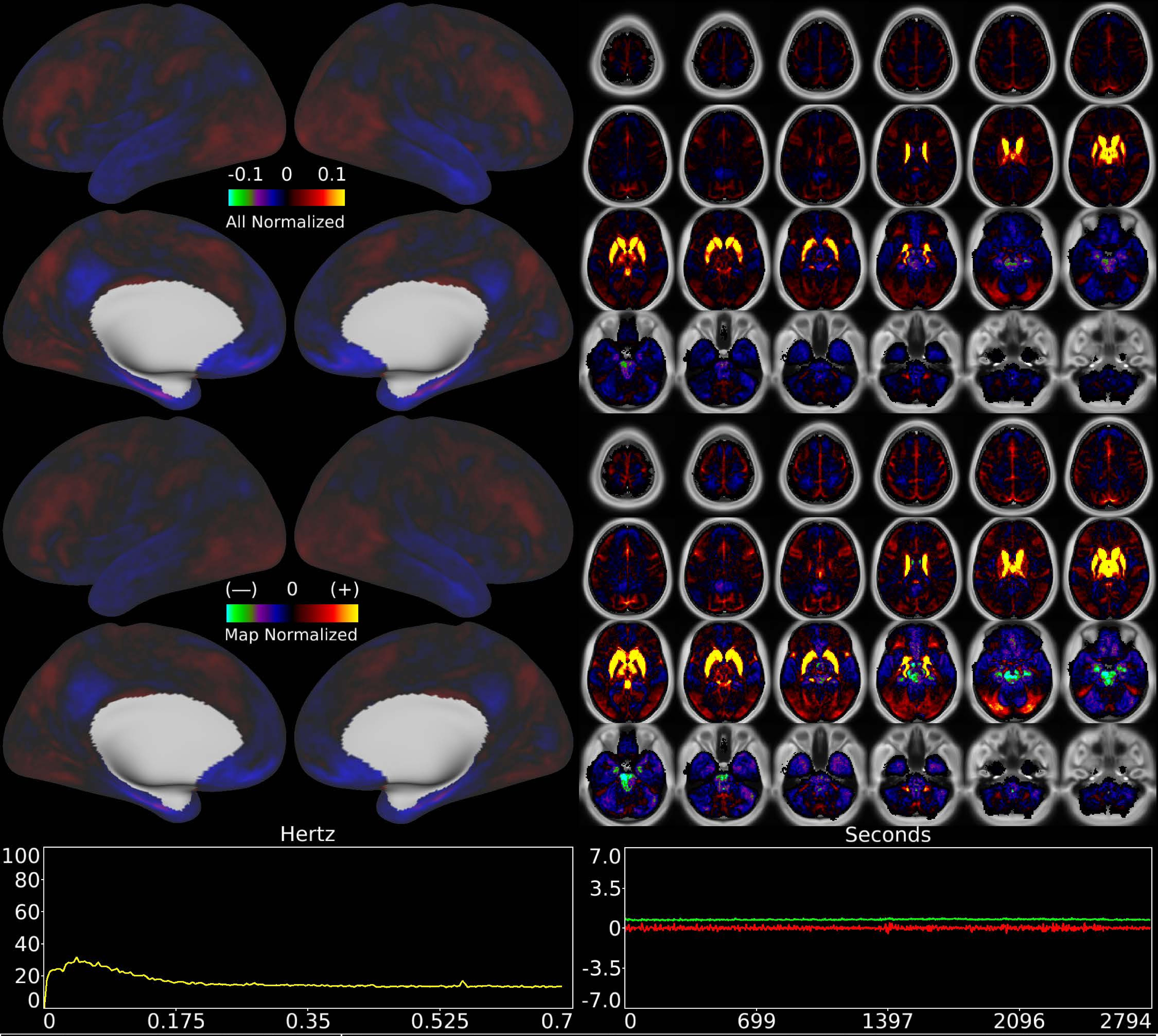


| | | |
|---|---|---|
| Number & Class: 36 Noise | Name: Striatum and Dorsal + Anterior Thalamus + WM | |
| RVT Correlated: No | DVARS Dip Associated: No | Cross-Subject Variable: No |
| Single Subject: No | % Variance Explained: 1.04 | Globality Index: 0.35 |
| Task Component: 42 | Rest Component: 44 | Task Modulated: No |
| Rationale: Controversial: Diencephalon together with surrounding white matter positive vs rest of brain negative; could be due to differing vascular supplies | | |

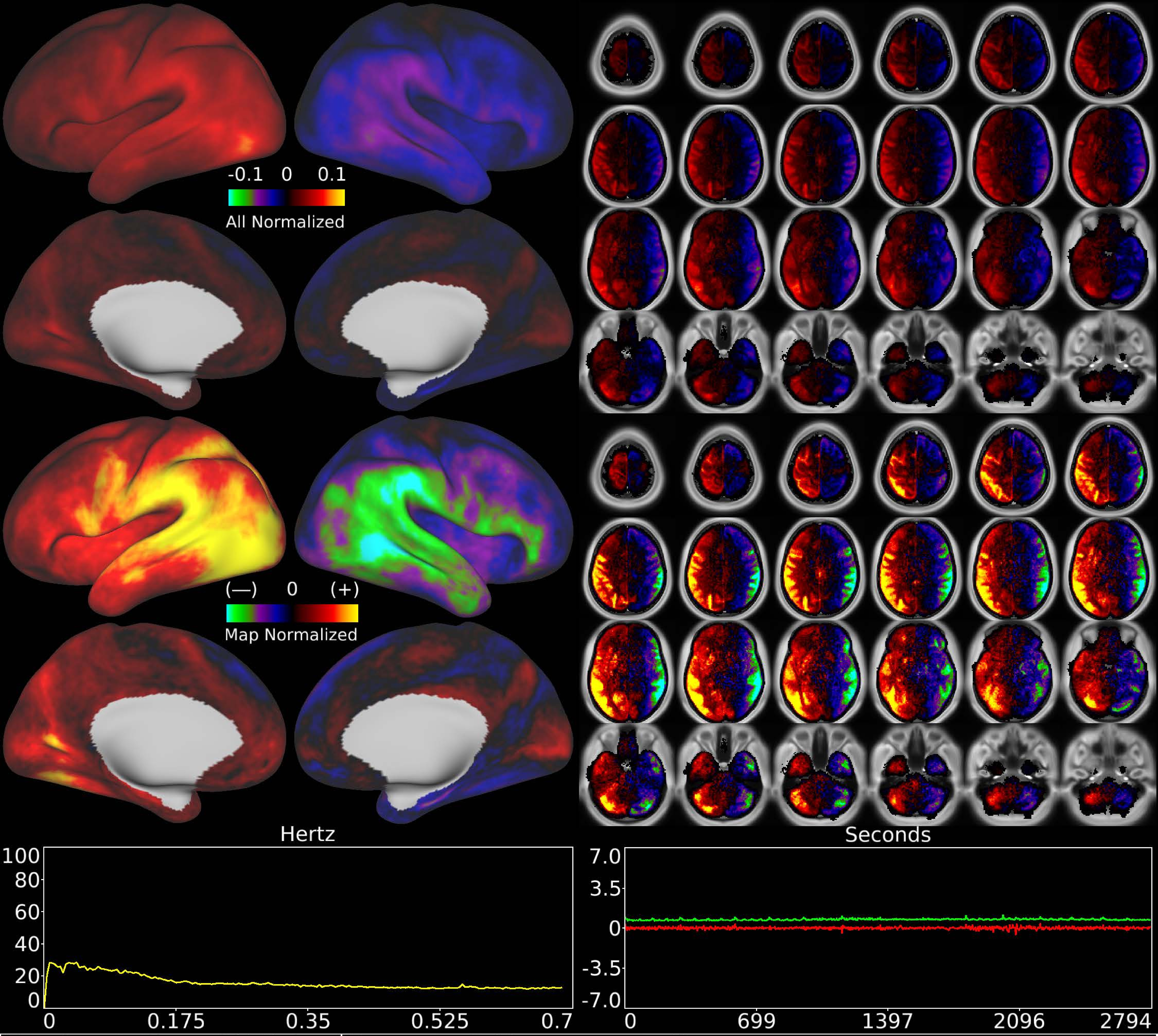


| | | |
|---|---|---|
| Number & Class: 37 Noise | Name: L>R Movement Artifact | |
| RVT Correlated: No | DVARS Dip Associated: Yes | Cross-Subject Variable: No |
| Single Subject: No | % Variance Explained: 1.05 | Globality Index: 0.47 |
| Task Component: 51 | Rest Component: 50 | Task Modulated: No |
| Rationale: Highly DVARS Dips associated component; looks like movement regressor beta map (Z rotation) | | |

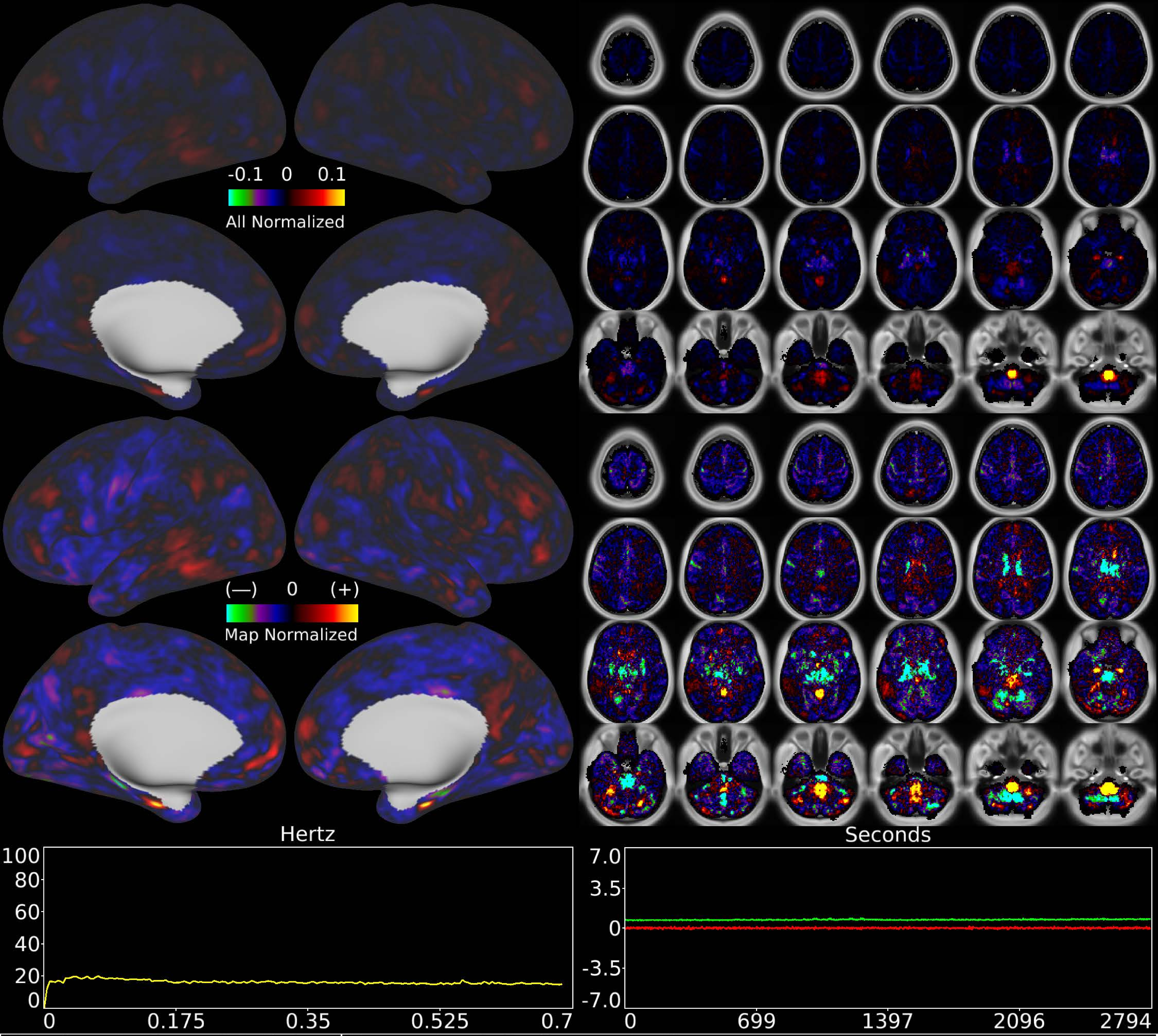


| | | |
|---|---|---|
| Number & Class: 38 Noise | Name: Medulla Recon Artifact | |
| RVT Correlated: No | DVARS Dip Associated: Yes | Cross-Subject Variable: Yes |
| Single Subject: No | % Variance Explained: 1.05 | Globality Index: 1.53 |
| Task Component: 43 | Rest Component: 46 | Task Modulated: No |
| Rationale: Spatial map not reflective of known areas or RSNs without connectivity to other brain structures; some banding in sagittal plane suggestive of multi-band recon | | |

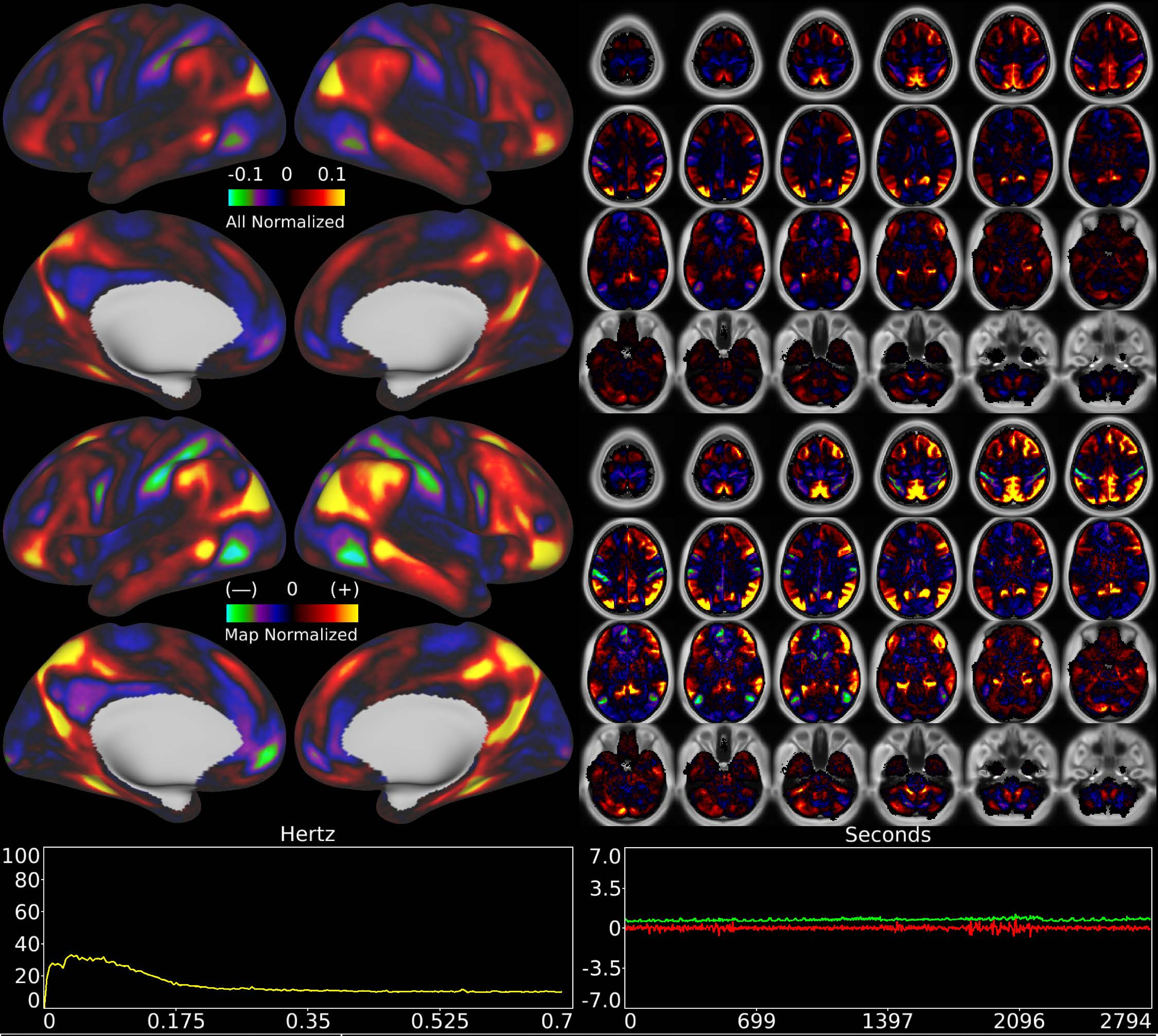


| | | |
|---|---|---|
| Number & Class: 39 Signal | Name: Parietal Occipital Network (Latest Modularity) | |
| RVT Correlated: No | DVARS Dip Associated: No | Cross-Subject Variable: No |
| Single Subject: No | % Variance Explained: 1.03 | Globality Index: 0.43 |
| Task Component: No | Rest Component: 32 | Task Modulated: No |
| Rationale: Spatial map includes positive and negative patches that respect known RSNs (e.g. Fronto-Parietal Network) | | |

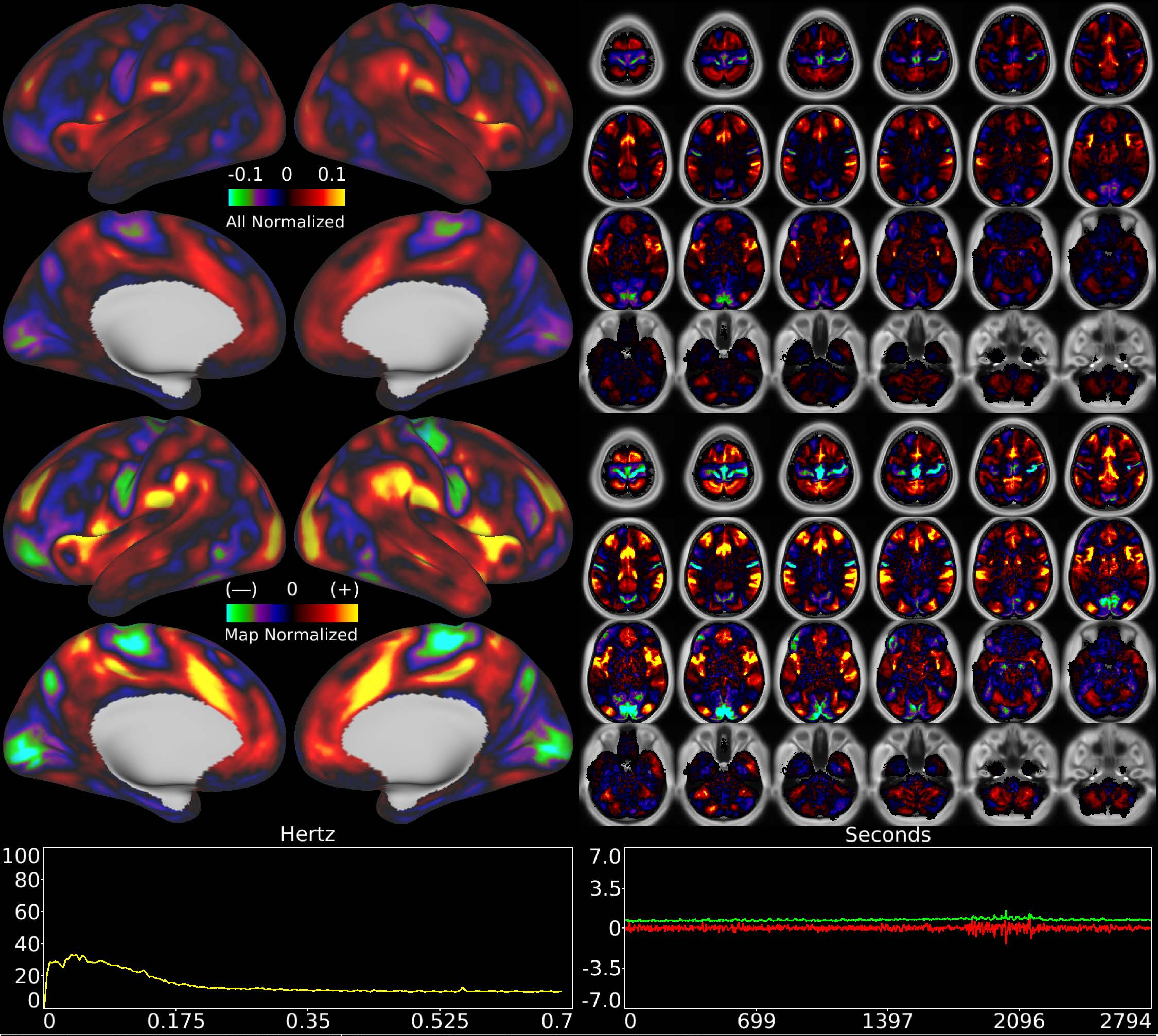


| | | |
|---|---|---|
| Number & Class: 40 Signal | Name: Cingular-Opercular-Like | |
| RVT Correlated: No | DVARS Dip Associated: No | Cross-Subject Variable: No |
| Single Subject: No | % Variance Explained: 0.99 | Globality Index: 0.55 |
| Task Component: No | Rest Component: No | Task Modulated: No |
| Rationale: Spatial map includes positive and negative patches that respect known RSNs (e.g. Cingulo-Opercular Network) | | |

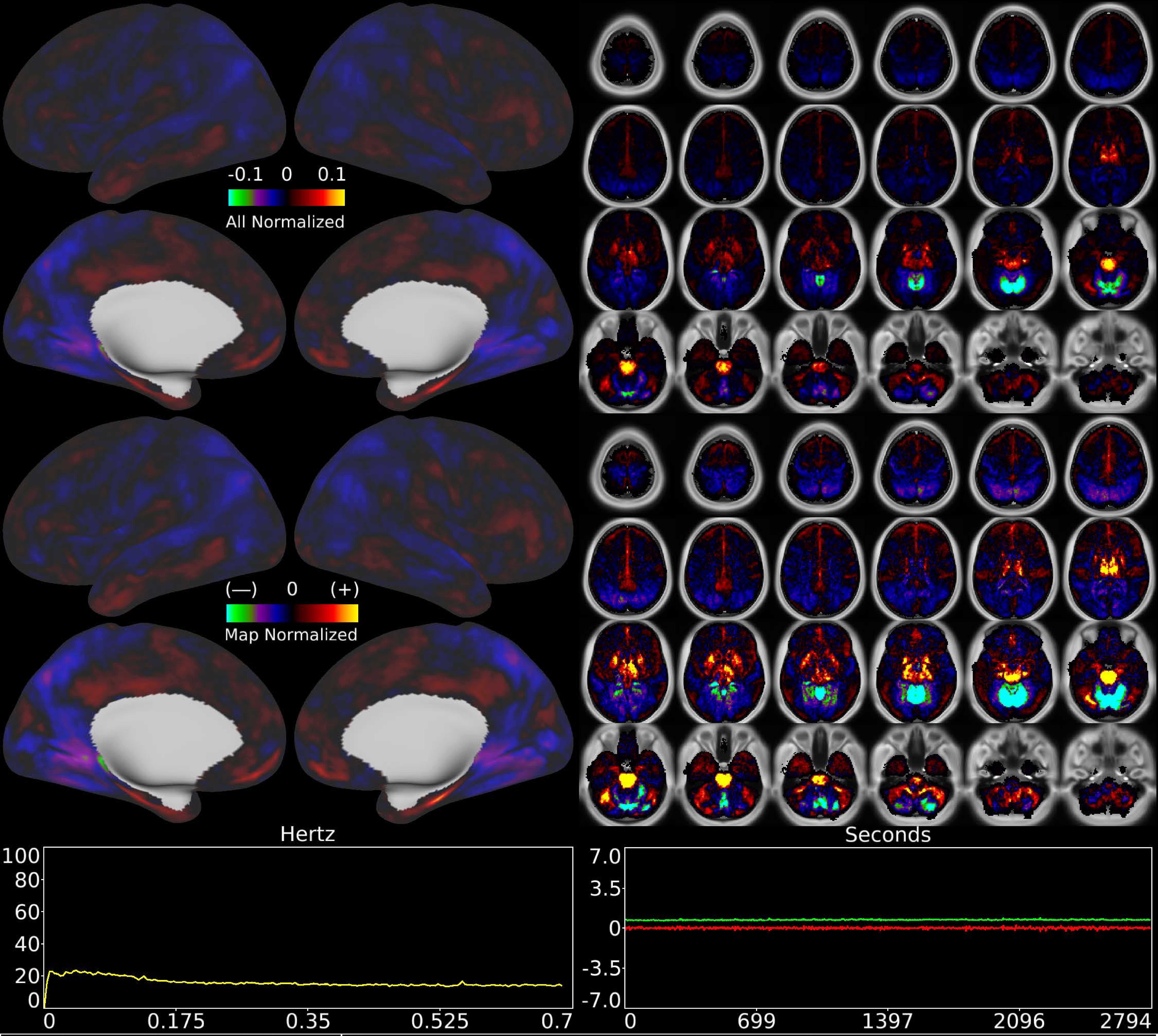


| | | |
|---|---|---|
| Number & Class: 41 Noise | Name: Pons + Thalamus > Superior Cerebellum | |
| RVT Correlated: No | DVARS Dip Associated: No | Cross-Subject Variable: No |
| Single Subject: No | % Variance Explained: 0.97 | Globality Index: 0.33 |
| Task Component: 45 | Rest Component: No | Task Modulated: No |
| Rationale: Spatial map not reflective of known areas or RSNs | | |

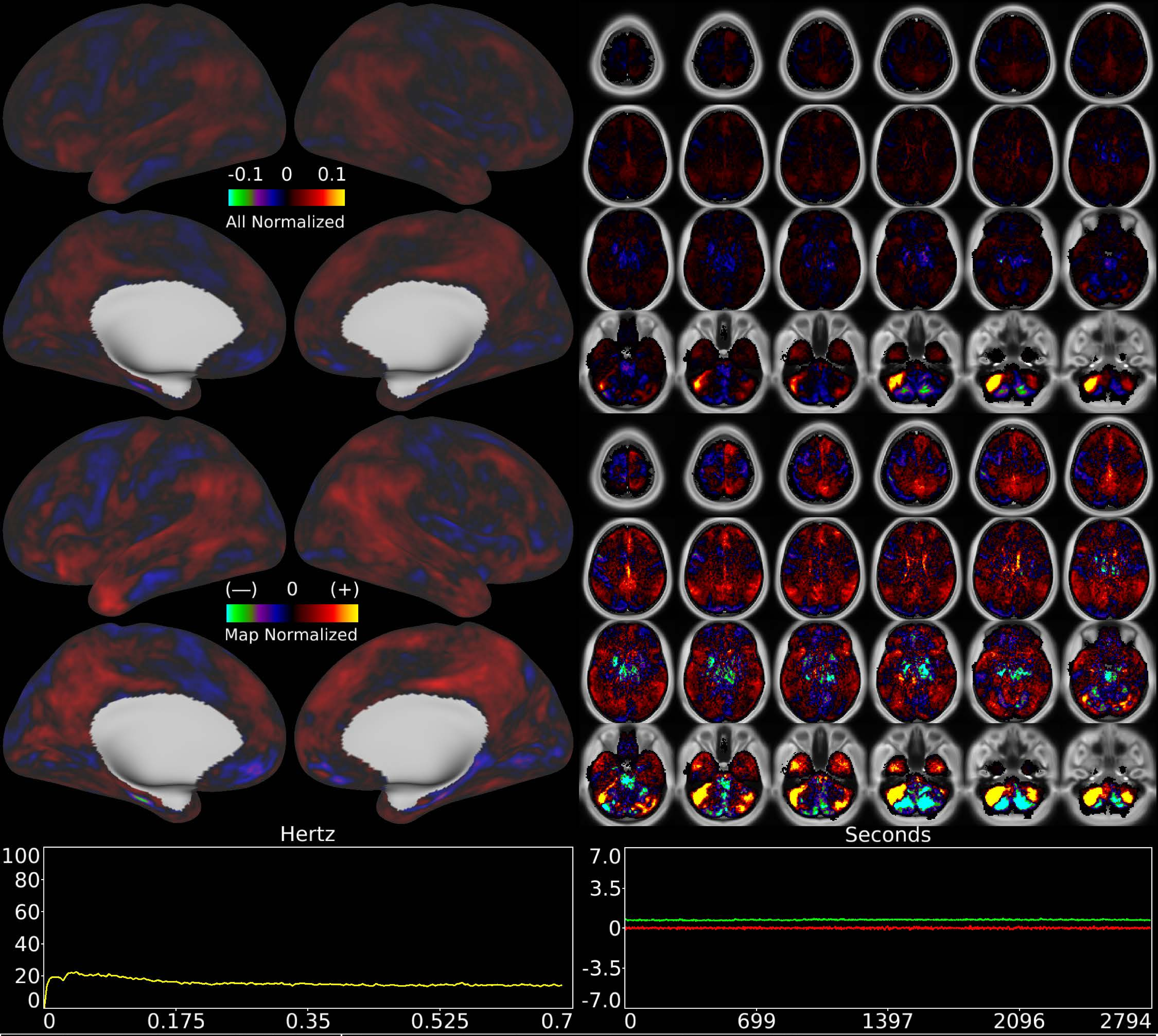


| | | |
|---|---|---|
| Number & Class: 42 Noise | Name: L Cerebellum Near Sigmoid Sinus | |
| RVT Correlated: No | DVARS Dip Associated: No | Cross-Subject Variable: No |
| Single Subject: Yes | % Variance Explained: 0.97 | Globality Index: 0.55 |
| Task Component: 46 | Rest Component: No | Task Modulated: No |
| Rationale: Single subject component not reflective of known areas or RSNs | | |

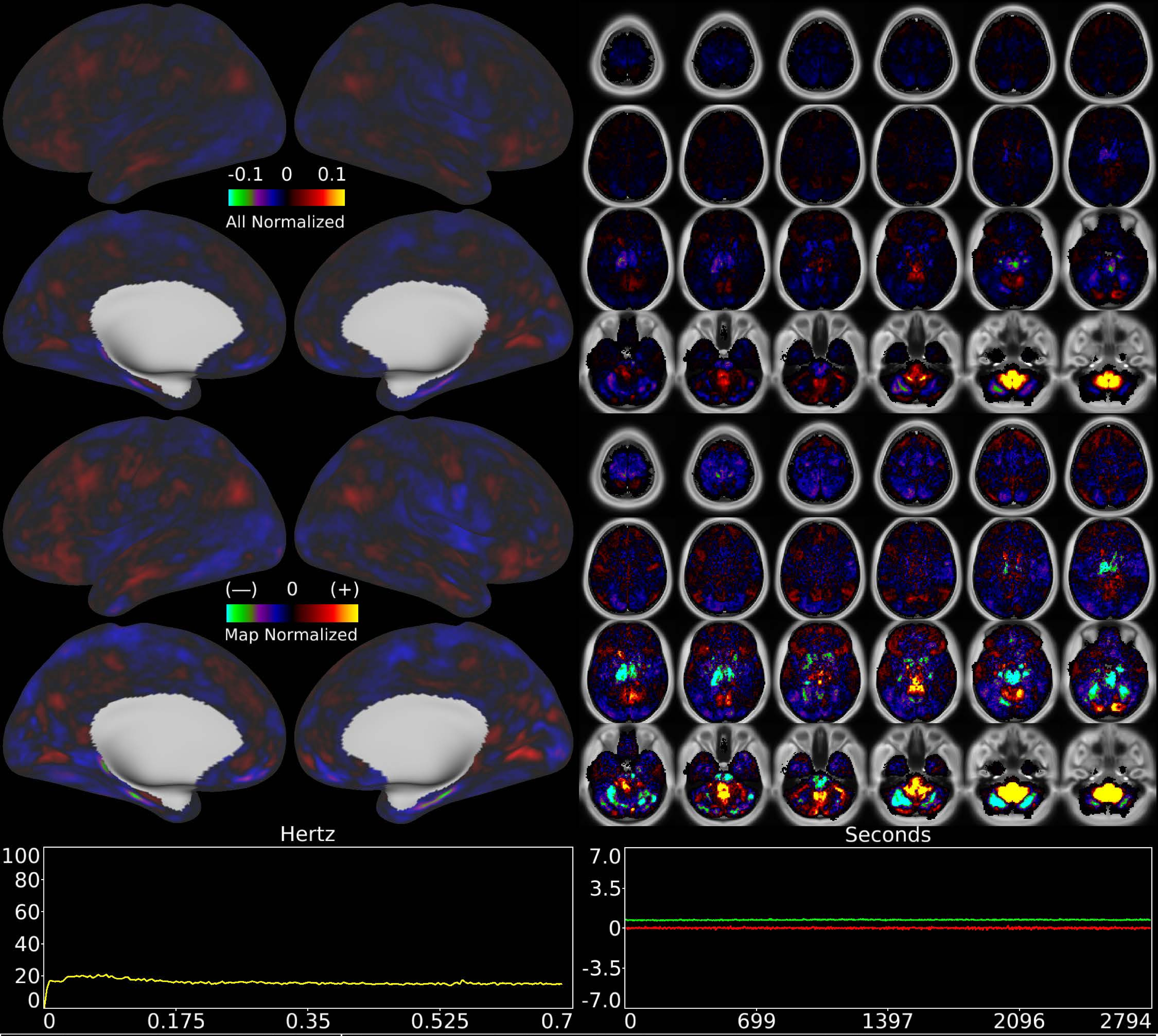


| Number & Class: 43 Noise | Name: Cerebellar + Brainstem Recon Artifact | |
|---|---|---|
| RVT Correlated: No | DVARS Dip Associated: No | Cross-Subject Variable: No |
| Single Subject: Yes | % Variance Explained: 0.96 | Globality Index: 0.83 |
| Task Component: 49 | Rest Component: No | Task Modulated: No |

Rationale: Spatial map not reflective of known areas or RSNs without connectivity to other brain structures; some banding in sagittal plane suggestive of multi-band recon

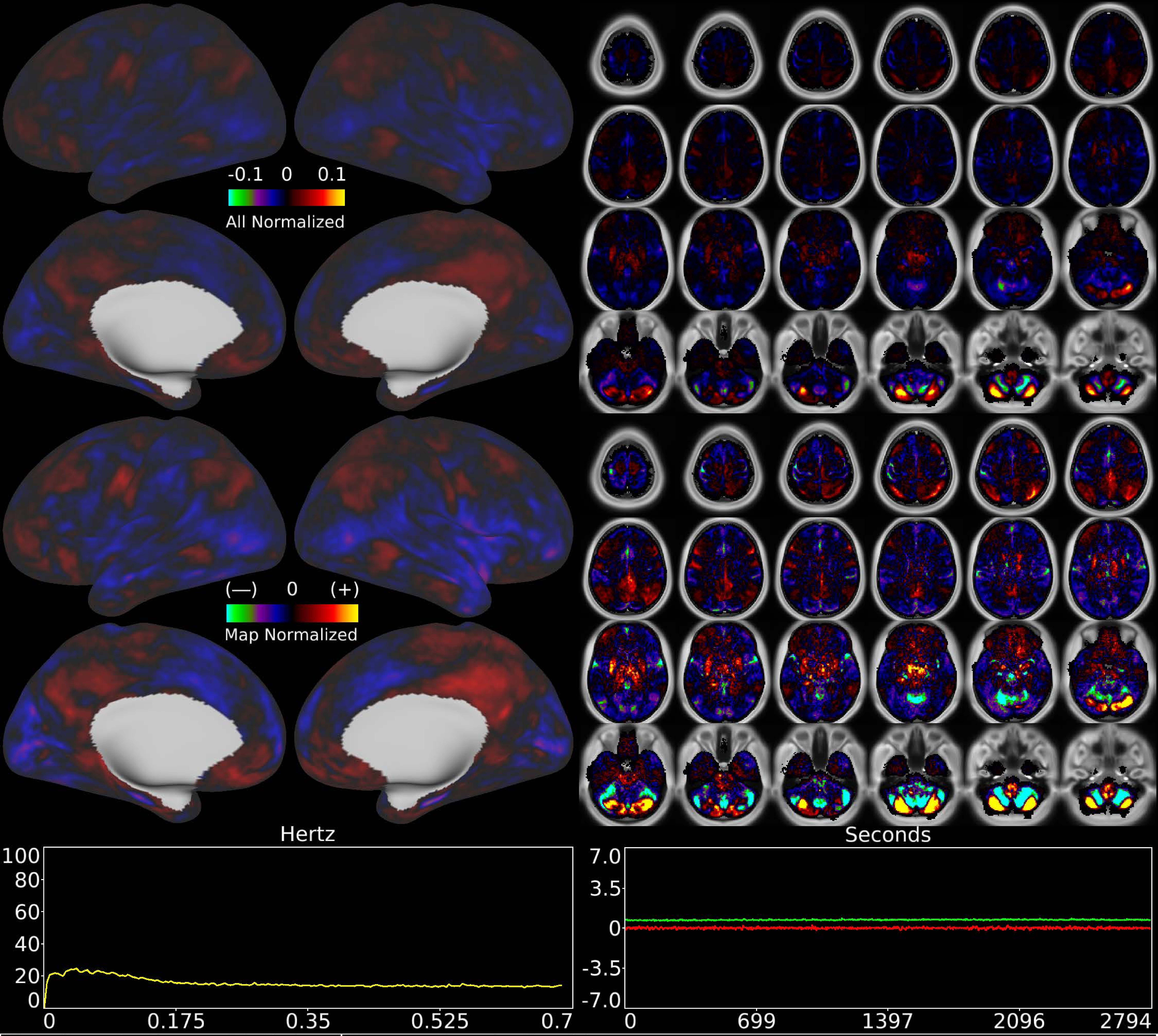


| | | |
|---|---|---|
| Number & Class: 44 Noise | Name: Cerebellar Unknown | |
| RVT Correlated: No | DVARS Dip Associated: No | Cross-Subject Variable: No |
| Single Subject: Yes | % Variance Explained: 0.96 | Globality Index: 0.39 |
| Task Component: No | Rest Component: No | Task Modulated: No |
| Rationale: Controversial: Clusters with other noise components in subject-wise amplitude correlations; spatial map does not match known areas or RSNs | | |

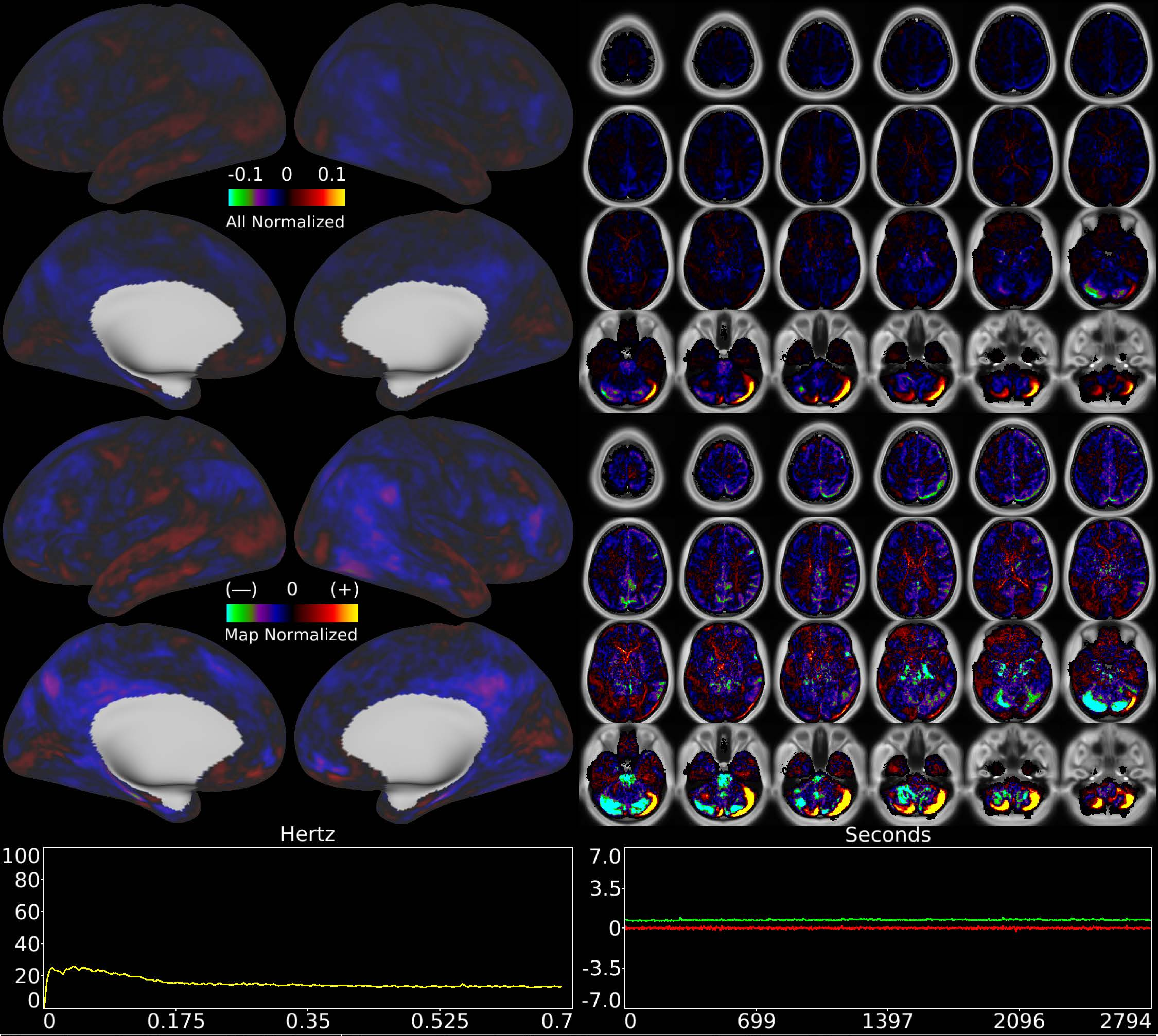


| | | |
|---|---|---|
| Number & Class: 45 Noise | Name: Cerebellar Movement Artifact Right | |
| RVT Correlated: No | DVARS Dip Associated: Yes | Cross-Subject Variable: No |
| Single Subject: Yes | % Variance Explained: 0.96 | Globality Index: 1.45 |
| Task Component: 47 | Rest Component: No | Task Modulated: No |
| Rationale: Cerebellar edge motion component with high correlation to DVARS dips; looks like movement regressor beta map (derivative of X translation) | | |

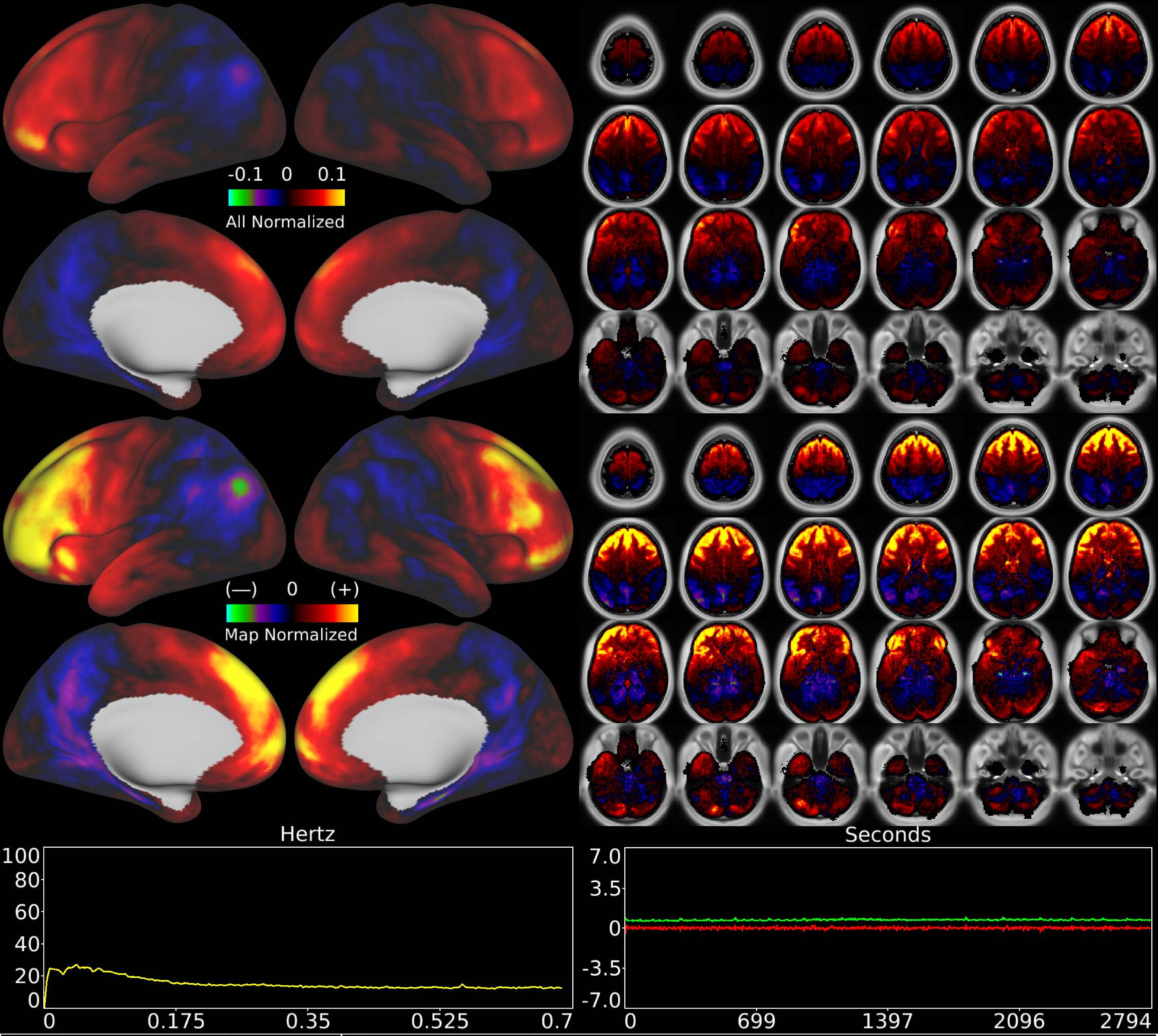


| | | |
|---|---|---|
| Number & Class: 46 Noise | Name: Coil or Movement | |
| RVT Correlated: No | DVARS Dip Associated: Yes | Cross-Subject Variable: No |
| Single Subject: No | % Variance Explained: 0.95 | Globality Index: 0.68 |
| Task Component: 60 | Rest Component: No | Task Modulated: No |
| Rationale: Substantial white matter signal; associated with DVARS dips; perhaps related to coil or motion | | |

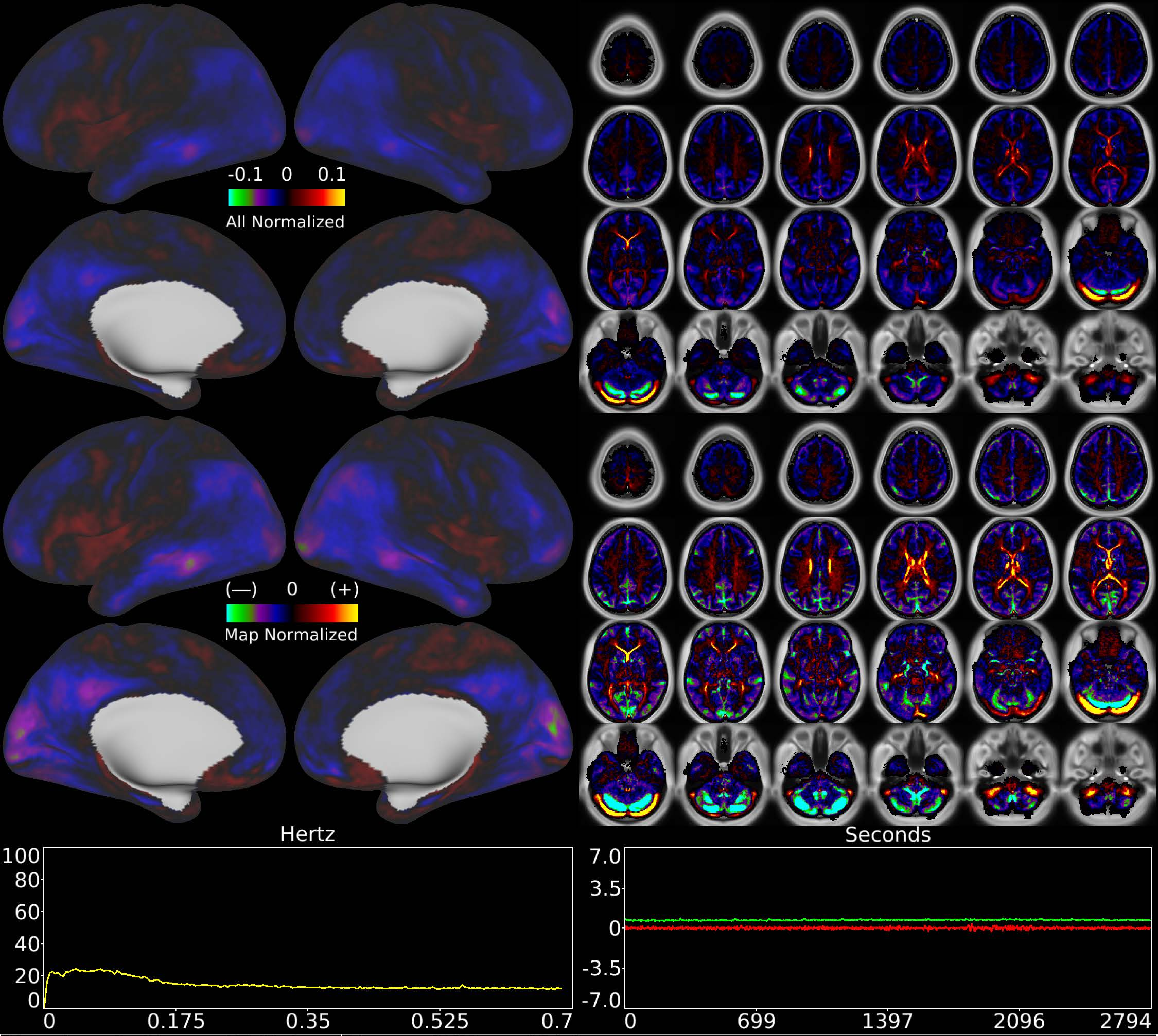


| | | |
|---|---|---|
| Number & Class: 47 Noise | Name: WM and Veins | |
| RVT Correlated: No | DVARS Dip Associated: No | Cross-Subject Variable: No |
| Single Subject: No | % Variance Explained: 0.91 | Globality Index: 1.42 |
| Task Component: No | Rest Component: No | Task Modulated: No |
| Rationale: Spatial map primarily white matter and veins | | |

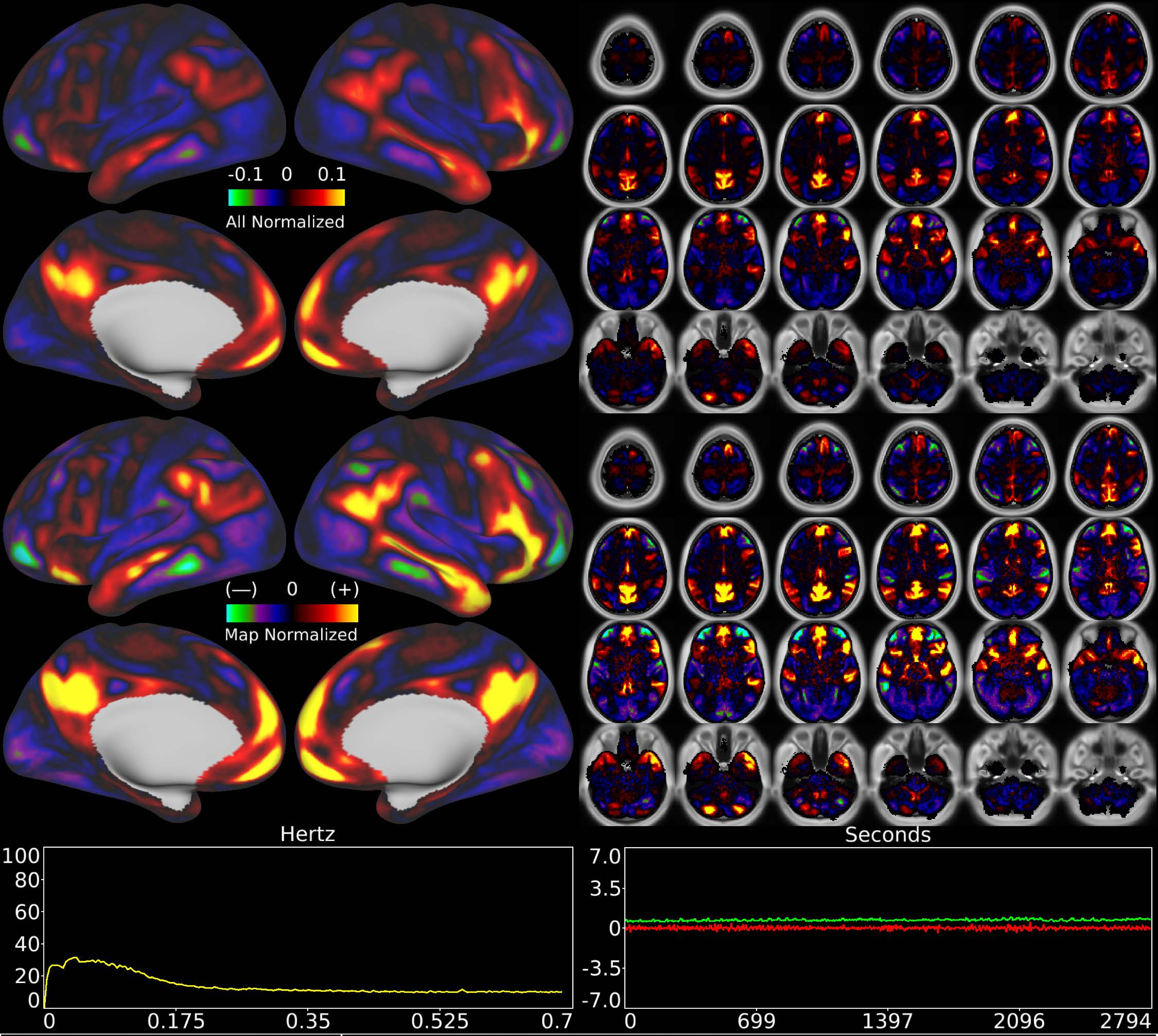


| | | |
|---|---|---|
| Number & Class: 48 Signal | Name: Subsidiary Default Mode | |
| RVT Correlated: No | DVARS Dip Associated: No | Cross-Subject Variable: No |
| Single Subject: No | % Variance Explained: 0.94 | Globality Index: 0.31 |
| Task Component: No | Rest Component: 24 | Task Modulated: No |
| Rationale: Spatial map includes positive and negative patches that respect known RSNs (e.g. Default Mode Network) | | |

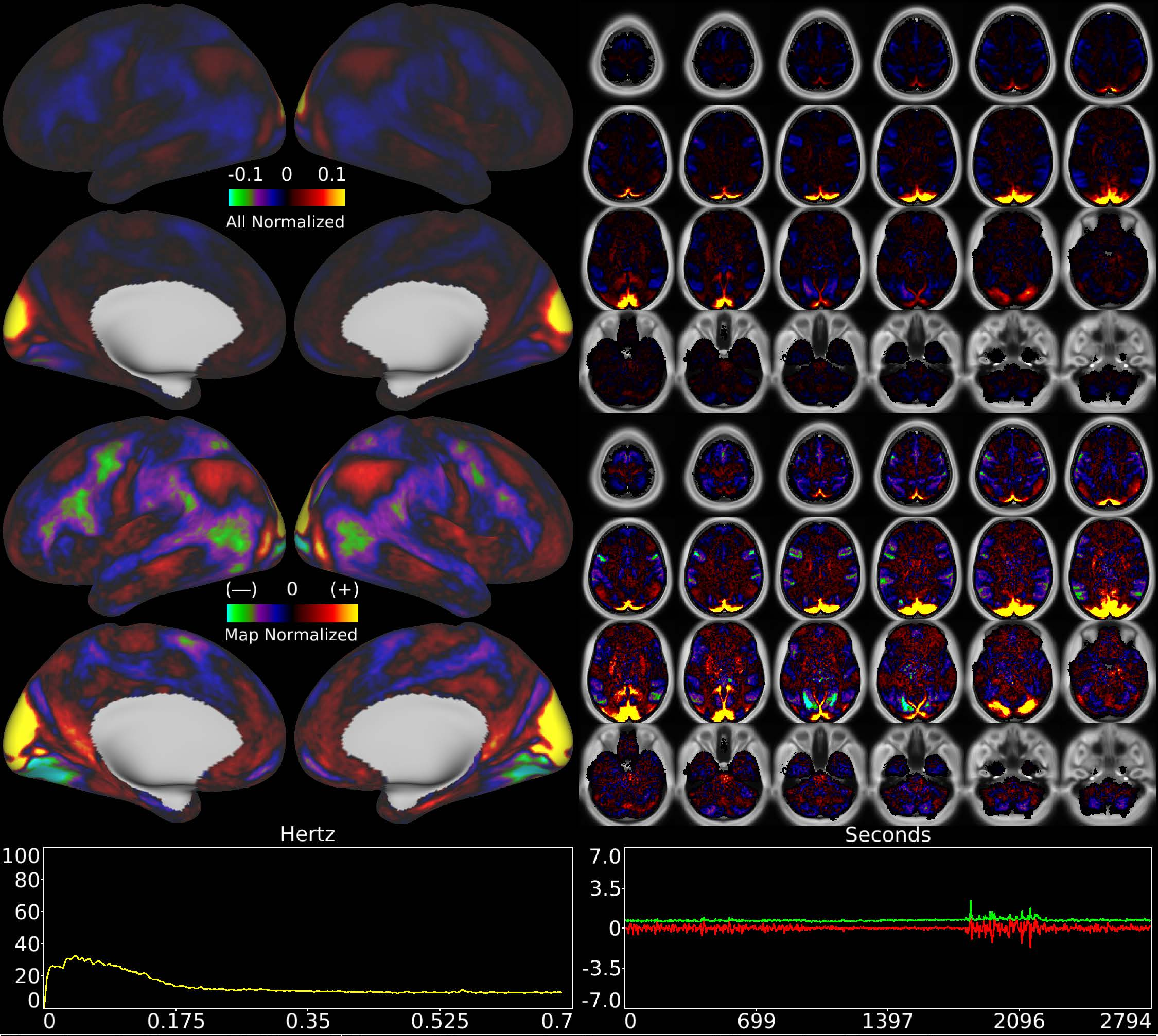


| | | |
|---|---|---|
| Number & Class: 49 Signal | Name: Visuotopic: Paracentral Dorsal > Ventral | |
| RVT Correlated: No | DVARS Dip Associated: No | Cross-Subject Variable: No |
| Single Subject: No | % Variance Explained: 0.93 | Globality Index: 0.09 |
| Task Component: 67 | Rest Component: 42+82 | Task Modulated: Social |
| Rationale: Spatial map includes positive and negative patches that respect known retinotopic visual organization (Lower vs Upper) | | |

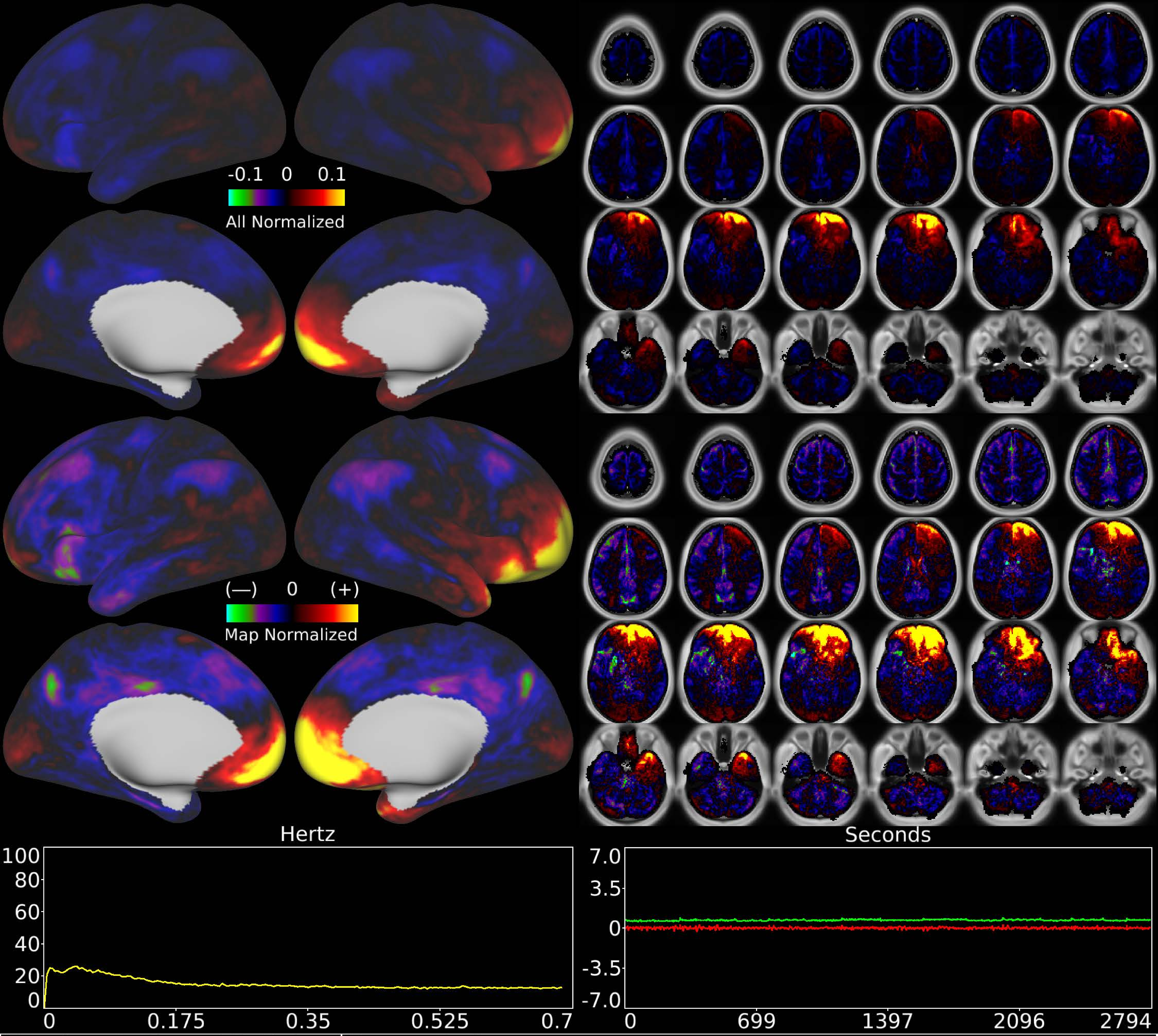


| | | |
|---|---|---|
| Number & Class: 50 Noise | Name: Coil | |
| RVT Correlated: No | DVARS Dip Associated: Yes | Cross-Subject Variable: No |
| Single Subject: No | % Variance Explained: 0.92 | Globality Index: 1.19 |
| Task Component: 50 | Rest Component: 47 | Task Modulated: No |
| Rationale: Known coil noise component | | |

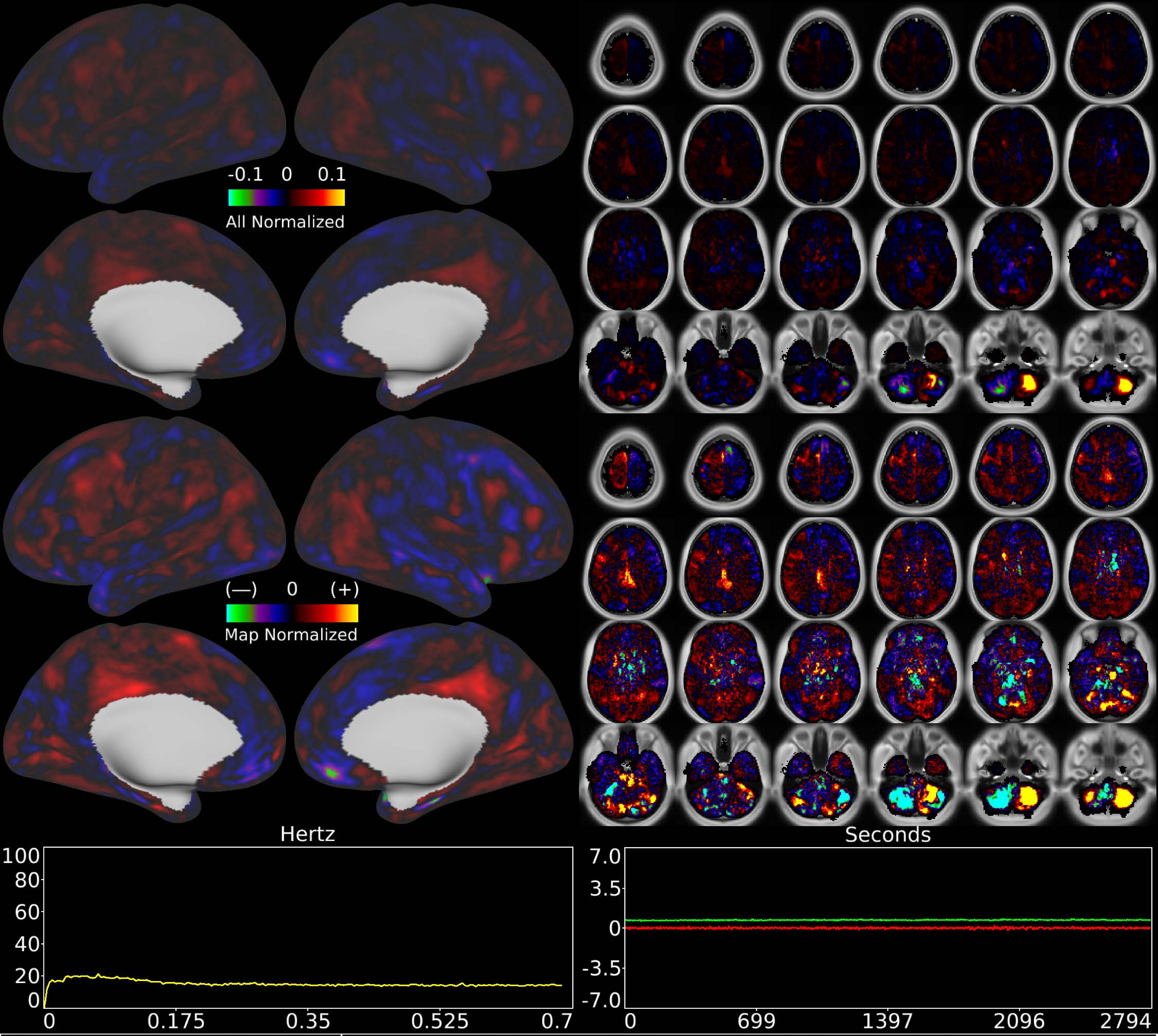


| | | |
|---|---|---|
| Number & Class: 51 Noise | Name: R Inferior Cerebellum > L Inferior Cerebellum | |
| RVT Correlated: No | DVARS Dip Associated: No | Cross-Subject Variable: No |
| Single Subject: Yes | % Variance Explained: 0.92 | Globality Index: 0.06 |
| Task Component: 54 | Rest Component: No | Task Modulated: No |
| Rationale: Classified same as TC54: single subject component whose spatial map is not reflective of known RSNs or areas | | |

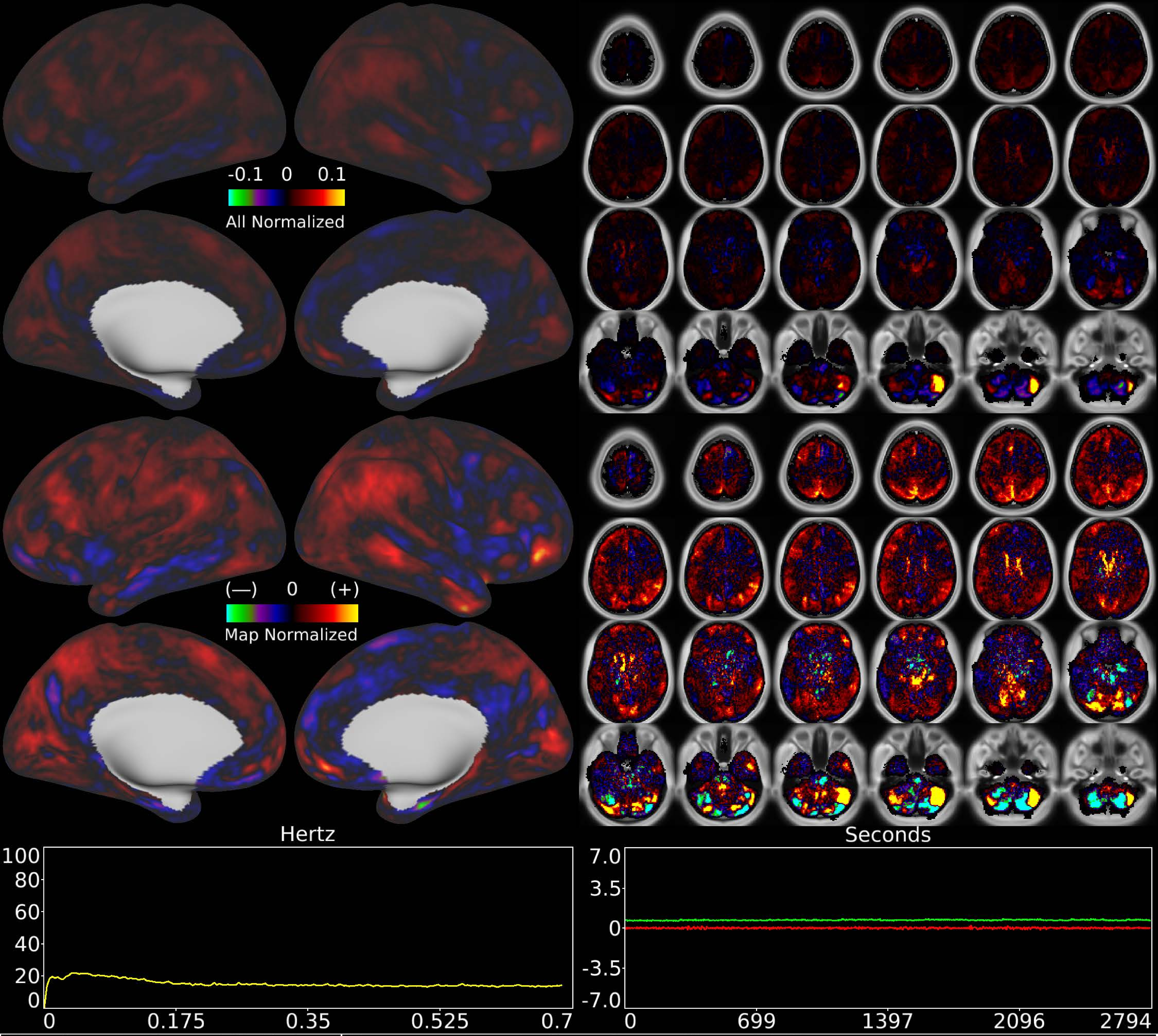


| | | |
|---|---|---|
| Number & Class: 52 Noise | Name: R Cerebellum Near Sigmoid Sinus | |
| RVT Correlated: No | DVARS Dip Associated: No | Cross-Subject Variable: No |
| Single Subject: Yes | % Variance Explained: 0.91 | Globality Index: 0.71 |
| Task Component: 48 | Rest Component: No | Task Modulated: No |
| Rationale: Classified same as TC48: single subject component whose spatial map is not reflective of known RSNs or areas | | |

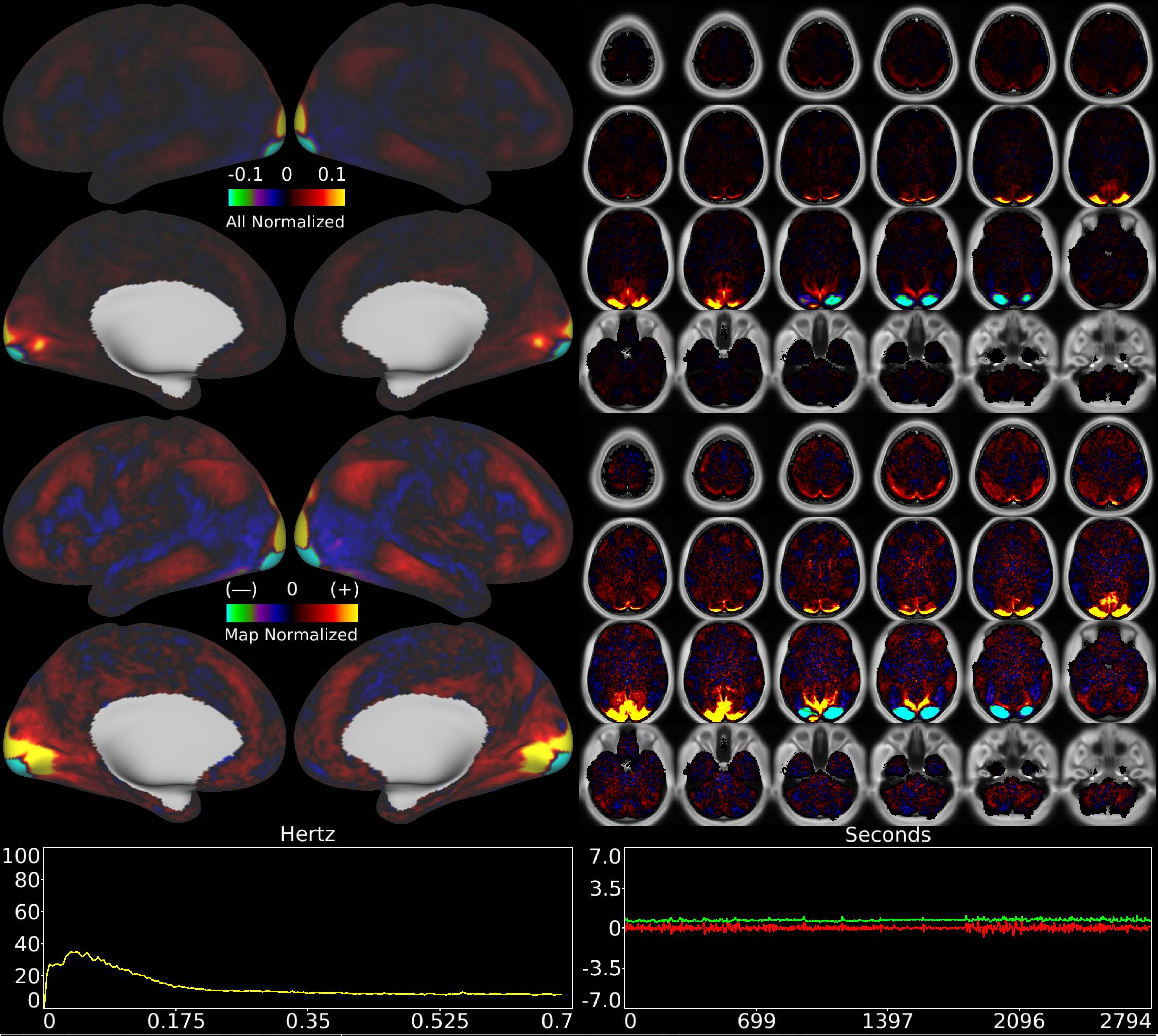


| | | |
|---|---|---|
| Number & Class: 53 Signal | Name: Visuotopic: Foveal Dorsal > Ventral | |
| RVT Correlated: No | DVARS Dip Associated: No | Cross-Subject Variable: No |
| Single Subject: No | % Variance Explained: 0.91 | Globality Index: 0.81 |
| Task Component: 65 | Rest Component: 76 | Task Modulated: No |
| Rationale: Spatial map includes positive and negative patches that respect known retinotopic visual organization (Foveal and Lower vs Upper) | | |

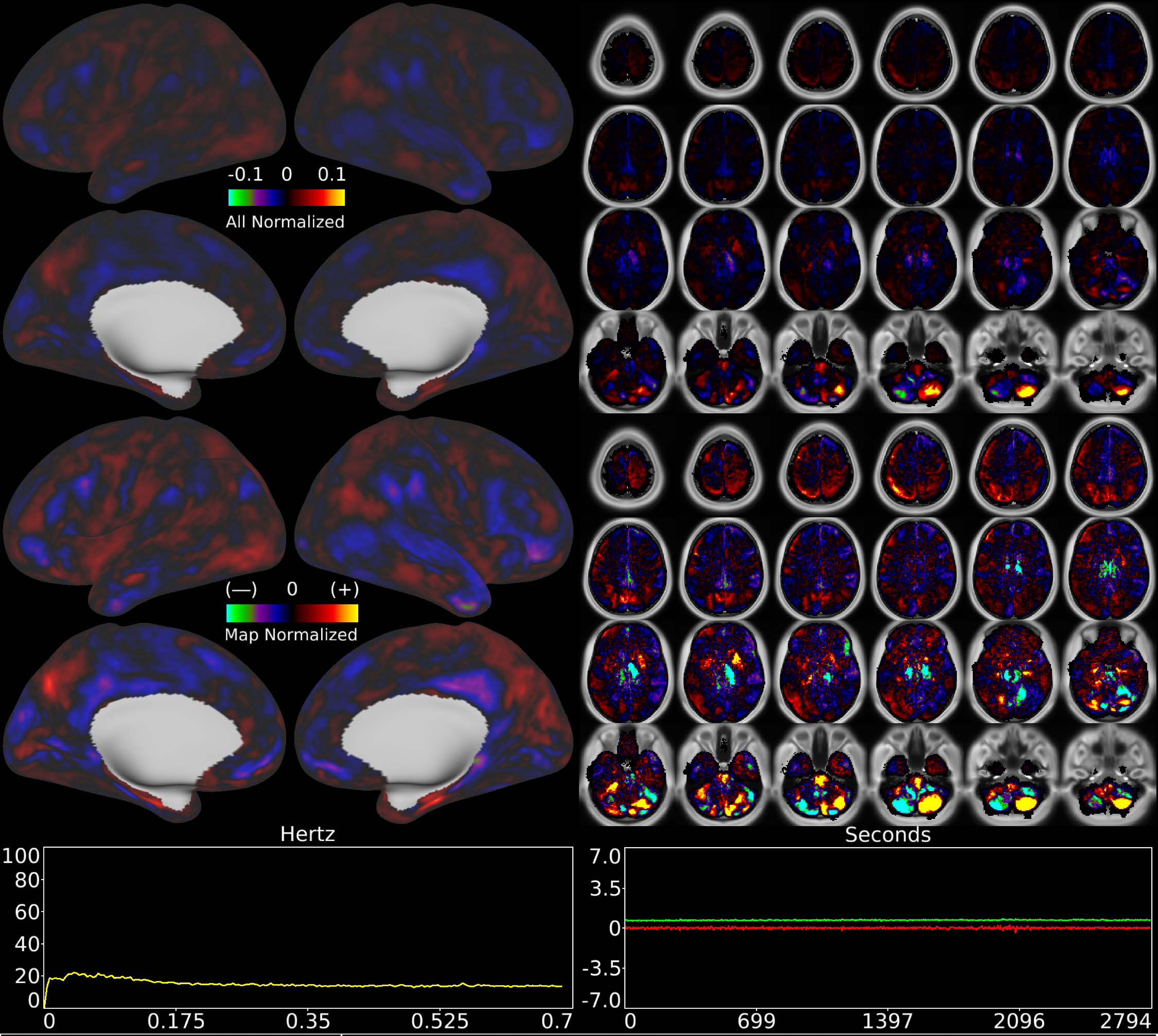


| | | |
|---|---|---|
| Number & Class: 54 Noise | Name: R Inferior Cerebellum > L Inferior Cerebellum | |
| RVT Correlated: No | DVARS Dip Associated: No | Cross-Subject Variable: No |
| Single Subject: Yes | % Variance Explained: 0.9 | Globality Index: 0.13 |
| Task Component: 55 | Rest Component: No | Task Modulated: No |
| Rationale: Single subject component not reflective of known areas or RSNs | | |

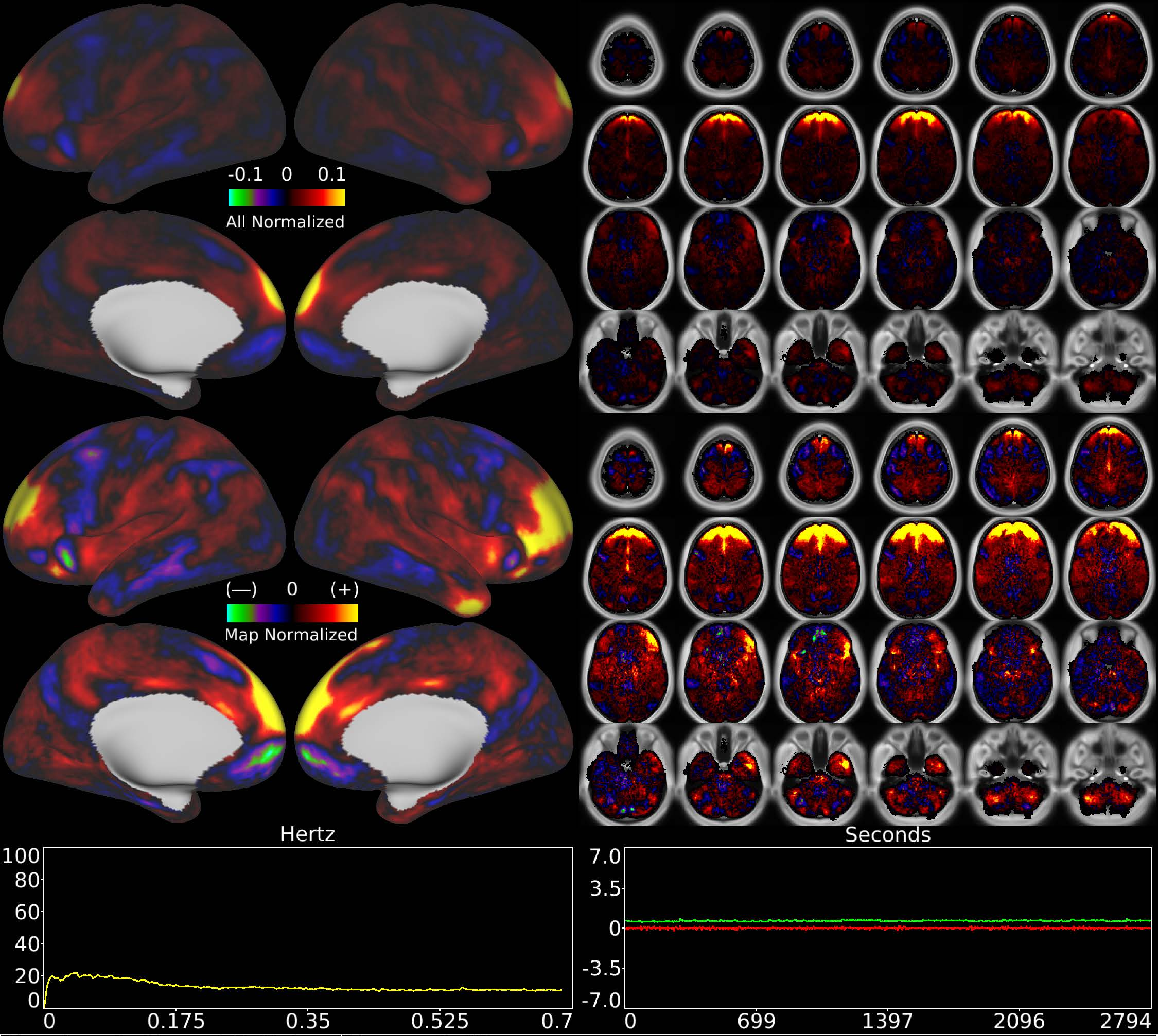


| | | |
|---|---|---|
| Number & Class: 55 Noise | Name: Coil? | |
| RVT Correlated: No | DVARS Dip Associated: Yes | Cross-Subject Variable: Yes |
| Single Subject: Yes | % Variance Explained: 0.78 | Globality Index: 1.25 |
| Task Component: 64 | Rest Component: 70 | Task Modulated: No |
| Rationale: Spatial map is a band across frontal cortex and white matter; could be related to coil; head motion; or eye motion | | |

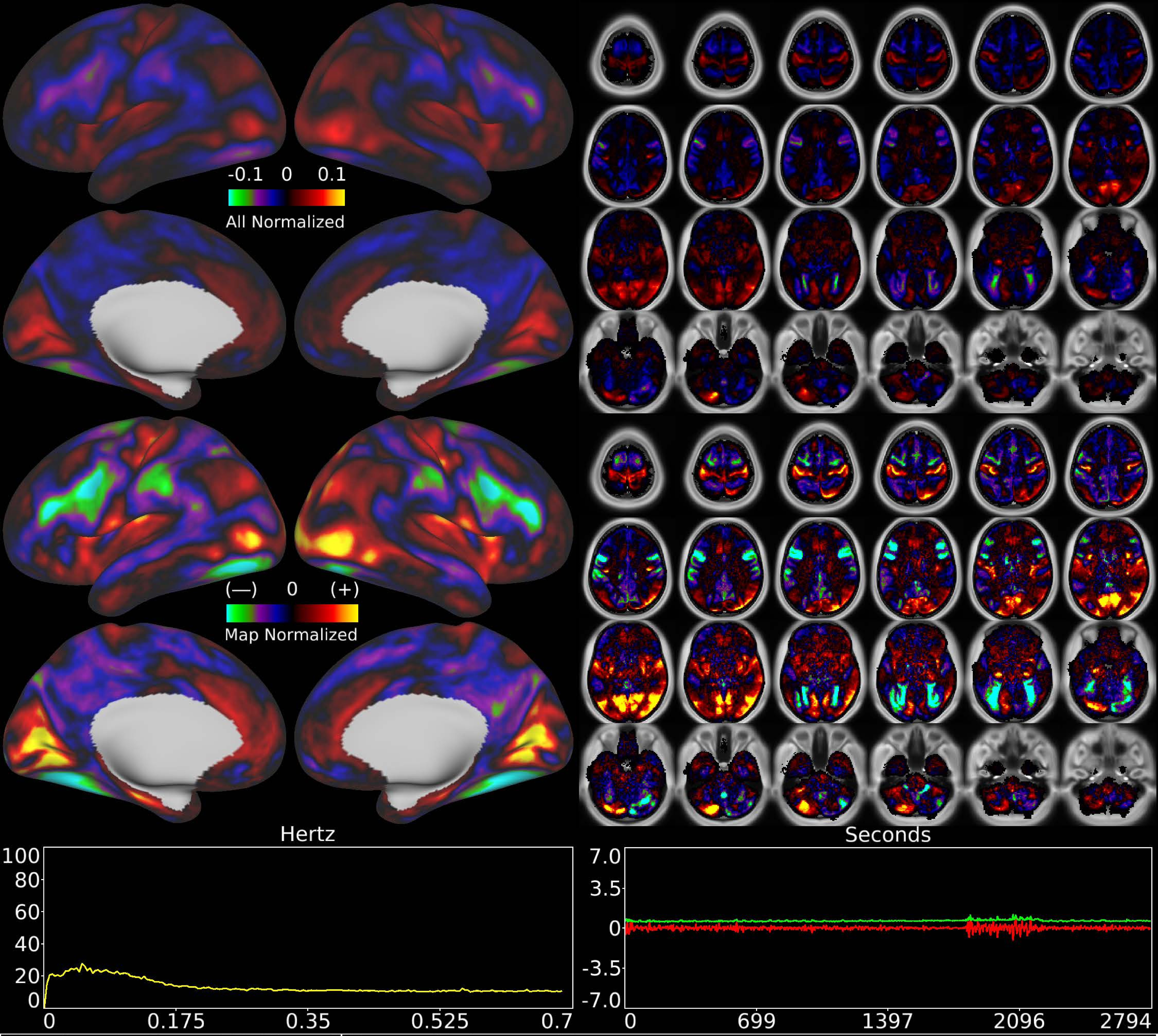


| | | |
|---|---|---|
| Number & Class: 56 Signal | Name: Unknown Network | |
| RVT Correlated: No | DVARS Dip Associated: No | Cross-Subject Variable: No |
| Single Subject: No | % Variance Explained: 0.77 | Globality Index: 0.12 |
| Task Component: No | Rest Component: No | Task Modulated: No |
| Rationale: Spatial map has elements of visuotopy and known RSNs though in an atypical arrangement | | |

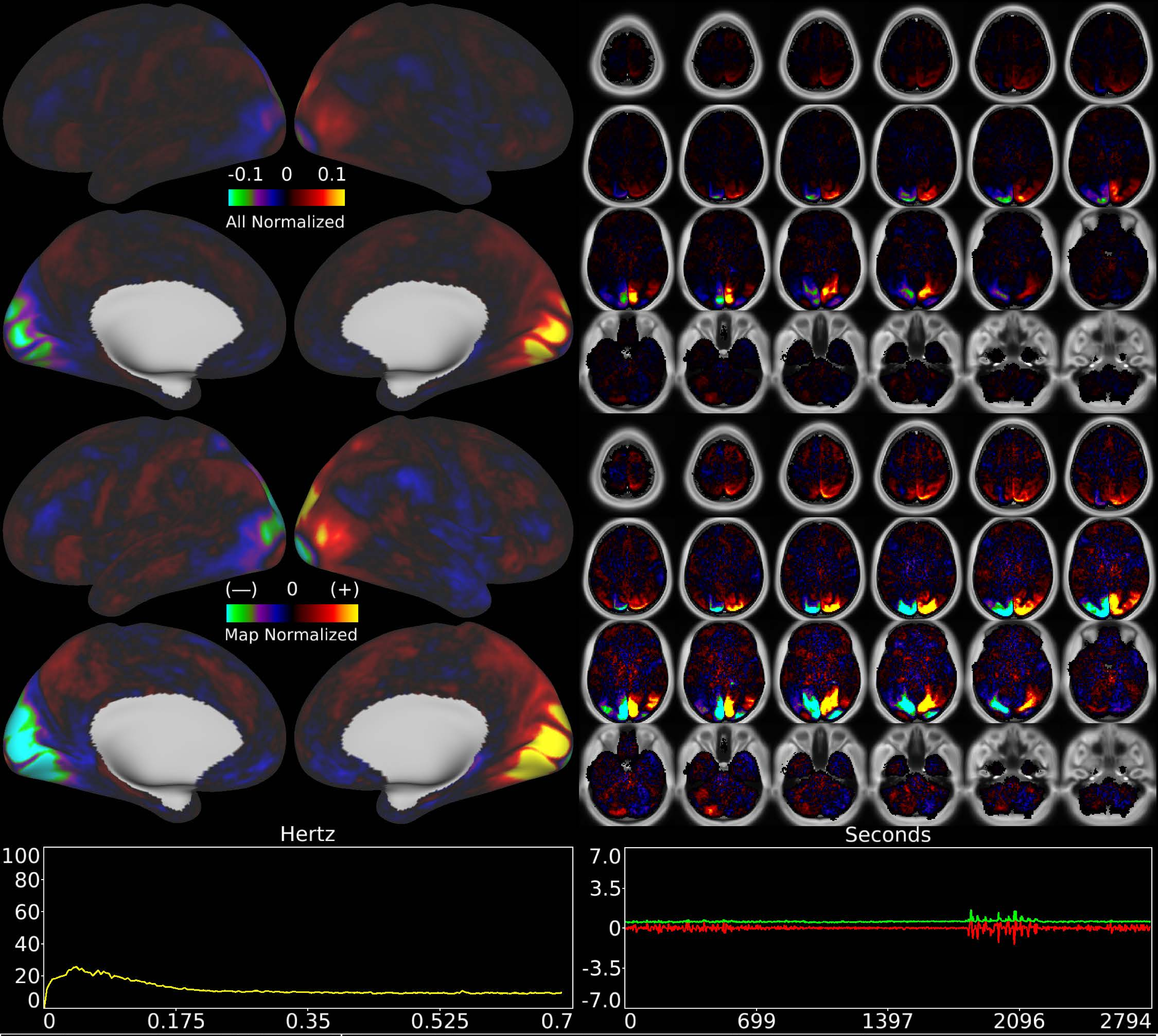


| | | |
|---|---|---|
| Number & Class: 57 Signal | Name: Visuotopic: Paracentral Right > Left | |
| RVT Correlated: No | DVARS Dip Associated: No | Cross-Subject Variable: No |
| Single Subject: No | % Variance Explained: 0.63 | Globality Index: 0.29 |
| Task Component: -69 | Rest Component: 84 | Task Modulated: Social |

Rationale: Spatial map includes positive and negative patches that respect known retinotopic visual organization (Peripheral Right vs Left)

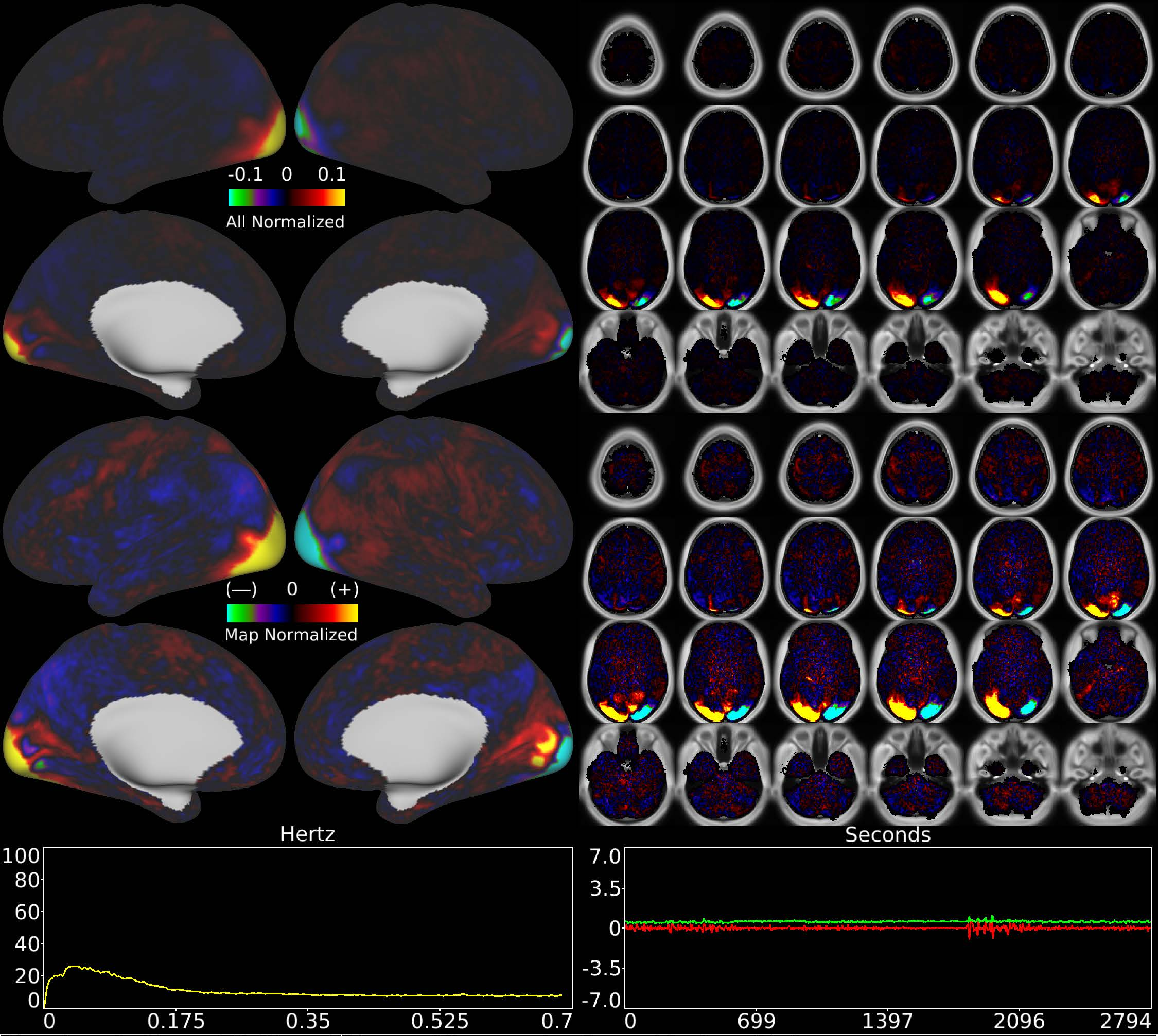


| | | |
|---|---|---|
| Number & Class: 58 Signal | Name: Visuotopic: Foveal Left > Right | |
| RVT Correlated: No | DVARS Dip Associated: No | Cross-Subject Variable: No |
| Single Subject: No | % Variance Explained: 0.61 | Globality Index: 0.02 |
| Task Component: 70 | Rest Component: 81 | Task Modulated: No |

Rationale: Spatial map includes positive and negative patches that respect known retinotopic visual organization (Foveal Right vs Left)
