## Supplementary Materials for "Using Temporal ICA to Selectively Remove Global Noise While Preserving Global Signal in Functional MRI Data"

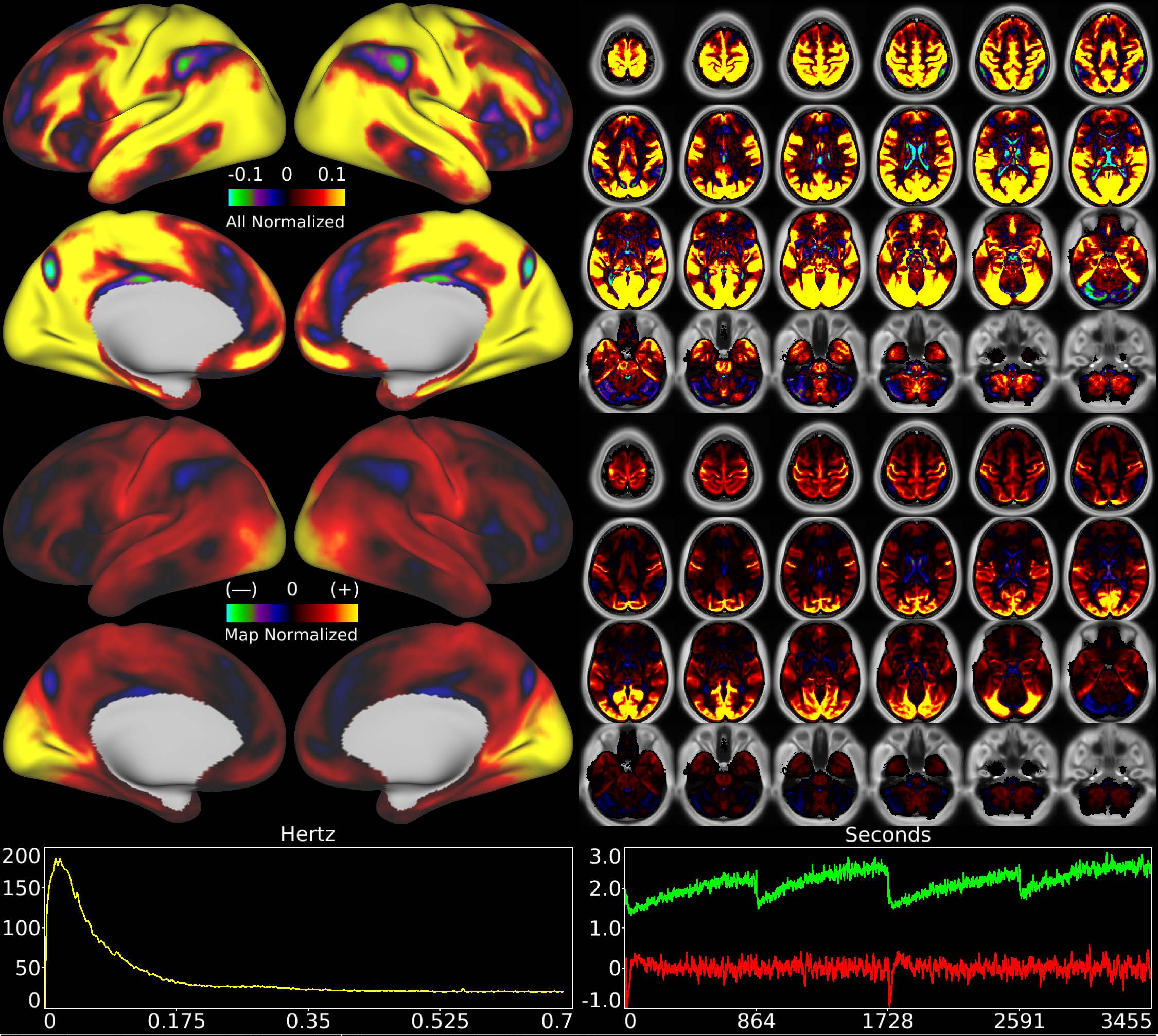


| | | |
|---|---|---|
| Number & Class: 1 Signal | Name: Main Sleep Component | |
| RVT Correlated: No | DVARS Dip Associated: No | Cross-Subject Variable: Yes |
| Single Subject: No | % Variance Explained: 8.47 | Globality Index: 2.35 |
| Task Component: No | Taskr Component: No | Sleep Noted: Yes |
| Rationale: Spatial map includes positive and negative patches that respect known areal boundaries (e.g. POS2 and RSC) | | |

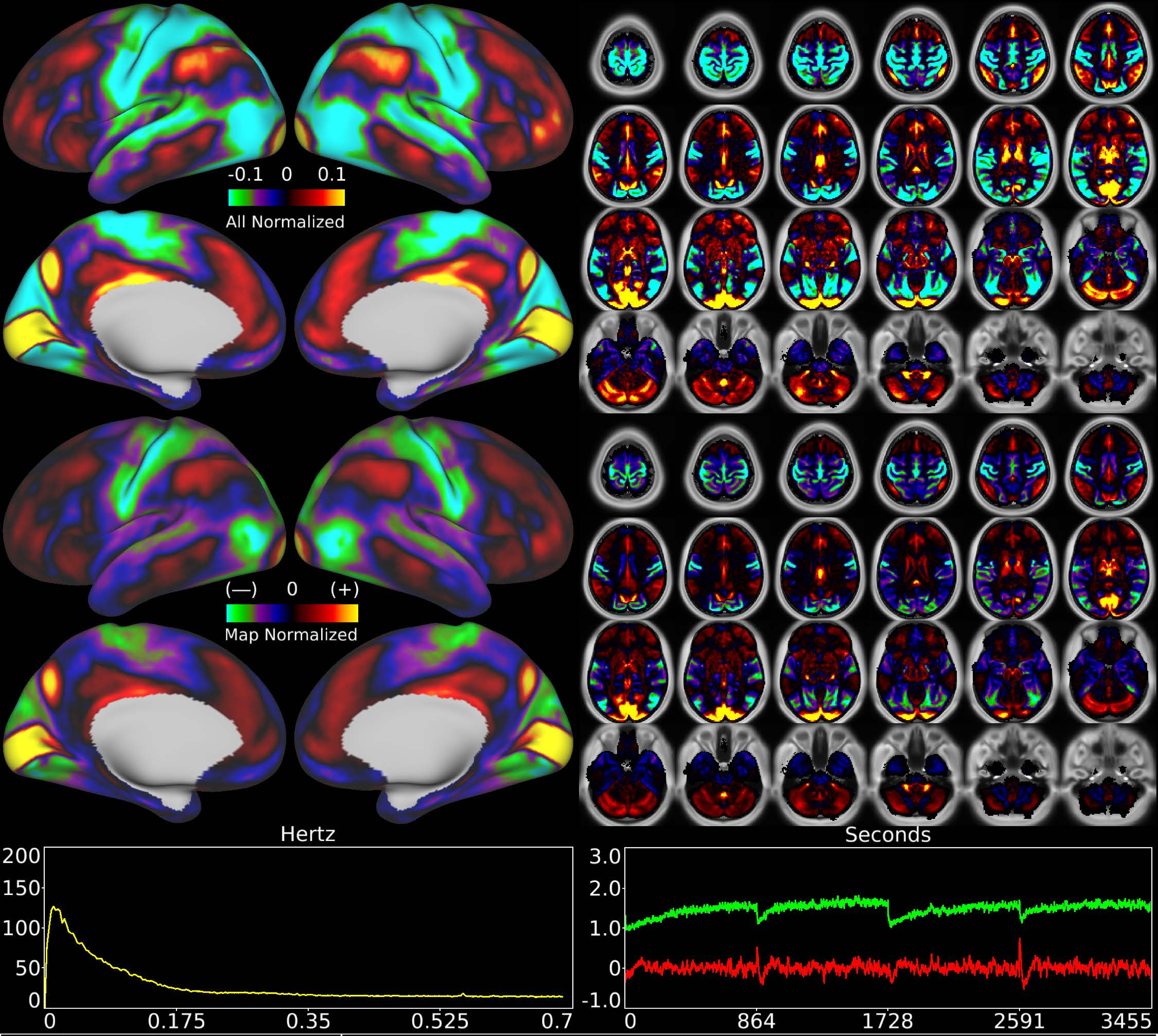


| | | |
|---|---|---|
| Number & Class: 2 Signal | Name: LGN to V1 Variable Component | |
| RVT Correlated: No | DVARS Dip Associated: No | Cross-Subject Variable: Yes |
| Single Subject: No | % Variance Explained: 3.64 | Globality Index: 0.24 |
| Task Component: 14 | Taskr Component: 14 | Sleep Noted: No |
| Rationale: Spatial map includes positive and negative patches that respect known areal boundaries (e.g. V1 and V2) | | |

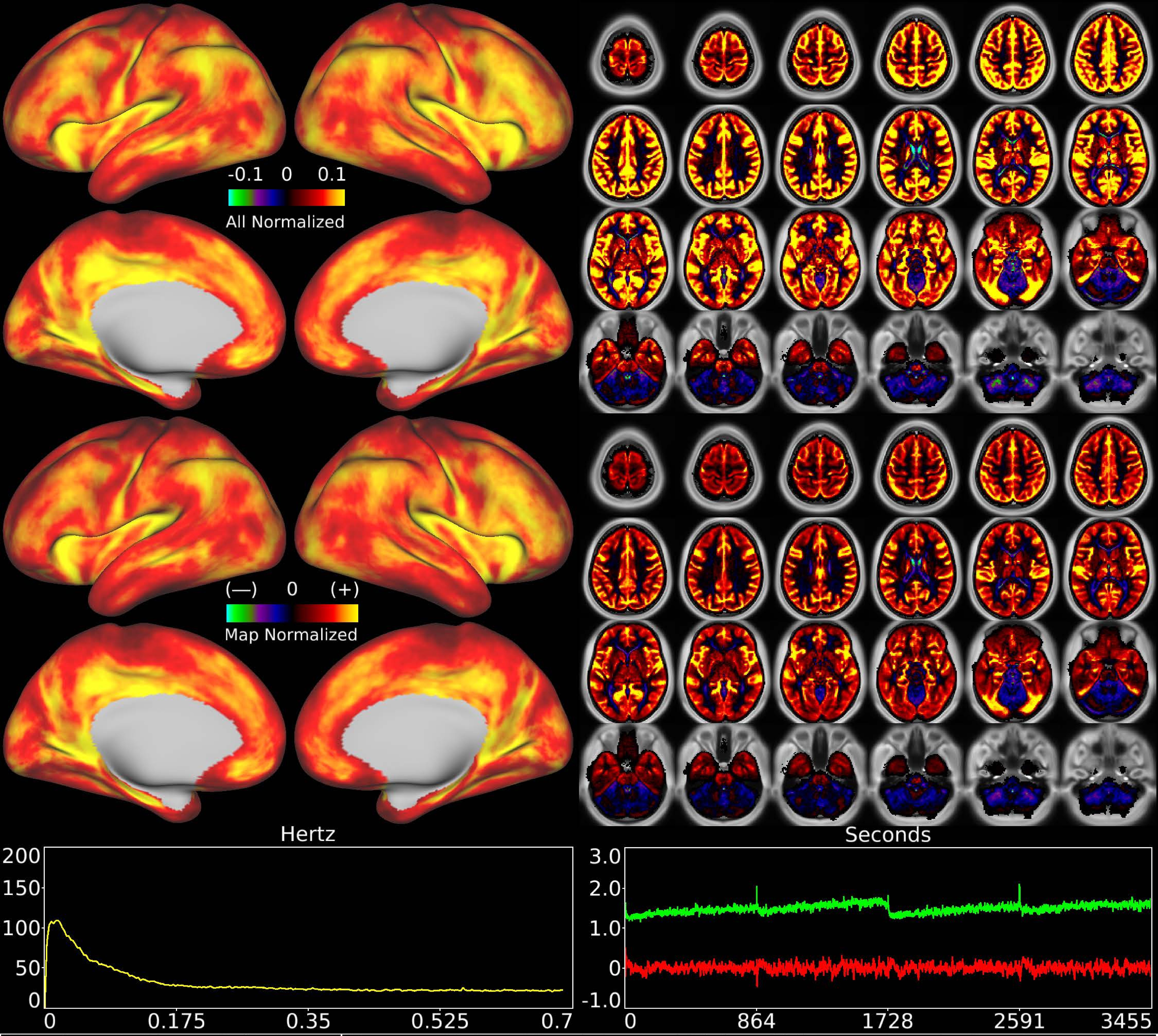


| | | |
|---|---|---|
| Number & Class: 3 Noise | Name: Global Physiological Noise (Without Cerebellum) | |
| RVT Correlated: Yes | DVARS Dip Associated: Yes | Cross-Subject Variable: No |
| Single Subject: No | % Variance Explained: 3.13 | Globality Index: 2.23 |
| Task Component: 1 | Taskr Component: 1 | Sleep Noted: No |

Rationale: Globally positive component except for cerebellum that is globally negative; which could be due to differing vascular supplies of these structures

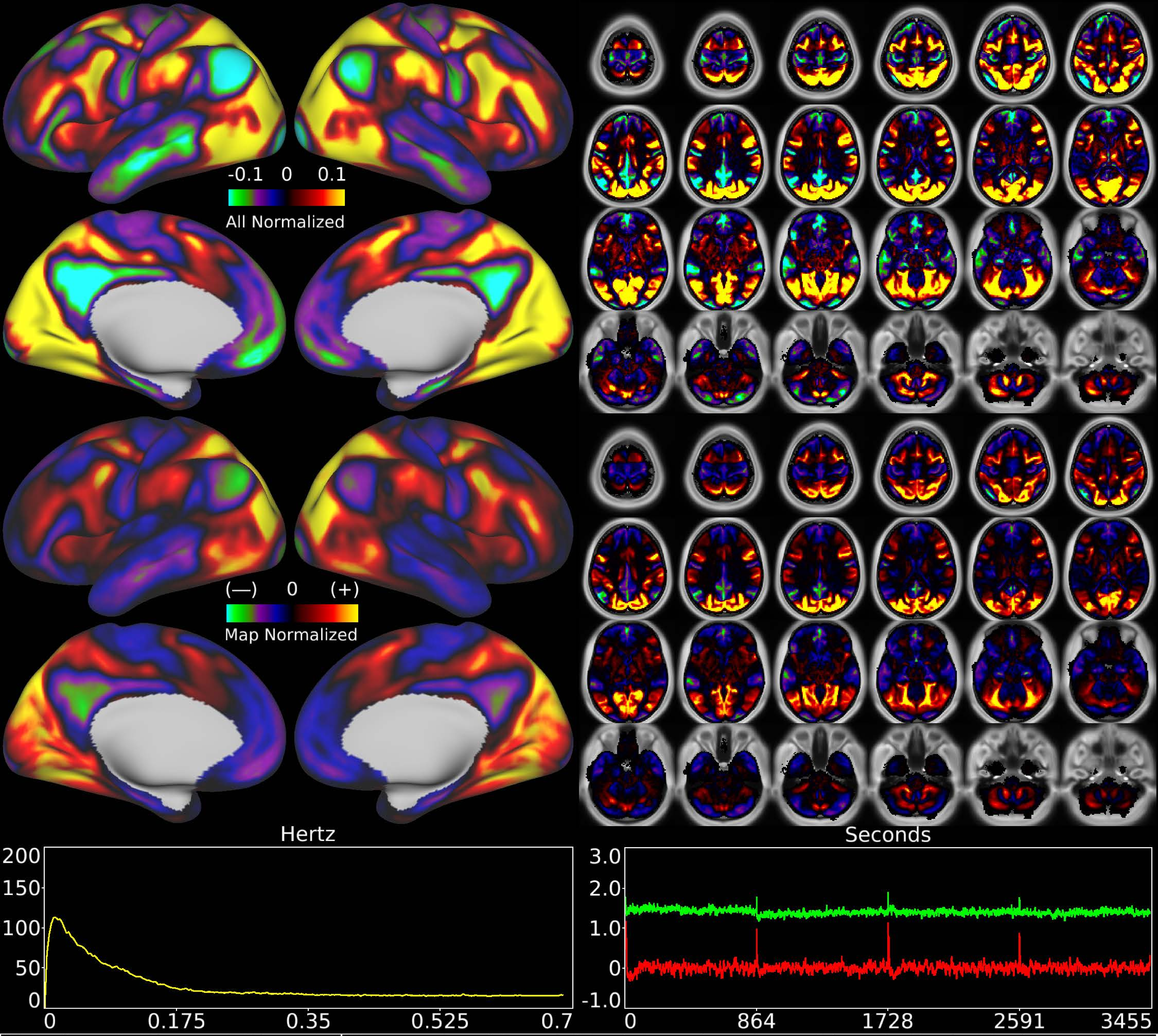


| | | |
|---|---|---|
| Number & Class: 4 Signal | Name: Pan Visual (Peripheral > FoveaL) > Default Mode | |
| RVT Correlated: No | DVARS Dip Associated: No | Cross-Subject Variable: No |
| Single Subject: No | % Variance Explained: 3.15 | Globality Index: 0.16 |
| Task Component: No | Taskr Component: 3 | Sleep Noted: No |
| Rationale: Spatial map includes positive and negative patches that respect known RSNs (e.g. Default Mode Network) | | |

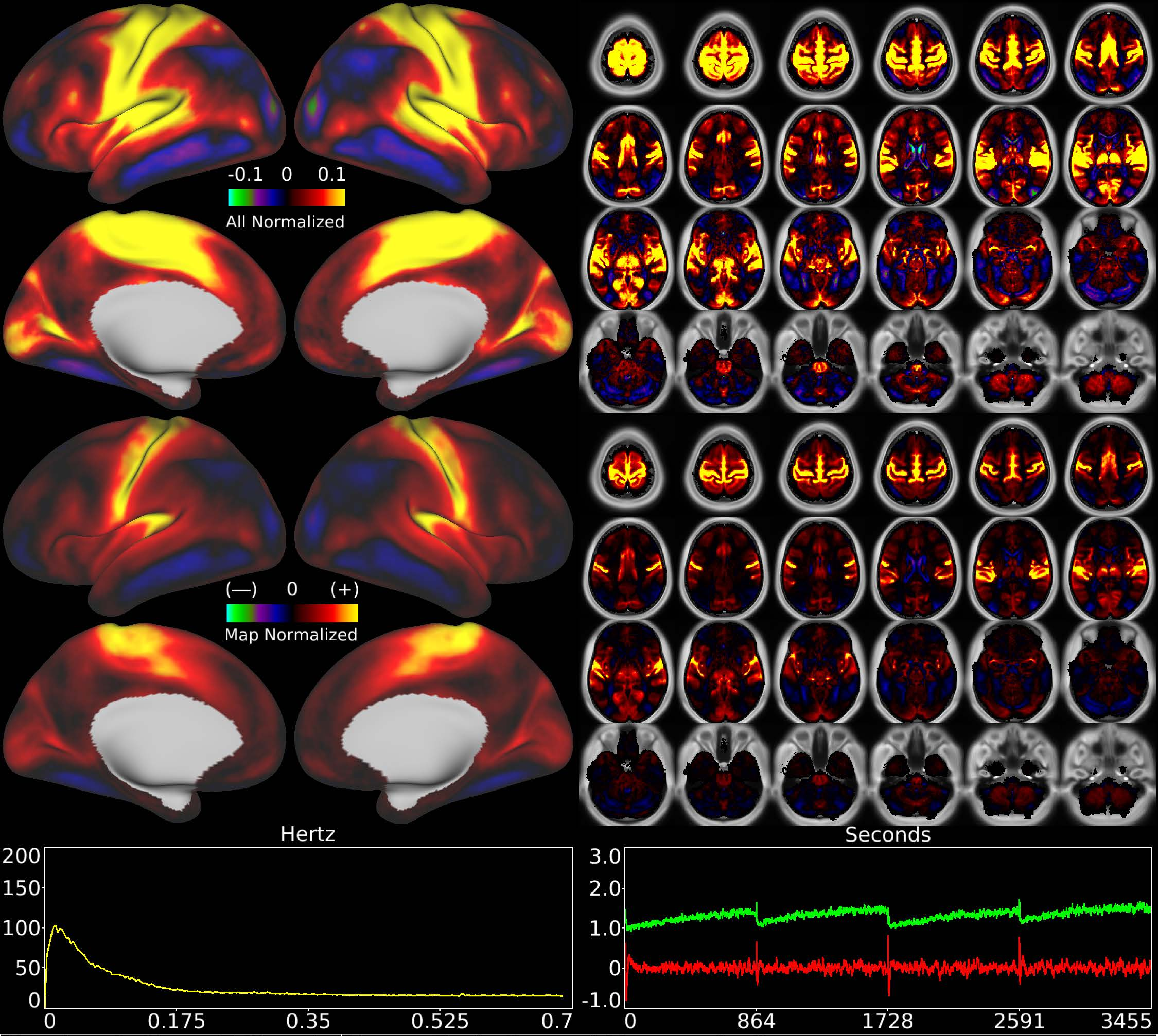


| | | |
|---|---|---|
| Number & Class: 5 Signal | Name: Early Sensori-Motor + Auditory | |
| RVT Correlated: No | DVARS Dip Associated: No | Cross-Subject Variable: Yes |
| Single Subject: No | % Variance Explained: 2.87 | Globality Index: 1.84 |
| Task Component: 23 | Taskr Component: No | Sleep Noted: Yes |
| Rationale: Spatial map includes positive and negative patches that respect known areal boundaries (e.g. around MT+); most specific to sensori-motor cortex | | |

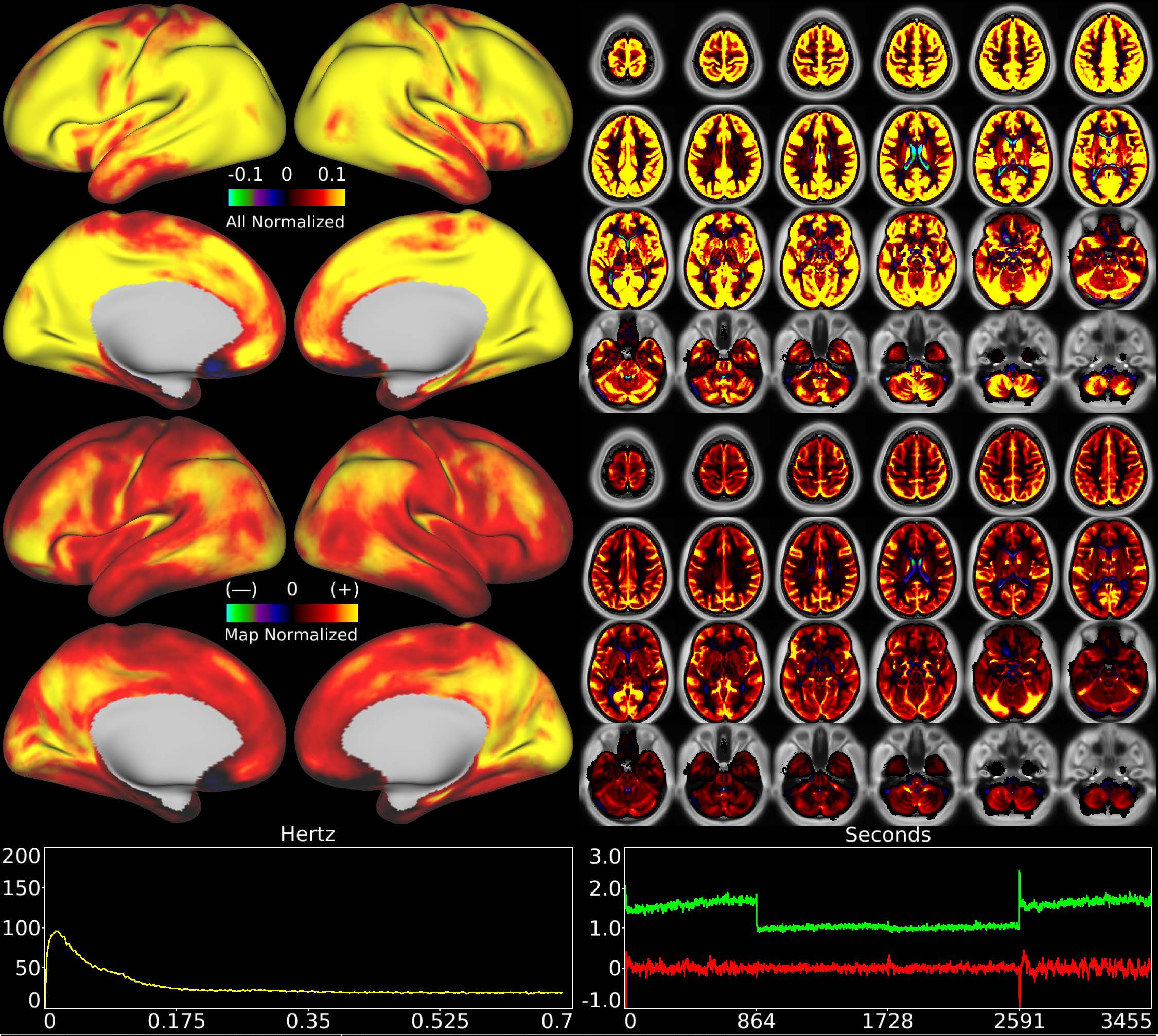


| | | |
|---|---|---|
| Number & Class: 6 Noise | Name: Global Physiological Noise | |
| RVT Correlated: Yes | DVARS Dip Associated: Yes | Cross-Subject Variable: No |
| Single Subject: No | % Variance Explained: 2.86 | Globality Index: 5.67 |
| Task Component: 1 | Taskr Component: 1 | Sleep Noted: No |

Rationale: Globally positive component matching hypothesized pattern for physiological artifact; does not clearly follow known areal boundaries

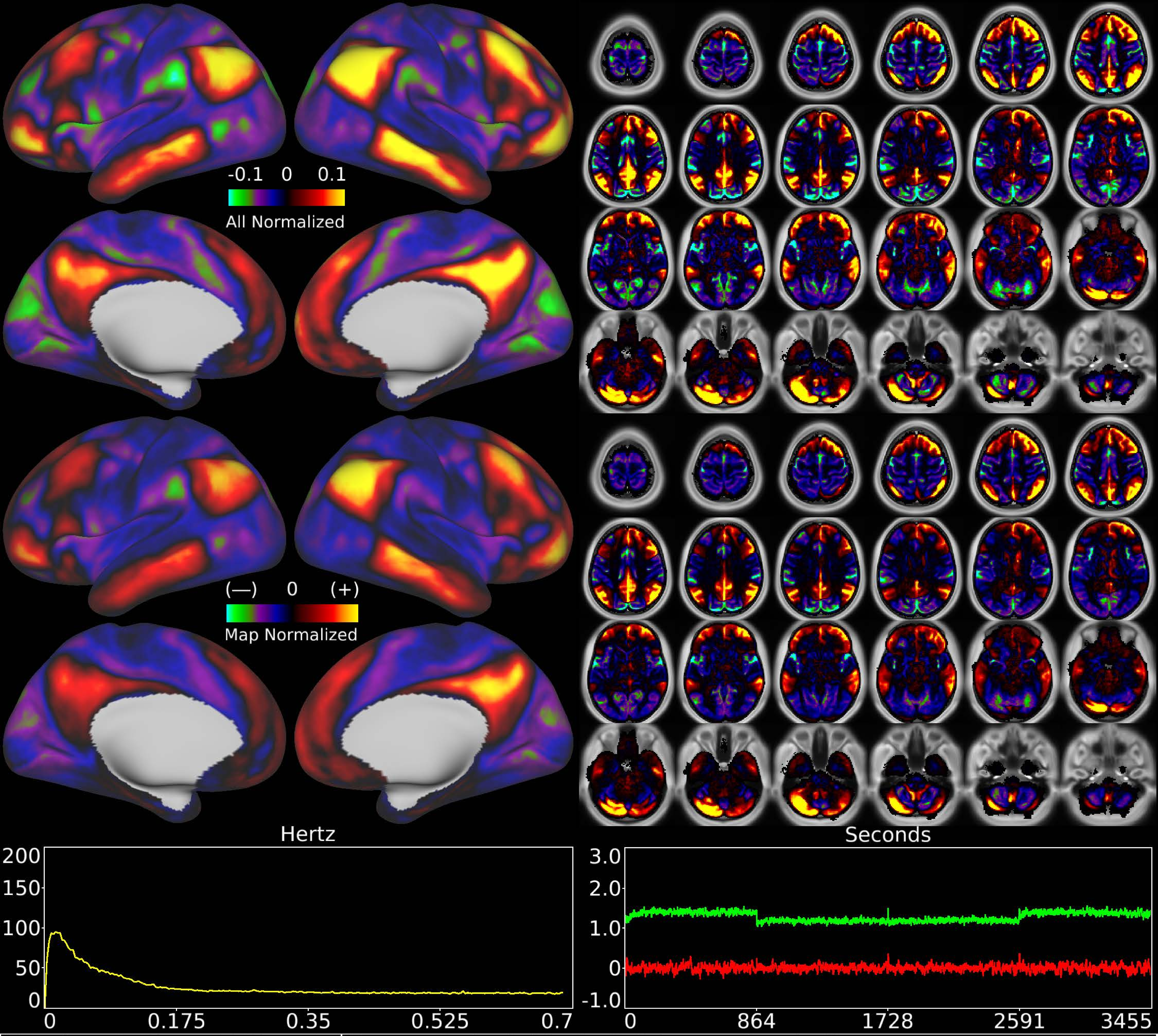


| | | |
|---|---|---|
| Number & Class: 7 Signal | Name: Default Mode R Primary Cerebellar | |
| RVT Correlated: No | DVARS Dip Associated: No | Cross-Subject Variable: No |
| Single Subject: No | % Variance Explained: 2.54 | Globality Index: 0.74 |
| Task Component: No | Taskr Component: No | Sleep Noted: No |
| Rationale: Spatial map includes positive and negative patches that respect known RSNs (e.g. Default Mode Network) | | |

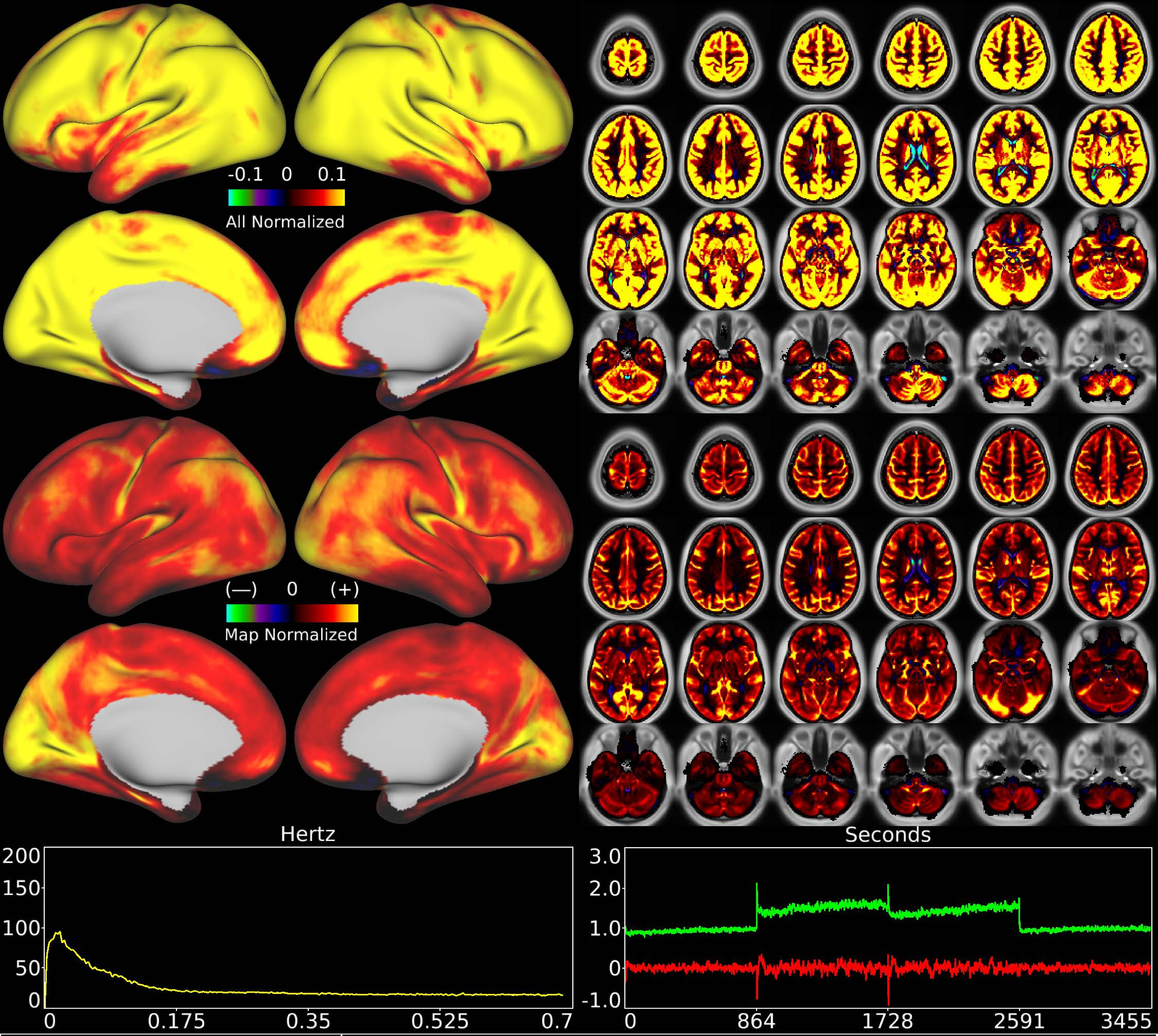


| | | |
|---|---|---|
| Number & Class: 8 Noise | Name: Global Physiological Noise | |
| RVT Correlated: Yes | DVARS Dip Associated: Yes | Cross-Subject Variable: No |
| Single Subject: No | % Variance Explained: 2.51 | Globality Index: 5.94 |
| Task Component: 1 | Taskr Component: 1 | Sleep Noted: No |

Rationale: Globally positive component matching hypothesized pattern for physiological artifact; does not clearly follow known areal boundaries

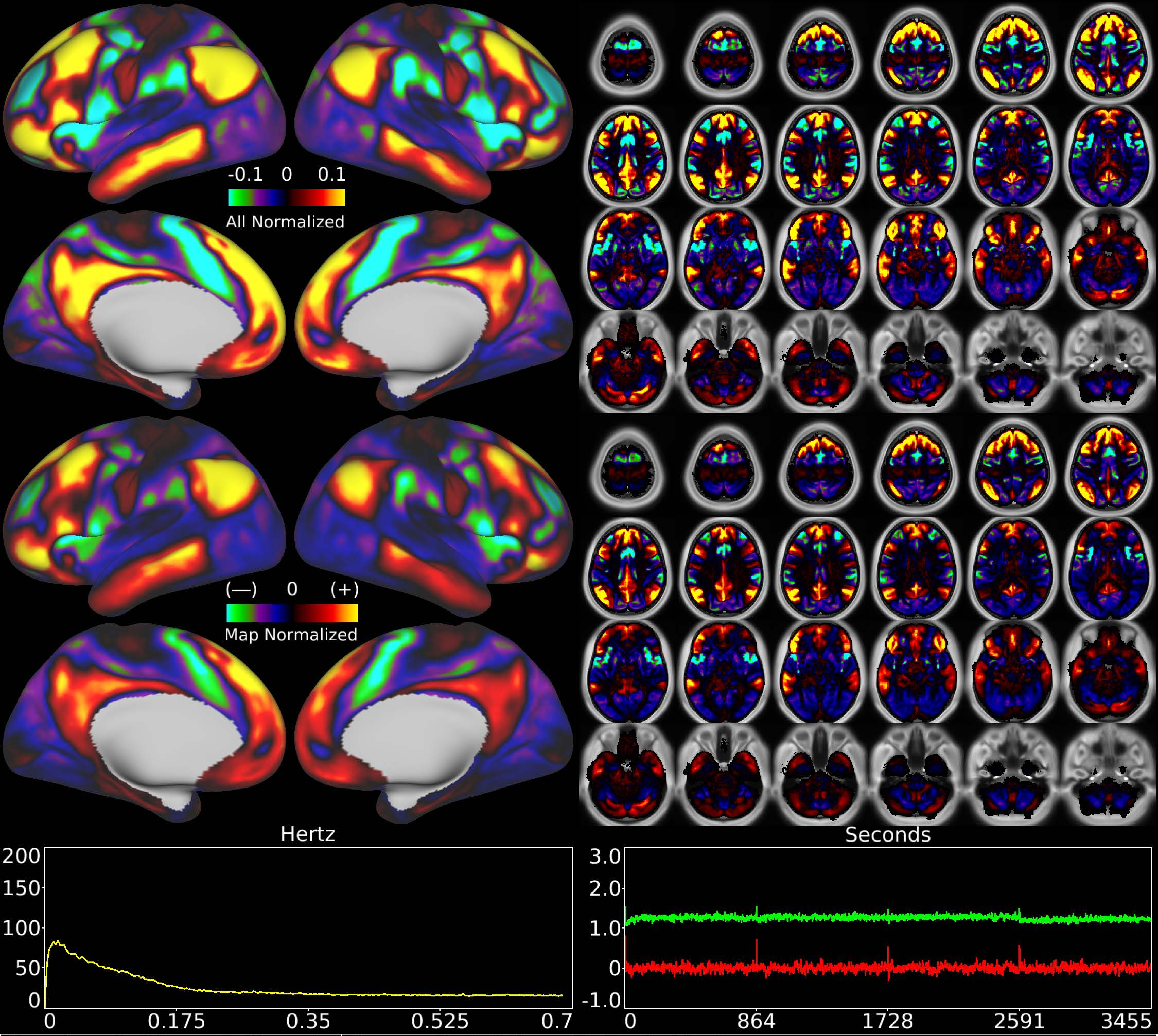


| | | |
|---|---|---|
| Number & Class: 9 Signal | Name: Subsidiary Default Mode | |
| RVT Correlated: No | DVARS Dip Associated: No | Cross-Subject Variable: No |
| Single Subject: No | % Variance Explained: 2.44 | Globality Index: 0.43 |
| Task Component: -9 | Taskr Component: 8 | Sleep Noted: No |
| Rationale: Spatial map includes positive and negative patches that respect known RSNs (e.g. Default Mode Network) | | |

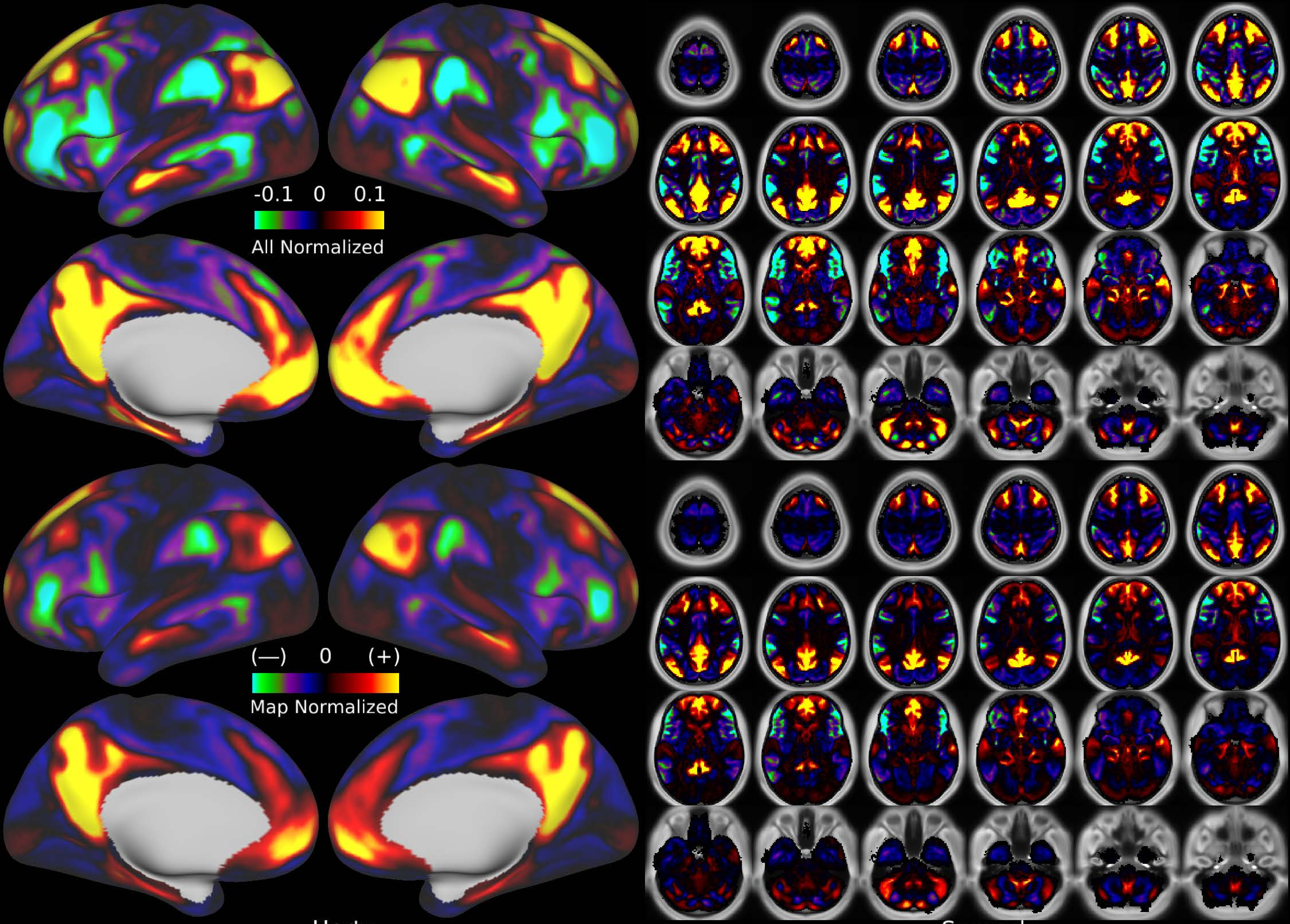


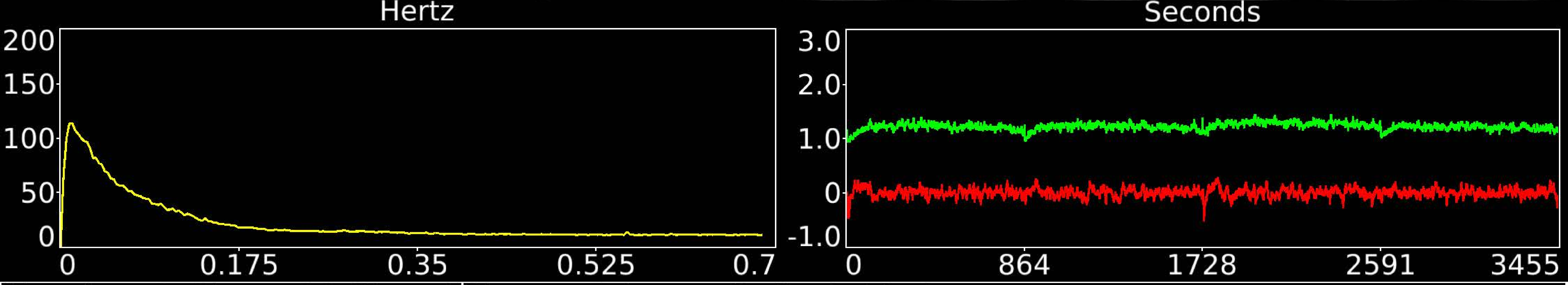


| | | |
|---|---|---|
| Number & Class: 10 Signal | Name: Primary Default Mode | |
| RVT Correlated: No | DVARS Dip Associated: No | Cross-Subject Variable: No |
| Single Subject: No | % Variance Explained: 2.39 | Globality Index: 0.35 |
| Task Component: 5 | Taskr Component: 2 | Sleep Noted: No |
| Rationale: Spatial map includes positive and negative patches that respect known RSNs (e.g. Default Mode Network) | | |

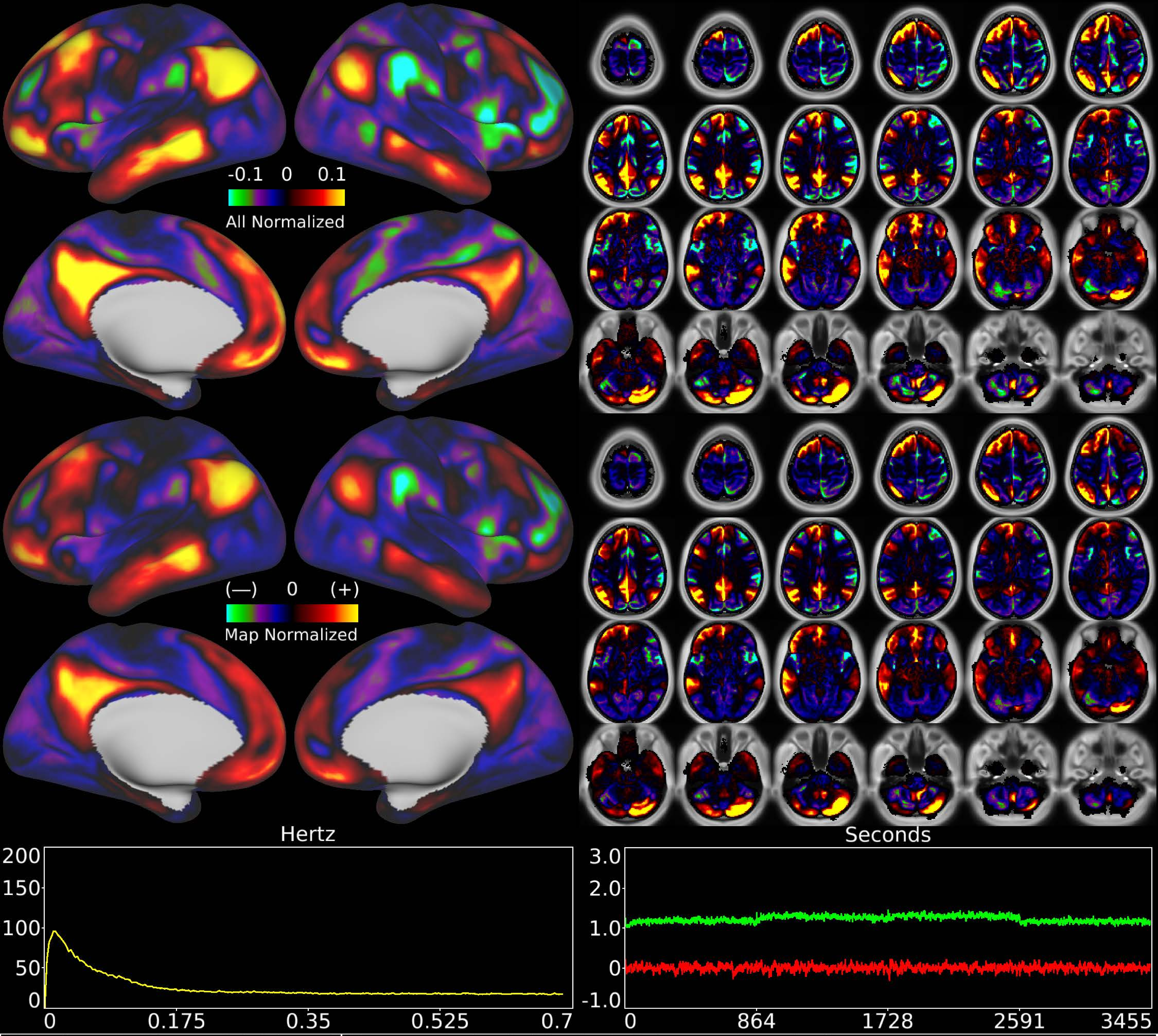


| | | |
|---|---|---|
| Number & Class: 11 Signal | Name: Default Mode L Primary Cerebellar | |
| RVT Correlated: No | DVARS Dip Associated: No | Cross-Subject Variable: No |
| Single Subject: No | % Variance Explained: 2.33 | Globality Index: 0.91 |
| Task Component: No | Taskr Component: No | Sleep Noted: No |
| Rationale: Spatial map includes positive and negative patches that respect known RSNs (e.g. Default Mode Network) | | |

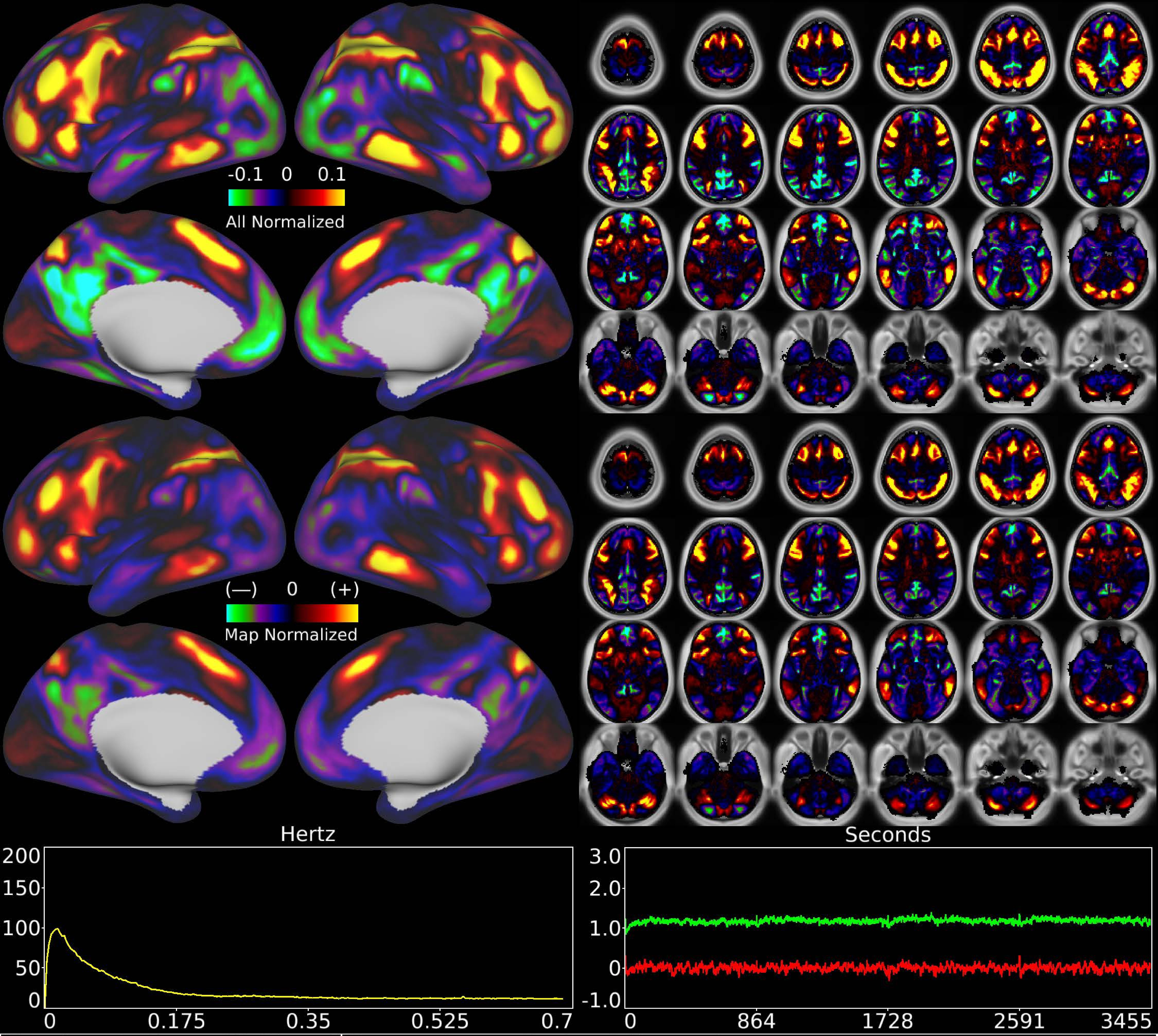


| | | |
|---|---|---|
| Number & Class: 12 Signal | Name: Fronto-Parietal Network | |
| RVT Correlated: No | DVARS Dip Associated: No | Cross-Subject Variable: No |
| Single Subject: No | % Variance Explained: 2.25 | Globality Index: 0.56 |
| Task Component: 4 | Taskr Component: 12+17+23 | Sleep Noted: No |
| Rationale: Spatial map includes positive and negative patches that respect known RSNs (e.g. Fronto-Parietal Network) | | |

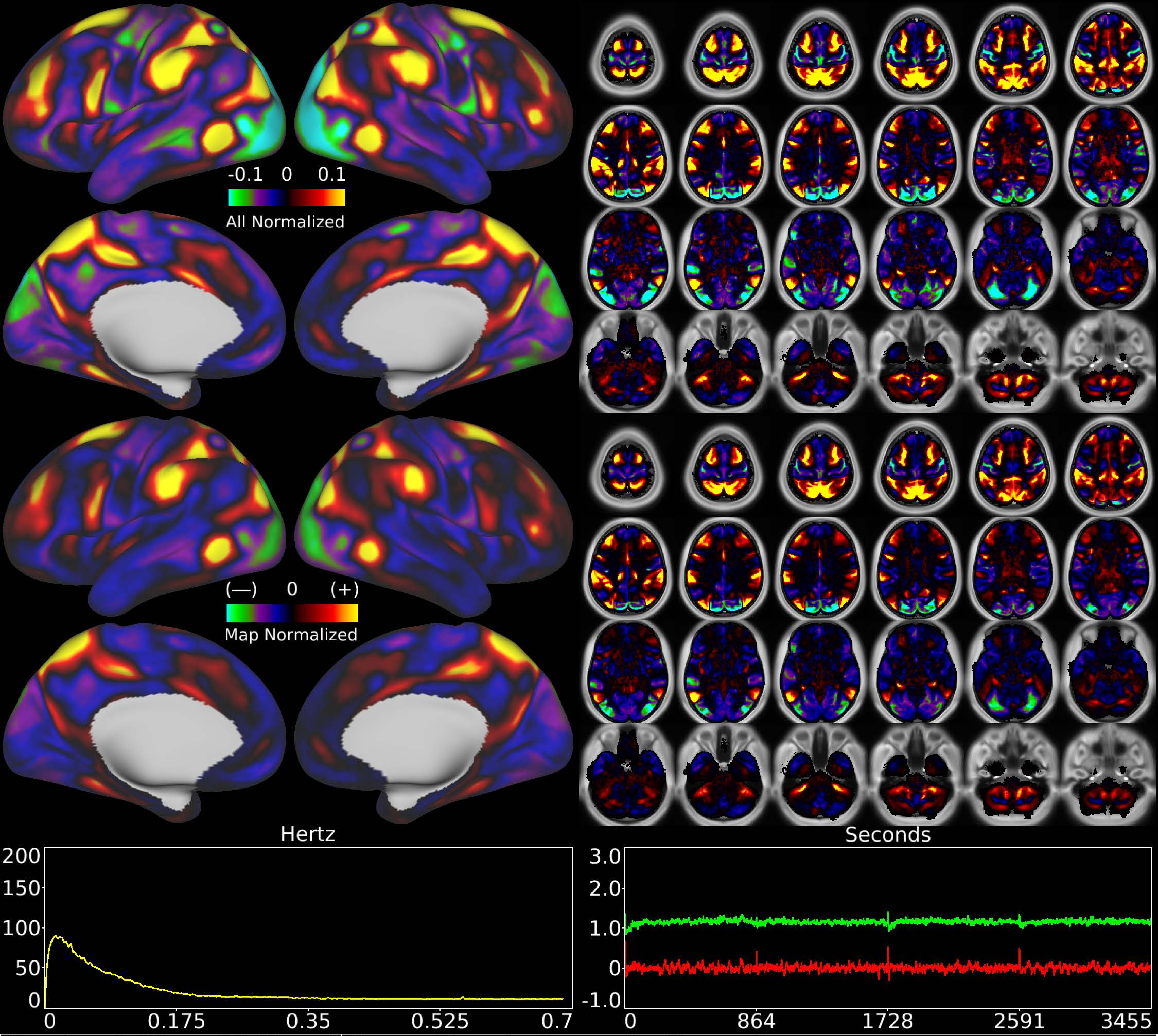


| | | |
|---|---|---|
| Number & Class: 13 Signal | Name: Cingulo-Opercular Network | |
| RVT Correlated: No | DVARS Dip Associated: No | Cross-Subject Variable: No |
| Single Subject: No | % Variance Explained: 2.15 | Globality Index: 0.34 |
| Task Component: 10 | Taskr Component: 9 | Sleep Noted: No |
| Rationale: Spatial map includes positive and negative patches that respect known RSNs (e.g. Cingulo-Opercular Network) | | |

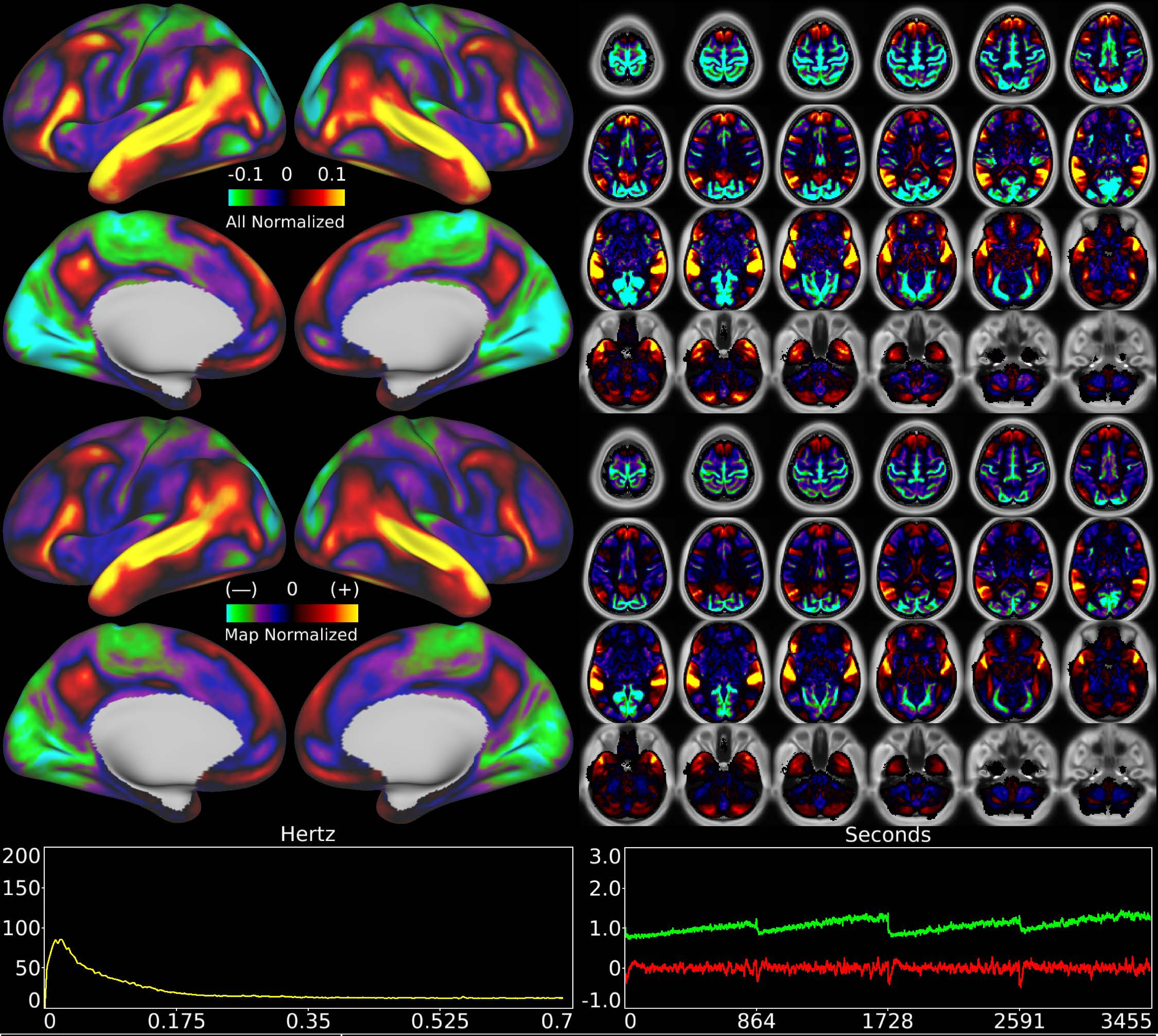


| | | |
|---|---|---|
| Number & Class: 14 Signal | Name: Subsidiary Sleep Component | |
| RVT Correlated: No | DVARS Dip Associated: No | Cross-Subject Variable: Yes |
| Single Subject: No | % Variance Explained: 2.11 | Globality Index: 1.07 |
| Task Component: No | Taskr Component: No | Sleep Noted: Yes |
| Rationale: Spatial map includes positive and negative patches that respect known RSNs (e.g. Ventral Attention; Language Network) and areal boundaries (44 and 45) | | |

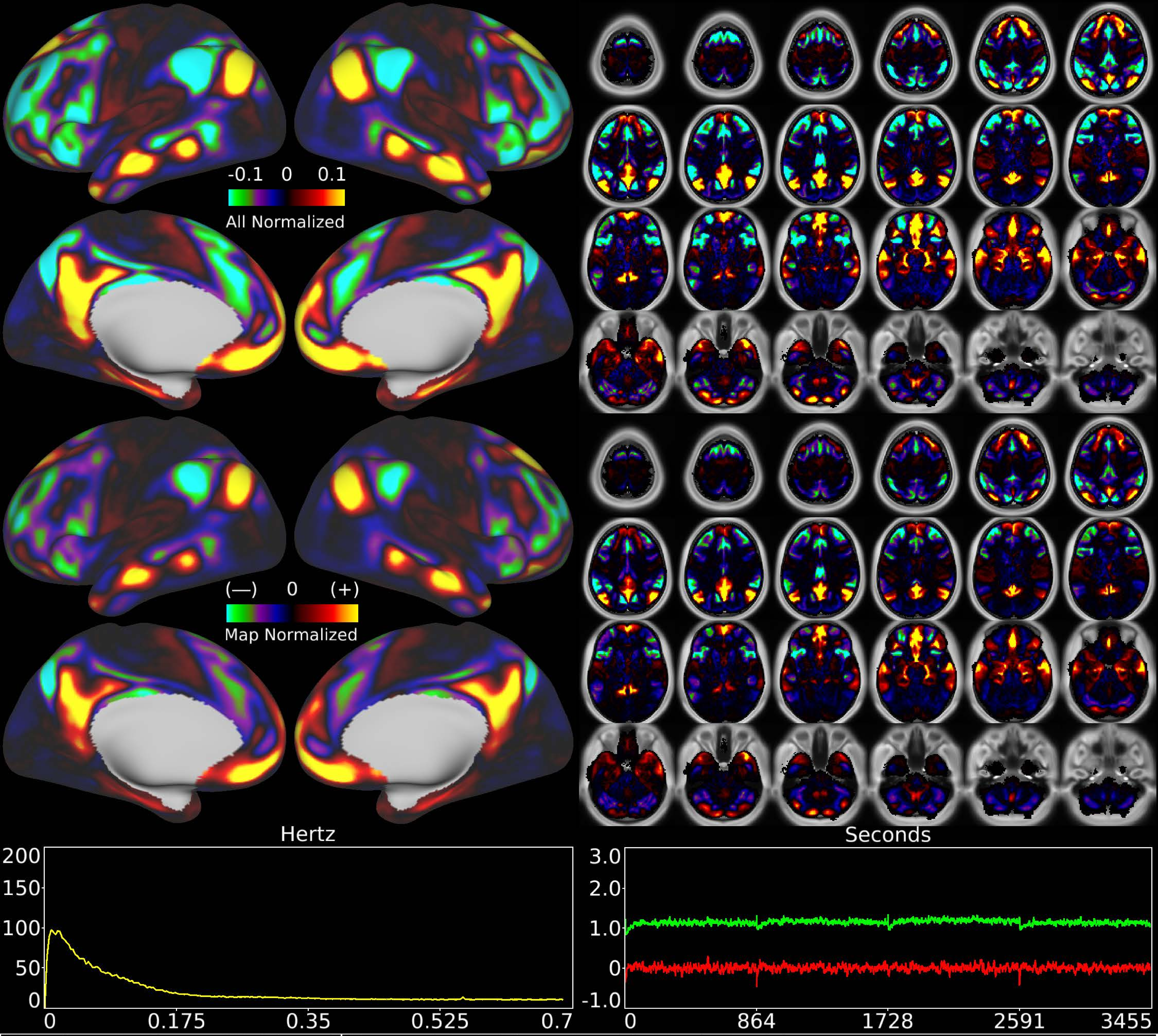


| | | |
|---|---|---|
| Number & Class: 15 Signal | Name: Task Negative Vs Task Positive | |
| RVT Correlated: No | DVARS Dip Associated: No | Cross-Subject Variable: No |
| Single Subject: No | % Variance Explained: 2.09 | Globality Index: 0.57 |
| Task Component: 26 | Taskr Component: 19 | Sleep Noted: No |
| Rationale: Spatial map includes positive and negative patches that respect known RSNs (e.g. Default Mode Network) | | |

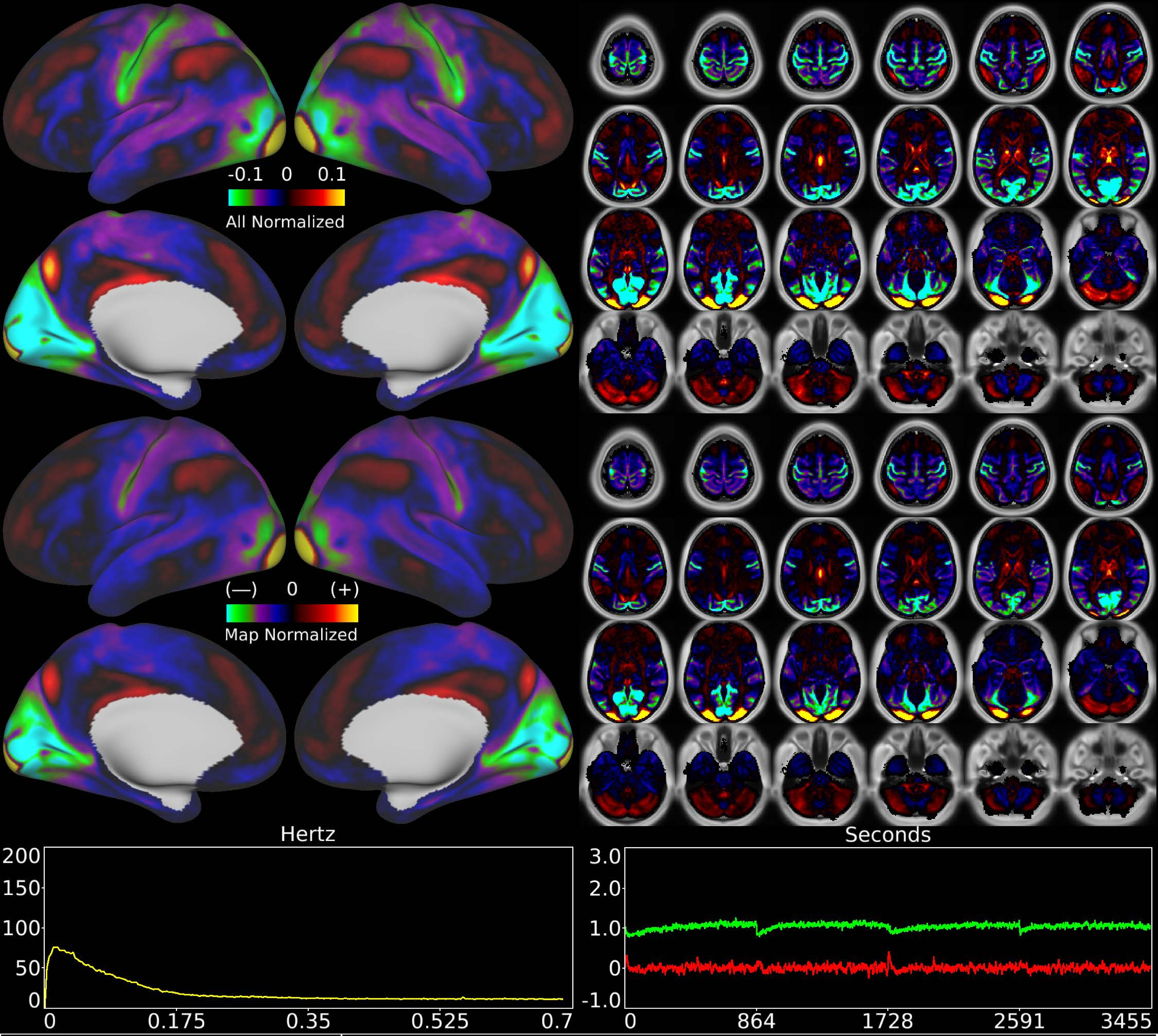


| Number & Class: 16 Signal | Name: Foveal > Peripheral Visual (Cross Hairs?) | |
|---|---|---|
| RVT Correlated: No | DVARS Dip Associated: No | Cross-Subject Variable: Yes |
| Single Subject: No | % Variance Explained: 1.71 | Globality Index: 1 |
| Task Component: 17 | Taskr Component: No | Sleep Noted: No |
| Rationale: Spatial map includes positive and negative patches that respect known retinotopic visual organization (Foveal vs Peripheral) | | |

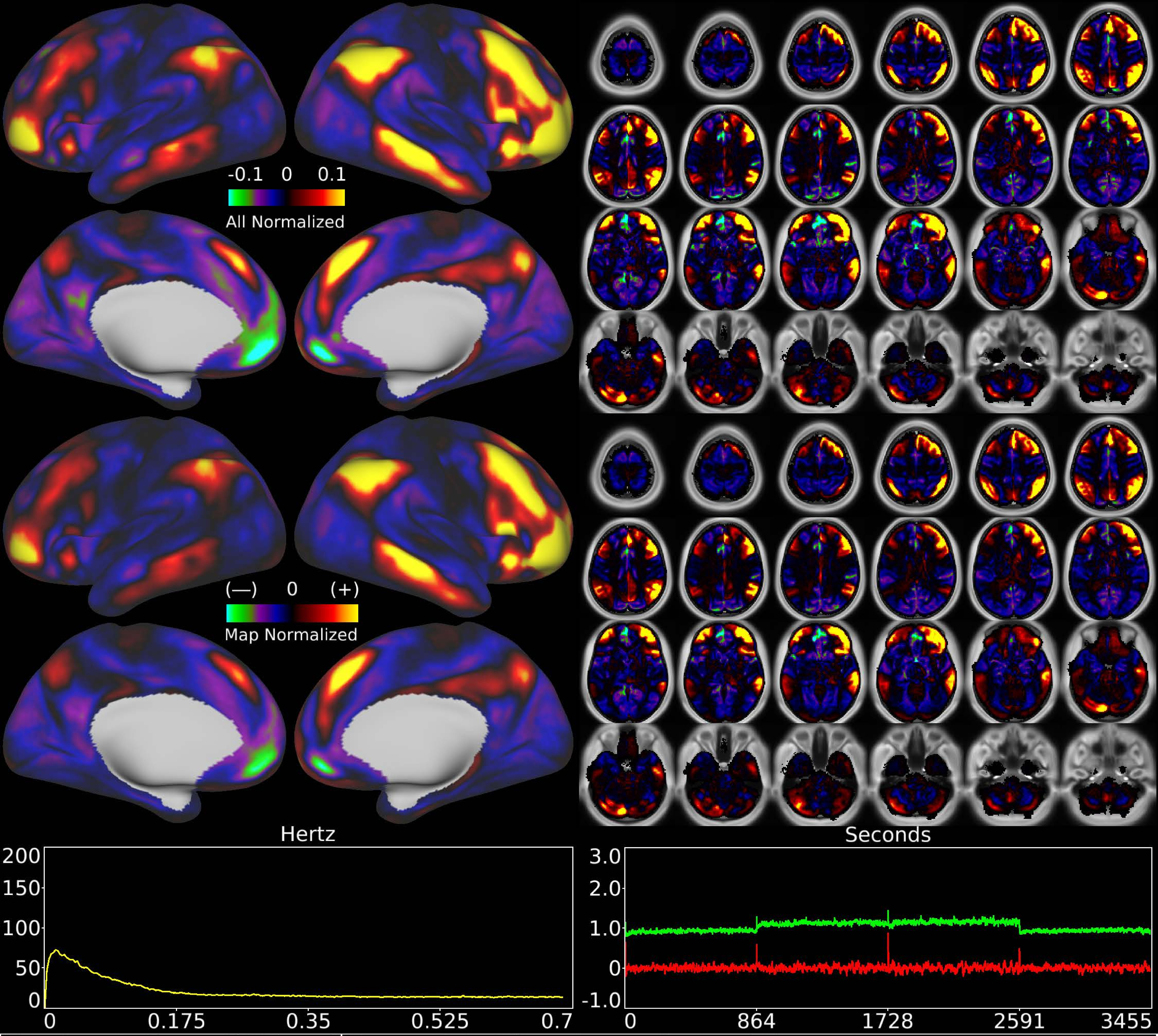


| | | |
|---|---|---|
| Number & Class: 17 Signal | Name: Subsidiary Fronto-Parietal Network | |
| RVT Correlated: No | DVARS Dip Associated: No | Cross-Subject Variable: No |
| Single Subject: No | % Variance Explained: 1.65 | Globality Index: 0.85 |
| Task Component: No | Taskr Component: No | Sleep Noted: No |
| Rationale: Spatial map includes positive and negative patches that respect known RSNs (e.g. Yeo Network 13 | | |

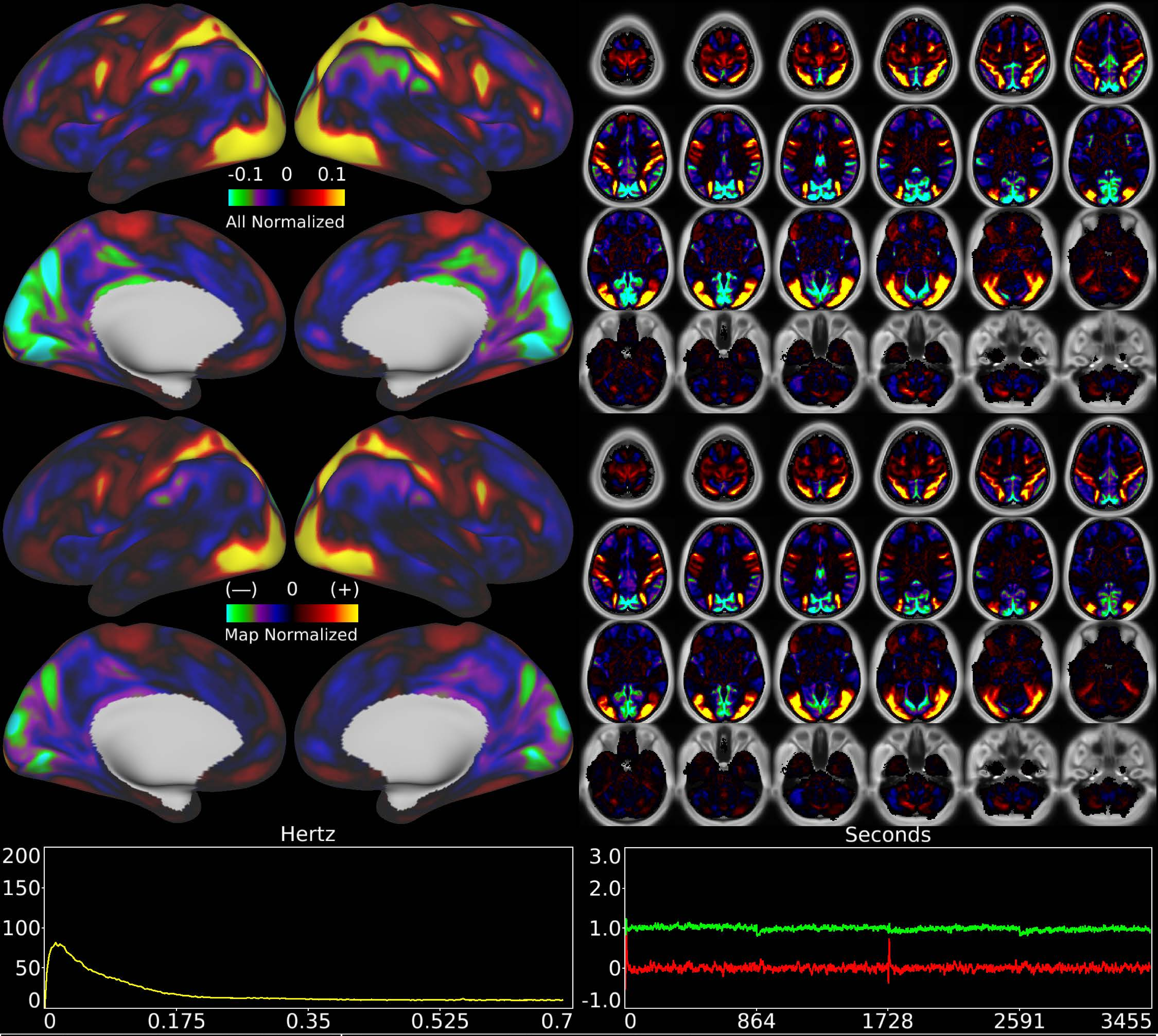


| | | |
|---|---|---|
| Number & Class: 18 Signal | Name: Dorsal Stream Visual | |
| RVT Correlated: No | DVARS Dip Associated: No | Cross-Subject Variable: No |
| Single Subject: No | % Variance Explained: 1.6 | Globality Index: 0.28 |
| Task Component: No | Taskr Component: 16 | Sleep Noted: No |
| Rationale: Spatial map includes positive and negative patches that respect known RSNs (e.g. Dorsal Attention Network) | | |

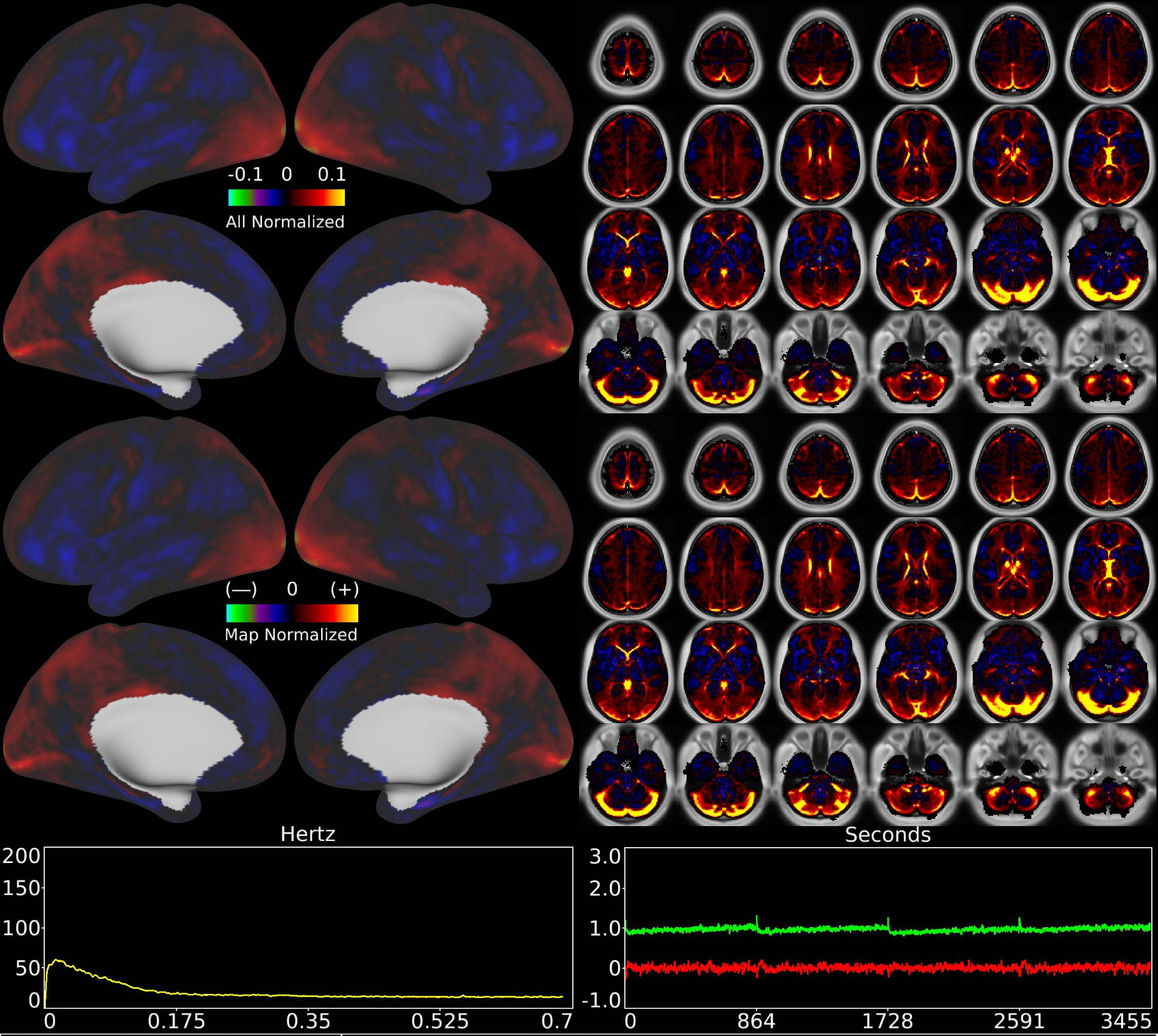


| | | |
|---|---|---|
| Number & Class: 19 Noise | Name: Veins + WM | |
| RVT Correlated: No | DVARS Dip Associated: Yes | Cross-Subject Variable: Yes |
| Single Subject: No | % Variance Explained: 1.46 | Globality Index: 0.34 |
| Task Component: 8 | Taskr Component: 5 | Sleep Noted: No |
| Rationale: Spatial map contains substantial white matter and venous signal | | |

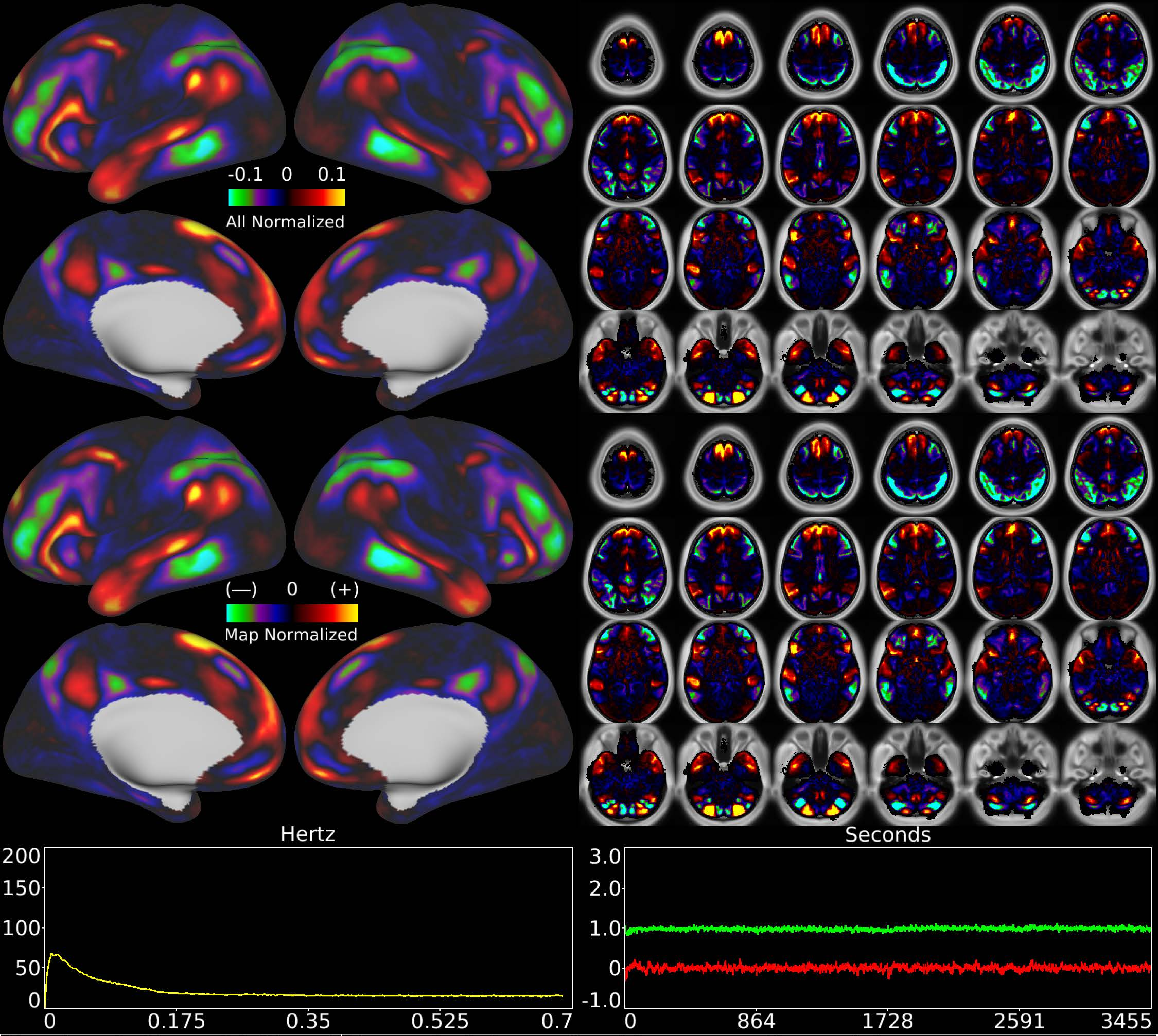


| | | |
|---|---|---|
| Number & Class: 20 Signal | Name: Left Lateralized Language Network I | |
| RVT Correlated: No | DVARS Dip Associated: No | Cross-Subject Variable: No |
| Single Subject: No | % Variance Explained: 1.49 | Globality Index: 1 |
| Task Component: 22+36 | Taskr Component: 30+32 | Sleep Noted: No |
| Rationale: Spatial map includes positive and negative patches that respect known areal boundaries (e.g. 44; 45; 55b; SFL) | | |

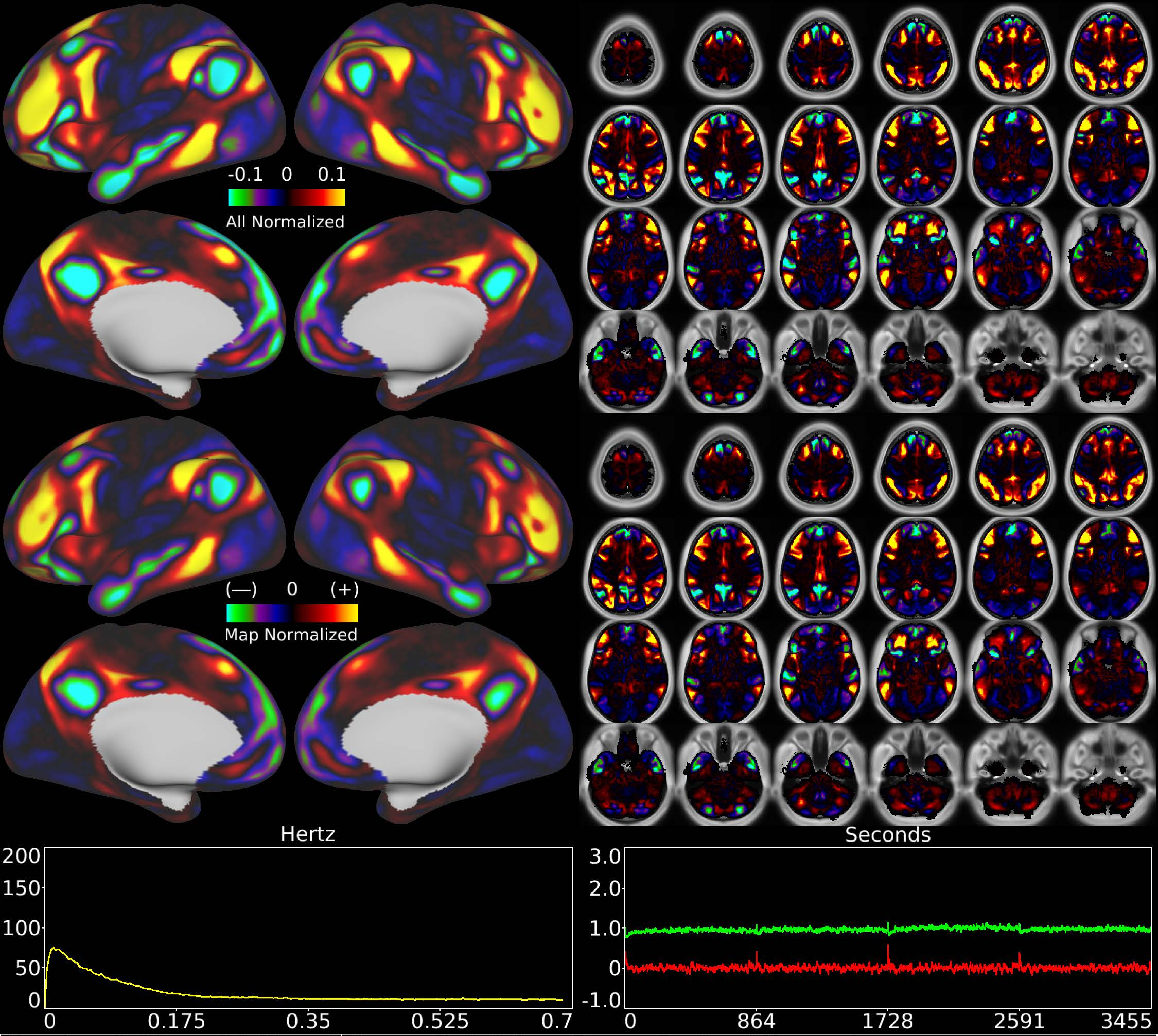


| | | |
|---|---|---|
| Number & Class: 21 Signal | Name: Subsidiary Fronto-Parietal Network | |
| RVT Correlated: No | DVARS Dip Associated: No | Cross-Subject Variable: No |
| Single Subject: No | % Variance Explained: 1.46 | Globality Index: 0.32 |
| Task Component: No | Taskr Component: -13 | Sleep Noted: No |

Rationale: Spatial map includes positive and negative patches that respect known RSNs (e.g. Fronto-Parietal Network)

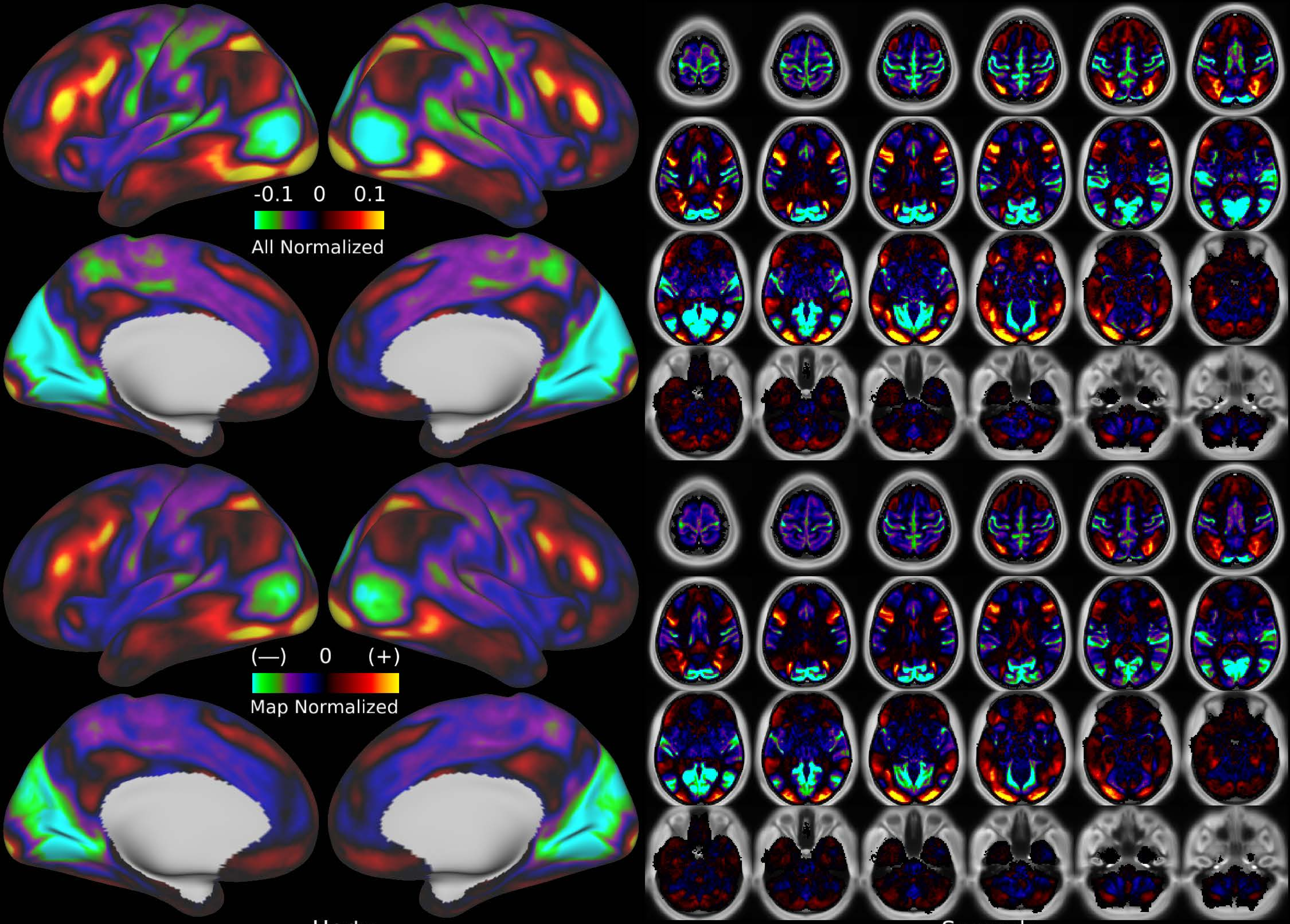


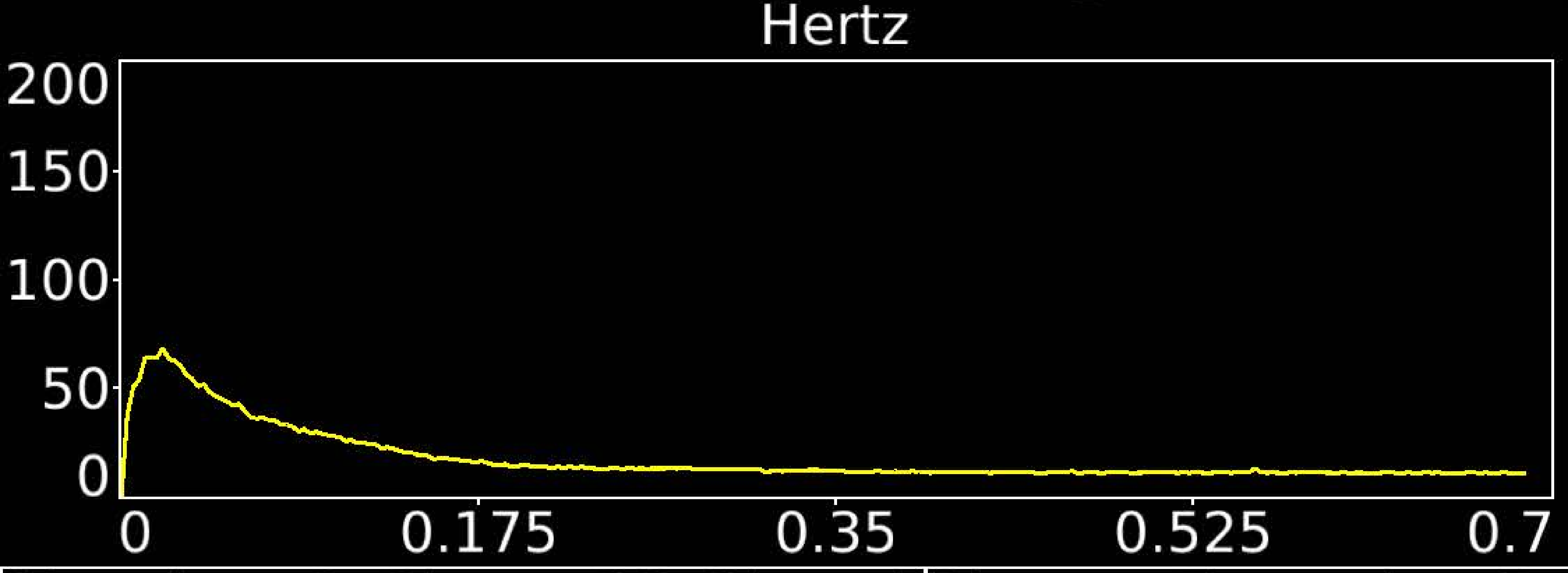


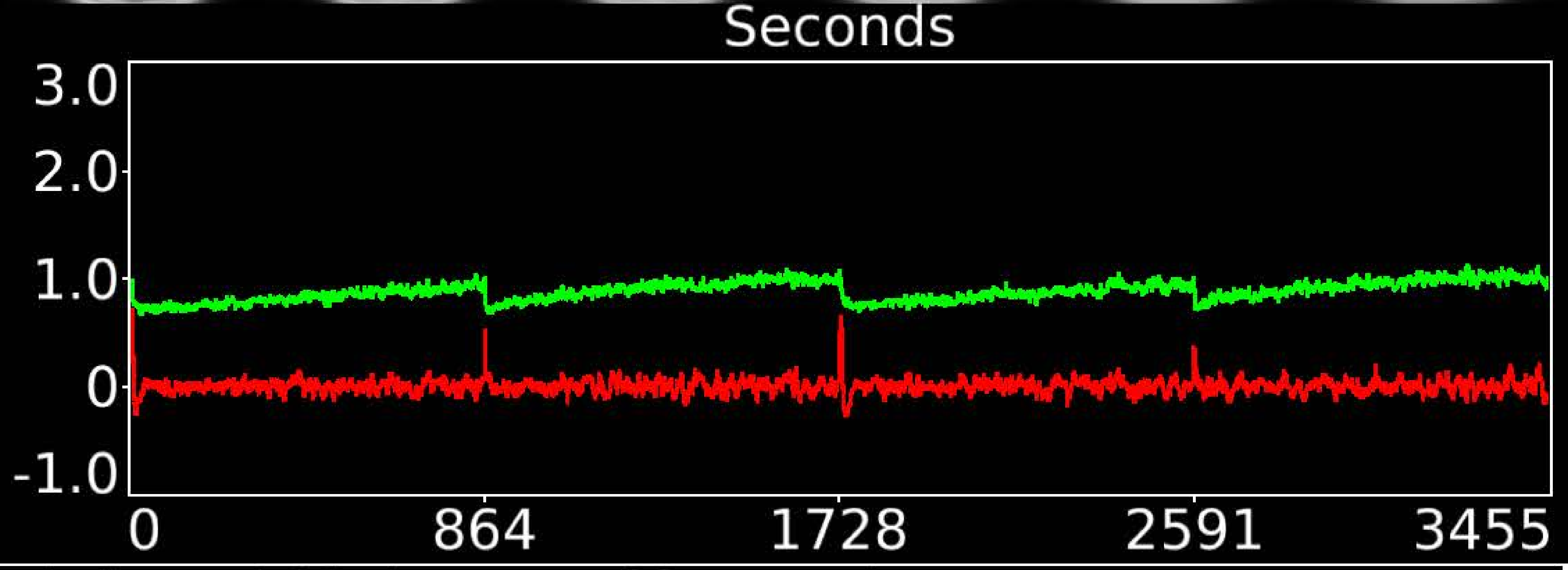


| | | |
|---|---|---|
| Number & Class: 22 Signal | Name: Foveal > Peripheral Unknown Network | |
| RVT Correlated: No | DVARS Dip Associated: No | Cross-Subject Variable: Yes |
| Single Subject: No | % Variance Explained: 1.35 | Globality Index: 0.92 |
| Task Component: No | Taskr Component: No | Sleep Noted: Yes |
| Rationale: Spatial map includes positive and negative patches that respect known retinotopic visual organization (Foveal vs Peripheral) | | |

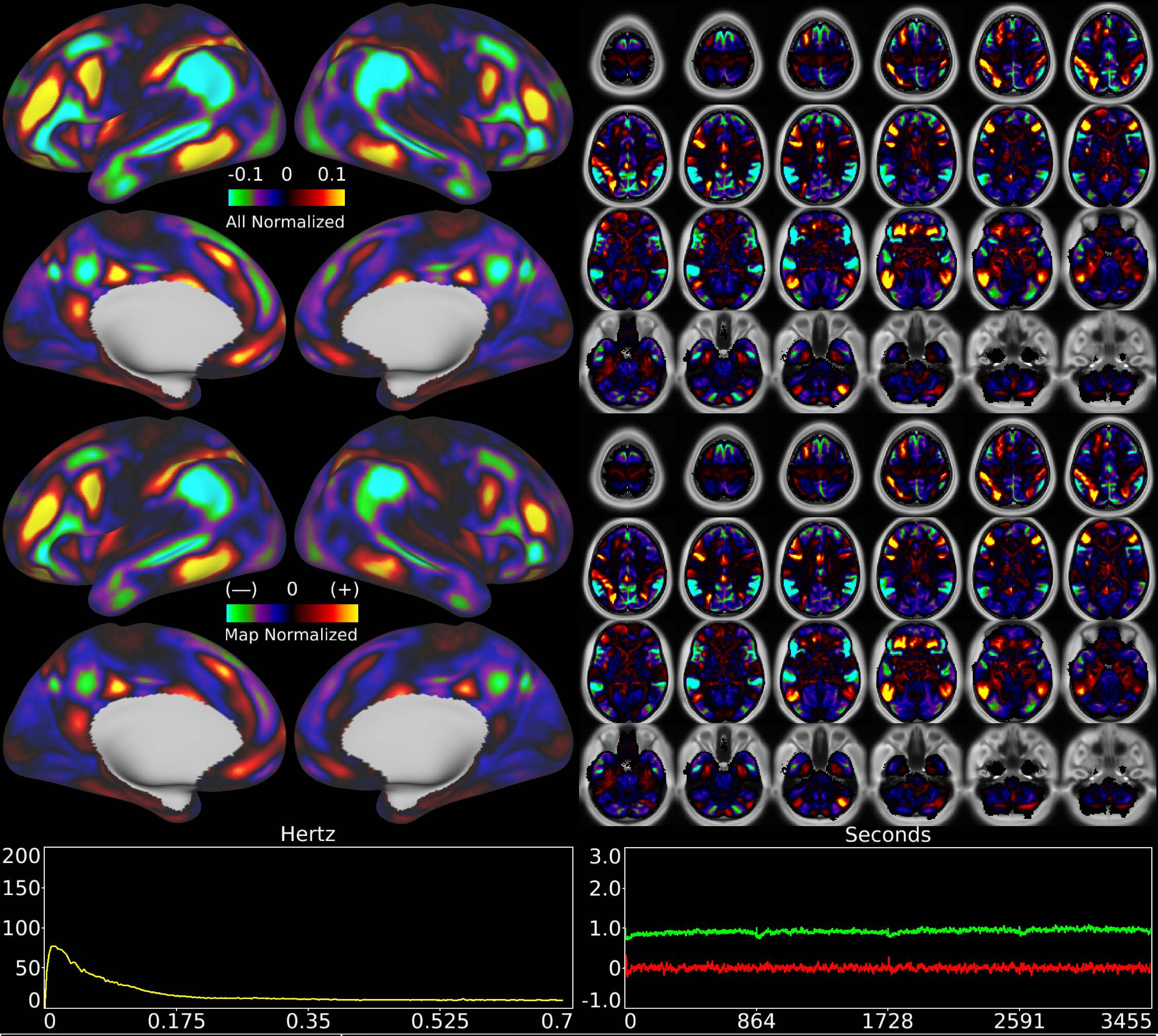


| | | |
|---|---|---|
| Number & Class: 23 Signal | Name: Subsidiary Fronto-Parietal Network | |
| RVT Correlated: No | DVARS Dip Associated: No | Cross-Subject Variable: No |
| Single Subject: No | % Variance Explained: 1.35 | Globality Index: 0.91 |
| Task Component: No | Taskr Component: No | Sleep Noted: No |
| Rationale: Spatial map includes positive and negative patches that respect known RSNs (e.g. Fronto-Parietal Network) | | |

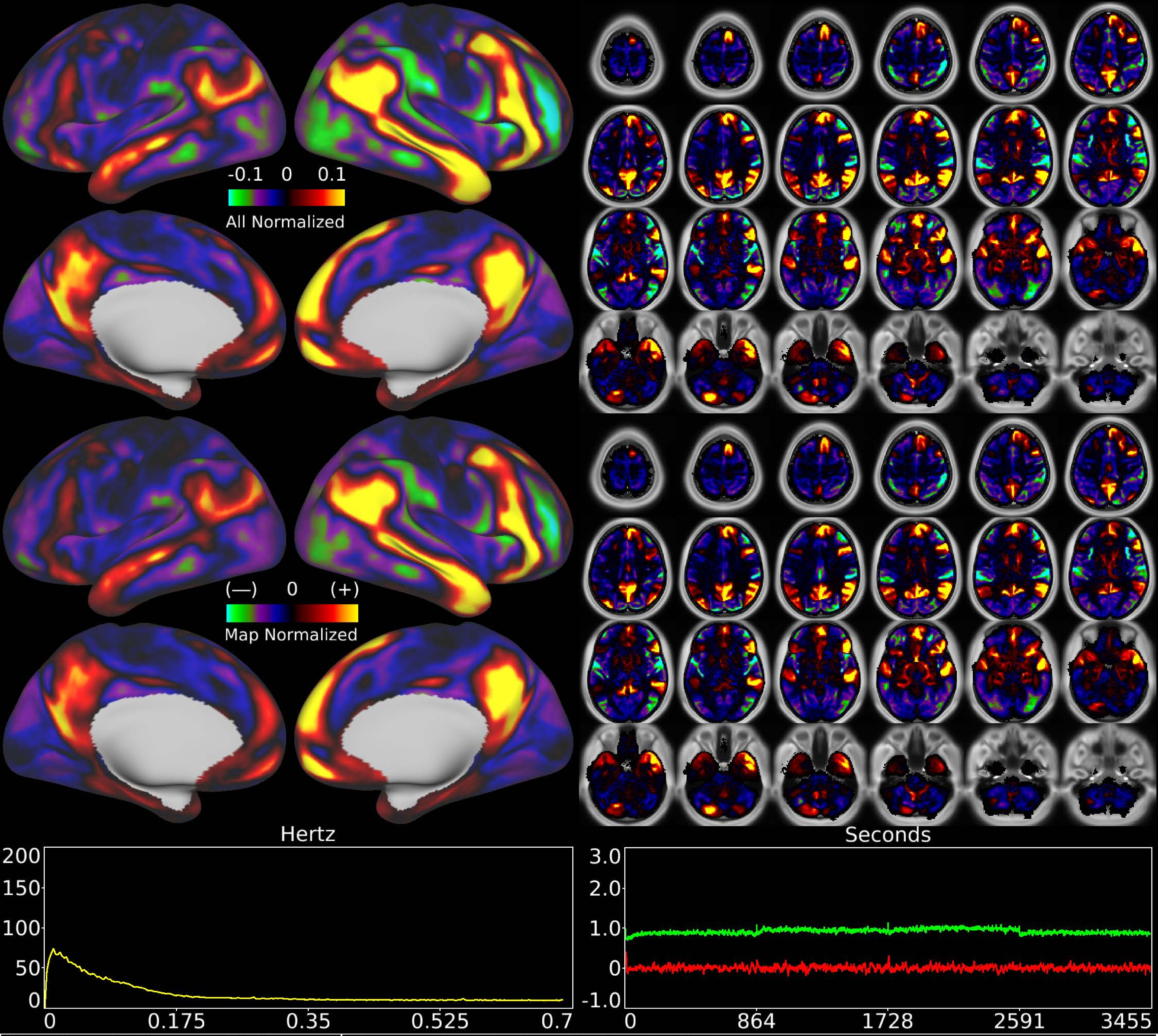


| | | |
|---|---|---|
| Number & Class: 24 Signal | Name: Subsidiary Default Mode | |
| RVT Correlated: No | DVARS Dip Associated: No | Cross-Subject Variable: No |
| Single Subject: No | % Variance Explained: 1.32 | Globality Index: 0.94 |
| Task Component: No | Taskr Component: 48 | Sleep Noted: No |
| Rationale: Spatial map includes positive and negative patches that respect known RSNs (e.g. Default Mode Network) | | |

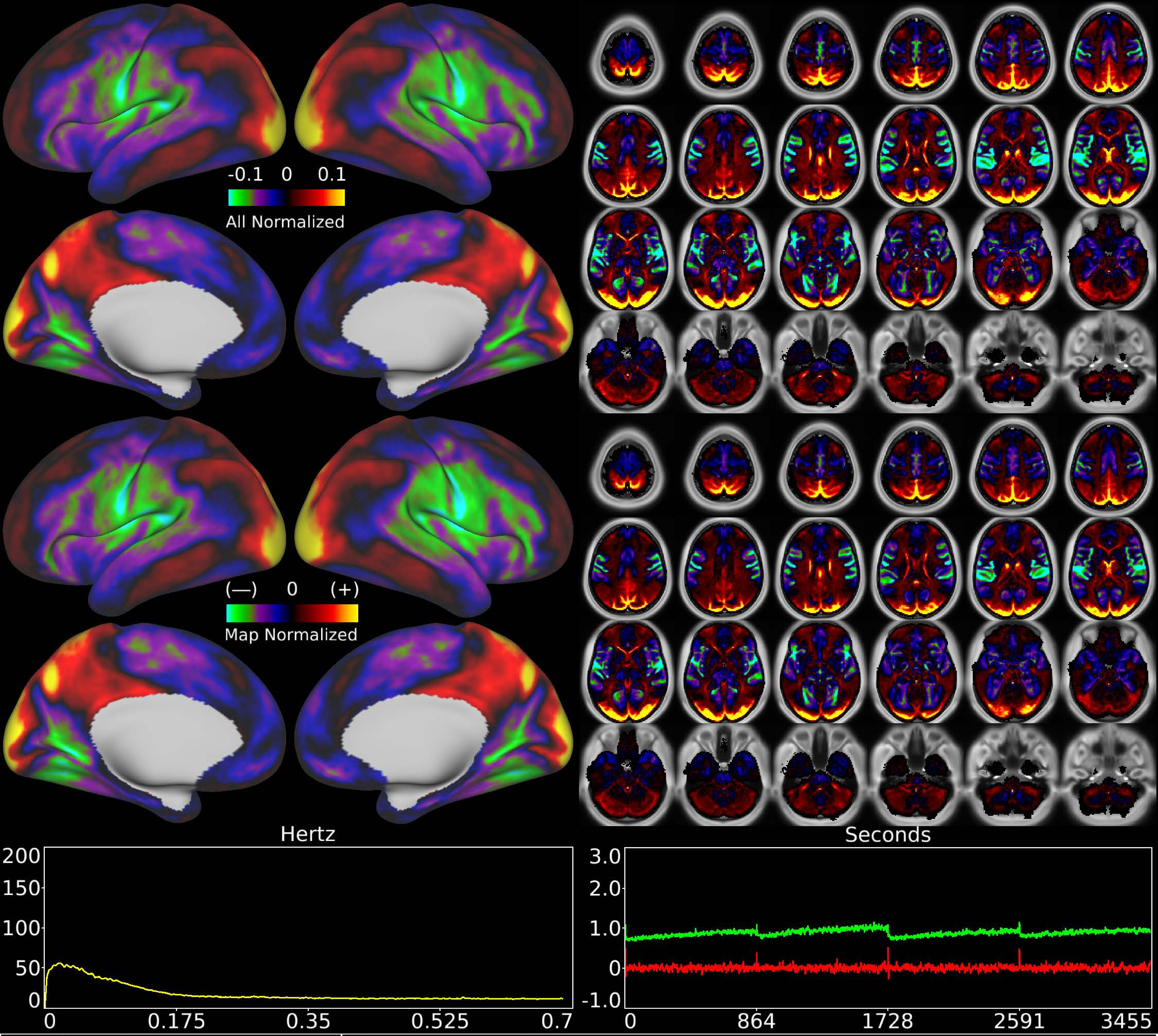


| | | |
|---|---|---|
| Number & Class: 25 Noise | Name: WM + Odd Cortical Maps | |
| RVT Correlated: No | DVARS Dip Associated: No | Cross-Subject Variable: Yes |
| Single Subject: No | % Variance Explained: 1.27 | Globality Index: 0.12 |
| Task Component: No | Taskr Component: 31 | Sleep Noted: No |
| Rationale: Spatial map contains substantial white matter signal; may also reflect vascular distribution | | |

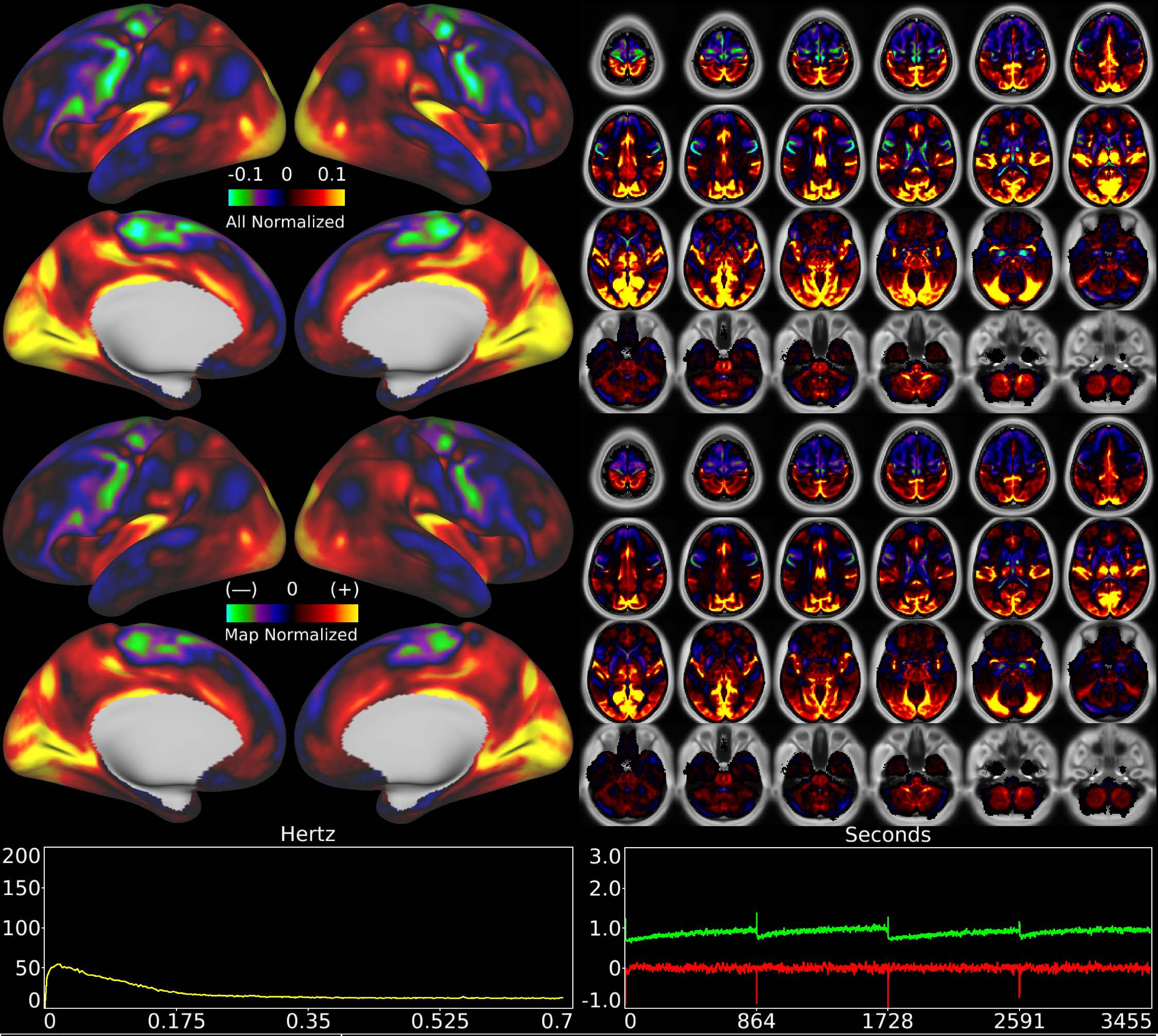


| | | |
|---|---|---|
| Number & Class: 26 Signal | Name: Sensory > Motor | |
| RVT Correlated: No | DVARS Dip Associated: No | Cross-Subject Variable: Yes |
| Single Subject: No | % Variance Explained: 1.29 | Globality Index: 1.17 |
| Task Component: No | Taskr Component: No | Sleep Noted: No |
| Rationale: Spatial map includes positive and negative patches that respect known areal boundaries (e.g. PSL; 45; 55b; FEF) | | |

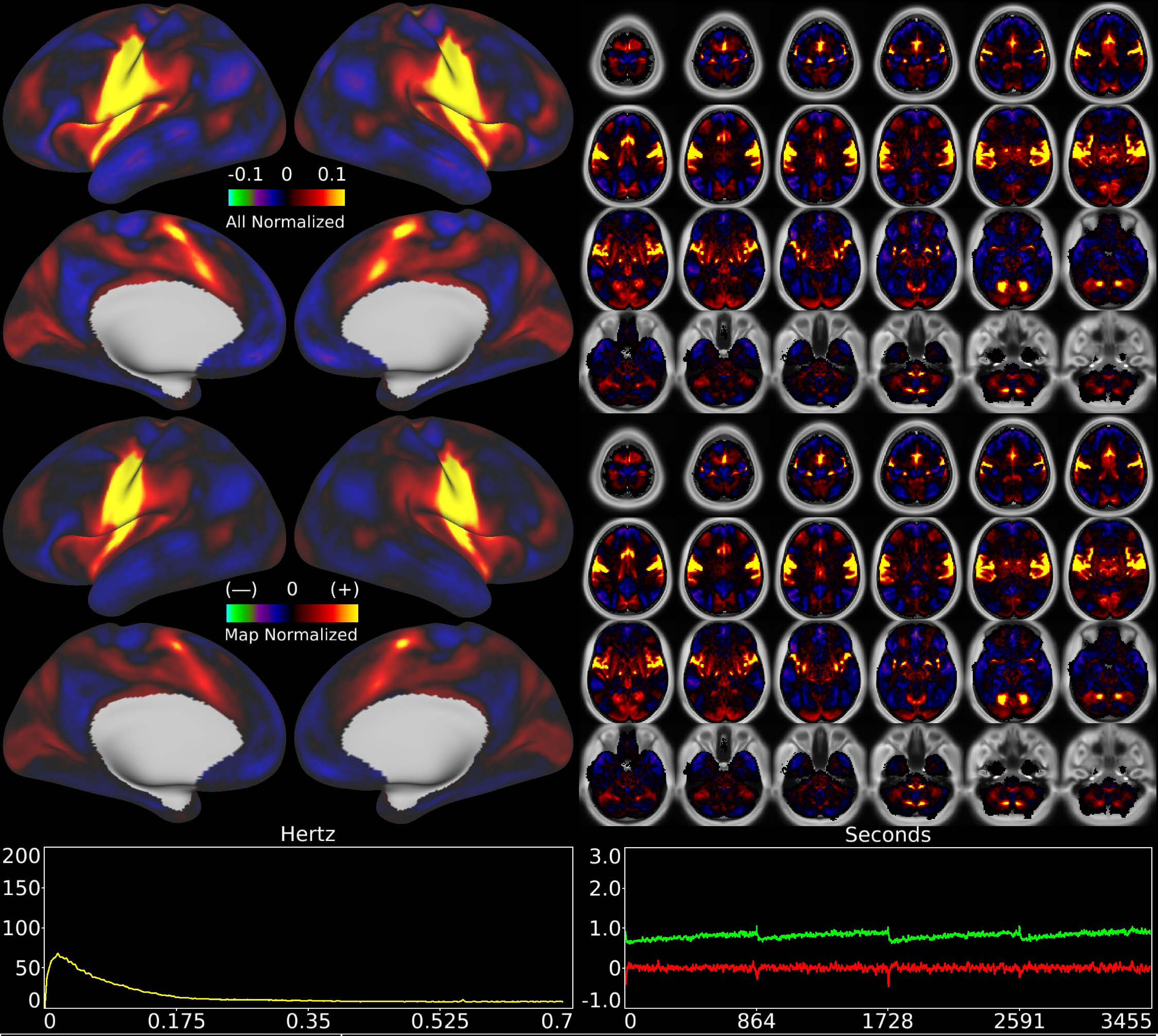


| | | |
|---|---|---|
| Number & Class: 27 Signal | Name: Head Motor Network | |
| RVT Correlated: No | DVARS Dip Associated: Yes | Cross-Subject Variable: Yes |
| Single Subject: No | % Variance Explained: 1.23 | Globality Index: 0.45 |
| Task Component: 16 | Taskr Component: 18 | Sleep Noted: No |
| Rationale: Spatial map includes positive and negative patches that respect known somatotopic sensori-motor organization (Face) | | |

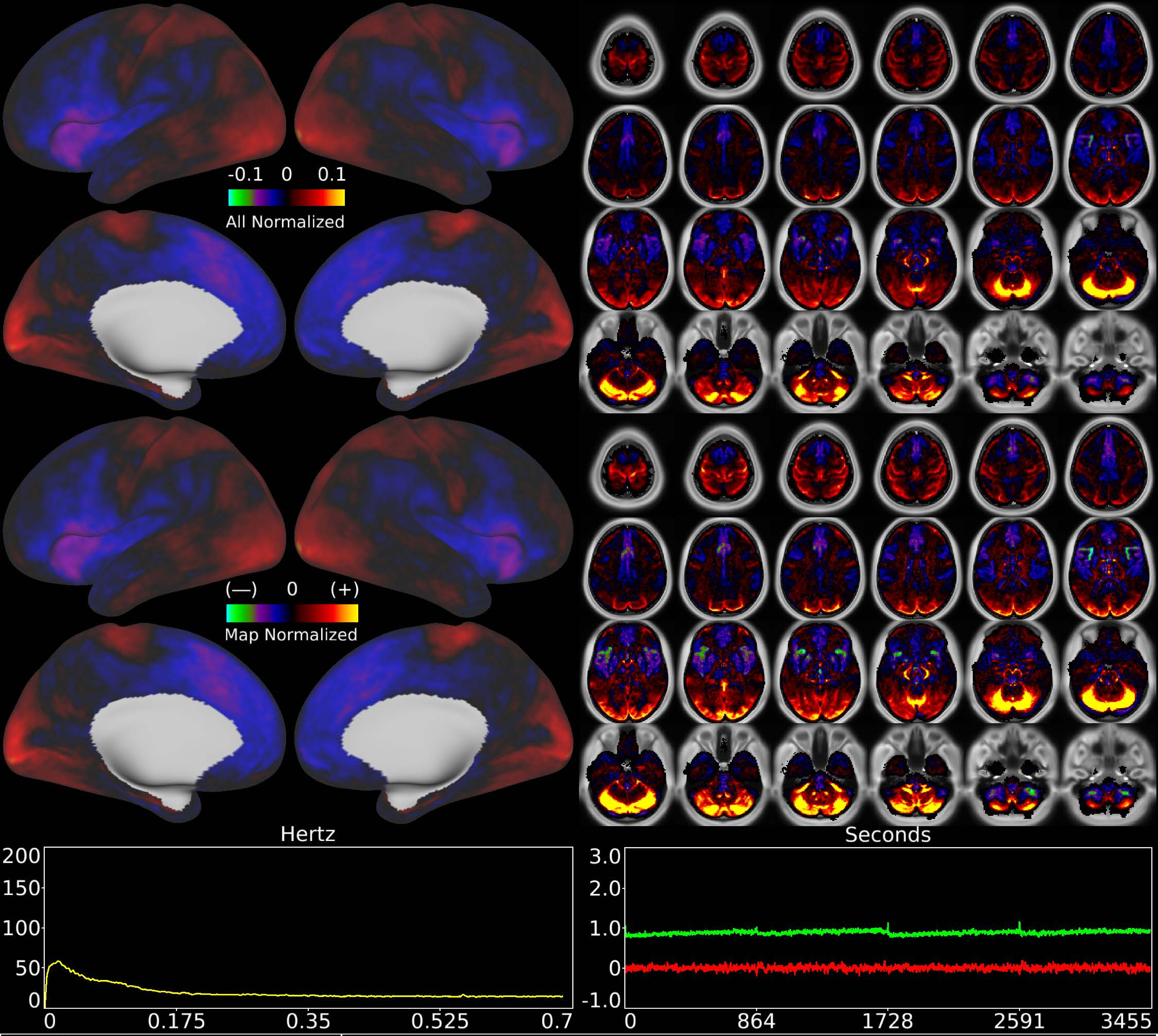


| | | |
|---|---|---|
| Number & Class: 28 Noise | Name: Veins + WM + Cerebellum | |
| RVT Correlated: No | DVARS Dip Associated: Yes | Cross-Subject Variable: No |
| Single Subject: No | % Variance Explained: 1.15 | Globality Index: 0.21 |
| Task Component: 8 | Taskr Component: 5 | Sleep Noted: No |
| Rationale: Spatial map contains substantial white matter and venous signal; positive and negative patches do not reflect known areal boundaries | | |

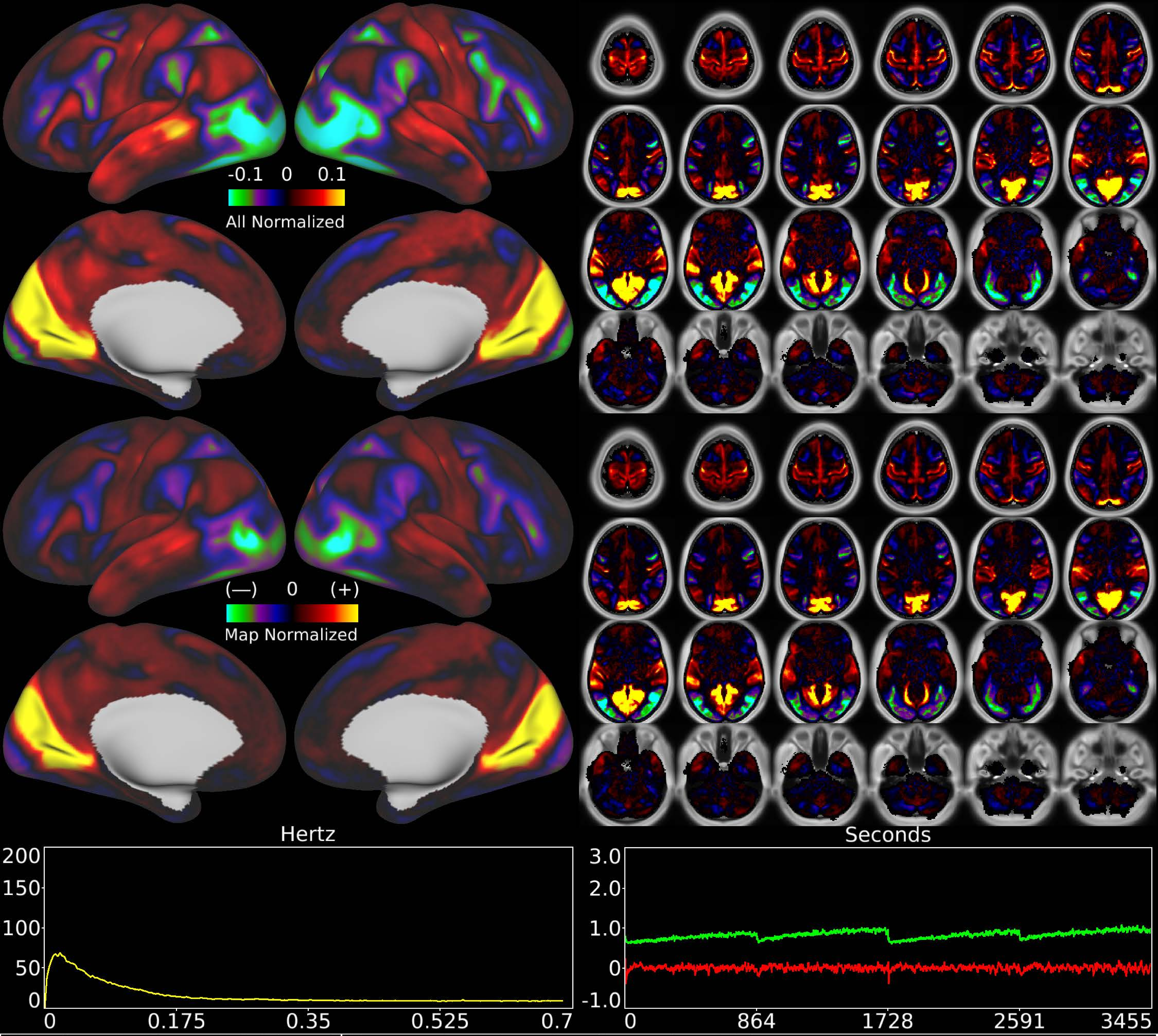


| | | |
|---|---|---|
| Number & Class: 29 Signal | Name: Peripheral > Foveal + MT+ | |
| RVT Correlated: No | DVARS Dip Associated: No | Cross-Subject Variable: Yes |
| Single Subject: No | % Variance Explained: 1.21 | Globality Index: 0.5 |
| Task Component: No | Taskr Component: No | Sleep Noted: Yes |

Rationale: Spatial map includes positive and negative patches that respect known retinotopic visual organization (Peripheral vs Foveal)

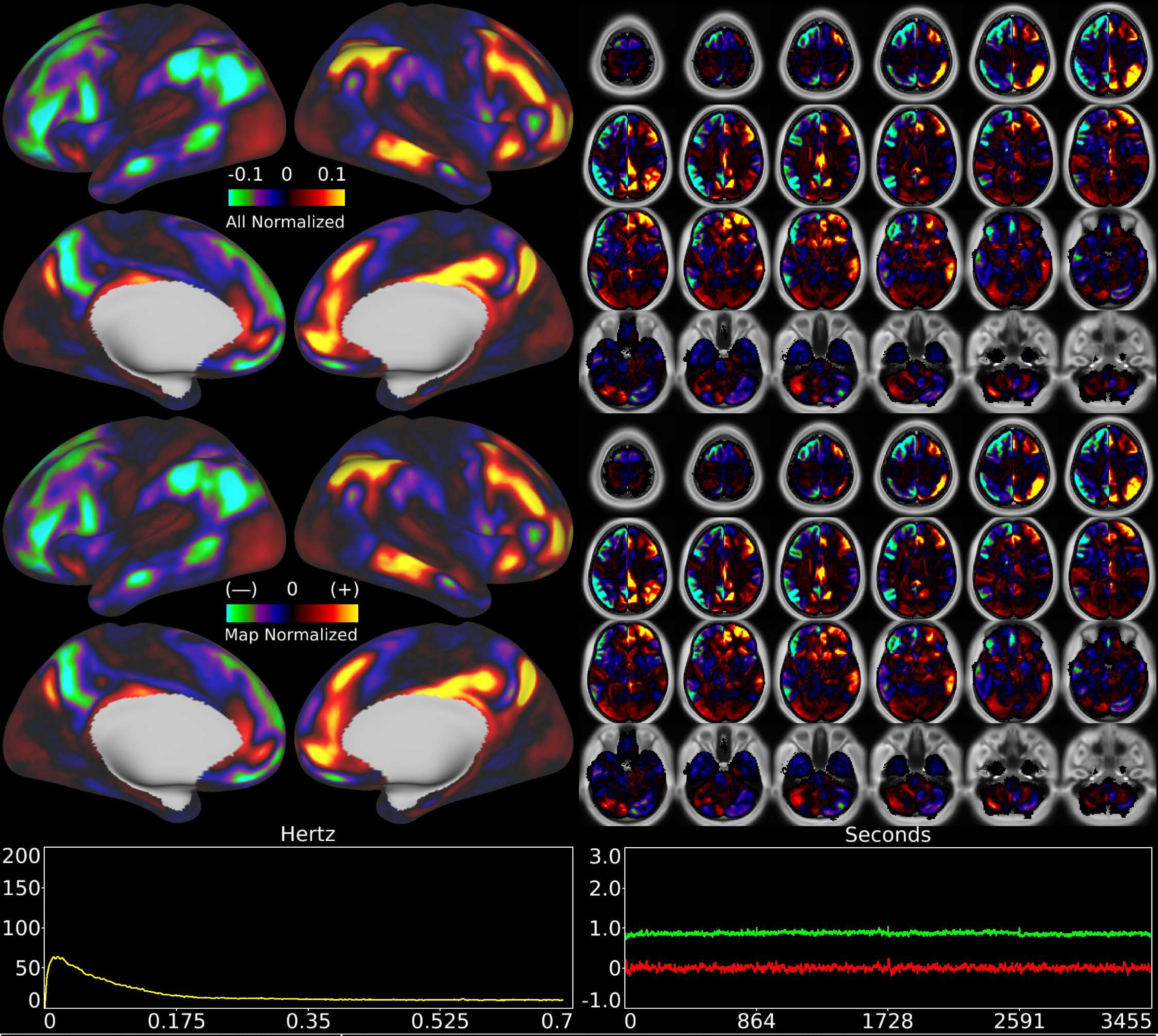


| | | |
|---|---|---|
| Number & Class: 30 Signal | Name: R Fronto-Parietal > L Default Mode | |
| RVT Correlated: No | DVARS Dip Associated: No | Cross-Subject Variable: No |
| Single Subject: No | % Variance Explained: 1.16 | Globality Index: 0.16 |
| Task Component: No | Taskr Component: No | Sleep Noted: No |
| Rationale: Spatial map includes positive and negative patches that respect known areal boundaries (e.g. POS2 and RSC) | | |

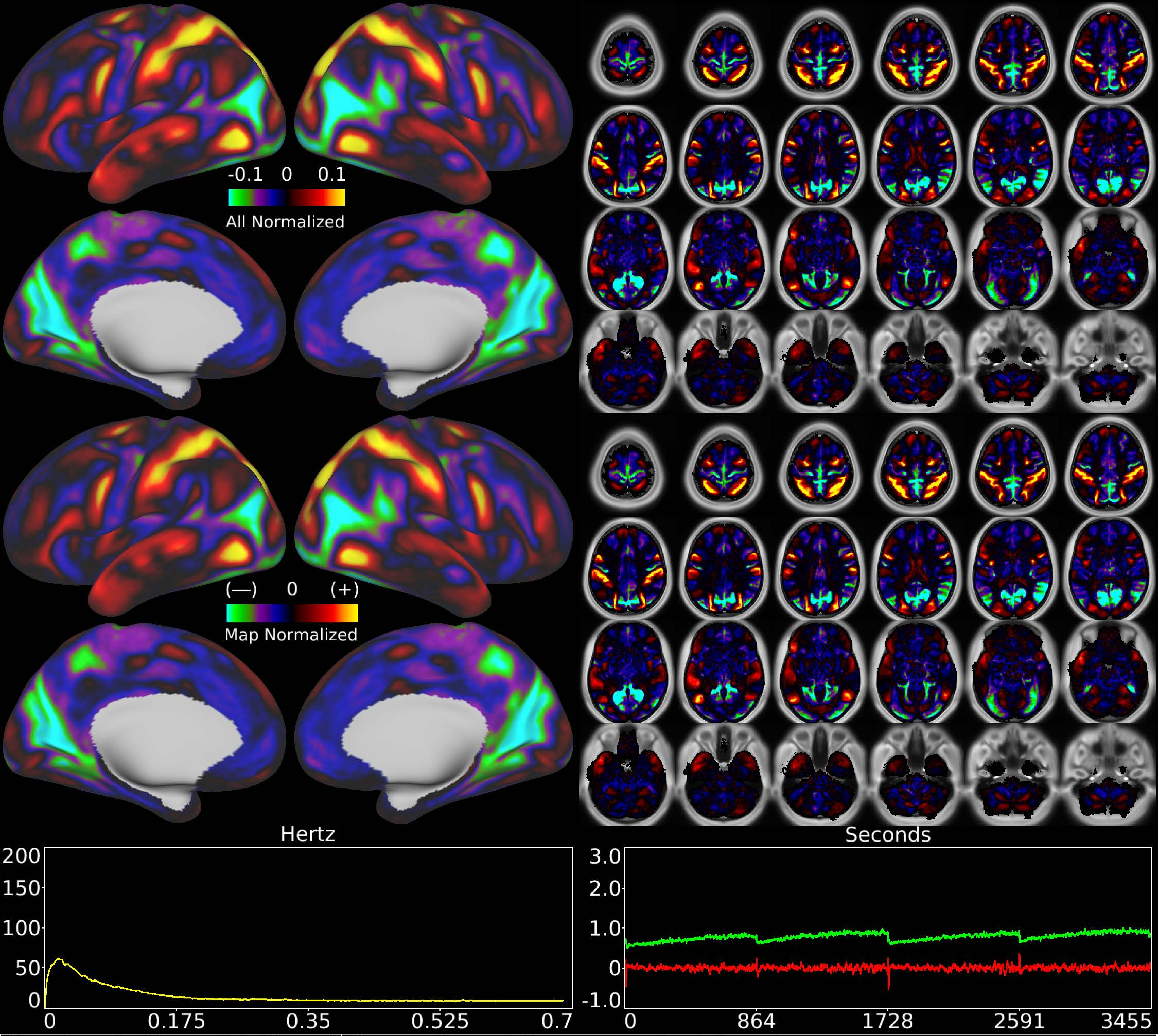


| | | |
|---|---|---|
| Number & Class: 31 Signal | Name: Dorsal Stream vs Peripheral | |
| RVT Correlated: No | DVARS Dip Associated: No | Cross-Subject Variable: Yes |
| Single Subject: No | % Variance Explained: 1.1 | Globality Index: 0.87 |
| Task Component: No | Taskr Component: No | Sleep Noted: Yes |
| Rationale: Spatial map includes positive and negative patches that respect known RSNs (e.g. Dorsal Attention Network) | | |

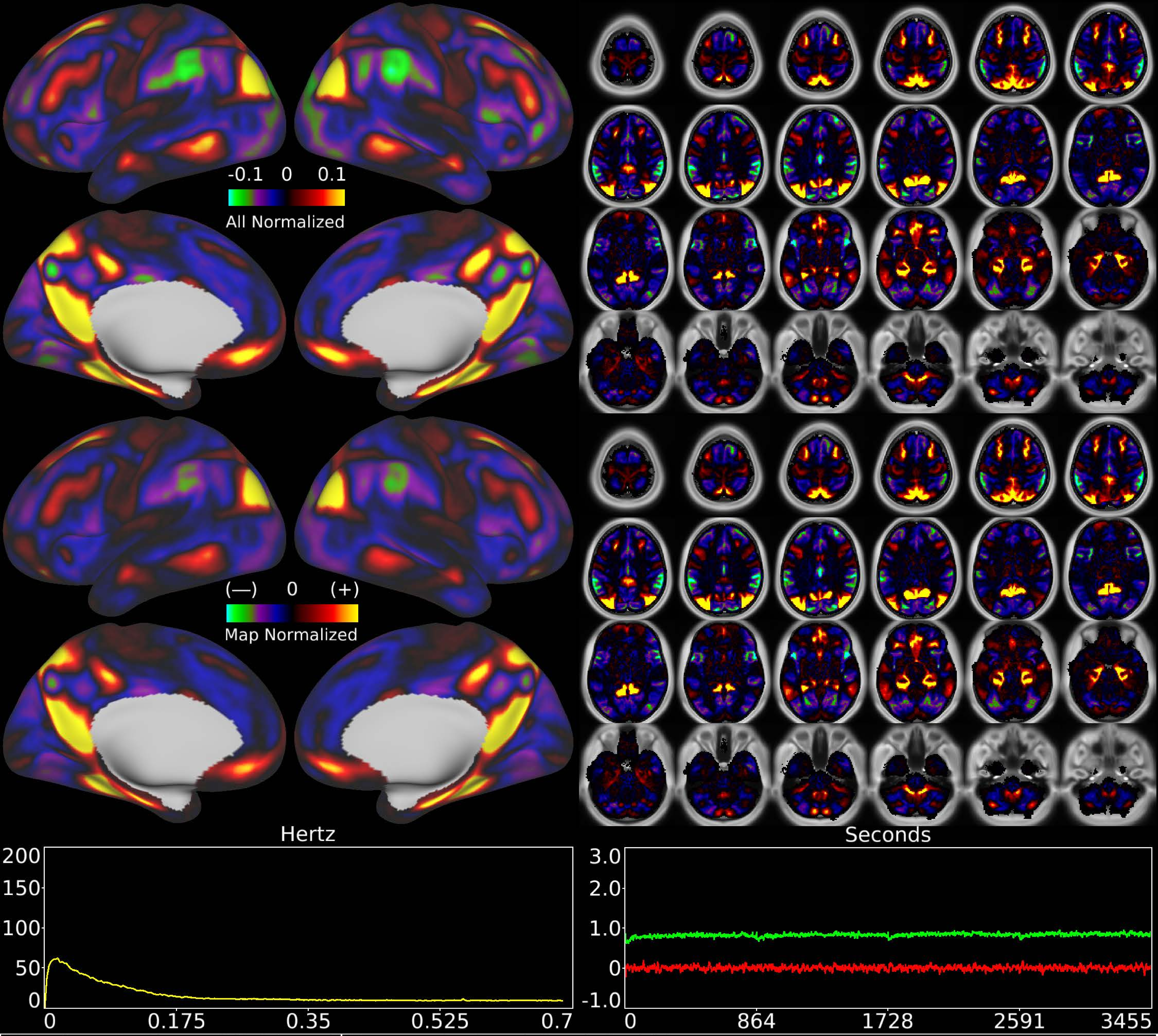


| | | |
|---|---|---|
| Number & Class: 32 Signal | Name: Parietal Occipital Network | |
| RVT Correlated: No | DVARS Dip Associated: No | Cross-Subject Variable: No |
| Single Subject: No | % Variance Explained: 1.1 | Globality Index: 0.55 |
| Task Component: No | Taskr Component: 39 | Sleep Noted: No |
| Rationale: Spatial map includes positive and negative patches that respect known RSNs (e.g. Parietal Occipital Network) | | |

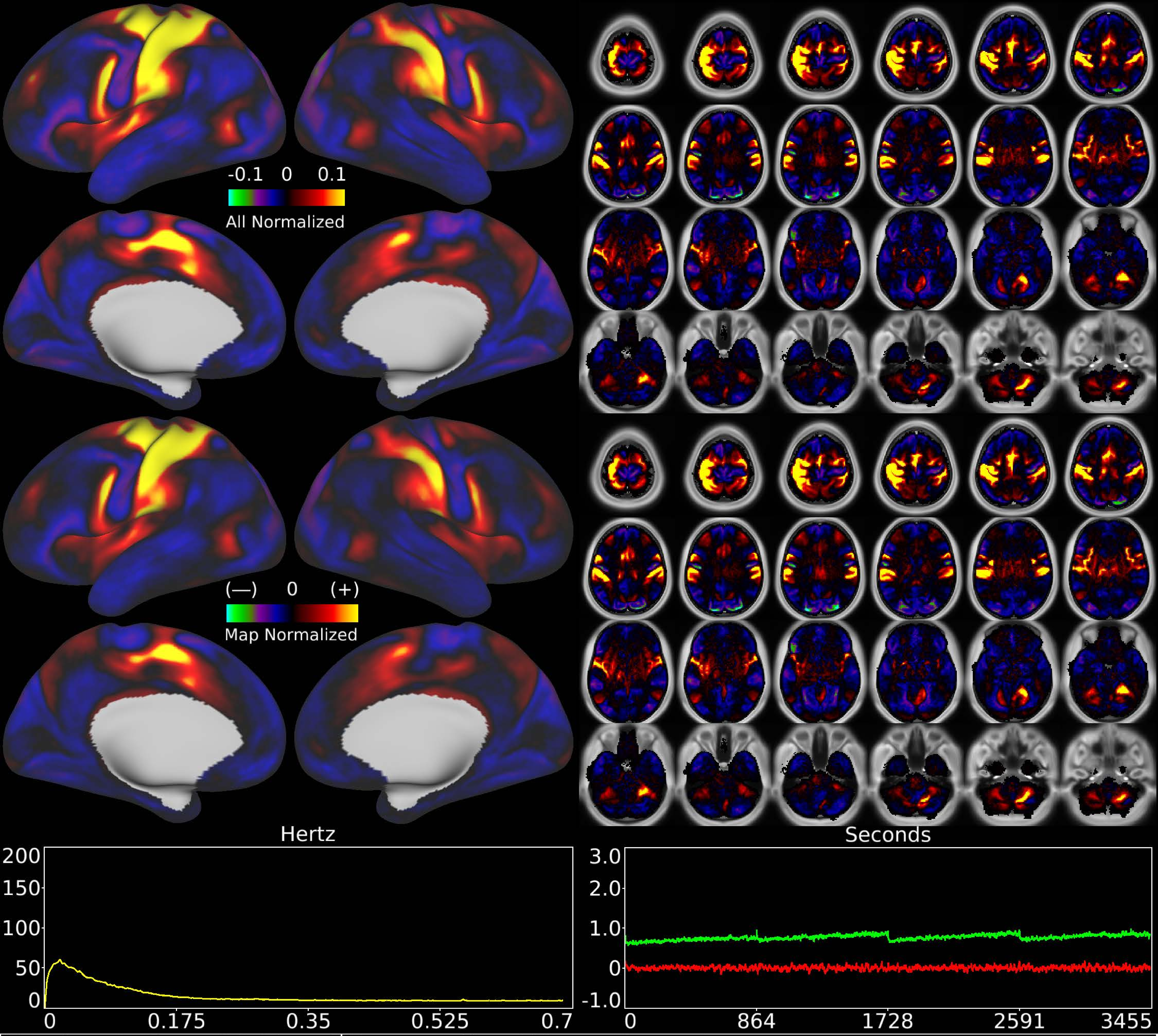


| | | |
|---|---|---|
| Number & Class: 33 Signal | Name: R Hand Motor Network | |
| RVT Correlated: No | DVARS Dip Associated: Yes | Cross-Subject Variable: Yes |
| Single Subject: No | % Variance Explained: 1.07 | Globality Index: 0.25 |
| Task Component: 6 | Taskr Component: 7 | Sleep Noted: No |

Rationale: Spatial map includes positive and negative patches that respect known somatotopic sensori-motor organization (Right Hand)

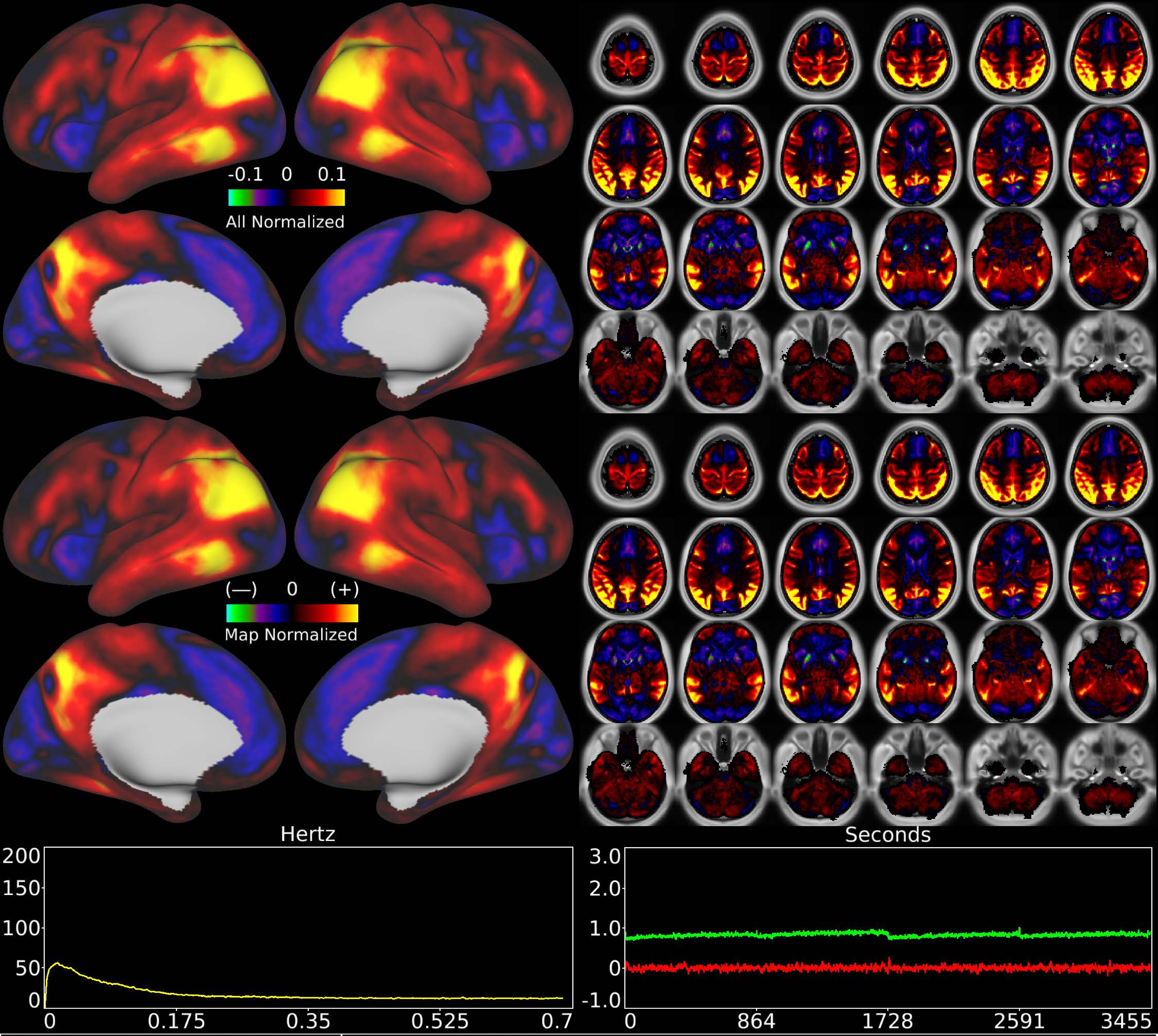


| | | |
|---|---|---|
| Number & Class: 34 Noise | Name: DVARS Assoc | |
| RVT Correlated: No | DVARS Dip Associated: Yes | Cross-Subject Variable: No |
| Single Subject: No | % Variance Explained: 1.03 | Globality Index: 1.64 |
| Task Component: No | Taskr Component: No | Sleep Noted: No |
| Rationale: Controversial: Spatial map is challenging to interpret so component was primarily classified on the basis of being associated with DVARS dips | | |

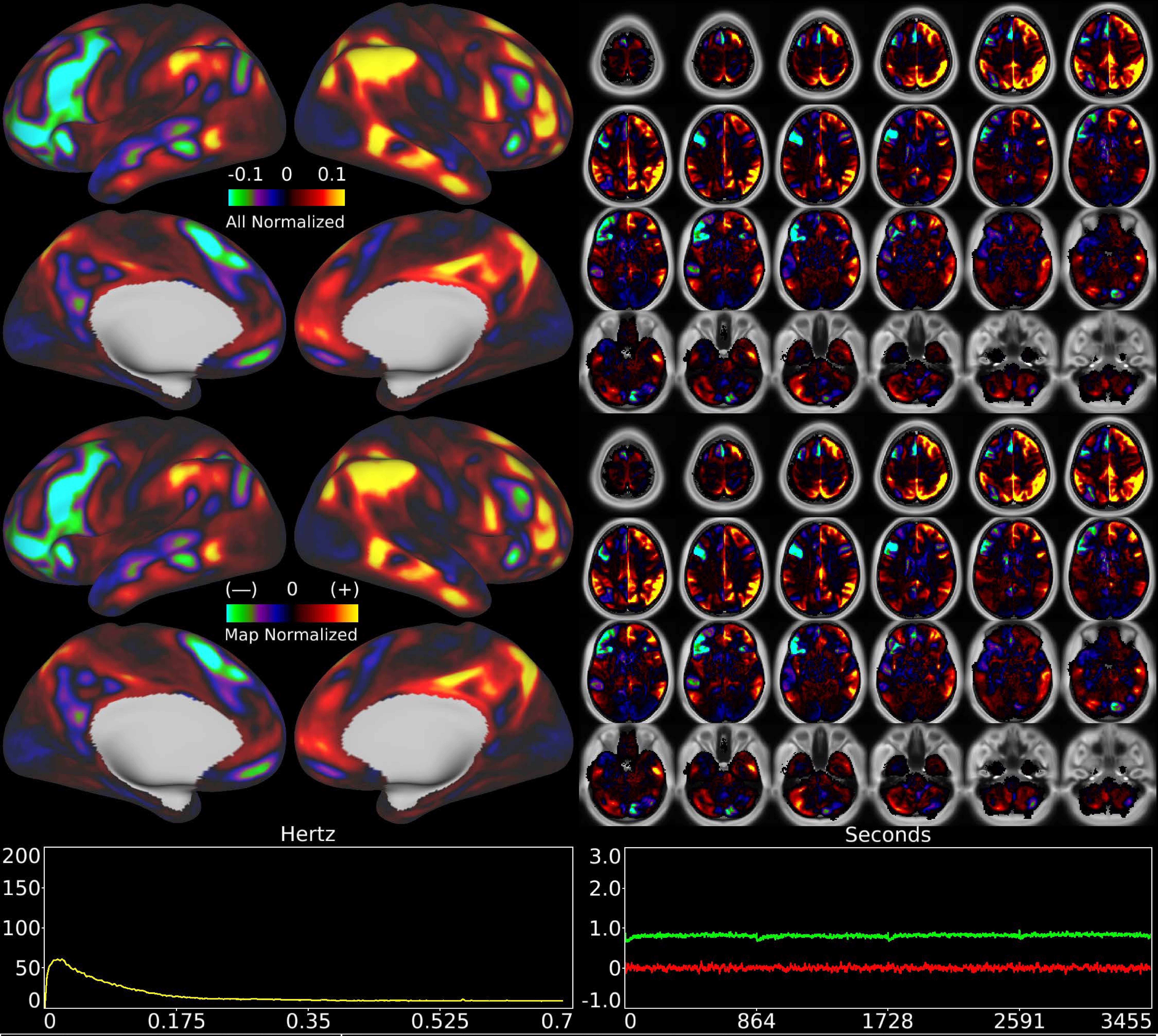


| | | |
|---|---|---|
| Number & Class: 35 Signal | Name: R Unknown Network > L Unknown Network | |
| RVT Correlated: No | DVARS Dip Associated: No | Cross-Subject Variable: No |
| Single Subject: No | % Variance Explained: 1.03 | Globality Index: 0.89 |
| Task Component: No | Taskr Component: No | Sleep Noted: No |
| Rationale: Spatial map includes positive and negative patches that respect known RSNs (e.g. Fronto-Parietal Network) | | |

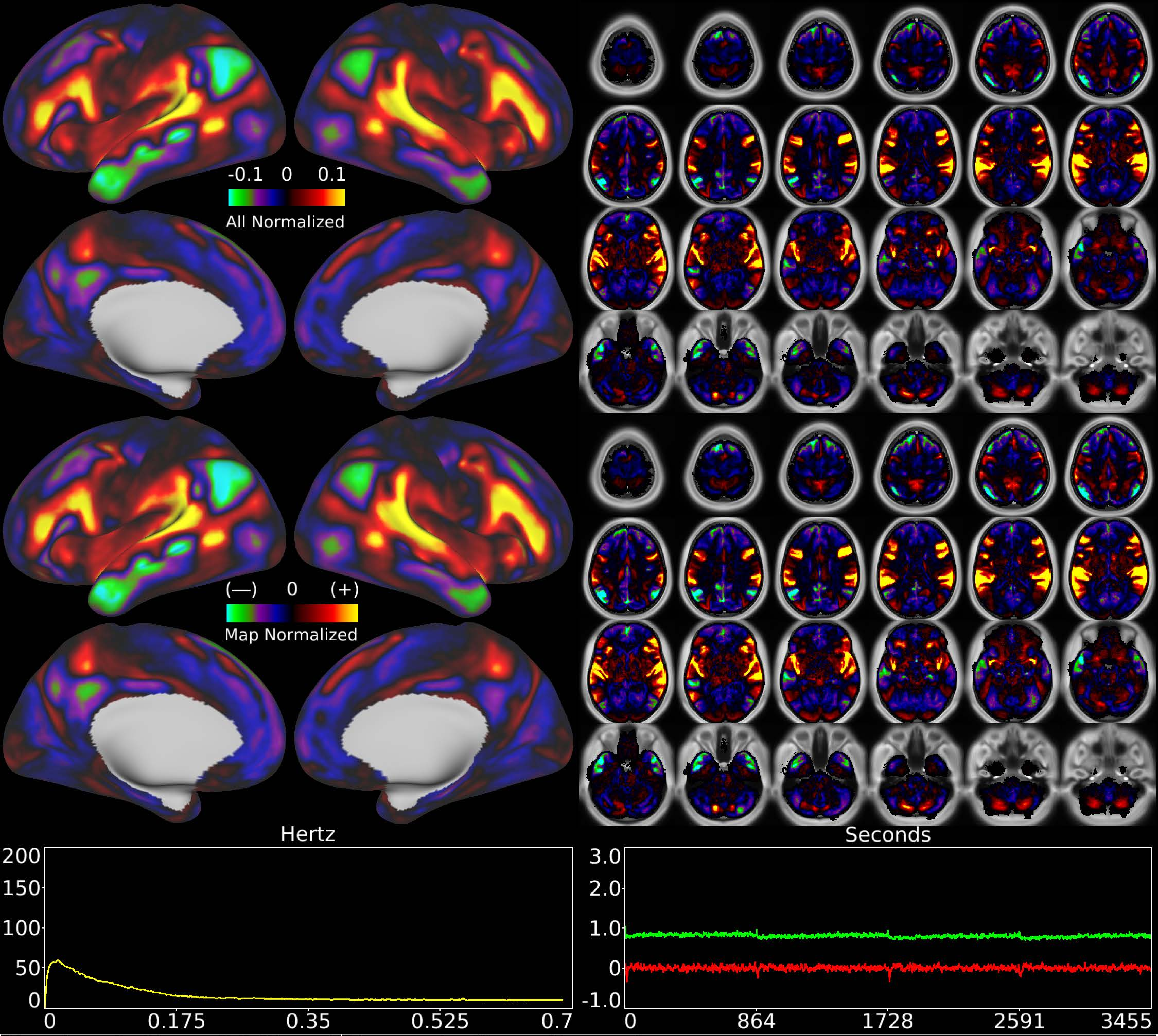


| | | |
|---|---|---|
| Number & Class: 36 Signal | Name: Auditory Network | |
| RVT Correlated: No | DVARS Dip Associated: No | Cross-Subject Variable: No |
| Single Subject: No | % Variance Explained: 1.01 | Globality Index: 0.08 |
| Task Component: No | Taskr Component: No | Sleep Noted: No |
| Rationale: Spatial map includes positive and negative patches that respect known RSNs (e.g. Auditory and Default Mode Network) | | |

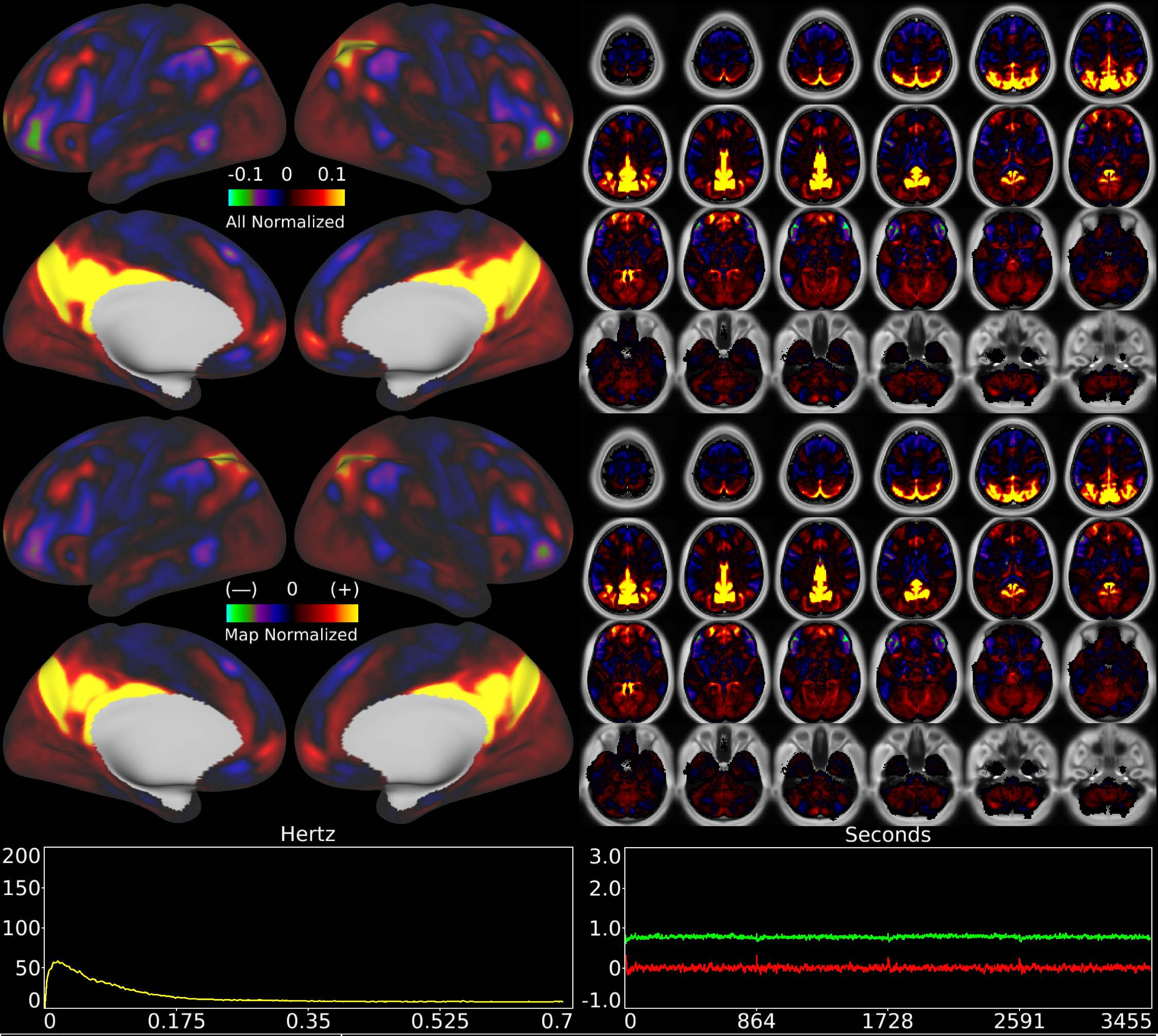


| | | |
|---|---|---|
| Number & Class: 37 Signal | Name: POS2 + RSC Network | |
| RVT Correlated: No | DVARS Dip Associated: No | Cross-Subject Variable: No |
| Single Subject: No | % Variance Explained: 0.94 | Globality Index: 0.93 |
| Task Component: 13 | Taskr Component: 21 | Sleep Noted: No |

Rationale: Spatial map includes positive and negative patches that respect known RSNs (e.g. Medial Parietal Network)

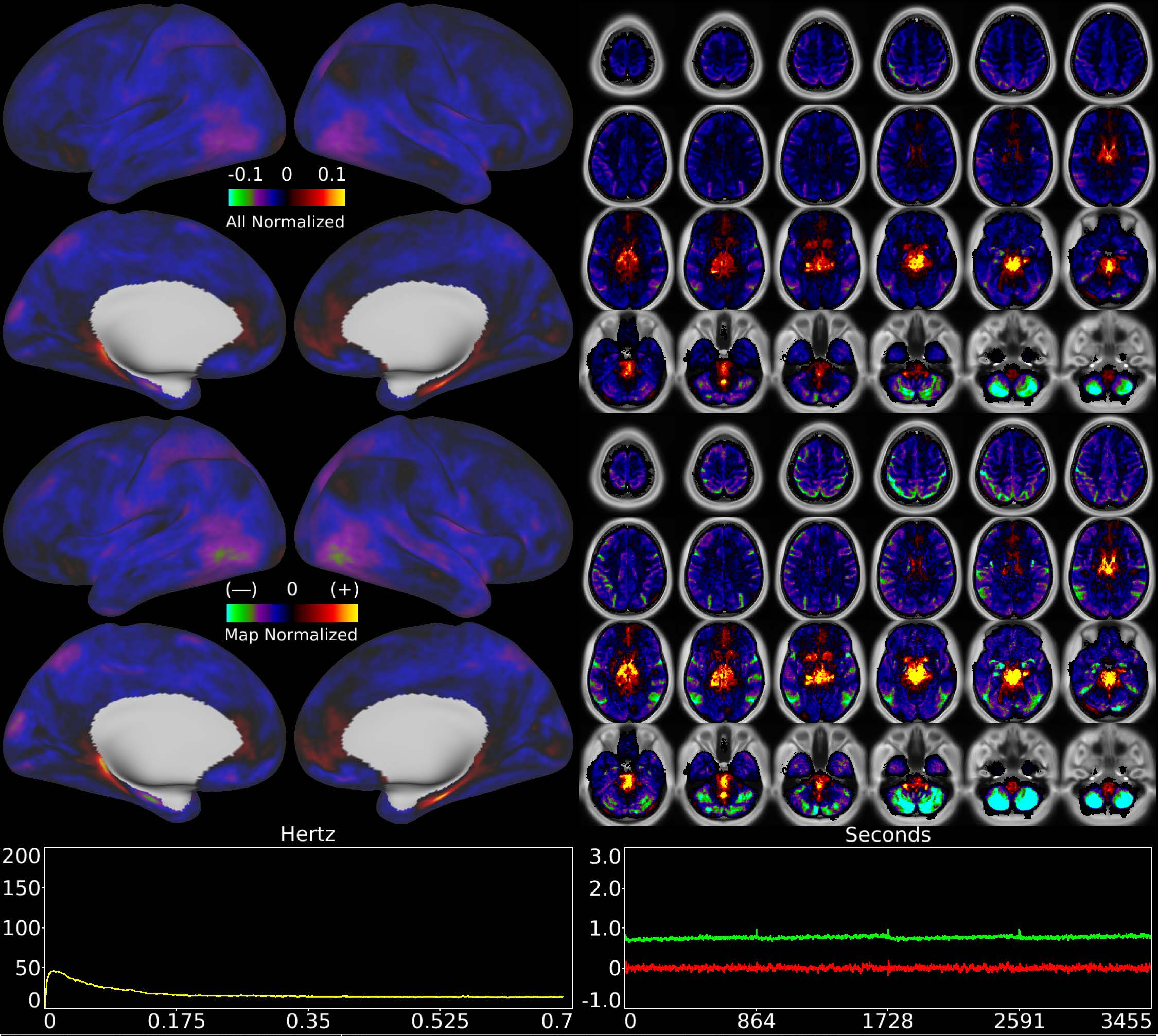


| | | |
|---|---|---|
| Number & Class: 38 Noise | Name: Pons > Inferior Cerebellum DVAR Assoc | |
| RVT Correlated: No | DVARS Dip Associated: Yes | Cross-Subject Variable: No |
| Single Subject: No | % Variance Explained: 0.89 | Globality Index: 2.15 |
| Task Component: No | Taskr Component: No | Sleep Noted: No |
| Rationale: Spatial map not reflective of known areas or RSNs; associated with DVARS Dips | | |

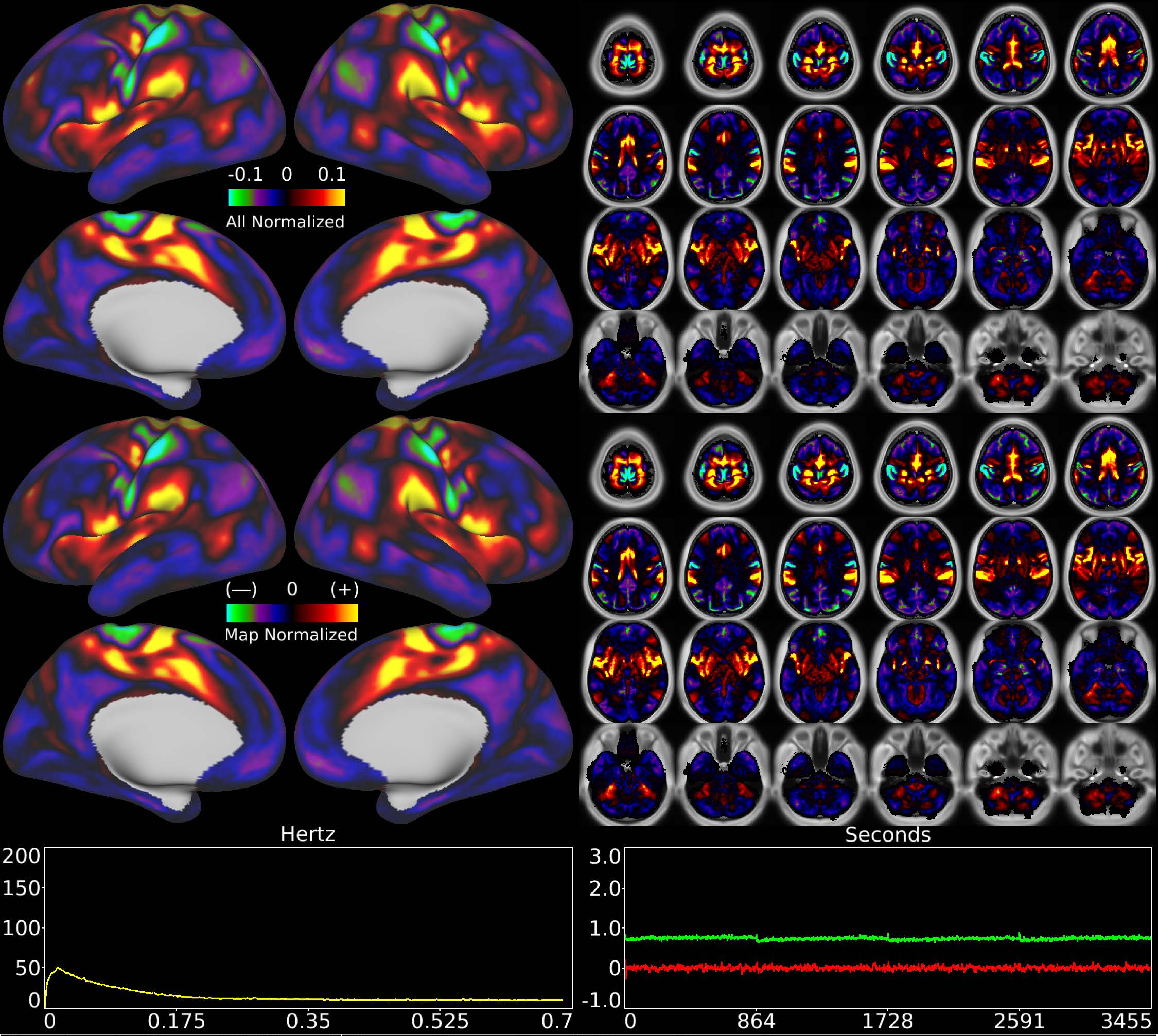


| | | |
|---|---|---|
| Number & Class: 39 Signal | Name: Eye + Trunk Motor Network | |
| RVT Correlated: No | DVARS Dip Associated: Yes | Cross-Subject Variable: No |
| Single Subject: No | % Variance Explained: 0.86 | Globality Index: 0.26 |
| Task Component: 29 | Taskr Component: 28 | Sleep Noted: No |
| Rationale: Spatial map includes positive and negative patches that respect known somatotopic sensori-motor organization (Eye and Trunk) | | |

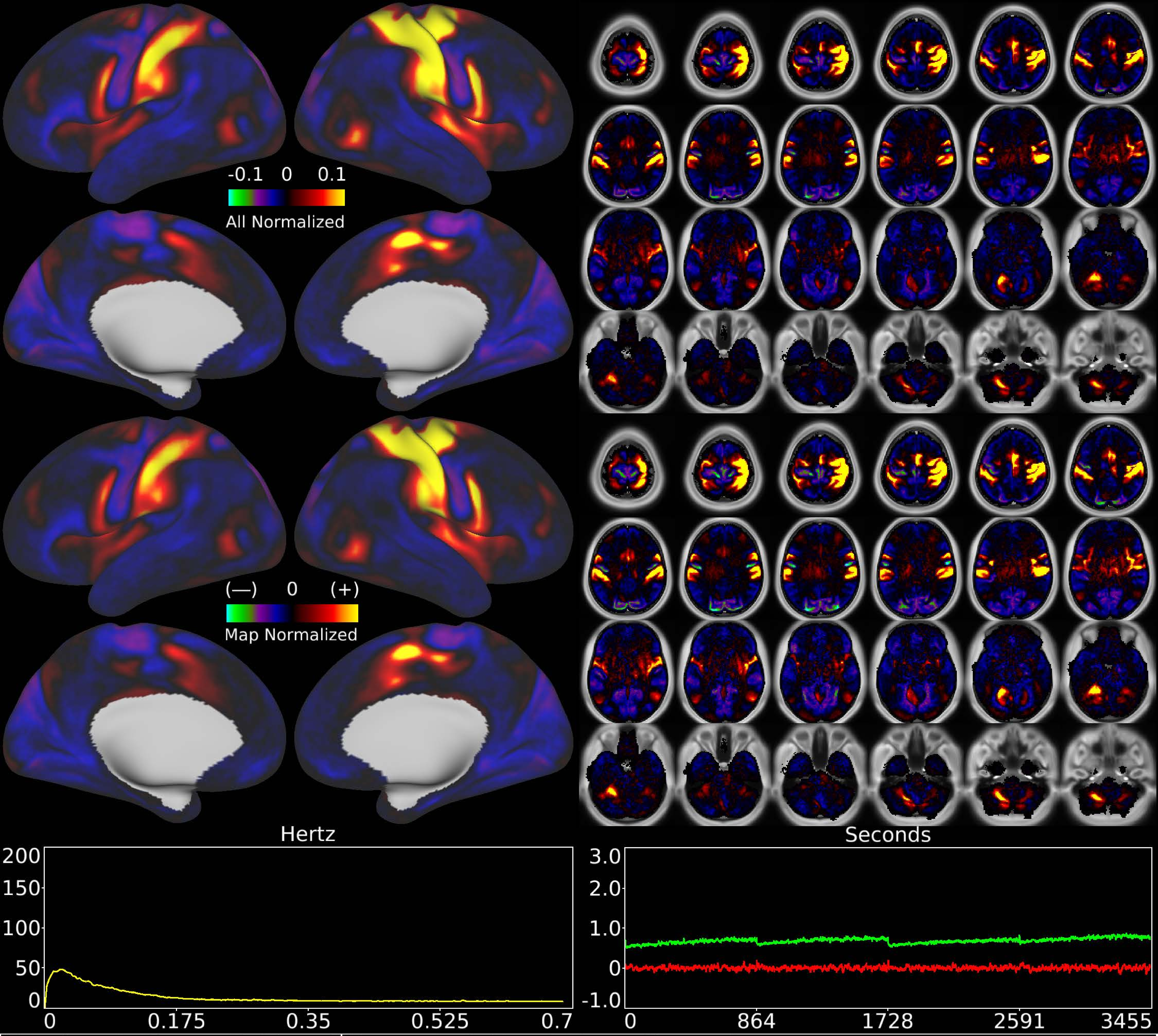


| | | |
|---|---|---|
| Number & Class: 40 Signal | Name: Left Hand Motor Network | |
| RVT Correlated: No | DVARS Dip Associated: No | Cross-Subject Variable: Yes |
| Single Subject: No | % Variance Explained: 0.83 | Globality Index: 0.76 |
| Task Component: 31 | Taskr Component: 26 | Sleep Noted: No |
| Rationale: Spatial map includes positive and negative patches that respect known somatotopic sensori-motor organization (Left Hand) | | |

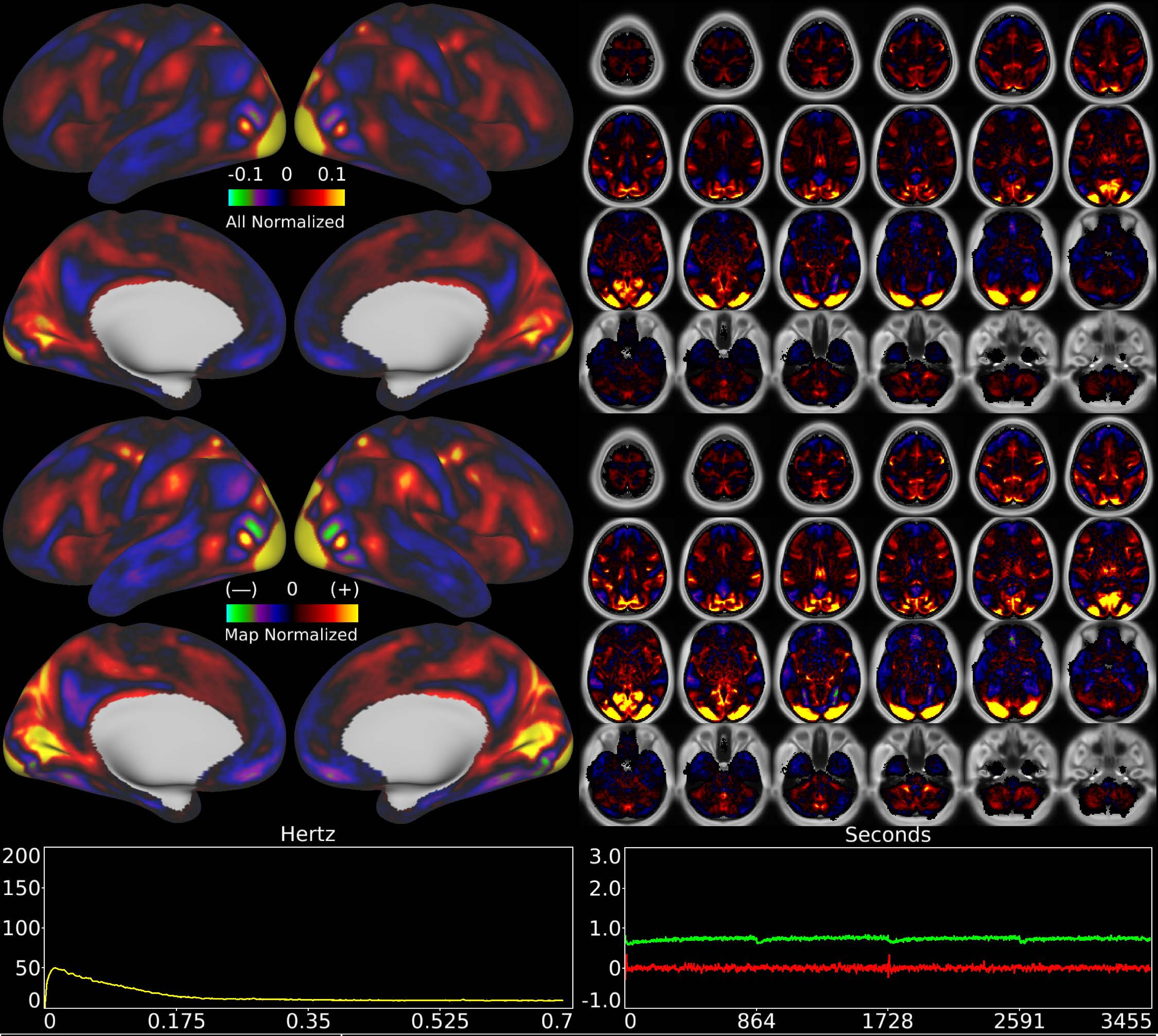


| | | |
|---|---|---|
| Number & Class: 41 Signal | Name: Pan Visual Foveal | |
| RVT Correlated: No | DVARS Dip Associated: No | Cross-Subject Variable: No |
| Single Subject: No | % Variance Explained: 0.83 | Globality Index: 0.73 |
| Task Component: 34 | Taskr Component: 15 | Sleep Noted: No |
| Rationale: Spatial map includes positive and negative patches that respect known retinotopic visual organization (Foveal vs ParaCentral) | | |

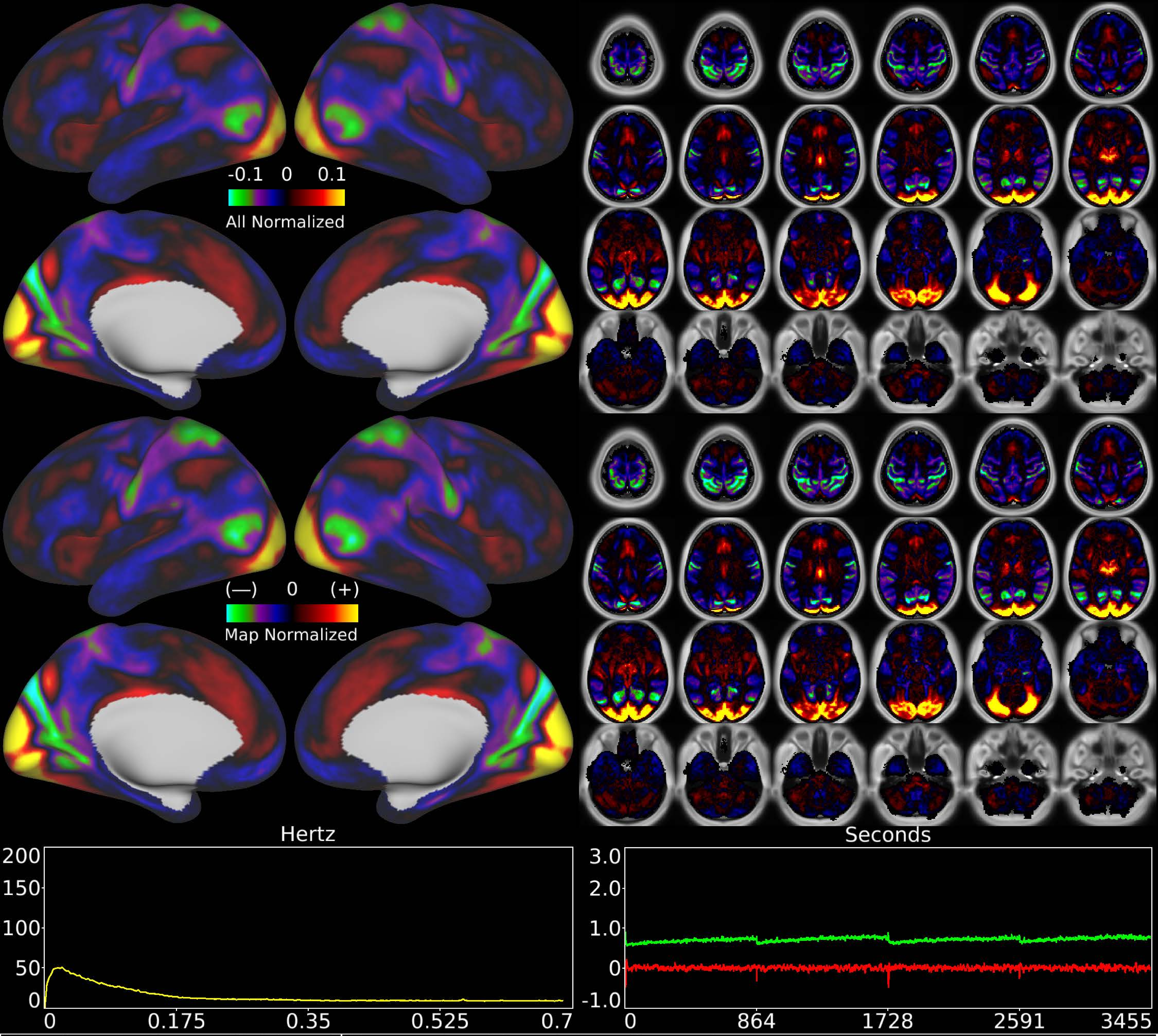


| | | |
|---|---|---|
| Number & Class: 42 Signal | Name: Pan Visual ParaCentral | |
| RVT Correlated: No | DVARS Dip Associated: No | Cross-Subject Variable: No |
| Single Subject: No | % Variance Explained: 0.79 | Globality Index: 0.46 |
| Task Component: No | Taskr Component: 49 | Sleep Noted: No |
| Rationale: Spatial map includes positive and negative patches that respect known retinotopic visual organization (ParaCentral vs Peripheral) | | |

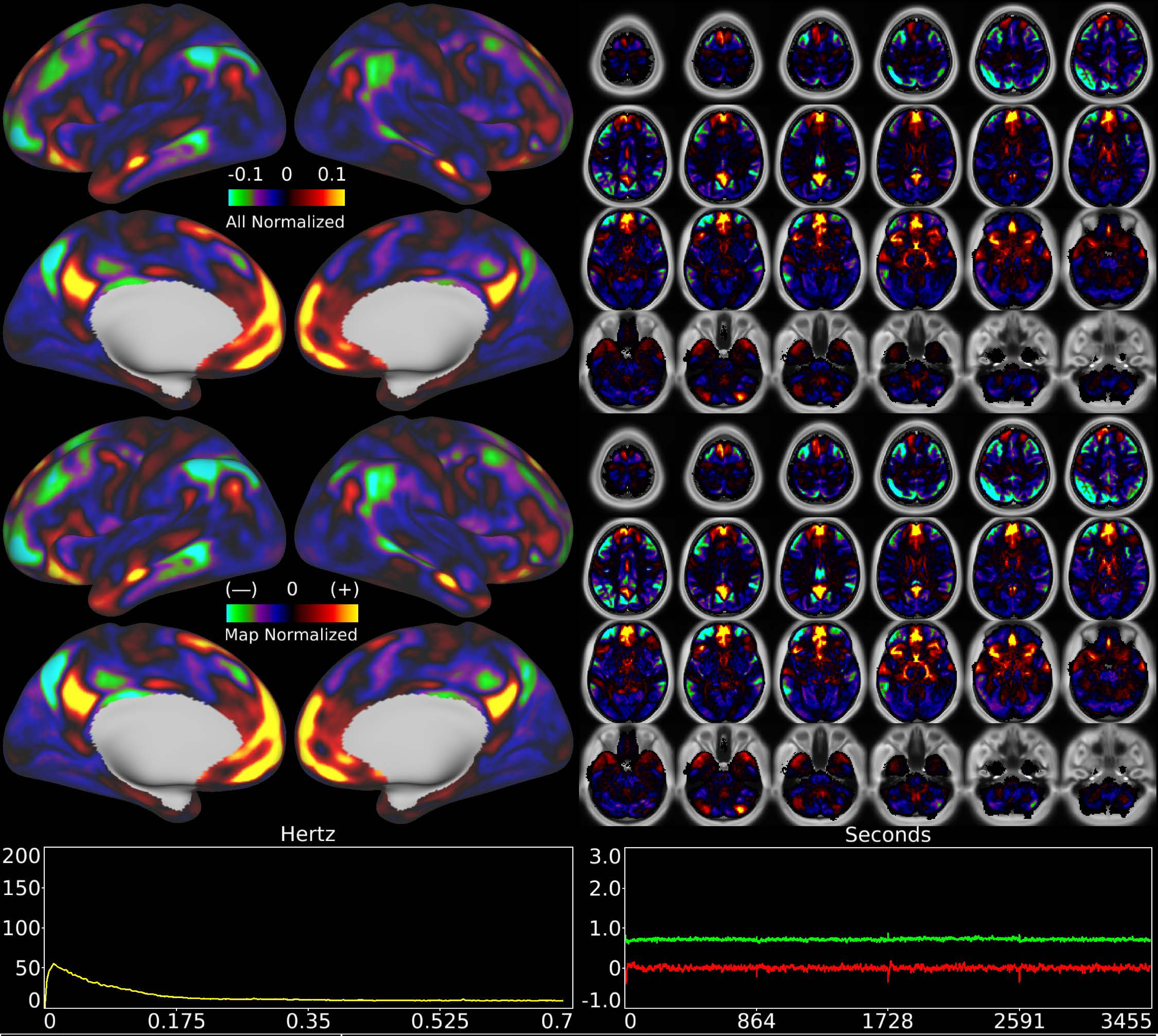


| | | |
|---|---|---|
| Number & Class: 43 Signal | Name: Subsidiary Default Mode | |
| RVT Correlated: No | DVARS Dip Associated: No | Cross-Subject Variable: No |
| Single Subject: No | % Variance Explained: 0.78 | Globality Index: 0.92 |
| Task Component: No | Taskr Component: No | Sleep Noted: No |
| Rationale: Spatial map includes positive and negative patches that respect known RSNs (e.g. Default Mode Network) | | |

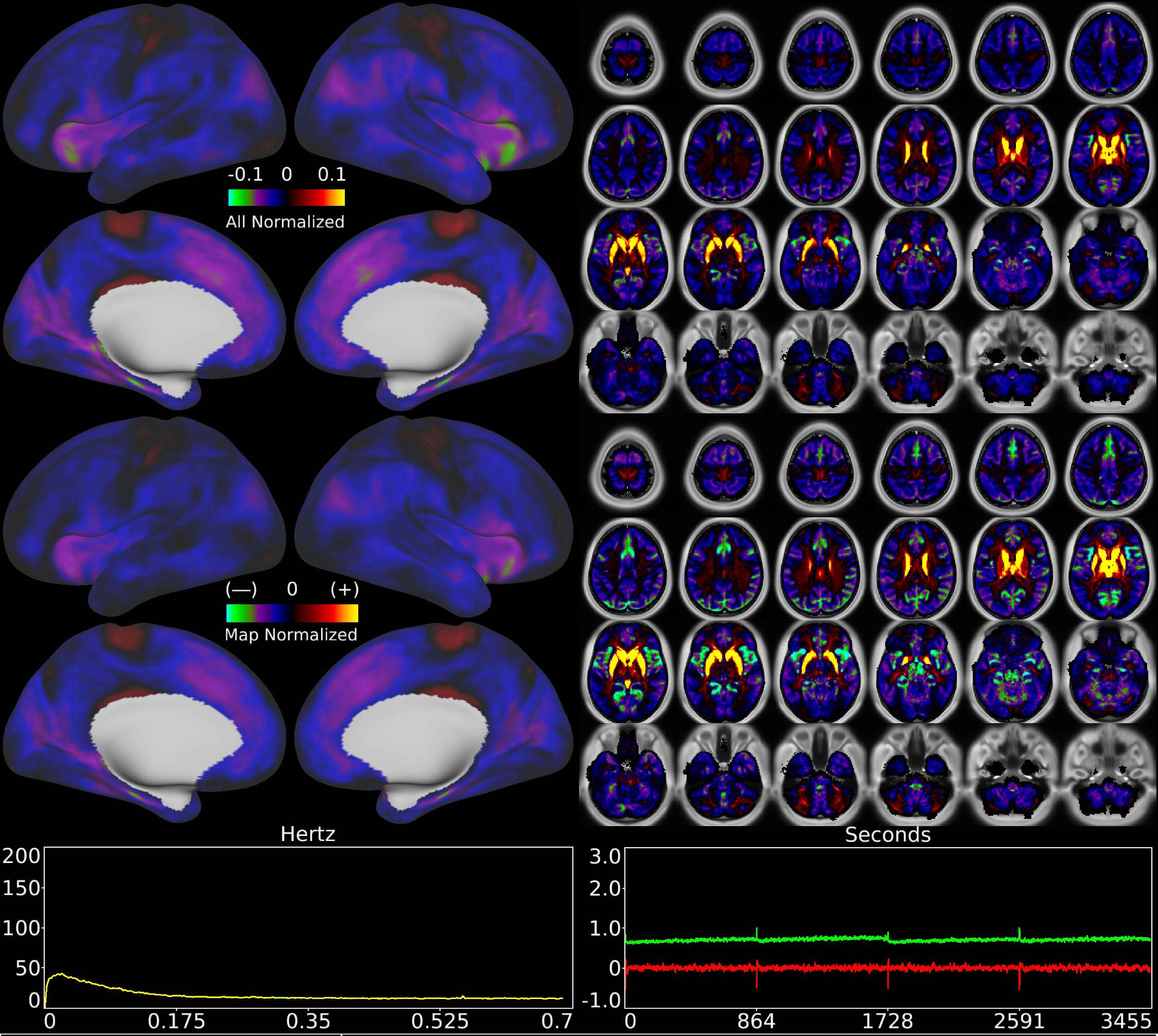


| | | |
|---|---|---|
| Number & Class: 44 Noise | Name: Striatum and Dorsal + Anterior Thalamus + WM | |
| RVT Correlated: No | DVARS Dip Associated: No | Cross-Subject Variable: No |
| Single Subject: No | % Variance Explained: 0.76 | Globality Index: 2.36 |
| Task Component: 42 | Taskr Component: 36 | Sleep Noted: No |
| Rationale: Controversial: Diencephalon together with surrounding white matter positive vs rest of brain negative; could be due to differing vascular supplies | | |

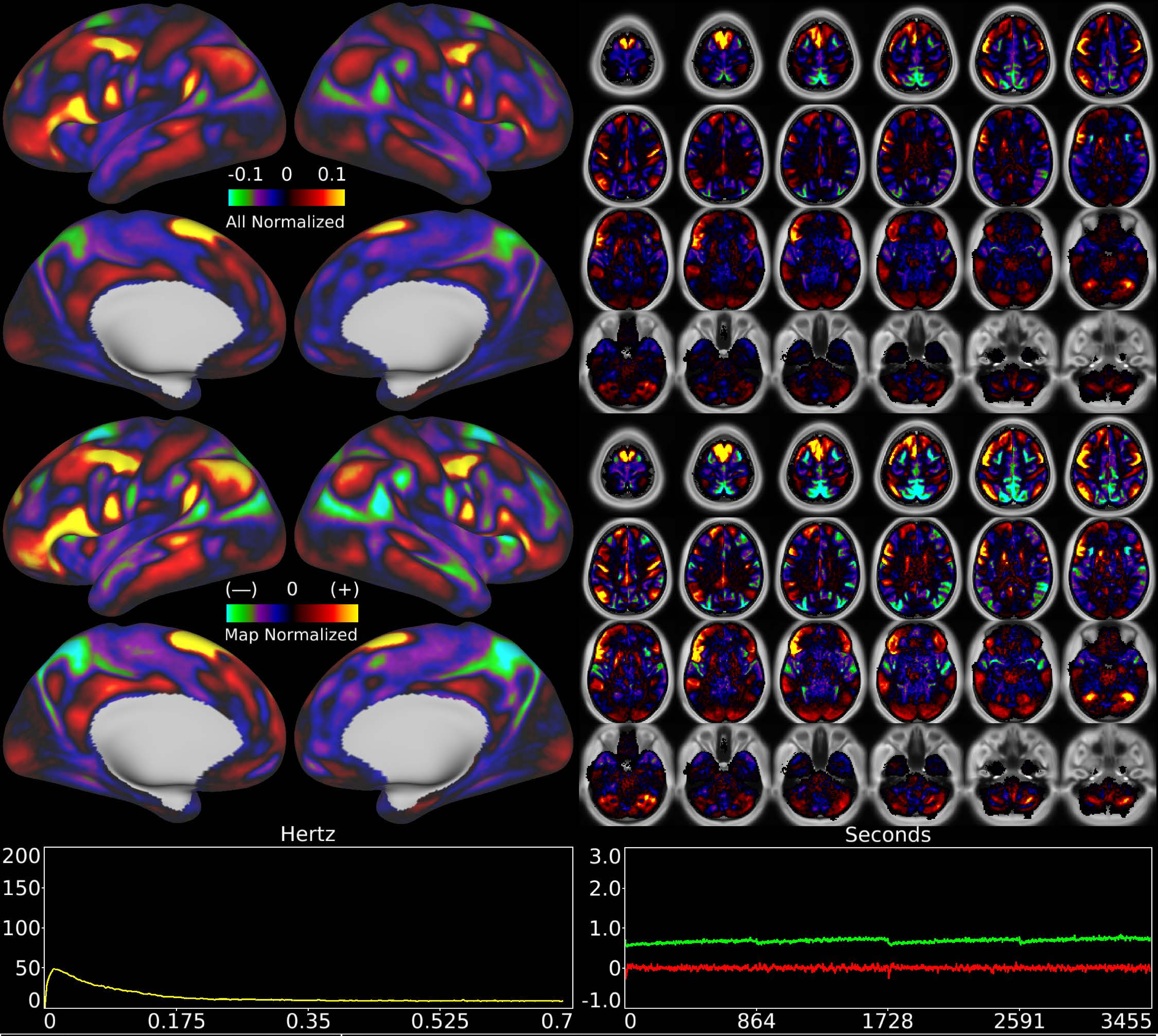


| | | |
|---|---|---|
| Number & Class: 45 Signal | Name: Left Lateralized Language Network II | |
| RVT Correlated: No | DVARS Dip Associated: No | Cross-Subject Variable: No |
| Single Subject: No | % Variance Explained: 0.73 | Globality Index: 0.47 |
| Task Component: No | Taskr Component: No | Sleep Noted: No |

Rationale: Spatial map includes positive and negative patches that respect known RSNs (e.g. Language Network) and areal boundaries (44; 45; 55b; PSL; SFL)

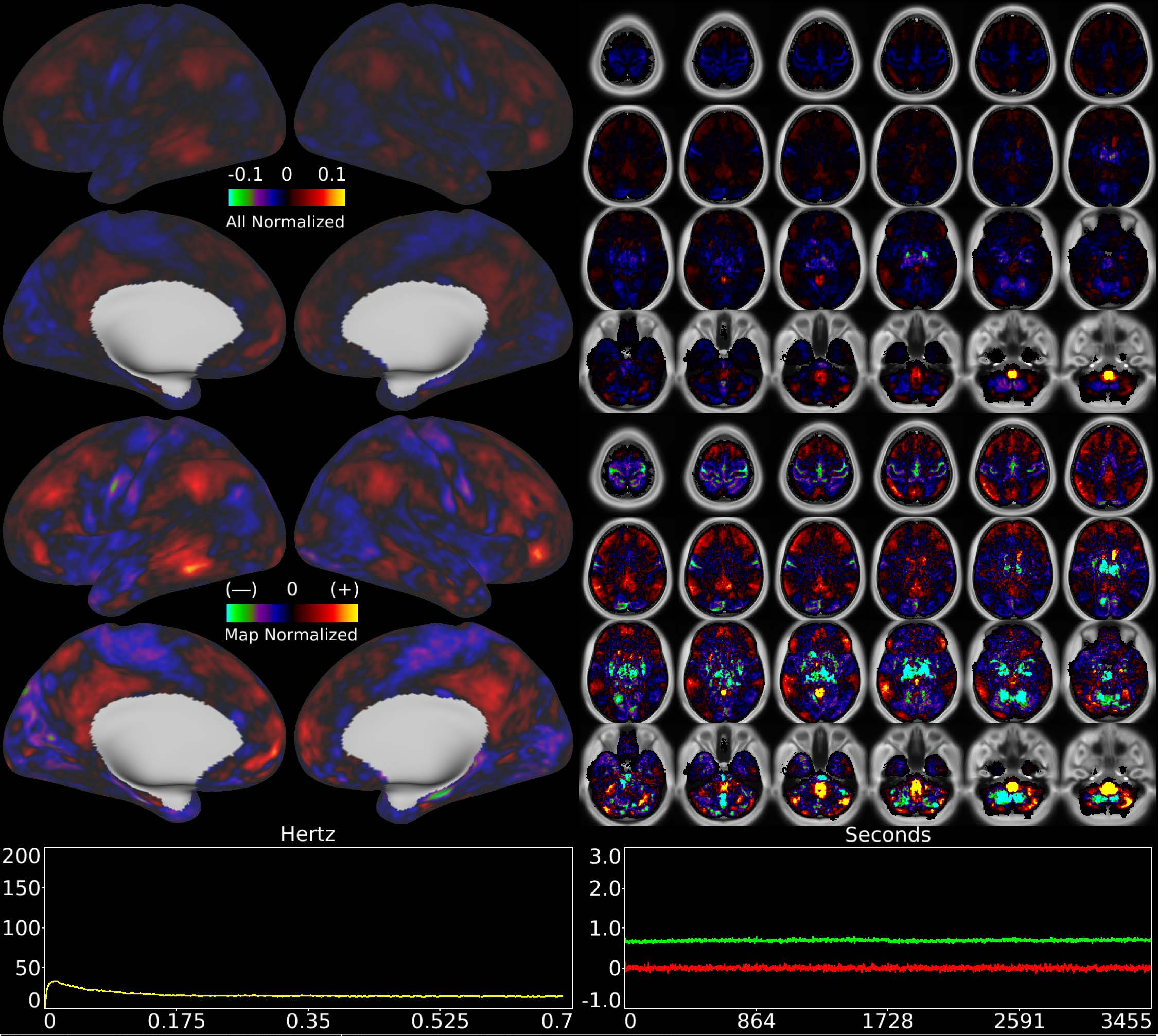


| | | |
|---|---|---|
| Number & Class: 46 Noise | Name: Medulla Recon Artifact | |
| RVT Correlated: No | DVARS Dip Associated: No | Cross-Subject Variable: No |
| Single Subject: No | % Variance Explained: 0.72 | Globality Index: 0.58 |
| Task Component: 43 | Taskr Component: 38 | Sleep Noted: No |
| Rationale: Spatial map not reflective of known areas or RSNs without connectivity other brain structures; some banding in sagittal plane suggestive of multi-band recon | | |

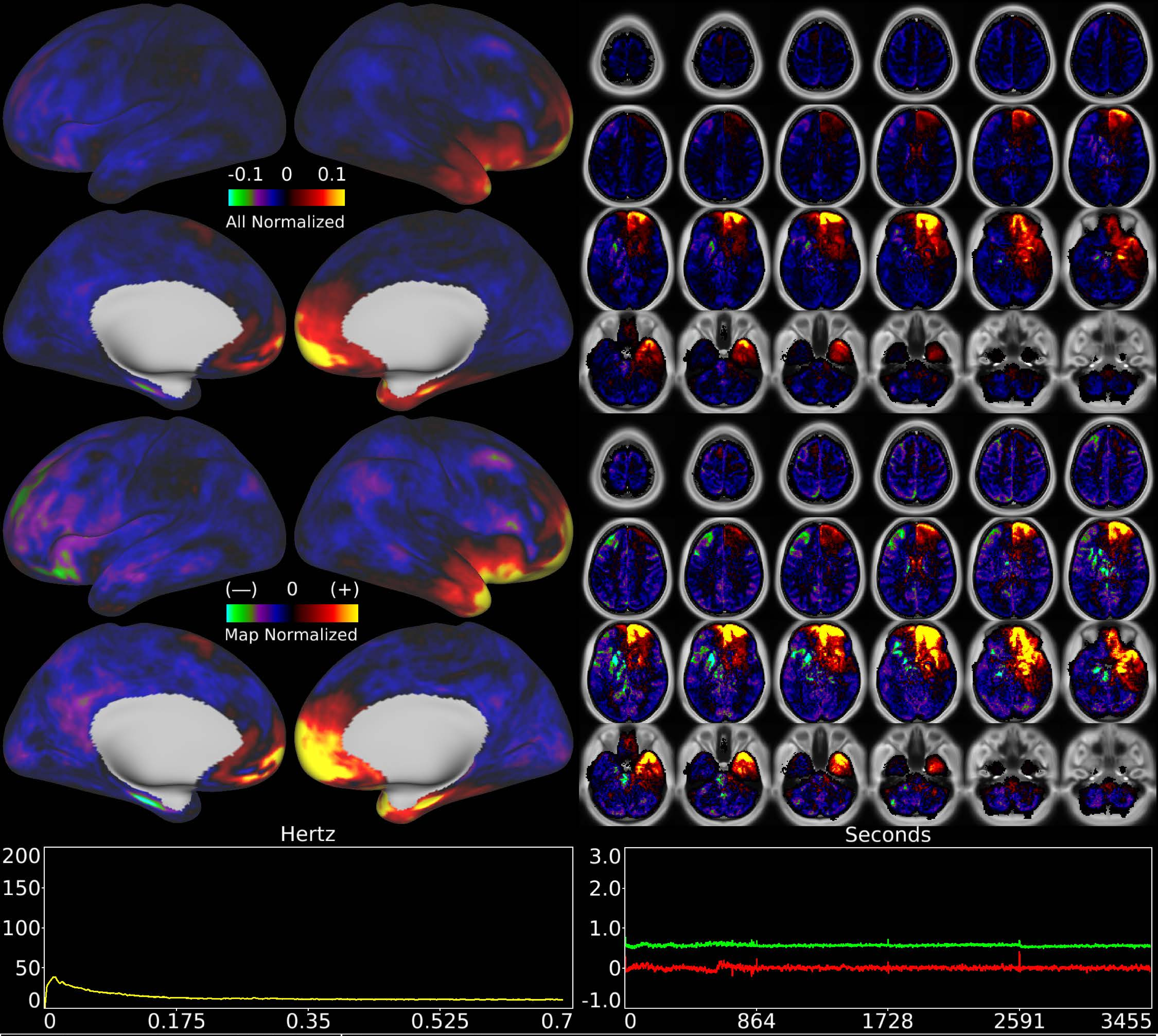


| | | |
|---|---|---|
| Number & Class: 47 Noise | Name: Coil | |
| RVT Correlated: No | DVARS Dip Associated: Yes | Cross-Subject Variable: Yes |
| Single Subject: Yes | % Variance Explained: 0.7 | Globality Index: 2.16 |
| Task Component: 50 | Taskr Component: 50 | Sleep Noted: Yes |
| Rationale: Known single run artifact due to coil element failure | | |

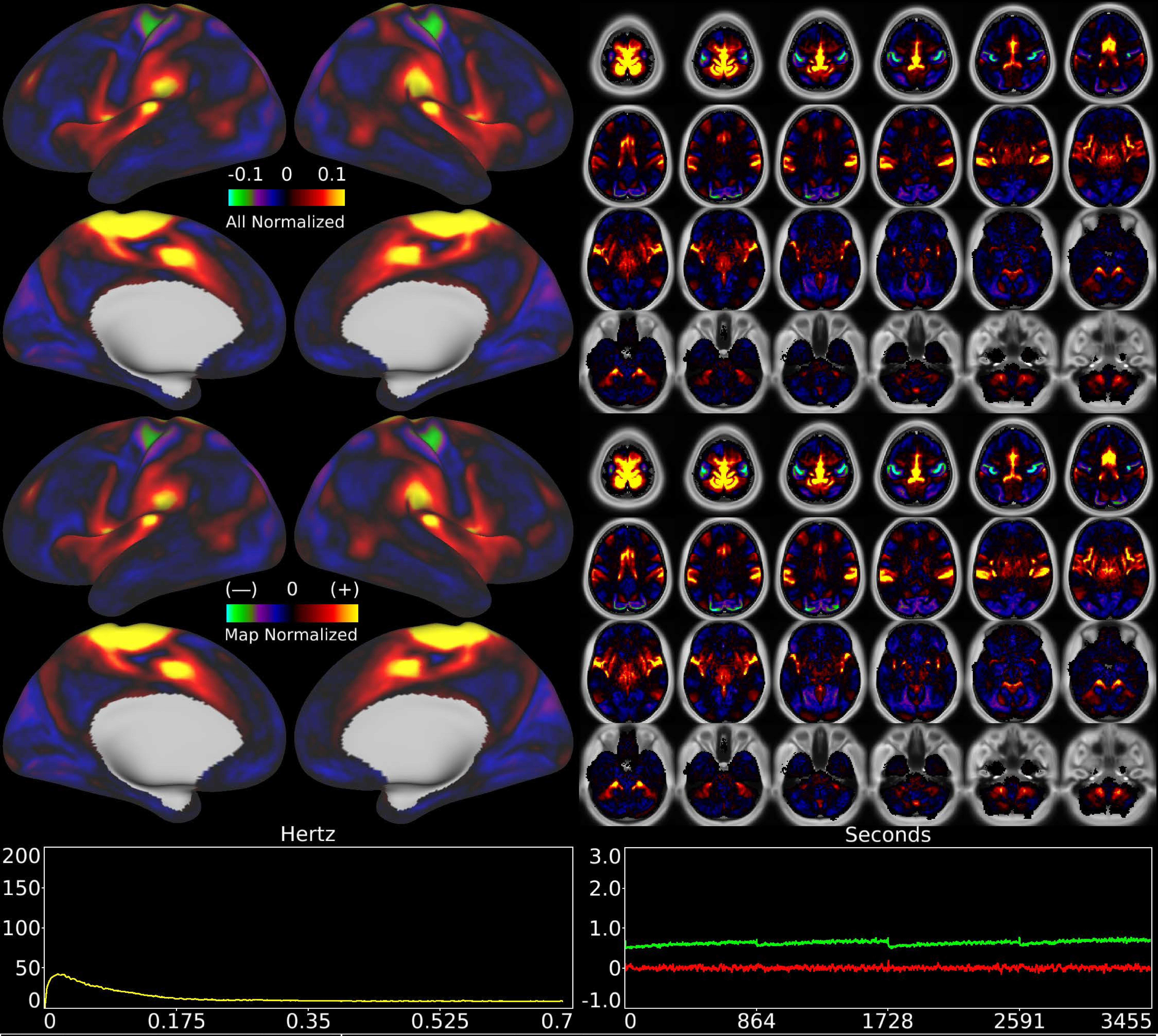


| | | |
|---|---|---|
| Number & Class: 48 Signal | Name: Feet Motor Network | |
| RVT Correlated: No | DVARS Dip Associated: Yes | Cross-Subject Variable: Yes |
| Single Subject: No | % Variance Explained: 0.67 | Globality Index: 0.21 |
| Task Component: 19 | Taskr Component: 24 | Sleep Noted: No |
| Rationale: Spatial map includes positive and negative patches that respect known somatotopic sensori-motor organization (Feet) | | |

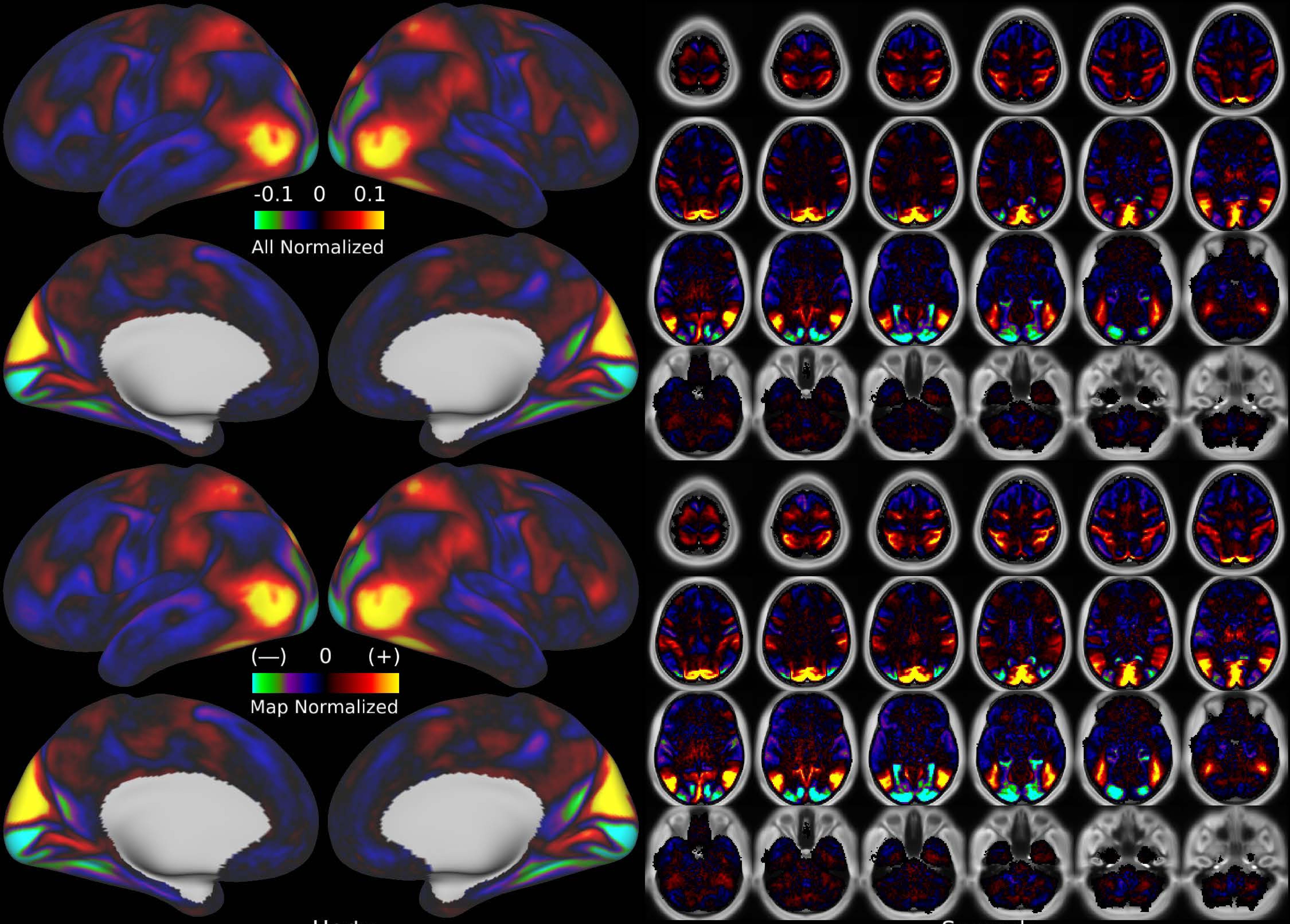


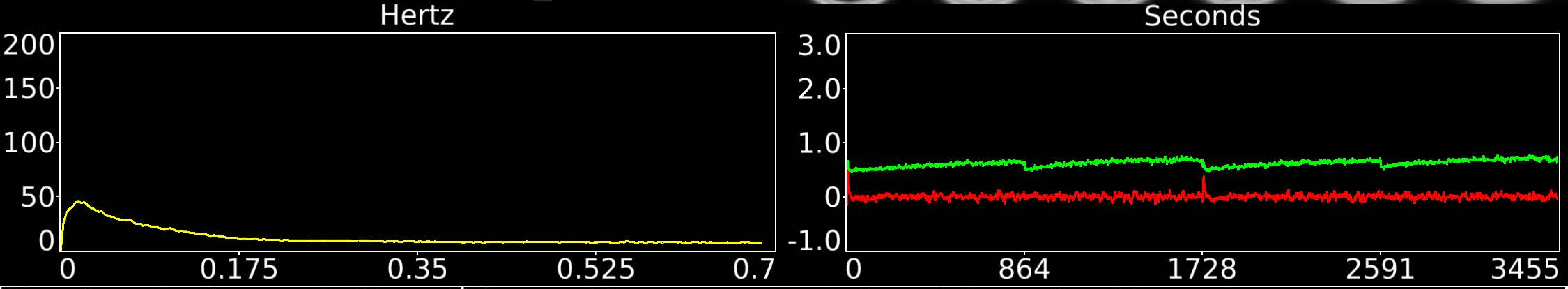


| Number & Class: 49 Signal | Name: Visuotopy: Peripheral Dorsal > Ventral | |
|---|---|---|
| RVT Correlated: No | DVARS Dip Associated: No | Cross-Subject Variable: Yes |
| Single Subject: No | % Variance Explained: 0.66 | Globality Index: 0.13 |
| Task Component: No | Taskr Component: No | Sleep Noted: No |
| Rationale: Spatial map includes positive and negative patches that respect known retinotopic visual organization (Lower vs Upper) | | |

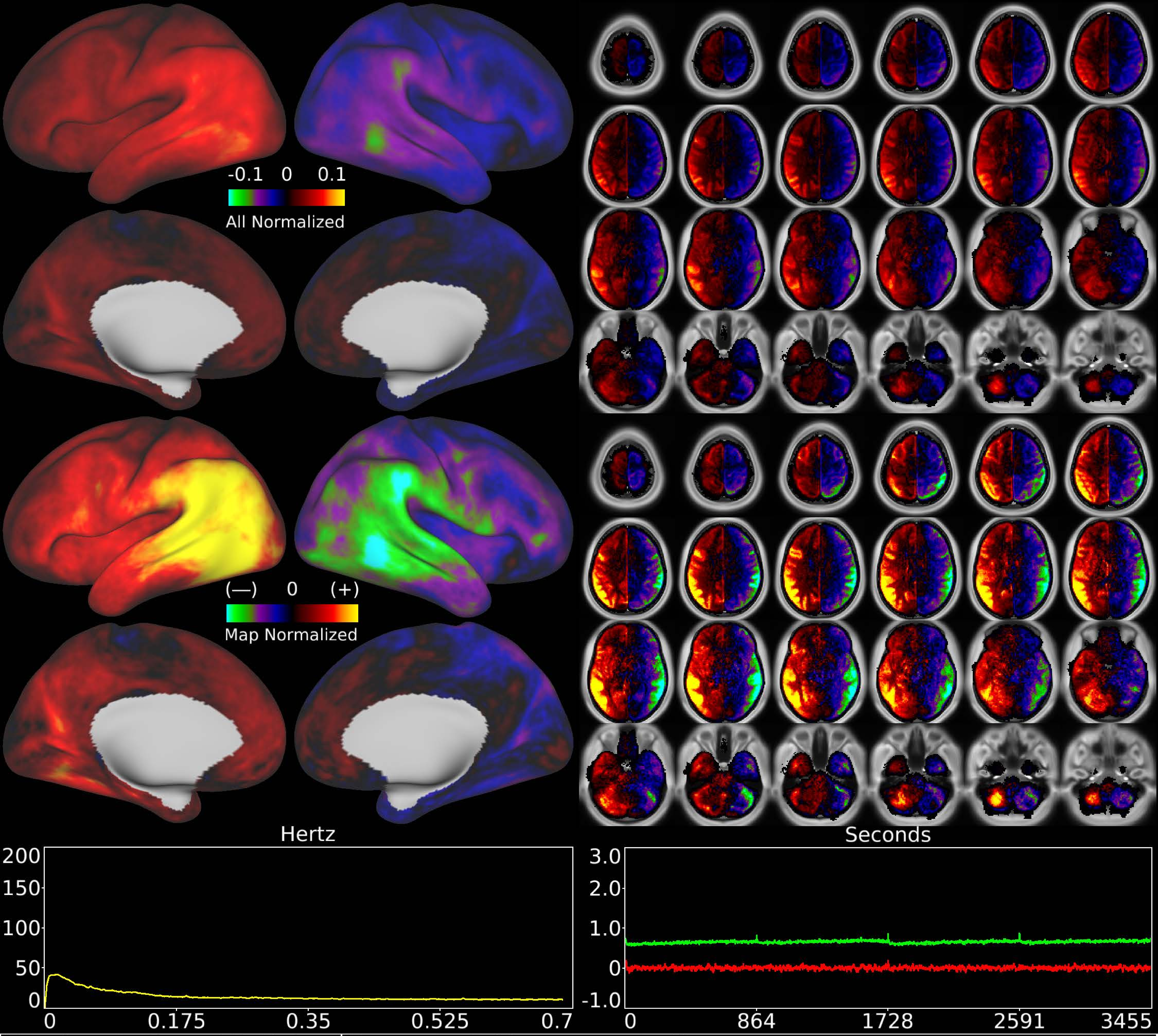


| | | |
|---|---|---|
| Number & Class: 50 Noise | Name: L>R Movement Artifact | |
| RVT Correlated: No | DVARS Dip Associated: Yes | Cross-Subject Variable: No |
| Single Subject: No | % Variance Explained: 0.65 | Globality Index: 0.29 |
| Task Component: 51 | Taskr Component: 37 | Sleep Noted: No |
| Rationale: Highly DVARS Dips associated component; looks like movement regressor beta map (Z rotation) | | |

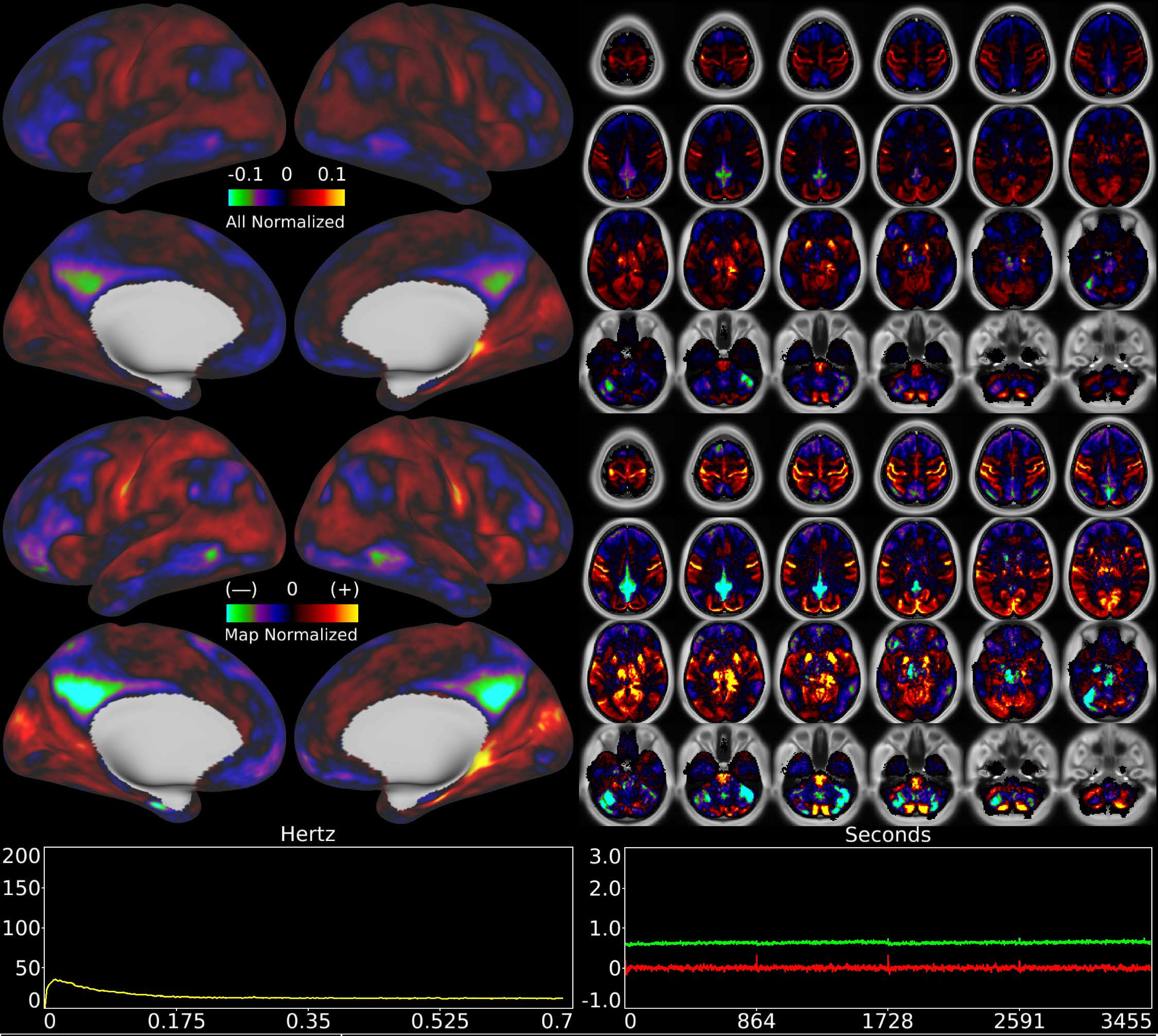


| | | |
|---|---|---|
| Number & Class: 51 Noise | Name: Single Run of Single Subject: Highly Globally Correlated | |
| RVT Correlated: No | DVARS Dip Associated: No | Cross-Subject Variable: No |
| Single Subject: Yes | % Variance Explained: 0.61 | Globality Index: 0.45 |
| Task Component: No | Taskr Component: No | Sleep Noted: No |
| Rationale: Single subject component with high correlation to global timecourse | | |

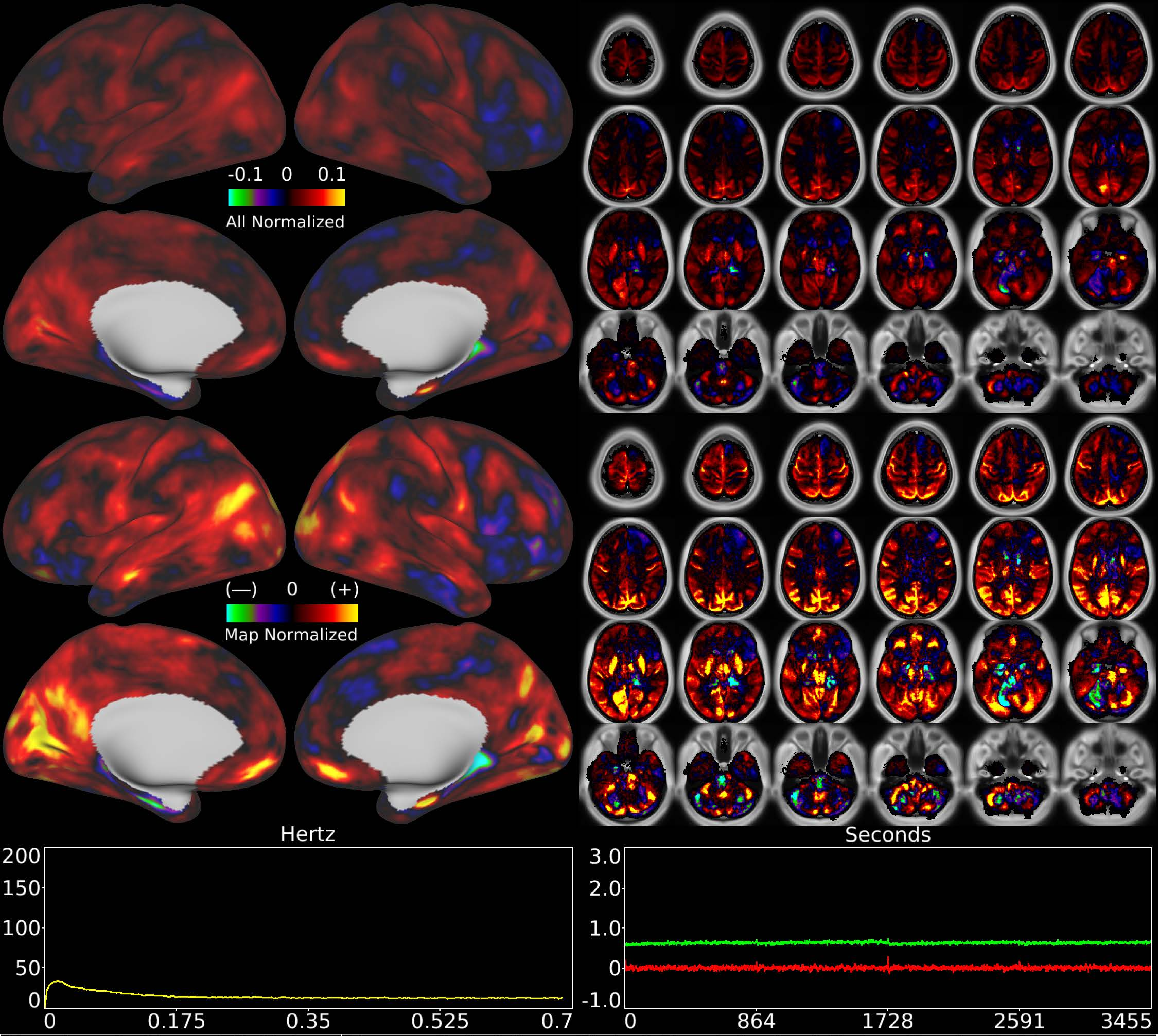


| | | |
|---|---|---|
| Number & Class: 52 Noise | Name: Subsidiary Global Physiological Noise Single Subject | |
| RVT Correlated: No | DVARS Dip Associated: No | Cross-Subject Variable: No |
| Single Subject: Yes | % Variance Explained: 0.6 | Globality Index: 1.51 |
| Task Component: No | Taskr Component: No | Sleep Noted: No |
| Rationale: Single subject component with high correlation to global timecourse | | |

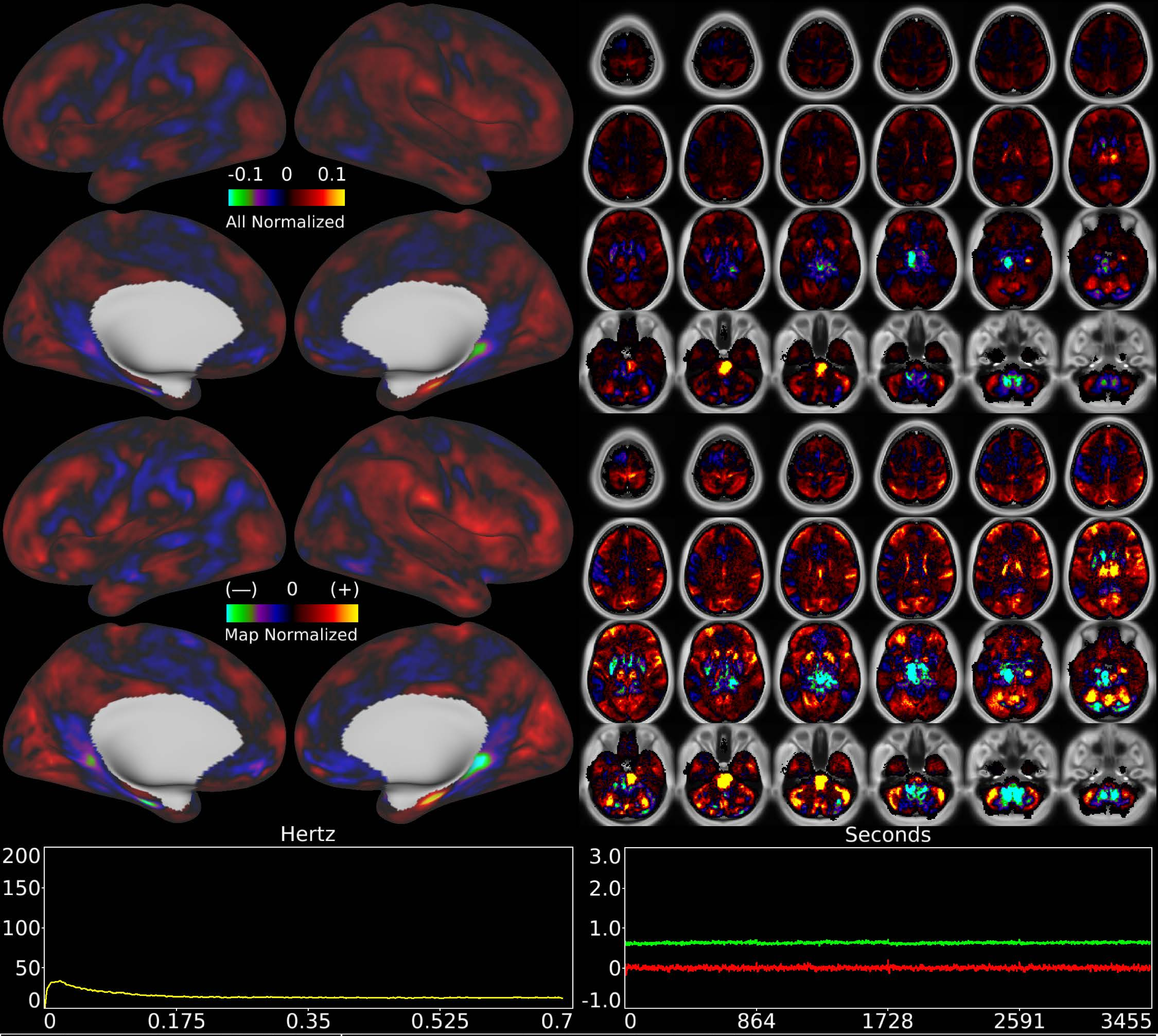


| | | |
|---|---|---|
| Number & Class: 53 Noise | Name: Recon Artifact Single Subject | |
| RVT Correlated: No | DVARS Dip Associated: No | Cross-Subject Variable: No |
| Single Subject: Yes | % Variance Explained: 0.6 | Globality Index: 0.85 |
| Task Component: No | Taskr Component: No | Sleep Noted: No |
| Rationale: Single subject component with obvious banding in sagittal plane suggestive of multi-band recon artifact along slice direction | | |

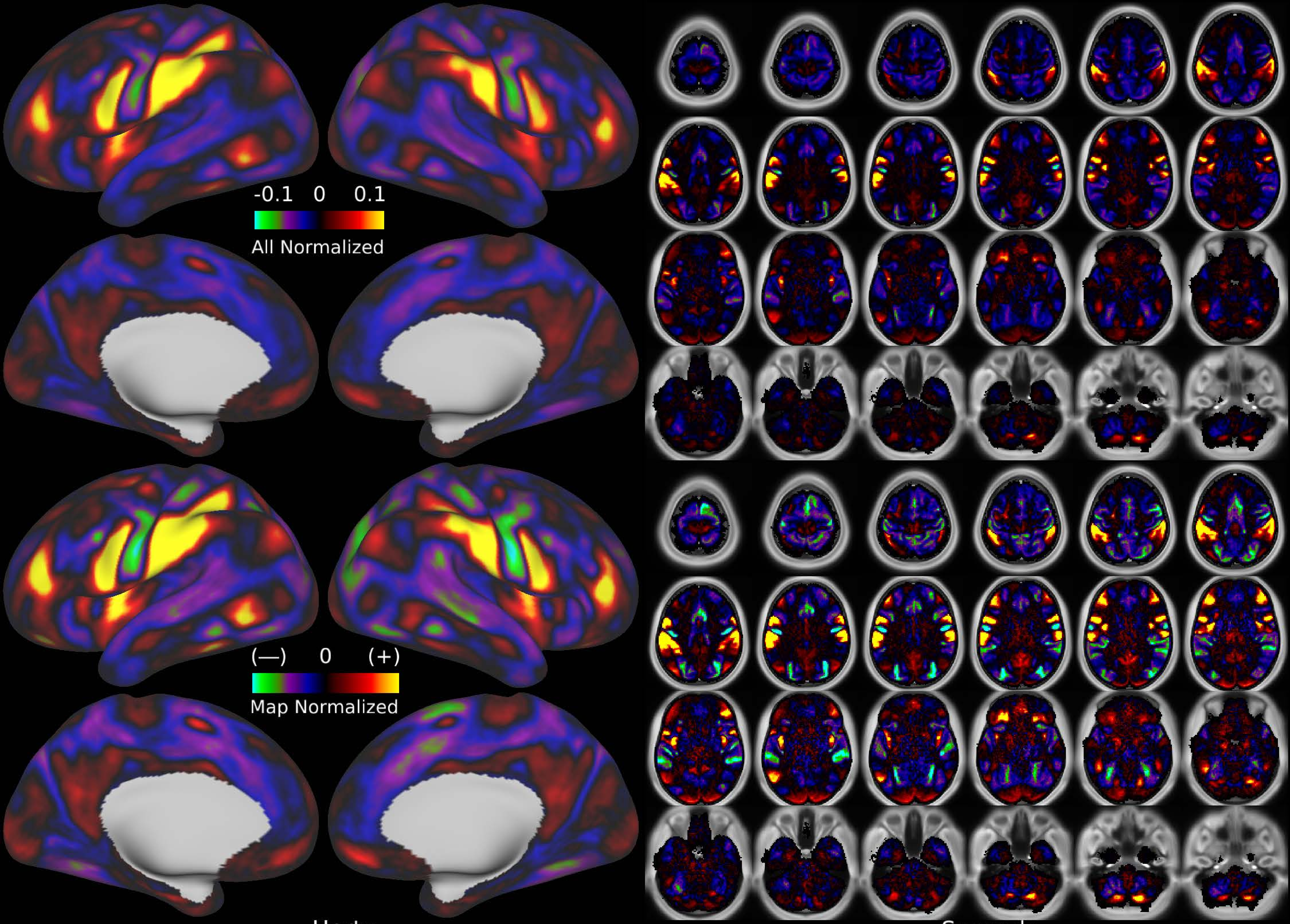


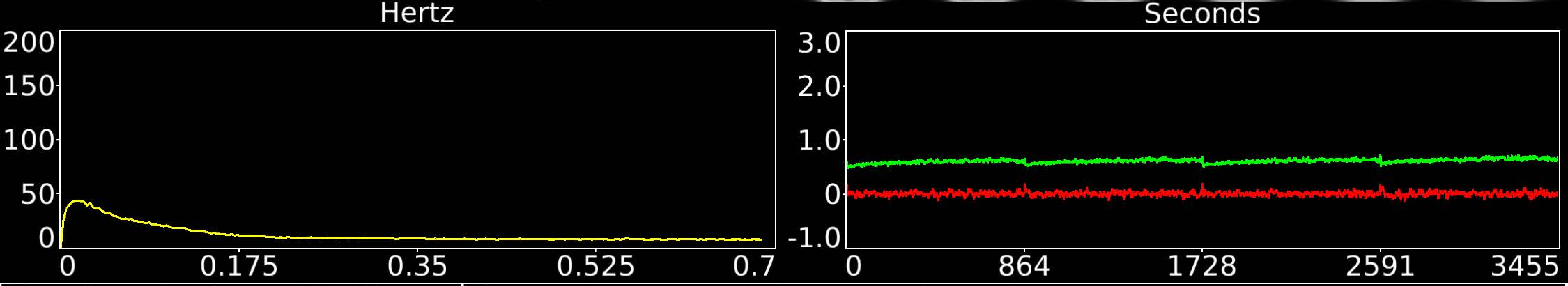


| | | |
|---|---|---|
| Number & Class: 54 Signal | Name: Motor and Sensory Association | |
| RVT Correlated: No | DVARS Dip Associated: No | Cross-Subject Variable: No |
| Single Subject: No | % Variance Explained: 0.58 | Globality Index: 0.48 |
| Task Component: No | Taskr Component: 35 | Sleep Noted: No |
| Rationale: Spatial map includes positive and negative patches that respect known areal boundaries (e.g. 4 and 1) | | |

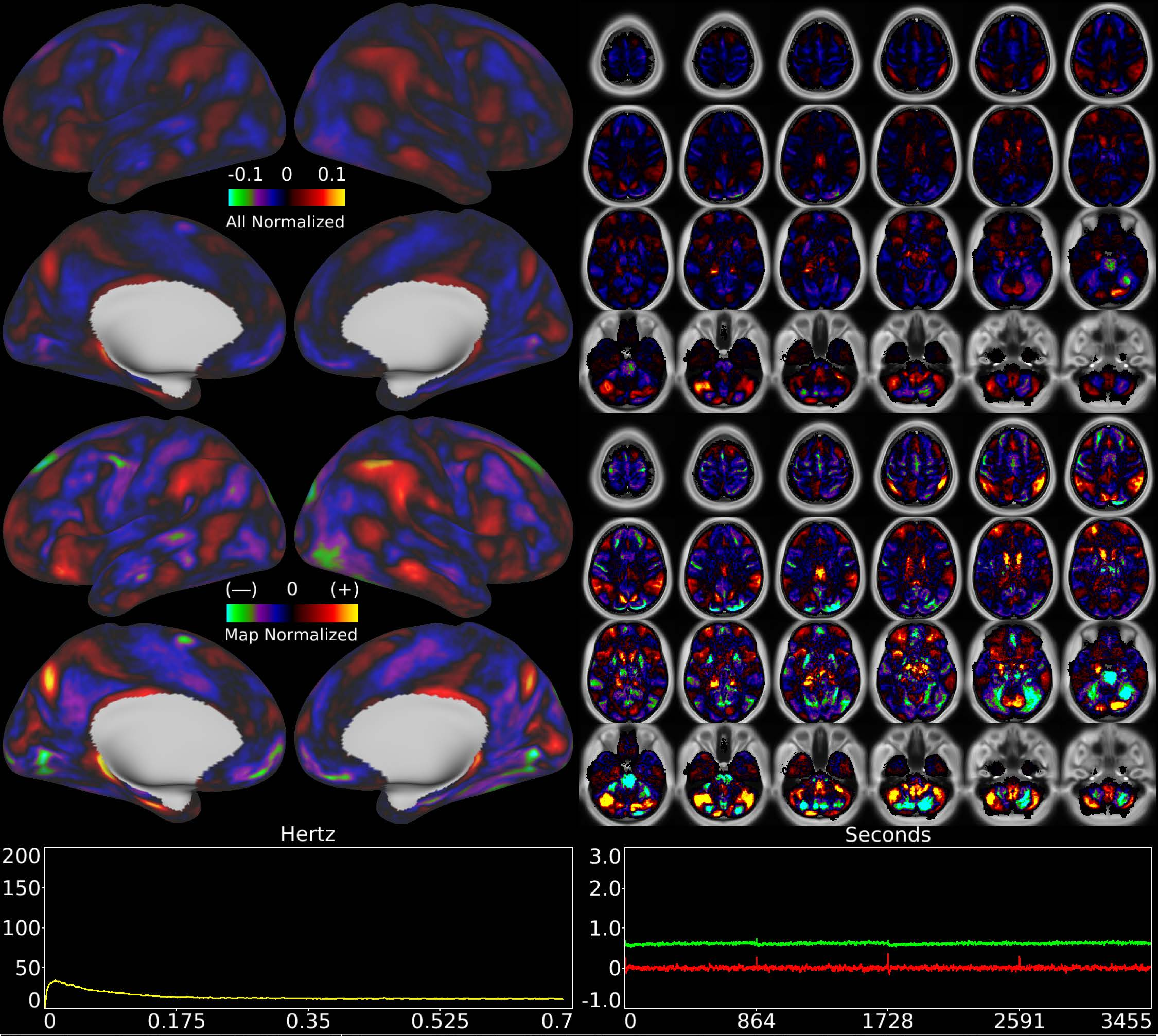


| | | |
|---|---|---|
| Number & Class: 55 Signal | Name: Highly Globally Correlated Single Subject: Sleep? | |
| RVT Correlated: No | DVARS Dip Associated: No | Cross-Subject Variable: No |
| Single Subject: Yes | % Variance Explained: 0.57 | Globality Index: 0.54 |
| Task Component: No | Taskr Component: No | Sleep Noted: No |
| Rationale: Controversial: Single subject component with high correlation to global timecourse; looked more like RC1 than global noise components) | | |

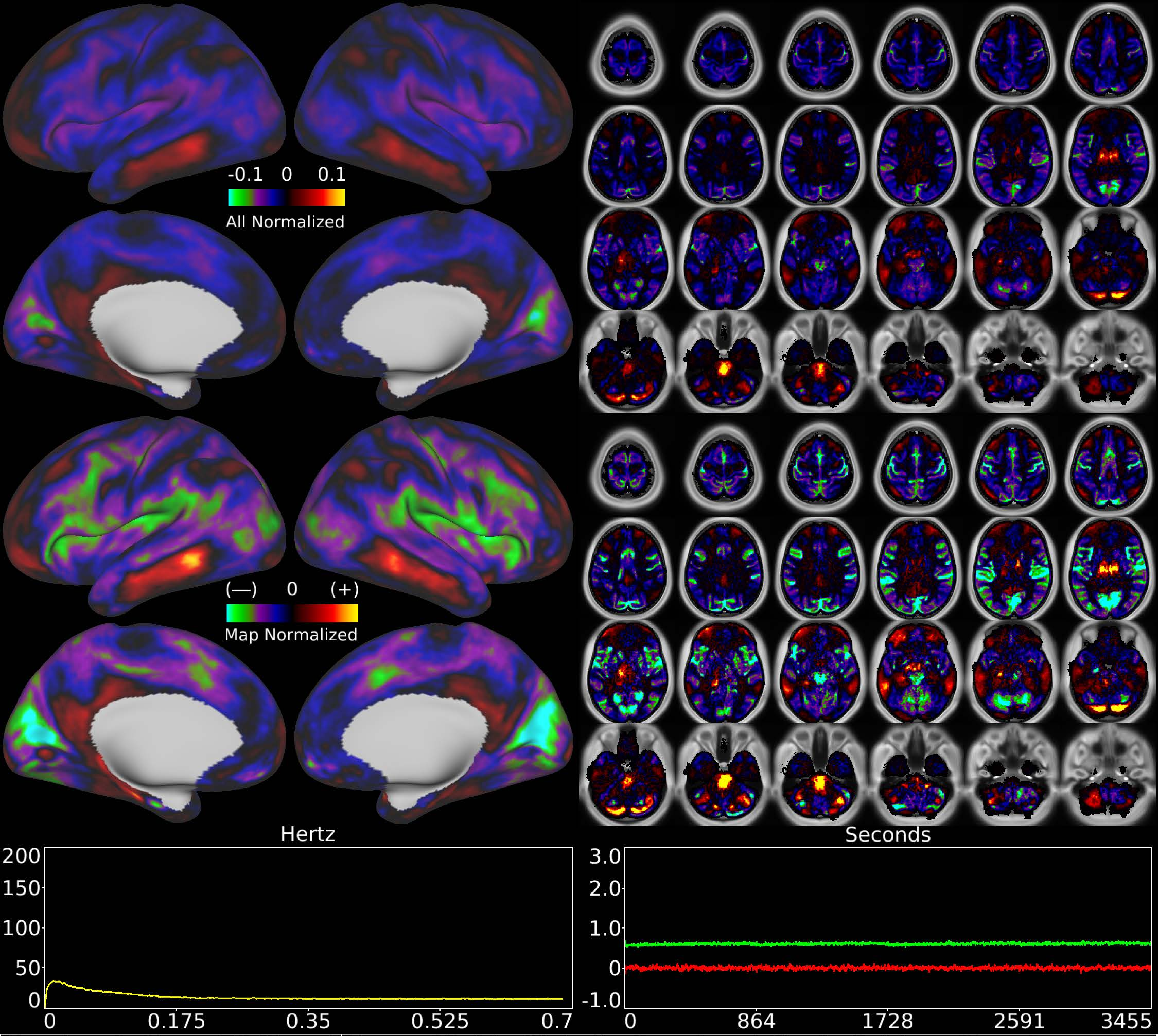


| Number & Class: 56 Signal | Name: Highly Globally Correlated Single Subject: Sleep? | |
|---|---|---|
| RVT Correlated: No | DVARS Dip Associated: No | Cross-Subject Variable: No |
| Single Subject: Yes | % Variance Explained: 0.56 | Globality Index: 1.4 |
| Task Component: No | Taskr Component: No | Sleep Noted: No |

Rationale: Controversial: Single subject component with high correlation to global timecourse; looked more like RC1 than global noise components)

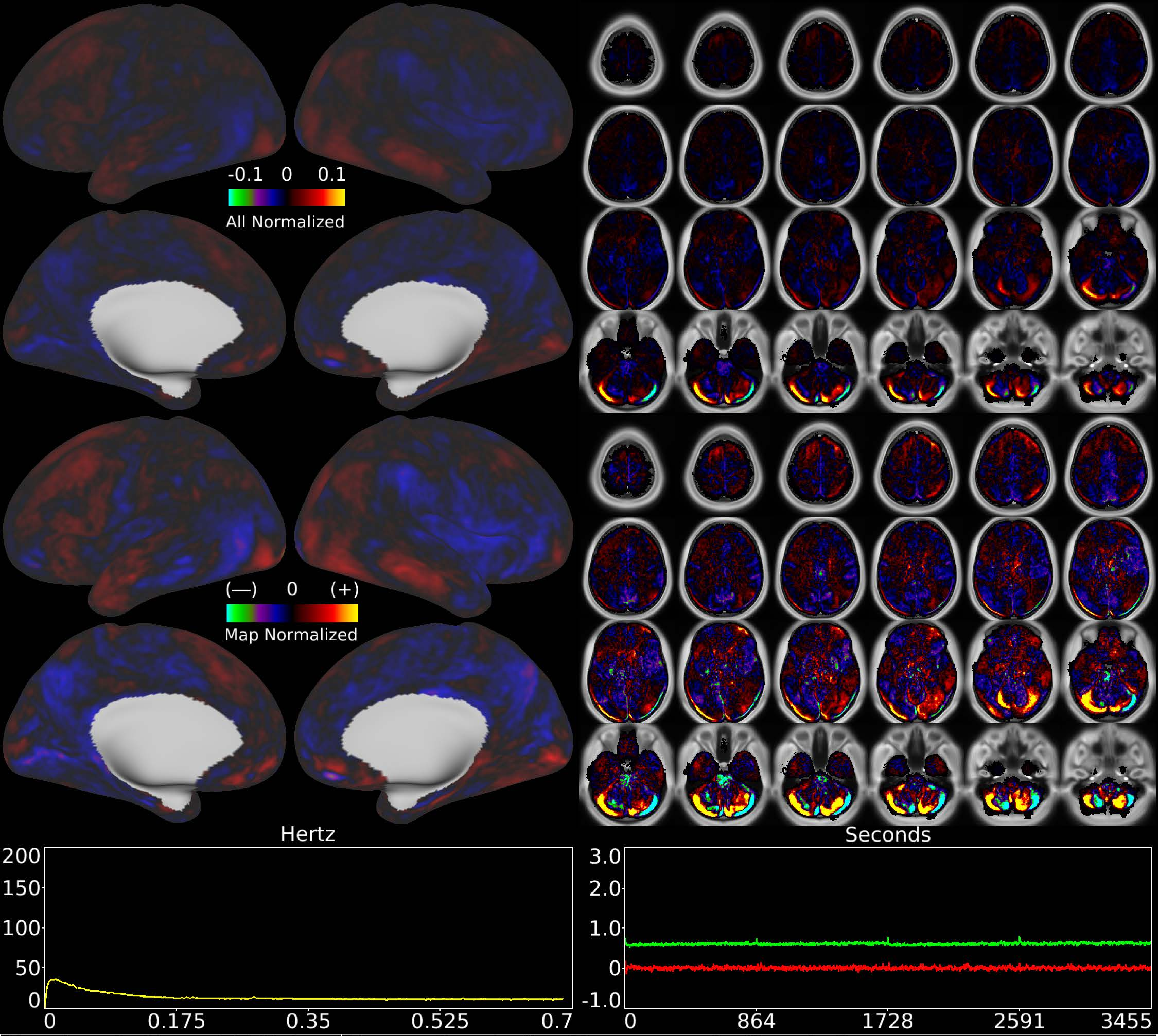


| | | |
|---|---|---|
| Number & Class: 57 Noise | Name: Cerebellar Movement Artifact Left | |
| RVT Correlated: No | DVARS Dip Associated: Yes | Cross-Subject Variable: No |
| Single Subject: No | % Variance Explained: 0.57 | Globality Index: 0.28 |
| Task Component: 44 | Taskr Component: 34 | Sleep Noted: No |

Rationale: Cerebellar edge motion component with high correlation to DVARS dips; looks like movement regressor beta map (derivative of X translation)

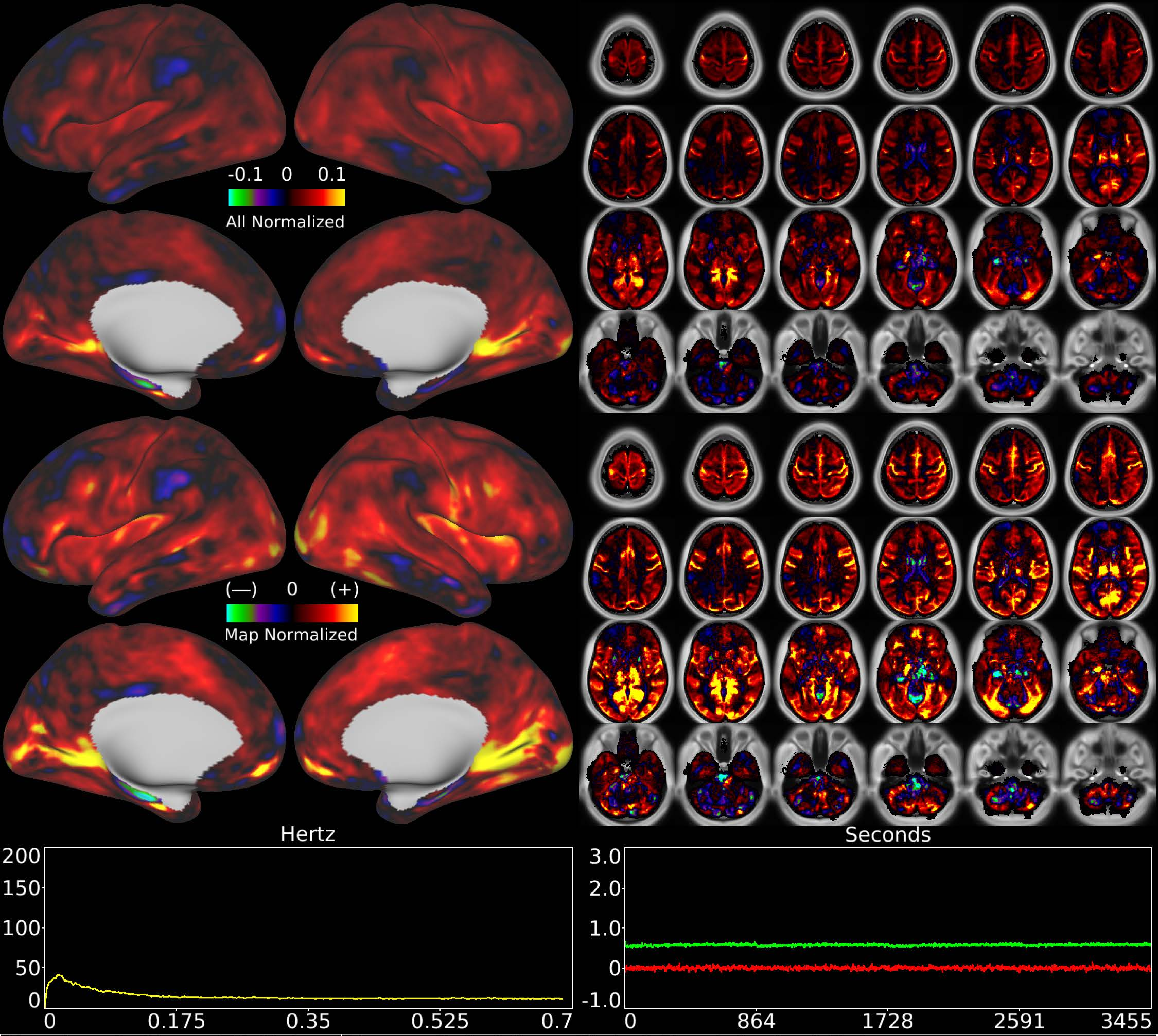


| | | |
|---|---|---|
| Number & Class: 58 Noise | Name: Subsidiary Global Physiological Noise | |
| RVT Correlated: No | DVARS Dip Associated: No | Cross-Subject Variable: Yes |
| Single Subject: Yes | % Variance Explained: 0.55 | Globality Index: 1.99 |
| Task Component: No | Taskr Component: No | Sleep Noted: Yes |
| Rationale: Single subject component with high correlation to global timecourse | | |

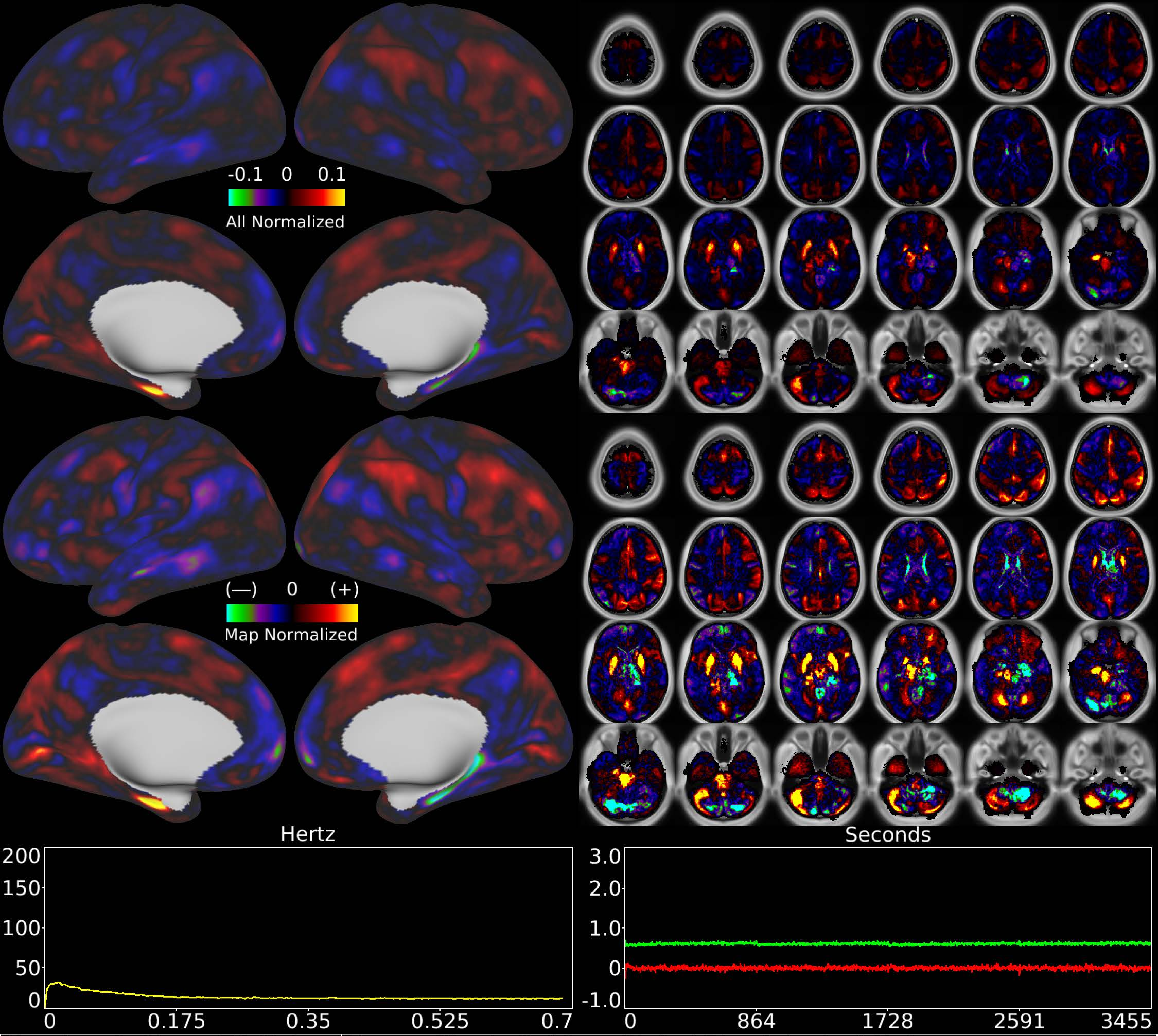


| | | |
|---|---|---|
| Number & Class: 59 Noise | Name: Single Subject Global Physiological Noise | |
| RVT Correlated: No | DVARS Dip Associated: No | Cross-Subject Variable: No |
| Single Subject: Yes | % Variance Explained: 0.56 | Globality Index: 0.13 |
| Task Component: No | Taskr Component: No | Sleep Noted: No |
| Rationale: Single subject component with high correlation to global timecourse | | |

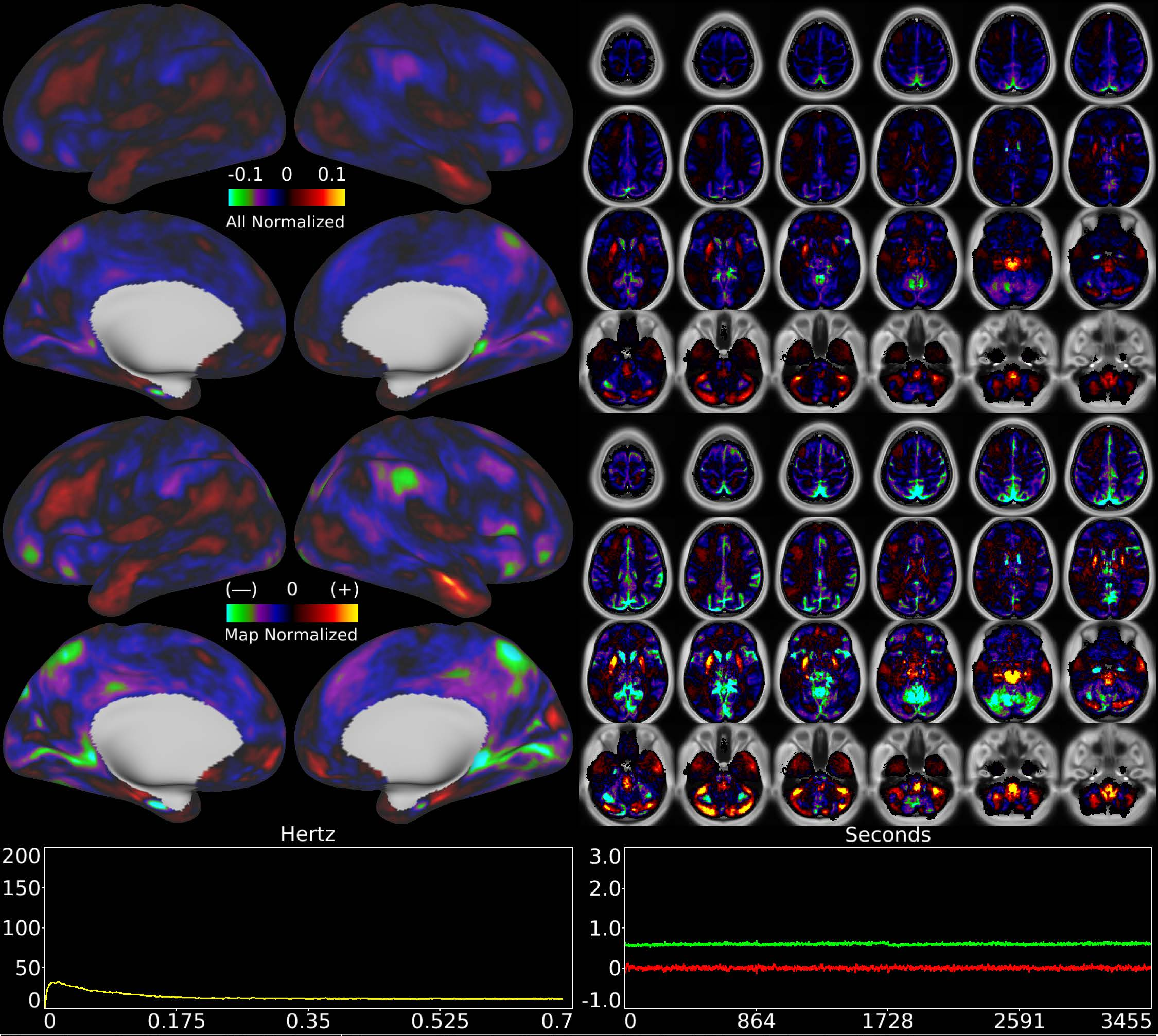


| | | |
|---|---|---|
| Number & Class: 60 Noise | Name: Single Subject Global Physiological Noise | |
| RVT Correlated: No | DVARS Dip Associated: No | Cross-Subject Variable: No |
| Single Subject: Yes | % Variance Explained: 0.54 | Globality Index: 1.05 |
| Task Component: No | Taskr Component: No | Sleep Noted: No |
| Rationale: Single subject component with high correlation to global timecourse | | |

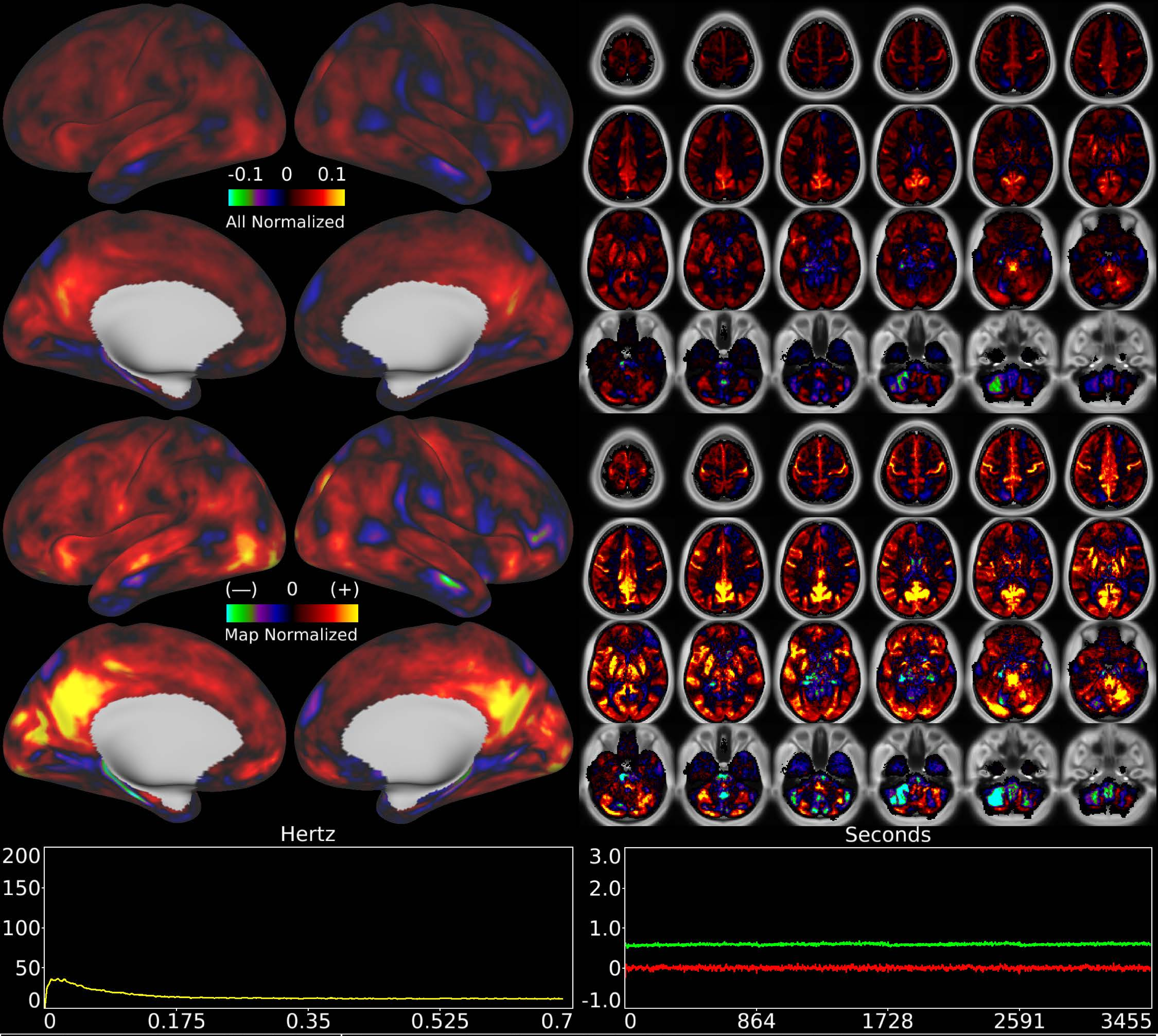


| | | |
|---|---|---|
| Number & Class: 61 Noise | Name: Subsidiary Global Physiological Noise | |
| RVT Correlated: No | DVARS Dip Associated: No | Cross-Subject Variable: Yes |
| Single Subject: Yes | % Variance Explained: 0.54 | Globality Index: 1.41 |
| Task Component: No | Taskr Component: No | Sleep Noted: No |
| Rationale: Single subject component with high correlation to global timecourse | | |

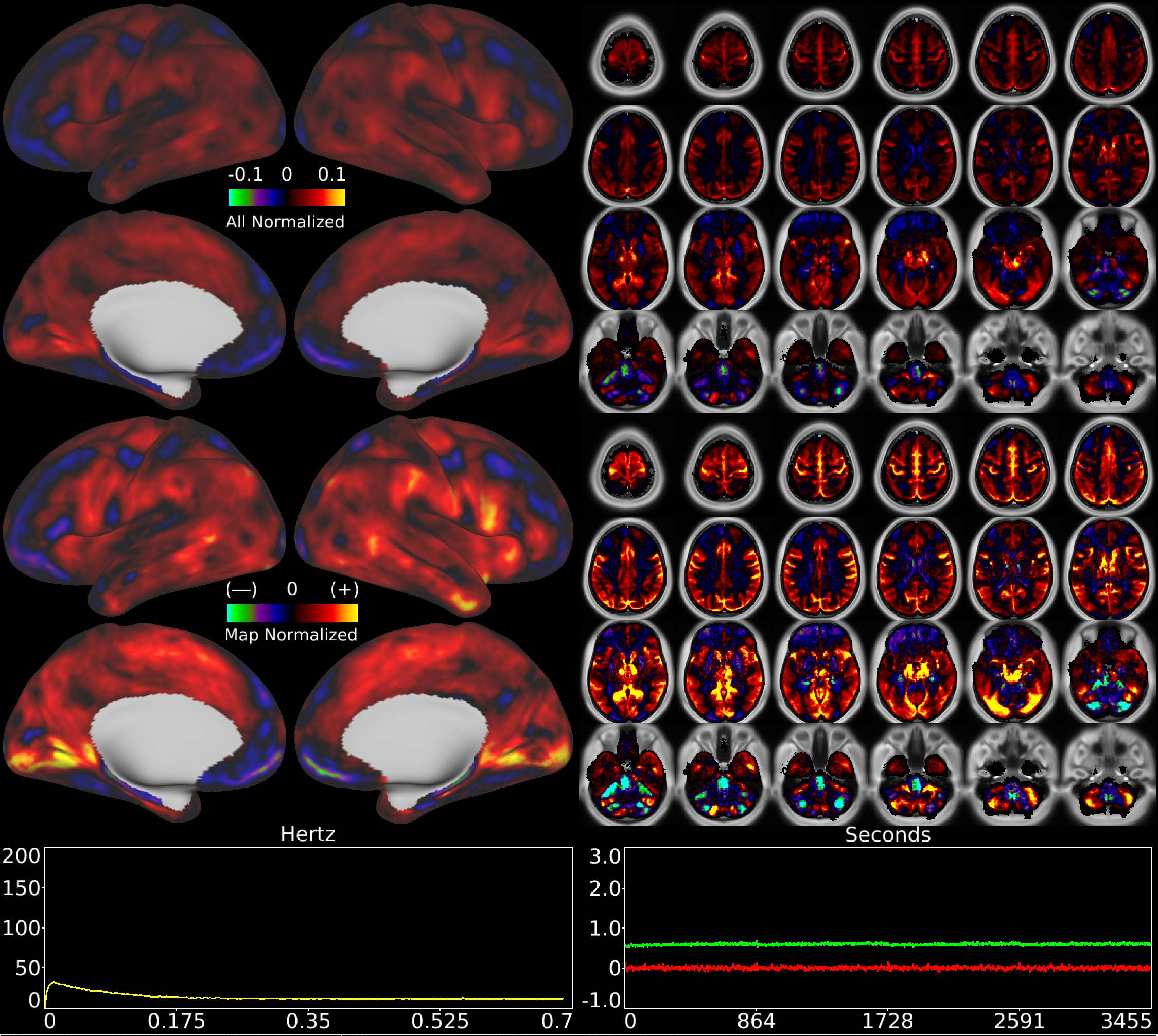


| | | |
|---|---|---|
| Number & Class: 62 Noise | Name: Subsidiary Global Physiological Noise | |
| RVT Correlated: No | DVARS Dip Associated: No | Cross-Subject Variable: No |
| Single Subject: Yes | % Variance Explained: 0.54 | Globality Index: 1.41 |
| Task Component: No | Taskr Component: No | Sleep Noted: No |
| Rationale: Single subject component with high correlation to global timecourse | | |

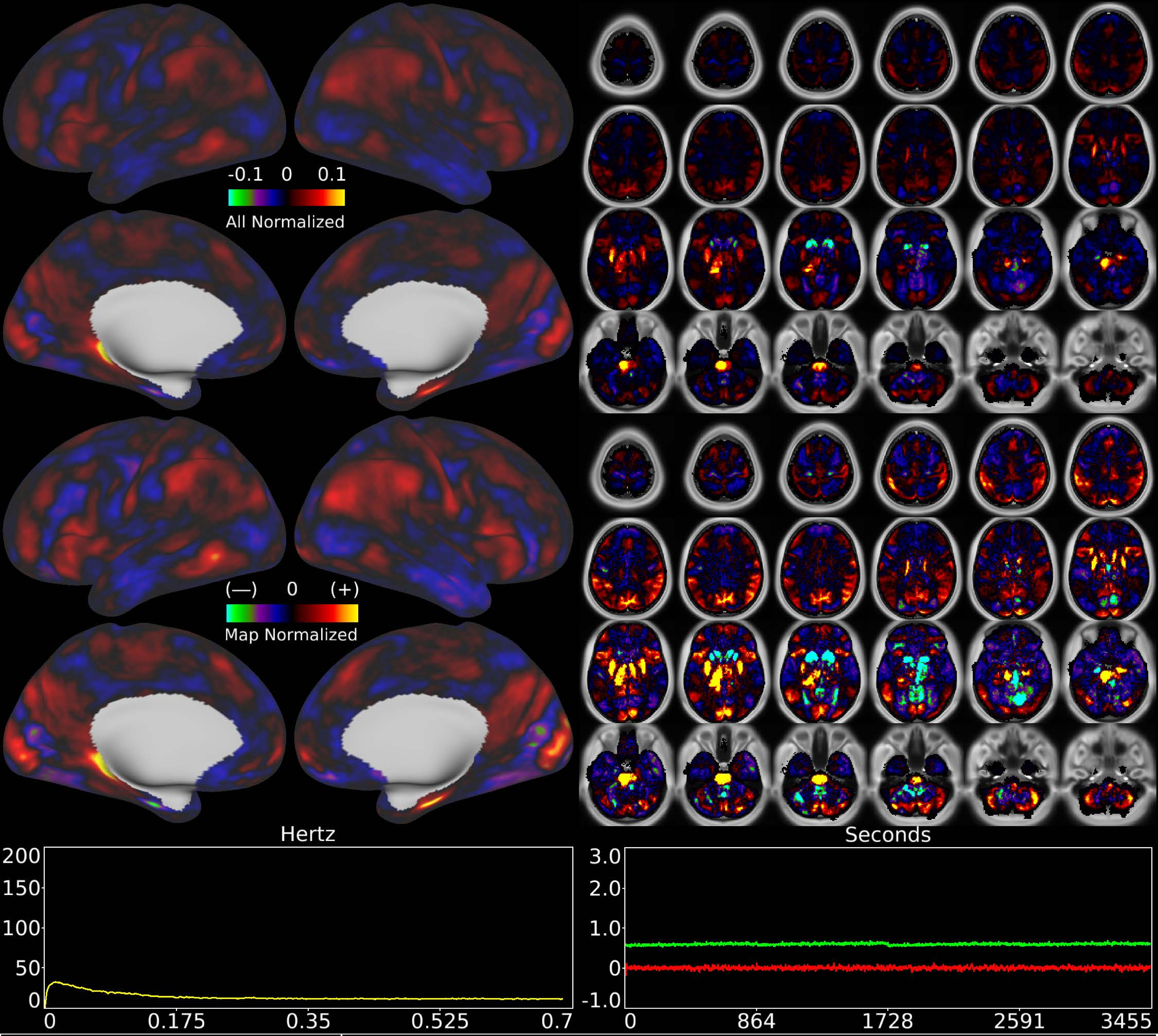


| | | |
|---|---|---|
| Number & Class: 63 Noise | Name: Pons + L Thalamus > Amygdala | |
| RVT Correlated: No | DVARS Dip Associated: No | Cross-Subject Variable: No |
| Single Subject: Yes | % Variance Explained: 0.54 | Globality Index: 0.01 |
| Task Component: No | Taskr Component: No | Sleep Noted: No |
| Rationale: Controversial: Single subject component with spatial map dominated by subcortical patches | | |

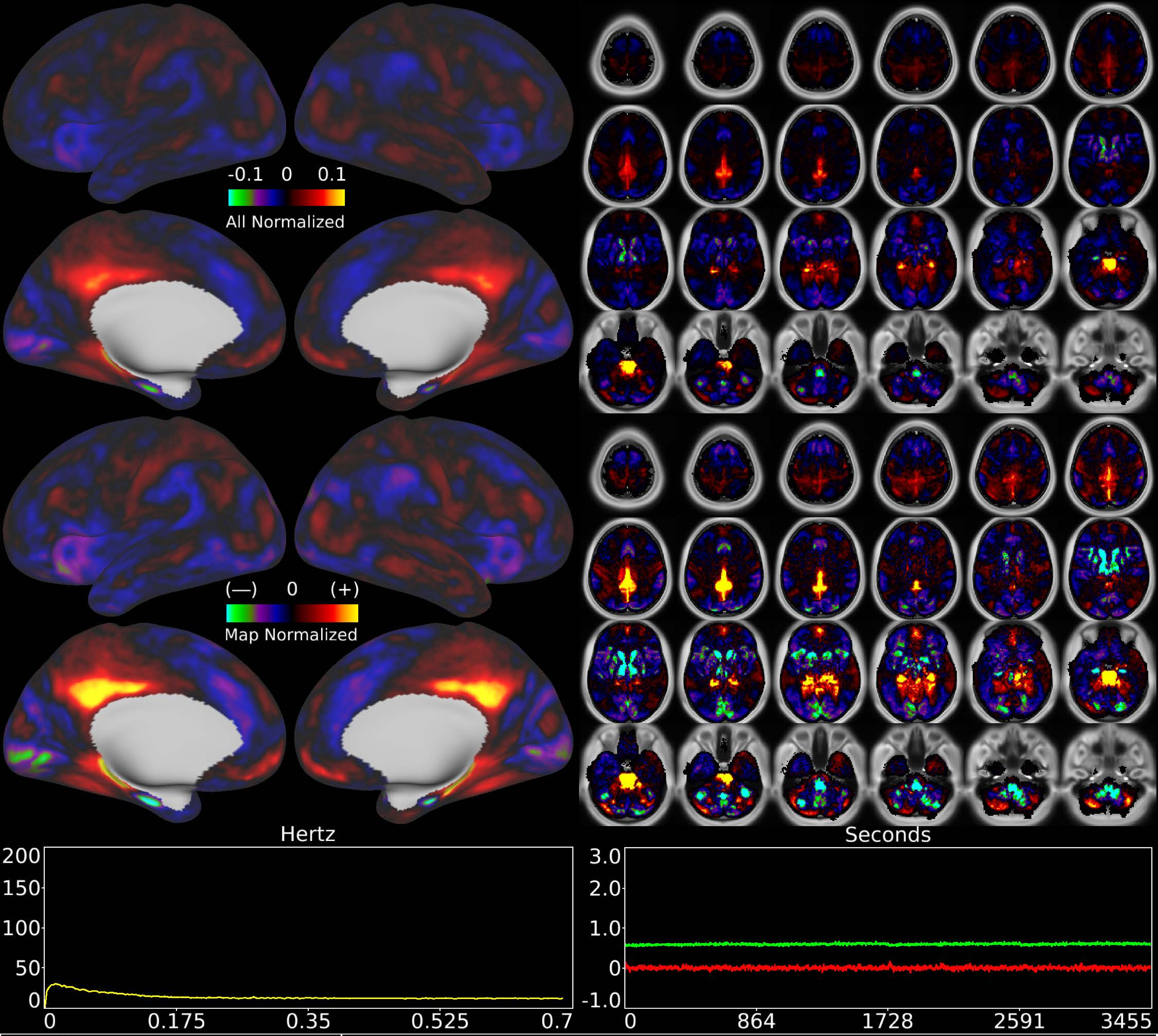


| Number & Class: 64 Noise | Name: Recon Artifact? | |
|---|---|---|
| RVT Correlated: No | DVARS Dip Associated: No | Cross-Subject Variable: No |
| Single Subject: Yes | % Variance Explained: 0.54 | Globality Index: 0.42 |
| Task Component: No | Taskr Component: No | Sleep Noted: No |
| Rationale: Single subject component with banding in sagittal plane; potentially a multi-band recon artifact | | |

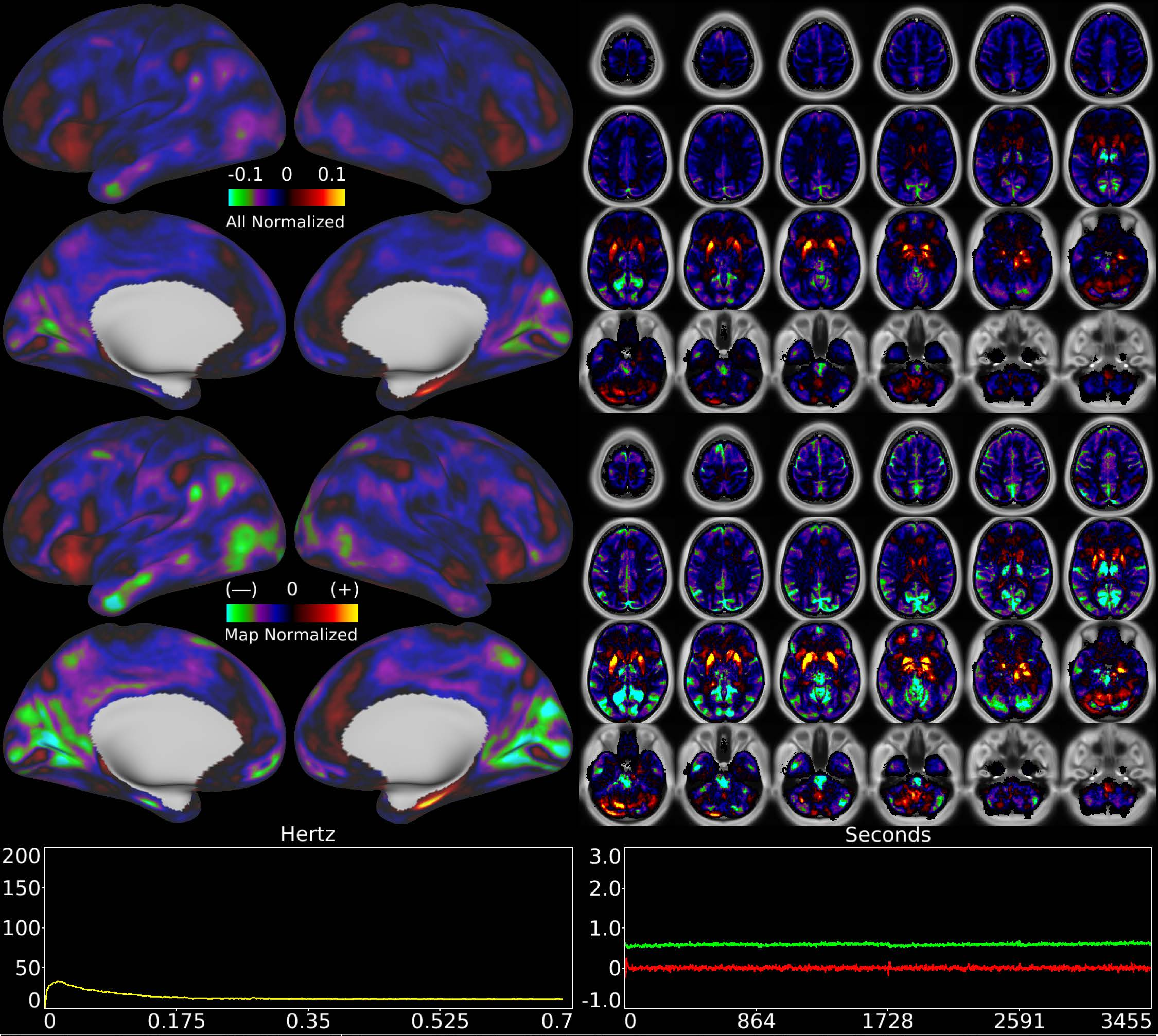


| | | |
|---|---|---|
| Number & Class: 65 Signal | Name: Highly Globally Correlated Single Subject: Sleep? | |
| RVT Correlated: No | DVARS Dip Associated: No | Cross-Subject Variable: No |
| Single Subject: Yes | % Variance Explained: 0.54 | Globality Index: 1.57 |
| Task Component: No | Taskr Component: No | Sleep Noted: No |
| Rationale: Controversial: Single subject component with high correlation to global timecourse; looked more like RC1 than global noise components) | | |

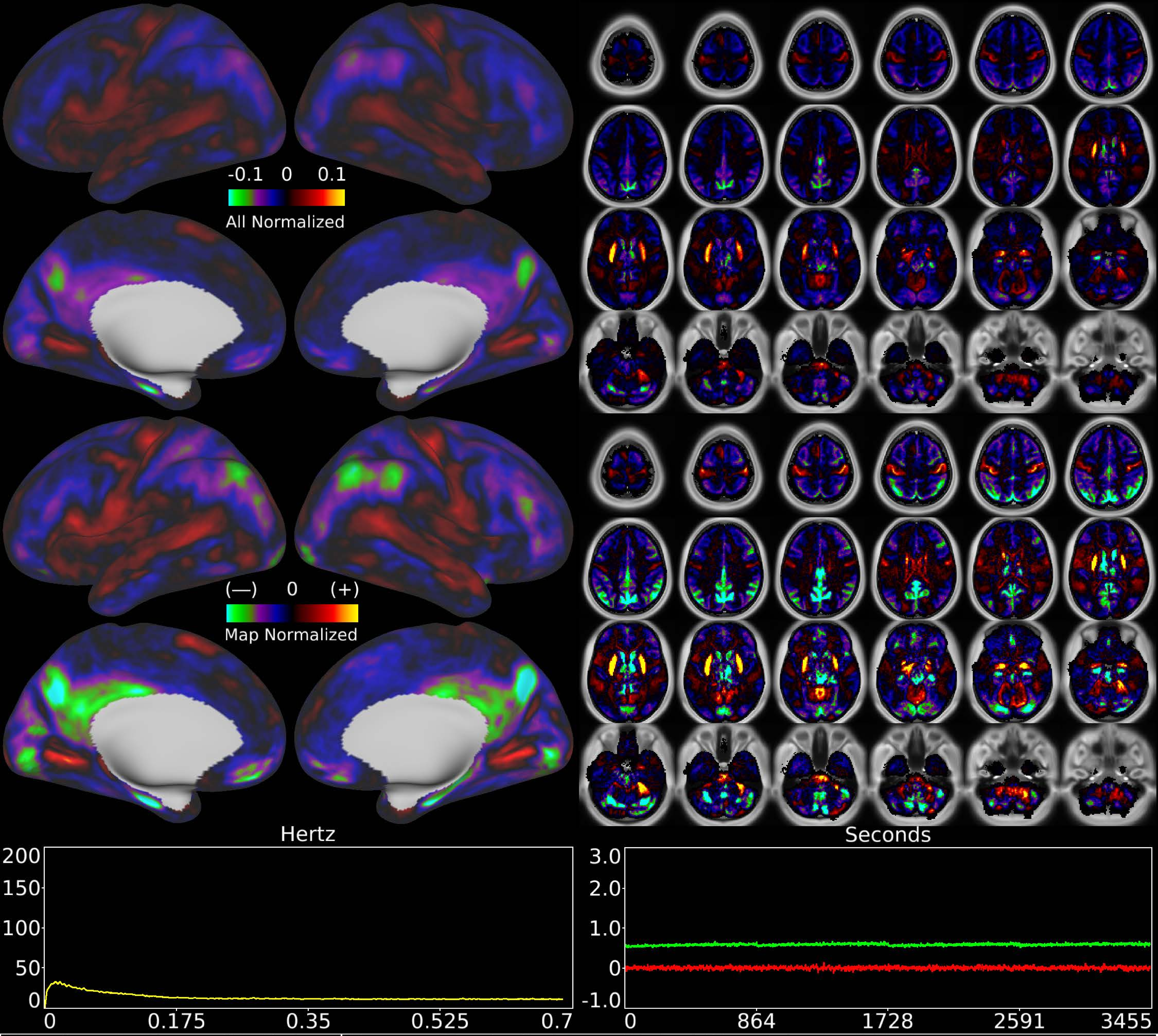


| | | |
|---|---|---|
| Number & Class: 66 Noise | Name: Single Subject Global Physiological Noise | |
| RVT Correlated: No | DVARS Dip Associated: No | Cross-Subject Variable: No |
| Single Subject: Yes | % Variance Explained: 0.53 | Globality Index: 1.07 |
| Task Component: No | Taskr Component: No | Sleep Noted: No |
| Rationale: Single subject component with high correlation to global timecourse | | |

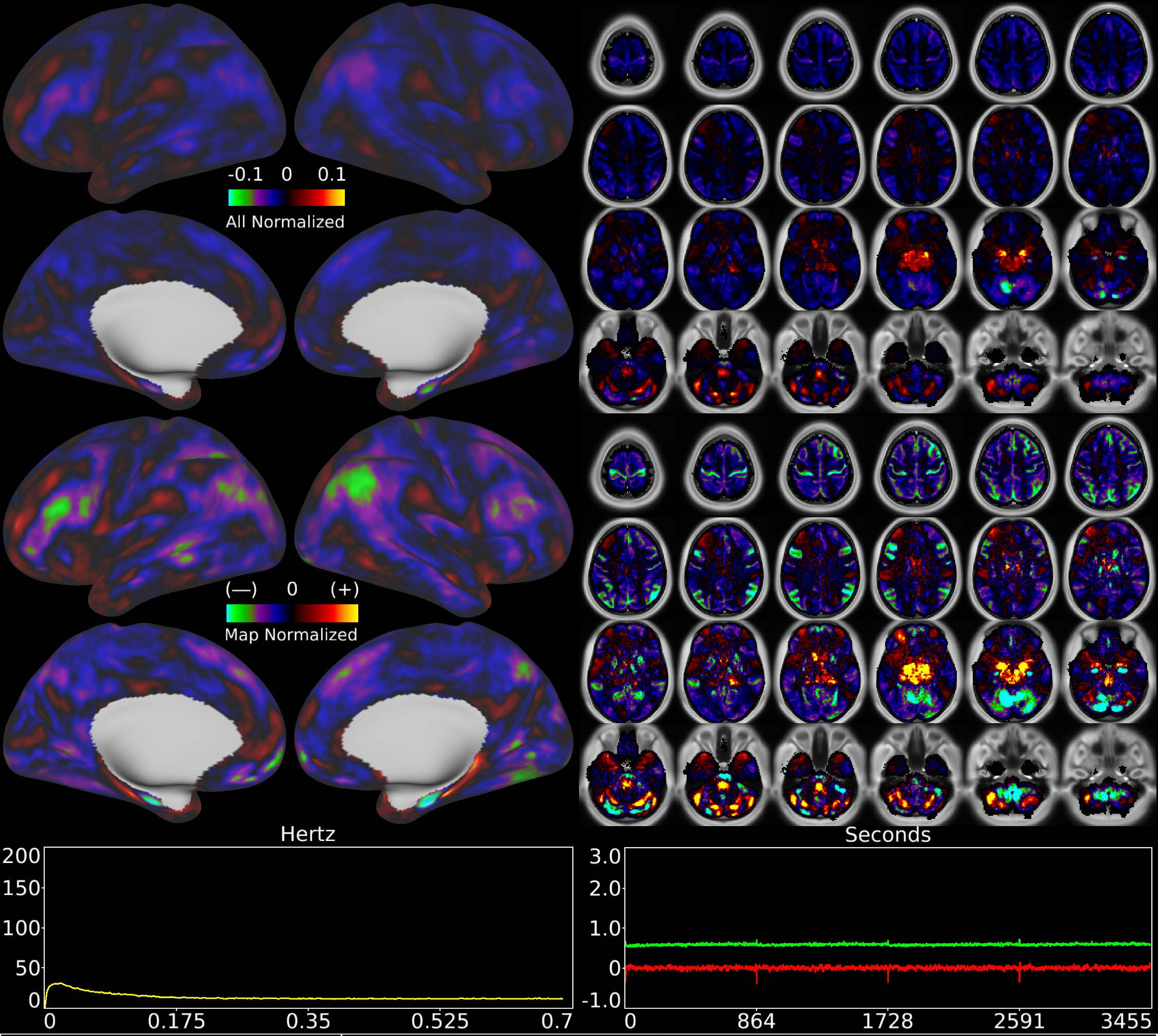


| | | |
|---|---|---|
| Number & Class: 67 Noise | Name: Single Subject Global Physiological Noise | |
| RVT Correlated: No | DVARS Dip Associated: No | Cross-Subject Variable: No |
| Single Subject: Yes | % Variance Explained: 0.52 | Globality Index: 1.3 |
| Task Component: No | Taskr Component: No | Sleep Noted: No |
| Rationale: Single subject component with high correlation to global timecourse | | |

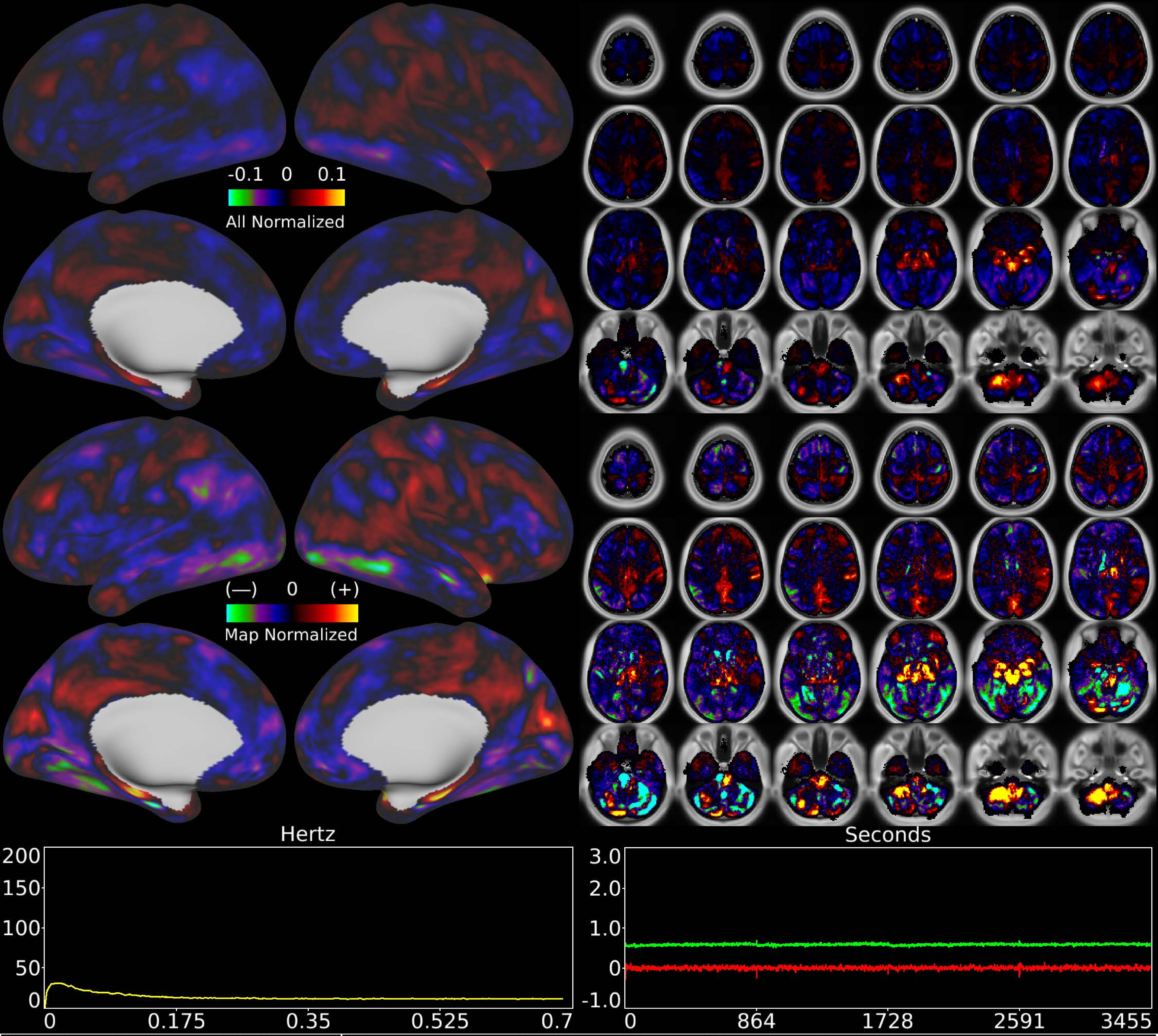


| | | |
|---|---|---|
| Number & Class: 68 Noise | Name: Single Subject Global Physiological Noise | |
| RVT Correlated: No | DVARS Dip Associated: No | Cross-Subject Variable: No |
| Single Subject: Yes | % Variance Explained: 0.52 | Globality Index: 0.62 |
| Task Component: No | Taskr Component: No | Sleep Noted: No |
| Rationale: Single subject component with high correlation to global timecourse | | |

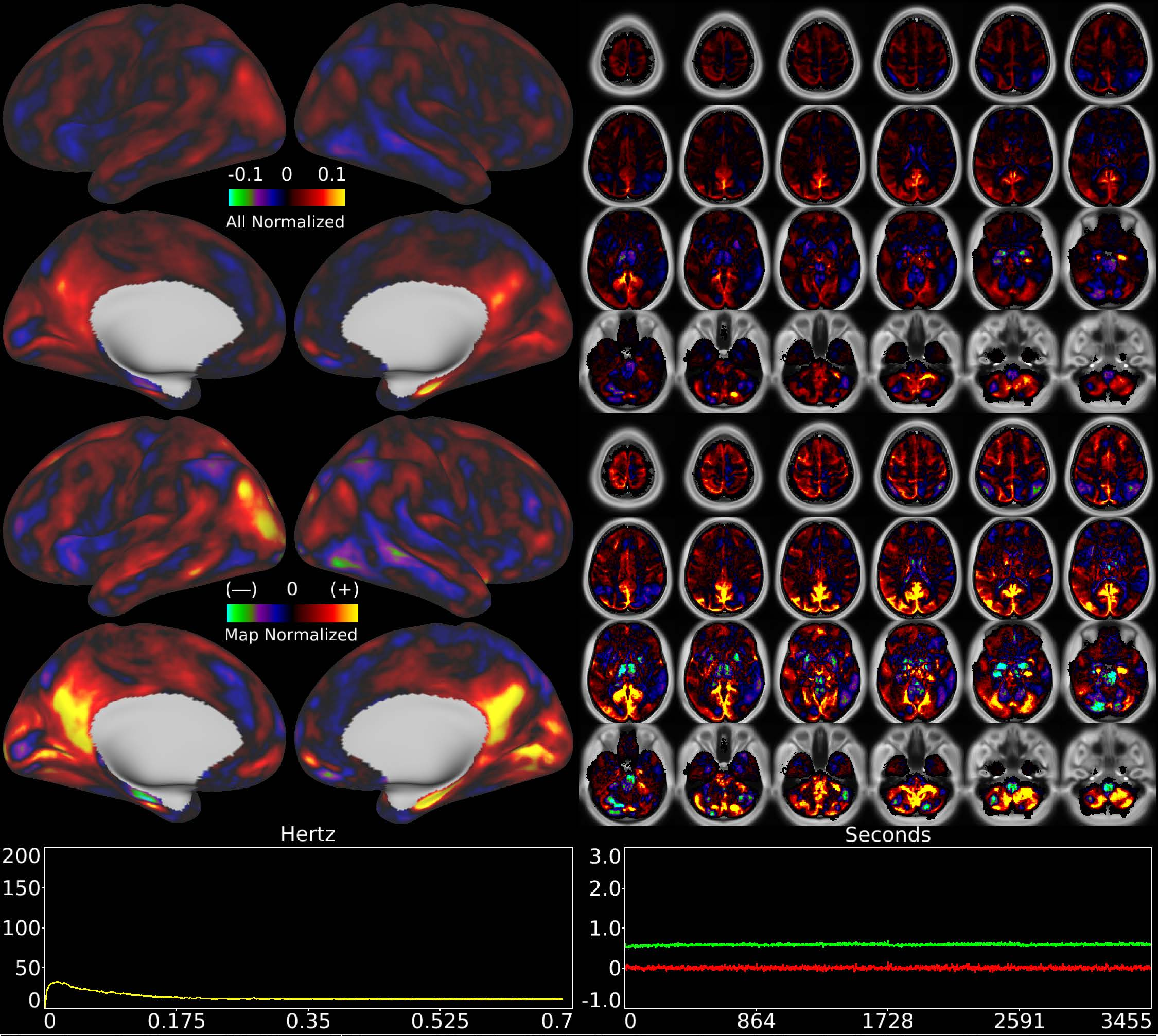


| | | |
|---|---|---|
| Number & Class: 69 Signal | Name: Highly Globally Correlated Single Subject: Sleep? | |
| RVT Correlated: No | DVARS Dip Associated: No | Cross-Subject Variable: No |
| Single Subject: Yes | % Variance Explained: 0.52 | Globality Index: 0.86 |
| Task Component: No | Taskr Component: No | Sleep Noted: No |
| Rationale: Controversial: Single subject component with high correlation to global timecourse; looked more like RC1 than global noise components) | | |

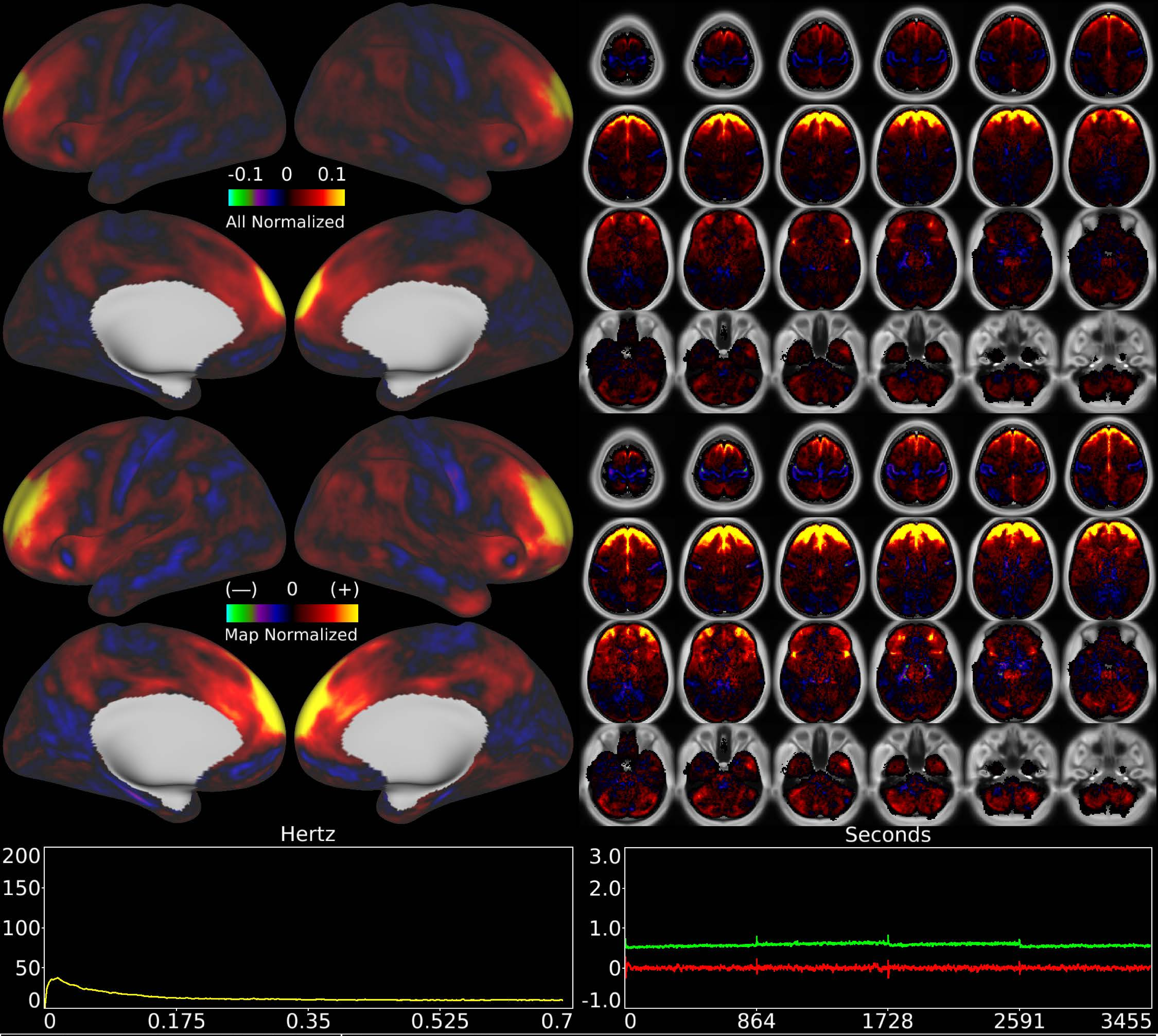


| | | |
|---|---|---|
| Number & Class: 70 Noise | Name: Coil? | |
| RVT Correlated: No | DVARS Dip Associated: Yes | Cross-Subject Variable: No |
| Single Subject: No | % Variance Explained: 0.52 | Globality Index: 1.42 |
| Task Component: 64 | Taskr Component: 55 | Sleep Noted: No |
| Rationale: Spatial map is a band across frontal cortex and white matter; could be related to coil; head motion; or eye motion | | |

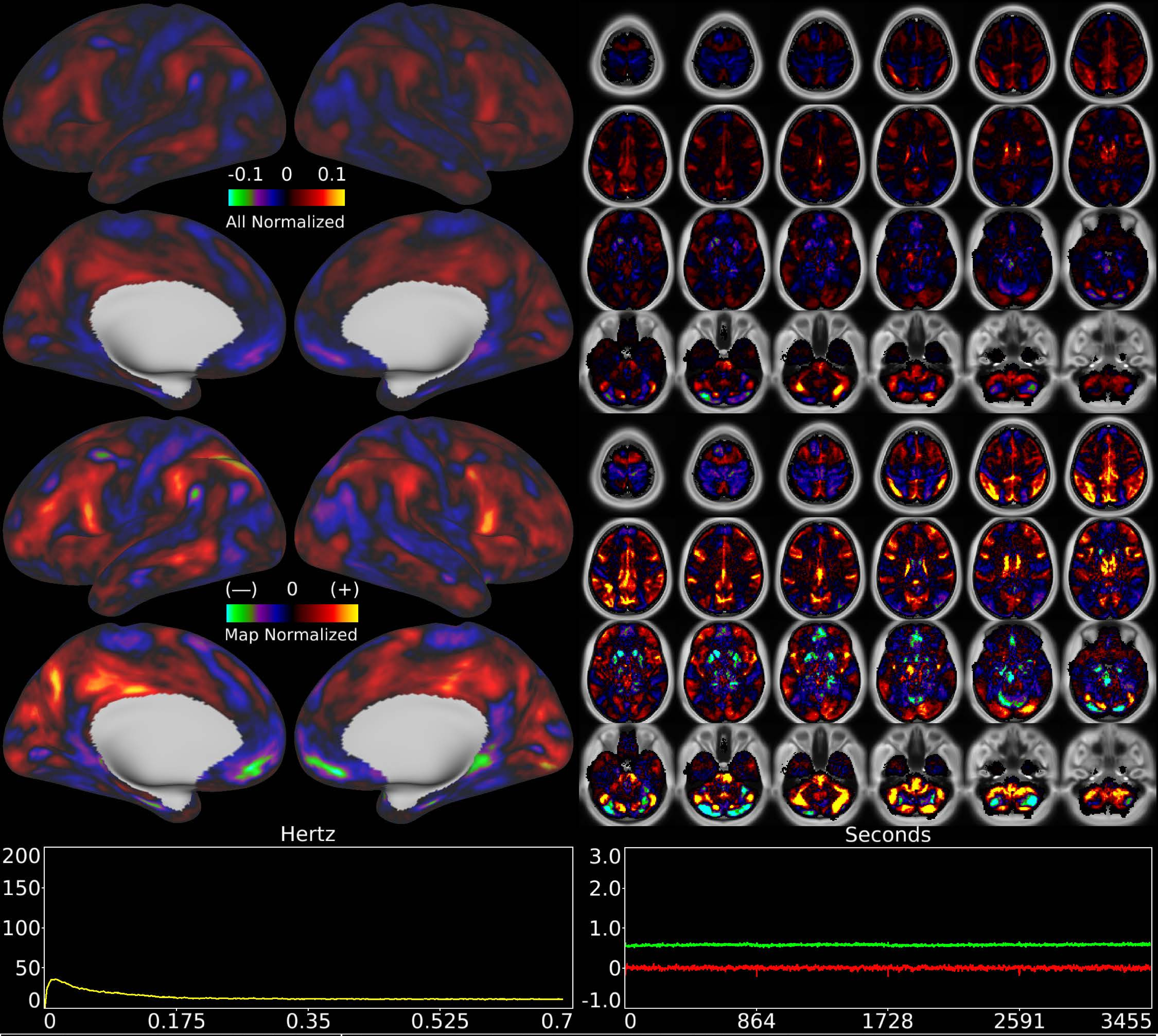


| | | |
|---|---|---|
| Number & Class: 71 Noise | Name: Single Subject Global Physiological Noise | |
| RVT Correlated: No | DVARS Dip Associated: No | Cross-Subject Variable: No |
| Single Subject: Yes | % Variance Explained: 0.51 | Globality Index: 0.32 |
| Task Component: No | Taskr Component: No | Sleep Noted: No |
| Rationale: Single subject component with high correlation to global timecourse | | |

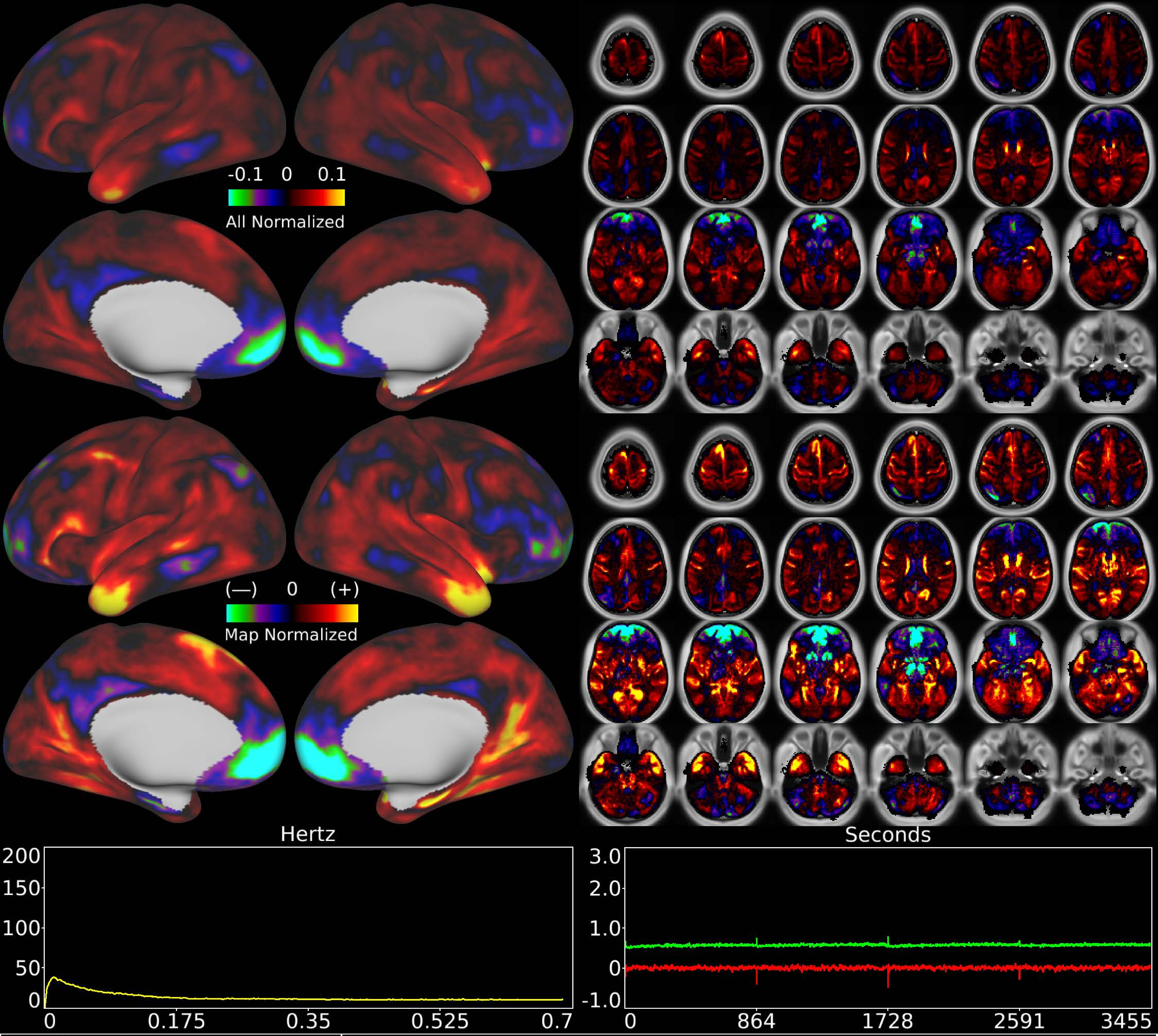


| | | |
|---|---|---|
| Number & Class: 72 Noise | Name: Coil? | |
| RVT Correlated: No | DVARS Dip Associated: No | Cross-Subject Variable: No |
| Single Subject: Yes | % Variance Explained: 0.5 | Globality Index: 1.36 |
| Task Component: No | Taskr Component: No | Sleep Noted: No |

Rationale: Single subject component with high correlation to global timecourse

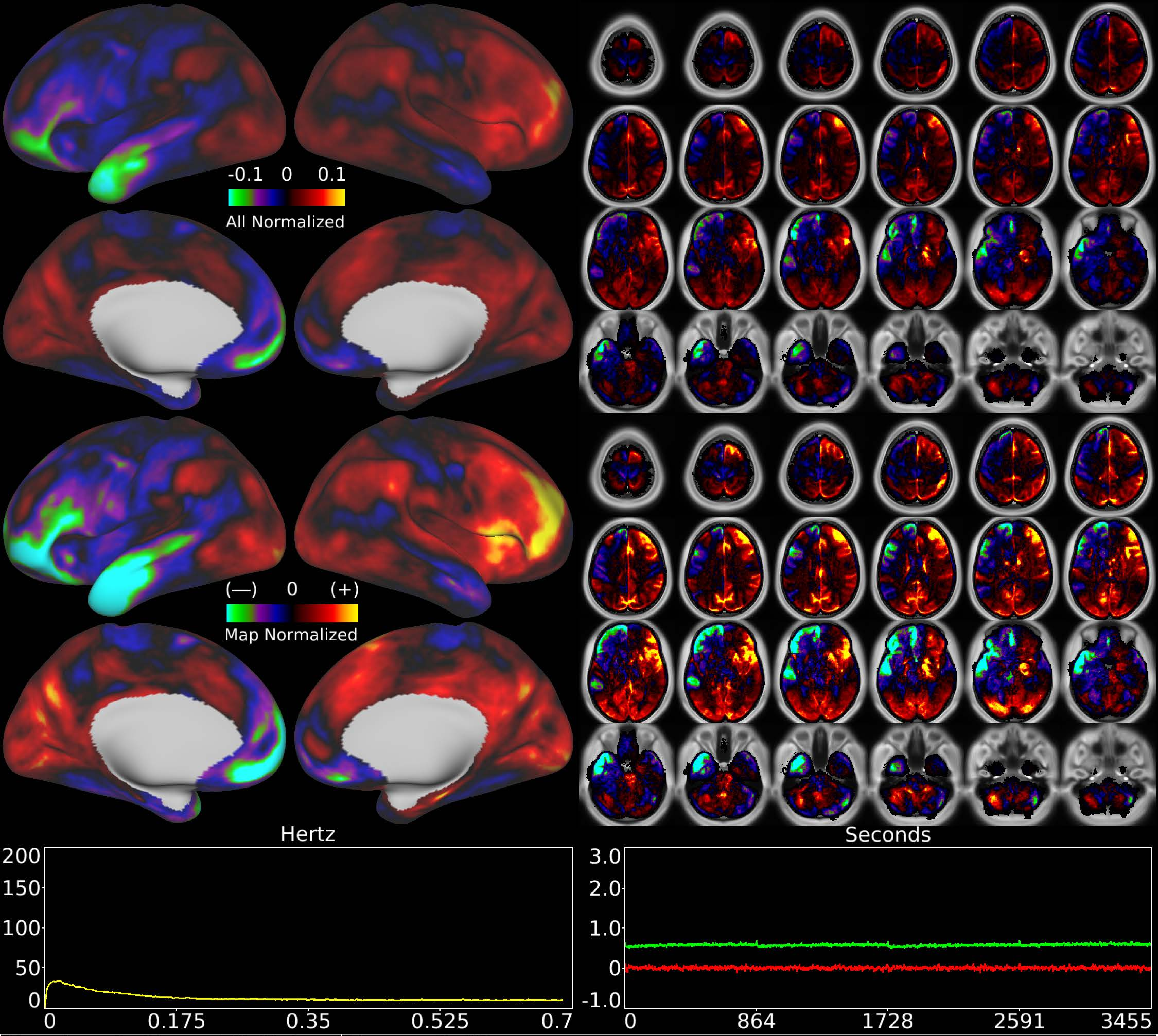


| | | |
|---|---|---|
| Number & Class: 73 Noise | Name: Coil? | |
| RVT Correlated: No | DVARS Dip Associated: Yes | Cross-Subject Variable: No |
| Single Subject: Yes | % Variance Explained: 0.5 | Globality Index: 0.66 |
| Task Component: No | Taskr Component: No | Sleep Noted: No |

Rationale: Single subject component with correlation to DVARS dips; potentially coil related or movement related

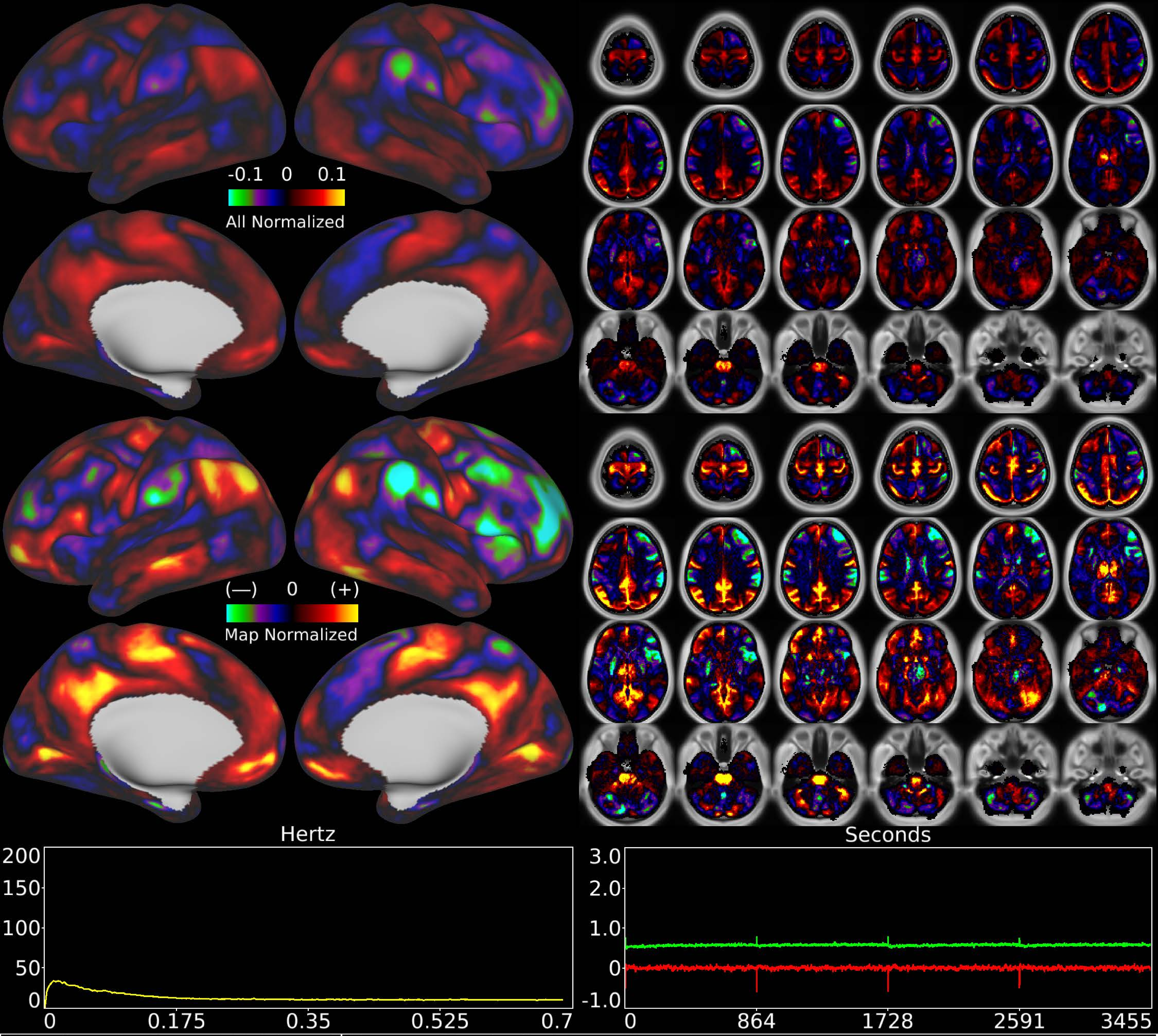


| | | |
|---|---|---|
| Number & Class: 74 Signal | Name: Single Subject Signal | |
| RVT Correlated: No | DVARS Dip Associated: No | Cross-Subject Variable: No |
| Single Subject: Yes | % Variance Explained: 0.5 | Globality Index: 0.12 |
| Task Component: No | Taskr Component: No | Sleep Noted: No |
| Rationale: Controversial: Single Subject Component with spatial map that includes positive and negative patches that respect known RSNs (e.g. Default Mode Network) | | |

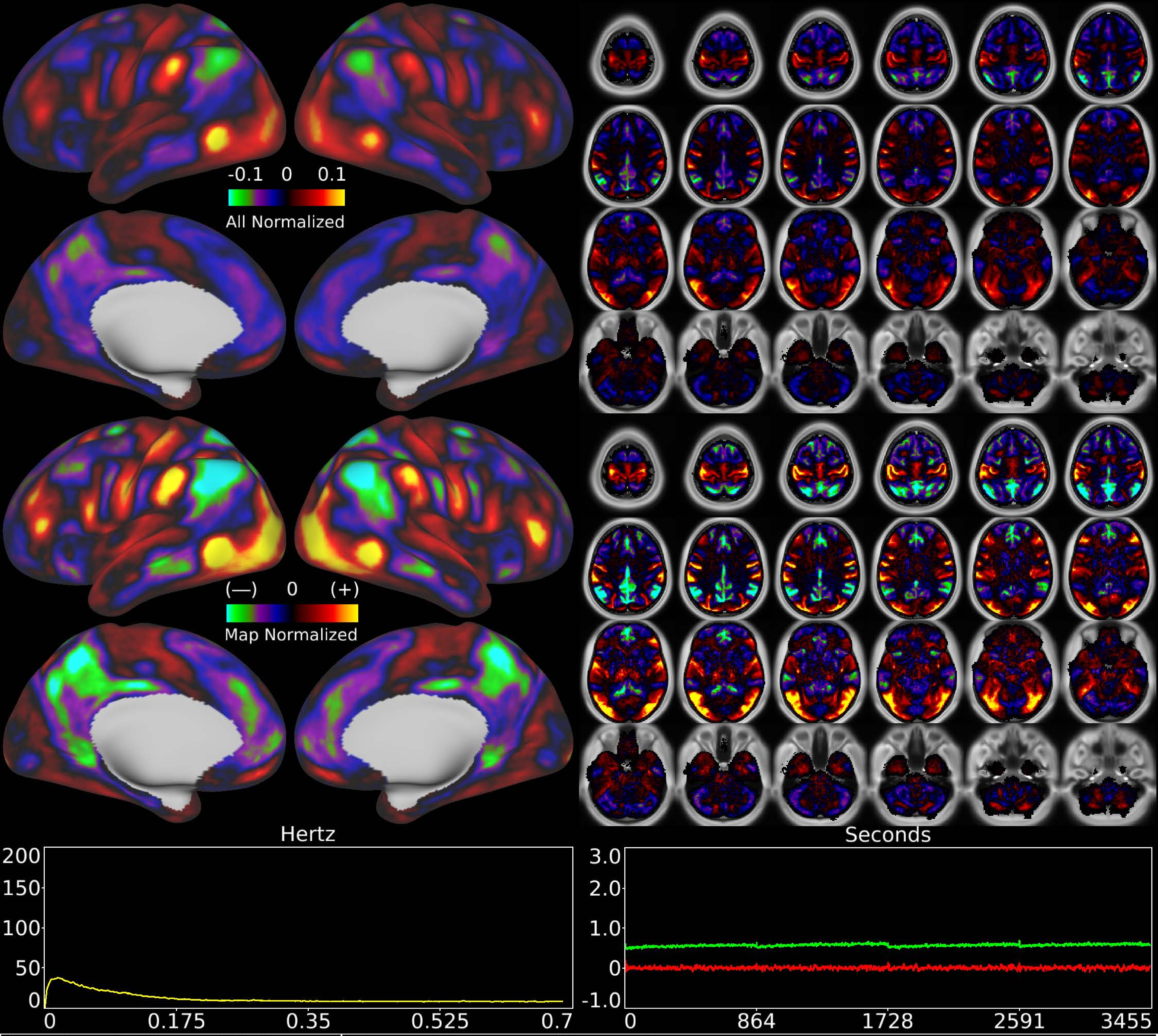


| | | |
|---|---|---|
| Number & Class: 75 Signal | Name: Unknown Network | |
| RVT Correlated: No | DVARS Dip Associated: No | Cross-Subject Variable: No |
| Single Subject: No | % Variance Explained: 0.5 | Globality Index: 0.18 |
| Task Component: No | Taskr Component: No | Sleep Noted: No |

Rationale: Spatial map includes positive and negative patches that respect known areal boundaries (e.g. PF and PFm)

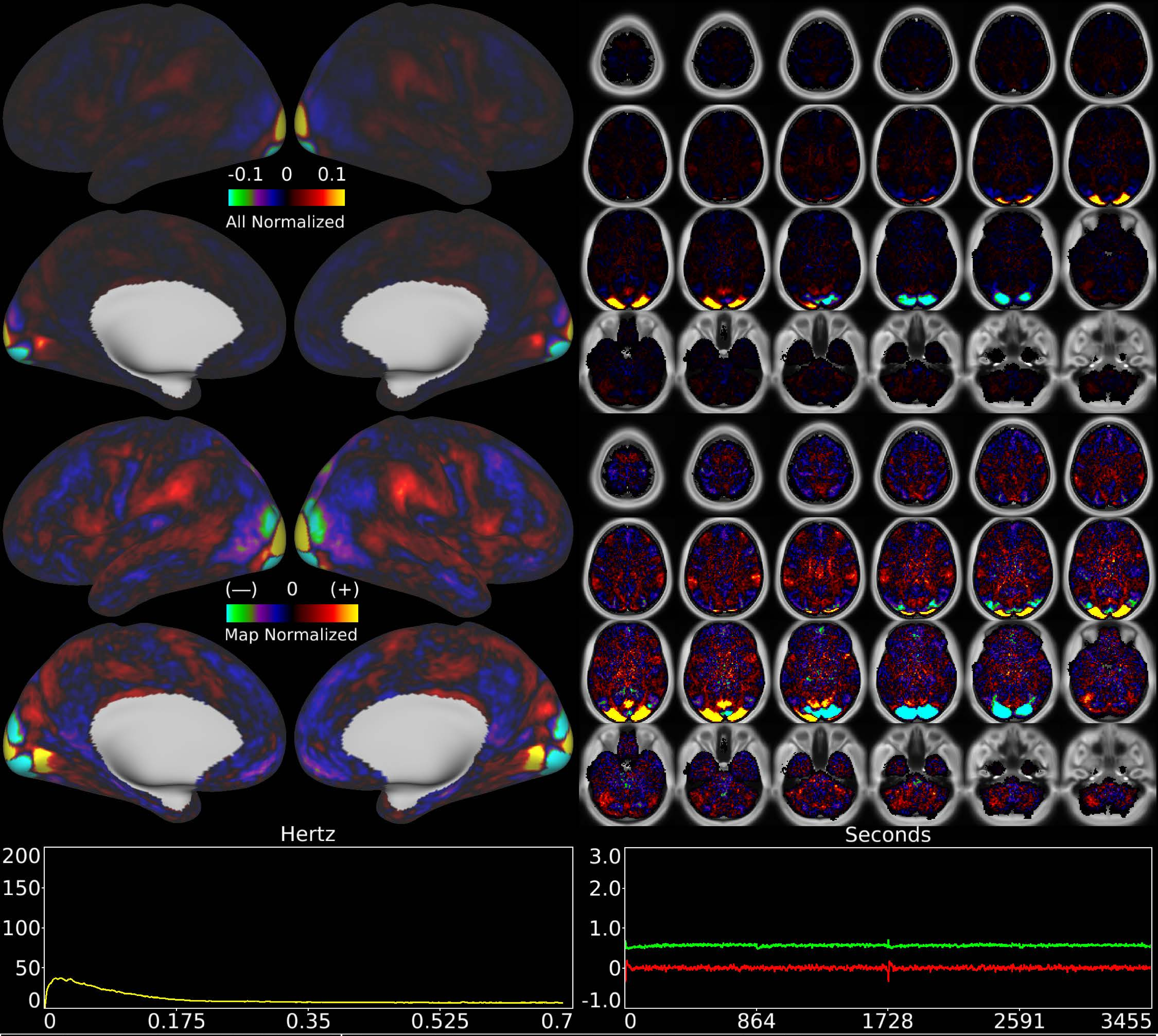


| | | |
|---|---|---|
| Number & Class: 76 Signal | Name: Visuotopic: Foveal Dorsal > Ventral | |
| RVT Correlated: No | DVARS Dip Associated: No | Cross-Subject Variable: No |
| Single Subject: No | % Variance Explained: 0.5 | Globality Index: 0.04 |
| Task Component: 65 | Taskr Component: 53 | Sleep Noted: No |
| Rationale: Spatial map includes positive and negative patches that respect known retinotopic visual organization (Foveal and Lower vs Upper) | | |

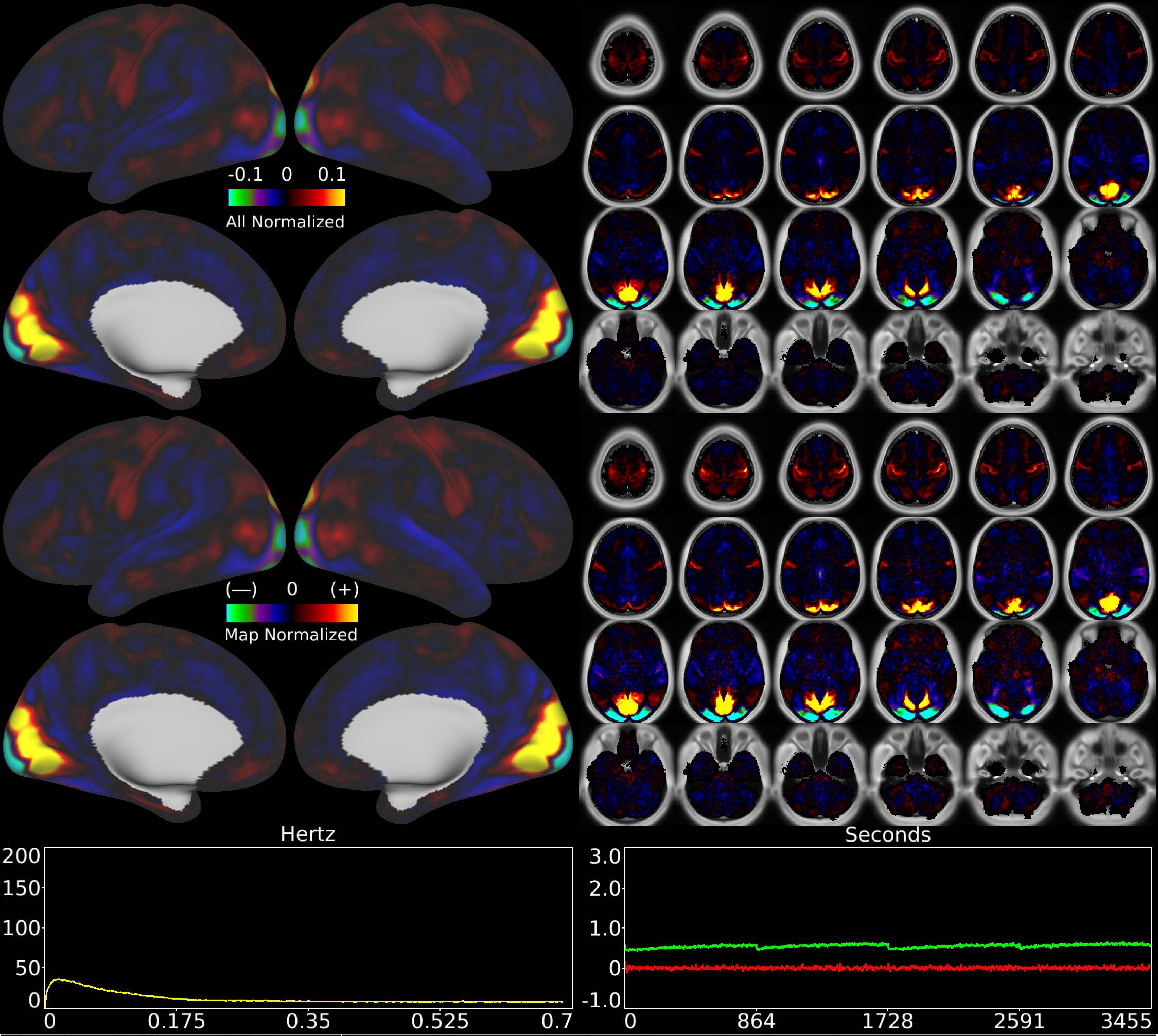


| | | |
|---|---|---|
| Number & Class: 77 Signal | Name: Visuotopic: ParaCentral > Foveal | |
| RVT Correlated: No | DVARS Dip Associated: No | Cross-Subject Variable: No |
| Single Subject: No | % Variance Explained: 0.48 | Globality Index: 0.52 |
| Task Component: No | Taskr Component: No | Sleep Noted: No |
| Rationale: Spatial map includes positive and negative patches that respect known retinotopic visual organization (Paracentral vs Foveal) | | |

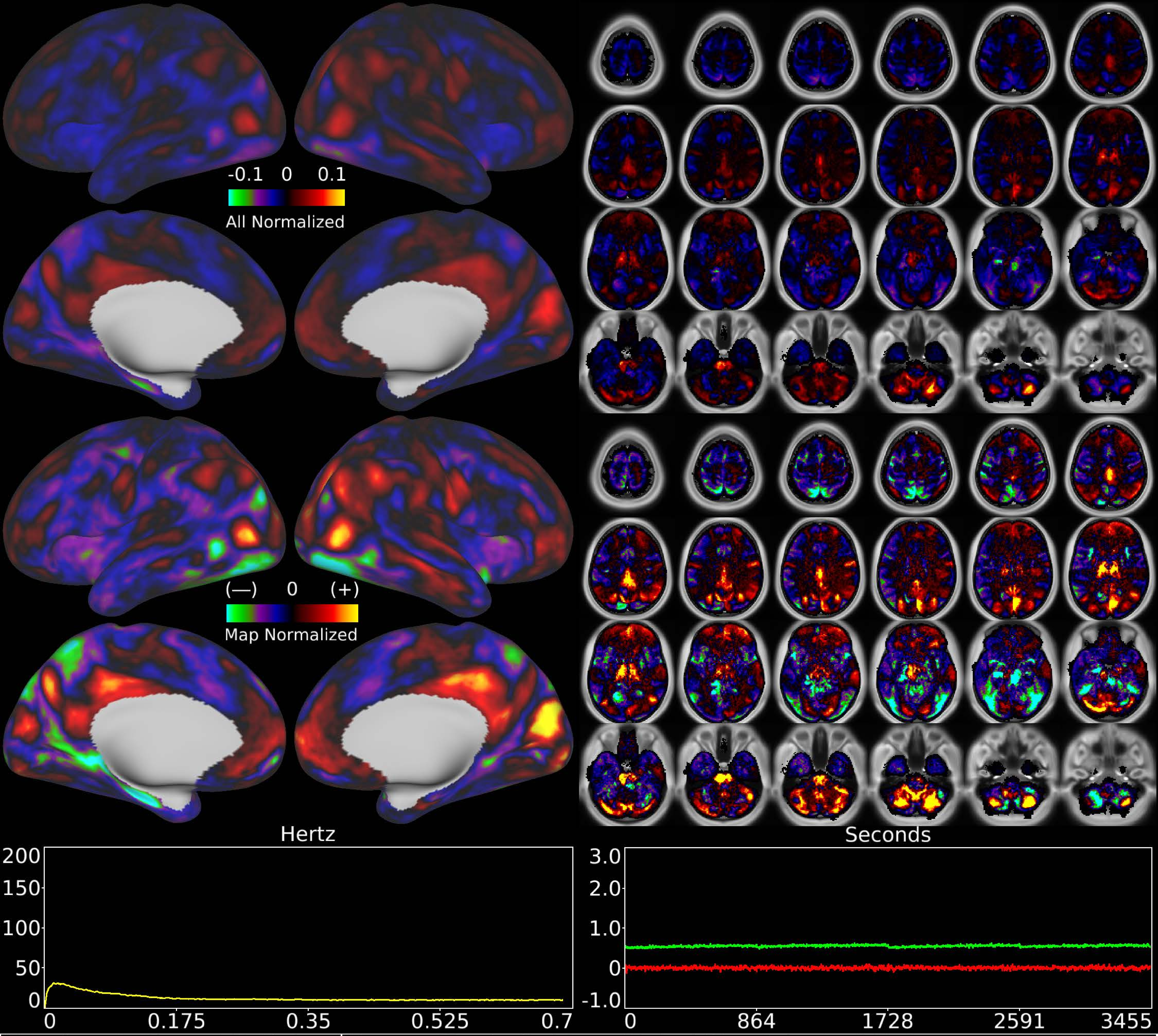


| | | |
|---|---|---|
| Number & Class: 78 Noise | Name: Single Subject Global Physiological Noise | |
| RVT Correlated: No | DVARS Dip Associated: No | Cross-Subject Variable: No |
| Single Subject: Yes | % Variance Explained: 0.46 | Globality Index: 0.47 |
| Task Component: No | Taskr Component: No | Sleep Noted: No |
| Rationale: Single subject component with high correlation to global timecourse | | |

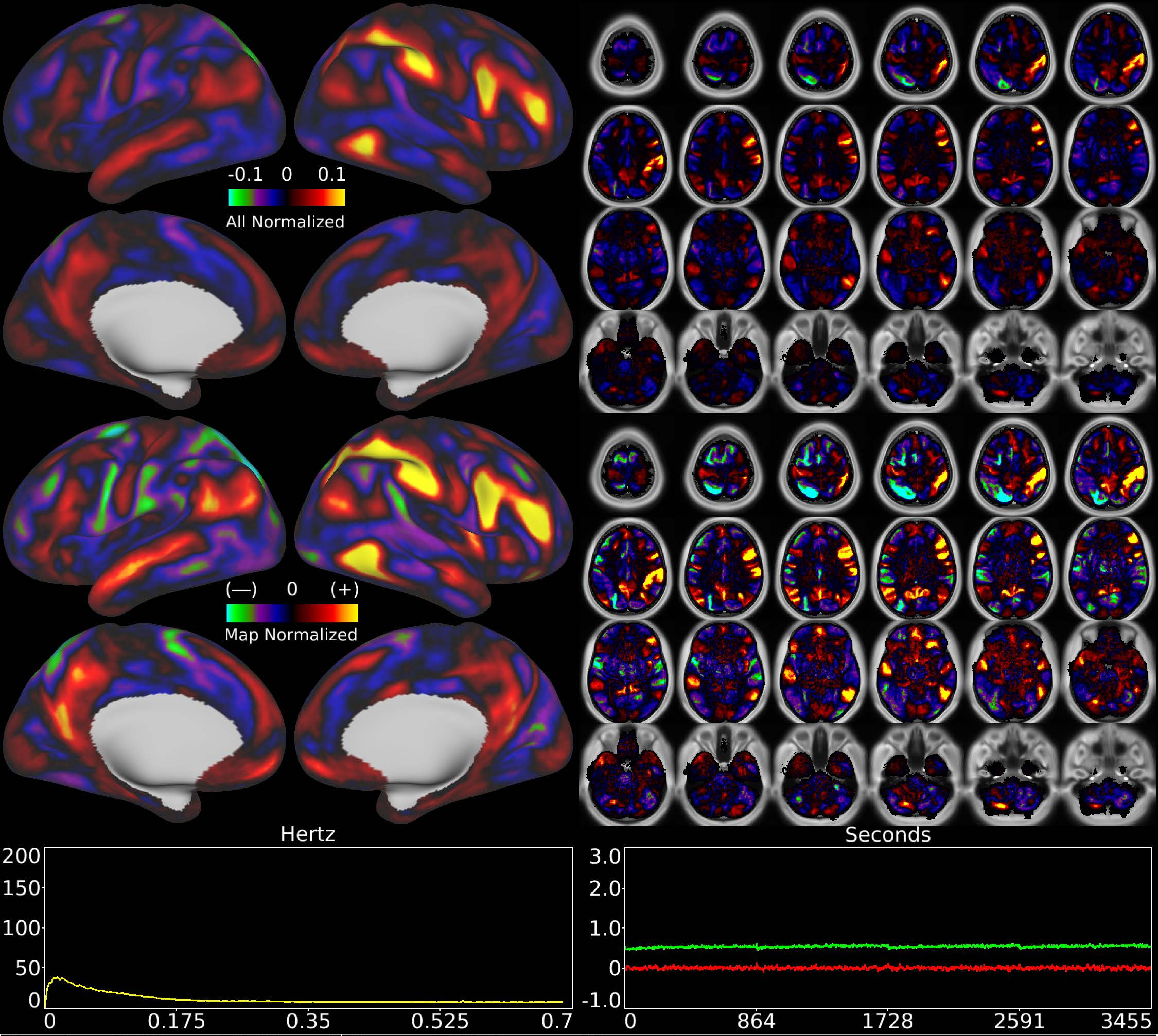


| | | |
|---|---|---|
| Number & Class: 79 Signal | Name: R Unknown Network | |
| RVT Correlated: No | DVARS Dip Associated: No | Cross-Subject Variable: No |
| Single Subject: No | % Variance Explained: 0.44 | Globality Index: 0.33 |
| Task Component: No | Taskr Component: No | Sleep Noted: No |
| Rationale: Spatial map includes positive and negative patches that respect known RSNs (e.g. Dorsal Attention Network) | | |

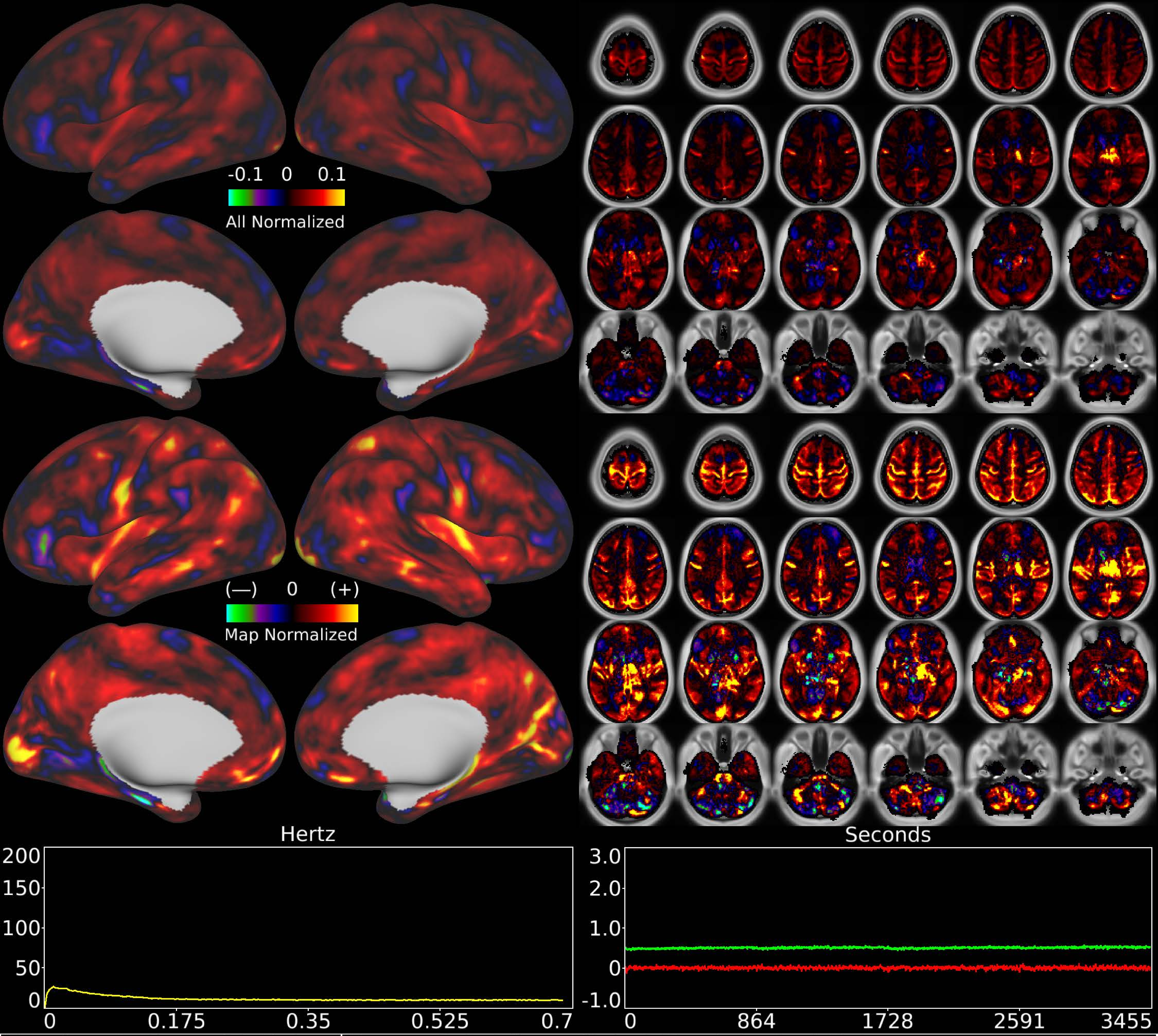


| | | |
|---|---|---|
| Number & Class: 80 Noise | Name: Subsidiary Global Physiological Noise | |
| RVT Correlated: No | DVARS Dip Associated: No | Cross-Subject Variable: Yes |
| Single Subject: Yes | % Variance Explained: 0.42 | Globality Index: 1.52 |
| Task Component: No | Taskr Component: No | Sleep Noted: Yes |
| Rationale: Single subject component with high correlation to global timecourse | | |

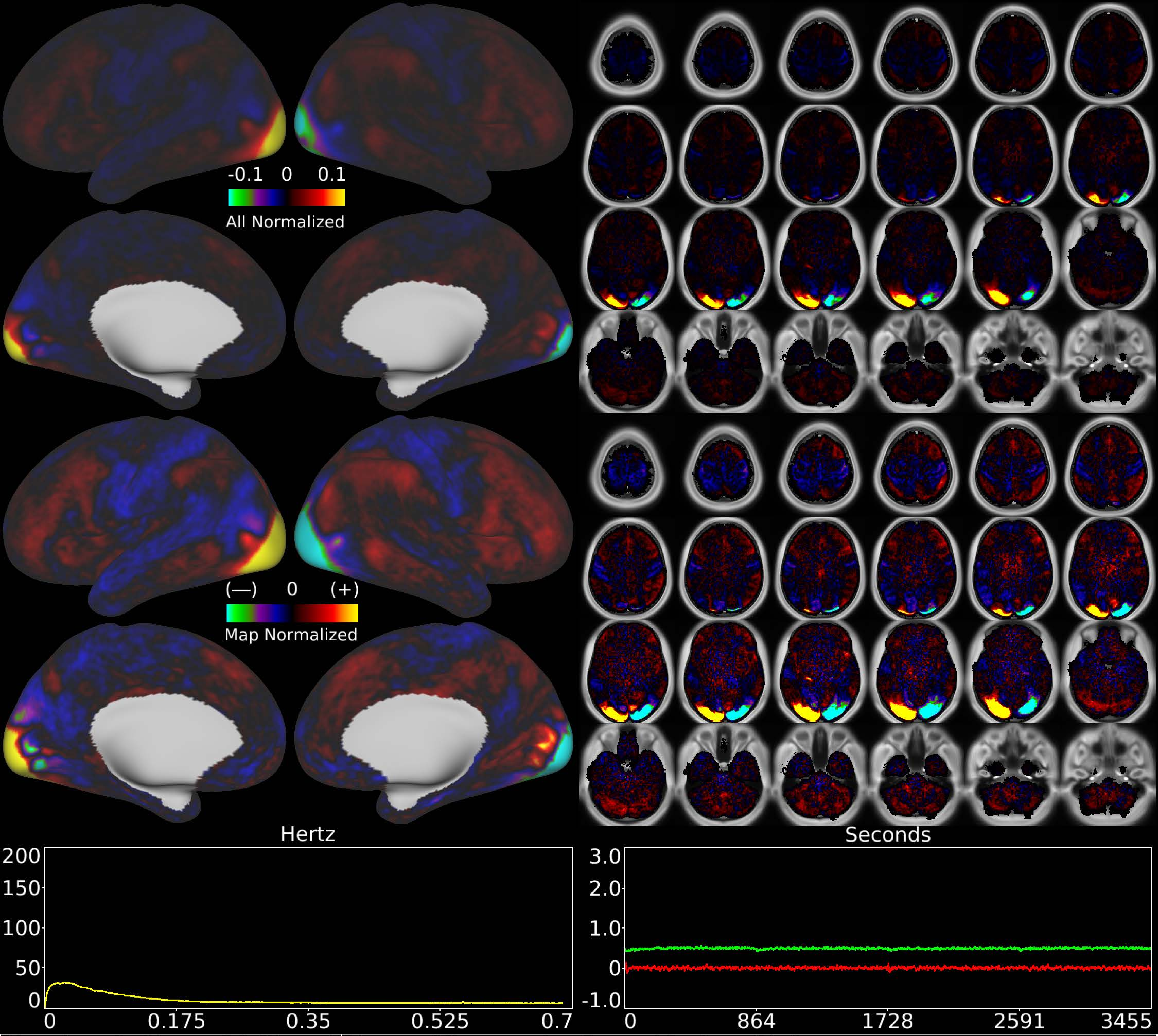


| | | |
|---|---|---|
| Number & Class: 81 Signal | Name: Visuotopic: Foveal Left > Right | |
| RVT Correlated: No | DVARS Dip Associated: No | Cross-Subject Variable: No |
| Single Subject: No | % Variance Explained: 0.37 | Globality Index: 0.29 |
| Task Component: 70 | Taskr Component: 58 | Sleep Noted: No |
| Rationale: Spatial map includes positive and negative patches that respect known retinotopic visual organization (Foveal Right vs Left) | | |

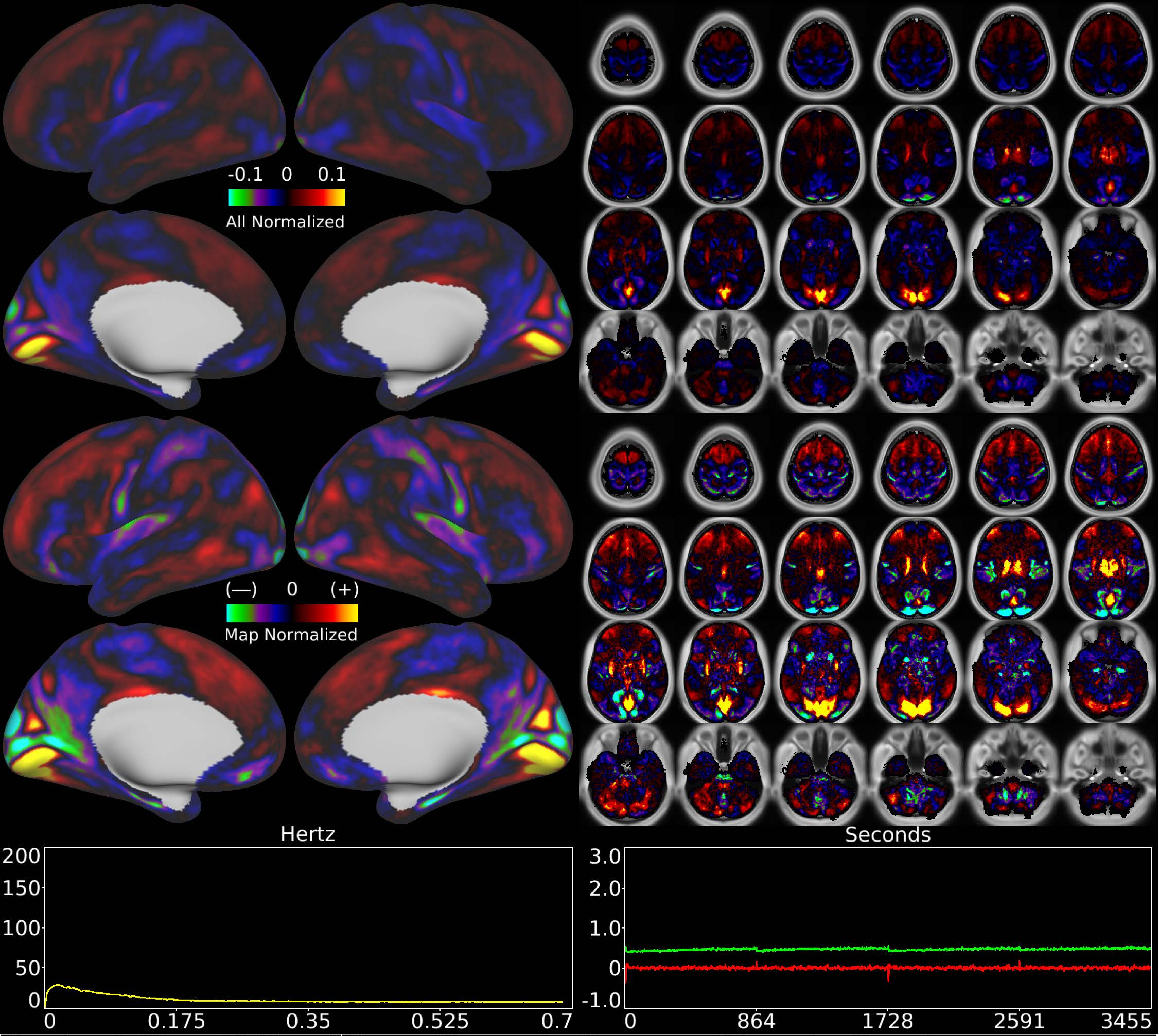


| | | |
|---|---|---|
| Number & Class: 82 Signal | Name: Visuotopic: Paracentral Ventral > Dorsal | |
| RVT Correlated: No | DVARS Dip Associated: No | Cross-Subject Variable: No |
| Single Subject: No | % Variance Explained: 0.33 | Globality Index: 0.22 |
| Task Component: -67 | Taskr Component: -49 | Sleep Noted: No |
| Rationale: Spatial map includes positive and negative patches that respect known retinotopic visual organization (Upper vs Lower) | | |

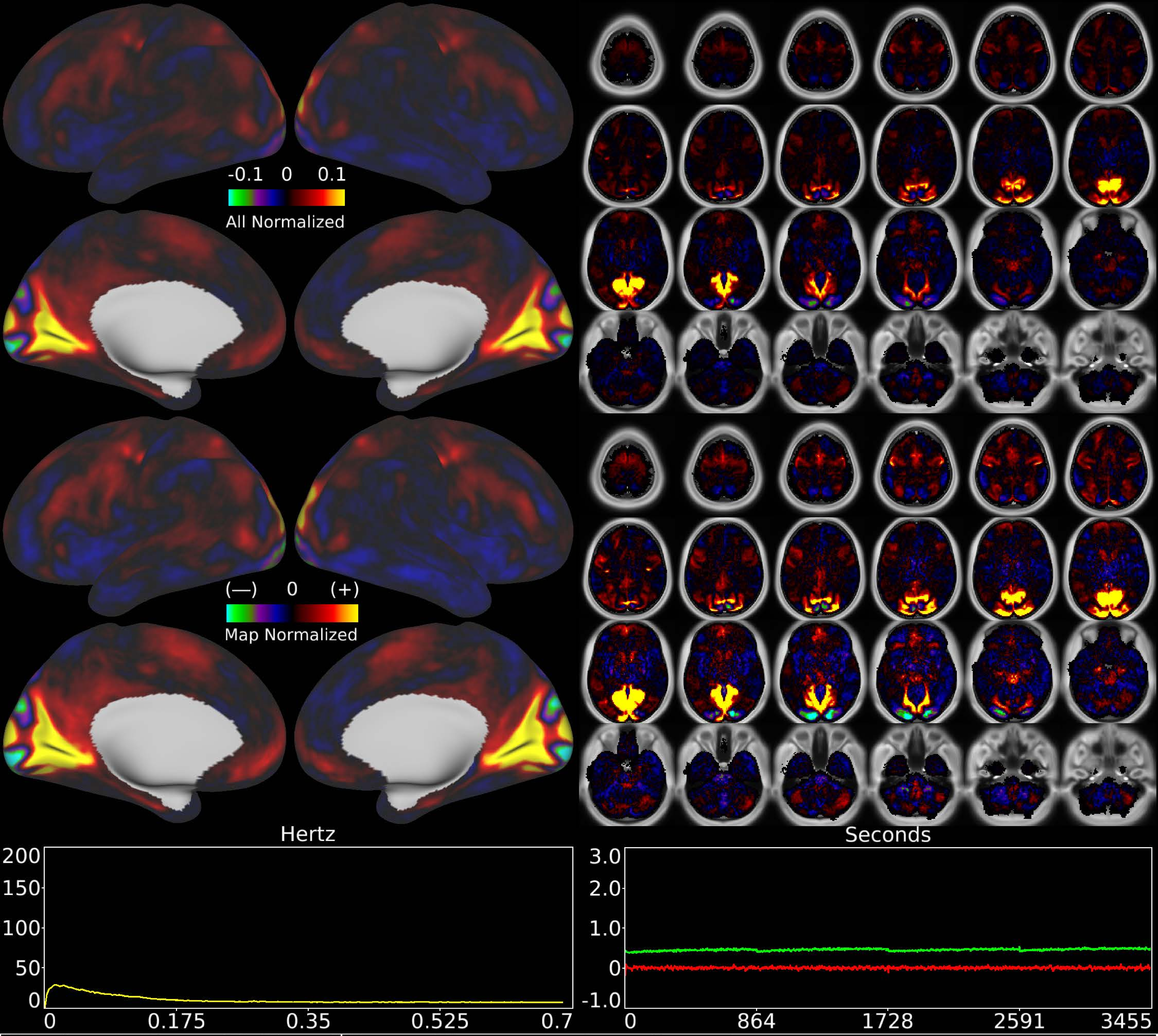


| | | |
|---|---|---|
| Number & Class: 83 Signal | Name: Visuotopic: Complicated | |
| RVT Correlated: No | DVARS Dip Associated: No | Cross-Subject Variable: No |
| Single Subject: No | % Variance Explained: 0.32 | Globality Index: 0.2 |
| Task Component: No | Taskr Component: No | Sleep Noted: No |
| Rationale: Spatial map includes positive and negative patches that respect known retinotopic visual organization (Peripheral vs Foveal) | | |

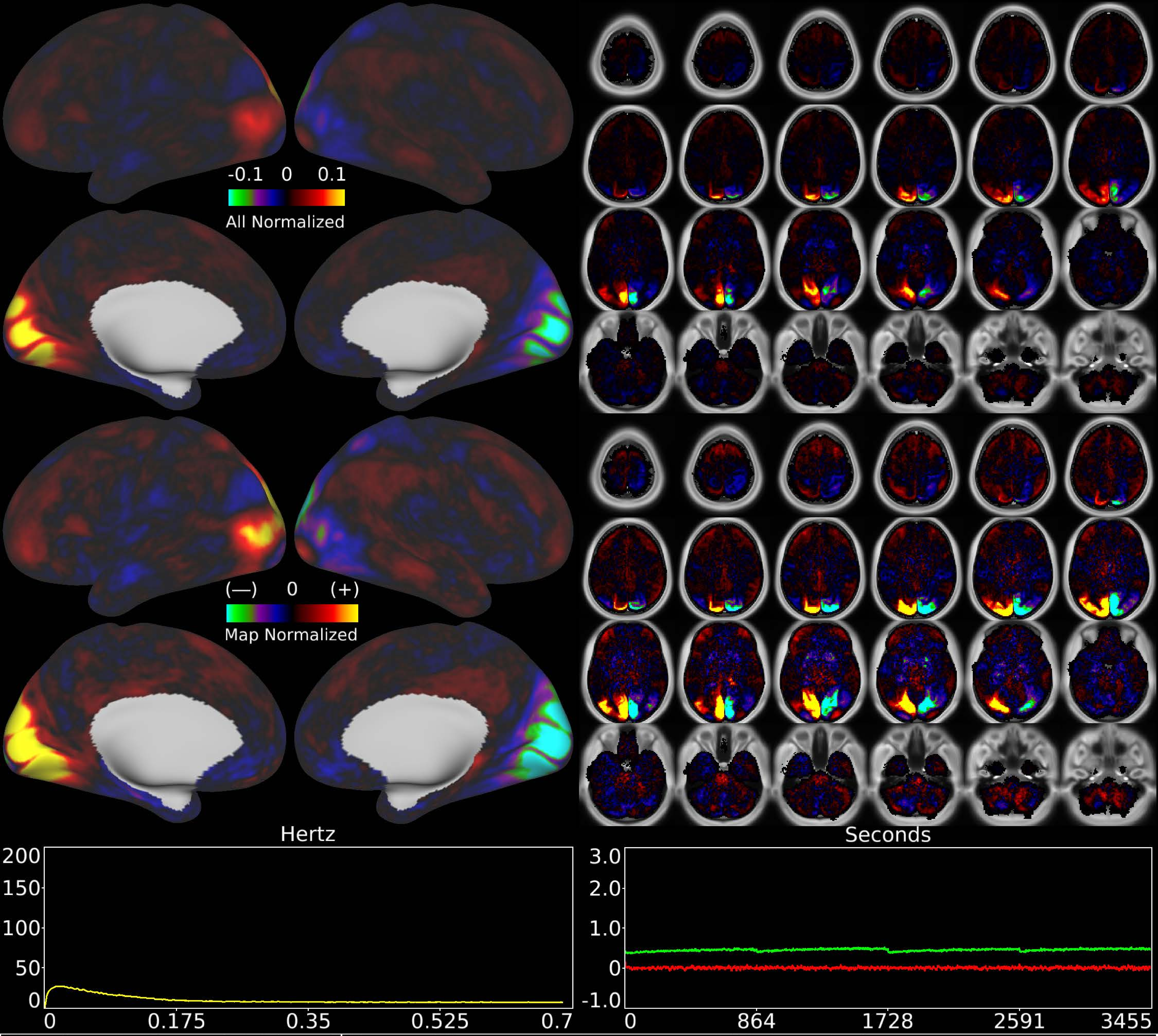


| | | |
|---|---|---|
| Number & Class: 84 Signal | Name: Visuotopic: Paracentral Right > Left | |
| RVT Correlated: No | DVARS Dip Associated: No | Cross-Subject Variable: No |
| Single Subject: No | % Variance Explained: 0.32 | Globality Index: 0.12 |
| Task Component: 69 | Taskr Component: 57 | Sleep Noted: No |

Rationale: Spatial map includes positive and negative patches that respect known retinotopic visual organization (Peripheral Right vs Left)
